# Methane emission of humans is explained by dietary habits, host genetics, local formate availability and a uniform archaeome

**DOI:** 10.1101/2020.12.21.423794

**Authors:** Christina Kumpitsch, Florian Ph. S. Fischmeister, Alexander Mahnert, Sonja Lackner, Marilena Wilding, Corina Sturm, Sandra Holasek, Christoph Högenauer, Ivan Berg, Veronika Schöpf, Christine Moissl-Eichinger

**Affiliations:** Diagnostic and Research Institute of Hygiene, Microbiology and Environmental Medicine, Medical University of Graz, Graz 8010, Austria; Department of Psychology, University of Graz, Graz 8010, Austria; Department of Biomedical Imaging and Image-guided Therapy, Medical University of Vienna, Vienna 1090, Austria; Division of Immunology and Pathophysiology, Medical University of Graz, Graz 8010, Austria; Division of Gastroenterology and Hepatology, Medical University of Graz, Graz, Austria; Institute for Molecular Microbiology and Biotechnology, University of Münster, Münster, Germany; BioTechMed, Graz, Graz 8010, Austria

**Keywords:** archaeome, microbiome, methanogens, methane, gut, gastrointestinal tract, metabolome, metagenome, *Methanobrevibacter*, Christensenellaceae

## Abstract

Archaea are responsible for methane production in the human gastrointestinal tract. Twenty percent of the Western populations exhale substantial amounts of this gas. The underlying principle determining high or low methane emission and its effect on human health was still not sufficiently understood.

In this study, we analysed the gastrointestinal microbiome, archaeome, metagenome, metabolome, and eating behaviour of 100 healthy young adults. We correlated high levels of human methane emission (5-75 ppm) with a 1000-fold increase in *Methanobrevibacter smithii*. This archaeon co-occurred with a bacterial community specialised on dietary fibre degradation, which included members of the Ruminococcaceae and Christensenellaceae. Methane production was negatively affected by high vitamin B12 and fat intake of the subjects, and was positively associated with increased formate concentrations in the gut. Overall, methane emission is explained by dietary habits, host genetics, local metabolite availability and microbiome/archaeome composition, emphasizing the unique biology of high methane-emitters which has potentially positive impact on human health.

Graphical abstract:

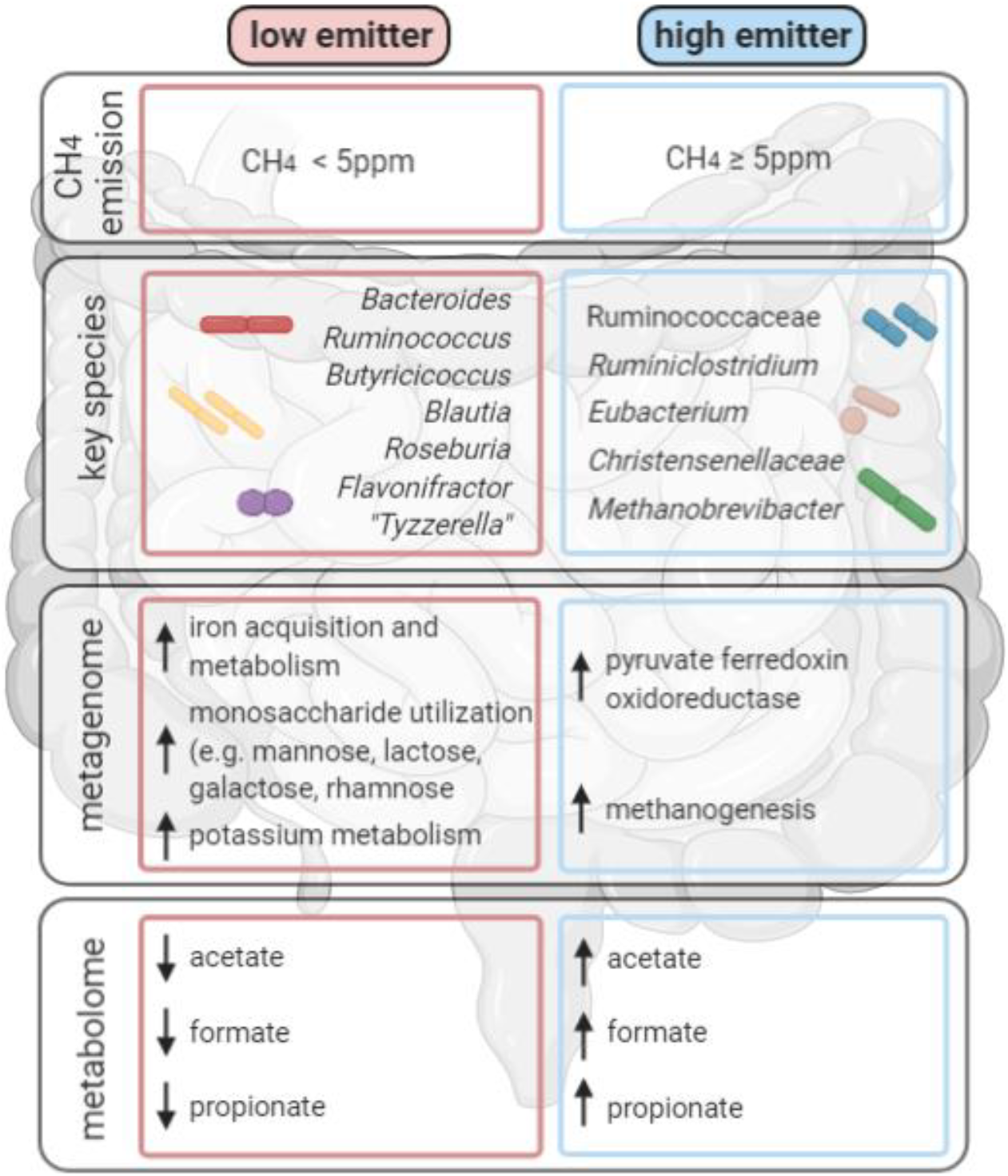

## Introduction

The composition and function of the human microbial community is closely linked with the fitness of the host. In addition to the predominant bacterial component, archaea, eukaryotes (in particular fungi) and viruses contribute important regulatory aspects, influencing the dynamics, host-interaction and metabolic output of the entire microbiome. However, the non-bacterial constituents are often overlooked and remain unconsidered, since standard molecular methods are currently not well developed to resolve this component of the microbiome (Mahnert *et al.*, 2018; Borrel *et al.*, 2020).

Methane-forming archaea (“methanogens”) in the gastrointestinal tract (GIT) were first observed long ago - through the detection of methane in the human breath and flatus. The most-prevalent archaeon in the human body (*Methanobrevibacter smithii*) was isolated nearly 40 years ago (Calloway, Colasito and Mathews, 1966; Miller *et al.*, 1982). In recent years, however, human-associated archaea that reside in the GIT as well as other body sites (e.g. skin, respiratory tract), have gained increased recognition (Koskinen *et al.*, 2017; Moissl-Eichinger *et al.*, 2018; Borrel *et al.*, 2020).

Although the average abundance of archaea in human faecal samples is low as compared to bacteria (Borrel *et al.*, 2020), methanogens are considered to represent key-stone species in the GIT. By maintaining numerous syntrophic relationships with bacteria, methanogens control the efficiency of the bacterial primary and secondary fermentation of complex organic molecules. By consuming by-products of bacterial metabolism they particularly keep the hydrogen concentration low, which would inhibit the fermentation activity and reduce the overall energy yield.

Methanogenic archaea have the unique metabolic capability to form methane (methanogenesis) using H_2_, CO_2_, formate, methyl-compounds, and acetate to thrive. Hydrogenotrophic methanogenesis (the most prevalent pathway in the human gut) uses hydrogen as the electron donor (applies to CO_2_ and methyl-reducing methanogens), whereas no external hydrogen source is needed in methylotrophic and acetoclastic methanogenic pathways (Adam, Borrel and Gribaldo, 2019; Mand and Metcalf, 2019). In the human GIT, methanogens are mainly represented by the Methanobacteriales (*M. smithii, Methanosphaera stadtmanae*) and Methanomassiliicoccales (*Ca.* Methanomassiliicoccus and *Ca.* Methanomethylophilus representatives). Studies based on next generation sequencing (NGS) have also detected signatures of *Methanobacterium* (Methanobacteriales), *Methanosarcina* (Methanosarcinales), *Methanoculleus* (Methanomicrobiales) and representatives of Methanocellales in samples from the human gut. These methanogens contribute to an average human body methane emission of about 0.35 l per day (Lewis *et al.*, 2018), released through the breath and flatus.

Notably, not all human subjects produce the same amount of methane. High methane emission in breath correlates with e.g. sex, age, environmental factors, diet, diseases, and geography, but also with ethnicity and genetic aspects (Polag and Keppler, 2019). The latter has been supported by correlations between *M. smithii* and a single nucleotide polymorphism in a long non-coding RNA of the human genome (Bonder *et al.*, 2016). The abundance of methanoarchaea correlates also with certain bacterial members of the microbiome, such as the Christensenellaceae, representing a highly-heritable bacterial clade in the human GIT (Goodrich *et al.*, 2014; Ruaud *et al.*, 2020). Specifically, the symbiosis of *M. smithii* and *Christensenella minuta* was found to be based on syntrophy, as driven by efficient H_2_ transfer via close physical interactions. Notably, Christensenellaceae have robustly been linked with a low host body mass index (Ruaud *et al.*, 2020).

Although the methods used to determine the individual’s methane production are not standardised with respect to the procedure and cut-off values, it has become clear that only a subpopulation is producing higher amounts of methane, ranging from 14 to 78% in Japanese and rural Africans, respectively (Borrel *et al.*, 2020). The Western adult population was found to range somewhere in-between (24-60%) (Polag and Keppler, 2019). The results of cultivation assays have indicated that a “positive” methane breath test correlates with cultivable methanogen concentrations greater than 10^8^ cells per gram stool (dry weight), and reaching up to 3× 10^10^ cells in high methane producers (Weaver *et al.*, 1986). Considering the measured value of 4× 10^11^ microbial cells per gram dry stool (Stephen and Cummings, 1980), this corresponds to 0.03 to 7.5% methanogens in methane-producing subjects.

Based on this substantial increase in the population of methanogens, as well as the massive methane production (sometimes reaching up to 300 ppm, pers. com. C. Högenauer) in high methane producers, a putative impact on the GIT microbiome and host physiology can be expected. Indeed, the gaseous product of methanogens, methane, is not only a potent greenhouse gas, but also has a physiological effect on the host. While its role as a gasotransmitter is controversially discussed (Boros *et al.*, 2015), methane is a causally linked to a slowed gastrointestinal motility (methane slows down the faeces transit time by up to 59%), probably caused by the direct action of methane on the cholinergic pathway of the enteric nervous system (Pimentel *et al.*, 2006). This fact also partially explains the continuous reports of methanogens being associated with constipation. Other effects were shown in rodent models, such as enhanced exercise capacity (Xin, Sun and Lou, 2016), increased secretion of GLP-1 (Laverdure *et al.*, 2018), or even anti-inflammatory and neuroprotective effects (Boros and Keppler, 2019).

Overall, the role of methanogens *per se* in health and disease is not yet clear, and analyses suffer from methodological pitfalls to correctly detect and characterize the human archaeome, as well as the contradictory information that appears in the literature (for further reading, please refer to (Borrel et al., 2020)).

In this study, we aim to understand the correlation between methane production in young, healthy subjects with the composition of the microbiome (archaeome and bacteriome), microbial function, and diet. We have recruited 100 young volunteers (age 18-37 years), profiled their GIT microbiome, and performed a standardised methane breath measurement. Fifteen subjects out of 100 showed methane levels ≥ 5 ppm and were categorised as high-methane producers. Metagenomic information from stool samples of this subgroup was compared to 15 matched (age, sex, and vegetarianism), low-methane producing controls. Microbial profile and functions were correlated with methane production and dietary habits. Herein, we show that human methane production is caused by a uniform, archaeome predominated by *M. smithii*, and linked to a specific bacterial community, which is specialised to degrade dietary fibres. *Methanobrevibacter smithii* has a key-stone role, as this archaeon consumes H_2_ and CO_2_, which leads to the lower availability of these compounds to the gut microbial community, which, in turn, has effects on overall metabolite production.

## Results (Section headings and subheadings)

### Description of the cohort and datasets

In total, 100 participants (female: *n* = 52, male: *n* = 48; mean age =24.1) were recruited in this study. Metadata information (sex, age, vegetarian yes/no, contraception yes/no, breath methane content as well as metabolite information) of all participants is provided in Supplementary Table 1. All participants provided one stool sample, one breath sample for methane measurements, and a completed dietary questionnaire (Supplementary Table 1). On the basis of the amount of methane emitted participants were grouped into high methane emitters (HE; CH_4_ value: 5-75 ppm) and low emitters (LE; CH_4_ value < 5 ppm). Fifteen percent of the participants were categorised as HEs (Supplementary Table 2), with the percentage in congruence with known levels of methane emission of young adult European cohorts (Polag and Keppler, 2019).

The following data sets were obtained through 16S rRNA gene amplicon sequencing, shotgun metagenomics sequencing, and a questionnaire: “universal” and archaeal 16S rRNA gene profiles for all stool samples (Supplementary Dataset 1 and 2; Supplementary Table 1), and a metagenomics dataset as well as metabolomic information (e.g. acetate, succinate, formate), and detailed dietary information (e.g. diversity, energy, protein, fat, carbohydrates) from matched participants (Supplementary Dataset 3 and 4; Supplementary Table 2).

### High-methane microbiomes are characterised by a specific microbial community and a 1000-fold increase in the *Methanobrevibacter* relative abundance

Using the “universal” approach to amplify 16S rRNA genes, we obtained 2,293,161 sequences after denoising with DADA2 (Callahan *et al.*, 2016) and processing through Qiime2 (Bolyen *et al.*, 2019) which were classified into 17 microbial phyla, 254 genera and 2,570 unique ribosomal sequence variants (RSVs). Using an archaea-targeted approach (Pausan *et al.*, 2019), we obtained 1,035,202 sequences grouped into 4 phyla, 6 genera and 41 unique RSVs. Respective RSV tables are provided in Supplementary Dataset 1 and 2.

We focused on the intact microbial community by applying a propidium monoazide (PMA) treatment to remove the background signals of free DNA (Nocker *et al.*, 2007; Young *et al.*, 2017). Within the “universal” data set, the phylum Firmicutes was found to be predominant (45.84%; 1,051,161 reads), followed by Bacteroidetes (40.46%; 927,837 reads) and Proteobacteria (7.53%; 172,656 reads); the main representative genera were *Bacteroides* (26.61% 610,127 reads), *Alistipes* (6.25%; 143,248 reads), and *Faecalibacterium* (5.82%; 133,442 reads) (Supplementary Figure 1; Supplementary Dataset 1).

When we compared the stool microbial profiles of HEs and LEs, we observed a significant (p = 0.00033; ANOVA) increase in alpha diversity (Shannon Index; Figure 1A.I). Redundancy analysis (RDA) results confirmed that methane production had a significant impact on the microbiome composition (p = 0.001; Figure 1A.II). Although the HE microbial profiles did not group separately in the PCoA plot, HEs formed a sub-cluster within the cloud of LE (Supplementary Figure 2A.I). On a phylum level, results of LEfSe (Linear Discriminant Analysis Effect Size) analyses revealed a significant association between Euryarchaeota and HE and between Bacteroidetes and LE (Figure 1.B, Supplementary Figure 2). On genus level, taxa belonging to *Ruminococcaceae UCG014* as well as the Christensenellaceae R7 group and *Methanobrevibacter* were shown to have potential associations with HEs, whereas *Bacteroides* and *Blautia* were associated with LEs (Figure 1.C). These associations on both taxonomic levels were confirmed through the results of independent ANOVA plot analyses (Supplementary Figure 2) and Spearman-based regression analyses, confirming a highly significant (p < 0.001) positive correlation of *Methanobrevibacter*, the Christensenellaceae_R7_group, Ruminococcaceae (UCG010 and UCG002), and *Desulfovibrio* with high methane emission, and a significant (p < 0.01) negative correlation among *Bacteroides*, the Ruminococcaceae gnavus group, *Flavonifractor*, and *Holdemania* (Supplementary Figure 3; individual *r*s-values provided in the Figure).

**Figure 1.**
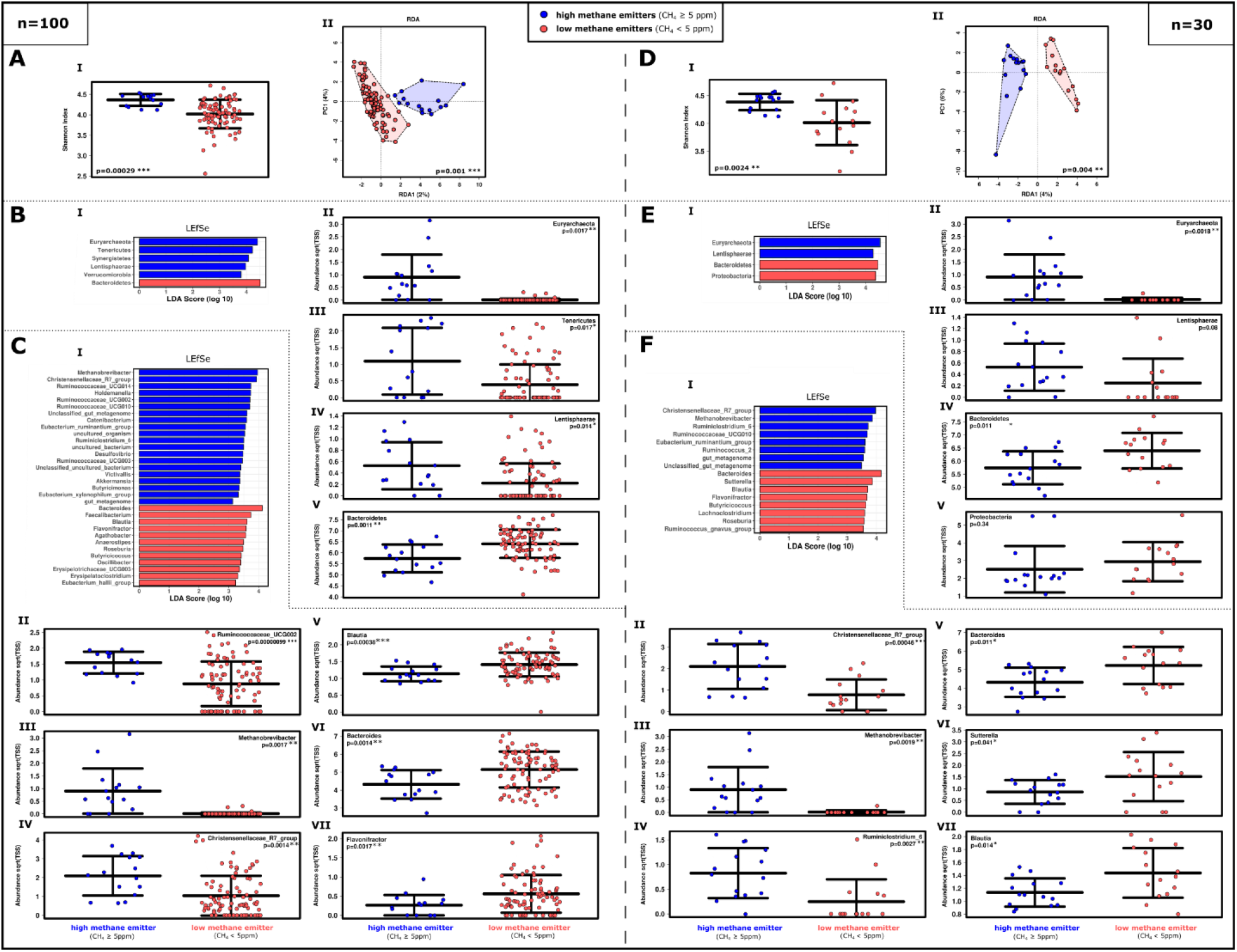
Differences in alpha and beta diversity based on the “universal” approach of 16S rRNA gene sequencing between high (HE) and low methane emitters (LE). **A-C.** Profiles of the whole study cohort (n=100). **D-F**. Profiles of the matched subset only (n=30). **A.I/D.I.** An examination of Shannon diversity index revealed significant differences in alpha diversity (RSV based; ANOVA). **A.II/D.II.** The microbiome of HEs clustered significantly differently in the RDA plot (RSV based). **B.I/E.I.** LEfSe analysis of the 100 most abundant phyla and **B.II/E.II-B.V/E.V.** Relative abundance of selected phyla in ANOVA plots. **C.I/F.I.** LEfSe analysis of the 100 most abundant genera and **C.II/F.II-C.VII/F.VII.** ANOVA plots of selected genera.

An overview of the taxonomic composition of HE and LE samples is given in a Krona Chart (Supplementary Item 1). This type of display confirms the different relative abundances of signatures from *Methanobrevibacter* (HE: 2%, LE: 0.002%), *Bacteroides* (HE: 19%, LE: 28%), the Christensenellaceae R7 group (HE: 6%, LE 2%), and *Ruminococcaceae UCGs* (HE: 22%, LE: 20%).

Moreover, a significant co-occurrence was observed for *Methanobrevibacter* and Christensenellaceae in HEs in every constellation of a Spearman’s rho -based network analysis. Those taxa formed a stable network with different *Ruminococcus*/Ruminococcaceae, *Holdemanella*, and the *Eubacterium ruminantium* group in HEs. On the contrary, LEs were characterised by a network of *Bacteroides, Lachnoclostridium, Sutterella, Flavonifractor, Blautia,* and *Anaerostipes* (Figure 2; Supplementary Figure 4).

**Figure 2.**
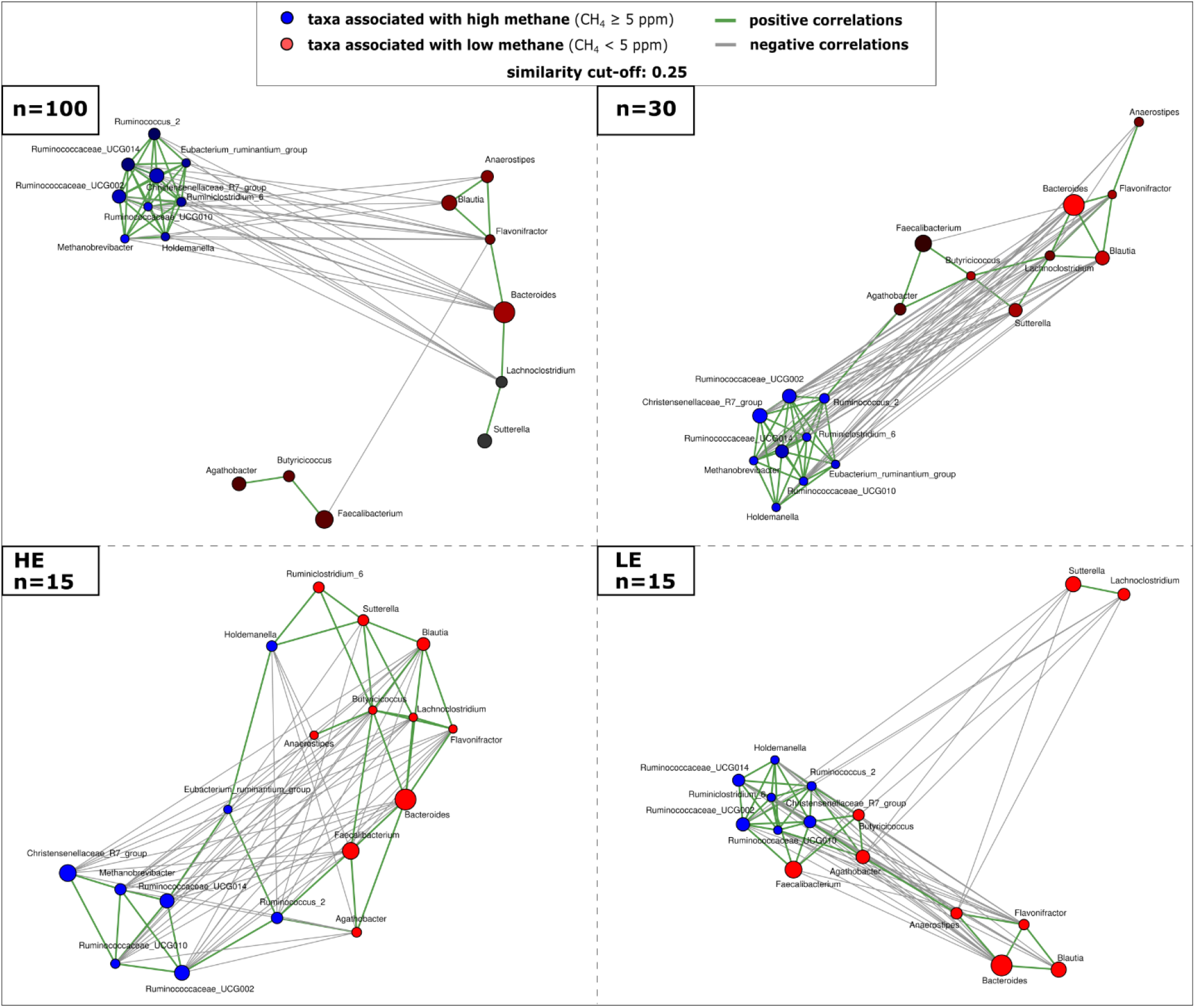
Co-occurrence networks based on Spearman’s rho correlation of selected genera in HE and LE microbiome samples. Taxa were selected based on significantly different relative abundances in both sample types and LEfSe analyses. Left, upper panel: Whole study cohort (n=100), right, upper panel: matched study subset (n=30). Lower panels show co-occurrence patterns in the HE (left) or the LE samples (right).

In order to obtain more detailed insights into the archaeal composition of the microbiomes, we performed archaea-targeted 16S rRNA gene amplicon sequencing. Notably, archaeal reads could not be obtained for all samples (10 out of 100 samples were negative and, namely, samples with IDs (all LEs): 28, 31, 47, 48, 89, 96, 115, 118, 120, and 123 (Supplementary Dataset 2).

Among the remaining samples, Euryarchaeota (99.07%; 1,025,526 reads) was shown to be the predominant phylum, followed by Thaumarchaeota, and Crenarchaeota. Euryarchaeota were represented by *Methanobrevibacter* (96.3%; 996,950 reads), *Methanosphaera* (1.26%; 13,092 reads), and *Methanobacterium* (1.44%; 14,937 reads). *Methanosarcina, Methanocorpusculum, Methanomassiliicoccus,* and unclassified representatives were detected in traces (Figure 3; Supplementary Dataset 2).

**Figure 3.**
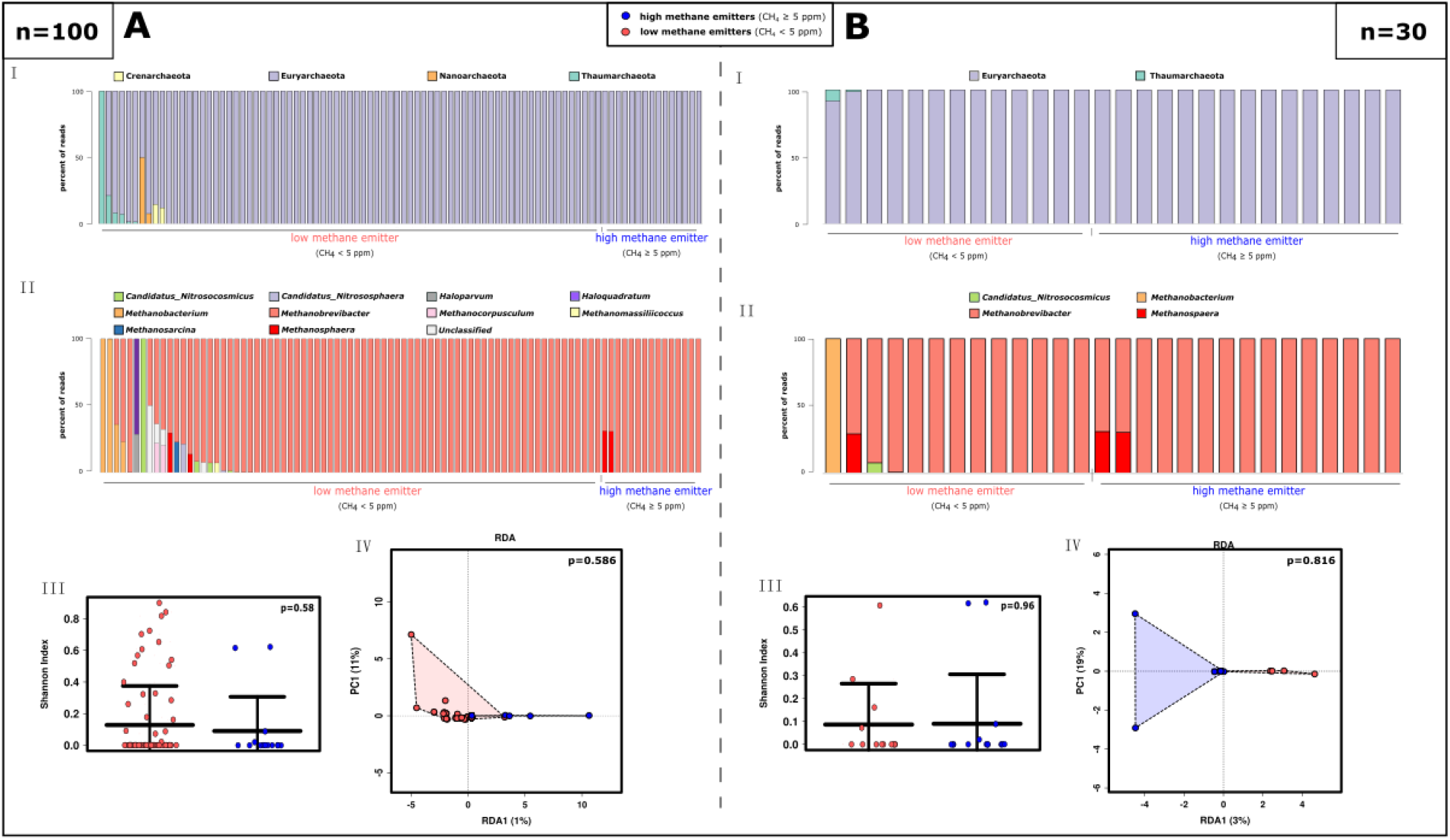
Archaeome profile of HE and LE samples, based on the “archaeal” approach of 16S rRNA gene sequencing. **A.** Profile of the whole study cohort (n=100). **B.** Matched study subset (n=30). **I.** Bar chart of the 20 most abundant taxa compared regarding their low or high methane emission at the phylum level and **II.** at the genus level. **III.** Shannon diversity, and **IV.** RDA plot at RSV level.

The archaeomes of HEs and LEs were not significantly different with respect to their alpha or beta diversity (Figure 3). Samples from HEs did not contain any archaeal signatures apart from the Euryarchaeota, which were represented solely by *Methanobrevibacter* and *Methanosphaera*. In the entire dataset, 21 *Methanobrevibacter* RSVs were observed, whereas *Methanosphaera* was represented by only two RSVs (both genera are represented by one RSV each in the universal dataset). The analysis of HEs samples resulted, on average, in a more than ten-fold, significantly increased number of archaeal reads per sample (HEs: 46,077 archaeal reads per sample, LEs: 4,048 reads (p<0.00001 (Mann-Whitney U Test)).

Based on our observations, the archaeal diversity profile of HE emitters is not significantly different *per se*; the methane emission is driven solely by the predominance of one particular *Methanobrevibacter* strain. The 16S rRNA gene sequence of this strain (dominant in HEs in the universal and archaeal datasets), matched the 16S rRNA gene sequence of *M. smithii* strain KB11 by 100% (NCBI blast), which is a representative of the *Methanobrevibacter*_A *smithii* group according to the GTDB classification (SILVA).

Due to the nature of the study set up, the number of recruited HE and LE subjects was divergent (n=15 and n=85, respectively). Thus, we then focused on an equal subset of the cohort to investigate the statistical relevance of our data regarding methane production. For this purpose, 15 HEs were matched to 15 LEs by age, sex, vegetarianism, and hormonal contraception method (Supplementary Table 2). This data set (n=30) was subjected to the same statistical analyses as described above for the entire cohort.

The overall profile of the reduced (n=30) “universal” dataset was highly similar to the profiles revealed for the non-matched volunteers, and the same predominant phyla and genera were also identified (Supplementary Figure 1B; Supplementary Dataset 1). In this data set, we could also confirm increased alpha diversity in HE (Figure 1D.I, p=0.0024), a the significant impact of methane on the microbiome composition (Figure 1D.II, p=0.004), the formation of a subcluster within the PCoA (Supplementary Figure 2B.I), and a significant difference in the relative abundance of Euryarchaeota and Bacteroidetes in HEs and LEs, respectively (Figure 1E, F; Supplementary Figure 2B).

The significant (p<0.01) co-occurrence of *Methanobrevibacter* and Christensenellaceae in HEs could also be confirmed (Figure 2). In addition, the findings on the archaeome (alpha, beta diversity) did not change (Figure 3B). Confirmation of the congruency of the n=30 data set was an important prerequisite for the subsequent shotgun metagenomic analyses.

### Shotgun metagenomic analyses reveal functional differences in the HE and LE microbiomes

Extracted DNA from stool samples of the matched study subset (Supplementary Table 2) was further used to conduct a shotgun-based metagenomic analysis, which was resolved with respect to taxonomic (see below) and functional information.

The functional analysis of the metagenomics dataset was based on 14,616,890 sequences which were categorised into 28 SEED subsystems and contained 6,956 actual function assignments and 6,589 unique features. The output was organised hierarchically into four levels; level one represented the SEED subsystem and level four represented the most detailed functional information. To gain additional clarity, the information on the detected functions was made available as interactive, hierarchical Krona charts (Supplementary Item 3).

The most prominent subsystems (level 1) identified were “carbohydrate” (17.41%), “clustering-based subsystems”, (12.21%) and “protein metabolism” (8.01%) (Supplementary Figure 5, Supplementary Dataset 3).

Like the profile information derived from 16S rRNA gene data, the diversity of unique functions was significantly higher in HEs as compared to LEs. The impact of methane emission on the overall functions was also found to be significant (Figure 4A, p-values included therein). At level 1, LEfSe analysis identified “protein metabolism”, “nucleosides and nucleotides” and “RNA metabolism” as being significantly correlated with HE samples, whereas LE microbiomes were significantly correlated with “iron acquisition and metabolism”, “carbohydrates”, and “sulfur metabolism” (Figure 4B; Supplementary Figure 6).

**Figure 4.**
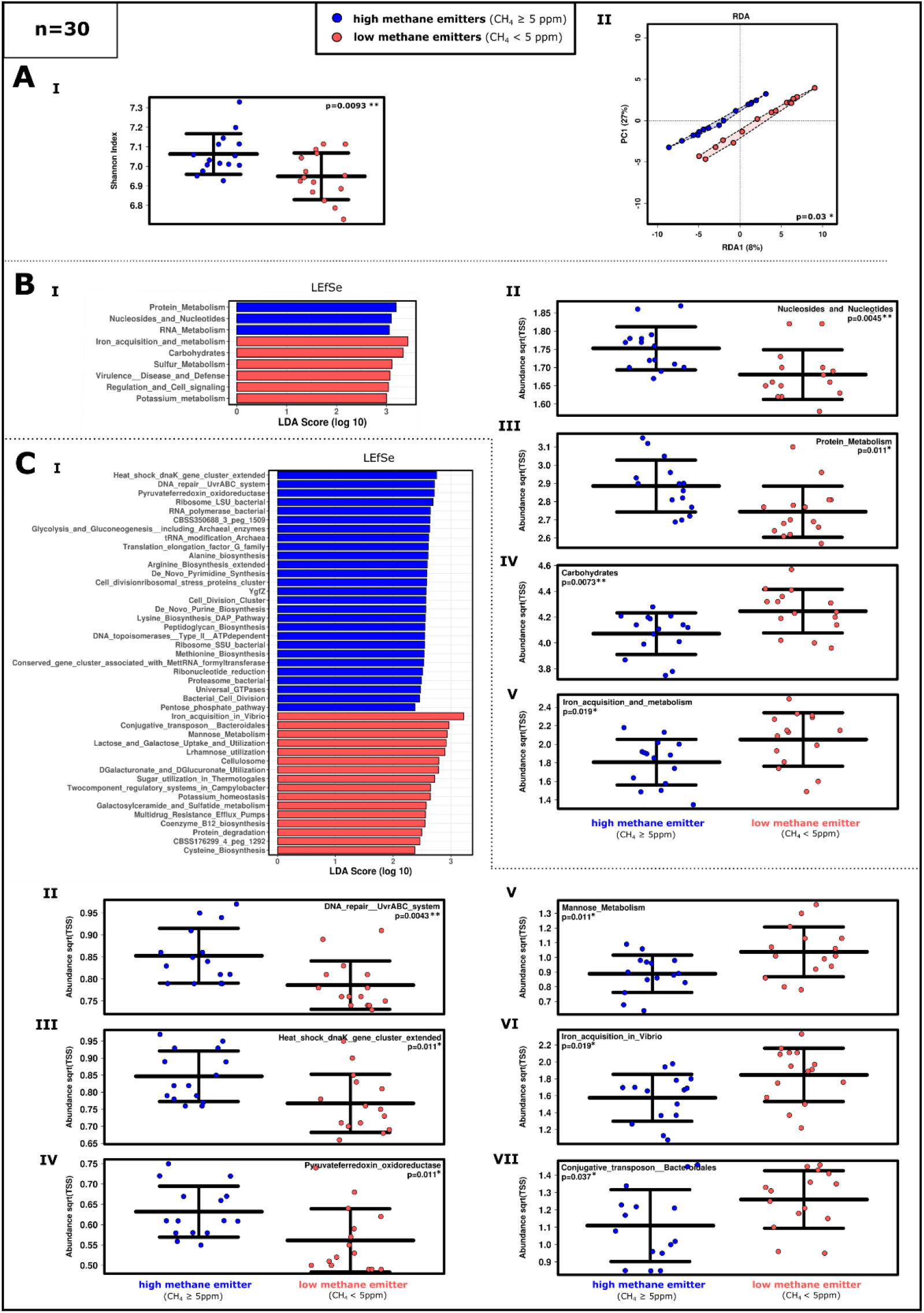
Overview of the divergent functions of the HE and LE based on the shotgun metagenome analysis (subsystems). **A.I.** Shannon diversity and **A.II.** RDA plot at feature level. **B.I.** LEfSe analysis and **B.II-.V.** ANOVA plots at highest subsystem level (level 1). **C.I.** LEfSe analysis and **C.II-V.II.** subsystem at level 3. (100 most abundant; n=30)

The increase in the relative abundance in genes involved in “protein metabolism” in HE (HE: 9%, LE: 8%) was mainly reflected by an increase in the genes associated with “protein biosynthesis” (level 2). This was caused by an increase in the relative abundance of the involved archaeal functions, such as archaea-specific elongation factors or translation initiation factors. The increased proportion of genes involved in RNA metabolism (HE: 5%, LE: 4%) can be explained similarly, as the proportion of archaeal-specific genes was increased (e.g. tRNA modification or transcription elongation factor in Archaea; level 3). The archaeal RNA polymerase was undetectable in LE samples (HE 0.06%, LE: 0%)(Figure, 4B; Supplementary Item 3). The relative abundance of genes involved in “iron acquisition and metabolism” was found to be reduced in the HE dataset (HE: 3%, LE: 4%), which was mainly caused by a relative increase in the number of genes involved in “iron acquisition in Vibrio” in the LE data set (HE: 2%, LE: 4%). These gene sets include the TonB-dependent transport of heme and several siderophores that are produced by a broad variety of bacteria (Wyckoff, Mey and Payne, 2007). Differences in the relative abundance of sulphur metabolism genes (HE: 0.7%, LE: 0.8%) were manifested by increased gene counts in the level of “galactosylceramide and sulfatide metabolism” (level 3; HE: 0.4%, LE: 0.5%), as well as “thioredoxin-disulfide reductase” (level 3; HE: 0.09%, LE: 0.1%).

Based on the results of the LEfSe analysis at level 3, we found that the “heat shock *dnaK* gene cluster” (in “Stress response”) (HE: 0.7%, LE: 0.6%), involved in chaperone *Hsp70* formation, and the “*UvrABC* system” involved in “DNA repair” (in “DNA metabolism”; HE: 0.7%, LE: 0.6%) were significantly associated with HEs. Moreover, we detected an increased contribution of genes involved in “Synthesis of osmoregulated periplasmic glucans”, indicating that Gram-negative bacteria made a high contribution to HE samples (0.04% of functional genes; 0.03% in LE) (Figure 4C; Supplementary Item 3).

Among the functions associated with “carbohydrate”, a particular increase in the LE dataset was observed in the “monosaccharide” (level 2) turnover-associated genes (HE: 3%, LE: 4%) (e.g. in D-galacturonate, L-rhamnose, xylose, L-arabinose, and L-fucose metabolism), as well as in the uptake of lactose and galactose. Especially mannose metabolism (level 3; HE: 0.8%, LE: 1%), including the metabolism of alpha-1,2-mannosidase (level 4; HE: 0.6%, LE: 0.9%), was found to be increased in LE samples.

Notably, the “Pyruvate ferredoxin oxidoreductase” (HE: 0.4%, LE: 0.3%; alpha and beta subunits; HE: 0.04% LE: 0.01% and HE: 0.02% LE: 0.01%, respectively), which is part of the “central carbohydrate metabolism” of pyruvate, propanoate, and butanoate, and the reductive carboxylate cycle, was found to be significantly increased in HE samples. This enzyme (also known as pyruvate synthase), catalyses the interconversion of pyruvate and acetyl-CoA, and thus is responsible for the binding or release of CO_2_ with the help of ferredoxin.

Genes involved in “methanogenesis” were rarely abundant in the LE dataset (0.00004%), but reached a 0.1% overall relative abundance in the HE dataset. This was also reflected by the methyl-coenzyme M reductase, which is responsible for the release of methane in the last step of methanogenesis, and whose alpha subunit was represented in a proportion of 0.01% in the HE dataset but only of 0.00001% in the LE dataset. Subunits beta and gamma were not detectable in the LE dataset. Notably, genes involved in “methanogenesis from methylated compounds” comprised 0.01% in the HE dataset, and 0.005% in the LE dataset, indicating that a similar proportion of these genes existed in both datasets, largely independent of methane emission.

### Shotgun metagenomics confirms taxonomic differences between HE and LE microbiomes

From the metagenomics dataset, 68,084,011 fragments of ribosomal RNA genes were obtained. These were classified into 57 phyla, 889 genera, 2,192 species, and 2,193 unique features. The samples were predominated by signatures of Bacteroidetes (62.69%; 42,678,473 reads), Firmicutes (28.80%; 19,611,159 reads), and Proteobacteria (3.98%; 2,706,855 reads) at the phylum level; *Bacteroides* (51.10%, 34,793,498 reads), *Clostridium* (6.54%; 4,456,393 reads), and *Eubacterium* (4.93%; 3,357,015 reads) at the genus level; and *Bacteroides vulgatus* (8.60%; 5,855,942 reads), *Bacteroides fragilis* (4.48%; 3,051,022 reads), and *Bacteroides sp. 4_3_47FAA* (4.14%; 2,817,577 reads) at the species level (Supplementary Figure 7; Supplementary Figure 8; Supplementary Dataset 4).

Overall, 488,550 sequences (0.72%) were assigned to the archaeal domain, 67,447,694 (99.07%) to the bacterial domain, 110,352 (0.16%) to Eukaryota, 35,836 (0.05%) to viruses, and 1,579 (0.002%) to other sequences. In the HE metagenomic dataset, 0.61% of all taxonomic information could be assigned to archaea, whereas 0.11% were archaeal reads in the LE dataset (see also Supplementary Dataset 4). An additional Krona chart based on archaeal and bacterial signatures only is provided in Supplementary Item 4 (Supplementary Item 4).

The taxonomic information that could be extracted from the metagenomics data was highly similar to the information that was derived from 16S rRNA gene amplicon sequencing. This information also revealed that the LEs and HEs were significantly different in terms of their alpha and beta diversity (Figure 5.I-.II; p-values provided within the figure; Supplementary Figure 9.I).

**Figure 5.**
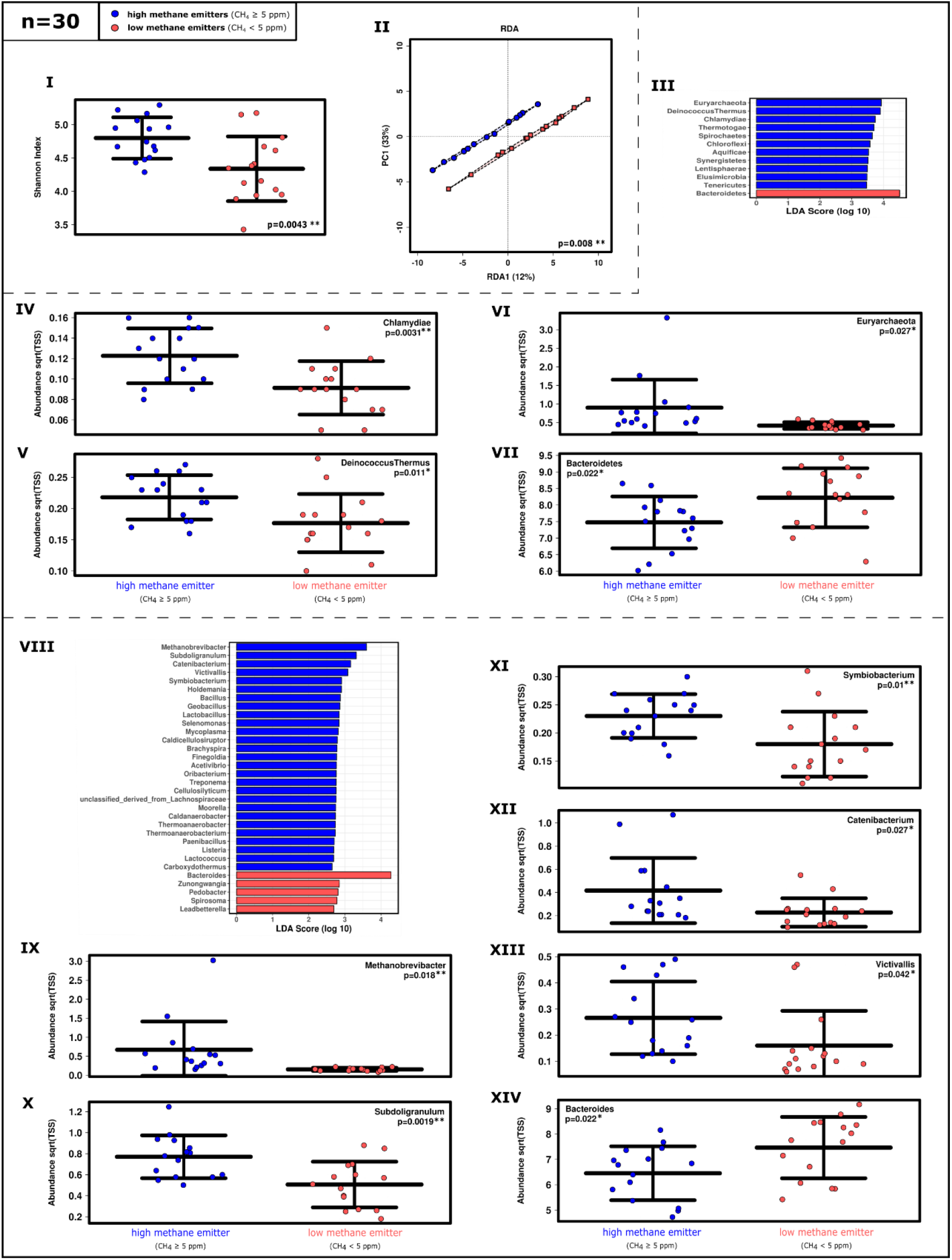
Shotgun metagenome-derived information on the microbial community composition in samples of HEs and LEs (RefSeq). **I** Shannon diversity and **II.** RDA plot based on strain level. **III.** LEfSe analysis and **IV-VII.** ANOVA plots at phylum level. **VIII.** LEfSe analysis and **IX-XIV.** ANOVA plots at the genus level (100 most abundant taxa; n=30).

The significantly higher abundance of archaea in HE samples was confirmed by the results of the LEfSe analysis and ANOVA plots at the super-kingdom level, whereas 14.64% of all archaeal reads were retrieved from the LE samples, and 85.36%, from the HE samples (Supplementary Figure 9.II-.III; Supplementary Dataset 4). Euryarchaeota, Deinococcus-Thermus, and Chlamydia were significantly more abundant in the microbiome of HEs and Bacteroidetes in the microbiome of LEs at phylum level. *Methanobrevibacter*, *Subdoligranulum*, and *Catenibacterium* were associated with HEs, and *Bacteroides*, *Zunongwangia,* and *Pedobacter* were associated with LEs at the genus level (Figure 5.III-.XIV; Supplementary Figure 9.IV-.V). At the species level, *Methanobrevibacter smithii*, *Eubacterium siraeum,* and *Subdoligranulum variabile*, and *Bacteroides vulgatus*, *Bacteroides sp. 4_3_47FAA*, and *Bacteroides sp. 2_2_4* were correlated with the microbiomes of HEs and LEs, respectively (Supplementary Figure 8.II-.III). Notably, signatures of Christensenellaceae could not be retrieved, a phenomenon that has been reported earlier (Ruaud *et al.*, 2020).

### Archaeal profiles can be used to predict methane emissions and are not associated with specific viral or eukaryotic signatures(n=30)

In the next step, we analysed whether it was possible to predict the methane emission levels of individuals based on compositional and functional information derived from their stool microbiomes. Specifically, we applied supervised learning methods that had been trained on the amplicon and metagenomic datasets. Although the individual datasets were rather small, which increases the risk of overfitting the learning model, the overall prediction accuracies reached 63.6% in case of 16S rRNA gene amplicons and up to 100% for RefSeq in the shotgun dataset. When we applied the classification model to a larger public dataset with unknown methane emissions, the estimators achieved 85% prediction accuracy. Hence, despite the obvious limitations of our classification model due to sample size and likely overfitting, these results indicate that it has a high potential for predicting methane emissions above 5 ppm.

Network analyses of the archaeome profile in HE and LE on the species level revealed again the predominance of *Methanobrevibacter* species under HE conditions, whereas LE samples were characterised by a more diverse but rarely abundant archaeome (Supplementary Figure 11). HE samples were characterised by the overwhelming predominance of *M. smithii* (70% of all archaeal taxonomic features; 9% in LE), with *M. stadtmanae* representing 1% (3% of all archaeal tax. features in LE), and Thermoplasmatales (Methanomassiliicoccales), 0.3% (1% in LE). An extraordinarily broad diversity of archaea was detected in both datasets, including members of the Methanobacteriales, Methanomicrobiales, Methanosarcinales, Methanococcales, Thermococcaceae, Halobacteriaceae, and Archaeoglobaceae, as well as unclassified reads from Thaumarchaeota, and including a number of taxa that had not been previously detected in the human microbiome (Supplementary Figure 10; Supplementary Dataset 4; Supplementary Item 5).

In order to identify other microbial variables that influence the archaeal profiles, and in particular the profile of the dominant *M. smithii* strain, eukaryotic signatures and viral/phage signatures were correlated accordingly. However, no archaeal viruses could be identified, and also no correlating eukaryotic/protist signatures could be observed, indicating that the detected methanogenic archaea are free-living nature.

### Diet modulates HE and LE keystone taxa and methane production

As indicated above, we identified a number of representative bacterial and archaeal genera which were indicative for HE and LE, respectively (Supplementary Table 3; Figure 6; see also Figure 2 co-occurrence patterns). To perform more detailed analyses on the RSV level, we proceeded with amplicon data (matched dataset) because taxonomic information for Christensenellaceae was missing from the metagenomics dataset. We identified 21 RSVs, revealing significantly discriminative abundances (identified through LEfSe analyses) and substantial mean abundances (top 600 taxa) (Supplementary Table 3, Figure 6). We found that the LE profile was mainly defined by four RSVs of *Bacteroides*, four RSVs of *Butyricicoccus*, and one RSV each of *Flavonifractor, Blautia*, “*Tyzzerella”, Ruminococcus* (*R. gnavus* group), and *Roseburia*, whereas the HE profile was driven by one RSV of *Methanobrevibacter*, three RSVs of the Christensenellaceae R7 group, two RSVs of *Ruminiclostridium*, one RSV of Ruminococcaceae UCG010, and one RSV of *Eubacterium* (*E. ruminantium* group) (Figure 6, Supplementary Table 3). This selection of keystone taxa was further supported by 84 dereplicated high quality MAGs (metagenome assembled genomes; mean completeness 90%, mean contamination 7%, Supplementary Table 4) with replication rates in the range of 1.3 to 2.6 (Methanobrevibacter smithii 4 MAGs, Bacteroides 32 MAGs, Christensenellales 19 MAGs, Ruminococcaceae 19 MAGs, Ruminiclostridium 2 MAGs, Ruminococcus 4 MAGs).

**Figure 6.**
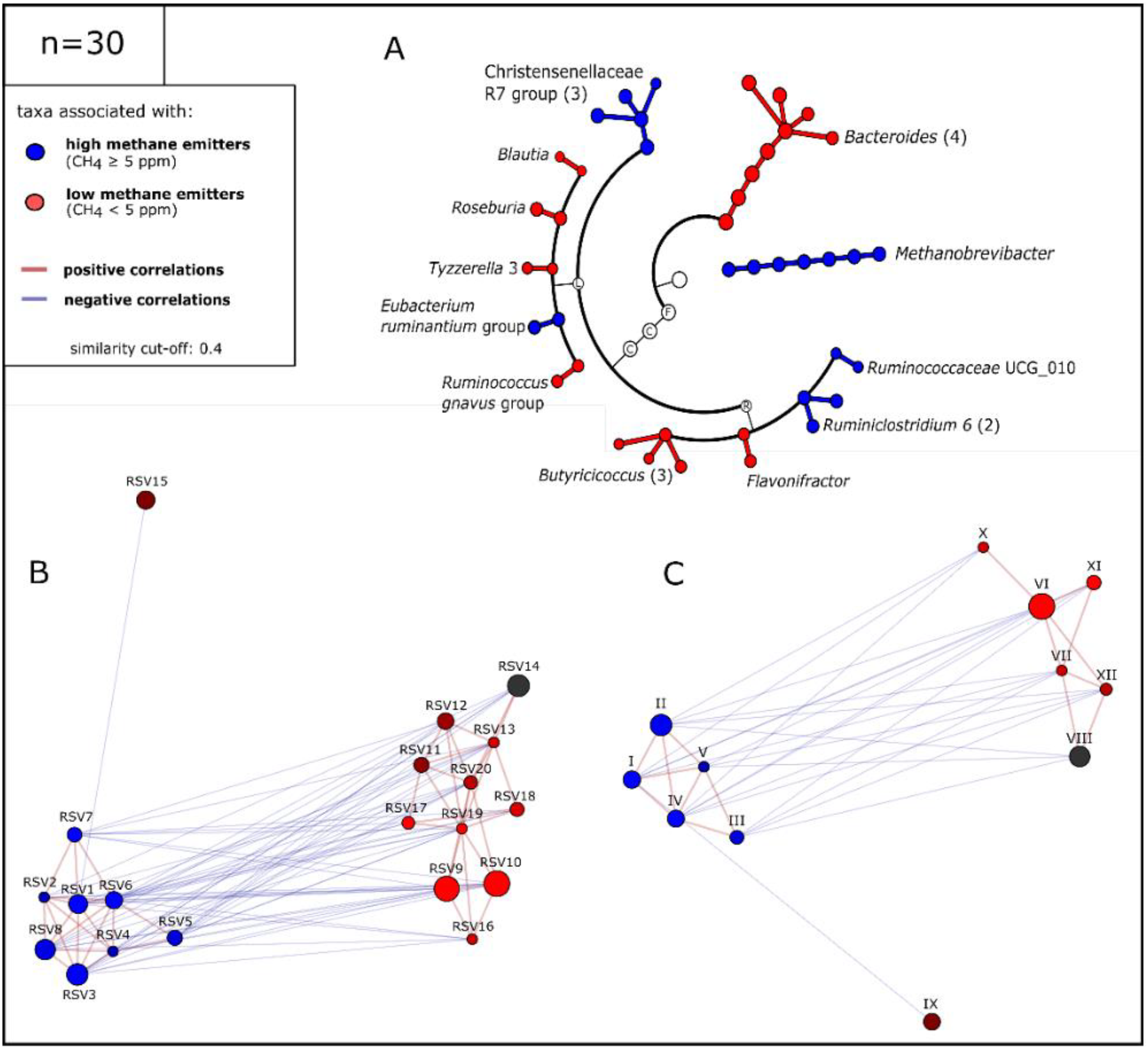
Identified keystone taxa in HE and LE subjects. **A.** Cladogram of LE and HE keystone taxa. F: Firmicutes, C: Clostridia/Clostridiales, L: Lachnospiraceae, R: Ruminococcaceae. Numbers in brackets indicate the number of contributing RSVs; **B.** and **C.** Network of keystone taxa of HE and LE at RSV and genus levels, respectively. I → RSV1: *Methanobrevibacter*; II→ RSV2-4: Christensenellaceae R7 group; III → RSV5: Eubacterium ruminantium group; IV → RSV6-7: *Ruminiclostridium*; V → RSV8: Ruminococcaceaea UCG010; VI → RSV9-12: *Bacteroides*; VII → RSV13: *Ruminococcus gnavus group*; VIII → RSV14: *Blautia*; IX → RSV15: *Roseburia*; X → RSV16: “*Tyzzerella*”; XI → RSV17-19: *Butyricicoccus*; XII → RSV20: *Flavonifractor* (also see Supplementary Table 3)

In a subsequent step, we were interested in examining the way diet correlates with the relative abundance of the identified keystone taxa and methane emissions. A Food Frequency Questionnaire (FFQ) (Haftenberger *et al.*, 2010) was used to assess the food habits of each participant during the four weeks prior to sampling. Overall, the daily intake of 19 food ingredients were tracked, including proteins, carbohydrates, fat/saturated fat/omega-3 fatty acids/omega-6-fatty acids, fibre, alcohol, sodium, vitamins C/B12/E/D, zinc, calcium, magnesium, potassium, iron, water, as well as three diet’s quality indicators, including overall energy intake (kcal), nutritional variety, and diversity (Supplementary Table 2).

Regarding the HE states, trends for negative correlations could be identified for *Methanobrevibacter* counts and energy, vitamin D, or calcium in a BioEnv plot (Spearman’s rho correlation; Supplementary Figure 12). The results of a correlation analysis revealed that a higher relative abundance of *Methanobrevibacter* (n=30 amplicon-based dataset) was negatively correlated with total fat (*r*s=-0.435, p=0.016; if not stated otherwise a Spearman’s correlation analysis was performed), saturated fat (*r*s=−0.421, 0.021) and omega-3 fatty acids (*r*s=−0.407, p=0.026). Trends indicating a correlation were observed for vitamin B12 intake (*r*s=−0.355, p=0.054). Similar trends for vitamin B12 (*r*s=−0.465, p=0.01) and omega-3 fatty acid (*r*s=−0.349, p=0.059) intake were seen when examining the relative abundance of the Christensenellaceae R7 group. Vitamin D intake was negatively correlated with the Christensenellaceae R7 group relative abundances (*r*s=−0.345, p=0.062;), whereas *Ruminococcaceae UCG10* was positively correlated with alcohol consumption (*r*s=0.390, p=0.033) (Supplementary Table 5).

Within the LE community cluster, an analysis of the genera *Bacteroides*, *Flavonifractor* and the *Ruminococcus gnavus group* revealed a trend with respect to a negative correlation with dietary fibre intake (*r*s=−0.379, p=0.039; *r*s=−0.517, p=0.003 and *r*s=−0.382, p=0.037, respectively;). The relative abundance of *Blautia* positively correlated with vitamin B12 levels (*r*s=0.505, p=0.004) and protein intake (R=0.422, p=0.020), whereas protein (*r*s=−0.375, p=0.041) as well as zinc (*r*s=−0.370, p=0.044) intake was negatively correlated with “*Tyzerrella”*. Interestingly, only the presence of the genus “*Tyzerella”* was also positively correlated with vegetarianism (*r*s=0.325, p=0.08). Apart from this, vegetarianism only correlated with different dietary compound intake, namely, vitamin C and sugar intake was positively correlated (*r*s=0.490, p=0.006 and *r*s=0.441, p=0.015, respectively), whereas food diversity and vitamin B12 levels (*r*s=−0.473, p=0.008 and *r*s=−0.449, p=0.013, respectively) were negatively correlated with vegetarianism (Figure 7, Supplementary Table 5).

**Figure 7.**
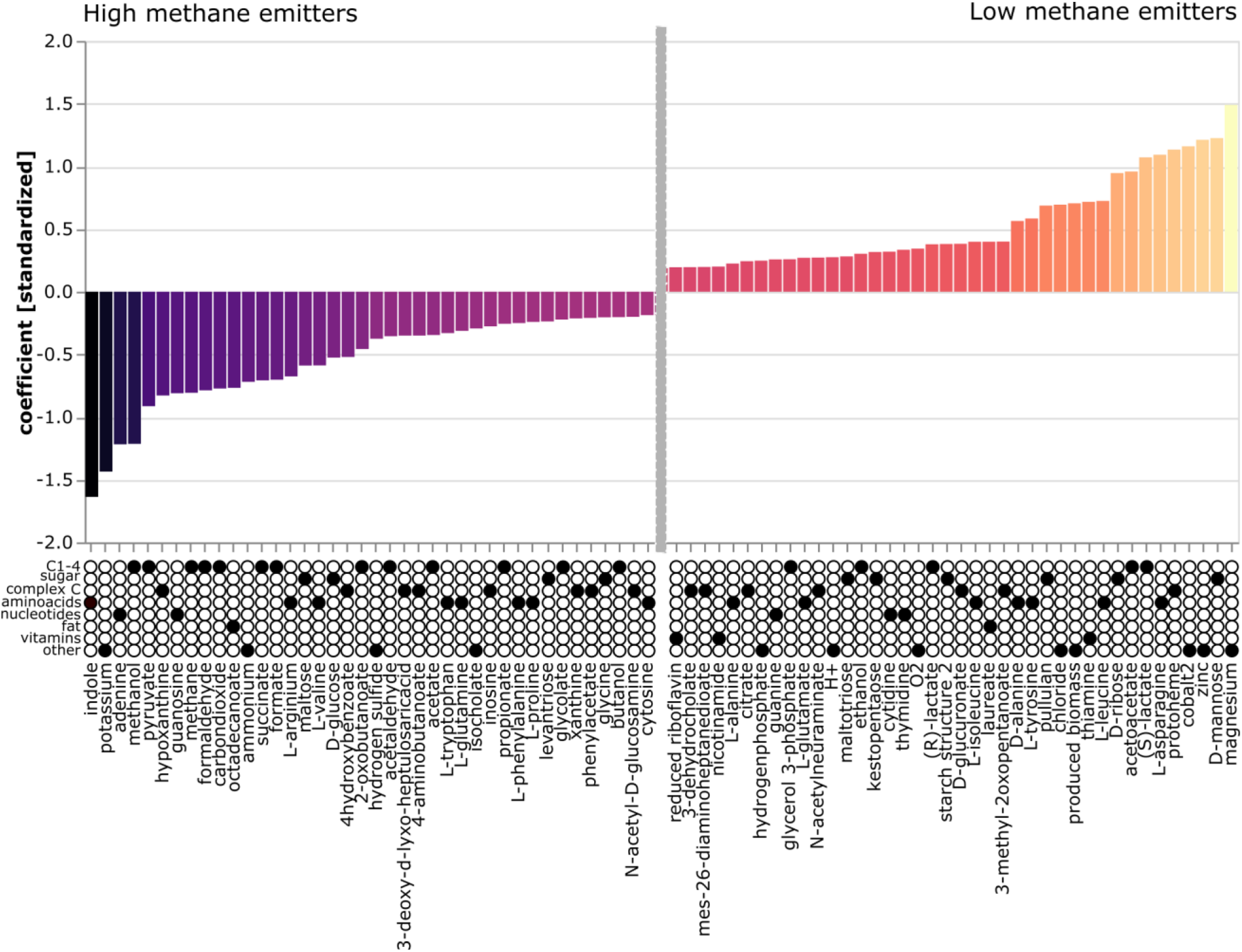
MICOM model-based flux balance analysis of keystone taxa. The 40 most significant metabolites are shown for each condition. Left: HE, right: LE.

Altogether, the information derived from the subjects’ dietary information revealed that one specific bundle of dietary compounds (high dietary fibre levels, low fat and low vitamin B12 intake) is associated with HE status, confirming the above-mentioned observation that the HE microbial community possesses a higher ability to degrade dietary fibres (Supplementary Table 5).

### Methane emission is driven by reduced vitamin B12 intake and fibre-derived, increased formate concentrations

HE and LE keystone communities are each metabolically highly interwoven. Overall, degradation of dietary carbohydrates results in metabolic cycles of short chain fatty acids and CO_2_/H_2_ (Supplementary Figure Metabolic Interaction). Under LE conditions, these metabolites are trapped in the cycle until they are uptaken by the host or used for biomass production. The conversion of H_2_/CO_2_/formate into methane by *Methanobrevibacter* under HE conditions, however, results in a metabolic “dead end”, as methane cannot further be metabolised by gut microbiota or human epithelial cells. Formate-based methanogenesis is widely distributed amongst human-associated methanogens, as e.g. all *Methanobrevibacter* species detected in a catalogue of 1,167 genomes have the capability to use formate for methanogenesis (Chibani et al., 2020).

To characterize the role of these metabolites in more detail, we performed NMR-based metabolomic analyses of the stool samples to assess the relative quantities of formate, acetate, lactate, butyrate, succinate, and propionate in samples from both groups. Indeed, we recognised a significant correlation between the formate concentration and methane emissions (in ppm, Spearman’s-rho correlation coefficient 0.491, p=0.006), confirming the important role of the C1 metabolite. Formate concentration was significantly correlated with acetate (spearman-rho correlation coefficient 0.628), butyrate (0.416) and propionate (0.448) abundance, whereas no correlations were found for lactate and succinate (0.204 and 0.258, respectively). In fact, we measured an increase in formate concentrations (1.5-fold, based on median concentrations per group), acetate (1.35-fold), and propionate (1.17-fold) under HE conditions, whereas the butyrate, lactate, and succinate concentrations remained largely equal (Supplementary Table 6).

To collect more information about the metabolic interaction and growth rates of the different microbial communities, we used MICOM (Diener, Gibbons and Resendis-Antonio, 2020) (XXXrefXXX), a tool that allows us to apply flux balance analysis (FBA) to entire microbial communities. Growth simulations (growth rates, growth niches, metabolite consumptions and phenotype associated fluxes; Supplementary Dataset 5 and 6) were based on the individual dietary information obtained from the donors (Supplementary Dataset 7), and community models were based on the AGORA 1.03 genus model (Magnúsdóttir *et al.*, 2017). The results of the analysis performed on previously identified keystone taxa confirmed a significant association between the HE conditions and an increased flux of C1 metabolites, such as methanol, formaldehyde, carbon dioxide and formate (Fig. 7), as well as acetate and propionate. LE conditions were associated with D-mannose, lactate, ribose levels, and overall a greater complexity of organic molecules. Notably, the hydrogen flux was only minimally associated with HE (−0.021595761). Fluxes in vitamin compounds (nicotinamide, riboflavin, thiamine, pyridoxin, menaquinone 8) were strongly associated with the LE conditions. In addition, LE conditions were significantly associated with fluxes in magnesium, zinc, cobalt, chloride, and biomass production, whereas HE conditions were significantly associated with higher fluxes in potassium, ammonium, and hydrogen sulphide.

Notably, in our model, all identified keystone members of both community types were involved in formate turnover, emphasizing the very important role of this C1 compound, whereas vitamin fluxes were mainly associated with *Blautia*, *Clostridium* (“*Tyzzerella*”) and *Ruminococcus* representatives, all of which are members of the LE community (Supplementary Dataset 6).

In our model, the HE community was strongly associated with increased indole fluxes. Indole is an important tryptophan break-down product, and controls a number of microbial processes, such as biofilm formation, drug resistance, and virulence (Lee and Lee, 2010).

## Discussion

In this study, we focused on performing detailed analyses of human methane emissions, which are strong indicators of the biological contribution of the methano-archaeome to human physiology. Using amplicon-based and metagenomic sequencing, NMR-based metabolomics, dietary intake analysis and metabolic modelling, we were able to show that: i) high methane emission is linked to a significantly higher microbial alpha diversity in the GIT, ii) the microbial community composition and function differs significantly between high- and low- methane emitters and is pronounced in specific key-taxa, iii) methane emission is driven by dietary habits, such as high fibre, low fat, and low vitamin B12 uptakes, iv) C1 compounds, short-chain fatty acids and particularly formate, are keystone metabolites associated with methane formation. Our analysis results confirmed that detectable methane formation is associated with a uniform archaeome, which is predominated by *M. smithii* (Goodrich *et al.*, 2014). Although this archaeal species is generally highly prevalent in the population (detectable in about 97.5% of all analysed subjects; (Dridi *et al.*, 2009)), its abundance in our study was highly variable. High methane emitters (HEs) revealed a relative abundance of approx. 2% (1.37% in the shotgun metagenomic dataset), whereas LEs were characterised by theextraordinarily low contribution of about 0.002% of this species (0.19% in the shotgun metagenomic dataset; see also (Borrel *et al.*, 2020).

The abundance of *Methanobrevibacter* was strongly correlated with a core group of keystone species, including various Ruminococcaceae and Christensenellaceae (see also: (Vojinovic *et al.*, 2019)). In our study, three Christenellaceae RSVs, which co-occurred stably with *Methanobrevibacter,* were indeed significantly associated with methane production. The interplay between *Methanobrevibacter* and Christensenellaceae is of great interest, as this syntrophic partnership has been associated with a lean phenotype (Goodrich *et al.*, 2014) and a reduced gain of fat tissue (Oki *et al.*, 2016; Alonso *et al.*, 2017) in earlier publications. Notably, both taxa are considered to be highly inheritable (Goodrich *et al.*, 2014; Waters and Ley, 2019). In co-culturing studies, the methanogenic partner shifted the *Christensenella minuta* metabolism, probably due to its potent hydrogen consumption, toward acetate production rather than toward butyrate production, leading to increased H2 and CO_2_ production (Goodrich *et al.*, 2014; Ruaud *et al.*, 2020). Although this observation would indicate a bilateral syntrophic relationship of both microorganisms, we observed in our study, that both partners were unevenly affected by LE and HE conditions: Christensenellaceae were present in both communities (2% in LE), and signatures increased only three-fold towards those observed under HE conditions, whereas *Methanobrevibacter* signatures increased 1000-fold, probably indicating a more complex underlying principle. Indeed, we could not identify any dietary-derived compound which had a direct, significantly stimulating or inhibiting effect on the Christensenellaceae population.

The complexity of ingested saccharides is an important modulator for the composition and functionality of a gastrointestinal microbiome, and an interesting link between cellulose degradation and methane emission was observed by other researchers. Chassared et al. (2010) described that dominant cellulose degraders isolated from non-methane-excreting subjects are mainly affiliated with Bacteroidetes, while they are predominantly represented by Firmicutes in methane-excreting individuals (Chassard *et al.*, 2010). In our study, we also identified *Bacteroides* and *Roseburia*, which belong to the phylum Bacteroidetes, as well as Christensenellaceae, *Ruminiclostridium* and *Ruminococcaceae* (Firmicutes), as important key taxa in LE and HE subjects, respectively. Notably, *Bacteroides* (which was shown to be significantly negatively correlated with dietary fibres in our study) and *Roseburia*, unlike high- H_2_- producing *Ruminococcus* sp., are not able to digest e.g. microcrystalline cellulose (Aminov *et al.*, 2006; Duncan *et al.*, 2006; Chassard *et al.*, 2010). This indicates that the type of dietary fibre has a potential modulating impact on methane production.

The negative correlations observed for fat intake and methanogen abundance are highly congruent with previous observations made in ruminants, where an increased fat (oil) concentration in the diet led to a reduced enteric methane production of up to 36% ((Alvarez-Hess *et al.*, 2019) and references therein). It is considered that dietary fat affects methane production in rumen because it reduces the hydrogen accumulation through fatty acid biohydrogenation, leading to the conversion of unsaturated fatty acids to saturated fatty acids, reducing the intake of fermentable organic matter and fibre digestion (Alvarez-Hess *et al.*, 2019).

*Methanobrevibacter* abundance was also negatively correlated with vitamin B12 intake. As vitamin B12 is solely found in animal products (meat, fish, but also eggs and milk products), this association was considered as indicative of vegetarianism, and this was statistically confirmed (vitamin B12 intake was negatively correlated with vegetarianism in our study, R=−0.449, p=0.013, Spearman correlation, Supplementary Table 5).

Another important finding, which was confirmed by the results of various analyses we carried out, was the keystone role of formate in methane emission. Notably, formate and vitamin B12 (cobalamin) metabolism are closely connected in humans. Cobalamin deficiency was associated with increased formate concentrations in urine and plasma (in rats, (MacMillan *et al.*, 2018)), due to the so-called methyl-folate trap (HERBERT and ZALUSKY, 1962; Scott and Weir, 1981; Lamarre *et al.*, 2013). Under these conditions, the cytosolic folate accumulates as 5-methyl-THF (thus reducing the concentration of THF), which impedes the incorporation of formate into the folate pool, and results in formate accumulation. In general, replenishing the THF pool also involves ALDH1L1 (10-formyltetrahydrofolate dehydrogenase), an enzyme involved in formate oxidation, which converts 10-formyl-THF to THF and CO_2_. Notably, an association between the Christensenellaceae/*Methanobrevibacter* abundance and the abundance of a certain SNP (rs2276731) in the ALDH1L1 gene was observed when genetic correlations with microbiome profiles were analysed in a large UK twin study (Goodrich *et al.*, 2016). SNP rs2276731 is characterised by a nucleotide exchange towards C (instead of G, T) in approx. 17% of the population (*rs2276731 RefSNP Report - dbSNP - NCBI*, 2020). This ratio is in high agreement with the percentage of methane producers observed in our (15%) and other studies (Polag and Keppler, 2019), however, a more detailed analysis of this complex relationship still needs to be carried out.

Based on these considerations and also the fact that we could measure an increased formate concentration in stool samples and observe an increased abundance of genes within the formate dehydrogenase cluster in HE samples (HE: 0.05%; LE: 0.03% of all genes), we conclude that formate represents a keystone metabolite in this entire process.

*Methanobrevibacter smithii* is highly specialised to perform methanogenesis on H_2_/CO_2_ and formate compounds, and the ability to consume formate appears to be an important specialisation displayed by methanogens in the human gastrointestinal tract (Chibani *et al.*, 2020). This hypothesis is supported by the observation that *M. smithii* upregulates formate utilisation gene clusters in syntrophic relationships (Samuel *et al.*, 2007), and methano-archaeal adhesin-like proteins are expressed differently in response to formate, indicating that the physical relationship with bacterial partners changes when different amounts of different metabolites are available (Hansen *et al.*, 2011).

The findings of this study are based on a relatively small sample size and a homogenous study group (e.g. neither elderly persons nor children were recruited), and thus no general conclusions can be drawn regarding the impact of methanogen presence on aging, health status, or obesity. Future studies are needed to collect data from more variable study groups and to examine the longitudinal dynamics of the HE microbiome in more detail in terms of its correlation with additional parameters (e.g. blood metabolites, concentration of H_2_ and CO_2_).

### Conclusions

At this point it appears too early to ask how gastrointestinal methanogenesis impacts the host and whether the presence and activity of methanogens could contribute to health or disease. However, higher formate levels (herein correlated with increased methane emission) correlated with positive foetal development, T-cell activation, a lean phenotype, and cardiovascular function (Pietzke, Meiser and Vazquez, 2020). Although we lack detailed data on this metabolite-microbiome interplay, our study and its results re-emphasize the importance of archaeome activity in the human body. This activity serves as an important mirror, modulator, and regulator of the microbiome and overall body processes.

## Methods

### Key Resources Table

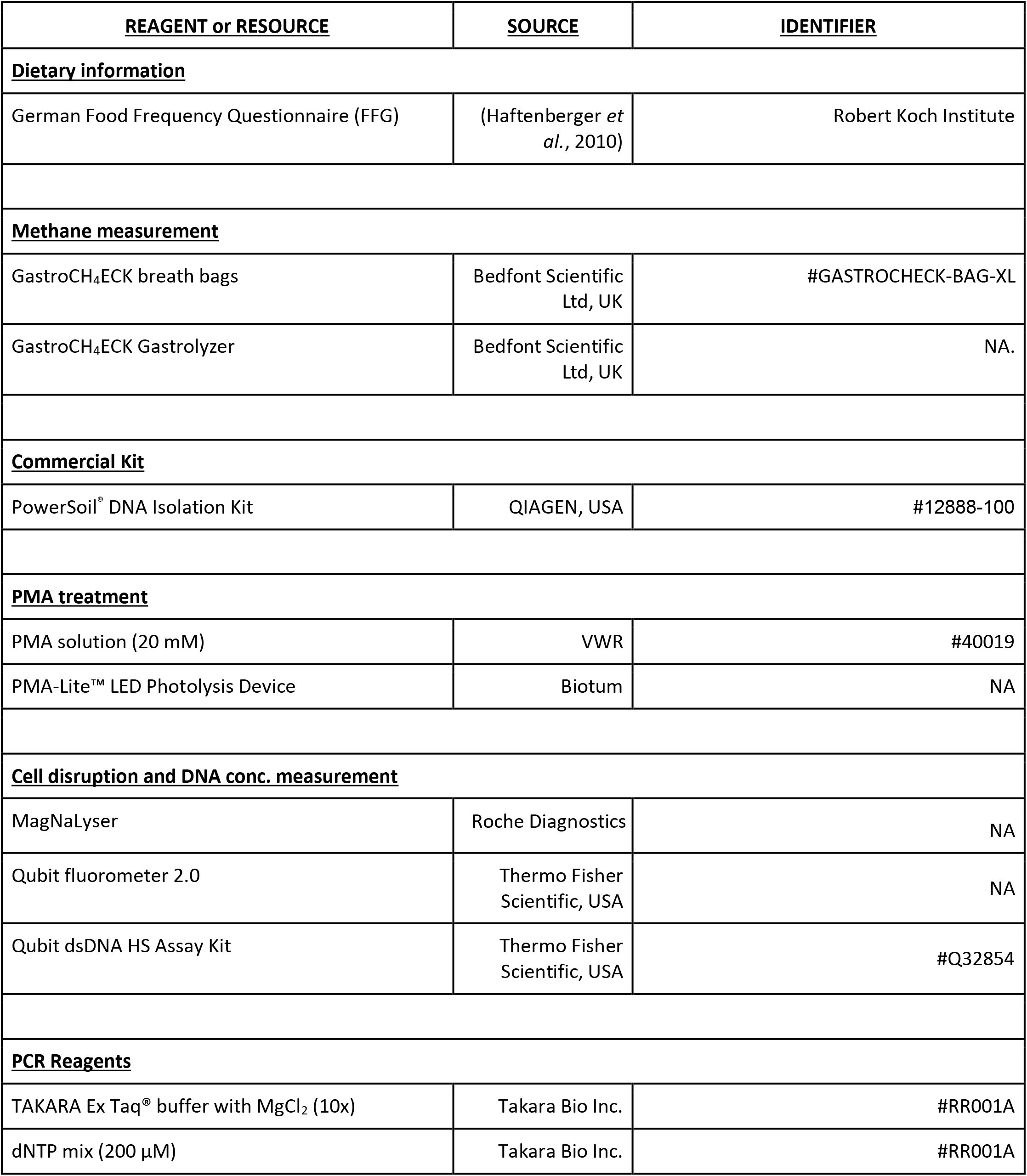
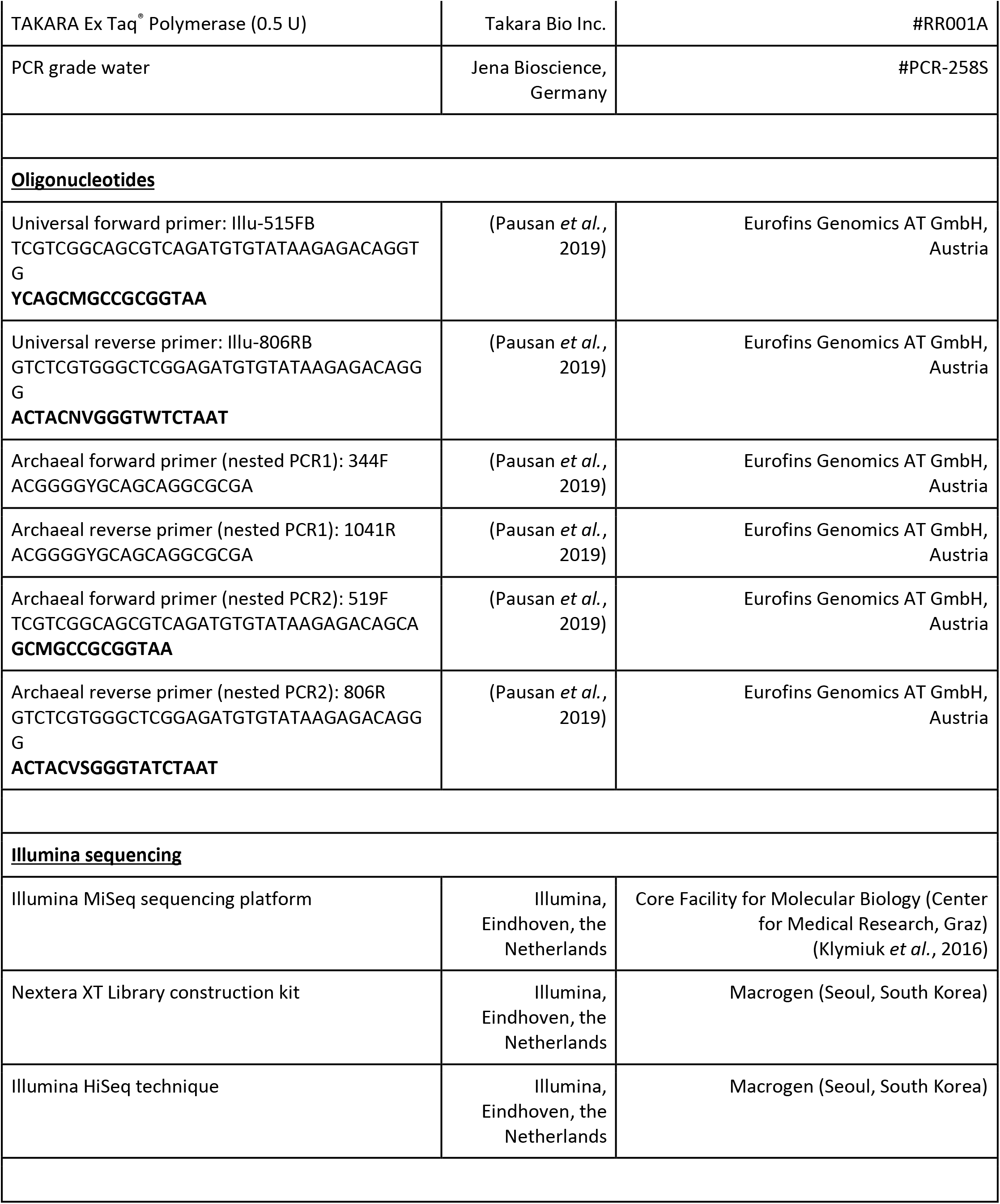
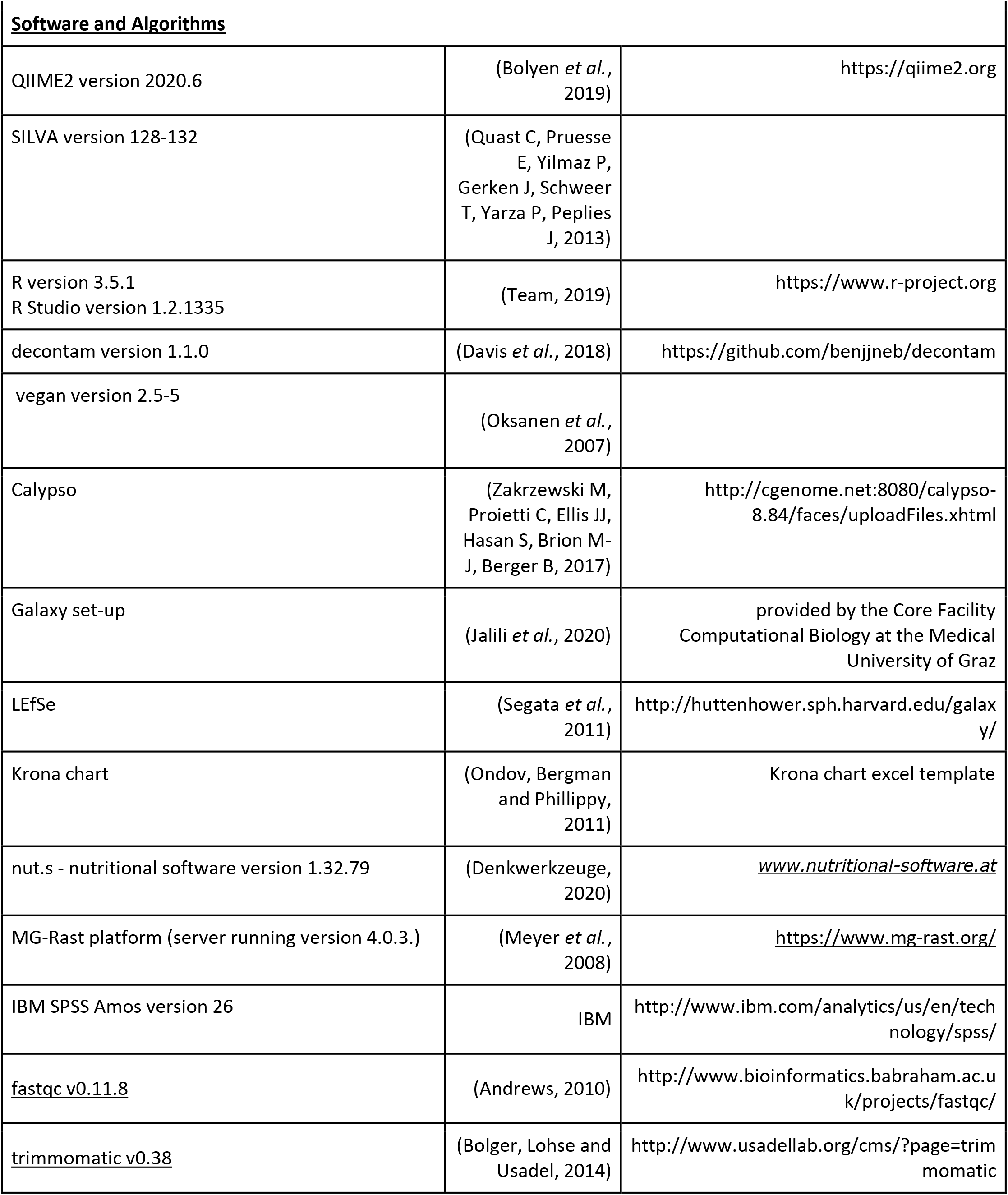
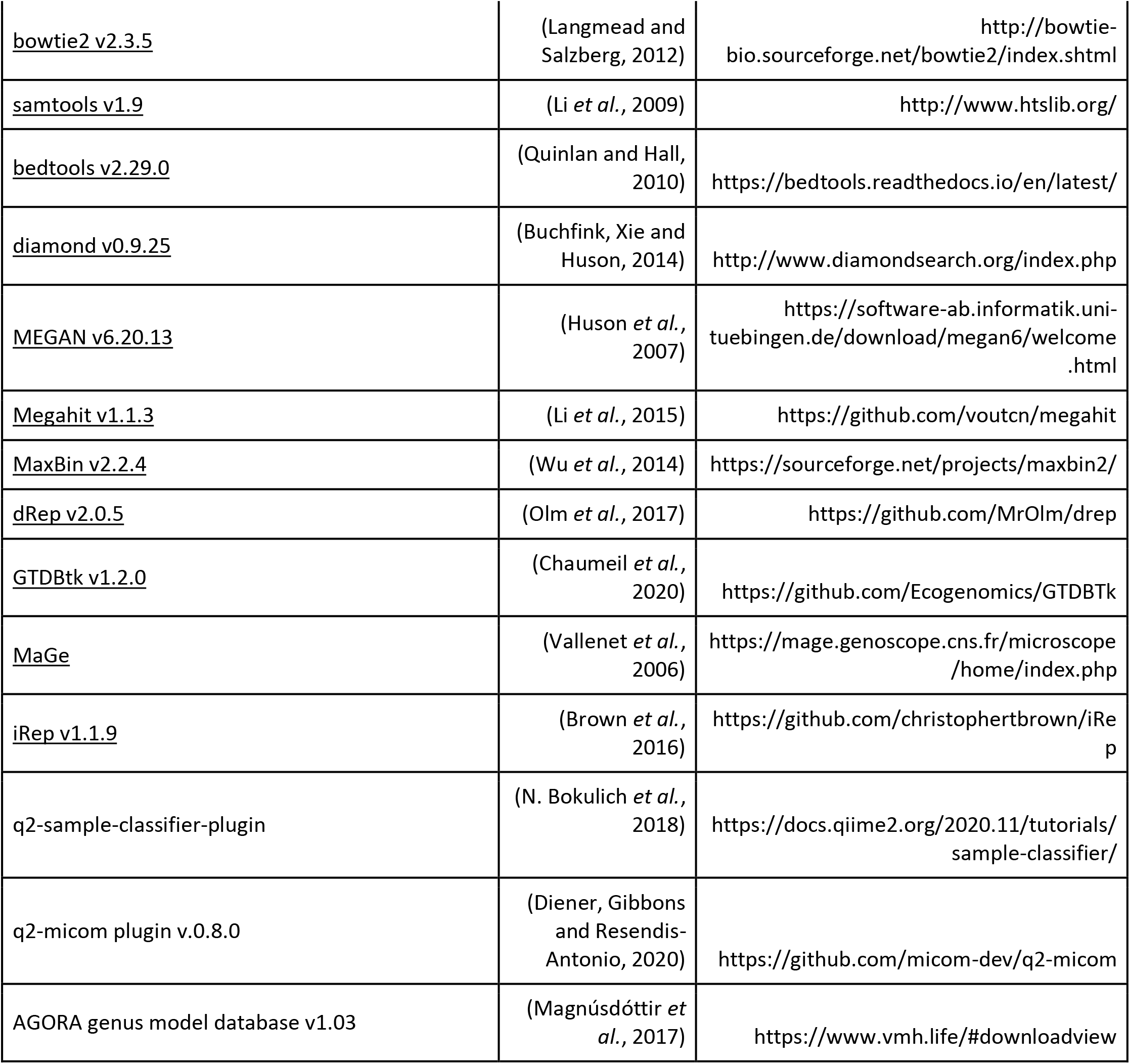

### PCR conditions

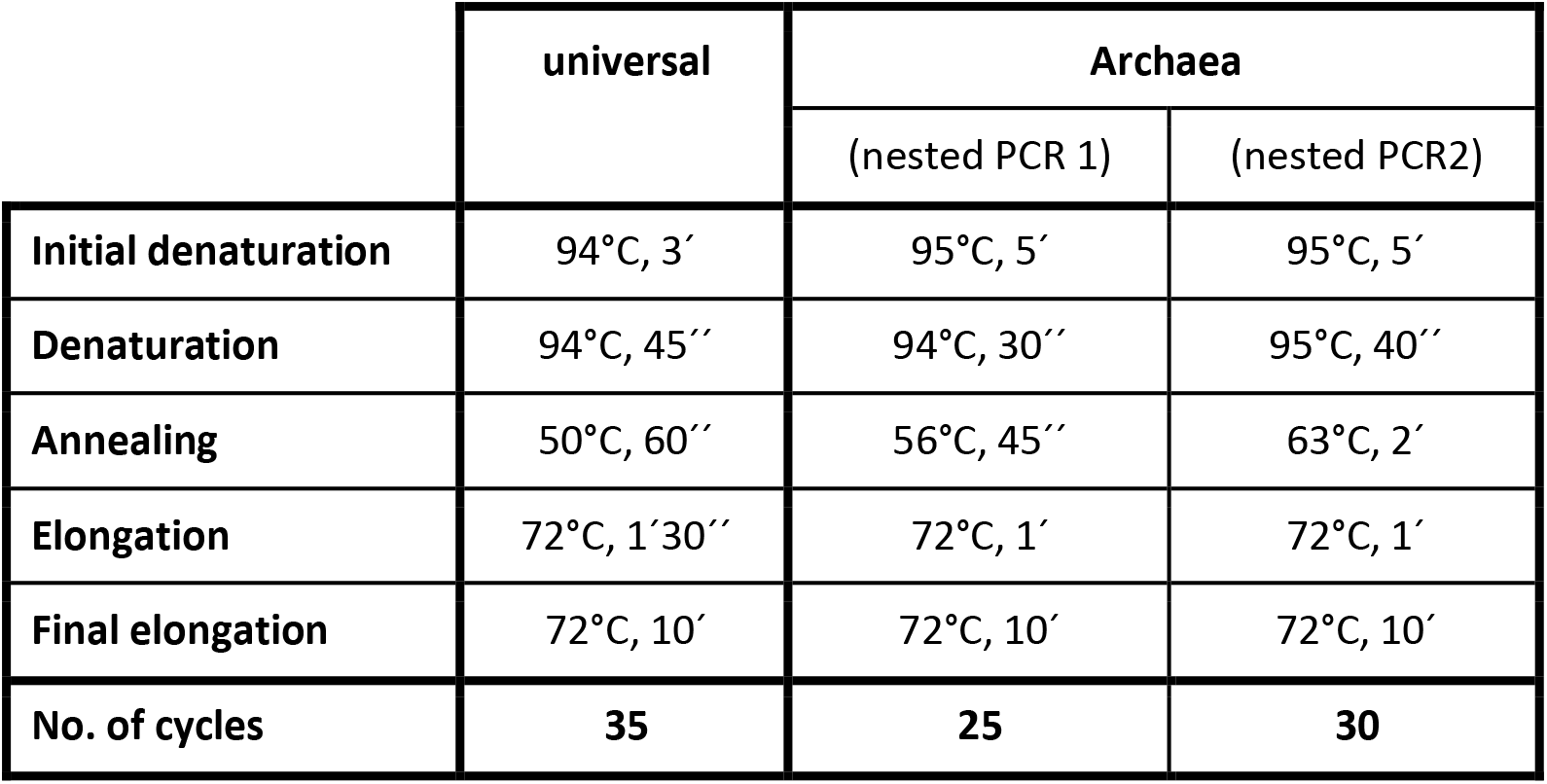

### Contact for Reagent and Resource Sharing

Further information and requests for resources should be directed to and will be fulfilled by the Lead Contact, Christine Moissl-Eichinger.

### Subject Details

#### n=100

One-hundred participants between 18-37 years were recruited at the University of Graz. Following exclusion criteria were set: smoker, left-handers, intake of antibiotics and probiotics within the last 3 months before sampling and neurological, psychiatric or internal diseases. The study was evaluated and approved according to the Declaration of Helsinki by the local ethics committee of the University of Graz (EK-Nr. GZ. 39/44/63 ex 2017/18). Before participation, all participants signed an informed consent.

## Method Details

### Methane measurement

All volunteers were asked to inhale deeply through the nose and hold their breath for 15 s before complete exhalation into the GastroCH_4_ECK breath bags (Bedfont Scientific Ltd, UK) via the mouth. Breath was collected on the same day as the stool sample in the morning before brushing their teeth and eating breakfast. Methane in the breath was measured by GastroCH4ECK Gastrolyzer (Bedfont Scientific Ltd, UK). Participants with CH_4_ values above 5 ppm were stated as methane producers. With these measurements 15% of the study group (n=15) were classified as high methane emitters (Ch_4_ value ≥ 5ppm).

### Matched subset (n=30)

15 high methane emitters were matched to 15 low methane emitters by sex, age, hormonal contraception, and vegetarianism (Supplementary Table 2). All other participants were excluded in this subset.

### Nutritional Assessment

Dietary habits and food intake information of the 4 weeks before the investigation were collected by a validated food frequency questionnaire (“German Food Frequency Questionnaire (FFG)” of the Robert Koch Institute) (Haftenberger *et al.*, 2010). The diet’s nutritive composition (e.g. intake of fat, protein, magnesium, zinc, etc) and dietary diversity indices were analyzed by a specific nutrition software using food and nutritive values specific for Austria (Denkwerkzeuge, 2020).

### Sample collection, DNA extraction and amplicon sequencing

#### Collection and PMA treatment

Stool samples were collected of every participant. To make sure that we analyse intact cells, a 10% stool suspension with 0.9% sodium chloride was treated with propidium monoazide (PMA) solution to mask freely accessible DNA. During PMA treatment, all steps were performed in the dark. PMA solution (final concentration: 50 uM) was added to the stool samples. Samples were vortexed briefly, incubated for 10 min on a shaker and 15 min in a PMA-Lite™ LED Photolysis Device (Biotum) afterwards. Samples were stored at −20 °C until further use.

#### DNA extraction

PMA-treated stool samples were used to extract microbial genomic DNA by using the DNeasy PowerSoil Kit (QIAGEN, USA) according to manufacturer’s protocol. Only modification was the use of MagNaLyser at 6500 rpm for 2 times 30 s instead of vortexing the samples. DNA concentration of extracted DNA was quantified via Qubit dsDNA HS Assay Kit (Thermo Fisher Scientific, USA).

#### 16S rRNA gene-based next generation sequencing (NGS) and sequence data processing

To determine the bacterial microbial diversity the variable region V4 of 16S rRNA gene was amplified using universal PCR primers 515FB and 806RB. For the archaeal set up a nested PCR approach was used, using the primer pair 344F and 1041R at the first and 519F and 806R for the second PCR. For detailed protocol and primer sequences see (Pausan *et al.*, 2019). Each PCR reaction was performed in triplicates. Triplicates were pooled after visualization in 3 % agarose gel. Fragments were sequenced using the Illumina MiSeq sequencing platform (Illumina, Eindhoven, the Netherlands) performed in cooperation with the Core Facility for Molecular Biology of the Center for Medical Research in Graz (Klymiuk *et al.*, 2016).

Raw reads were analyzed with QIIME2 (Quantitative Insights Into Microbial Ecology) version 2019.1 using DADA2 (Divisive Amplicon Denoising Algorithm) to denoise sequences (Callahan *et al.*, 2016; Bolyen *et al.*, 2019). Briefly, paired end reads were joined together before a quality check of the produced sequences was performed. Afterwards, taxonomic assignment was determined with SILVA v128 (universal approach), and SILVA v132 (archaeal approach)(Quast C, Pruesse E, Yilmaz P, Gerken J, Schweer T, Yarza P, Peplies J, 2013) as a reference database for a Naïve-Bayes classifier (N. A. Bokulich *et al.*, 2018). For phylogenetic metrics and analysis a rooted tree was generated with FastTree 2 (Price, Dehal and Arkin, 2010).

LEfSe (LDA Effect Size) (Segata *et al.*, 2011) was used to identify genomic features characterizing the differences between two given conditions. In our case, the LEfSe tool was integrated in a user-friendly Galaxy set-up provided by the Core Facility Computational Biology at the Medical University of Graz. The cladogram was created by the “Plot Cladogram” function, and further-on optimized using Inkscape (inkscape.org).

#### Controls

Extraction blanks and PCR negative controls were processed in parallel. All controls were removed using the R package decontam (Davis *et al.*, 2018) with the prevalence method and threshold set to 0.5 (https://github.com/benjjneb/decontam). Unassigned sequences mitochondrial and chloroplast signatures as well as features with zero or only one read were also removed. Remaining RSV tables (Supplementary Dataset 1-3) were processed in Calypso (Zakrzewski M, Proietti C, Ellis JJ, Hasan S, Brion M-J, Berger B, 2017) to generate RDA, Shannon, PCoA, ANOVA plots as well as networks and correlation analysis.

### BioEnv

R Studio version 1.2.1335 (2018-07-02) and R package vegan 2.5-5 (Oksanen *et al.*, 2007) was used to generate a BioEnv diagram with environmental variables (dietary information, CH4 emission, ..) with maximum correlation with microbial community dissimilarities.

### Metagenome Analysis

#### Shotgun metagenome sequencing

200 ng extracted DNA (PMA treated) of each of the 30 matched samples was sent for sequencing to Macrogen (Seoul, South Korea). Library was extracted via Nextera XT Library construction kit (Illumina, Eindhoven, the Netherlands) and sequenced using Illumina HiSeq technique (Illumina, Eindhoven, the Netherlands).

#### Metagenomics analysis via MG-Rast

Raw data was quality controlled, sequences were paired and analyzed with the open-submission data portal MG-Rast platform (server running version 4.0.3.)(Meyer *et al.*, 2008). Features with zero or one read were removed before feature tables (RefSeq and SEED) were uploaded in Calypso (Zakrzewski M, Proietti C, Ellis JJ, Hasan S, Brion M-J, Berger B, 2017).

#### Metagenome assembled genomes (MAGs)

After checking quality with fastqc (v0.11.8)(Andrews, 2010), raw shotgun reads were filtered accordingly with trimmomatic (v0.38) (Bolger, Lohse and Usadel, 2014) by using a minimal length of 50 bp and a Phred quality score of 20 in a sliding window of 5 bp. Quality filtered sequences were then mapped against the human chromosome hg19 with bowtie2 (v2.3.5) (Langmead and Salzberg, 2012) to remove sequences of the human host by retaining all unmapped reads with samtools (v1.9, settings: -b -f 12 -F 256)(Li *et al.*, 2009). Host removed forward and reverse fastq files were then extracted from sorted bam files with bedtools (v2.29.0) (Quinlan and Hall, 2010). Reads were then analyzed in a gene and genome-centric manner. For the gene-centric analysis, host removed quality filtered reads were annotated by blastX searches against the NCBInr database (release of Sep. 9th 2020) using diamond (v0.9.25) (Buchfink, Xie and Huson, 2014). Resulting m8 files were then visualized in MEGAN (v6.20.13) (Huson *et al.*, 2007). For the genome-centric analysis host removed quality filtered reads were co-assembled in Megahit (v1.1.3) (Li *et al.*, 2015) by using the preset meta-sensitive. Resulting contigs were binned with MaxBin v2.2.4 (Wu *et al.*, 2014). Further on, bins were quality scored (based on CheckM (Parks *et al.*, 2015) estimates for completeness, contamination and strain heterogeneity as well as N50 based assembly continuity) and de-replicated to pick representative MAGs (metagenome assembled genomes) with dRep (v2.0.5) (Olm *et al.*, 2017). Quality MAGs were then classified with GTDBtk (v1.2.0) (Chaumeil *et al.*, 2020). Identified key MAGs were further annotated and analyzed including gene synteny in MaGe (Vallenet *et al.*, 2006). Finally, replication rates were determined with iRep (v1.1.9) (Brown *et al.*, 2016).

### Prediction model

Raw metagenome data was used to create prediction models in QIIME2 (Caporaso *et al.*, 2010).

#### Supervised metadata classifications and regressions

The q2-sample-classifier-plugin (N. Bokulich *et al.*, 2018) was used to predict high and low methane emitters from feature table compositions. To determine accuracy by comparing predicted values the data set was randomly split by 5 into a training set (⅘) and a test set (⅕). The training set was used for the learning model including settings for optimized feature-selection, parameter tuning and K-fold cross validation based on RandomForest. The resulting sample estimator (trained classification model) was also used to predict methane emissions between the shotgun (RefSeqs) and amplicon dataset.

### Krona charts

Datasets (amplicon and metagenome) were normalized and Krona chart templates (Ondov, Bergman and Phillippy, 2011) were used to visualize the differences between HE and LE.

#### Metabolic quantification using NMR

Nuclear magnetic resonance spectroscopy (NMR) analysis was used to analyze concentrations of Acetate, Succinate, Formate, Lactate and Propionate in stool samples (PMA untreated) performed at the Gottfried Schatz Research Center for Cell Signaling, Metabolism and Aging, Molecular Biology and Biochemistry, Medical University of Graz. To quench enzymatic reactions and remove proteins, methanol-water solution was added to the stool sample (2:1), cells were lysed using a precellys homogenizer and stored at −20°C for 1 hour until further processing. Samples were centrifuged (4°C, 30 min, 17949 rcf) and supernatants were lyophilized afterwards. Samples were then mixed with 500 μl NMR buffer in D_2_O and transferred into NMR tubes. NMR was performed on an AVANCE™ Neo Bruker Ultrashield 600 MHz spectrometer equipped with a TXI probe head at 310 K and processed as described elsewhere (Alkan *et al.*, 2018).

The 1D CPMG (Carr-Purcell_Meiboom_Gill) pulse sequence (cpmgpr1d, 512 scans, 73728 points in F1, 11904.76 HZ spectral width, 512 transients, recycle delays 4 s) with water suppression using pre-saturation, was used for 1H 1D NMR experiments. Bruker Topspin version 4.0.2 was used for NMR data acquisition. The spectra for all samples were automatically processed (exponential line broadening of 0.3 Hz), phased, and referenced using TSP at 0.0 ppm using Bruker Topspin 4.0.2 software (Bruker GmbH, Rheinstetten, Germany).

Spectra pre-processing and data analysis have been carried out using the state-of-the-art data analysis pipeline (group of Prof. Jeremy Nicholson at Imperials College London) using Matlab^®^ scripts and MetaboAnalyst 4.0 (Chong *et al.*, 2018). NMR data were imported to Matlab^®^ vR2014a (Mathworks, Natick, Massachusetts, United States), regions around the water, TSP, and remaining methanol signals excluded, and to correct for sample metabolite dilution probabilistic quotient normalization (Dieterle *et al.*, 2006) was performed.

Stated concentrations correspond to normalized concentrations after probabilistic quotient normalization.

#### Metabolic predictions

Potential metabolites were predicted with the q2-micom plugin (v. 0.8.0, (Diener, Gibbons and Resendis-Antonio, 2020). All analysis were conducted with the AGORA genus model database (v1.03) (Magnúsdóttir *et al.*, 2017) and covered the entire dataset (n=100) and the matched dataset (n=30) as well as all and selected key features. In addition, the standard western diet gut medium was adapted (with provided jupyter notebooks from the developers) according to measured nutrients to provide a per sample diet model as well. No abundance cutoff was used for all and selected features. In addition, a leave one out strategy was included for selected features to determine the behaviour of the established metabolic models in absence of a potential microbial key-player. The growth simulation was performed with individual settings for the tradeoff between community growth rate and individual taxon growth rate. This pressure to the model was determined by an evaluation of the tradeoff from 0-1 (zero to maximum enforced growth) and was set between 0.1 and 0.7 accordingly (all features and selected features respectively). Resulting growth rates could be partly verified with calculated replication rates using iRep of representative key MAGs. Subsequent visualizations and analysis included potential metabolite consumptions, growth niches, and metabolite fluxes in dependence of measured methane emissions. Finally, a minimal medium was determined for selected key features of matched samples.

### Quantification and Statistical Analysis

#### Statistical test on metadata, metabolomics and amplicon data

Statistical tests (Spearman rho’s and Pearson’s correlation) were performed using IBM SPSS Amos version 26. Different parameters were checked for normal distribution. Correlations were calculated based on distribution of the compared parameters via Spearman’s rho and Pearson’s correlation, respectively. In the manuscript, non-corrected p values were used to describe specific trends, however Bonferroni corrected p values can be found in Supplementary Table 5.

## Supporting information

Supplementary Item 1

Supplementary Item 2

Supplementary Item 3

Supplementary Item 4

Supplementary Item 5

Supplementary Item 6

Supplementary Item 7

Supplementary Item 8

Supplementary Item 9

Supplementary Item 10

Supplementary Item 11

Supplementary Item 12

Supplementary Item 13

## Data and Software Availability

Raw sequencing data obtained from amplicon-based sequencing and metagenomics sequencing data (technical sequences including adaptor sequences, linker sequences and barcode sequences as well as human reads were removed) used in this paper can be found in the European Nucleotide Archive (ENA): PRJEB41867.

## Acknowledgments

We appreciate the financial support provided by FWF through grant P 32697 given to CME and through grant KLI 639 given to CME, VS, and FF. We are thankful to all participants of the study for providing sample material and questionnaires.

The authors acknowledge computational resources of the MedBioNode at the Medical University Graz and the support of the ZMF Galaxy Team: Core Facility Computational Bioanalytics, Medical University of Graz, funded by the Austrian Federal Ministry of Education, Science and Research, Hochschulraum-Strukturmittel 2016 grant as part of BioTechMed Graz. We would like to thank Christian Diener and other developers of the micom tool for their valuable input and support to predict metabolites, adapt diet models, or evaluate growth and replication rates. We appreciate critical proofreading by Marcus Blohs.

## Author Contributions

CK performed sampling, DNA extraction, PCR, data analysis (microbiome, metagenome, metabolome, correlation analysis), produced most of the display items and wrote the manuscript. FF, MW and CS were responsible for recruitment, sample and questionnaire/cohort metadata collection. AM supported data analysis, performed metabolic prediction, genome centric analysis, supervised learning methods, and wrote the manuscript. SL and SH performed analysis of the dietary information, provided dietary indices and contributed to manuscript writing. CH and IB provided valuable contributions on the study design and research questions, and critically read the manuscript. CH supported methane breath measurements. FF, VS contributed to manuscript writing. VS and CME initiated this project and were responsible for the study design. CME supervised all activities, performed analyses (e.g. LefSe cladogram), and wrote the manuscript. All authors read and corrected the manuscript.

## Declaration of Interests

The authors declare no conflict of interest

## Supplementary Figures

**Supplementary Figure 1. Bubble plots of the 50 most abundant taxa based on the “universal” approach of 16S rRNA gene sequencing. A.** Microbiome profiles of the whole study cohort (n=100). **B.** Microbiome profiles of matched study subset (n=30). **AI/BI.** Phylum level. **AII/BII.** Genus level. Christensenellaceae_R7_group and *Methanobrevibacter* are highlighted.

**Supplementary Figure 2. Microbiome profiles and differences in abundances of specific taxa in HEs compared to LEs based on the “universal” approach (16S rRNA gene sequencing)**. **A.** Whole study cohort (n=100). **B.** Matched study subset (n=30). **AI/BI.** PCoA plots (RSV based); **AII/BII.** ANOVA analysis at phylum level and **AIII/BIII**at genus level on the 100 most abundant taxa. **AIV-VII/BIV-VII.** Relative abundances of individual genera.

**Supplementary Figure 3. Significant positive and negative correlation of specific taxa with emitted methane concentrations based on “universal” approach 16S rRNA gene sequencing, Spearman-based regression analysis. A.** Whole study cohort (n=100). **B.** matched study subset only (n=30). **I-V.** Significant positive correlation with emitted methane. **VI-X.** Significant negative correlation with emitted methane. (100 most abundant genera; Spearman); r=Spearman’s rho correlation coefficient (*r*s)

**Supplementary Figure 4. Co-correlation network of taxa associated with HE and LE based on “universal” approach 16S rRNA gene sequencing and Spearman’s rho**. Networks showing connections of the 100 most abundant genera of **A.** the whole study cohort (n=100), **B.** our matched study subset (n=30), **C.** HE only (n=15) and **D.** LE only (n=15). Taxa highlighted in red and blue were shown to be most significantly different in LEfSe and ANOVA analysis.

**Supplementary Figure 5. Bubble plot overview on subsystems at the highest (I.) and at functional level (II.) based on shotgun metagenome analysis.** In II., the 50 most abundant features are shown.; n=30.

**Supplementary Figure 6. Relative abundance of the most significantly different subsystems of HEs compared to LEs shown in ANOVA plots based on shotgun metagenome analysis (Subsystems). I.** At highest subsystem level (level 1) and **II.** level3. (100 most features; n=30)

**Supplementary Figure 7. Bubble plots of gut microbiome of HEs and LEs based on shotgun metagenome (RefSeq). I.** visualized at phylum level and **II.** genus level. (50 most abundant taxa; n=30)

**Supplementary Figure 8**. Significant differences were also observed at species level based on shotgun metagenome analysis (RefSeq). I. Bubble plot of the 50 most abundant taxa. II. LefSe analysis and III. ANOVA plot of 100 most abundant taxa. (n=30)

**Supplementary Figure 9. Microbial community differs significantly with respect to methane production based on shotgun metagenome analysis (RefSeq). I.** LEfSe analysis and **II.** ANOVA plot at superkingdom level. **III.** PCoA plot at RSV level. **IV.** ANOVA plot showing significant differences at phylum (100 most abundant) and **V.** genus level (50 most abundant taxa). (n=30)

**Supplementary Figure 10. Diversity and composition of the archaeal community as detected in HE and LE samples based on shotgun metagenomic analyses (RefSeq). I** Alpha diversity based on Shannon index, **II.** RDA plot, **III.** PCoA plot, **IV:** LEfSe analysis on genus level.

**Supplementary Figure 11. Archaeal network in LE and HE (blue) based on shotgun metagenomics information (RefSeq).**

**Supplementary Figure 12. Correlations with dietary intake. BIOENV analysis** showing explanatory variables triggering the microbial communities of HEs (blue) and LEs (red) based on Euclidean distances that were superimposed on a Non-metric multidimensional scaling (NMDS) plot derived from Bray-Curtis dissimilarities of HE and LE samples (stress:0.1939). *Methanobrevibacter* read counts were included as a variable for better orientation.

**Supplementary Figure 13. Metabolic network of keystone taxa in LE and HE microbiomes**. Metabolites measured in stool samples are indicated by arrows; respective increase or decrease of the median by >5% is displayed.

## Supplementary Tables

**Supplementary Table 1. Characteristics of all participants (n=100).**

**Supplementary Table 2. Characteristics of the matched subset (n=30).**

**Supplementary Table 3. Keystone taxa of high and low methane emitters (n=30).** Identified key taxa based on LEfSe analysis of the 600 most abundant genera/RSVs. Numbers in column 2 and 3 refer to Figure 6b and c.

**Supplementary Table 4**. **High quality dereplicated key MAGs** including quality and replication estimates as well as taxonomic classification according to GTDB.

**Supplementary Table 5. Correlations of different parameters (general, keystone taxa, metabolites and diet) of this study among each other.**

**Supplementary Table 6. Metabolite concentrations in high and low methane emitters (n=30).**

## Supplementary Items

**Supplementary Item 1. Krona chart based on amplicon data (universal, n=100).**

**Supplementary Item 2. Krona chart based on amplicon data (archaea, n=100).**

**Supplementary Item 3. Krona chart based on metagenomic data (SEED, n=30).**

**Supplementary Item 4. Krona chart based on metagenomic data (RefSeq, archaea and bacteria only, n=30).**

**Supplementary Item 5. Krona chart based on metagenomic data (RefSeq, archaea only, n=30).**

**Supplementary Item 6. Heatmap of amino acid flux predictions according to MICOM (universal primer: 515F-806R; n=30)**

**Supplementary Item 7. Heatmap of C1-C4 flux predictions according to MICOM (universal primer: 515F-806R; n=30)**

**Supplementary Item 8. Heatmap of complex compound flux predictions according to MICOM (universal primer: 515F-806R; n=30)**

**Supplementary Item 9. Heatmap of fat flux predictions according to MICOM (universal primer: 515F-806R; n=30)**

**Supplementary Item 10. Heatmap of nucleotide flux predictions according to MICOM (universal primer: 515F-806R; n=30)**

**Supplementary Item 11. Heatmap of other metabolite flux predictions according to MICOM (universal primer: 515F-806R; n=30)**

**Supplementary Item 12. Heatmap of sugar flux predictions according to MICOM (universal primer: 515F-806R; n=30)**

**Supplementary Item 13. Heatmap of vitamine flux predictions according to MICOM (universal primer: 515F-806R; n=30)**

## Supplementary Datasets

**Supplementary Dataset 1. Feature table amplicon data of universal approach (universal primer: 515F-806R; n=100).**

**Supplementary Dataset 2. Feature table amplicon data of archaeal approach (nested PCR: 344F-1041R, 519F-806R; (n=100).**

**Supplementary Dataset 3. Feature table metagenomic data showing functional gene information (SEED; n=30).**

**Supplementary Dataset 4. Feature table metagenomic data showing taxonomic information (RefSeq; n=30).**

**Supplementary Dataset 5. MICOM growth rate predictions (universal primer: 515F-806R; n=30).**

**Supplementary Dataset 6. MICOM metabolite flux predictions (universal primer: 515F-806R; n=30).**

**Supplementary Dataset 7. Adapted per sample diet model for MICOM (n=30).**

## Notes

**Conflict of interests:** No conflict of interest and financial disclosures.

### Competing Interest Statement

The authors have declared no competing interest.

## References

Adam, P. S., Borrel, G. and Gribaldo, S. (2019) ‘An archaeal origin of the Wood–Ljungdahl H4MPT branch and the emergence of bacterial methylotrophy.’, Nature Microbiology, 4, pp. 2155–2163. doi: https://doi.org/10.1038/s41564-019-0534-2.

Alkan, H. F. et al. (2018) ‘Cytosolic Aspartate Availability Determines Cell Survival When Glutamine Is Limiting’, Cell Metabolism. Cell Press, 28(5), pp. 706–720.e6. doi: 10.1016/j.cmet.2018.07.021.

Alonso, B. L. et al. (2017) ‘First report of human infection by Christensenella minuta, a gram-negative, strickly anaerobic rod that inhabits the human intestine’, Anaerobe. Academic Press, 44, pp. 124–125. doi: 10.1016/j.anaerobe.2017.03.007.

Alvarez-Hess, P. S. et al. (2019) ‘Effect of dietary fat supplementation on methane emissions from dairy cows fed wheat or corn’, Journal of Dairy Science. Elsevier Inc., 102(3), pp. 2714–2723. doi: 10.3168/jds.2018-14721.

Aminov, R. I. et al. (2006) ‘Molecular diversity, cultivation, and improved detection by fluorescent in situ hybridization of a dominant group of human gut bacteria related to Roseburia spp. or Eubacterium rectale’, Applied and Environmental Microbiology. American Society for Microbiology (ASM), 72(9), pp. 6371–6376. doi: 10.1128/AEM.00701-06.

Andrews, S. (2010) FastQC A Quality Control tool for High Throughput Sequence Data. Available at: http://www.bioinformatics.babraham.ac.uk/projects/fastqc/. (Accessed: 21 December 2020).

Bokulich, N. et al. (2018) ‘q2-sample-classifier: machine-learning tools for microbiome classification and regression’, Journal of Open Source Software. The Open Journal, 3(30), p. 934. doi: 10.21105/joss.00934.

Bokulich, N. A. et al. (2018) ‘Optimizing taxonomic classification of marker-gene amplicon sequences with QIIME 2’s q2-feature-classifier plugin’, Microbiome. BioMed Central Ltd., 6(1). doi: 10.1186/s40168-018-0470-z.

Bolger, A. M., Lohse, M. and Usadel, B. (2014) ‘Trimmomatic: a flexible trimmer for Illumina sequence data’, Bioinformatics. Oxford University Press, 30(15), pp. 2114–2120. doi: 10.1093/bioinformatics/btu170.

Bolyen, E. et al. (2019) ‘Reproducible, interactive, scalable and extensible microbiome data science using QIIME 2’, Nature Biotechnology. Nature Publishing Group, pp. 852–857. doi: 10.1038/s41587-019-0209-9.

Bonder, M. J. et al. (2016) ‘The effect of host genetics on the gut microbiome’, Nature Genetics. Nature Publishing Group, 48(11), pp. 1407–1412. doi: 10.1038/ng.3663.

Boros, M. et al. (2015) ‘The role of methane in mammalian physiology - Is it a gasotransmitter?’, Journal of Breath Research. Institute of Physics Publishing, 9(1). doi: 10.1088/1752-7155/9/1/014001.

Boros, M. and Keppler, F. (2019) ‘Methane production and bioactivity-A link to oxido-reductive stress’, Frontiers in Physiology. Frontiers Media S.A. doi: 10.3389/fphys.2019.01244.

Borrel, G. et al. (2020) ‘The host-associated archaeome’, Nature Reviews Microbiology. Nature Research, pp. 622–636. doi: 10.1038/s41579-020-0407-y.

Brown, C. T. et al. (2016) ‘Measurement of bacterial replication rates in microbial communities’, Nature Biotechnology. Nature Publishing Group, 34(12), pp. 1256–1263. doi: 10.1038/nbt.3704.

Buchfink, B., Xie, C. and Huson, D. H. (2014) ‘Fast and sensitive protein alignment using DIAMOND’, Nature Methods. Nature Publishing Group, pp. 59–60. doi: 10.1038/nmeth.3176.

Callahan, B. J. et al. (2016) ‘DADA2: High-resolution sample inference from Illumina amplicon data’, Nature Methods. Nature Publishing Group, 13(7), pp. 581–583. doi: 10.1038/nmeth.3869.

Calloway, D. H., Colasito, D. J. and Mathews, R. D. (1966) ‘Gases produced by human intestinal microflora’, Nature. Nature, 212(5067), pp. 1238–1239. doi: 10.1038/2121238a0.

Caporaso, J. et al. (2010) ‘QIIME allows analysis of high-throughput community sequencing data.’, Nat. Methods, 7, pp. 335–336. doi: https://doi.org/10.1038/nmeth.f.303.

Chassard, C. et al. (2010) ‘The cellulose-degrading microbial community of the human gut varies according to the presence or absence of methanogens’, FEMS Microbiology Ecology. Oxford Academic, 74(1), pp. 205–213. doi: 10.1111/j.1574-6941.2010.00941.x.

Chaumeil, P. A. et al. (2020) ‘GTDB-Tk: A toolkit to classify genomes with the genome taxonomy database’, Bioinformatics. Oxford University Press, 36(6), pp. 1925–1927. doi: 10.1093/bioinformatics/btz848.

Chibani, C. M. et al. (2020) ‘A comprehensive analysis of the global human gut archaeome from a thousand genome catalogue’, bioRxiv. Cold Spring Harbor Laboratory, p. 2020.11.21.392621. doi: 10.1101/2020.11.21.392621.

Chong, J. et al. (2018) ‘MetaboAnalyst 4.0: Towards more transparent and integrative metabolomics analysis’, Nucleic Acids Research. Oxford University Press, 46(W1), pp. W486–W494. doi: 10.1093/nar/gky310.

Davis, N. M. et al. (2018) ‘Simple statistical identification and removal of contaminant sequences in marker-gene and metagenomics data.’, Microbiome, 6(1), p. 226. doi: 10.1186/s40168-018-0605-2.

Denkwerkzeuge, D. (2020) ‘Software:nut.s science v1.32.79’. Vienna. Available at: www.nutritional-software.at.

Diener, C., Gibbons, S. M. and Resendis-Antonio, O. (2020) ‘MICOM: Metagenome-Scale Modeling To Infer Metabolic Interactions in the Gut Microbiota’, mSystems. American Society for Microbiology, 5(1). doi: 10.1128/msystems.00606-19.

Dieterle, F. et al. (2006) ‘Probabilistic quotient normalization as robust method to account for dilution of complex biological mixtures. Application in1H NMR metabonomics’, Analytical Chemistry. Anal Chem, 78(13), pp. 4281–4290. doi: 10.1021/ac051632c.

Dridi, B. et al. (2009) ‘High prevalence of Methanobrevibacter smithii and Methanosphaera stadtmanae detected in the human gut using an improved DNA detection protocol’, PLoS ONE. PLoS One, 4(9). doi: 10.1371/journal.pone.0007063.

Duncan, S. H. et al. (2006) ‘Proposal of Roseburia faecis sp. nov., Roseburia hominis sp. nov. and Roseburia inulinivorans sp. nov., based on isolates from human faeces’, International Journal of Systematic and Evolutionary Microbiology. Microbiology Society, 56(10), pp. 2437–2441. doi: 10.1099/ijs.0.64098-0.

Goodrich, J. K. et al. (2014) ‘Human genetics shape the gut microbiome’, Cell. Cell Press, 159(4), pp. 789–799. doi: 10.1016/j.cell.2014.09.053.

Goodrich, J. K. et al. (2016) ‘Genetic Determinants of the Gut Microbiome in UK Twins’, Cell Host and Microbe, 19(5), pp. 731–743. doi: 10.1016/j.chom.2016.04.017.

Haftenberger, M. et al. (2010) ‘Relative validation of a food frequency questionnaire for national health and nutrition monitoring’, Nutrition Journal. Nutr J, 9(1). doi: 10.1186/1475-2891-9-36.

Hansen, E. E. et al. (2011) ‘Pan-genome of the dominant human gut-associated archaeon, Methanobrevibacter smithii, studied in twins’, Proceedings of the National Academy of Sciences of the United States of America. National Academy of Sciences, 108(SUPPL. 1), pp. 4599–4606. doi: 10.1073/pnas.1000071108.

Herbert, V. and Zalusky, R. (1962) ‘Interrelations of vitamin B12 and folic acid metabolism: folic acid clearance studies.’, The Journal of clinical investigation. J Clin Invest, 41(6), pp. 1263–1276. doi: 10.1172/JCI104589.

Huson, D. H. et al. (2007) ‘MEGAN analysis of metagenomic data’, Genome Research. Genome Res, 17(3), pp. 377–386. doi: 10.1101/gr.5969107.

Jalili, V. et al. (2020) ‘The Galaxy platform for accessible, reproducible and collaborative biomedical analyses: 2020 update’, Nucleic acids research. NLM (Medline), 48(W1), pp. W395–W402. doi: 10.1093/nar/gkaa434.

Klymiuk, I. et al. (2016) ‘16S based microbiome analysis from healthy subjects’ skin swabs stored for different storage periods reveal phylum to genus level changes’, Frontiers in Microbiology. Frontiers Media S.A., 7(DEC). doi: 10.3389/fmicb.2016.02012.

Koskinen, K. et al. (2017) ‘First Insights into the Diverse Human Archaeome : Specific Detection of Archaea in the Gastrointestinal Tract … crossm First Insights into the Diverse Human Archaeome : Specific Detection of Archaea in the Gastrointestinal Tract’, mBio, 8(6), pp. 1–17. doi: 10.1128/mBio.00824-17.

Lamarre, S. G. et al. (2013) ‘Formate: An essential metabolite, a biomarker, or more?’, in Clinical Chemistry and Laboratory Medicine. De Gruyter, pp. 571–578. doi: 10.1515/cclm-2012-0552.

Langmead, B. and Salzberg, S. L. (2012) ‘Fast gapped-read alignment with Bowtie 2’, Nature Methods. Nat Methods, 9(4), pp. 357–359. doi: 10.1038/nmeth.1923.

Laverdure, R. et al. (2018) ‘A role for methanogens and methane in the regulation of GLP-1’, Endocrinology, Diabetes & Metabolism. Wiley, 1(1), p. e00006. doi: 10.1002/edm2.6.

Lee, J. H. and Lee, J. (2010) ‘Indole as an intercellular signal in microbial communities’, FEMS Microbiology Reviews. Blackwell Publishing Ltd, pp. 426–444. doi: 10.1111/j.1574-6976.2009.00204.x.

Lewis, W. H. et al. (2018) ‘Morphology and phylogeny of a new species of anaerobic ciliate, Trimyema finlayi n. sp., with endosymbiotic methanogens’, Frontiers in Microbiology. Frontiers Media S.A., 9(FEB). doi: 10.3389/fmicb.2018.00140.

Li, D. et al. (2015) ‘MEGAHIT: An ultra-fast single-node solution for large and complex metagenomics assembly via succinct de Bruijn graph’, Bioinformatics. Oxford University Press, 31(10), pp. 1674–1676. doi: 10.1093/bioinformatics/btv033.

Li, H. et al. (2009) ‘The Sequence Alignment/Map format and SAMtools’, Bioinformatics. Bioinformatics, 25(16), pp. 2078–2079. doi: 10.1093/bioinformatics/btp352.

MacMillan, L. et al. (2018) ‘Cobalamin deficiency results in increased production of formate secondary to decreased mitochondrial oxidation of one-carbon units in rats’, Journal of Nutrition. Oxford University Press, 148(3), pp. 358–363. doi: 10.1093/jn/nxx057.

Magnúsdóttir, S. et al. (2017) ‘Generation of genome-scale metabolic reconstructions for 773 members of the human gut microbiota.’, Nature Microbiology, 35, pp. 81–89. doi: https://doi.org/10.1038/nbt.3703.

Mahnert, A. et al. (2018) ‘The human archaeome: methodological pitfalls and knowledge gaps’, Emerging Topics in Life Sciences. Portland Press Journals portal, 2(4), pp. 469–482. doi: 10.1042/ETLS20180037.

Mand, T. D. and Metcalf, W. W. (2019) ‘Energy Conservation and Hydrogenase Function in Methanogenic Archaea, in Particular the Genus Methanosarcina’, Microbiology and Molecular Biology Reviews. American Society for Microbiology, 83(4). doi: 10.1128/mmbr.00020-19.

Meyer, F. et al. (2008) ‘The metagenomics RAST server - A public resource for the automatic phylogenetic and functional analysis of metagenomes’, BMC Bioinformatics. BMC Bioinformatics, 9. doi: 10.1186/1471-2105-9-386.

Miller, T. L. et al. (1982) ‘Isolation of Methanobrevibacter smithii from human feces’, Applied and Environmental Microbiology. American Society for Microbiology (ASM), 43(1), pp. 227–232. doi: 10.1128/aem.43.1.227-232.1982.

Moissl-Eichinger, C. et al. (2018) ‘Archaea Are Interactive Components of Complex Microbiomes’, Trends in Microbiology. Elsevier Ltd, pp. 70–85. doi: 10.1016/j.tim.2017.07.004.

Nocker, A. et al. (2007) ‘Use of propidium monoazide for live/dead distinction in microbial ecology’, Applied and Environmental Microbiology. American Society for Microbiology (ASM), 73(16), pp. 5111–5117. doi: 10.1128/AEM.02987-06.

Oki, K. et al. (2016) ‘Comprehensive analysis of the fecal microbiota of healthy Japanese adults reveals a new bacterial lineage associated with a phenotype characterized by a high frequency of bowel movements and a lean body type’, BMC Microbiology. BioMed Central, 16(1), pp. 1–13. doi: 10.1186/s12866-016-0898-x.

Oksanen, J. et al. (2007) ‘The vegan package -Community ecology package’, pp. 631–637. Available at: https://scholar.google.com/citations?user=2WBRFVIAAAAJ&hl=sv#d=gs_md_cita-d&u=%2Fcitations%3Fview_op%3Dview_citation%26hl%3Dsv%26user%3D2WBRFVIAAAAJ%26citation_for_view%3D2WBRFVIAAAAJ%3Au5HHmVD_uO8C%26tzom%3D-60.

Olm, M. R. et al. (2017) ‘DRep: A tool for fast and accurate genomic comparisons that enables improved genome recovery from metagenomes through de-replication’, ISME Journal. Nature Publishing Group, 11(12), pp. 2864–2868. doi: 10.1038/ismej.2017.126.

Ondov, B. D., Bergman, N. H. and Phillippy, A. M. (2011) ‘Interactive metagenomic visualization in a Web browser’, BMC Bioinformatics. BMC Bioinformatics, 12. doi: 10.1186/1471-2105-12-385.

Parks, D. H. et al. (2015) ‘CheckM: Assessing the quality of microbial genomes recovered from isolates, single cells, and metagenomes’, Genome Research. Cold Spring Harbor Laboratory Press, 25(7), pp. 1043–1055. doi: 10.1101/gr.186072.114.

Pausan, M. R. et al. (2019) ‘Exploring the Archaeome: Detection of Archaeal Signatures in the Human Body’, Frontiers in Microbiology. Frontiers Media S.A., 10. doi: 10.3389/fmicb.2019.02796.

Pietzke, M., Meiser, J. and Vazquez, A. (2020) ‘Formate metabolism in health and disease’, Molecular Metabolism. Elsevier GmbH, pp. 23–37. doi: 10.1016/j.molmet.2019.05.012.

Pimentel, M. et al. (2006) ‘Methane, a gas produced by enteric bacteria, slows intestinal transit and augments small intestinal contractile activity’, American Journal of Physiology - Gastrointestinal and Liver Physiology. Am J Physiol Gastrointest Liver Physiol, 290(6). doi: 10.1152/ajpgi.00574.2004.

Polag, D. and Keppler, F. (2019) ‘Global methane emissions from the human body: Past, present and future’, Atmospheric Environment. Elsevier Ltd, 214, p. 116823. doi: 10.1016/j.atmosenv.2019.116823.

Price, M. N., Dehal, P. S. and Arkin, A. P. (2010) ‘FastTree 2 - Approximately maximum-likelihood trees for large alignments’, PLoS ONE. PLoS One, 5(3). doi: 10.1371/journal.pone.0009490.

Quast C, Pruesse E, Yilmaz P, Gerken J, Schweer T, Yarza P, Peplies J, G. F. (2013) ‘The SILVA ribosomal RNA gene database project: improved data processing and web-based tools’, Nucleic Acids Res 41, 41, pp. D590–D596.

Quinlan, A. R. and Hall, I. M. (2010) ‘BEDTools: A flexible suite of utilities for comparing genomic features’, Bioinformatics. Bioinformatics, 26(6), pp. 841–842. doi: 10.1093/bioinformatics/btq033.

rs2276731 RefSNP Report - dbSNP - NCBI (2020). Available at: https://www.ncbi.nlm.nih.gov/snp/rs2276731) (Accessed: 21 December 2020).

Ruaud, A. et al. (2020) ‘Syntrophy via interspecies H2 transfer between christensenella and methanobrevibacter underlies their global cooccurrence in the human gut’, mBio. American Society for Microbiology, 11(1). doi: 10.1128/mBio.03235-19.

Samuel, B. S. et al. (2007) ‘Genomic and metabolic adaptations of Methanobrevibacter smithii to the human gut’, Proceedings of the National Academy of Sciences of the United States of America. National Academy of Sciences, 104(25), pp. 10643–10648. doi: 10.1073/pnas.0704189104.

Scott, J. M. and Weir, D. G. (1981) ‘THE METHYL FOLATE TRAP. A physiological response in man to prevent methyl group deficiency in kwashiorkor (methionine deficiency) and an explanation for folic-acid-induced exacerbation of subacute combined degeneration in pernicious anaemia’, The Lancet. Lancet, 318(8242), pp. 337–340. doi: 10.1016/S0140-6736(81)90650-4.

Segata, N. et al. (2011) ‘Metagenomic biomarker discovery and explanation’, Genome Biology. Genome Biol, 12(6). doi: 10.1186/gb-2011-12-6-r60.

Stephen, A. M. and Cummings, J. H. (1980) ‘The microbial contribution to human faecal mass’, Journal of Medical Microbiology. J Med Microbiol, 13(1), pp. 45–56. doi: 10.1099/00222615-13-1-45.

Team, R. D. C. (2019) ‘R: A Language and Environment for Statistical Computing’. Available at: https://www.r-project.org.

Vallenet, D. et al. (2006) ‘MaGe: A microbial genome annotation system supported by synteny results’, Nucleic Acids Research. Nucleic Acids Res, 34(1), pp. 53–65. doi: 10.1093/nar/gkj406.

Vojinovic, D. et al. (2019) ‘Relationship between gut microbiota and circulating metabolites in population-based cohorts’, Nature Communications. Nature Research, 10(1). doi: 10.1038/s41467-019-13721-1.

Waters, J. L. and Ley, R. E. (2019) ‘The human gut bacteria Christensenellaceae are widespread, heritable, and associated with health’, BMC Biology. BioMed Central Ltd., pp. 1–11. doi: 10.1186/s12915-019-0699-4.

Weaver, G. A. et al. (1986) ‘Incidence of methanogenic bacteria in a sigmoidoscopy population: An association of methanogenic bacteria and diverticulosis’, Gut. Gut, 27(6), pp. 698–704. doi: 10.1136/gut.27.6.698.

Wu, Y. W. et al. (2014) ‘MaxBin: An automated binning method to recover individual genomes from metagenomes using an expectation-maximization algorithm’, Microbiome. BioMed Central Ltd., 2(1). doi: 10.1186/2049-2618-2-26.

Wyckoff, E. E., Mey, A. R. and Payne, S. M. (2007) ‘Iron acquisition in Vibrio cholerae’, in BioMetals. Biometals, pp. 405–416. doi: 10.1007/s10534-006-9073-4.

Xin, L., Sun, X. and Lou, S. (2016) ‘Effects of Methane-Rich Saline on the Capability of One-Time Exhaustive Exercise in Male SD Rats’, PLOS ONE. Edited by G. López Lluch. Public Library of Science, 11(3), p. e0150925. doi: 10.1371/journal.pone.0150925.

Young, G. R. et al. (2017) ‘Reducing Viability Bias in Analysis of Gut Microbiota in Preterm Infants at Risk of NEC and Sepsis’, Frontiers in Cellular and Infection Microbiology. Frontiers Media S.A., 7(JUN), p. 237. doi: 10.3389/fcimb.2017.00237.

Zakrzewski M, Proietti C, Ellis JJ, Hasan S, Brion M-J, Berger B, K. L. (2017) ‘Calypso: a user-friendly web-server for mining and visualizing microbiome-environment interactions.’, Bioinformatics, 33, pp. 782–783.

