## Supplementary Figures for "Methane emission of humans is explained by dietary habits, host genetics, local formate availability and a uniform archaeome"

^4^ Division of [Immunology and Pathophysiology](https://online.medunigraz.at/mug_online/pl/ui/$ctx;design=ca2;header=max;lang=en;rbacId=/wborg.display?pOrgNr=14014), Medical University of Graz, Graz 8010, Austria

**Running title:** Methane emission of humans

This study was approved by the ethics committee of the University of Graz (EK-Nr. GZ. 39/44/63 ex 2017/18). Experimental protocols were approved by the ethics committee and the Department of Otorhinolaryngology of the Medical University of Graz.

**Conflict of interests:** No conflict of interest and financial disclosures.

**Key Words**: archaeome, microbiome, methanogens, methane, gut, gastrointestinal tract, metabolome, metagenome, *Methanobrevibacter*, *Christensenellaceae*

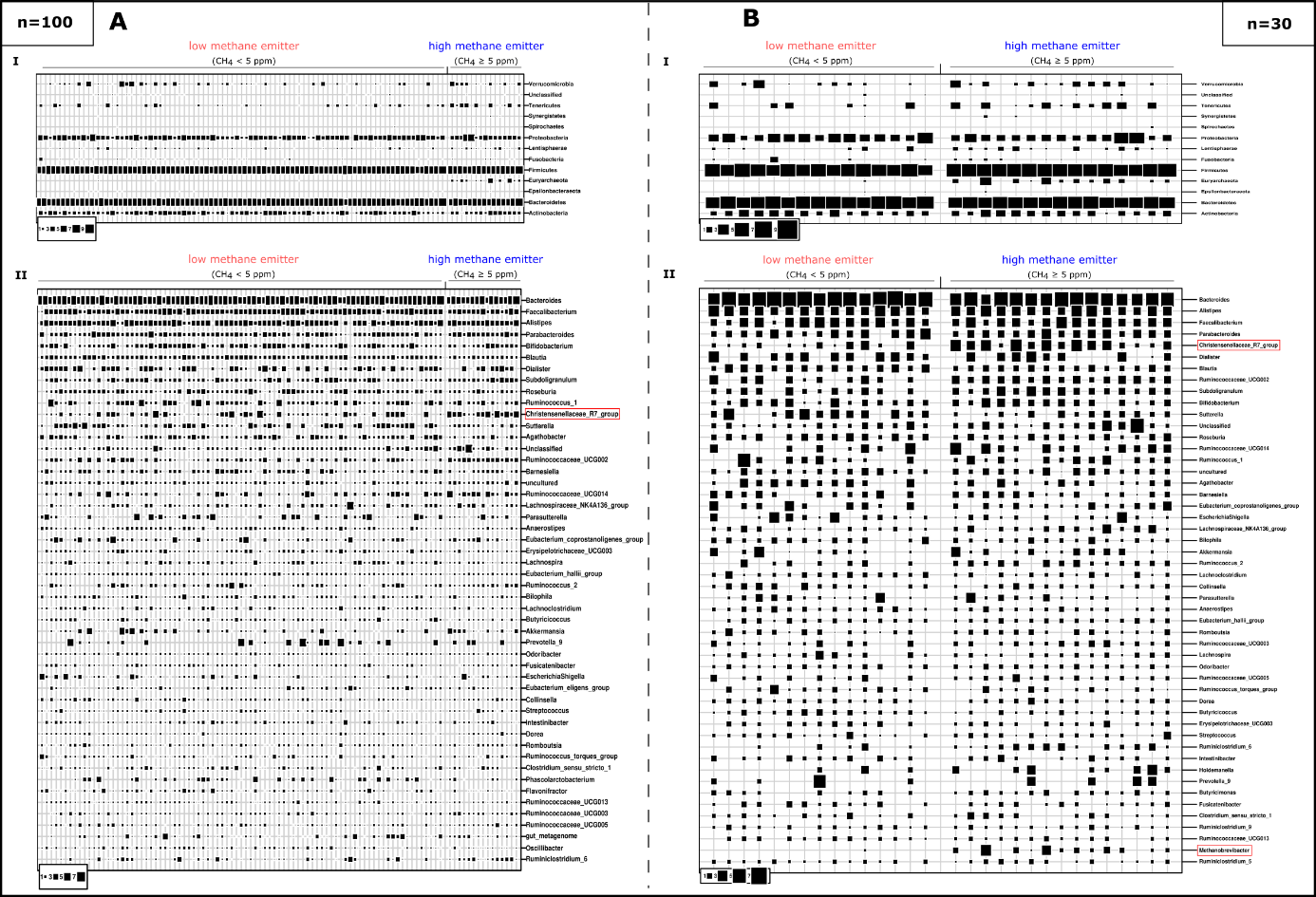

### Supplementary Figure 1. Bubble plots of the 50 most abundant taxa based on the “universal” approach of 16S rRNA gene sequencing. A. Microbiome profiles of the whole study cohort (n=100). B. Microbiome profiles of matched study subset (n=30). AI/BI. Phylum level. AII/BII. Genus level. Christensenellaceae_R7_group and *Methanobrevibacter* are highlighted.

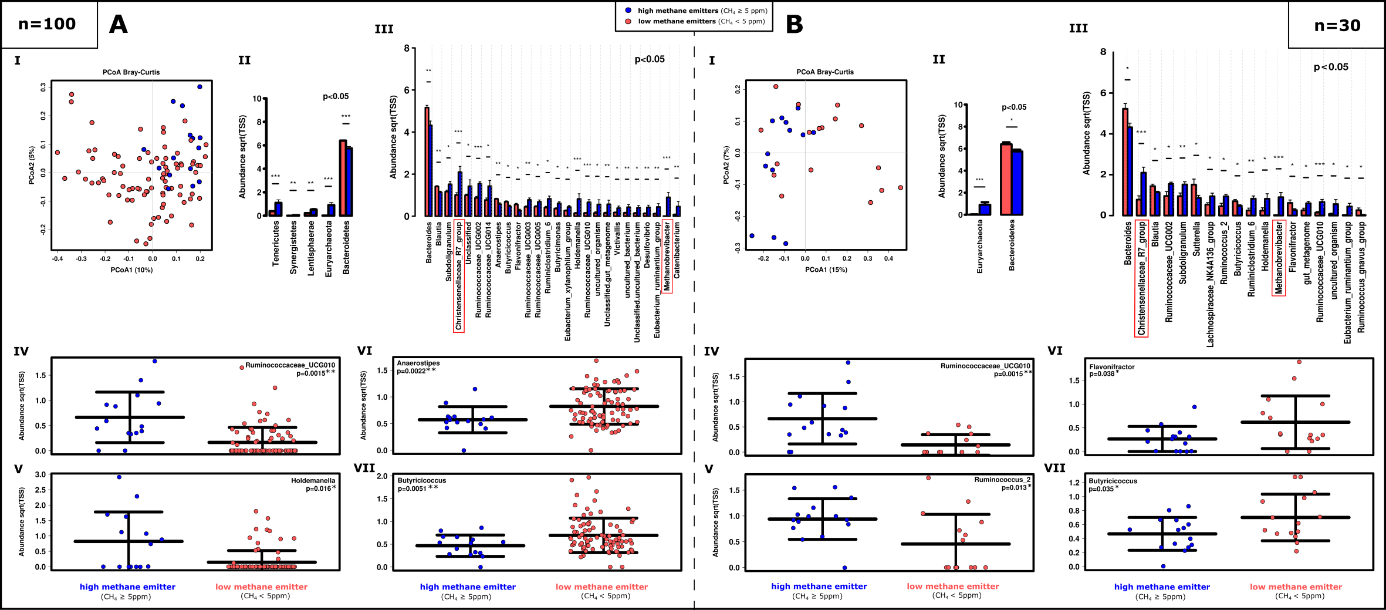

**Supplementary Figure 2. Microbiome profiles and differences in abundances of specific taxa in HEs compared to LEs based on the “universal” approach (16S rRNA gene sequencing)**. **A.** Whole study cohort (n=100). **B.** Matched study subset (n=30). **AI/BI.** PCoA plots (RSV based); **AII/BII.** ANOVA analysis at phylum level and **AIII/BIII** at genus level on the 100 most abundant taxa. **AIV-VII/BIV-VII.** Relative abundances of individual genera.

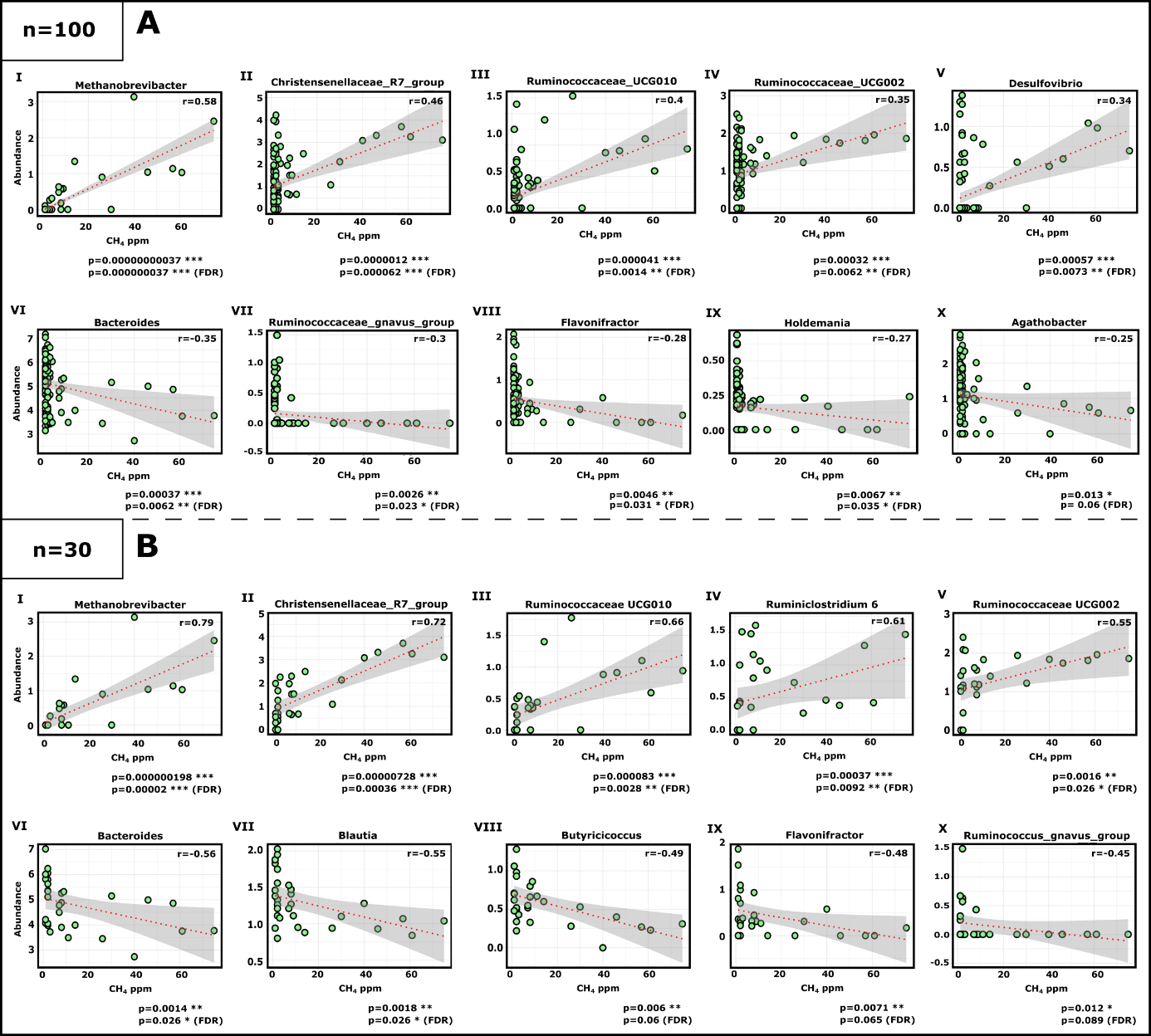

### Supplementary Figure 3. Significant positive and negative correlation of specific taxa with emitted methane concentrations based on “universal” approach 16S rRNA gene sequencing, Spearman-based regression analysis. A. Whole study cohort (n=100). B. matched study subset only (n=30). I-V. Significant positive correlation with emitted methane. VI-X. Significant negative correlation with emitted methane. (100 most abundant genera; Spearman); r=Spearman’s rho correlation coefficient (*r*s)

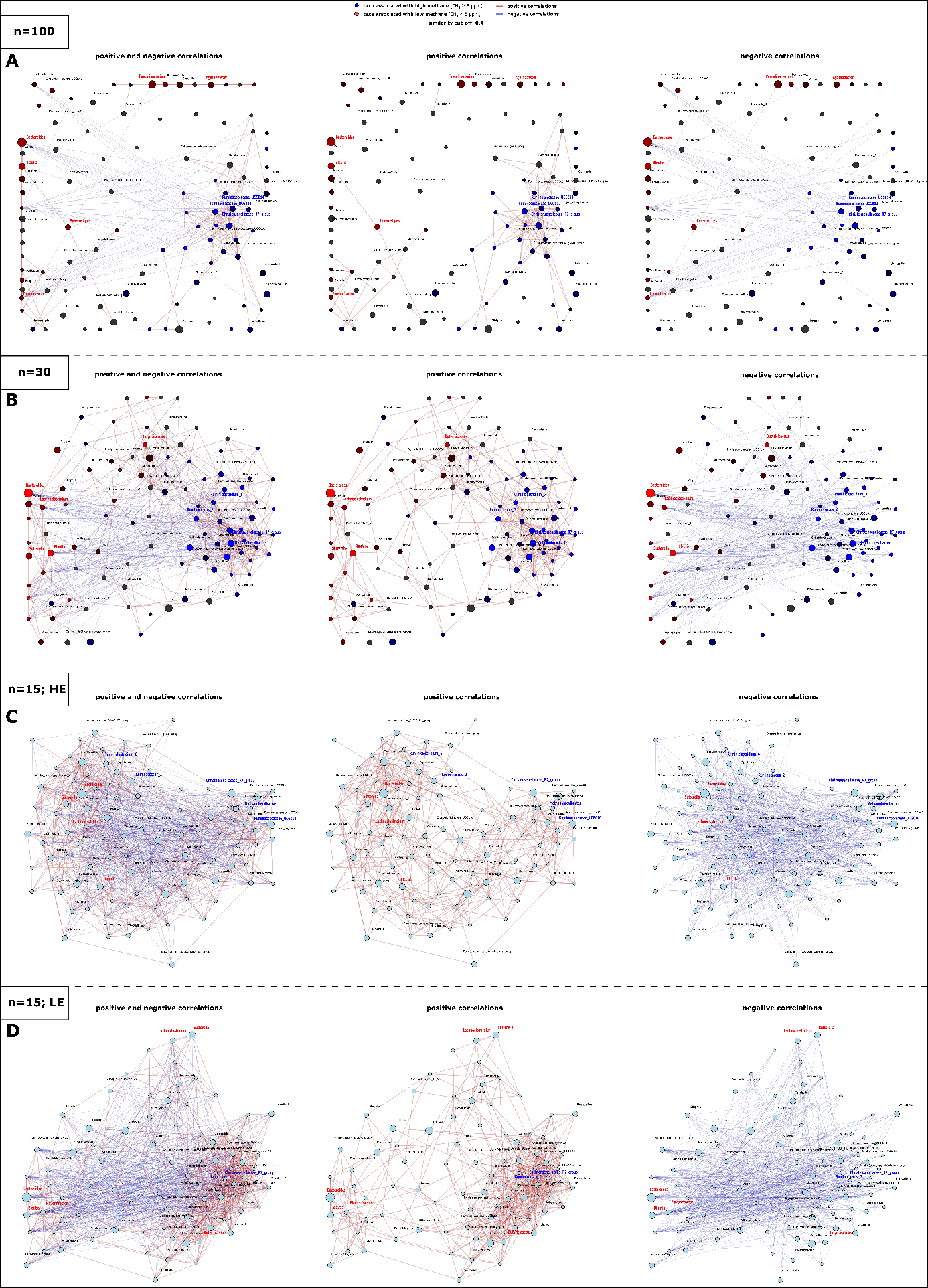

**Supplementary Figure 4. Co-correlation network of taxa associated with HE and LE based on “universal” approach 16S rRNA gene sequencing and Spearman’s rho**. Networks showing connections of the 100 most abundant genera of **A.** the whole study cohort (n=100), **B.** our matched study subset (n=30), **C.** HE only (n=15) and **D.** LE only (n=15). Taxa highlighted in red and blue were shown to be most significantly different in LEfSe and ANOVA analysis.

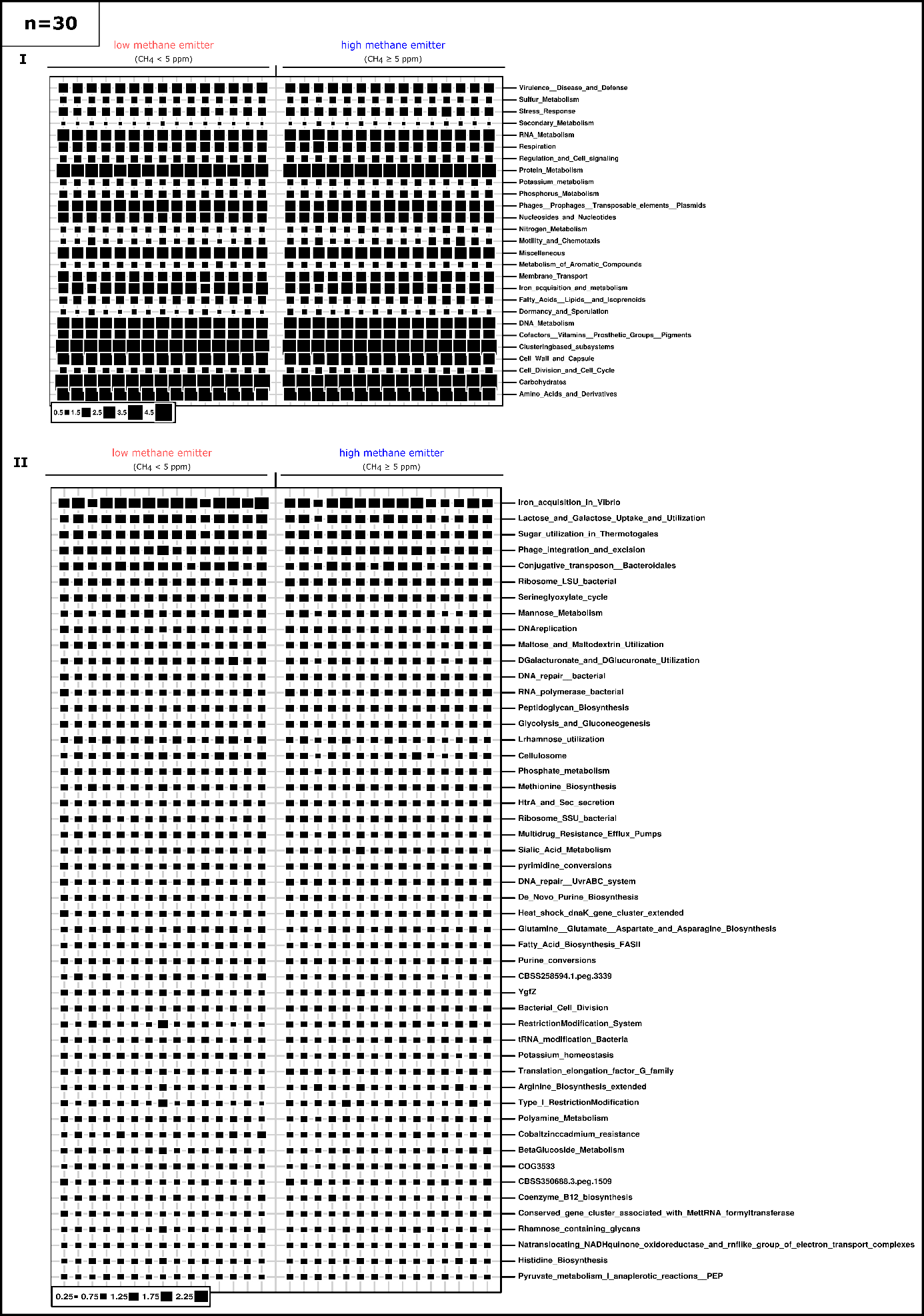

**Supplementary Figure 5. Bubble plot overview on subsystems at the highest (I.) and at functional level (II.) based on shotgun metagenome analysis.** In II., the 50 most abundant features are shown.; n=30.

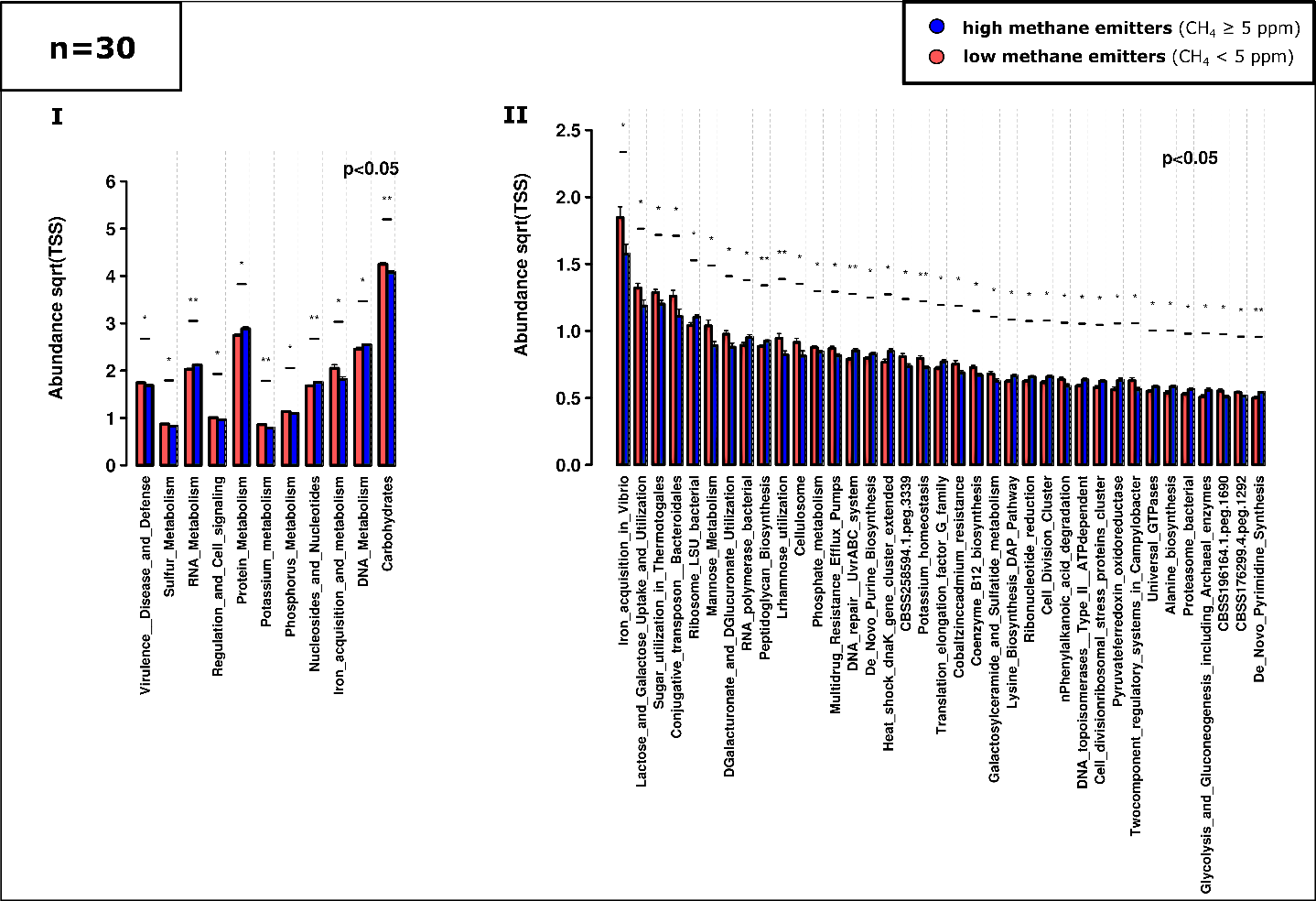

**Supplementary Figure 6. Relative abundance of the most significantly different subsystems of HEs compared to LEs shown in ANOVA plots based on shotgun metagenome analysis (Subsystems). I.** At highest subsystem level (level 1) and **II.** level3. (100 most features; n=30)

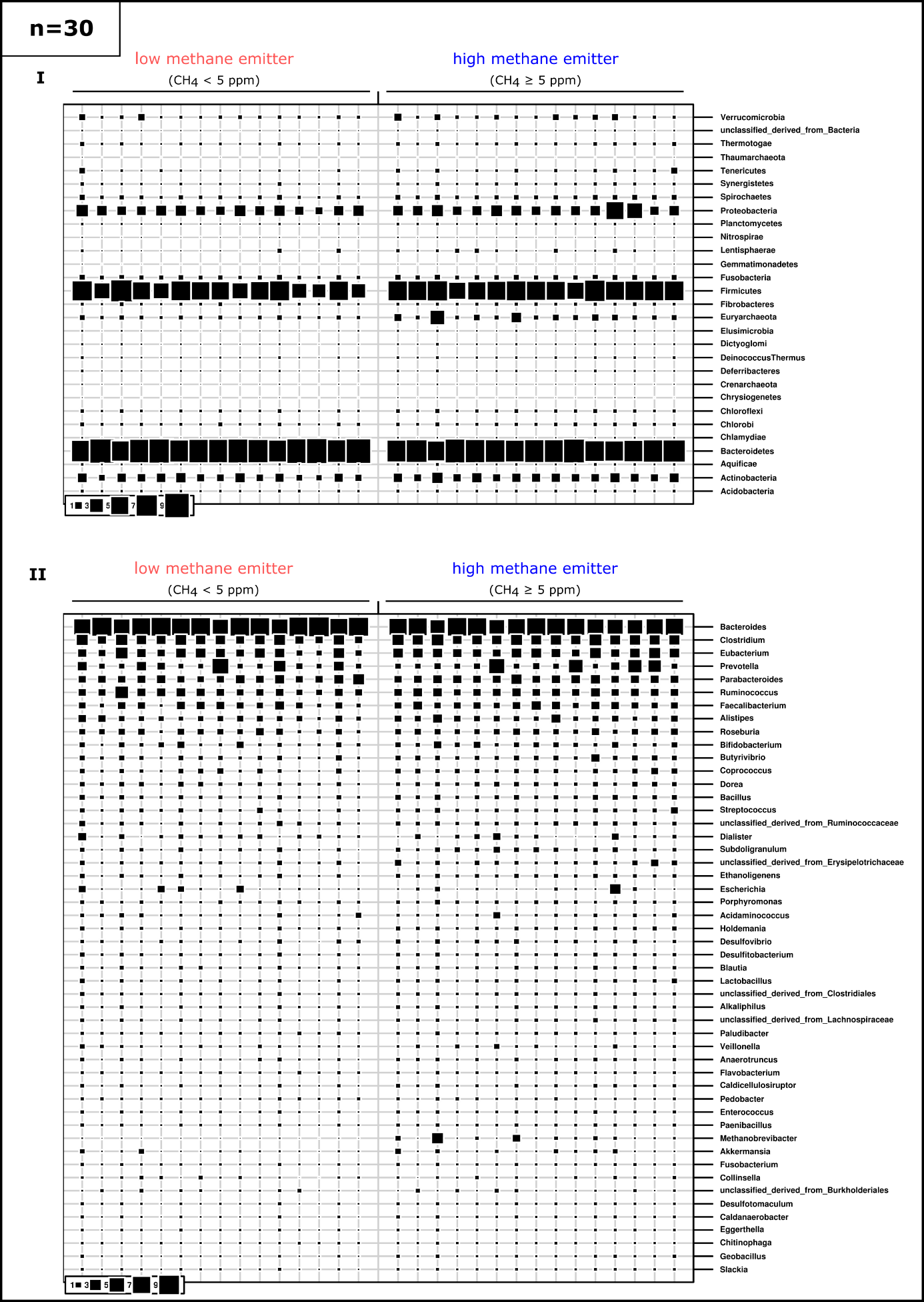

### Supplementary Figure 7. Bubble plots of gut microbiome of HEs and LEs based on shotgun metagenome (RefSeq). I. visualized at phylum level and II. genus level. (50 most abundant taxa; n=30)

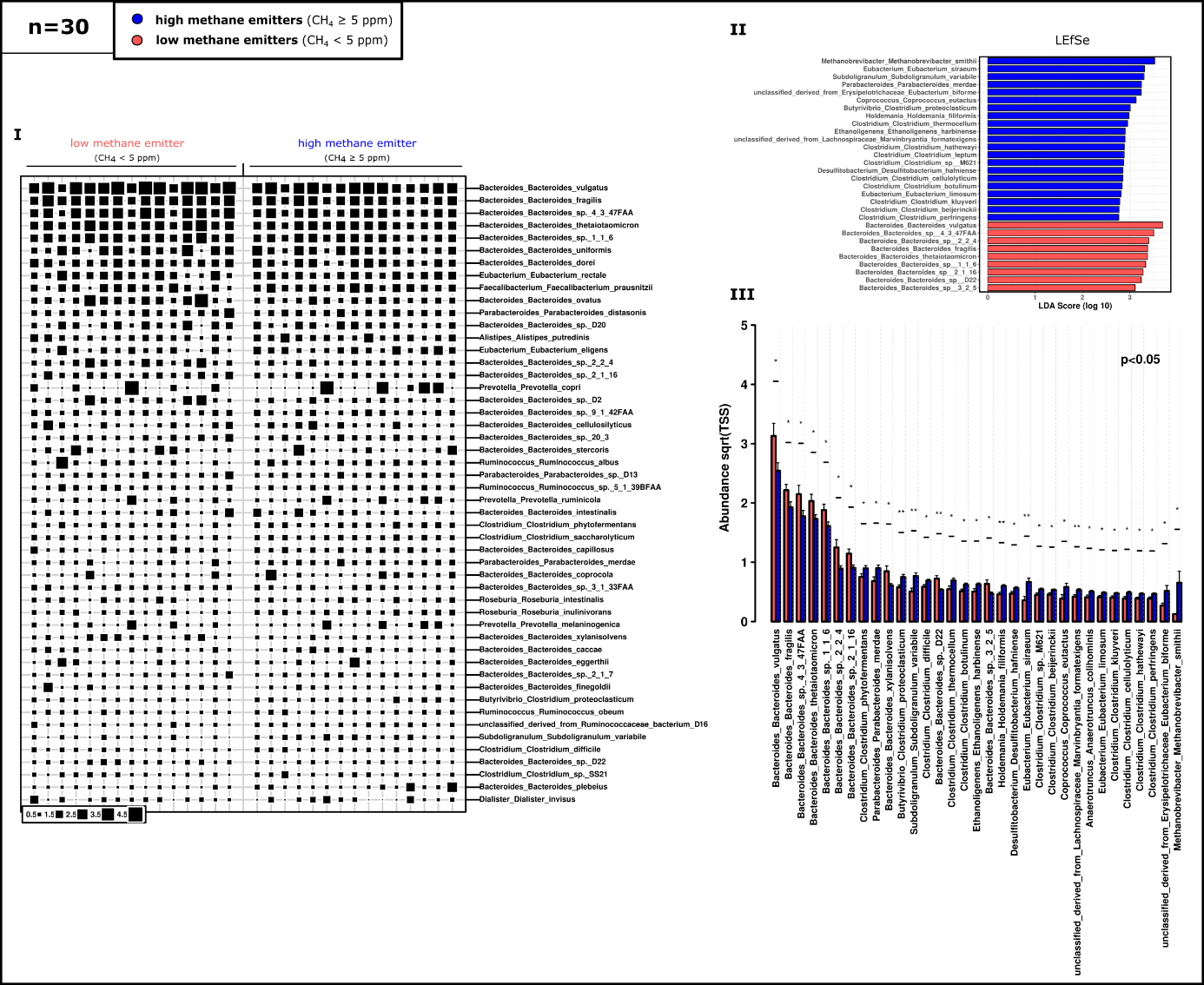
**Supplementary Figure 8**. Significant differences were also observed at species level based on shotgun metagenome analysis (RefSeq). I. Bubble plot of the 50 most abundant taxa. II. LefSe analysis and III. ANOVA plot of 100 most abundant taxa. (n=30)

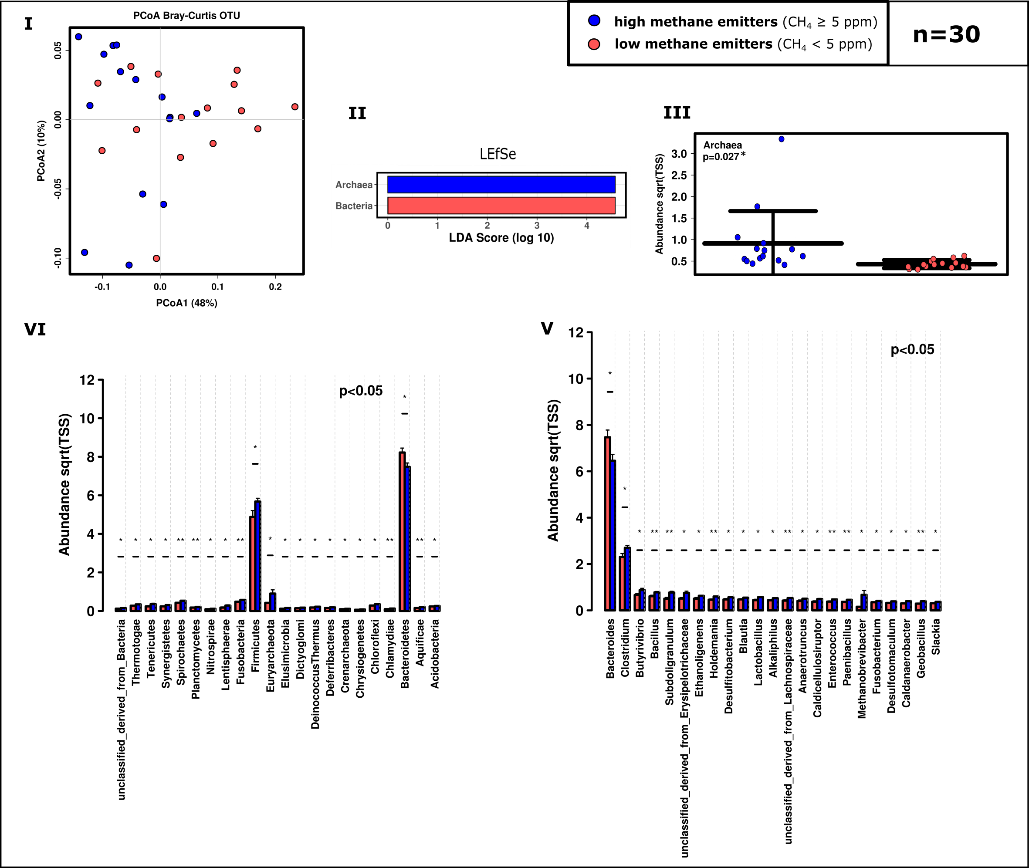

### Supplementary Figure 9. Microbial community differs significantly with respect to methane production based on shotgun metagenome analysis (RefSeq). I. LEfSe analysis and II. ANOVA plot at superkingdom level. III. PCoA plot at RSV level. IV. ANOVA plot showing significant differences at phylum (100 most abundant) and V. genus level (50 most abundant taxa). (n=30)

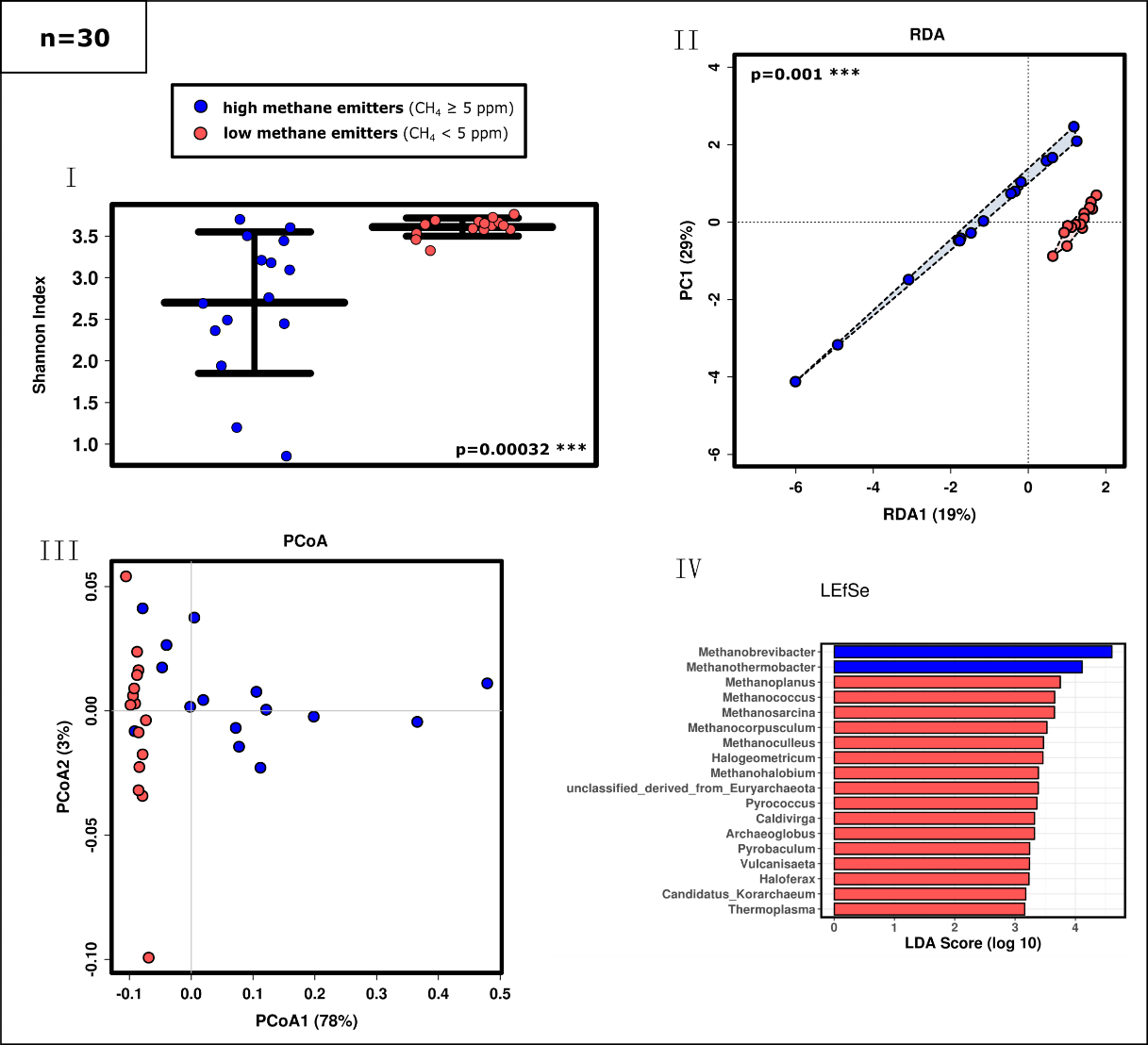

**Supplementary Figure 10.** **Diversity and composition of the archaeal community as detected in HE and LE samples based on shotgun metagenomic analyses (RefSeq). I** Alpha diversity based on Shannon index**, II.** RDA plot**, III.** PCoA plot**, IV:** LEfSe analysis on genus level**.**

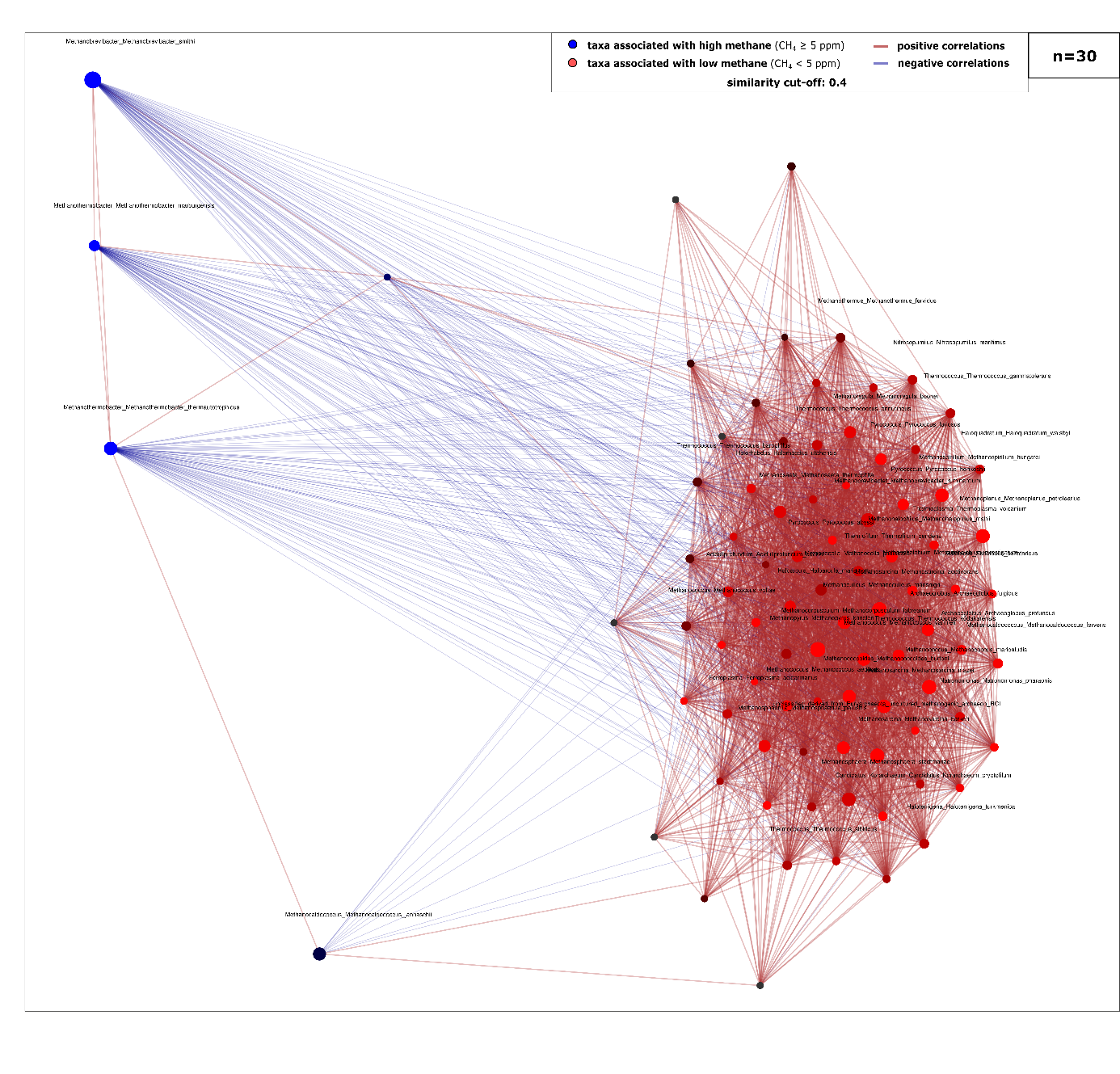

**Supplementary Figure 11. Archaeal network in LE and HE (blue) based on shotgun metagenomics information (RefSeq).**

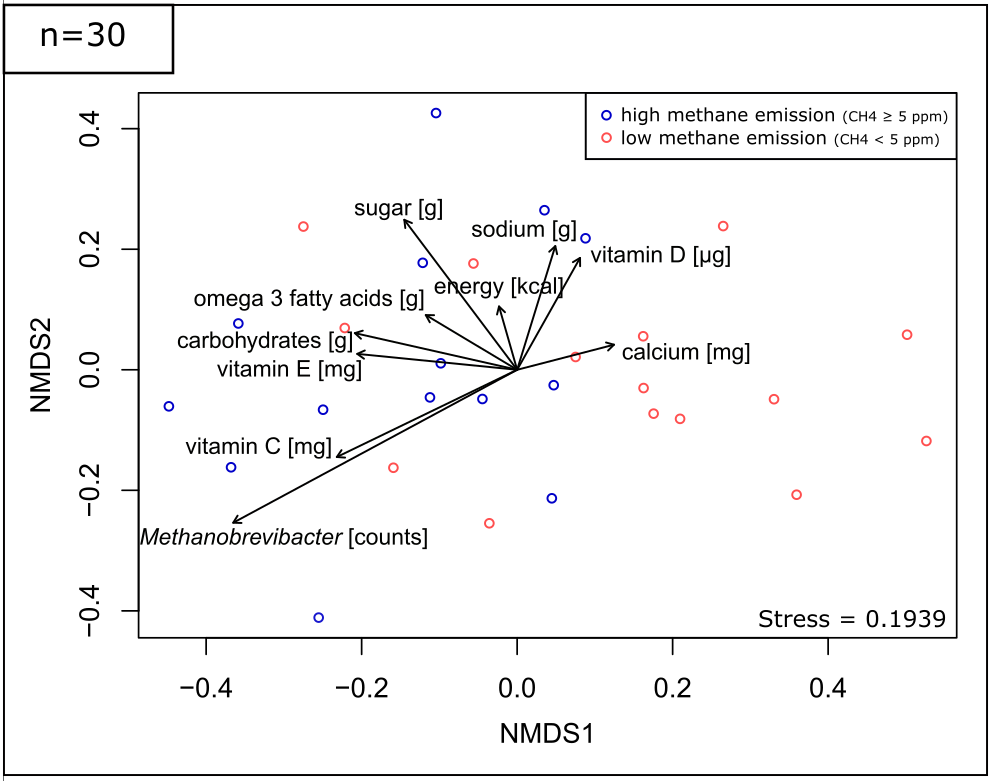

**Supplementary Figure 12.** **Correlations with dietary intake. BIOENV analysis** showing explanatory variables triggering the microbial communities of HEs (blue) and LEs (red) based on Euclidean distances that were superimposed on a Non-metric multidimensional scaling (NMDS) plot derived from Bray-Curtis dissimilarities of HE and LE samples (stress:0.1939). *Methanobrevibacter* read counts were included as a variable for better orientation.

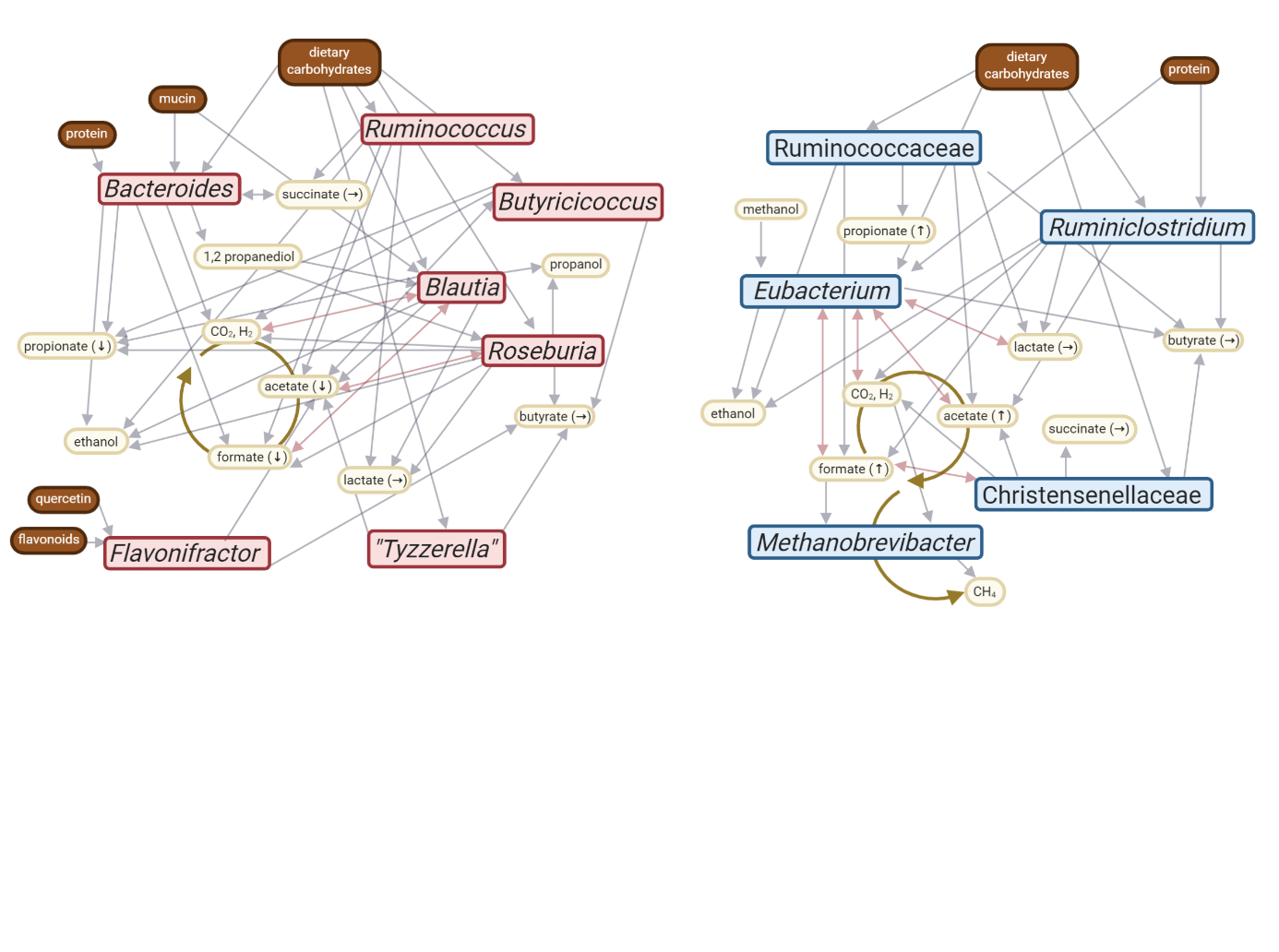

**Supplementary Figure 13.** **Metabolic network of key-stone taxa in LE and HE microbiomes**. Metabolites measured in stool samples are indicated by arrows; respective increase or decrease of the median by >5% is displayed.
