## Supplementary Item 1 for "Methane emission of humans is explained by dietary habits, host genetics, local formate availability and a uniform archaeome"

Javascript must be enabled to view this page.

magnitude

HE
LE

 100
 99.9999999999997

 98.0984828192295
 99.9973481916194

 50.1021787089066
 45.1173068234089

 42.0515799326434
 38.2555996503692

 41.9980075904184
 38.2497860704585

 22.4970730841116
 20.0831647905122

 1.68542200258231E-02
 9.63830353618647E-03

 1.12411628207945
 1.09698173210004

 3.48972644998962
 1.37950131723482

 .635644869545327
 .10178864475253

 .257327823608549
 .413733103645931

 .298861437243613
 .20235337794491

 .262745251473992
 .37400697425604

 2.59524891576201
 1.25639621280966

 .563412498006086
 .454581151965959

 .261240410400258
 .266710727482832

 .145969584152218
 .388183949827785

 6.35042933115834E-02
 .153753889743927

 .604645143426403
 .378596642606605

 2.49803618239878E-02
 4.81405213659261E-02

 4.42423275677856
 6.05540443647542

 .642567138484505
 .377270738416442

 1.56503471668357E-02
 7.08848778587259E-03

 0
 1.72877507871281E-02

 2.67861711124688E-02
 7.59845093593537E-03

 5.71839608018997E-03
 1.72367544721218E-02

 .248298777166143
 .632711280282887

 2.50676426062644
 1.6945565513436

 1.51928754804205
 2.38326178550338

 .049057819003735
 1.73897434171407E-02

 1.68542200258231E-02
 1.78487102521972E-02

 2.52813300387346E-02
 1.47379350368142E-02

 1.26406650193673E-02
 5.55859833568426E-03

 9.36011147862674E-02
 7.71064282894918E-02

 .128513427696901
 .590690316717714

 2.40774571797472E-02
 2.65180838032644E-03

 1.35435696636078E-02
 2.94248737586222E-02

 1.07897104986742
 .94735854387162

 2.70871393272157E-03
 8.05741777099187E-03

 7.22323715392417E-03
 6.93549884085376E-03

 2.70871393272157E-03
 1.17291524514439E-03

 .874914600269066
 .419189709351602

 2.40774571797472E-03
 1.01992630012555E-04

 0
 3.05977890037666E-04

 6.01936429493681E-04
 2.70280469533272E-03

 8.51740047733559E-02
 1.85116623472788E-02

 .118882444825002
 6.03796369674327E-02

 2.40774571797472E-03
 1.17291524514439E-03

 0
 1.42789682017577E-03

 1.20387285898736E-02
 1.92766070723729E-02

 1.20387285898736E-02
 2.44782312030133E-03

 0
 9.28132933114253E-03

 0
 0

 0
 2.80479732534527E-03

 0
 2.39682680529505E-03

 0
 2.34583049028877E-03

 10.7247013642889
 14.1322008071697

 1.33419209597274
 2.10405696084401

 .379520918795766
 .659229364086151

 .940525671083877
 1.83153265345046

 .442122307463109
 .340553391611922

 6.62130072443049E-02
 5.02823665961898E-02

 .470112351434565
 .510218131637808

 5.11645965069629E-02
 8.49088644854523E-02

 1.15692181748686
 1.72989699764295

 .405705153478741
 .480436283674142

 .222716478912662
 .204750204750205

 0
 7.13438446937824E-02

 .148979266299686
 .129020676965882

 4.03297407760766E-02
 1.77977139371909E-02

 9.33001465715206E-03
 7.95542514097931E-03

 .539335040826338
 .124380012300311

 .410520644914691
 .815278088005361

 6.13975158083555E-02
 .125195953340412

 .42105453243083
 .784170335851531

 3.91258679170893E-03
 4.72225876958131E-02

 .155600567024117
 .119127391854665

 .237463921435257
 .171908577886162

 0
 4.7936536105901E-03

 .018960997529051
 3.17197079339047E-02

 3.25045671926588E-02
 .126623850160587

 8.72807822765838E-03
 .135446212656673

 .479442366091717
 .604153343879371

 3.00968214746841E-04
 4.06950593750096E-02

 0
 7.13948410087887E-04

 6.32033250968365E-03
 3.72273099545827E-03

 1.10455334812091
 .763159854068945

 .106542748020382
 .18919632867329

 4.63491050710135E-02
 3.05977890037666E-03

 0
 1.17291524514439E-03

 1.92619657437978E-02
 2.13674559876303E-02

 1.23396968046205E-02
 4.54377166705934E-02

 .387045124164437
 .329538187570566

 .201648703880383
 .19919160641452

 4.42423275677856E-02
 3.41165347391997E-02

 0
 2.80479732534527E-03

 .020766806817532
 1.59618465969649E-02

 .163726708822281
 .164004149060189

 1.02329193013926E-02
 3.49834720943065E-02

 2.70871393272157E-03
 0

 .314210816195702
 .49374632189078

 2.40774571797472E-03
 4.48767572055243E-03

 1.62522835963294E-02
 0

 1.26406650193673E-02
 3.56974205043944E-03

 0
 4.58966835056499E-04

 0
 2.03985260025111E-04

 7.52420536867101E-03
 0

 1.08348557308863E-02
 1.03012556312681E-02

 0
 1.98885628524483E-03

 0
 2.14184523026366E-03

 1.14367921603799E-02
 2.78949843084339E-02

 1.14367921603799E-02
 2.78949843084339E-02

 1.04556357803052
 1.04358859028847

 .570033798730516
 .405063730094863

 .425870023866779
 .530616657640319

 0
 6.62952095081609E-04

 3.16016625484183E-02
 .098269899017097

 0
 9.17933670112998E-04

 .887555265288433
 .16685994270054

 .208871941034307
 5.37501160166166E-02

 8.06594815521533E-02
 .07384266412909

 3.37084400516461E-02
 0

 2.79900439714562E-02
 3.36575679041432E-03

 6.53101026000644E-02
 2.26933601777936E-02

 .466199764642856
 9.07734407111742E-03

 0
 5.60959465069054E-04

 5.93599609945194
 2.01970905582363

 2.04658386027852E-02
 1.33610345316447E-02

 5.91553026084915
 2.00634802129198

 .239871667153232
 .192307103888673

 9.39020830010143E-02
 8.49598608004586E-02

 8.54749729881027E-02
 6.52242868930291E-02

 3.00968214746841E-03
 1.10662003563622E-02

 0
 5.09963150062776E-04

 0
 2.44782312030133E-03

 0
 7.64944725094165E-03

 0
 1.52988945018833E-04

 .566121211938807
 .531942561830482

 0
 3.16177153038921E-03

 .566121211938807
 .528780790300093

 6.65139754590518E-02
 .027741995363415

 1.92619657437978E-02
 1.47889313518205E-02

 2.43784253944941E-02
 1.29530640115945E-02

 2.28735843207599E-02
 0

 0
 1.73387471021344E-03

 1.5048410737342E-03
 0

 1.5048410737342E-03
 0

 0
 3.56974205043944E-04

 0
 3.56974205043944E-04

 5.35723422249376E-02
 5.81357991071565E-03

 5.35723422249376E-02
 5.81357991071565E-03

 3.72147197534468
 1.90751716280981

 3.72147197534468
 1.90751716280981

 3.72147197534468
 1.90751716280981

 2.43784253944941E-02
 6.15525522125771E-02

 9.6309828718989E-03
 4.82425139959387E-02

 2.52813300387346E-02
 3.60543947094383E-02

 .428879706014248
 .784017346906512

 0
 2.28463491228124E-02

 2.79900439714562E-02
 .153396915538883

 8.1261417981647E-03
 2.68750580083083E-02

 1.26406650193673E-02
 1.03012556312681E-02

 0
 1.01992630012555E-02

 1.49611299550654
 .160383410694743

 .182988674566079
 .324795530274982

 6.32033250968365E-03
 1.74917360471532E-02

 9.96204790812042E-02
 2.15204449326492E-02

 .898089152804572
 .102502593162618

 0
 5.09963150062776E-04

 .495092713258553
 .125144957025405

 6.32033250968365E-03
 0

 0
 1.68287839520716E-03

 3.44969767742829
 4.29807142135909

 3.44969767742829
 4.29807142135909

 2.82849928219081
 3.12235137888936

 2.4622209648439
 2.6506864613963

 6.35042933115834E-02
 8.27670192551886E-02

 .12098922232823
 .152631970813789

 2.40774571797472E-02
 .02534516855812

 .144464743078483
 .209492862045789

 .013242601448861
 0

 0
 5.09963150062776E-04

 .621198395237479
 1.17572004246973

 .25431814146108
 .234838030603909

 .295550786881397
 .940882011865822

 7.13294668950012E-02
 0

 .879429123490268
 .656118588870768

 .809303529454254
 .619452238381255

 .117678571966015
 .110611007248616

 .117678571966015
 .107908202553283

 0
 2.70280469533272E-03

 .558897974784883
 .427757090272657

 .545354405121275
 .414243066795993

 1.35435696636078E-02
 1.35140234766636E-02

 0
 2.4988194353076E-03

 0
 2.4988194353076E-03

 .11527082624804
 7.39956530741089E-02

 .11527082624804
 7.39956530741089E-02

 3.61161857696209E-03
 3.4167531054206E-03

 3.61161857696209E-03
 3.4167531054206E-03

 1.38445378783547E-02
 1.17291524514439E-03

 1.38445378783547E-02
 1.17291524514439E-03

 7.01255940360139E-02
 3.66663504895136E-02

 5.50771832986718E-02
 1.56558687069272E-02

 0
 0

 5.50771832986718E-02
 1.56558687069272E-02

 1.11358239456331E-02
 0

 1.11358239456331E-02
 0

 0
 4.89564624060265E-03

 0
 4.89564624060265E-03

 3.91258679170893E-03
 1.61148355419837E-02

 3.91258679170893E-03
 1.61148355419837E-02

 7.29306177974544
 7.56907705849175

 5.76715293097896
 5.3079004511134

 1.95749726871345
 4.086079739878

 1.94997306334478
 4.07792032947699

 .993195108664574
 1.80853331538263

 .847526492727103
 2.21925763644319

 0
 3.56974205043944E-04

 0
 0

 0
 0

 8.93875597798116E-02
 4.79875324209073E-02

 0
 0

 0
 0

 0
 0

 3.00968214746841E-03
 6.11955780075332E-04

 2.70871393272157E-03
 8.15941040100442E-03

 2.70871393272157E-03
 8.15941040100442E-03

 0
 0

 0
 0

 2.10677750322788E-03
 0

 2.10677750322788E-03
 0

 .79636189622014
 2.75380101033899E-03

 4.21355500645577E-03
 2.14184523026366E-03

 4.21355500645577E-03
 2.14184523026366E-03

 .792148341213684
 6.11955780075332E-04

 .788837690851469
 6.11955780075332E-04

 0
 0

 3.31065036221525E-03
 0

 1.05368971982869
 .894271379950085

 1.05368971982869
 .894271379950085

 7.82517358341786E-03
 0

 .827963558768558
 .841796171808625

 0
 1.22391156015066E-03

 0
 .001835867340226

 .466500732857603
 .322041729264643

 .466500732857603
 .322041729264643

 .414734199921146
 .308221727897942

 5.41742786544313E-03
 1.93785997023855E-03

 5.41742786544313E-03
 1.93785997023855E-03

 5.41742786544313E-03
 1.93785997023855E-03

 0
 0

 1.48768588549363
 8.15941040100442E-04

 1.48768588549363
 8.15941040100442E-04

 2.34755207502536E-02
 3.05977890037666E-04

 1.5048410737342E-03
 0

 0
 0

 0
 0

 0
 0

 .591703510192289
 1.372871796284

 9.02904644240522E-04
 5.60959465069054E-04

 9.02904644240522E-04
 5.60959465069054E-04

 9.02904644240522E-04
 0

 .590800605548048
 1.37174987735386

 .590800605548048
 1.37174987735386

 .377113173077791
 .5653961444746

 5.98926747346213E-02
 .729349297219783

 .10955243016785
 0

 0
 0

 0
 0

 0
 0

 0
 0

 0
 0

 0
 0

 0
 0

 0
 0

 0
 0

 0
 0

 0
 0

 .934205338574193
 .88830481109435

 .933302433929953
 .88830481109435

 .933302433929953
 .88830481109435

 .35153087482431
 .123768056520236

 .51826726579406
 .737253726045756

 3.91258679170893E-02
 2.57531390781702E-02

 2.43784253944941E-02
 1.07092261513183E-03

 9.02904644240522E-04
 0

 9.02904644240522E-04
 0

 9.02904644240522E-04
 0

 33.1335907614797
 41.7020836074385

 33.1335907614797
 41.7020836074385

 32.9322430258141
 41.6725057447349

 19.0269095680805
 27.6860014095381

 19.0269095680805
 27.6860014095381

 .610062571291846
 .399964098594235

 .457170718200451
 .273901207898717

 0
 7.30267230889896E-02

 0
 1.32590419016322E-03

 .136037633065572
 4.20719598801791E-02

 1.68542200258231E-02
 1.47889313518205E-03

 0
 5.91557254072821E-03

 6.20205200128814
 6.30095169323065

 6.19121714555726
 6.25607493602512

 1.08348557308863E-02
 4.48767572055243E-02

 1.59121895136655
 2.35246001123959

 3.61161857696209E-03
 1.63188208020088E-03

 .096309828718989
 .054362071796692

 1.18671767074679
 1.77885346004898

 .223017447127409
 .282315599874753

 2.67861711124688E-02
 .120198314469796

 1.62522835963294E-02
 2.14694486176429E-02

 0
 3.21276784539549E-03

 2.70871393272157E-02
 4.49277535205306E-02

 1.14367921603799E-02
 1.11171966713685E-02

 0
 2.46822164630384E-02

 .976039920424004
 .700791360816267

 .466199764642856
 .272677296338567

 .4953936814733
 .425819230302418

 1.44464743078483E-02
 2.29483417528249E-03

 2.98801243600663
 2.58622711922836

 2.98801243600663
 2.58622711922836

 1.45367647722724
 1.63060717232573

 .217299051047219
 .097708939552028

 .133930855562344
 .182566807722474

 1.10244657061768
 1.35033142505123

 3.64171539843677E-02
 4.99763887061521E-03

 3.64171539843677E-02
 4.99763887061521E-03

 0
 1.42789682017577E-03

 0
 1.42789682017577E-03

 2.70871393272157E-02
 9.07734407111742E-03

 2.70871393272157E-02
 0

 0
 9.07734407111742E-03

 .201347735665636
 2.88639142935531E-02

 .198338053518168
 2.76909990484088E-02

 9.02904644240522E-04
 0

 .197435148873927
 2.74360174733774E-02

 0
 2.54981575031388E-04

 3.00968214746841E-03
 1.17291524514439E-03

 2.10677750322788E-03
 1.17291524514439E-03

 9.02904644240522E-04
 0

 0
 1.52988945018833E-04

 0
 1.52988945018833E-04

 0
 1.52988945018833E-04

 0
 5.60959465069054E-04

 0
 4.07970520050221E-04

 0
 4.07970520050221E-04

 0
 1.52988945018833E-04

 0
 1.52988945018833E-04

 2.61842346829751E-02
 3.05977890037666E-04

 2.61842346829751E-02
 3.05977890037666E-04

 2.61842346829751E-02
 3.05977890037666E-04

 2.61842346829751E-02
 3.05977890037666E-04

 2.61842346829751E-02
 3.05977890037666E-04

 .546257309765516
 .387673986677723

 3.61161857696209E-03
 8.46538829104209E-03

 3.61161857696209E-03
 8.46538829104209E-03

 3.61161857696209E-03
 3.00878258537038E-03

 0
 0

 .542645691188554
 .379208598386681

 .542645691188554
 .379208598386681

 .101125320154938
 3.70233246945576E-02

 0
 5.66059096569682E-03

 0
 2.03985260025111E-04

 0
 4.64066466557127E-03

 2.70871393272157E-03
 0

 2.21091250553029
 .537603152796179

 2.21091250553029
 .537603152796179

 1.63154869214262
 .454377166705934

 1.02329193013926E-02
 9.28132933114253E-03

 .136940537709812
 9.96467995222665E-02

 .76686701117495
 .194907915953993

 1.35435696636078E-02
 0

 .442122307463109
 3.10567558388231E-02

 1.71551882405699E-02
 2.65180838032644E-03

 0
 3.72273099545827E-03

 0
 9.79129248120531E-03

 2.55822982534815E-02
 0

 .47342300179678
 4.01340999099405E-02

 3.73200586286082E-02
 1.42789682017577E-03

 .436102943168172
 3.87062030897647E-02

 .105940811590888
 1.52988945018833E-02

 .105940811590888
 1.52988945018833E-02

 .105940811590888
 1.52988945018833E-02

 0
 2.77929916784213E-02

 0
 2.77929916784213E-02

 0
 2.03985260025111E-03

 0
 2.57531390781702E-02

 2.90675101802499
 3.34474630863174

 2.22746575734137
 2.52227774021049

 1.14367921603799E-02
 3.4167531054206E-03

 1.14367921603799E-02
 3.4167531054206E-03

 9.33001465715206E-03
 3.4167531054206E-03

 2.10677750322788E-03
 0

 2.17870890655238
 2.48979308755149

 2.17870890655238
 2.48979308755149

 2.17870890655238
 2.47643205301985

 0
 1.33610345316447E-02

 3.73200586286082E-02
 2.90678995535783E-02

 0
 0

 0
 0

 3.73200586286082E-02
 2.90678995535783E-02

 3.73200586286082E-02
 2.90678995535783E-02

 0
 0

 0
 0

 0
 0

 0
 0

 0
 0

 0
 0

 0
 0

 0
 0

 0
 0

 0
 0

 0
 0

 0
 0

 0
 0

 0
 0

 0
 0

 0
 0

 0
 0

 0
 0

 .679285260683619
 .822213586846214

 .679285260683619
 .822213586846214

 .462889114280641
 .529086768190131

 .462889114280641
 .528627801355074

 0
 4.58966835056499E-04

 .151086043802914
 .184606660322725

 4.99607236479755E-02
 4.37548382753862E-02

 2.67861711124688E-02
 1.14231745614062E-02

 1.62522835963294E-02
 1.14231745614062E-02

 3.06987579041777E-02
 3.94711478148589E-02

 3.00968214746841E-03
 2.54981575031388E-03

 1.08348557308863E-02
 2.35602975329003E-02

 0
 5.7115872807031E-03

 0
 9.17933670112998E-04

 0
 3.16177153038921E-03

 1.35435696636078E-02
 4.18169783051477E-02

 0
 8.15941040100442E-04

 4.63491050710135E-02
 8.13391224350128E-02

 4.63491050710135E-02
 8.13391224350128E-02

 1.59513153815826E-02
 2.39172717379442E-02

 7.22323715392417E-03
 1.26470861215569E-02

 3.00968214746841E-03
 6.98649515586004E-03

 1.20387285898736E-03
 4.28369046052732E-03

 4.51452322120261E-03
 0

 3.00968214746841E-03
 3.26376416040177E-03

 3.00968214746841E-03
 3.26376416040177E-03

 0
 2.54981575031388E-04

 0
 2.54981575031388E-04

 0
 2.54981575031388E-04

 0
 0

 0
 0

 0
 0

 0
 0

 4.75529779300008E-02
 .058951740147257

 4.75529779300008E-02
 .058951740147257

 4.75529779300008E-02
 .058951740147257

 3.28055354074056E-02
 .051251296581309

 3.28055354074056E-02
 .051251296581309

 1.47474425225952E-02
 7.70044356594792E-03

 1.47474425225952E-02
 3.92671625548338E-03

 0
 3.77372731046455E-03

 .45536490891197
 .167114924275572

 .45536490891197
 .167114924275572

 .45536490891197
 .167114924275572

 .119484381254496
 4.13070151550849E-02

 4.00287725613298E-02
 2.69770506383209E-02

 7.94556086931659E-02
 .014329964516764

 .335880527657474
 .125807909120487

 2.97958532599372E-02
 6.11955780075332E-03

 .306084674397537
 .107347243088214

 0
 1.16271598214313E-02

 1.35826955315249
 1.09514586475981

 1.35826955315249
 1.09514586475981

 .064106229741077
 8.77136618107975E-03

 .064106229741077
 8.77136618107975E-03

 5.14655647217097E-02
 8.77136618107975E-03

 1.29416332341141
 1.08617051331871

 1.29416332341141
 1.08617051331871

 1.29416332341141
 1.08617051331871

 0
 2.03985260025111E-04

 0
 2.03985260025111E-04

 0
 2.03985260025111E-04

 0
 2.03985260025111E-04

 0
 2.03985260025111E-04

 0
 2.03985260025111E-04

 0
 1.01992630012555E-04

 0
 1.01992630012555E-04

 0
 1.01992630012555E-04

 6.01936429493681E-04
 0

 6.01936429493681E-04
 0

 6.01936429493681E-04
 0

 6.01936429493681E-04
 0

 6.01936429493681E-04
 0

 5.11645965069629E-03
 1.73387471021344E-03

 5.11645965069629E-03
 1.73387471021344E-03

 5.11645965069629E-03
 1.73387471021344E-03

 5.11645965069629E-03
 1.73387471021344E-03

 5.11645965069629E-03
 1.73387471021344E-03

 6.62130072443049E-03
 1.93785997023855E-03

 6.62130072443049E-03
 1.58088576519461E-03

 6.62130072443049E-03
 1.58088576519461E-03

 6.62130072443049E-03
 1.58088576519461E-03

 6.62130072443049E-03
 1.58088576519461E-03

 0
 3.56974205043944E-04

 0
 3.56974205043944E-04

 0
 3.56974205043944E-04

 0
 3.56974205043944E-04

 0
 5.60959465069054E-04

 0
 5.60959465069054E-04

 0
 5.60959465069054E-04

 0
 5.60959465069054E-04

 0
 5.60959465069054E-04

 1.90151718077054
 2.65180838032644E-03

 1.9006142761263
 2.65180838032644E-03

 1.9006142761263
 2.24383786027622E-03

 1.9006142761263
 2.24383786027622E-03

 1.9006142761263
 2.24383786027622E-03

 1.89248813432813
 2.24383786027622E-03

 8.1261417981647E-03
 0

 0
 4.07970520050221E-04

 0
 4.07970520050221E-04

 0
 4.07970520050221E-04

 0
 4.07970520050221E-04

 9.02904644240522E-04
 0

 9.02904644240522E-04
 0

 9.02904644240522E-04
 0

 9.02904644240522E-04
 0

 9.02904644240522E-04
 0
