## Supplementary Item 2 for "Methane emission of humans is explained by dietary habits, host genetics, local formate availability and a uniform archaeome"

Javascript must be enabled to view this page.

magnitude

HE
LE

 100
 100

 100
 100

 0
 2.90662186593497E-03

 0
 2.90662186593497E-03

 0
 2.32529749274798E-03

 100
 97.1875526825213

 0
 4.06927061230896E-03

 0
 4.06927061230896E-03

 0
 4.06927061230896E-03

 0
 1.16264874637399E-03

 0
 1.16264874637399E-03

 0
 2.90662186593497E-03

 100
 97.0285604664547

 100
 97.0285604664547

 100
 97.0285604664547

 98.1256149082702
 92.6471186657443

 7.23421494299439E-04
 0

 0
 4.34162108114707

 0
 3.48794623912197E-03

 0
 .111904941838496

 1.87438509172985
 3.98207195633091E-02

 0
 4.65059498549596E-03

 0
 1.68584068224228E-02

 0
 1.68584068224228E-02

 0
 1.68584068224228E-02

 0
 1.68584068224228E-02

 0
 2.32529749274798E-03

 0
 1.45331093296749E-02

 0
 .138064538631911

 0
 1.42424471430814E-02

 0
 1.42424471430814E-02

 0
 1.42424471430814E-02

 0
 .12382209148883

 0
 .12382209148883

 0
 .12382209148883

 0
 8.71986559780492E-03

 0
 8.71986559780492E-03

 0
 8.71986559780492E-03

 0
 2.80082083001494

 0
 2.80082083001494

 0
 2.80082083001494

 0
 2.80082083001494

 0
 2.49969480470408E-02

 0
 2.49969480470408E-02

 0
 2.7758238819679

 0
 3.54607867644067E-02
