## Supplementary Item 3 for "Methane emission of humans is explained by dietary habits, host genetics, local formate availability and a uniform archaeome"

Javascript must be enabled to view this page.

magnitude

HE
LE

 100
 100

 7.6658676273669
 7.36616001902648

 .601333974659557
 .67352950517889

 2.73731890188192E-02
 .018568827327912

 1.5106616456302E-05
 1.25042608268768E-05

 7.553308228151E-05
 1.87563912403151E-04

 9.8193006965963E-04
 6.62725823824468E-04

 1.11788961776635E-03
 8.25281214573866E-04

 1.08767638485374E-03
 9.50323822842634E-04

 1.38980871397978E-03
 1.03785364863077E-03

 2.20556600262009E-03
 1.28793886516831E-03

 3.14217622291082E-03
 3.0135268592773E-03

 .017357502308291
 1.05911089203646E-02

 3.0213232912604E-05
 0

 3.0213232912604E-05
 0

 4.13921290902675E-03
 2.53836494785598E-03

 3.0213232912604E-05
 1.25042608268768E-05

 4.5319849368906E-05
 0

 4.5319849368906E-05
 6.25213041343838E-05

 2.41705863300832E-04
 1.00034086615014E-04

 4.98518343057966E-04
 2.75093738191289E-04

 3.27813577101753E-03
 2.08821155808842E-03

 .568159844921518
 .651221903863741

 4.5319849368906E-05
 1.25042608268768E-05

 3.0213232912604E-05
 2.50085216537535E-05

 3.0213232912604E-05
 2.50085216537535E-05

 7.553308228151E-05
 1.25042608268768E-05

 1.05746315194114E-04
 0

 2.80983066087217E-03
 2.97601407679667E-03

 2.88687440479931E-02
 3.23610270199571E-02

 3.90052836901718E-02
 4.48277750643532E-02

 4.99575806209907E-02
 6.35716620438414E-02

 5.16646282805528E-02
 7.20745594061176E-02

 6.27982046088474E-02
 6.48095838657022E-02

 7.89320709841779E-02
 9.44321777645733E-02

 .253836476315242
 .276094079057439

 1.63151457728062E-03
 1.20040903938017E-03

 1.63151457728062E-03
 1.20040903938017E-03

 1.54149424981751
 1.4671499313391

 .511026621483784
 .528255002892235

 0
 2.50085216537535E-05

 3.0213232912604E-05
 2.50085216537535E-05

 2.94579020897889E-03
 4.6265765059444E-04

 2.58323141402764E-03
 7.62759910439482E-04

 1.75236750893103E-03
 1.7756050374165E-03

 3.06664314062931E-03
 8.12776953746989E-04

 1.82790059121254E-03
 2.87597999018165E-03

 8.76183754465516E-04
 3.9263378996393E-03

 2.46237848237723E-03
 3.25110781498796E-03

 3.64069456596878E-03
 4.76412337504004E-03

 5.83115395213257E-03
 4.73911485338629E-03

 6.7828707888796E-03
 1.06161174420184E-02

 .014608098113244
 1.48550618623296E-02

 1.41246863866424E-02
 1.77685546349919E-02

 2.59984869212957E-02
 2.19574820119956E-02

 2.42914392617336E-02
 2.74843652974751E-02

 2.73429757859066E-02
 2.93725086823335E-02

 4.14374489396364E-02
 5.25429039945361E-02

 4.95497019766706E-02
 4.81664127051293E-02

 .050274819566573
 4.94293430486438E-02

 6.47922779810793E-02
 6.11083226609467E-02

 7.12125899750076E-02
 7.75264171266359E-02

 .095594668935479
 9.40070328964595E-02

 2.85515051024108E-03
 6.50221562997591E-04

 4.5319849368906E-05
 2.50085216537535E-05

 7.85544055727704E-04
 2.87597999018165E-04

 2.02428660514447E-03
 3.37615042325672E-04

 .563552326902346
 .490667194846644

 7.553308228151E-05
 1.25042608268768E-05

 7.25117589902496E-04
 3.25110781498796E-04

 2.32641893427051E-03
 1.71308373328212E-03

 5.89158041795778E-03
 1.9381604281659E-03

 1.81732595969313E-02
 7.59008632191419E-03

 2.29167371642101E-02
 .016743205247188

 .021557141683143
 2.57212645208855E-02

 2.43367591111025E-02
 2.52461026094642E-02

 5.34018891730276E-02
 4.74161570555167E-02

 5.03956724982235E-02
 6.48971136914904E-02

 7.10162039610757E-02
 5.91951707544346E-02

 8.31468169754862E-02
 5.85074364089564E-02

 9.35250624809657E-02
 7.41002496600717E-02

 .116064134233768
 .107261549372949

 1.11788961776635E-02
 8.57792292723746E-03

 1.05746315194114E-04
 0

 1.10731498624694E-02
 8.57792292723746E-03

 .292947506320608
 .31328175075657

 3.0213232912604E-05
 8.75298257881373E-05

 4.38091877232758E-04
 2.25076694883782E-04

 3.7766541140755E-04
 2.75093738191289E-04

 5.89158041795778E-04
 1.75059651576275E-04

 4.68305110145362E-04
 3.37615042325672E-04

 4.98518343057966E-04
 3.62623563979426E-04

 6.7979774053359E-04
 3.37615042325672E-04

 6.64691124077288E-04
 3.50119303152549E-04

 6.94904356989892E-04
 3.75127824806303E-04

 4.07878644320154E-04
 6.37717302170715E-04

 6.49584507620986E-04
 4.87666172248194E-04

 7.70437439271402E-04
 4.37649128940687E-04

 6.94904356989892E-04
 5.25178954728824E-04

 6.49584507620986E-04
 6.00204519690084E-04

 1.02724991902854E-03
 3.25110781498796E-04

 6.64691124077288E-04
 6.87734345478222E-04

 2.00917998868817E-03
 7.75264171266359E-04

 1.05746315194114E-03
 1.81311781989713E-03

 2.82493727732847E-03
 1.88814338485839E-03

 1.66172781019322E-03
 3.95134642129306E-03

 1.06199513687803E-02
 8.36535049318055E-03

 1.24025321106239E-02
 1.12038177008816E-02

 8.52013168135433E-03
 1.47925405581952E-02

 1.45929914967877E-02
 1.53677365562315E-02

 2.70861633061495E-02
 3.23485227591302E-02

 3.58631074672609E-02
 4.17267183792877E-02

 3.90354969230844E-02
 5.66693100674055E-02

 .127968148001334
 .118152760553158

 .10302712423198
 8.45788202329944E-02

 7.553308228151E-05
 5.0017043307507E-05

 3.0213232912604E-04
 1.00034086615014E-04

 1.93364690640666E-03
 1.15039199607266E-03

 .100715811914165
 8.32783771069992E-02

 9.27546250416943E-03
 5.73945571953643E-03

 2.87025712669738E-04
 7.50255649612605E-05

 3.17238945582342E-04
 2.50085216537535E-04

 1.32938224815458E-03
 6.12708780516961E-04

 1.32938224815458E-03
 8.37785475400743E-04

 1.5408748785428E-03
 8.5028973622762E-04

 2.13003292033858E-03
 1.35046016930269E-03

 2.34152555072681E-03
 1.76310077658962E-03

 8.71651769528625E-03
 7.30248832289603E-03

 2.87025712669738E-04
 1.50051129922521E-04

 5.43838192426872E-04
 3.12606520671919E-04

 6.7979774053359E-04
 2.87597999018165E-04

 9.36610220290724E-04
 4.50153389767563E-04

 1.55598149499911E-03
 7.37751388785729E-04

 1.72215427601843E-03
 1.30044312599518E-03

 2.9911100583478E-03
 4.06388476873495E-03

 3.78118609901239E-02
 2.78845016439352E-02

 3.92772027863852E-04
 6.25213041343838E-05

 3.62558794951248E-04
 1.00034086615014E-04

 3.92772027863852E-04
 6.37717302170715E-04

 1.05746315194114E-03
 1.00034086615014E-04

 1.40491533043609E-03
 4.87666172248194E-04

 2.03939322160077E-03
 1.37546869095644E-04

 1.38980871397978E-03
 6.87734345478222E-04

 2.09981968742598E-03
 2.37580955710658E-04

 1.91854028995035E-03
 5.62691737209454E-04

 2.47748509883353E-03
 2.12572434056905E-04

 1.11788961776635E-03
 1.36296443012957E-03

 2.09981968742598E-03
 2.21325416635719E-03

 3.32345562038644E-03
 4.02637198625432E-03

 6.40520537747205E-03
 6.01454945772772E-03

 1.13299623422265E-02
 1.10412623101322E-02

 1.10278300131005E-03
 2.12572434056905E-04

 1.10278300131005E-03
 2.12572434056905E-04

 1.14243786950784
 1.10626445961461

 4.13770224738112E-02
 3.87882170849717E-02

 1.5106616456302E-05
 1.25042608268768E-05

 3.0213232912604E-05
 1.25042608268768E-05

 1.35959548106718E-04
 3.00102259845042E-04

 6.64691124077288E-04
 3.25110781498796E-04

 6.94904356989892E-04
 5.0017043307507E-04

 6.94904356989892E-04
 6.25213041343838E-04

 6.64691124077288E-04
 8.37785475400743E-04

 1.29916901524197E-03
 1.67557095080149E-03

 3.95793351155112E-03
 1.65056242914773E-03

 3.32194495874081E-02
 3.28486931922052E-02

 .106637605565036
 8.62168784013153E-02

 3.0213232912604E-05
 0

 1.05746315194114E-04
 1.25042608268768E-05

 9.0639698737812E-05
 3.75127824806303E-05

 1.20852931650416E-04
 1.12538347441891E-04

 2.11492630388228E-04
 1.75059651576275E-04

 8.61077138009214E-04
 3.37615042325672E-04

 4.38091877232758E-03
 4.66408928842503E-03

 1.65266384031944E-02
 4.57655946263689E-03

 .036165239796387
 3.53245368359268E-02

 4.81447866462345E-02
 4.09764627296751E-02

 7.34785824434529E-02
 4.60782011470409E-02

 1.5106616456302E-05
 1.25042608268768E-05

 1.5106616456302E-05
 6.25213041343838E-05

 2.11492630388228E-04
 1.25042608268768E-05

 7.32368765801521E-02
 4.59906713212527E-02

 .293385598197841
 .291611866743593

 3.0213232912604E-05
 3.75127824806303E-05

 5.78583410276366E-03
 8.90303370873625E-03

 8.20289273577198E-03
 9.54075101090697E-03

 1.45174584145062E-02
 2.06070218426929E-02

 1.78711272678053E-02
 .02242013966259

 2.75242551833822E-02
 2.62214349539606E-02

 4.10899967611414E-02
 3.56621518782525E-02

 4.80541469474966E-02
 4.66033801017697E-02

 6.10005172505475E-02
 5.76571466727287E-02

 6.93091563015136E-02
 6.39592941294746E-02

 .107000164359987
 .102709998431966

 1.5106616456302E-04
 1.25042608268768E-05

 2.41705863300832E-04
 5.0017043307507E-05

 1.81279397475624E-04
 2.50085216537535E-04

 3.32345562038644E-04
 2.37580955710658E-04

 6.64691124077288E-04
 4.75161911421317E-04

 8.68630446237365E-03
 8.32783771069992E-03

 1.78258074184364E-02
 2.05820133210391E-02

 3.55458685216786E-02
 3.13356776321532E-02

 .043371095846043
 4.14391203802696E-02

 1.35657415777592E-02
 1.04785705729227E-02

 3.47452178494946E-04
 2.25076694883782E-04

 1.32182893992642E-02
 1.02534938780389E-02

 .153286837182096
 .182062037639326

 8.45970521552912E-04
 2.50085216537535E-05

 1.95630683109111E-02
 2.62589477364412E-02

 2.85515051024108E-02
 3.43617087522573E-02

 .104326293247222
 .121416372628973

 .284155455543041
 .277719632964933

 1.17831608359156E-03
 0

 1.93364690640666E-03
 2.03819451478091E-03

 4.83411726601664E-03
 7.37751388785729E-04

 6.76776417242329E-03
 7.62759910439482E-04

 3.70112103179399E-03
 4.72661059255941E-03

 7.40224206358798E-03
 5.38933641638388E-03

 .018656671323533
 .012479252305223

 4.63319926714782E-02
 5.02921370456983E-02

 5.42931795439494E-02
 5.36807917297819E-02

 6.59857006811271E-02
 7.25372170567121E-02

 7.30707037991328E-02
 7.50755820045681E-02

 6.95508621648144E-02
 7.05990566285462E-02

 6.95508621648144E-02
 7.05990566285462E-02

 2.28916601808927
 2.13466488706027

 2.61344464694025E-03
 1.11287921359203E-03

 3.0213232912604E-05
 0

 3.0213232912604E-05
 0

 1.5106616456302E-05
 1.25042608268768E-05

 4.5319849368906E-05
 0

 3.0213232912604E-05
 1.25042608268768E-05

 4.5319849368906E-05
 5.0017043307507E-05

 1.5106616456302E-04
 1.62555390749398E-04

 2.11492630388228E-04
 1.37546869095644E-04

 2.05449983805707E-03
 7.37751388785729E-04

 6.91580901369505E-02
 8.49039310144932E-02

 0
 2.50085216537535E-05

 3.0213232912604E-05
 1.25042608268768E-05

 6.0426465825208E-05
 1.25042608268768E-05

 1.05746315194114E-04
 7.50255649612605E-05

 6.0426465825208E-05
 1.37546869095644E-04

 0
 6.62725823824468E-04

 9.51716836747026E-04
 1.25042608268768E-05

 1.05746315194114E-03
 8.00272692920113E-04

 8.3086390509661E-04
 1.70057947245524E-03

 6.7979774053359E-04
 2.3132882529722E-03

 2.97600344189149E-03
 1.20040903938017E-03

 1.37470209752348E-03
 3.02603112010418E-03

 6.72244432305439E-03
 4.0888932903887E-03

 7.08500311800564E-03
 6.48971136914904E-03

 .0472232830424
 6.43469262151078E-02

 3.23885856823115E-02
 2.10446709716336E-02

 1.5106616456302E-05
 1.25042608268768E-05

 8.76183754465516E-04
 3.37615042325672E-04

 9.92504701179041E-03
 4.85165320082818E-03

 2.15722482995993E-02
 1.58428984676529E-02

 .762672638412863
 .704002388813988

 9.0639698737812E-05
 0

 4.68305110145362E-04
 4.37649128940687E-04

 4.68305110145362E-04
 4.75161911421317E-04

 1.34448886461088E-03
 5.75195998036331E-04

 7.85544055727704E-04
 1.0753664311114E-03

 2.05449983805707E-03
 7.25247127958852E-04

 6.57137815849137E-03
 4.66408928842503E-03

 5.57434147237544E-03
 9.54075101090697E-03

 .018158152980475
 1.07286557894603E-02

 2.17384210806186E-02
 1.51551641221746E-02

 4.51838898207993E-02
 5.16551014758279E-02

 .054776591270551
 5.23178272996524E-02

 6.59101675988456E-02
 5.84074023223413E-02

 6.55929286532633E-02
 6.64476420340231E-02

 7.81163136955376E-02
 6.46470284749528E-02

 .106395899701735
 .101046931741991

 .135400603297835
 .10613616589853

 .154042168004911
 .159967008758234

 .381638451535557
 .409364490950291

 1.81279397475624E-04
 1.25042608268768E-05

 1.55598149499911E-03
 9.00306779535127E-04

 2.85515051024108E-03
 2.02569025395404E-03

 3.14217622291082E-03
 2.93850129431604E-03

 7.35692221421907E-03
 6.52722415162967E-03

 3.57422545356105E-02
 3.90508065623361E-02

 7.27987847029193E-02
 6.77981022033258E-02

 7.59107476929175E-02
 7.02739458470474E-02

 7.94759091766048E-02
 9.84585497508276E-02

 .102619245587659
 .121378859846493

 7.23909060585992E-02
 .04900419818053

 3.0213232912604E-04
 0

 1.5106616456302E-03
 1.25042608268768E-03

 6.99436341926782E-03
 4.87666172248194E-03

 1.61791862246994E-02
 8.74047831798685E-03

 2.19348070945505E-02
 1.74059310710124E-02

 2.54697553453252E-02
 1.67307009863611E-02

 4.81901064956034E-03
 4.17642311617684E-03

 2.56812479757134E-04
 2.75093738191289E-04

 8.15757288640308E-04
 6.12708780516961E-04

 7.25117589902496E-04
 9.37819562015757E-04

 3.0213232912604E-03
 2.35080103545283E-03

 .133618022555991
 .132770241459777

 6.49584507620986E-04
 3.25110781498796E-04

 1.07256976839744E-03
 2.00068173230028E-04

 1.28406239878567E-03
 3.37615042325672E-04

 2.96089682543519E-03
 2.56337346950974E-03

 3.31288098886703E-02
 3.12106350238844E-02

 4.58787941777892E-02
 4.31772126352054E-02

 4.86433049892924E-02
 5.49562263341234E-02

 .446324983201443
 .382130210869354

 1.01214330257223E-03
 4.12640607286933E-04

 2.58323141402764E-03
 1.18790477855329E-03

 5.18156944451159E-03
 3.45117598821799E-03

 .010106326409266
 5.52688328547953E-03

 8.97333017504339E-03
 7.71512893018296E-03

 1.09220836979063E-02
 6.51471989080279E-03

 3.38690340950291E-02
 3.60998010071932E-02

 5.33112494742897E-02
 4.82414382700905E-02

 .070729178248406
 6.18585783105593E-02

 8.55185557591256E-02
 6.05831437062179E-02

 7.71645968587906E-02
 7.47504712230693E-02

 8.69536843224743E-02
 .0757883248717

 .159208630832967
 .131832421897762

 2.52280494820243E-03
 1.87563912403151E-03

 5.07582312931747E-03
 3.93884216046618E-03

 6.90372372053001E-03
 5.03921711323133E-03

 7.5835214610636E-03
 5.30180659059575E-03

 .012553598275187
 7.61509484356795E-03

 .124569159298666
 .108061822065869

 .224333254376085
 .214323030572668

 4.15885151041994E-02
 4.17767354225952E-02

 .075382016116947
 7.17244401029651E-02

 .107362723154938
 .100821855047107

 2.56510347428008E-02
 2.96976194638323E-02

 4.5319849368906E-05
 1.25042608268768E-05

 3.0213232912604E-05
 0

 1.5106616456302E-05
 1.25042608268768E-05

 2.32490827262488E-02
 2.83596635553565E-02

 1.66172781019322E-04
 0

 1.45023517980499E-03
 1.4004772126102E-03

 1.91854028995035E-03
 1.51301556005209E-03

 3.59537471659988E-03
 2.90098851183541E-03

 1.61187597588742E-02
 2.25451822708588E-02

 2.35663216718311E-03
 1.32545164764894E-03

 2.35663216718311E-03
 1.32545164764894E-03

 .077632901968936
 7.41502667033792E-02

 3.63465191938626E-02
 2.39956765267765E-02

 3.0213232912604E-05
 0

 6.0426465825208E-05
 0

 1.5106616456302E-05
 3.75127824806303E-05

 6.0426465825208E-05
 1.25042608268768E-05

 4.5319849368906E-05
 6.25213041343838E-05

 4.5319849368906E-05
 8.75298257881373E-05

 1.5106616456302E-04
 2.50085216537535E-05

 1.5106616456302E-04
 8.75298257881373E-05

 2.71919096213436E-04
 2.75093738191289E-04

 5.43838192426872E-04
 1.62555390749398E-04

 4.22985260776456E-04
 4.12640607286933E-04

 1.01214330257223E-03
 5.0017043307507E-05

 7.10010973446194E-04
 4.6265765059444E-04

 5.74051425339476E-04
 1.60054538584023E-03

 1.31427563169827E-03
 1.41298147343707E-03

 1.60130134436801E-03
 1.21291330020705E-03

 1.5408748785428E-03
 1.28793886516831E-03

 5.74051425339476E-03
 4.00136346460056E-03

 .00980419408014
 5.35182363390325E-03

 1.22514659460609E-02
 7.45253945281855E-03

 4.12863827750734E-02
 5.01545901766027E-02

 4.12863827750734E-02
 5.01545901766027E-02

 .901970748756423
 .881525379773158

 7.56992550625293E-02
 7.60008973057569E-02

 1.5106616456302E-05
 2.50085216537535E-05

 .075684148446073
 7.59758887841032E-02

 .345911303616403
 .284259361377389

 9.0639698737812E-05
 3.75127824806303E-05

 2.16024615325119E-03
 6.75230084651345E-04

 1.73726089247473E-03
 1.08787069193828E-03

 2.91557697606629E-03
 8.8780251870825E-04

 3.41409531912425E-03
 2.16323712304968E-03

 1.20248666992164E-02
 7.76514597349047E-03

 6.74208292444758E-02
 6.71478806403282E-02

 .076983317461315
 6.53347628204311E-02

 .179164471171742
 .139159918742311

 .187593963154358
 .1952165200292

 4.38091877232758E-04
 1.00034086615014E-04

 7.70437439271402E-04
 1.87563912403151E-04

 1.04235653548484E-03
 8.12776953746989E-04

 8.88269047630557E-03
 6.73979658568657E-03

 3.68903573862895E-02
 3.99386090810444E-02

 .044020680353664
 4.57780988871958E-02

 4.48364376423043E-02
 5.00295475683339E-02

 5.07129114438058E-02
 5.16300929541741E-02

 .208561946795705
 .230428518517685

 9.23014265480052E-03
 5.77696850201706E-03

 1.48346973600886E-02
 9.72831492331012E-03

 2.14665019844051E-02
 2.35205146153552E-02

 2.73882956352755E-02
 2.20575160986106E-02

 .058326646137782
 6.67102315113875E-02

 7.73156630233536E-02
 .102634972867004

 8.42042801274273E-02
 9.56200825431266E-02

 3.16936813253216E-02
 3.43867172739111E-02

 5.25105988021057E-02
 6.12333652692155E-02

 .484499402986518
 .487466104074964

 4.50781435056052E-02
 6.08207246619285E-02

 1.5106616456302E-05
 2.50085216537535E-05

 7.553308228151E-05
 0

 3.17238945582342E-04
 1.25042608268768E-05

 4.22985260776456E-04
 1.50051129922521E-04

 6.94904356989892E-04
 5.25178954728824E-04

 4.35523752435187E-02
 6.01079817947966E-02

 .439421259480912
 .426645379413035

 1.99407337223186E-03
 1.33795590847581E-03

 1.28859438372256E-02
 1.16914838731298E-02

 1.50159767575642E-02
 1.53677365562315E-02

 1.97292410919304E-02
 .014504942559177

 2.34001488908118E-02
 .01805615263401

 3.84010190319197E-02
 3.97135323861606E-02

 4.97460879906025E-02
 4.36648788074536E-02

 5.63476793820065E-02
 4.53904668015626E-02

 5.62117198338997E-02
 5.74695827603256E-02

 8.04427326298081E-02
 7.61884612181601E-02

 8.52466366629122E-02
 .103260185908348

 .601681426838052
 .511711865818278

 5.41270067629301E-02
 5.34056979915906E-02

 7.553308228151E-05
 0

 1.05746315194114E-04
 7.50255649612605E-05

 5.39457273654544E-02
 5.33306724266294E-02

 8.76183754465516E-03
 5.15175546067322E-03

 7.553308228151E-05
 2.50085216537535E-05

 3.62558794951248E-04
 1.25042608268768E-05

 4.09389305965784E-03
 2.06320303643467E-03

 4.22985260776456E-03
 3.05103964175793E-03

 4.35825884764313E-02
 1.79436142865681E-02

 1.20852931650416E-04
 0

 4.38091877232758E-04
 0

 5.74051425339476E-04
 2.50085216537535E-05

 2.11492630388228E-04
 4.50153389767563E-04

 4.04857321028894E-03
 0

 6.60159139140397E-03
 1.20040903938017E-03

 7.05478988509303E-03
 4.00136346460056E-03

 1.23723188777113E-02
 3.25110781498796E-04

 1.21608262473231E-02
 1.19415690896673E-02

 1.29161570701382E-02
 7.84017153845173E-03

 1.20852931650416E-04
 5.0017043307507E-05

 4.07878644320154E-04
 1.12538347441891E-04

 8.45970521552912E-04
 5.37683215555701E-04

 8.91290370921818E-04
 5.0017043307507E-04

 1.10278300131005E-03
 4.2514486811381E-04

 9.54738160038286E-03
 6.21461763095775E-03

 1.69194104310582E-03
 1.10037495276515E-03

 2.2659924684453E-04
 6.25213041343838E-05

 1.35959548106718E-04
 3.25110781498796E-04

 5.74051425339476E-04
 2.00068173230028E-04

 7.553308228151E-04
 5.12674693901947E-04

 5.43838192426872E-03
 1.05035790945765E-03

 2.56812479757134E-04
 8.75298257881373E-05

 1.32938224815458E-03
 7.62759910439482E-04

 3.85218719635701E-03
 2.00068173230028E-04

 .12675961868483
 .104185501209537

 7.70437439271402E-04
 7.62759910439482E-04

 .125989181245559
 .103422741299098

 .348403895331693
 .321034392469234

 1.07256976839744E-03
 5.62691737209454E-04

 9.98547347761562E-03
 2.40081807876034E-03

 4.14374489396364E-02
 3.60998010071932E-02

 6.20428737860323E-02
 6.88859728952641E-02

 .104703958658629
 .096995551234083

 .129161570701382
 .116089557516724

 16.5134050071787
 18.1523479125512

 3.15087763398965
 2.92728497235433

 4.78879741664773E-03
 6.70228380320594E-03

 1.5106616456302E-05
 1.25042608268768E-05

 1.20852931650416E-04
 1.25042608268768E-04

 6.0426465825208E-05
 1.87563912403151E-04

 9.0639698737812E-05
 5.12674693901947E-04

 1.35959548106718E-04
 6.25213041343838E-04

 2.87025712669738E-04
 6.12708780516961E-04

 4.98518343057966E-04
 1.0753664311114E-03

 9.8193006965963E-04
 7.00238606305098E-04

 2.59833803048394E-03
 2.8509714685279E-03

 5.19063341438537E-02
 5.95828028400677E-02

 3.0213232912604E-05
 0

 1.20852931650416E-04
 1.25042608268768E-05

 1.35959548106718E-04
 0

 4.22985260776456E-04
 2.12572434056905E-04

 5.11963231704075E-02
 .059357726145184

 .107664855484064
 8.55916653599714E-02

 3.0213232912604E-05
 0

 4.5319849368906E-05
 1.25042608268768E-05

 6.0426465825208E-05
 1.25042608268768E-05

 3.0213232912604E-05
 6.25213041343838E-05

 7.553308228151E-05
 2.50085216537535E-05

 7.553308228151E-05
 3.75127824806303E-05

 1.20852931650416E-04
 1.25042608268768E-05

 4.38091877232758E-04
 2.50085216537535E-05

 5.2873157597057E-04
 7.50255649612605E-05

 6.0426465825208E-04
 6.00204519690084E-04

 1.29916901524197E-03
 1.13788773524579E-03

 1.99407337223186E-03
 1.16289625689954E-03

 2.46237848237723E-03
 1.6130496466671E-03

 9.72866099785849E-03
 1.07286557894603E-02

 1.36865945094096E-02
 8.77799110046748E-03

 1.25838115080996E-02
 1.12413304833622E-02

 6.39009876101574E-02
 5.00670603508145E-02

 .808536325974196
 .783054325761503

 1.5106616456302E-05
 2.50085216537535E-05

 1.5106616456302E-04
 7.50255649612605E-05

 8.61077138009214E-04
 7.00238606305098E-04

 1.11788961776635E-03
 5.37683215555701E-04

 4.21474599130826E-03
 2.90098851183541E-03

 6.34477891164684E-03
 1.42923701251201E-02

 5.30544369945326E-02
 6.74354786393464E-02

 .068281906382485
 6.15959888331949E-02

 9.76189555406235E-02
 .108111839109176

 .116683505508477
 .11050015292711

 .125596409217695
 .109199709801115

 .141367716798074
 .109662367451709

 .193228731092559
 .19801747445442

 5.51391500655023E-02
 .059557794318414

 4.5319849368906E-05
 2.50085216537535E-05

 2.11492630388228E-04
 2.75093738191289E-04

 1.26895578232937E-03
 9.6282808366951E-04

 2.41705863300832E-03
 1.38797295178332E-03

 6.0426465825208E-03
 3.1760822500267E-03

 6.23903259645273E-03
 5.08923415653884E-03

 1.81128331311061E-02
 .022520173749205

 2.08018108603279E-02
 2.61214008673455E-02

 6.16349951417121E-02
 6.86358876787265E-02

 7.553308228151E-05
 0

 6.15594620594306E-02
 6.86358876787265E-02

 .453878291429593
 .428883642101046

 4.5319849368906E-05
 5.0017043307507E-05

 1.35959548106718E-04
 3.75127824806303E-05

 2.87025712669738E-04
 3.12606520671919E-04

 4.07878644320154E-04
 3.62623563979426E-04

 4.5319849368906E-04
 6.62725823824468E-04

 1.67683442664952E-03
 1.03785364863077E-03

 2.94579020897889E-03
 3.11356094589231E-03

 8.76183754465516E-03
 9.00306779535127E-04

 6.29945906227793E-03
 4.11390181204245E-03

 1.75538883222229E-02
 5.4893705029989E-03

 1.92307227488724E-02
 1.17540051772642E-02

 2.76148948821201E-02
 9.47822970677258E-03

 .026663178045373
 1.91190148042946E-02

 2.28412040819286E-02
 2.35830359194896E-02

 7.09859907281631E-02
 7.66761273904083E-02

 8.52919565122811E-02
 9.22564363806967E-02

 .162683152617916
 .179936313298757

 4.60600735752648E-02
 3.05229006784062E-02

 1.05746315194114E-04
 8.75298257881373E-05

 2.41705863300832E-04
 1.25042608268768E-05

 2.41705863300832E-04
 1.12538347441891E-04

 3.0213232912604E-04
 2.37580955710658E-04

 2.56812479757134E-04
 3.00102259845042E-04

 6.0426465825208E-04
 2.75093738191289E-04

 1.04235653548484E-03
 6.50221562997591E-04

 1.28406239878567E-03
 9.75332344496387E-04

 1.55598149499911E-03
 1.28793886516831E-03

 2.22067261907639E-03
 1.57553686418647E-03

 3.09685637354191E-03
 1.56303260335959E-03

 4.10899967611414E-03
 1.53802408170584E-03

 7.14542958383084E-03
 4.10139755121558E-03

 7.41734868004428E-03
 1.10412623101322E-02

 1.64359987044566E-02
 6.76480510734033E-03

 1.86415647070767E-02
 1.74309395926662E-02

 1.5106616456302E-04
 1.50051129922521E-04

 1.96386013931926E-04
 2.25076694883782E-04

 5.2873157597057E-04
 1.75059651576275E-04

 9.97036686115932E-04
 9.00306779535127E-04

 1.28406239878567E-03
 9.37819562015757E-04

 1.5106616456302E-03
 1.32545164764894E-03

 3.08174975708561E-03
 1.52551982087896E-03

 2.85515051024108E-03
 2.92599703348916E-03

 2.97600344189149E-03
 2.96350981596979E-03

 5.06071651286117E-03
 6.30214745674589E-03

 .320909853381223
 .257625285816142

 1.5106616456302E-04
 2.87597999018165E-04

 3.7766541140755E-04
 3.25110781498796E-04

 1.64662119373692E-03
 1.22541756103392E-03

 4.5621981698032E-03
 0

 5.68008778756955E-03
 0

 4.41113200524018E-03
 4.21393589865747E-03

 .126140247410122
 9.99965738325334E-02

 .177940835238781
 .1515766497434

 9.46731653316446E-02
 .114689080304114

 1.96386013931926E-04
 2.87597999018165E-04

 7.553308228151E-04
 8.00272692920113E-04

 9.0639698737812E-04
 7.75264171266359E-04

 1.5257682620865E-03
 1.08787069193828E-03

 6.0728598154334E-03
 2.1507328622228E-03

 8.52164234299996E-02
 .109587341886748

 6.71640167647187E-02
 6.68727869021369E-02

 4.83411726601664E-04
 3.12606520671919E-04

 7.70437439271402E-04
 6.25213041343838E-04

 1.66172781019322E-03
 1.16289625689954E-03

 3.09685637354191E-03
 1.11287921359203E-03

 4.42623862169648E-03
 7.37751388785729E-04

 2.96089682543519E-03
 2.42582660041409E-03

 4.80390403310403E-03
 3.03853538093105E-03

 4.81901064956034E-03
 4.52654241932939E-03

 4.41415332853144E-02
 5.29305360801693E-02

 .267054765714507
 .26800382230245

 6.94904356989892E-04
 7.00238606305098E-04

 9.0639698737812E-04
 7.37751388785729E-04

 9.97036686115932E-04
 6.75230084651345E-04

 9.66823453203328E-04
 7.50255649612605E-04

 2.50769833174613E-03
 1.06286217028452E-03

 3.8068673469881E-03
 6.50221562997591E-04

 3.55005486723097E-03
 2.45083512206784E-03

 4.5319849368906E-03
 2.92599703348916E-03

 5.81604733567627E-03
 5.50187476382577E-03

 5.39306207489981E-03
 7.79015449514422E-03

 2.41252664807143E-02
 2.68591522561313E-02

 6.60008072975834E-02
 8.41036583215731E-02

 .14775781555909
 .133795590847581

 3.17238945582342E-04
 1.37546869095644E-03

 3.17238945582342E-04
 1.37546869095644E-03

 .414223423231801
 .321422024554867

 2.13003292033858E-03
 1.25042608268768E-05

 2.20556600262009E-03
 8.12776953746989E-04

 2.37173878363941E-02
 9.61577657586823E-03

 3.78722874559491E-02
 1.03410237038271E-02

 .348298149016499
 .300639943060598

 7.29649574839386E-03
 6.16460058765024E-03

 7.29649574839386E-03
 6.16460058765024E-03

 .370988286933864
 .351169661062007

 7.26628251548126E-03
 9.39069988098445E-03

 1.16925211371777E-02
 9.85335753157889E-03

 2.78868139783335E-02
 3.95259684737574E-02

 3.02585527619729E-02
 3.85381318684342E-02

 .055426175778172
 4.80413700968605E-02

 5.69368374238022E-02
 5.00795646116414E-02

 8.55185557591256E-02
 7.11867568874094E-02

 9.60025475797992E-02
 8.45538117113406E-02

 .205706796285464
 .176522650093019

 .188243547661979
 .160617230321232

 3.0213232912604E-05
 0

 4.5319849368906E-05
 0

 2.56812479757134E-04
 6.87734345478222E-04

 5.58944808883174E-04
 6.50221562997591E-04

 6.0577531989771E-03
 0

 1.59979068272238E-02
 1.34420803888925E-02

 .062027767169576
 6.02455286638922E-02

 .10326883009528
 8.55916653599714E-02

 6.19371274708382E-04
 9.87836605323264E-04

 3.0213232912604E-05
 0

 0
 2.50085216537535E-05

 3.0213232912604E-05
 0

 0
 3.75127824806303E-05

 1.5106616456302E-05
 3.75127824806303E-05

 3.0213232912604E-05
 2.50085216537535E-05

 1.5106616456302E-04
 3.75127824806303E-05

 3.62558794951248E-04
 8.25281214573866E-04

 3.0213232912604E-05
 1.25042608268768E-05

 3.0213232912604E-05
 1.25042608268768E-05

 1.68136641158641E-02
 1.49050789056371E-02

 4.5319849368906E-05
 0

 1.46534179626129E-03
 9.12811040362003E-04

 1.22363593296046E-03
 1.18790477855329E-03

 2.84004389378478E-03
 2.80095442522039E-03

 1.12393226434887E-02
 1.00034086615014E-02

 3.55600687411475
 4.0232459210476

 .260120828761064
 .27953275078483

 0
 2.50085216537535E-05

 2.85515051024108E-03
 3.00102259845042E-03

 1.09371903143626E-02
 7.54006927860669E-03

 1.00761131763534E-02
 1.07036472678065E-02

 1.44117120993121E-02
 1.57928814243453E-02

 1.58468406626608E-02
 1.73934268101856E-02

 2.51978362491117E-02
 2.77094419923589E-02

 .029865780734109
 3.08605157207318E-02

 6.78438145052523E-02
 7.24246787092702E-02

 .083086390509661
 9.40820584614207E-02

 1.40721153613744
 1.76401358762998

 1.5106616456302E-05
 1.25042608268768E-05

 4.5319849368906E-05
 2.50085216537535E-05

 7.553308228151E-05
 1.25042608268768E-05

 1.20852931650416E-04
 2.50085216537535E-05

 2.2659924684453E-04
 1.00034086615014E-04

 2.56812479757134E-04
 1.12538347441891E-04

 4.38091877232758E-04
 2.50085216537535E-04

 3.17238945582342E-04
 3.75127824806303E-04

 6.0426465825208E-04
 5.0017043307507E-04

 6.64691124077288E-04
 6.75230084651345E-04

 1.34448886461088E-03
 2.62589477364412E-04

 1.32938224815458E-03
 1.2379218218608E-03

 1.32938224815458E-03
 1.4004772126102E-03

 2.2810990849016E-03
 2.57587773033661E-03

 3.0364299077167E-03
 2.13822860139593E-03

 2.94579020897889E-03
 2.56337346950974E-03

 2.00464800375127E-02
 2.01193556704447E-02

 2.86421448011486E-02
 2.10196624499798E-02

 3.18447474898846E-02
 2.10196624499798E-02

 .037268022797697
 2.44583341773709E-02

 1.27437905763718
 1.66512989301104

 .889326510782499
 .920963818421127

 3.0213232912604E-05
 0

 3.0213232912604E-05
 1.25042608268768E-05

 1.5106616456302E-05
 6.25213041343838E-05

 2.41705863300832E-04
 2.50085216537535E-04

 3.7766541140755E-04
 2.62589477364412E-04

 3.92772027863852E-04
 2.87597999018165E-04

 5.2873157597057E-04
 2.75093738191289E-04

 4.98518343057966E-04
 3.00102259845042E-04

 5.89158041795778E-04
 2.87597999018165E-04

 6.19371274708382E-04
 2.75093738191289E-04

 3.62558794951248E-04
 5.12674693901947E-04

 7.40224206358798E-04
 3.25110781498796E-04

 5.13624959514268E-04
 5.62691737209454E-04

 6.19371274708382E-04
 5.87700258863208E-04

 1.4804484127176E-03
 8.8780251870825E-04

 1.75236750893103E-03
 2.1507328622228E-03

 2.97600344189149E-03
 2.95100555514291E-03

 4.75858418373513E-03
 2.96350981596979E-03

 5.42327530781242E-03
 2.52586068702911E-03

 9.0941831066938E-03
 4.66408928842503E-03

 8.62587799654844E-03
 6.26463467426526E-03

 1.08918704649937E-02
 7.81516301679797E-03

 1.22212527131483E-02
 7.82766727762485E-03

 1.24327453435365E-02
 1.33045335197969E-02

 1.34448886461088E-02
 1.51551641221746E-02

 1.76294214045044E-02
 1.31544823898744E-02

 1.99407337223186E-02
 2.21825587068794E-02

 2.85817183353234E-02
 1.52301896871359E-02

 2.92161962264881E-02
 2.08446027984036E-02

 2.90198102125561E-02
 2.74093397325139E-02

 3.58480008508046E-02
 2.63964946055368E-02

 3.84312322648323E-02
 2.56587432167511E-02

 5.16797348970091E-02
 3.68875694392864E-02

 4.58939007942455E-02
 5.71819847613074E-02

 7.64545858853444E-02
 8.04649184209519E-02

 .103072444081349
 .125005095486287

 .12385914832522
 .159079206239526

 .201038851800467
 .240957106133915

 6.92336232192321E-02
 .07995224372705

 0
 2.50085216537535E-05

 6.92336232192321E-02
 7.99272352053962E-02

 .518534609862566
 .511799395644066

 3.0213232912604E-05
 1.25042608268768E-05

 3.0213232912604E-05
 1.25042608268768E-05

 4.5319849368906E-05
 1.25042608268768E-05

 3.0213232912604E-05
 8.75298257881373E-05

 1.81279397475624E-04
 3.75127824806303E-05

 1.05746315194114E-04
 1.12538347441891E-04

 2.41705863300832E-04
 1.25042608268768E-05

 2.71919096213436E-04
 1.25042608268768E-04

 4.83411726601664E-04
 4.00136346460056E-04

 7.85544055727704E-04
 1.75059651576275E-04

 6.34477891164684E-04
 3.37615042325672E-04

 9.66823453203328E-04
 5.0017043307507E-04

 8.45970521552912E-04
 1.18790477855329E-03

 1.26895578232937E-03
 1.02534938780389E-03

 1.31427563169827E-03
 1.22541756103392E-03

 1.58619472791171E-03
 1.22541756103392E-03

 1.40491533043609E-03
 1.55052834253272E-03

 2.2810990849016E-03
 1.97567321064653E-03

 5.19667606096789E-03
 5.07672989571196E-03

 5.37795545844351E-03
 5.38933641638388E-03

 6.72244432305439E-03
 5.93952389276646E-03

 7.46266852941319E-03
 7.50255649612605E-03

 7.05478988509303E-03
 8.14027379829677E-03

 6.8432972547048E-03
 8.3903590148343E-03

 1.26291313574685E-02
 .009740819184137

 4.30840701333733E-02
 2.79845357305502E-02

 7.63790528030629E-02
 5.71444719788268E-02

 .091591415574559
 .130081825381999

 .243684830056608
 .236393050932105

 .232294441248556
 .279520246524003

 4.5319849368906E-05
 0

 1.96386013931926E-04
 2.50085216537535E-04

 1.19342270004786E-03
 2.75093738191289E-04

 1.28406239878567E-03
 5.12674693901947E-04

 2.32339761097925E-02
 2.02318940178866E-02

 .033914353944398
 3.05979262433674E-02

 3.45790450684753E-02
 .048091387140168

 4.01080666914818E-02
 6.18585783105593E-02

 9.77398084722739E-02
 .117702607163391

 5.65591720123947E-02
 .056106618330196

 6.0426465825208E-05
 0

 3.0213232912604E-05
 1.37546869095644E-04

 4.5319849368906E-04
 8.75298257881373E-05

 1.02724991902854E-03
 1.00034086615014E-04

 9.97036686115932E-04
 7.75264171266359E-04

 9.97036686115932E-04
 1.16289625689954E-03

 2.09981968742598E-03
 1.43798999509083E-03

 3.08174975708561E-03
 2.82596294687415E-03

 3.73133426470659E-03
 3.15107372837294E-03

 3.67090779888139E-03
 3.88882511715867E-03

 6.49584507620986E-03
 6.38967728253402E-03

 8.44459859907282E-03
 7.74013745183671E-03

 2.54697553453252E-02
 .028409680598664

 1.75387817057666E-02
 1.83437506330282E-02

 7.553308228151E-05
 6.25213041343838E-05

 1.35959548106718E-04
 2.12572434056905E-04

 1.73272890753784E-02
 1.80686568948369E-02

 5.92330431251601E-02
 6.82232470714396E-02

 2.2659924684453E-04
 3.25110781498796E-04

 5.58944808883174E-04
 5.37683215555701E-04

 1.31427563169827E-03
 4.12640607286933E-04

 1.05746315194114E-03
 6.50221562997591E-04

 6.43541861038465E-03
 6.70228380320594E-03

 4.96403416754084E-02
 5.95953071008946E-02

 3.03038726113418E-02
 2.99727132020236E-02

 3.32345562038644E-04
 2.50085216537535E-04

 5.13624959514268E-04
 8.25281214573866E-04

 8.3086390509661E-04
 6.50221562997591E-04

 1.37470209752348E-03
 2.11322007974217E-03

 3.17238945582342E-03
 2.50085216537535E-03

 2.85515051024108E-03
 3.02603112010418E-03

 7.11521635091824E-03
 4.67659354925191E-03

 1.41095797701861E-02
 .015930428293441

 1.56504546487289E-02
 .014817549079849

 9.97036686115932E-04
 1.75059651576275E-03

 1.46534179626129E-02
 1.30669525640862E-02

 3.3508137027888
 4.33390177703052

 1.73726089247473E-02
 1.64806157698236E-02

 3.0213232912604E-05
 0

 1.5106616456302E-05
 1.25042608268768E-05

 4.5319849368906E-05
 0

 4.5319849368906E-05
 3.75127824806303E-05

 6.0426465825208E-05
 5.0017043307507E-05

 1.20852931650416E-04
 2.25076694883782E-04

 2.11492630388228E-04
 1.75059651576275E-04

 1.96386013931926E-04
 2.37580955710658E-04

 6.0426465825208E-05
 6.00204519690084E-04

 1.5106616456302E-04
 6.62725823824468E-04

 6.49584507620986E-04
 3.00102259845042E-04

 8.15757288640308E-04
 1.87563912403151E-04

 2.87025712669738E-04
 8.62793997054496E-04

 6.7979774053359E-04
 6.75230084651345E-04

 9.21503603834422E-04
 5.75195998036331E-04

 1.07256976839744E-03
 5.75195998036331E-04

 1.02724991902854E-03
 6.25213041343838E-04

 1.78258074184364E-03
 1.81311781989713E-03

 2.53791156465874E-03
 3.02603112010418E-03

 6.66201785722918E-03
 5.83948980615145E-03

 8.15757288640308E-03
 7.17744571462726E-03

 0
 2.50085216537535E-05

 7.553308228151E-05
 2.50085216537535E-05

 1.20852931650416E-04
 2.37580955710658E-04

 3.0213232912604E-05
 5.87700258863208E-04

 2.02428660514447E-03
 1.47550277757146E-03

 5.90668703441408E-03
 4.82664467917443E-03

 .199301590907992
 .260626308414592

 1.5106616456302E-05
 1.25042608268768E-05

 3.0213232912604E-05
 3.75127824806303E-05

 1.5106616456302E-05
 6.25213041343838E-05

 9.0639698737812E-05
 6.25213041343838E-05

 1.05746315194114E-04
 1.50051129922521E-04

 7.64394792688881E-03
 4.52654241932939E-03

 5.49880839009393E-03
 6.67727528155219E-03

 1.07256976839744E-02
 .015730360120211

 1.85811382412515E-02
 2.25826950533394E-02

 .025862527373189
 3.36364616242985E-02

 .130732658812837
 .177147863134363

 .122695938858085
 .172971440018186

 0
 2.50085216537535E-05

 1.35959548106718E-04
 1.25042608268768E-04

 1.81279397475624E-04
 2.12572434056905E-04

 1.81279397475624E-04
 2.25076694883782E-04

 1.66172781019322E-04
 2.87597999018165E-04

 3.7766541140755E-04
 2.25076694883782E-04

 3.92772027863852E-04
 2.25076694883782E-04

 6.0426465825208E-04
 5.62691737209454E-04

 2.77961742795957E-03
 1.65056242914773E-03

 2.52280494820243E-03
 3.90132937798555E-03

 5.3175289926183E-03
 8.35284623235368E-03

 8.71651769528625E-03
 9.22814449023505E-03

 1.11335763282946E-02
 1.82062037639326E-02

 3.08477108037687E-02
 5.03171455673521E-02

 5.93387894403542E-02
 7.94270647723212E-02

 .319791963763457
 .390095425016074

 4.5319849368906E-05
 0

 1.5106616456302E-05
 2.50085216537535E-05

 6.0426465825208E-05
 2.50085216537535E-05

 1.5106616456302E-04
 5.0017043307507E-05

 1.20852931650416E-04
 1.12538347441891E-04

 1.5106616456302E-04
 1.25042608268768E-04

 1.96386013931926E-04
 1.25042608268768E-04

 3.32345562038644E-04
 6.25213041343838E-05

 8.45970521552912E-04
 4.37649128940687E-04

 6.0426465825208E-04
 7.37751388785729E-04

 8.3086390509661E-04
 6.75230084651345E-04

 1.26895578232937E-03
 1.15039199607266E-03

 2.06960645451337E-03
 1.25042608268768E-03

 2.9911100583478E-03
 2.27577547049157E-03

 2.94579020897889E-03
 2.73843312108601E-03

 5.71030102048216E-03
 4.13891033369621E-03

 3.38388208621165E-03
 6.28964319591901E-03

 1.38980871397978E-02
 1.28168673475487E-02

 1.79315537336305E-02
 1.78435801999531E-02

 2.01522263527069E-02
 2.41207191350453E-02

 3.05304718581863E-02
 2.83596635553565E-02

 3.56214016039601E-02
 .04900419818053

 4.76311616867202E-02
 5.55064138105059E-02

 .132303746924293
 .182224593030075

 .77273364497276
 .952562085530644

 0
 5.0017043307507E-05

 1.5106616456302E-05
 3.75127824806303E-05

 4.5319849368906E-05
 2.50085216537535E-05

 6.0426465825208E-05
 2.50085216537535E-05

 9.0639698737812E-05
 3.75127824806303E-05

 1.05746315194114E-04
 2.50085216537535E-05

 9.0639698737812E-05
 5.0017043307507E-05

 1.96386013931926E-04
 0

 2.11492630388228E-04
 2.50085216537535E-04

 3.47452178494946E-04
 3.00102259845042E-04

 3.47452178494946E-04
 4.75161911421317E-04

 6.94904356989892E-04
 4.37649128940687E-04

 7.85544055727704E-04
 5.87700258863208E-04

 8.91290370921818E-04
 6.12708780516961E-04

 9.36610220290724E-04
 7.00238606305098E-04

 9.36610220290724E-04
 9.2531530118888E-04

 1.22363593296046E-03
 9.75332344496387E-04

 1.90343367349405E-03
 1.08787069193828E-03

 1.96386013931926E-03
 1.36296443012957E-03

 2.2659924684453E-03
 1.56303260335959E-03

 2.32641893427051E-03
 2.32579251379908E-03

 3.45941516849316E-03
 3.0135268592773E-03

 4.10899967611414E-03
 .002638399034471

 7.5684148446073E-03
 .011266339005016

 1.20701865485853E-02
 .018768895501142

 4.96101284424958E-02
 5.44935686835289E-02

 .050773337909631
 5.80447787583619E-02

 5.29335840628822E-02
 5.70569421530387E-02

 .052873157597057
 5.97953752741247E-02

 5.23444260210864E-02
 6.03330584896804E-02

 .060275399660645
 6.62475738607931E-02

 7.56992550625293E-02
 9.05683611690684E-02

 8.04276260133518E-02
 .100996914698684

 .112710465380469
 .153277229215855

 .142440286566472
 .204207083563724

 .803551142543616
 1.15241768632662

 4.5319849368906E-05
 1.25042608268768E-05

 4.5319849368906E-05
 2.50085216537535E-05

 6.0426465825208E-05
 3.75127824806303E-05

 4.5319849368906E-05
 7.50255649612605E-05

 3.62558794951248E-04
 2.12572434056905E-04

 7.85544055727704E-04
 2.37580955710658E-04

 1.16320946713525E-03
 1.37546869095644E-04

 7.70437439271402E-04
 7.00238606305098E-04

 8.61077138009214E-04
 1.63805816832086E-03

 2.91557697606629E-03
 2.48834790454847E-03

 6.35988552810314E-03
 6.30214745674589E-03

 8.02161333829636E-03
 5.83948980615145E-03

 1.02724991902854E-02
 7.91519710341299E-03

 1.19795468498475E-02
 2.45833767856397E-02

 .047570735220895
 4.81163956618218E-02

 9.33286764670337E-02
 .146462407065207

 .618963396064062
 .907634276379676

 5.13624959514268E-04
 6.12708780516961E-04

 7.553308228151E-05
 0

 9.0639698737812E-05
 5.0017043307507E-05

 9.0639698737812E-05
 7.50255649612605E-05

 2.56812479757134E-04
 4.87666172248194E-04

 .683362902017277
 .927828657615082

 7.553308228151E-05
 0

 6.0426465825208E-05
 2.50085216537535E-05

 4.5319849368906E-05
 5.0017043307507E-05

 7.553308228151E-05
 1.25042608268768E-04

 1.05746315194114E-04
 1.25042608268768E-04

 1.81279397475624E-04
 6.25213041343838E-05

 1.66172781019322E-04
 1.25042608268768E-04

 3.17238945582342E-04
 8.75298257881373E-05

 1.96386013931926E-04
 2.12572434056905E-04

 3.92772027863852E-04
 2.25076694883782E-04

 4.22985260776456E-04
 2.12572434056905E-04

 4.22985260776456E-04
 2.50085216537535E-04

 3.92772027863852E-04
 3.62623563979426E-04

 4.83411726601664E-04
 4.00136346460056E-04

 4.38091877232758E-04
 4.87666172248194E-04

 4.98518343057966E-04
 5.12674693901947E-04

 9.66823453203328E-04
 7.50255649612605E-04

 1.04235653548484E-03
 8.00272692920113E-04

 1.93364690640666E-03
 2.06320303643467E-03

 8.52013168135433E-03
 7.6401033652217E-03

 8.71651769528625E-03
 1.09037154410365E-02

 1.32636092486332E-02
 2.08696113200573E-02

 3.01528064467788E-02
 3.72001759599584E-02

 3.42769127393492E-02
 4.08639243822332E-02

 3.38992473279417E-02
 4.60156798429065E-02

 5.02446063336604E-02
 .056719327110713

 4.84771322082731E-02
 8.15027720695827E-02

 .057374929301035
 9.06683952556834E-02

 6.84178659305918E-02
 8.19029084160428E-02

 .071182376742095
 8.28032151955779E-02

 .118692685497165
 .175509804966042

 .131926081512885
 .188351680835245

 5.60153338199678E-02
 8.40911540607462E-02

 6.0426465825208E-05
 2.50085216537535E-05

 1.05746315194114E-04
 1.00034086615014E-04

 1.35959548106718E-04
 2.12572434056905E-04

 5.57132014908418E-02
 8.37535390184205E-02

 1.21910394802357E-02
 1.42923701251201E-02

 3.0213232912604E-05
 6.25213041343838E-05

 0
 2.25076694883782E-04

 3.92772027863852E-04
 2.75093738191289E-04

 1.96386013931926E-04
 5.25178954728824E-04

 4.07878644320154E-04
 3.62623563979426E-04

 2.87025712669738E-04
 5.37683215555701E-04

 5.43838192426872E-04
 4.12640607286933E-04

 1.66172781019322E-04
 7.50255649612605E-04

 4.38091877232758E-04
 5.37683215555701E-04

 7.70437439271402E-04
 7.62759910439482E-04

 7.85544055727704E-04
 1.05035790945765E-03

 1.08767638485374E-03
 1.52551982087896E-03

 2.32641893427051E-03
 2.18824564470343E-03

 2.14513953679488E-03
 2.40081807876034E-03

 2.61344464694025E-03
 2.67591181695163E-03

 4.18302209675002E-02
 5.27804849502468E-02

 9.0639698737812E-05
 6.25213041343838E-05

 1.96386013931926E-04
 2.50085216537535E-05

 4.98518343057966E-04
 4.6265765059444E-04

 8.15757288640308E-04
 8.12776953746989E-04

 9.8193006965963E-04
 7.00238606305098E-04

 9.0639698737812E-04
 7.75264171266359E-04

 7.553308228151E-04
 9.75332344496387E-04

 3.89750704572592E-03
 3.60122711814051E-03

 4.59241140271581E-03
 4.13891033369621E-03

 2.90953432948376E-02
 4.12265479462127E-02

 5.32055031590956E-02
 4.95668899177395E-02

 3.0213232912604E-05
 1.25042608268768E-04

 9.0639698737812E-05
 3.12606520671919E-04

 3.0213232912604E-04
 3.37615042325672E-04

 4.22985260776456E-04
 3.62623563979426E-04

 4.98518343057966E-04
 3.75127824806303E-04

 6.49584507620986E-04
 2.62589477364412E-04

 4.22985260776456E-04
 4.6265765059444E-04

 7.25117589902496E-04
 4.12640607286933E-04

 6.49584507620986E-04
 5.87700258863208E-04

 6.94904356989892E-04
 9.00306779535127E-04

 9.0639698737812E-04
 7.50255649612605E-04

 2.59833803048394E-03
 1.53802408170584E-03

 2.56812479757134E-03
 3.02603112010418E-03

 3.29324238747384E-03
 2.86347572935478E-03

 3.93527358686667E-02
 3.72501930032659E-02

 .125838115080996
 .111362946924164

 1.5106616456302E-04
 7.50255649612605E-05

 1.66172781019322E-04
 1.50051129922521E-04

 2.87025712669738E-04
 1.00034086615014E-04

 2.87025712669738E-04
 4.87666172248194E-04

 6.19371274708382E-04
 4.12640607286933E-04

 1.58619472791171E-03
 9.87836605323264E-04

 2.40195201655202E-03
 1.87563912403151E-03

 8.94311694213078E-03
 7.67761614770233E-03

 1.47138444284381E-02
 1.34295761280656E-02

 3.21317732025544E-02
 1.61680092491516E-02

 6.45505721177784E-02
 6.99988521088561E-02

 3.62709861115811E-02
 3.06354390258481E-02

 2.11492630388228E-04
 8.75298257881373E-05

 4.22985260776456E-04
 2.50085216537535E-05

 7.553308228151E-04
 3.62623563979426E-04

 6.7979774053359E-04
 5.0017043307507E-04

 6.52605830912246E-03
 6.0770707618621E-03

 8.85247724339297E-03
 6.80231788982096E-03

 1.88228441045523E-02
 1.67807180296686E-02

 5.2873157597057E-03
 5.05172137405821E-03

 9.0639698737812E-05
 2.62589477364412E-04

 4.68305110145362E-04
 4.37649128940687E-04

 6.34477891164684E-04
 7.50255649612605E-04

 6.49584507620986E-04
 7.87768432093236E-04

 1.61640796082431E-03
 1.37546869095644E-03

 1.82790059121254E-03
 1.43798999509083E-03

 1.25687048916433E-02
 1.65931541172655E-02

 3.32345562038644E-04
 2.37580955710658E-04

 3.62558794951248E-04
 3.12606520671919E-04

 6.0426465825208E-04
 7.87768432093236E-04

 6.94904356989892E-04
 7.50255649612605E-04

 1.26895578232937E-03
 8.62793997054496E-04

 8.45970521552912E-04
 1.48800703839833E-03

 1.38980871397978E-03
 2.41332233958721E-03

 2.2810990849016E-03
 2.11322007974217E-03

 2.17535276970749E-03
 2.95100555514291E-03

 2.61344464694025E-03
 4.67659354925191E-03

 4.60751801917211E-02
 5.35307405998594E-02

 4.83411726601664E-04
 2.75093738191289E-04

 8.76183754465516E-04
 8.75298257881373E-04

 1.45023517980499E-03
 6.50221562997591E-04

 1.10278300131005E-03
 1.00034086615014E-03

 9.78908746368369E-03
 1.06161174420184E-02

 1.16320946713525E-02
 1.34670889105463E-02

 2.07413843945026E-02
 2.66465798220744E-02

 3.40503134925047E-02
 3.52245027493118E-02

 4.38091877232758E-04
 5.25178954728824E-04

 7.10010973446194E-04
 4.75161911421317E-04

 8.91290370921818E-04
 6.12708780516961E-04

 2.06960645451337E-03
 1.03785364863077E-03

 2.43216524946462E-03
 2.12572434056905E-03

 9.15460957251901E-03
 1.38297124745257E-02

 1.83545389944069E-02
 1.66181626389192E-02

 1.62532086453353
 1.8677239269281

 1.43218277313971
 1.65890277111926

 1.5106616456302E-05
 1.25042608268768E-05

 4.5319849368906E-05
 0

 3.0213232912604E-05
 1.25042608268768E-05

 3.0213232912604E-05
 1.25042608268768E-05

 3.0213232912604E-05
 2.50085216537535E-05

 6.0426465825208E-05
 0

 7.553308228151E-05
 1.25042608268768E-05

 3.0213232912604E-05
 6.25213041343838E-05

 3.0213232912604E-05
 1.00034086615014E-04

 1.35959548106718E-04
 1.25042608268768E-05

 1.20852931650416E-04
 6.25213041343838E-05

 6.0426465825208E-05
 1.12538347441891E-04

 7.553308228151E-05
 1.12538347441891E-04

 1.81279397475624E-04
 5.0017043307507E-05

 1.81279397475624E-04
 6.25213041343838E-05

 6.0426465825208E-05
 1.75059651576275E-04

 1.05746315194114E-04
 2.12572434056905E-04

 1.96386013931926E-04
 1.75059651576275E-04

 1.20852931650416E-04
 3.00102259845042E-04

 3.32345562038644E-04
 1.87563912403151E-04

 4.22985260776456E-04
 3.62623563979426E-04

 9.36610220290724E-04
 5.50187476382577E-04

 1.05746315194114E-03
 5.87700258863208E-04

 2.32641893427051E-03
 1.42548573426395E-03

 2.13003292033858E-03
 2.13822860139593E-03

 2.13003292033858E-03
 2.33829677462595E-03

 1.99407337223186E-03
 2.60088625199037E-03

 2.50769833174613E-03
 2.22575842718406E-03

 3.59537471659988E-03
 3.15107372837294E-03

 .005302422376162
 3.15107372837294E-03

 5.15135621159898E-03
 4.52654241932939E-03

 5.69519440402585E-03
 4.43901259354125E-03

 8.80715739402406E-03
 5.15175546067322E-03

 9.45674190164505E-03
 6.88984771560909E-03

 .011752947603003
 9.92838309654015E-03

 1.18738005346534E-02
 1.04660663120958E-02

 .015907267128486
 .011766509438091

 1.79013405007179E-02
 1.30794568249131E-02

 1.26442379739248E-02
 1.75059651576275E-02

 2.26146048350841E-02
 .014404908472562

 2.13154358198421E-02
 2.55462048693092E-02

 2.44425054262966E-02
 2.31078740080682E-02

 .028460865403673
 2.00568343663103E-02

 3.01981262961477E-02
 .036624979961922

 3.16785747088653E-02
 3.69375864825939E-02

 3.85218719635701E-02
 .039976121863525

 3.87937910597835E-02
 5.59815757219272E-02

 4.59090074107018E-02
 5.22928187779986E-02

 .067028057216612
 5.10173841736572E-02

 5.46859515718132E-02
 6.45970114316453E-02

 8.05937987943712E-02
 6.17960570064249E-02

 7.15147223041337E-02
 .072549721317539

 .158000101516463
 .188514236225994

 .200857572402991
 .273430671501314

 .394056090262638
 .538083351902161

 7.06989650154934E-03
 5.58940458961391E-03

 1.35959548106718E-04
 1.25042608268768E-04

 2.41705863300832E-04
 2.62589477364412E-04

 3.0213232912604E-04
 3.00102259845042E-04

 4.38091877232758E-04
 4.12640607286933E-04

 6.64691124077288E-04
 4.50153389767563E-04

 5.43838192426872E-04
 7.12742867131975E-04

 8.61077138009214E-04
 5.12674693901947E-04

 1.02724991902854E-03
 4.6265765059444E-04

 7.553308228151E-04
 7.62759910439482E-04

 9.21503603834422E-04
 6.75230084651345E-04

 1.17831608359156E-03
 9.12811040362003E-04

 8.15757288640308E-04
 3.23860355416108E-03

 7.553308228151E-05
 3.12606520671919E-04

 3.0213232912604E-05
 3.50119303152549E-04

 1.5106616456302E-04
 2.62589477364412E-04

 9.0639698737812E-05
 3.25110781498796E-04

 1.05746315194114E-04
 4.2514486811381E-04

 1.20852931650416E-04
 4.12640607286933E-04

 6.0426465825208E-05
 6.00204519690084E-04

 1.81279397475624E-04
 5.50187476382577E-04

 .05302422376162
 .065134694647201

 3.0213232912604E-04
 2.87597999018165E-04

 2.56812479757134E-04
 3.50119303152549E-04

 3.0213232912604E-04
 4.6265765059444E-04

 5.13624959514268E-04
 3.37615042325672E-04

 6.19371274708382E-04
 6.87734345478222E-04

 1.61640796082431E-03
 7.75264171266359E-04

 4.94137424285638E-02
 6.22337061353656E-02

 4.88245843867681E-02
 6.44844730842034E-02

 3.0213232912604E-04
 3.25110781498796E-04

 3.92772027863852E-04
 3.37615042325672E-04

 2.2659924684453E-04
 5.87700258863208E-04

 5.43838192426872E-04
 4.75161911421317E-04

 5.2873157597057E-04
 5.12674693901947E-04

 4.38091877232758E-04
 6.25213041343838E-04

 1.01214330257223E-03
 6.25213041343838E-04

 8.3086390509661E-04
 8.12776953746989E-04

 1.46534179626129E-03
 1.46299851674458E-03

 4.30840701333733E-02
 5.87200088430133E-02

 4.38091877232758E-04
 2.12572434056905E-04

 4.38091877232758E-04
 2.12572434056905E-04

 4.42925994498775E-02
 3.74877739589765E-02

 4.68305110145362E-04
 2.37580955710658E-04

 4.07878644320154E-04
 3.25110781498796E-04

 6.49584507620986E-04
 4.2514486811381E-04

 6.7979774053359E-04
 4.2514486811381E-04

 .010106326409266
 8.30282918904617E-03

 3.19807070379913E-02
 2.77719632964933E-02

 2.80831999922654E-02
 2.29203100956651E-02

 4.07878644320154E-04
 3.00102259845042E-04

 6.19371274708382E-04
 3.00102259845042E-04

 7.25117589902496E-04
 3.62623563979426E-04

 7.25117589902496E-04
 4.6265765059444E-04

 7.40224206358798E-04
 8.37785475400743E-04

 7.85544055727704E-04
 9.87836605323264E-04

 9.51716836747026E-04
 1.08787069193828E-03

 1.13299623422265E-03
 1.17540051772642E-03

 2.74940419504696E-03
 2.20074990553031E-03

 2.76451081150327E-03
 2.71342459943226E-03

 3.70112103179399E-03
 2.58838199116349E-03

 3.62558794951248E-03
 2.95100555514291E-03

 4.30538569004607E-03
 2.92599703348916E-03

 4.84922388247294E-03
 4.02637198625432E-03

 7.25117589902496E-04
 1.00034086615014E-04

 7.25117589902496E-04
 1.00034086615014E-04

 9.8646205459652E-03
 9.65328935834886E-03

 8.3086390509661E-04
 5.87700258863208E-04

 9.03375664086859E-03
 9.06558909948565E-03

 1.26330590777471
 1.199121100515

 1.07377829771395
 1.14480259148305

 3.0213232912604E-05
 0

 3.0213232912604E-05
 0

 3.0213232912604E-05
 6.25213041343838E-05

 1.20852931650416E-04
 1.25042608268768E-05

 6.94904356989892E-04
 1.00034086615014E-04

 4.68305110145362E-04
 7.00238606305098E-04

 6.64691124077288E-04
 5.87700258863208E-04

 1.02724991902854E-03
 6.12708780516961E-04

 1.73726089247473E-03
 1.17540051772642E-03

 1.19342270004786E-03
 1.82562208072401E-03

 2.64365787985285E-03
 2.21325416635719E-03

 1.76898478703296E-02
 5.1642597215001E-03

 1.59828002107675E-02
 1.40547891694095E-02

 2.10284101071724E-02
 1.46299851674458E-02

 1.79315537336305E-02
 1.99568002796953E-02

 4.07727578155591E-02
 5.97453582308171E-02

 4.60902868081774E-02
 6.74104701176926E-02

 5.68159844921518E-02
 6.50471648214129E-02

 7.94305893272359E-02
 8.73172533540804E-02

 9.93713230495545E-02
 .081477763547929

 9.10475773821321E-02
 .110350101797187

 .10784613488154
 .101134461567779

 .109825101637316
 .109112179975327

 .361304945785375
 .402112019670703

 1.31427563169827E-02
 5.3268151122495E-03

 6.0426465825208E-05
 0

 3.0213232912604E-05
 3.75127824806303E-05

 1.96386013931926E-04
 5.0017043307507E-05

 3.17238945582342E-04
 2.37580955710658E-04

 9.15460957251901E-03
 1.00034086615014E-04

 3.38388208621165E-03
 4.90167024413569E-03

 .121275916911192
 3.75127824806303E-05

 7.553308228151E-05
 0

 4.5319849368906E-04
 0

 4.68305110145362E-04
 0

 4.83411726601664E-04
 0

 4.98518343057966E-04
 0

 8.45970521552912E-04
 0

 9.36610220290724E-04
 0

 1.55598149499911E-03
 0

 2.03939322160077E-03
 0

 2.06960645451337E-03
 0

 2.43216524946462E-03
 0

 2.68897772922176E-03
 0

 3.33856223684274E-03
 0

 3.39898870266795E-03
 0

 3.45941516849316E-03
 0

 3.89750704572592E-03
 0

 4.18453275839565E-03
 0

 4.45645185460909E-03
 0

 5.3175289926183E-03
 0

 5.51391500655023E-03
 0

 5.92179365087038E-03
 0

 6.75265755596699E-03
 0

 7.5835214610636E-03
 0

 8.70141107882995E-03
 0

 9.03375664086859E-03
 1.25042608268768E-05

 9.90994039533411E-03
 1.25042608268768E-05

 1.21003997814979E-02
 1.25042608268768E-05

 .013157862933439
 0

 9.54738160038286E-03
 2.58838199116349E-03

 1.81279397475624E-04
 1.00034086615014E-04

 1.81279397475624E-04
 3.37615042325672E-04

 5.13624959514268E-04
 1.00034086615014E-04

 1.61640796082431E-03
 8.12776953746989E-04

 1.88832705703775E-03
 1.22541756103392E-03

 5.16646282805528E-03
 1.25042608268768E-05

 4.55615552322068E-02
 .046365799146059

 2.53791156465874E-03
 0

 2.47295311389664E-02
 .019781740628119

 1.82941125285817E-02
 .02658405851794

 .357573611520668
 .419605480567503

 5.65289587794821E-02
 7.33374897496322E-02

 3.0213232912604E-05
 1.25042608268768E-05

 1.81279397475624E-04
 0

 1.81279397475624E-04
 5.0017043307507E-05

 2.2659924684453E-04
 3.75127824806303E-05

 3.32345562038644E-04
 1.87563912403151E-04

 3.0213232912604E-04
 4.87666172248194E-04

 6.0426465825208E-04
 3.00102259845042E-04

 6.7979774053359E-04
 4.37649128940687E-04

 5.43838192426872E-04
 6.62725823824468E-04

 6.34477891164684E-04
 7.00238606305098E-04

 6.94904356989892E-04
 7.00238606305098E-04

 8.15757288640308E-04
 6.25213041343838E-04

 1.42002194689239E-03
 9.75332344496387E-04

 1.32938224815458E-03
 1.4504942559177E-03

 2.94579020897889E-03
 1.91315190651214E-03

 3.55005486723097E-03
 1.76310077658962E-03

 4.48666508752169E-03
 1.41298147343707E-03

 3.47452178494946E-03
 2.88848425100853E-03

 5.75562086985106E-03
 5.18926824315386E-03

 2.83400124720225E-02
 5.35432448606863E-02

 3.0213232912604E-04
 1.50051129922521E-04

 0
 5.0017043307507E-05

 3.0213232912604E-04
 1.00034086615014E-04

 .259833803048394
 .310743385808714

 7.553308228151E-05
 0

 4.5319849368906E-05
 3.75127824806303E-05

 4.5319849368906E-05
 5.0017043307507E-05

 7.553308228151E-05
 2.50085216537535E-05

 1.5106616456302E-04
 1.25042608268768E-05

 1.66172781019322E-04
 3.75127824806303E-05

 1.35959548106718E-04
 1.75059651576275E-04

 3.0213232912604E-04
 5.0017043307507E-05

 1.5106616456302E-04
 2.25076694883782E-04

 4.22985260776456E-04
 1.25042608268768E-05

 3.62558794951248E-04
 2.50085216537535E-04

 3.47452178494946E-04
 3.00102259845042E-04

 6.49584507620986E-04
 6.12708780516961E-04

 6.19371274708382E-04
 7.50255649612605E-04

 7.85544055727704E-04
 7.12742867131975E-04

 8.15757288640308E-04
 7.25247127958852E-04

 8.15757288640308E-04
 7.50255649612605E-04

 9.36610220290724E-04
 7.50255649612605E-04

 2.08471307096968E-03
 3.50119303152549E-04

 1.16320946713525E-03
 1.20040903938017E-03

 1.34448886461088E-03
 1.11287921359203E-03

 2.03939322160077E-03
 1.43798999509083E-03

 4.15431952548305E-03
 3.30112485829546E-03

 1.16774145207214E-02
 4.31396998527248E-03

 1.36110614271281E-02
 7.27747980124227E-03

 1.57713075803793E-02
 9.19063170775442E-03

 1.73121824589221E-02
 .022520173749205

 1.90494433513968E-02
 3.27236505839365E-02

 4.39451472713825E-02
 6.06831777928329E-02

 4.77520146183706E-02
 6.72229062052895E-02

 7.30253839497639E-02
 9.39320073314982E-02

 3.32194495874081E-02
 3.14607202404219E-02

 4.5319849368906E-05
 3.75127824806303E-05

 7.553308228151E-05
 2.50085216537535E-05

 1.35959548106718E-04
 1.87563912403151E-04

 1.96386013931926E-04
 1.87563912403151E-04

 2.41705863300832E-04
 2.12572434056905E-04

 2.11492630388228E-04
 3.62623563979426E-04

 3.62558794951248E-04
 3.50119303152549E-04

 2.32641893427051E-03
 2.23826268801094E-03

 2.96240748708082E-02
 2.78594931222814E-02

 7.68926777625772E-03
 3.91383363881243E-03

 4.83411726601664E-04
 1.00034086615014E-04

 8.76183754465516E-04
 1.50051129922521E-04

 1.79768735829994E-03
 1.12538347441891E-04

 1.25384916587307E-03
 9.75332344496387E-04

 3.27813577101753E-03
 2.57587773033661E-03

 .947199958426591
 .9117856909742

 8.49596109502424E-02
 7.12992952348513E-02

 0
 3.75127824806303E-05

 9.0639698737812E-05
 1.00034086615014E-04

 5.58944808883174E-04
 5.37683215555701E-04

 1.55144951006222E-02
 .012279184131993

 2.12550093540169E-02
 1.26293034351455E-02

 4.75405219879824E-02
 4.57155775830614E-02

 8.98088348327154E-02
 8.44037605814181E-02

 1.5106616456302E-05
 3.75127824806303E-05

 4.83411726601664E-04
 5.0017043307507E-05

 4.38091877232758E-04
 2.62589477364412E-04

 8.91290370921818E-04
 6.25213041343838E-05

 2.00917998868817E-03
 9.2531530118888E-04

 2.59833803048394E-03
 2.11322007974217E-03

 3.20260268873602E-03
 2.28827973131845E-03

 2.71919096213436E-03
 2.73843312108601E-03

 3.82197396344441E-03
 2.91349277266228E-03

 2.56812479757134E-02
 2.61589136498262E-02

 4.79484006323025E-02
 4.68534653183072E-02

 .200253307744739
 .187213793099999

 1.5106616456302E-04
 1.00034086615014E-04

 1.5106616456302E-04
 1.25042608268768E-04

 2.2659924684453E-04
 1.50051129922521E-04

 2.56812479757134E-04
 2.25076694883782E-04

 4.98518343057966E-04
 6.25213041343838E-05

 5.74051425339476E-04
 2.00068173230028E-04

 5.2873157597057E-04
 4.87666172248194E-04

 1.45023517980499E-03
 3.25110781498796E-04

 3.14217622291082E-03
 2.13822860139593E-03

 1.19795468498475E-02
 1.48050448190221E-02

 5.63778926149191E-02
 6.67727528155219E-02

 .124916611477161
 .101822195913257

 .284805040050662
 .295338136470002

 1.10278300131005E-03
 7.87768432093236E-04

 1.12242160270324E-02
 6.43969432584153E-03

 6.16803149910811E-02
 7.04990225419312E-02

 .210797726031238
 .217611651170136

 .172366493766406
 .165743977260251

 7.62884131043251E-03
 6.35216450005339E-03

 8.98843679149969E-03
 6.45219858666841E-03

 .015257682620865
 5.06422563488509E-03

 7.00644871243287E-02
 7.11992611482363E-02

 7.04270459192799E-02
 7.66761273904083E-02

 .115006671081827
 .107786728327678

 2.70861633061495E-02
 2.79595272088964E-02

 3.59688537824551E-02
 2.49334960887923E-02

 5.19516539932226E-02
 .054893705029989

 .412969574065928
 .432797475739858

 5.57132014908418E-02
 4.46026983694694E-02

 3.0213232912604E-05
 1.25042608268768E-05

 1.35959548106718E-04
 1.25042608268768E-05

 3.0213232912604E-05
 1.50051129922521E-04

 3.7766541140755E-04
 4.00136346460056E-04

 1.57108811145541E-03
 8.12776953746989E-04

 5.35680619540469E-02
 4.32147254176861E-02

 1.81279397475624E-04
 1.75059651576275E-04

 9.0639698737812E-05
 1.25042608268768E-05

 3.0213232912604E-05
 8.75298257881373E-05

 6.0426465825208E-05
 7.50255649612605E-05

 5.83115395213257E-03
 4.52654241932939E-03

 3.0213232912604E-05
 8.75298257881373E-05

 1.35959548106718E-04
 5.0017043307507E-05

 7.553308228151E-05
 1.25042608268768E-04

 2.2659924684453E-04
 8.75298257881373E-05

 3.32345562038644E-04
 1.75059651576275E-04

 5.74051425339476E-04
 4.6265765059444E-04

 1.05746315194114E-03
 5.37683215555701E-04

 1.82790059121254E-03
 1.26293034351455E-03

 1.57108811145541E-03
 1.73809225493587E-03

 8.97181951339776E-02
 .095269963239974

 6.0426465825208E-05
 1.00034086615014E-04

 5.89158041795778E-04
 4.87666172248194E-04

 6.0426465825208E-04
 5.12674693901947E-04

 1.4955550291739E-03
 4.37649128940687E-04

 1.34448886461088E-03
 8.12776953746989E-04

 8.56243020743197E-02
 9.29191622045212E-02

 8.74068828161634E-02
 .11410138004525

 1.81279397475624E-04
 3.75127824806303E-05

 1.5106616456302E-04
 7.50255649612605E-05

 5.43838192426872E-04
 3.00102259845042E-04

 7.553308228151E-04
 5.50187476382577E-04

 7.553308228151E-04
 6.00204519690084E-04

 1.48649105930012E-02
 1.55427962078078E-02

 1.65115317867381E-02
 2.61214008673455E-02

 2.45331451250344E-02
 3.12356435455381E-02

 2.91104499112939E-02
 3.96385068211993E-02

 4.5319849368906E-05
 1.87563912403151E-04

 4.5319849368906E-05
 1.87563912403151E-04

 1.72970758424658E-02
 .020294415322021

 3.32345562038644E-04
 1.12538347441891E-04

 7.40224206358798E-04
 8.00272692920113E-04

 1.62245060740683E-02
 .019381604281659

 7.44756191295688E-03
 6.10207928351586E-03

 8.76183754465516E-04
 0

 1.32938224815458E-03
 7.25247127958852E-04

 2.20556600262009E-03
 3.12606520671919E-04

 3.0364299077167E-03
 5.06422563488509E-03

 .149328903670545
 .147537773496319

 3.73133426470659E-03
 2.61339051281724E-03

 7.73458762562662E-03
 5.71444719788268E-03

 1.69949435133397E-02
 2.58463071291543E-02

 .120868038266872
 .113363628656465

 .415643445178693
 .407463843304606

 1.35959548106718E-04
 1.25042608268768E-04

 3.0213232912604E-05
 1.25042608268768E-05

 0
 6.25213041343838E-05

 6.0426465825208E-05
 1.25042608268768E-05

 4.5319849368906E-05
 3.75127824806303E-05

 .193138091393821
 .175509804966042

 3.0213232912604E-05
 1.25042608268768E-05

 6.7979774053359E-04
 4.6265765059444E-04

 2.16024615325119E-03
 1.88814338485839E-03

 2.44727186592092E-03
 2.07570729726154E-03

 3.24792253810493E-03
 2.86347572935478E-03

 4.62262463562841E-03
 2.27577547049157E-03

 5.18156944451159E-03
 3.32613337994922E-03

 6.31456567873423E-03
 4.88916598330881E-03

 8.82226401048037E-03
 6.55223267328342E-03

 8.91290370921818E-03
 7.11492441049288E-03

 5.64836389301132E-02
 7.20620551452908E-02

 9.42350734544119E-02
 7.19870295803295E-02

 1.82336860627565E-02
 1.64931200306504E-02

 0
 3.75127824806303E-05

 6.0426465825208E-05
 1.75059651576275E-04

 3.7766541140755E-04
 2.87597999018165E-04

 5.58944808883174E-04
 3.25110781498796E-04

 6.49584507620986E-04
 7.12742867131975E-04

 1.91854028995035E-03
 1.41298147343707E-03

 2.68897772922176E-03
 1.57553686418647E-03

 1.13299623422265E-03
 3.60122711814051E-03

 4.18453275839565E-03
 3.16357798919982E-03

 6.66201785722918E-03
 5.20177250398073E-03

 .101607102285087
 .106748874679047

 3.0213232912604E-05
 5.0017043307507E-05

 6.0426465825208E-05
 5.0017043307507E-05

 4.5319849368906E-05
 6.25213041343838E-05

 1.05746315194114E-04
 1.75059651576275E-04

 2.71919096213436E-04
 1.37546869095644E-04

 1.5106616456302E-04
 2.87597999018165E-04

 1.35959548106718E-04
 3.62623563979426E-04

 3.7766541140755E-04
 2.62589477364412E-04

 5.13624959514268E-04
 3.50119303152549E-04

 4.98518343057966E-04
 5.62691737209454E-04

 5.2873157597057E-04
 7.25247127958852E-04

 6.7979774053359E-04
 6.25213041343838E-04

 6.64691124077288E-04
 7.37751388785729E-04

 1.42002194689239E-03
 1.25042608268768E-03

 1.43512856334869E-03
 1.31294738682206E-03

 1.25384916587307E-03
 1.72558799410899E-03

 1.5257682620865E-03
 1.81311781989713E-03

 1.69194104310582E-03
 2.1507328622228E-03

 2.13003292033858E-03
 2.1507328622228E-03

 1.90343367349405E-03
 2.56337346950974E-03

 3.42920193558055E-03
 4.06388476873495E-03

 6.22392597999642E-03
 7.13993293214663E-03

 2.58927406061016E-02
 3.34113849294147E-02

 5.06373783615243E-02
 4.47777580210457E-02

 5.74202491504039E-02
 5.87950344079745E-02

 0
 1.37546869095644E-04

 3.17238945582342E-04
 1.50051129922521E-04

 5.2873157597057E-04
 2.25076694883782E-04

 3.0213232912604E-04
 7.00238606305098E-04

 7.553308228151E-04
 6.25213041343838E-04

 6.19371274708382E-04
 9.37819562015757E-04

 1.23874254941676E-03
 1.02534938780389E-03

 1.63151457728062E-03
 1.36296443012957E-03

 1.07256976839744E-03
 1.86313486320464E-03

 1.61640796082431E-03
 2.21325416635719E-03

 1.5106616456302E-03
 2.38831381793346E-03

 2.71919096213436E-03
 1.6130496466671E-03

 2.20556600262009E-03
 .002638399034471

 1.73726089247473E-03
 4.11390181204245E-03

 5.95200688378299E-03
 3.7637825088899E-03

 6.52605830912246E-03
 7.87768432093236E-03

 8.11225303703417E-03
 8.59042718806433E-03

 8.26331920159719E-03
 9.01557205617814E-03

 1.23118924118861E-02
 9.55325527173384E-03

 4.09540372130347E-02
 4.50903645417176E-02

 7.553308228151E-05
 1.25042608268768E-04

 1.66172781019322E-04
 1.87563912403151E-04

 2.41705863300832E-04
 1.62555390749398E-04

 3.7766541140755E-04
 2.12572434056905E-04

 3.17238945582342E-04
 2.87597999018165E-04

 4.07878644320154E-04
 2.37580955710658E-04

 1.96386013931926E-04
 4.75161911421317E-04

 4.07878644320154E-04
 4.00136346460056E-04

 4.98518343057966E-04
 4.2514486811381E-04

 5.89158041795778E-04
 4.00136346460056E-04

 5.13624959514268E-04
 4.75161911421317E-04

 5.58944808883174E-04
 5.50187476382577E-04

 5.74051425339476E-04
 7.50255649612605E-04

 1.26895578232937E-03
 1.48800703839833E-03

 1.79768735829994E-03
 1.53802408170584E-03

 2.06960645451337E-03
 1.78810929824338E-03

 1.81279397475624E-03
 2.07570729726154E-03

 1.61640796082431E-03
 2.27577547049157E-03

 1.94875352286296E-03
 3.13856946754607E-03

 2.9911100583478E-03
 3.30112485829546E-03

 3.29324238747384E-03
 3.82630381302429E-03

 3.55005486723097E-03
 5.12674693901947E-03

 9.23014265480052E-03
 6.76480510734033E-03

 6.45052522684095E-03
 9.07809336031253E-03

 4.15431952548305E-03
 4.70160207090566E-03

 1.20852931650416E-04
 1.12538347441891E-04

 1.20852931650416E-04
 2.12572434056905E-04

 1.96386013931926E-04
 2.50085216537535E-04

 3.62558794951248E-04
 4.12640607286933E-04

 1.37470209752348E-03
 1.56303260335959E-03

 1.97896675577556E-03
 2.1507328622228E-03

 1.04312697292411
 1.21438880298462

 5.58944808883174E-04
 1.63805816832086E-03

 0
 5.0017043307507E-05

 3.0213232912604E-05
 5.0017043307507E-05

 1.5106616456302E-05
 1.25042608268768E-04

 4.5319849368906E-05
 1.75059651576275E-04

 6.0426465825208E-05
 3.50119303152549E-04

 1.66172781019322E-04
 3.62623563979426E-04

 2.41705863300832E-04
 5.25178954728824E-04

 1.81279397475624E-03
 3.75127824806303E-04

 6.0426465825208E-05
 1.25042608268768E-05

 1.75236750893103E-03
 3.62623563979426E-04

 .368238882738817
 .367775319440099

 6.0426465825208E-05
 1.25042608268768E-05

 1.07256976839744E-03
 8.12776953746989E-04

 1.57108811145541E-03
 5.75195998036331E-04

 1.73726089247473E-03
 1.2379218218608E-03

 1.16774145207214E-02
 1.73809225493587E-03

 8.50502506489802E-03
 6.7773093681672E-03

 2.00766932704254E-02
 1.89314508918914E-02

 4.79030807829336E-02
 .033486410494376

 5.35378487211343E-02
 6.45845071708185E-02

 .222097475140552
 .239619150225439

 1.20852931650416E-04
 1.25042608268768E-05

 1.20852931650416E-04
 1.25042608268768E-05

 .672395498470002
 .844587793290564

 2.87025712669738E-04
 2.62589477364412E-04

 2.68897772922176E-03
 2.38831381793346E-03

 5.43838192426872E-03
 4.42650833271437E-03

 7.22096266611235E-03
 1.44174127333889E-02

 1.87473110222708E-02
 2.53586409569061E-02

 4.56975147803135E-02
 6.31465171757276E-02

 5.25408120350183E-02
 .06189609109304

 6.00488004138004E-02
 8.59792974456046E-02

 8.84794525845608E-02
 .11170056196649

 .100323039886302
 .129269048428252

 .290923219715464
 .345742811863142

 .184859665575768
 .238506271011847

 .184859665575768
 .238506271011847

 1.63151457728062E-03
 6.37717302170715E-04

 2.06960645451337E-03
 1.03785364863077E-03

 3.58026810014357E-03
 4.93918302661632E-03

 7.98233613550998E-02
 .118802982116156

 9.77549150887302E-02
 .113088534918273

 5.44679630963488
 5.59443130246632

 2.39235931510227
 2.50772950883013

 .237279624679135
 .278444880092892

 1.5106616456302E-05
 1.25042608268768E-05

 9.0639698737812E-05
 8.75298257881373E-05

 2.56812479757134E-04
 3.75127824806303E-05

 3.17238945582342E-04
 1.25042608268768E-04

 3.62558794951248E-04
 1.12538347441891E-04

 4.68305110145362E-04
 2.50085216537535E-05

 2.11492630388228E-04
 3.62623563979426E-04

 7.553308228151E-04
 2.50085216537535E-05

 6.94904356989892E-04
 3.62623563979426E-04

 2.41705863300832E-04
 8.00272692920113E-04

 1.29916901524197E-03
 5.0017043307507E-04

 1.23874254941676E-03
 6.50221562997591E-04

 1.16320946713525E-03
 8.37785475400743E-04

 7.70437439271402E-04
 1.53802408170584E-03

 1.17831608359156E-03
 1.27543460434143E-03

 1.13299623422265E-03
 1.70057947245524E-03

 2.76451081150327E-03
 1.32545164764894E-03

 1.91854028995035E-03
 3.42616746656423E-03

 3.0062166748041E-03
 4.55155094098314E-03

 2.38835606174135E-02
 .034399221534738

 .19550983017746
 .226289608183989

 5.37946612008914E-02
 5.77071637160362E-02

 1.5106616456302E-05
 1.25042608268768E-05

 1.5106616456302E-04
 7.50255649612605E-05

 5.36284884198721E-02
 5.76196338902481E-02

 7.61373469397621E-03
 5.97703667524709E-03

 3.0213232912604E-05
 0

 3.0213232912604E-05
 1.25042608268768E-05

 1.81279397475624E-04
 1.12538347441891E-04

 2.87025712669738E-04
 1.25042608268768E-04

 1.61640796082431E-03
 8.75298257881373E-05

 1.46534179626129E-03
 1.35046016930269E-03

 2.03939322160077E-03
 1.87563912403151E-03

 1.96386013931926E-03
 2.41332233958721E-03

 .169133677844757
 .195266537072507

 4.5319849368906E-05
 0

 1.28406239878567E-03
 5.62691737209454E-04

 1.58619472791171E-03
 6.87734345478222E-04

 1.76747412538733E-03
 1.63805816832086E-03

 2.31131231781421E-03
 4.83914894000131E-03

 3.56516148368727E-03
 4.81414041834755E-03

 2.06205314628522E-02
 3.29362230179934E-02

 3.98361475952684E-02
 5.09923756520034E-02

 9.81174738836815E-02
 9.87961647931533E-02

 6.90372372053001E-03
 3.03853538093105E-03

 6.0426465825208E-05
 0

 1.5106616456302E-05
 1.37546869095644E-04

 4.07878644320154E-04
 2.37580955710658E-04

 1.46534179626129E-03
 1.88814338485839E-03

 4.95497019766705E-03
 7.75264171266359E-04

 .684012486524898
 .695724568146596

 4.5319849368906E-05
 1.25042608268768E-05

 6.0426465825208E-05
 1.25042608268768E-05

 2.41705863300832E-04
 1.37546869095644E-04

 4.5319849368906E-04
 2.75093738191289E-04

 6.19371274708382E-04
 6.37717302170715E-04

 1.81279397475624E-04
 1.0753664311114E-03

 6.94904356989892E-04
 8.5028973622762E-04

 6.0426465825208E-04
 9.2531530118888E-04

 7.10010973446194E-04
 1.0753664311114E-03

 1.40491533043609E-03
 1.05035790945765E-03

 1.08767638485374E-03
 1.87563912403151E-03

 2.31131231781421E-03
 1.68807521162836E-03

 2.2659924684453E-03
 2.36330529627971E-03

 3.7464408811629E-03
 2.98851833762355E-03

 4.74347756727883E-03
 5.63942163292142E-03

 6.8130840217922E-03
 5.93952389276646E-03

 8.08203980412157E-03
 7.44003519199167E-03

 1.28708372207693E-02
 8.79049536129436E-03

 1.16018814384399E-02
 1.81936995031057E-02

 1.54993884841658E-02
 1.53427280345778E-02

 1.88681639539212E-02
 1.34295761280656E-02

 .013354248947371
 .023132882529722

 2.11190498059102E-02
 .024658402350601

 7.99140010538376E-02
 6.95111859366079E-02

 .068432972547048
 .085629178142452

 9.79059812532932E-02
 .111888125878893

 .167109391239613
 .133958146238331

 .143271150471568
 .157203567115495

 2.19045938616379E-03
 2.20074990553031E-03

 3.0213232912604E-05
 2.50085216537535E-05

 4.5319849368906E-04
 5.62691737209454E-04

 6.49584507620986E-04
 5.12674693901947E-04

 1.05746315194114E-03
 1.10037495276515E-03

 2.08622373261531E-02
 1.39422508219676E-02

 3.0213232912604E-05
 3.75127824806303E-05

 4.5319849368906E-05
 1.12538347441891E-04

 1.66172781019322E-03
 1.2379218218608E-03

 3.06664314062931E-03
 1.82562208072401E-03

 1.87322044058145E-03
 .003551210074833

 5.61966132174434E-03
 2.51335642620223E-03

 8.56545153072323E-03
 4.66408928842503E-03

 4.28121510371599E-02
 .046165730972829

 1.5106616456302E-05
 6.25213041343838E-05

 9.0639698737812E-05
 6.25213041343838E-05

 4.5319849368906E-05
 1.25042608268768E-04

 9.0639698737812E-05
 2.00068173230028E-04

 2.2659924684453E-04
 8.75298257881373E-05

 1.5106616456302E-04
 2.75093738191289E-04

 3.47452178494946E-04
 1.50051129922521E-04

 5.13624959514268E-04
 1.87563912403151E-04

 5.58944808883174E-04
 3.87632085633179E-04

 1.16320946713525E-03
 1.38797295178332E-03

 1.10278300131005E-03
 1.70057947245524E-03

 1.79768735829994E-03
 1.17540051772642E-03

 1.87322044058145E-03
 1.46299851674458E-03

 1.46534179626129E-03
 2.83846720770102E-03

 4.27517245713347E-03
 4.32647424609936E-03

 5.3326356090746E-03
 5.25178954728824E-03

 7.73458762562662E-03
 7.87768432093236E-03

 1.60281200601364E-02
 1.86063401103926E-02

 .113737715299498
 .127105811305202

 1.20852931650416E-04
 5.0017043307507E-05

 5.96711350023929E-03
 3.65124416144801E-03

 7.40224206358798E-03
 4.9516872874432E-03

 7.77990747499553E-03
 1.02159810955583E-02

 1.34448886461088E-02
 1.56678388160766E-02

 2.57718876744512E-02
 3.36989829284329E-02

 5.32508230084645E-02
 5.88700599729358E-02

 .024563358357947
 .019681706541504

 1.81279397475624E-04
 1.50051129922521E-04

 2.43820789604714E-02
 1.95316554115815E-02

 1.83998588437758E-02
 1.82062037639326E-02

 3.0213232912604E-04
 1.37546869095644E-04

 2.56812479757134E-04
 4.00136346460056E-04

 1.22363593296046E-03
 1.03785364863077E-03

 2.35663216718311E-03
 2.92599703348916E-03

 2.38684540009572E-03
 2.91349277266228E-03

 5.15135621159898E-03
 3.25110781498796E-03

 6.72244432305439E-03
 7.54006927860669E-03

 9.79513011026621E-02
 .122504243320912

 2.56812479757134E-04
 2.25076694883782E-04

 4.83411726601664E-04
 3.25110781498796E-04

 5.2873157597057E-04
 3.37615042325672E-04

 4.38091877232758E-04
 4.87666172248194E-04

 6.94904356989892E-04
 4.6265765059444E-04

 6.19371274708382E-04
 6.75230084651345E-04

 8.76183754465516E-04
 5.62691737209454E-04

 1.40491533043609E-03
 1.37546869095644E-04

 1.10278300131005E-03
 9.6282808366951E-04

 3.94282689509482E-03
 7.52756501777981E-03

 3.18296408734283E-02
 3.11731222414038E-02

 .055773627956667
 7.96271329455512E-02

 .026209979551684
 2.61714179106531E-02

 2.41705863300832E-04
 2.50085216537535E-04

 5.68008778756955E-03
 4.90167024413569E-03

 2.02881859008136E-02
 2.10196624499798E-02

 .24122245157423
 .238143647447868

 5.2873157597057E-04
 7.50255649612605E-05

 6.49584507620986E-04
 1.01284512697702E-03

 9.8193006965963E-04
 1.02534938780389E-03

 1.88832705703775E-03
 5.61441311126766E-03

 4.63470992879345E-02
 6.10958184001198E-02

 8.38266147160198E-02
 6.83232811580546E-02

 .107000164359987
 .100996914698684

 2.93068359252259E-03
 5.12674693901947E-03

 9.0639698737812E-05
 4.75161911421317E-04

 4.5319849368906E-04
 8.5028973622762E-04

 3.32345562038644E-04
 9.87836605323264E-04

 2.05449983805707E-03
 2.81345868604727E-03

 .445161773734307
 .466709031102348

 7.10010973446194E-04
 2.37580955710658E-04

 1.20852931650416E-03
 9.37819562015757E-04

 2.96089682543519E-03
 2.27577547049157E-03

 6.72244432305439E-03
 6.28964319591901E-03

 3.05002586252737E-02
 3.01852856360805E-02

 2.47295311389664E-02
 .036725014048537

 3.73737691128911E-02
 3.07604816341168E-02

 4.50781435056052E-02
 4.75411996637854E-02

 6.22392597999642E-02
 6.59849843834286E-02

 6.88408511913682E-02
 7.27747980124227E-02

 .164798078921798
 .17299644853984

 .121426983075755
 .119053067332694

 1.19342270004786E-03
 1.25042608268768E-04

 2.70408434567806E-03
 2.05069877560779E-03

 3.48811773976013E-02
 3.73877398723615E-02

 8.26482986324282E-02
 7.94895860764556E-02

 1.19342270004786E-03
 1.87563912403151E-04

 1.19342270004786E-03
 1.87563912403151E-04

 2.85212918694982E-02
 1.51051470788671E-02

 1.38980871397978E-03
 7.25247127958852E-04

 2.71314831555184E-02
 1.43798999509083E-02

 4.64377389866723E-02
 5.12674693901947E-02

 7.00947003572413E-03
 7.55257353943356E-03

 3.94282689509482E-02
 4.37148958507611E-02

 .98331987837361
 1.02414897876451

 7.553308228151E-05
 3.75127824806303E-04

 1.5106616456302E-05
 1.25042608268768E-05

 6.0426465825208E-05
 5.0017043307507E-05

 0
 3.12606520671919E-04

 5.67102381769577E-02
 5.83198724965532E-02

 1.5106616456302E-05
 1.25042608268768E-05

 4.5319849368906E-05
 8.75298257881373E-05

 9.21503603834422E-04
 3.50119303152549E-04

 2.37324944528504E-02
 1.72308714194362E-02

 3.19958136544476E-02
 4.06388476873495E-02

 4.51989964372556E-02
 5.74570784994987E-02

 3.0213232912604E-05
 0

 0
 1.50051129922521E-04

 4.22985260776456E-04
 1.87563912403151E-04

 3.62558794951248E-04
 3.75127824806303E-04

 2.56812479757134E-04
 5.87700258863208E-04

 4.5319849368906E-04
 4.37649128940687E-04

 7.40224206358798E-04
 4.37649128940687E-04

 5.43838192426872E-04
 6.25213041343838E-04

 8.76183754465516E-04
 1.0753664311114E-03

 4.5319849368906E-04
 2.07570729726154E-03

 1.76747412538733E-03
 1.32545164764894E-03

 3.61048133305618E-03
 2.41332233958721E-03

 3.56818280697853E-02
 4.77662763586692E-02

 3.45941516849316E-02
 3.59372456164438E-02

 3.0213232912604E-05
 0

 1.35959548106718E-04
 1.25042608268768E-05

 1.35959548106718E-04
 8.75298257881373E-05

 9.51716836747026E-04
 0

 8.15757288640308E-04
 6.00204519690084E-04

 1.19342270004786E-03
 8.00272692920113E-04

 2.41705863300832E-03
 1.62555390749398E-03

 2.76451081150327E-03
 2.97601407679667E-03

 9.66823453203328E-03
 5.27679806894199E-03

 1.64813185538255E-02
 .024558368263986

 .153437903346659
 .158028848330068

 6.0426465825208E-05
 2.50085216537535E-05

 3.47452178494946E-04
 2.00068173230028E-04

 5.58944808883174E-04
 3.00102259845042E-04

 7.25117589902496E-04
 3.00102259845042E-04

 6.94904356989892E-04
 3.50119303152549E-04

 8.45970521552912E-04
 5.37683215555701E-04

 1.17831608359156E-03
 4.6265765059444E-04

 1.10278300131005E-03
 9.75332344496387E-04

 1.75236750893103E-03
 5.37683215555701E-04

 2.9911100583478E-03
 8.62793997054496E-04

 3.17238945582342E-03
 1.22541756103392E-03

 4.13921290902675E-03
 1.75059651576275E-03

 5.18156944451159E-03
 1.32545164764894E-03

 4.74347756727883E-03
 2.00068173230028E-03

 6.96415018635522E-03
 5.5393875463064E-03

 8.61077138009214E-03
 4.36398702857999E-03

 .110368939829742
 .137271775357453

 .164616799524323
 .148162986537663

 7.553308228151E-05
 2.50085216537535E-05

 1.05746315194114E-04
 1.25042608268768E-05

 2.11492630388228E-04
 3.75127824806303E-05

 5.74051425339476E-04
 4.12640607286933E-04

 2.87025712669738E-04
 9.12811040362003E-04

 6.49584507620986E-04
 6.50221562997591E-04

 6.7979774053359E-04
 6.50221562997591E-04

 6.64691124077288E-04
 9.6282808366951E-04

 1.4955550291739E-03
 6.75230084651345E-04

 1.13299623422265E-03
 1.65056242914773E-03

 2.19045938616379E-03
 3.0135268592773E-03

 3.38388208621165E-03
 2.40081807876034E-03

 5.68008778756955E-03
 2.00068173230028E-03

 6.28435244582163E-03
 7.49005223529918E-03

 7.73458762562662E-03
 6.46470284749528E-03

 1.54389620183406E-02
 .00892804223039

 6.14839289771491E-02
 5.37683215555701E-02

 5.65440653959384E-02
 5.81073000624963E-02

 9.69391578000899E-02
 9.96964715726884E-02

 9.0639698737812E-05
 6.25213041343838E-05

 1.96386013931926E-04
 2.25076694883782E-04

 2.11492630388228E-04
 2.25076694883782E-04

 3.62558794951248E-04
 3.75127824806303E-04

 6.49584507620986E-04
 5.37683215555701E-04

 7.70437439271402E-04
 9.2531530118888E-04

 1.28406239878567E-03
 1.01284512697702E-03

 2.35663216718311E-03
 1.9881774714734E-03

 2.75695750327511E-02
 3.80004486528785E-02

 6.34477891164684E-02
 5.63441992859067E-02

 1.02876058067417E-02
 8.46538457979557E-03

 3.0213232912604E-04
 2.50085216537535E-04

 7.10010973446194E-04
 0

 6.19371274708382E-04
 6.12708780516961E-04

 9.0639698737812E-04
 4.00136346460056E-04

 1.26895578232937E-03
 1.21291330020705E-03

 1.38980871397978E-03
 1.52551982087896E-03

 2.80983066087217E-03
 1.92565616733902E-03

 2.2810990849016E-03
 2.53836494785598E-03

 3.0213232912604E-03
 2.07570729726154E-03

 5.43838192426872E-04
 2.25076694883782E-04

 1.07256976839744E-03
 6.00204519690084E-04

 1.40491533043609E-03
 1.25042608268768E-03

 4.38091877232758E-03
 4.67659354925191E-03

 4.38091877232758E-04
 4.50153389767563E-04

 4.38091877232758E-04
 4.6265765059444E-04

 6.0426465825208E-04
 7.25247127958852E-04

 7.70437439271402E-04
 7.62759910439482E-04

 9.66823453203328E-04
 1.08787069193828E-03

 1.16320946713525E-03
 1.18790477855329E-03

 .276904279644016
 .298001544026127

 7.70437439271402E-04
 7.12742867131975E-04

 1.29916901524197E-03
 9.87836605323264E-04

 1.58619472791171E-03
 1.00034086615014E-03

 1.13299623422265E-03
 1.82562208072401E-03

 5.2722091432494E-03
 1.66306668997461E-03

 2.85061852530419E-02
 1.67932222904955E-02

 3.19504938050787E-02
 3.42241618831617E-02

 2.85666117188671E-02
 3.72752015249196E-02

 .033566901765903
 3.48118621420249E-02

 3.52739494254652E-02
 4.68284567966535E-02

 5.15588819653587E-02
 5.24303656470942E-02

 5.74202491504039E-02
 6.94486646324735E-02

 2.77055345808579E-02
 3.65874671794414E-02

 1.11788961776635E-03
 4.50153389767563E-04

 2.65876449630915E-02
 3.61373137896738E-02

 .109447436225908
 .116364651254915

 2.16024615325119E-03
 1.62555390749398E-03

 3.61048133305618E-03
 6.51471989080279E-03

 1.16320946713525E-02
 1.31294738682206E-02

 9.20446140682481E-02
 9.50949035883977E-02

 .447941391162267
 .471285590564985

 3.12706960645451E-03
 2.96350981596979E-03

 3.0213232912604E-05
 0

 3.0213232912604E-05
 1.25042608268768E-05

 7.553308228151E-05
 3.75127824806303E-05

 7.553308228151E-05
 5.0017043307507E-05

 1.66172781019322E-04
 7.50255649612605E-05

 1.20852931650416E-04
 9.6282808366951E-04

 1.16320946713525E-03
 2.50085216537535E-04

 7.25117589902496E-04
 6.75230084651345E-04

 7.40224206358798E-04
 9.00306779535127E-04

 .194452367025519
 .203394306609977

 1.5106616456302E-05
 1.25042608268768E-05

 4.5319849368906E-05
 0

 1.5106616456302E-05
 2.50085216537535E-05

 1.5106616456302E-05
 2.50085216537535E-05

 6.0426465825208E-05
 0

 1.05746315194114E-04
 2.50085216537535E-05

 2.41705863300832E-04
 6.25213041343838E-05

 2.2659924684453E-04
 1.25042608268768E-04

 6.64691124077288E-04
 3.37615042325672E-04

 7.10010973446194E-04
 4.00136346460056E-04

 9.66823453203328E-04
 4.00136346460056E-04

 1.16320946713525E-03
 6.25213041343838E-04

 1.17831608359156E-03
 6.12708780516961E-04

 2.00917998868817E-03
 8.00272692920113E-04

 1.60130134436801E-03
 1.60054538584023E-03

 2.62855126339655E-03
 3.27611633664171E-03

 4.5319849368906E-03
 2.3132882529722E-03

 4.27517245713347E-03
 2.82596294687415E-03

 2.2962057013579E-03
 4.66408928842503E-03

 4.13921290902675E-03
 4.71410633173254E-03

 5.71030102048216E-03
 4.53904668015626E-03

 7.03968326863673E-03
 6.81482215064783E-03

 1.18133740688282E-02
 1.71683501153018E-02

 2.30526967123168E-02
 3.15107372837294E-02

 5.93387894403542E-02
 5.92326835369152E-02

 6.06077452226836E-02
 .061283382312523

 2.71919096213436E-04
 3.37615042325672E-04

 6.0426465825208E-05
 2.50085216537535E-05

 2.11492630388228E-04
 3.12606520671919E-04

 8.45970521552912E-03
 1.00284171831552E-02

 1.05746315194114E-04
 6.25213041343838E-05

 .008353958900335
 9.96589587902078E-03

 4.19963937485195E-03
 7.66511188687545E-03

 1.96386013931926E-04
 1.50051129922521E-04

 4.00325336092003E-03
 7.51506075695293E-03

 .237430690843698
 .246896630026682

 2.71919096213436E-04
 6.62725823824468E-04

 7.77990747499553E-03
 8.14027379829677E-03

 8.73162431174255E-03
 9.35318709850382E-03

 .220647239960747
 .228740443306057

 1.51242911975559
 1.47290189131947

 .101380503038243
 .110425127362149

 3.0213232912604E-05
 0

 8.3086390509661E-04
 2.37580955710658E-04

 .020106906503338
 2.52586068702911E-02

 2.71919096213436E-02
 2.71592545159763E-02

 5.32206097755519E-02
 5.77696850201706E-02

 .857270270662226
 .78546764810109

 4.5319849368906E-05
 2.50085216537535E-05

 9.36610220290724E-04
 5.25178954728824E-04

 1.13299623422265E-03
 1.38797295178332E-03

 3.56516148368727E-03
 3.16357798919982E-03

 4.19963937485195E-03
 3.1760822500267E-03

 4.69815771790992E-03
 4.43901259354125E-03

 1.19644402333912E-02
 7.87768432093236E-03

 .043567481859975
 4.76162252287467E-02

 6.39463074595264E-02
 6.28589191767095E-02

 7.64696925018007E-02
 6.70853593361938E-02

 8.29202177286417E-02
 7.56632822634313E-02

 .106622498948579
 9.50949035883977E-02

 .11336004988809
 9.43321436779583E-02

 .163876575317964
 .127518451912489

 .179965121843926
 .194703845335298

 3.71622764825029E-03
 3.56371433565988E-03

 7.553308228151E-05
 0

 2.41705863300832E-04
 1.25042608268768E-04

 3.62558794951248E-04
 1.87563912403151E-04

 3.7766541140755E-04
 3.37615042325672E-04

 2.65876449630915E-03
 2.91349277266228E-03

 .295485417885267
 .298664269849951

 9.0639698737812E-05
 3.75127824806303E-05

 3.7766541140755E-04
 3.37615042325672E-04

 1.99407337223186E-03
 1.50051129922521E-03

 2.2357792355327E-03
 1.41298147343707E-03

 2.2659924684453E-03
 1.97567321064653E-03

 6.59857006811271E-02
 5.93077091018765E-02

 9.30265441379077E-02
 8.60543230105659E-02

 .129509022879877
 .148037943929394

 .191279977569696
 .21230984457954

 1.05746315194114E-04
 2.25076694883782E-04

 6.34477891164684E-04
 3.37615042325672E-04

 9.21503603834422E-04
 6.37717302170715E-04

 4.01835997737633E-03
 3.28862059746859E-03

 7.79501409145183E-03
 3.18858651085357E-03

 2.59682736883831E-02
 3.56621518782525E-02

 5.24803855691931E-02
 .054481064422702

 9.93562164330982E-02
 .114489012130884

 1.24478519599928E-02
 9.07809336031253E-03

 2.41705863300832E-04
 3.25110781498796E-04

 1.85811382412515E-03
 1.71308373328212E-03

 1.82790059121254E-03
 1.91315190651214E-03

 2.87025712669738E-03
 1.32545164764894E-03

 5.64987455465695E-03
 3.80129529137053E-03

 5.86136718504517E-03
 3.06354390258481E-03

 3.32345562038644E-04
 3.87632085633179E-04

 3.47452178494946E-04
 3.75127824806303E-04

 4.83411726601664E-04
 4.50153389767563E-04

 6.0426465825208E-04
 4.6265765059444E-04

 1.35959548106718E-03
 8.75298257881373E-04

 2.73429757859066E-03
 5.12674693901947E-04

 4.49875038068673E-02
 5.03296498281789E-02

 1.75236750893103E-03
 1.95066468899277E-03

 6.85840387116111E-03
 5.83948980615145E-03

 3.63767324267752E-02
 4.25394953330347E-02

 .11074660524115
 .118365332987215

 .11074660524115
 .118365332987215

 4.5319849368906E-05
 2.50085216537535E-05

 4.5319849368906E-05
 3.75127824806303E-05

 0
 2.37580955710658E-04

 5.89158041795778E-04
 0

 1.66172781019322E-04
 5.87700258863208E-04

 4.68305110145362E-03
 2.58838199116349E-03

 5.24803855691931E-02
 5.65192589374829E-02

 5.27371980489503E-02
 5.83698895398607E-02

 12.4943652320618
 11.9813951604009

 9.97036686115932E-04
 3.8763208563318E-04

 9.97036686115932E-04
 3.8763208563318E-04

 3.0213232912604E-05
 0

 6.0426465825208E-05
 1.25042608268768E-05

 6.0426465825208E-05
 5.0017043307507E-05

 3.92772027863852E-04
 0

 4.5319849368906E-04
 3.25110781498796E-04

 .516993734984023
 .47695002071956

 5.89913372618593E-02
 3.73127143074002E-02

 1.5106616456302E-05
 1.25042608268768E-05

 4.07878644320154E-04
 1.75059651576275E-04

 7.5382016116947E-03
 6.27713893509213E-03

 5.10301503893881E-02
 .030848011459905

 .223124725059581
 .204057032433802

 1.05746315194114E-04
 1.00034086615014E-04

 2.17686343135312E-02
 1.51551641221746E-02

 3.96246549648801E-02
 3.69750992650746E-02

 5.07431246767184E-02
 4.79038232277649E-02

 .110882564789257
 .103922911732173

 6.25564987455466E-02
 5.66317972849248E-02

 1.66172781019322E-04
 1.87563912403151E-04

 3.32345562038644E-04
 1.37546869095644E-04

 2.11492630388228E-04
 2.87597999018165E-04

 5.2873157597057E-04
 2.87597999018165E-04

 6.49584507620986E-04
 4.6265765059444E-04

 3.92772027863852E-04
 7.75264171266359E-04

 3.7766541140755E-04
 7.87768432093236E-04

 9.97036686115932E-04
 3.50119303152549E-04

 6.7979774053359E-04
 6.50221562997591E-04

 6.0426465825208E-04
 7.50255649612605E-04

 4.22985260776456E-04
 9.12811040362003E-04

 1.13299623422265E-03
 4.2514486811381E-04

 1.26895578232937E-03
 3.75127824806303E-04

 6.0426465825208E-04
 1.08787069193828E-03

 7.85544055727704E-04
 1.18790477855329E-03

 1.35959548106718E-03
 1.2379218218608E-03

 8.91290370921818E-04
 1.65056242914773E-03

 1.35959548106718E-03
 1.4504942559177E-03

 1.61640796082431E-03
 1.43798999509083E-03

 1.35959548106718E-03
 1.7756050374165E-03

 4.68154043980799E-02
 4.04137709924657E-02

 .171686696025872
 .178673382955242

 2.87025712669738E-04
 2.62589477364412E-04

 3.71018500166777E-02
 4.49028006293144E-02

 4.10748901446851E-02
 4.50778602808907E-02

 .0472232830424
 4.28896146361873E-02

 4.59996471094396E-02
 4.55405179314852E-02

 6.34477891164684E-04
 2.75093738191289E-04

 6.34477891164684E-04
 2.75093738191289E-04

 .33125788565379
 .322659946376728

 .319882603462195
 .305954253912021

 3.0213232912604E-05
 0

 1.35959548106718E-04
 7.50255649612605E-05

 4.68305110145362E-04
 3.00102259845042E-04

 4.98518343057966E-03
 3.7137654655824E-03

 .011707627753634
 1.00659299656358E-02

 .02250885851989
 3.17233097177863E-02

 7.33275162788899E-02
 5.03796668714865E-02

 .079687401806993
 6.69102996846175E-02

 .127031537781043
 .142786154382106

 1.13752821915954E-02
 1.67056924647074E-02

 1.13752821915954E-02
 1.67056924647074E-02

 .182638992956691
 .188926876833281

 7.23909060585992E-02
 8.90928583914969E-02

 0
 2.50085216537535E-05

 3.24792253810493E-03
 1.87563912403151E-03

 3.01679130632351E-02
 3.34363934510684E-02

 3.89750704572592E-02
 5.37558172947432E-02

 .110248086898092
 .099834018441784

 .110248086898092
 .099834018441784

 6.89663871741201
 6.41609878566122

 5.2873157597057E-04
 3.37615042325672E-04

 3.0213232912604E-05
 0

 2.11492630388228E-04
 1.62555390749398E-04

 2.87025712669738E-04
 1.75059651576275E-04

 9.8344073130526E-03
 1.49425916881177E-02

 3.0213232912604E-05
 0

 1.5106616456302E-05
 1.25042608268768E-05

 4.22985260776456E-04
 2.12572434056905E-04

 9.36610220290724E-03
 1.47175149932339E-02

 8.76183754465516E-04
 3.00102259845042E-04

 3.0213232912604E-05
 0

 1.5106616456302E-05
 2.50085216537535E-05

 4.5319849368906E-05
 2.50085216537535E-05

 1.35959548106718E-04
 7.50255649612605E-05

 1.5106616456302E-04
 1.37546869095644E-04

 4.98518343057966E-04
 3.75127824806303E-05

 1.66172781019322E-04
 1.00034086615014E-04

 0
 2.50085216537535E-05

 3.0213232912604E-05
 2.50085216537535E-05

 1.35959548106718E-04
 5.0017043307507E-05

 4.22985260776456E-03
 2.65090329529787E-03

 0
 2.50085216537535E-05

 6.0426465825208E-05
 2.50085216537535E-05

 9.36610220290724E-04
 4.87666172248194E-04

 3.23281592164863E-03
 2.11322007974217E-03

 4.80390403310403E-03
 6.82732641147471E-03

 4.5319849368906E-05
 0

 4.5319849368906E-05
 3.75127824806303E-05

 6.0426465825208E-05
 1.25042608268768E-04

 5.58944808883174E-04
 6.25213041343838E-05

 1.5106616456302E-04
 4.50153389767563E-04

 2.41705863300832E-04
 4.50153389767563E-04

 1.25384916587307E-03
 5.50187476382577E-04

 6.19371274708382E-04
 1.27543460434143E-03

 5.13624959514268E-04
 1.37546869095644E-03

 1.31427563169827E-03
 2.50085216537535E-03

 2.71919096213436E-04
 4.50153389767563E-04

 1.5106616456302E-05
 2.50085216537535E-05

 1.20852931650416E-04
 8.75298257881373E-05

 1.35959548106718E-04
 3.37615042325672E-04

 4.38091877232758E-04
 3.75127824806303E-05

 3.0213232912604E-05
 1.25042608268768E-05

 1.20852931650416E-04
 2.50085216537535E-05

 2.87025712669738E-04
 0

 3.74493021951727E-02
 2.15323371438818E-02

 0
 5.0017043307507E-05

 3.74493021951727E-02
 2.14823201005743E-02

 1.40491533043609E-03
 8.8780251870825E-04

 3.0213232912604E-05
 2.50085216537535E-05

 5.89158041795778E-04
 1.87563912403151E-04

 7.85544055727704E-04
 6.75230084651345E-04

 1.96386013931926E-04
 1.62555390749398E-04

 1.5106616456302E-05
 3.75127824806303E-05

 1.81279397475624E-04
 1.25042608268768E-04

 6.0426465825208E-05
 7.50255649612605E-05

 0
 5.0017043307507E-05

 6.0426465825208E-05
 2.50085216537535E-05

 4.22985260776456E-04
 2.87597999018165E-04

 6.0426465825208E-05
 0

 7.553308228151E-05
 0

 6.0426465825208E-05
 6.25213041343838E-05

 2.2659924684453E-04
 2.25076694883782E-04

 1.19342270004786E-03
 3.75127824806303E-04

 3.0213232912604E-05
 3.75127824806303E-05

 4.5319849368906E-05
 3.75127824806303E-05

 1.20852931650416E-04
 2.50085216537535E-05

 1.05746315194114E-04
 8.75298257881373E-05

 1.81279397475624E-04
 3.75127824806303E-05

 1.81279397475624E-04
 1.12538347441891E-04

 5.2873157597057E-04
 3.75127824806303E-05

 1.66172781019322E-03
 1.12538347441891E-03

 4.5319849368906E-05
 2.50085216537535E-05

 6.0426465825208E-05
 8.75298257881373E-05

 3.62558794951248E-04
 1.25042608268768E-05

 2.11492630388228E-04
 2.87597999018165E-04

 9.8193006965963E-04
 7.12742867131975E-04

 1.45023517980499E-03
 5.0017043307507E-05

 3.0213232912604E-05
 5.0017043307507E-05

 2.11492630388228E-04
 0

 2.87025712669738E-04
 0

 4.5319849368906E-04
 0

 4.68305110145362E-04
 0

 1.81279397475624E-04
 5.0017043307507E-05

 9.0639698737812E-05
 0

 9.0639698737812E-05
 5.0017043307507E-05

 3.30230635734762E-02
 3.59372456164438E-02

 4.5319849368906E-05
 5.0017043307507E-05

 3.15728283936712E-03
 7.40252240951104E-03

 2.98204608847401E-02
 2.84847061636253E-02

 .122318273446677
 8.12151740705645E-02

 6.0426465825208E-05
 3.75127824806303E-05

 .122257846980852
 8.11776612880839E-02

 .070381726069911
 4.18517609875565E-02

 7.553308228151E-05
 5.0017043307507E-05

 1.05746315194114E-04
 1.00034086615014E-04

 3.7766541140755E-04
 2.50085216537535E-05

 6.34477891164684E-04
 1.50051129922521E-04

 1.07256976839744E-03
 1.00034086615014E-03

 1.29916901524197E-03
 2.40081807876034E-03

 1.66021714854759E-02
 6.85233493312846E-03

 5.02143931007478E-02
 3.12731563280188E-02

 .172668626095532
 .129344073993213

 9.0639698737812E-05
 6.25213041343838E-05

 2.11492630388228E-04
 1.50051129922521E-04

 1.06803778346055E-02
 3.70126120475552E-03

 2.35361084389185E-02
 1.72933927235706E-02

 6.30248038556919E-02
 4.34523063733967E-02

 7.51252036371898E-02
 6.46845412574335E-02

 1.16320946713525E-03
 5.25178954728824E-04

 9.0639698737812E-05
 7.50255649612605E-05

 5.43838192426872E-04
 1.62555390749398E-04

 5.2873157597057E-04
 2.87597999018165E-04

 1.20852931650416E-04
 5.0017043307507E-05

 1.20852931650416E-04
 5.0017043307507E-05

 8.71651769528625E-03
 6.75230084651345E-03

 1.05746315194114E-04
 7.50255649612605E-05

 2.2659924684453E-04
 2.00068173230028E-04

 4.5319849368906E-04
 1.62555390749398E-04

 1.82790059121254E-03
 1.56303260335959E-03

 6.10307304834601E-03
 4.75161911421317E-03

 .121230597061824
 .136383972838745

 1.5106616456302E-04
 7.50255649612605E-05

 1.94875352286296E-03
 1.47550277757146E-03

 4.33559892295867E-03
 3.07604816341168E-03

 1.17227343700903E-02
 1.19415690896673E-02

 1.92156161324161E-02
 8.79049536129436E-03

 8.38568279489324E-02
 .111025331881839

 1.75236750893103E-03
 1.27543460434143E-03

 1.05746315194114E-04
 1.25042608268768E-04

 1.66172781019322E-04
 1.25042608268768E-04

 2.56812479757134E-04
 6.25213041343838E-05

 1.81279397475624E-04
 2.12572434056905E-04

 3.32345562038644E-04
 3.62623563979426E-04

 7.10010973446194E-04
 3.87632085633179E-04

 .403089846903506
 .334576506944741

 1.66172781019322E-04
 7.50255649612605E-05

 4.24495922422086E-03
 1.7756050374165E-03

 1.45023517980499E-02
 1.45424553416577E-02

 2.22520460401328E-02
 2.39581637442959E-02

 3.50775634115332E-02
 .033073769887089

 4.00476402256566E-02
 3.78253890013022E-02

 4.57277280132261E-02
 5.02171114807371E-02

 .241071385409667
 .173108986887282

 2.56812479757134E-04
 0

 2.56812479757134E-04
 0

 1.35959548106718E-04
 1.12538347441891E-04

 1.35959548106718E-04
 1.12538347441891E-04

 5.83115395213257E-03
 5.72695145870955E-03

 0
 2.37580955710658E-04

 1.76747412538733E-03
 1.4504942559177E-03

 4.06367982674524E-03
 4.03887624708119E-03

 3.11196298999821E-03
 1.22541756103392E-03

 1.35959548106718E-04
 1.25042608268768E-04

 3.0213232912604E-04
 2.12572434056905E-04

 1.26895578232937E-03
 3.87632085633179E-04

 1.40491533043609E-03
 5.0017043307507E-04

 3.95793351155112E-03
 3.05103964175793E-03

 1.96386013931926E-04
 8.75298257881373E-05

 2.11492630388228E-04
 2.12572434056905E-04

 3.32345562038644E-04
 1.25042608268768E-04

 2.41705863300832E-04
 2.00068173230028E-04

 2.56812479757134E-04
 2.12572434056905E-04

 3.7766541140755E-04
 2.12572434056905E-04

 4.22985260776456E-04
 1.87563912403151E-04

 5.13624959514268E-04
 2.75093738191289E-04

 5.2873157597057E-04
 2.62589477364412E-04

 4.5319849368906E-04
 5.75195998036331E-04

 4.22985260776456E-04
 7.00238606305098E-04

 4.38091877232758E-04
 1.05035790945765E-03

 1.81279397475624E-04
 1.50051129922521E-04

 2.56812479757134E-04
 9.00306779535127E-04

 1.16774145207214E-02
 1.31544823898744E-02

 1.96386013931926E-04
 1.37546869095644E-04

 4.07878644320154E-04
 2.37580955710658E-04

 3.0213232912604E-04
 4.00136346460056E-04

 4.38091877232758E-04
 5.0017043307507E-04

 4.98518343057966E-04
 4.75161911421317E-04

 3.08174975708561E-03
 7.37751388785729E-04

 6.75265755596699E-03
 1.06661344853259E-02

 2.71919096213436E-04
 8.75298257881373E-05

 2.71919096213436E-04
 8.75298257881373E-05

 1.73726089247473E-03
 1.68807521162836E-03

 1.05746315194114E-04
 2.25076694883782E-04

 1.20852931650416E-04
 2.87597999018165E-04

 2.87025712669738E-04
 2.25076694883782E-04

 4.83411726601664E-04
 3.62623563979426E-04

 7.40224206358798E-04
 5.87700258863208E-04

 4.03346659383263E-03
 1.70057947245524E-03

 2.2659924684453E-04
 1.25042608268768E-04

 3.8068673469881E-03
 1.57553686418647E-03

 6.29945906227793E-03
 1.16289625689954E-02

 1.66172781019322E-04
 1.87563912403151E-04

 2.71919096213436E-04
 3.62623563979426E-04

 3.62558794951248E-04
 3.12606520671919E-04

 5.49880839009393E-03
 1.07661685719409E-02

 .117861821592068
 .125517770180189

 2.2659924684453E-04
 1.37546869095644E-04

 1.32938224815458E-03
 7.50255649612605E-04

 1.32938224815458E-03
 9.6282808366951E-04

 1.32485026321769E-02
 1.19790818721479E-02

 1.43512856334869E-02
 1.57178558593841E-02

 .023460575356637
 2.47209236547354E-02

 6.39160942266137E-02
 7.12492781915438E-02

 1.60130134436801E-03
 1.18790477855329E-03

 2.11492630388228E-04
 1.75059651576275E-04

 3.7766541140755E-04
 1.87563912403151E-04

 5.13624959514268E-04
 2.12572434056905E-04

 4.98518343057966E-04
 6.12708780516961E-04

 1.78258074184364E-03
 1.60054538584023E-03

 2.56812479757134E-04
 1.50051129922521E-04

 1.35959548106718E-04
 2.50085216537535E-04

 1.38980871397978E-03
 1.20040903938017E-03

 2.08169174767842E-02
 2.71842630376301E-02

 6.0426465825208E-05
 3.25110781498796E-04

 1.5106616456302E-04
 2.50085216537535E-04

 2.71919096213436E-04
 2.00068173230028E-04

 5.89158041795778E-04
 4.12640607286933E-04

 1.61640796082431E-03
 1.22541756103392E-03

 1.37470209752348E-03
 1.43798999509083E-03

 1.63151457728062E-03
 1.22541756103392E-03

 1.51217230727583E-02
 2.21075331419181E-02

 .511298540579997
 .45016589402839

 5.2873157597057E-04
 1.25042608268768E-05

 4.68305110145362E-04
 3.50119303152549E-04

 4.98518343057966E-04
 4.6265765059444E-04

 6.19371274708382E-03
 6.63976249907156E-03

 1.66172781019322E-02
 1.09162197018634E-02

 1.62698259234373E-02
 1.86813656753539E-02

 2.33699356578992E-02
 1.73559140277049E-02

 2.82040529239158E-02
 1.75309736792812E-02

 4.11655298434229E-02
 3.04478751134449E-02

 4.65434853018665E-02
 .029122423465796

 4.21474599130826E-02
 5.03046413065252E-02

 5.04258857311361E-02
 4.57530903655421E-02

 5.84021792200635E-02
 4.59906713212527E-02

 6.39312008430701E-02
 6.22587146570194E-02

 .116532439343914
 .114338961000961

 5.64987455465695E-03
 6.10207928351586E-03

 2.87025712669738E-04
 2.25076694883782E-04

 1.45023517980499E-03
 9.6282808366951E-04

 3.91261366218222E-03
 4.91417450496257E-03

 5.93538960568105E-02
 .055193807289834

 3.32345562038644E-04
 2.25076694883782E-04

 8.89779709276188E-03
 6.10207928351586E-03

 1.63000391563499E-02
 1.89439551527183E-02

 3.38237142456602E-02
 2.99226961587161E-02

 7.02457665218043E-02
 6.79356490724214E-02

 5.89158041795778E-04
 2.50085216537535E-05

 5.58944808883174E-04
 2.50085216537535E-04

 1.43512856334869E-03
 4.00136346460056E-04

 5.66498117111325E-03
 3.36364616242985E-03

 2.52582627149369E-02
 3.13981989362875E-02

 3.67392912217265E-02
 3.24985738890527E-02

 3.38388208621165E-03
 2.42582660041409E-03

 3.32345562038644E-04
 2.37580955710658E-04

 .003051536524173
 2.18824564470343E-03

 7.91586702310225E-03
 9.74081918413699E-03

 4.5319849368906E-05
 5.25178954728824E-04

 7.87054717373334E-03
 9.21564022940817E-03

 9.85706723773705E-02
 8.67420573560441E-02

 3.47452178494946E-04
 2.87597999018165E-04

 9.8344073130526E-03
 4.4140040718875E-03

 .088388812885823
 8.20404552851384E-02

 3.23281592164863E-03
 2.70092033860538E-03

 4.38091877232758E-04
 2.25076694883782E-04

 3.62558794951248E-04
 3.37615042325672E-04

 4.68305110145362E-04
 5.50187476382577E-04

 6.49584507620986E-04
 4.2514486811381E-04

 5.58944808883174E-04
 6.12708780516961E-04

 7.553308228151E-04
 5.50187476382577E-04

 5.93690026732668E-03
 4.51403815850251E-03

 3.0213232912604E-04
 4.00136346460056E-04

 7.10010973446194E-04
 3.50119303152549E-04

 6.64691124077288E-04
 5.25178954728824E-04

 8.00650672184006E-04
 5.75195998036331E-04

 9.97036686115932E-04
 8.5028973622762E-04

 1.02724991902854E-03
 8.5028973622762E-04

 1.43512856334869E-03
 9.6282808366951E-04

 1.57108811145541E-03
 8.37785475400743E-04

 4.98518343057966E-04
 2.50085216537535E-04

 1.07256976839744E-03
 5.87700258863208E-04

 9.92353635014478E-02
 9.52074419358396E-02

 4.07878644320154E-04
 3.25110781498796E-04

 9.88274848571277E-02
 9.48823311543408E-02

 5.91424034264223E-02
 .052755476428593

 3.32345562038644E-04
 4.12640607286933E-04

 8.61077138009214E-04
 5.25178954728824E-04

 1.20852931650416E-03
 7.12742867131975E-04

 4.22985260776456E-03
 7.21495849710789E-03

 5.25105988021057E-02
 4.38899555023374E-02

 .182729632655429
 .15197678608986

 8.45970521552912E-04
 0

 9.53227498392656E-03
 7.84017153845173E-03

 .172351387149949
 .144136614551408

 2.94579020897889E-03
 2.65090329529787E-03

 4.98518343057966E-04
 3.37615042325672E-04

 2.44727186592092E-03
 2.3132882529722E-03

 1.97896675577556E-03
 1.08787069193828E-03

 4.5319849368906E-04
 3.75127824806303E-04

 1.5257682620865E-03
 7.12742867131975E-04

 .622709836945225
 .578309558982223

 5.89158041795778E-04
 3.50119303152549E-04

 9.0639698737812E-04
 8.00272692920113E-04

 6.19371274708382E-04
 1.10037495276515E-03

 3.15728283936712E-03
 1.90064764568527E-03

 3.41409531912425E-03
 2.42582660041409E-03

 7.25117589902496E-03
 4.97669580909695E-03

 2.77357478137705E-02
 2.84346891203177E-02

 4.70571102613807E-02
 6.42719006501465E-02

 6.34175758835558E-02
 6.26588510034794E-02

 7.01249135901539E-02
 6.50471648214129E-02

 7.59560675422864E-02
 6.86233834178997E-02

 9.03828862580549E-02
 7.11492441049288E-02

 .115762001904642
 8.26281555440016E-02

 .116336053329982
 .123942233316002

 .178952978541353
 .144961895765982

 8.45970521552912E-04
 3.12606520671919E-04

 .178107008019801
 .14464928924531

 9.97036686115932E-03
 8.51540162310307E-03

 5.89158041795778E-04
 5.87700258863208E-04

 2.17535276970749E-03
 2.35080103545283E-03

 7.20585604965605E-03
 5.57690032878703E-03

 .108964024499306
 .104948261119977

 4.98518343057966E-04
 6.87734345478222E-04

 1.78862338842616E-02
 1.90439892393333E-02

 .0254093288795
 3.09730540681737E-02

 2.84759720201293E-02
 2.86722700760284E-02

 3.66939713723576E-02
 .025571213390963

 3.71622764825029E-03
 1.85063060237776E-03

 5.58944808883174E-04
 7.50255649612605E-04

 3.15728283936712E-03
 1.10037495276515E-03

 .067828707888796
 7.55632481768163E-02

 7.25117589902496E-04
 6.50221562997591E-04

 2.40195201655202E-03
 1.00034086615014E-03

 2.94579020897889E-03
 8.09025675498926E-03

 2.50769833174613E-02
 2.61839221714799E-02

 3.66788647559012E-02
 3.96385068211993E-02

 2.06960645451337E-03
 9.00306779535127E-04

 1.08767638485374E-03
 3.75127824806303E-04

 9.8193006965963E-04
 5.25178954728824E-04

 .409343986116415
 .4464021115195

 9.66823453203328E-04
 4.87666172248194E-04

 3.47452178494946E-03
 3.65124416144801E-03

 6.64691124077288E-03
 8.45288031896869E-03

 8.67119784591735E-03
 .009740819184137

 2.02730792843573E-02
 .016843239333803

 2.06809579286774E-02
 2.39956765267765E-02

 3.22375195177485E-02
 4.38524427198568E-02

 5.85079255352576E-02
 7.00488691521636E-02

 7.87205783537897E-02
 7.78140151256541E-02

 .179164471171742
 .191515258824444

 1.13299623422265E-03
 3.62623563979426E-04

 1.13299623422265E-03
 3.62623563979426E-04

 2.72825493200814E-02
 2.04194579302897E-02

 4.22985260776456E-04
 1.00034086615014E-03

 3.64069456596878E-03
 3.81379955219741E-03

 2.32188694933362E-02
 1.56053175119422E-02

 6.59252742153019E-02
 5.01921029590833E-02

 1.04235653548484E-03
 5.37683215555701E-04

 6.48829176798171E-02
 4.96544197435276E-02

 .24992386265306
 .224376456277477

 8.61077138009214E-04
 7.62759910439482E-04

 .249062785515051
 .223613696367037

 .132092254293905
 .153552322954047

 9.51716836747026E-04
 8.12776953746989E-04

 .032161986435467
 3.09355412856931E-02

 9.89785510216907E-02
 .121804004714607

 5.27976245147755E-02
 4.78162934019767E-02

 1.23874254941676E-03
 9.6282808366951E-04

 5.15588819653587E-02
 4.68534653183072E-02

 .573583120229331
 .475512030724469

 1.10278300131005E-03
 1.22541756103392E-03

 2.06960645451337E-03
 2.78845016439352E-03

 4.57730478625951E-03
 4.13891033369621E-03

 5.42327530781242E-03
 4.40149981106062E-03

 .011254429259945
 8.04023971168176E-03

 2.41705863300832E-02
 2.37205827885852E-02

 2.84155455543041E-02
 .02658405851794

 4.74800955221572E-02
 4.80163615752068E-02

 .449089494012946
 .356596510260871

 .064777171364623
 5.38683556421851E-02

 1.61640796082431E-03
 1.08787069193828E-03

 1.99558403387749E-02
 2.63214690405756E-02

 4.32049230650237E-02
 2.64590159096712E-02

 .228215654805354
 .291799430655996

 2.11492630388228E-03
 8.5028973622762E-04

 2.64365787985285E-02
 2.96601066813517E-02

 .022810990849016
 3.28611974530321E-02

 2.59833803048394E-02
 3.33738721469341E-02

 2.79472404441587E-02
 3.43367002306036E-02

 2.93219425416822E-02
 3.59247413556169E-02

 3.33856223684274E-02
 4.22769058556703E-02

 6.02149731948198E-02
 8.25156171965597E-02

 2.16024615325119E-03
 1.52551982087896E-03

 2.16024615325119E-03
 1.52551982087896E-03

 2.2357792355327E-03
 1.91315190651214E-03

 2.2357792355327E-03
 1.91315190651214E-03

 3.59537471659988E-03
 2.26327120966469E-03

 3.59537471659988E-03
 2.26327120966469E-03

 5.71936499035594E-02
 5.66818143282323E-02

 3.12706960645451E-03
 2.6884160777785E-03

 5.40665802971048E-02
 5.39933982504538E-02

 9.88576980900403E-02
 .114764105869075

 3.86729381281331E-03
 2.32579251379908E-03

 4.11806364598792E-02
 4.18267524659028E-02

 5.38097678173477E-02
 7.06115608893731E-02

 4.27517245713347E-03
 2.77594590356664E-03

 4.27517245713347E-03
 2.77594590356664E-03

 2.75242551833822E-02
 3.10980966764425E-02

 3.7917607305318E-03
 3.28862059746859E-03

 2.37324944528504E-02
 2.78094760789739E-02

 4.5470915533469E-03
 4.11390181204245E-03

 4.5470915533469E-03
 4.11390181204245E-03

 4.17395812687624E-02
 4.30396657661098E-02

 9.26035588771312E-03
 3.61373137896738E-03

 3.24792253810493E-02
 3.94259343871424E-02

 8.57149417730575E-02
 9.04308142999727E-02

 6.75265755596699E-03
 5.82698554532457E-03

 4.09842504459473E-02
 4.07638902956182E-02

 3.79780337711432E-02
 4.38399384590299E-02

 6.49584507620986E-03
 7.97771840754737E-03

 6.49584507620986E-03
 7.97771840754737E-03

 2.24635386705211E-02
 1.64180944656892E-02

 9.95526024470302E-03
 6.78981362899408E-03

 1.25082784258181E-02
 9.6282808366951E-03

 6.99889540420472E-02
 7.23121403618283E-02

 1.06199513687803E-02
 8.1777865807774E-03

 2.39137738503261E-02
 2.30453527039339E-02

 3.54552288229408E-02
 .041089001077117

 1.49102304423701E-02
 1.04160492687883E-02

 1.49102304423701E-02
 1.04160492687883E-02

 1.87473110222708E-02
 2.00068173230028E-02

 1.87473110222708E-02
 2.00068173230028E-02

 2.04543586818329E-02
 1.86313486320464E-02

 2.04543586818329E-02
 1.86313486320464E-02

 7.62279866384999E-02
 .085129007709377

 2.43669723440151E-02
 1.66806839430536E-02

 5.18610142944848E-02
 6.84483237663234E-02

 4.61507132740026E-02
 6.99988521088561E-02

 1.49102304423701E-02
 2.46458980897741E-02

 3.12404828316325E-02
 .045352954019082

 1.88530573374649E-02
 2.44583341773709E-02

 1.88530573374649E-02
 2.44583341773709E-02

 .206129781546241
 .17616002652904

 3.11800563658073E-02
 2.68966650386119E-02

 7.53215896511218E-02
 6.54973182111805E-02

 9.96281355293117E-02
 8.37660432792474E-02

 .185675422864408
 .161480024318286

 3.83556991825508E-02
 3.09855583290006E-02

 6.63180462431658E-02
 5.87200088430133E-02

 8.10016774386913E-02
 7.17744571462726E-02

 .344022976559365
 .32197221203125

 5.69217308073459E-02
 4.13015735111739E-02

 .138316180273901
 .139760123262002

 .148785065478118
 .140910515258074

 6.58195279001078E-02
 5.97828710132978E-02

 6.58195279001078E-02
 5.97828710132978E-02

 .062178833334139
 6.86608962003803E-02

 .062178833334139
 6.86608962003803E-02

 .109281263444889
 9.48073055893796E-02

 .109281263444889
 9.48073055893796E-02

 .209468343783083
 .176872769396172

 .209468343783083
 .176872769396172

 .117317983399641
 .126880734610318

 .117317983399641
 .126880734610318

 3.0213232912604E-05
 0

 .117287770166729
 .126880734610318

 6.11666900315668E-02
 5.84074023223413E-02

 6.11666900315668E-02
 5.84074023223413E-02

 3.0213232912604E-05
 0

 3.23281592164863E-03
 3.1760822500267E-03

 8.39927874970391E-03
 6.33966023922652E-03

 1.36110614271281E-02
 1.37671911703913E-02

 3.58933207001735E-02
 3.51244686626968E-02

 6.40520537747205E-03
 5.02671285240446E-03

 6.40520537747205E-03
 5.02671285240446E-03

 4.5319849368906E-05
 0

 6.35988552810314E-03
 5.02671285240446E-03

 8.83737062693667E-03
 4.76412337504005E-03

 8.83737062693667E-03
 4.76412337504005E-03

 3.0213232912604E-05
 1.25042608268768E-05

 2.11492630388228E-04
 0

 3.0213232912604E-04
 5.0017043307507E-05

 3.47452178494946E-04
 2.50085216537535E-04

 6.19371274708382E-04
 3.25110781498796E-04

 5.89158041795778E-04
 5.75195998036331E-04

 1.91854028995035E-03
 7.12742867131975E-04

 4.81901064956034E-03
 2.83846720770102E-03

 6.32816163354491E-02
 6.77230766383645E-02

 6.32816163354491E-02
 6.77230766383645E-02

 0
 3.75127824806303E-05

 0
 5.0017043307507E-05

 6.0426465825208E-05
 1.25042608268768E-05

 4.5319849368906E-05
 2.50085216537535E-05

 1.5106616456302E-05
 7.50255649612605E-05

 6.0426465825208E-05
 6.25213041343838E-05

 0
 1.25042608268768E-04

 1.20852931650416E-04
 8.75298257881373E-05

 1.96386013931926E-04
 5.0017043307507E-05

 2.2659924684453E-04
 1.37546869095644E-04

 2.11492630388228E-04
 1.87563912403151E-04

 2.2659924684453E-04
 1.87563912403151E-04

 9.0639698737812E-05
 3.87632085633179E-04

 1.07256976839744E-03
 1.41298147343707E-03

 2.70408434567806E-03
 5.03921711323133E-03

 4.48666508752169E-03
 7.91519710341299E-03

 5.37644479679788E-02
 5.19301952140192E-02

 .142379860100646
 .117352487860238

 9.62291468266437E-02
 7.54757183510281E-02

 3.0213232912604E-05
 2.50085216537535E-05

 1.5106616456302E-04
 0

 9.0639698737812E-05
 6.25213041343838E-05

 1.35959548106718E-04
 7.50255649612605E-05

 1.66172781019322E-04
 1.25042608268768E-04

 3.47452178494946E-04
 2.50085216537535E-05

 1.81279397475624E-03
 9.2531530118888E-04

 1.42002194689239E-03
 1.32545164764894E-03

 9.20748273011607E-02
 7.29123448815184E-02

 4.00778534585692E-02
 3.77628676971678E-02

 1.5106616456302E-04
 3.12606520671919E-04

 3.41409531912425E-03
 6.25213041343838E-04

 4.12410629257045E-03
 2.56337346950974E-03

 6.26924582936533E-03
 6.35216450005339E-03

 2.61193398529462E-02
 2.79095101655889E-02

 6.0728598154334E-03
 4.11390181204245E-03

 8.3086390509661E-04
 5.25178954728824E-04

 1.22363593296046E-03
 1.18790477855329E-03

 2.05449983805707E-03
 7.25247127958852E-04

 1.96386013931926E-03
 1.67557095080149E-03

 1.98198807906682E-02
 1.28168673475487E-02

 1.24931718093618E-02
 1.27543460434143E-02

 4.5319849368906E-05
 1.25042608268768E-05

 1.66172781019322E-04
 7.50255649612605E-05

 3.47452178494946E-04
 2.50085216537535E-05

 1.19342270004786E-02
 1.26418076959724E-02

 7.32670898130647E-03
 6.25213041343838E-05

 1.64662119373692E-03
 6.25213041343838E-05

 5.68008778756955E-03
 0

 1.08012307662559E-02
 8.64044423137184E-03

 1.08012307662559E-02
 8.64044423137184E-03

 6.0426465825208E-05
 0

 9.0639698737812E-05
 3.87632085633179E-04

 1.06501646016929E-02
 8.25281214573866E-03

 .102105620628145
 .090405805778319

 .102105620628145
 .090405805778319

 4.5319849368906E-05
 3.75127824806303E-05

 1.05746315194114E-04
 1.00034086615014E-04

 2.2659924684453E-04
 1.12538347441891E-04

 2.11492630388228E-04
 2.25076694883782E-04

 3.92772027863852E-04
 2.00068173230028E-04

 2.87025712669738E-04
 3.12606520671919E-04

 8.15757288640308E-04
 5.12674693901947E-04

 .100020907557176
 8.89052944790937E-02

 .01960838816028
 1.58679069893066E-02

 .01960838816028
 1.58679069893066E-02

 6.0426465825208E-05
 2.50085216537535E-05

 8.00650672184006E-04
 5.87700258863208E-04

 7.65905454334511E-03
 7.28998406206915E-03

 1.10882564789257E-02
 7.9652141467205E-03

 .266148368727129
 .296413502901114

 .266148368727129
 .296413502901114

 9.0639698737812E-05
 0

 4.5319849368906E-05
 1.12538347441891E-04

 6.25413921290903E-03
 5.72695145870956E-03

 1.95932815438237E-02
 2.58838199116349E-02

 5.26465583502125E-02
 5.76821551943825E-02

 6.38405611443322E-02
 8.51665204918576E-02

 .123677868927744
 .121841517497087

 1.25384916587307E-03
 9.2531530118888E-04

 1.25384916587307E-03
 9.2531530118888E-04

 9.0639698737812E-05
 0

 4.5319849368906E-05
 8.75298257881373E-05

 1.11788961776635E-03
 8.37785475400743E-04

 7.84184460246637E-02
 9.62327913236435E-02

 7.84184460246637E-02
 9.62327913236435E-02

 4.5319849368906E-05
 5.0017043307507E-05

 6.0426465825208E-05
 7.50255649612605E-05

 3.32345562038644E-04
 1.50051129922521E-04

 5.13624959514268E-04
 2.12572434056905E-04

 5.13624959514268E-04
 3.75127824806303E-04

 6.49584507620986E-04
 7.25247127958852E-04

 1.66172781019322E-03
 1.81311781989713E-03

 4.72837095082252E-03
 7.11492441049288E-03

 6.99134209597656E-02
 8.57167079682402E-02

 2.41705863300832E-04
 1.62555390749398E-04

 2.41705863300832E-04
 1.62555390749398E-04

 6.0426465825208E-05
 3.75127824806303E-05

 1.81279397475624E-04
 1.25042608268768E-04

 1.24327453435365E-02
 1.37921996920451E-02

 1.24327453435365E-02
 1.37921996920451E-02

 9.0639698737812E-05
 2.50085216537535E-05

 1.23421056447987E-02
 1.37671911703913E-02

 6.0426465825208E-05
 5.0017043307507E-05

 6.0426465825208E-05
 5.0017043307507E-05

 6.0426465825208E-05
 5.0017043307507E-05

 3.79629271546869E-02
 4.80038573143799E-02

 3.79629271546869E-02
 4.80038573143799E-02

 6.0426465825208E-05
 6.25213041343838E-05

 9.66823453203328E-04
 3.87632085633179E-04

 3.69356772356584E-02
 4.75537039246123E-02

 7.78443945993242E-02
 4.77287635761886E-02

 7.78443945993242E-02
 4.77287635761886E-02

 1.05746315194114E-04
 3.75127824806303E-05

 1.05746315194114E-04
 6.25213041343838E-05

 6.64691124077288E-04
 7.12742867131975E-04

 1.16320946713525E-03
 4.75161911421317E-04

 1.5257682620865E-03
 1.26293034351455E-03

 2.64365787985285E-03
 1.47550277757146E-03

 7.34181559776277E-03
 5.23928528646136E-03

 7.77990747499553E-03
 5.10173841736572E-03

 5.65138521630258E-02
 3.33613678861072E-02

 4.22985260776456E-03
 1.66306668997461E-03

 4.22985260776456E-03
 1.66306668997461E-03

 1.5106616456302E-04
 5.0017043307507E-05

 4.07878644320154E-03
 1.6130496466671E-03

 4.57730478625951E-03
 4.56405520181002E-03

 4.57730478625951E-03
 4.56405520181002E-03

 2.11492630388228E-04
 2.50085216537535E-05

 2.56812479757134E-04
 3.00102259845042E-04

 4.10899967611414E-03
 4.23894442031122E-03

 .183016658368099
 .190589943523256

 .183016658368099
 .190589943523256

 1.66172781019322E-04
 1.25042608268768E-04

 .027010630223868
 2.74593567758214E-02

 6.90523438217564E-02
 6.56098565586224E-02

 .086787511541455
 9.73956875805431E-02

 2.11492630388228E-04
 1.25042608268768E-04

 2.11492630388228E-04
 1.25042608268768E-04

 2.11492630388228E-04
 1.25042608268768E-04

 2.85515051024108E-03
 1.33795590847581E-03

 2.85515051024108E-03
 1.33795590847581E-03

 2.87025712669738E-04
 7.50255649612605E-05

 4.98518343057966E-04
 1.12538347441891E-04

 6.49584507620986E-04
 3.62623563979426E-04

 1.42002194689239E-03
 7.87768432093236E-04

 3.64824787419693E-02
 2.44083171340634E-02

 3.64824787419693E-02
 2.44083171340634E-02

 1.96386013931926E-04
 1.62555390749398E-04

 1.5106616456302E-03
 1.20040903938017E-03

 3.47754310824072E-02
 2.30453527039339E-02

 6.63935793254473E-02
 6.41593623027047E-02

 6.63935793254473E-02
 6.41593623027047E-02

 1.96386013931926E-04
 1.87563912403151E-04

 5.2873157597057E-04
 4.12640607286933E-04

 2.40950532478017E-02
 2.60963923456918E-02

 4.15734084877431E-02
 3.74627654373228E-02

 .258398674485046
 .319496368387528

 5.74051425339476E-04
 6.75230084651345E-04

 2.41705863300832E-04
 1.62555390749398E-04

 3.32345562038644E-04
 5.12674693901947E-04

 .257824623059706
 .318821138302877

 4.39753605042951E-02
 3.66124757010951E-02

 5.41572199958427E-02
 4.90667194846644E-02

 .159692042559568
 .233141943117117

 8.95822355858708E-03
 5.11424267819259E-03

 8.95822355858708E-03
 5.11424267819259E-03

 3.32345562038644E-04
 1.75059651576275E-04

 1.22363593296046E-03
 3.75127824806303E-05

 7.10010973446194E-04
 5.62691737209454E-04

 6.69223109014178E-03
 4.33897850692623E-03

 .243473337426219
 .202831614872768

 .239107525270348
 .202081359223155

 2.2659924684453E-04
 3.00102259845042E-04

 4.68305110145362E-04
 1.12538347441891E-04

 5.13624959514268E-04
 3.75127824806303E-04

 6.49584507620986E-04
 3.87632085633179E-04

 4.98518343057966E-04
 8.00272692920113E-04

 6.57137815849137E-03
 4.26395294196497E-03

 7.49288176232579E-03
 5.76446424119019E-03

 1.22816791789735E-02
 8.94054649121688E-03

 2.47144245225101E-02
 2.57837858250199E-02

 6.63180462431658E-02
 6.00329562298353E-02

 .119372483237698
 9.53199802832815E-02

 4.36581215587128E-03
 7.50255649612605E-04

 4.36581215587128E-03
 7.50255649612605E-04

 3.17238945582342E-04
 2.50085216537535E-04

 3.17238945582342E-04
 2.50085216537535E-04

 3.17238945582342E-04
 2.50085216537535E-04

 1.40491533043609E-02
 5.0017043307507E-05

 1.40491533043609E-02
 5.0017043307507E-05

 6.34477891164684E-04
 0

 6.64691124077288E-04
 0

 1.22363593296046E-03
 0

 1.29916901524197E-03
 0

 1.4955550291739E-03
 0

 4.18453275839565E-03
 0

 4.5470915533469E-03
 5.0017043307507E-05

 6.10458370999164E-02
 5.84074023223413E-02

 6.10458370999164E-02
 5.84074023223413E-02

 4.07878644320154E-04
 2.00068173230028E-04

 5.89158041795778E-04
 4.2514486811381E-04

 7.553308228151E-04
 4.75161911421317E-04

 5.92934695909853E-02
 5.73070273695762E-02

 9.65312791557698E-03
 8.82800814377499E-03

 9.65312791557698E-03
 8.82800814377499E-03

 2.87025712669738E-04
 3.50119303152549E-04

 9.97036686115932E-04
 7.75264171266359E-04

 8.36906551679131E-03
 7.70262466935608E-03

 1.57108811145541E-03
 5.12674693901947E-04

 1.57108811145541E-03
 5.12674693901947E-04

 4.98518343057966E-04
 2.12572434056905E-04

 1.07256976839744E-03
 3.00102259845042E-04

 .695372662100037
 .820992253110247

 .543898618892697
 .65010902465015

 9.51716836747026E-04
 2.50085216537535E-05

 1.84300720766884E-03
 1.36296443012957E-03

 5.95200688378299E-03
 7.32749684454978E-03

 .056226826450356
 7.37126175744385E-02

 .478925061514142
 .567680937279378

 .15147404320734
 .170883228460098

 1.73726089247473E-03
 1.63805816832086E-03

 7.74969424208292E-03
 5.26429380811512E-03

 5.98222011669559E-03
 1.15914497865148E-02

 1.16774145207214E-02
 1.31169696073937E-02

 1.45023517980499E-02
 1.31794909115281E-02

 4.55615552322068E-02
 .053668287468955

 6.42635464051087E-02
 7.24246787092702E-02

 .070079593740785
 6.78231107249795E-02

 .070079593740785
 6.78231107249795E-02

 8.00650672184006E-04
 2.75093738191289E-04

 7.70437439271402E-04
 7.50255649612605E-04

 6.94904356989892E-04
 1.05035790945765E-03

 1.26895578232937E-03
 9.00306779535127E-04

 2.2659924684453E-03
 1.43798999509083E-03

 4.5621981698032E-03
 3.7637825088899E-03

 5.97164548517618E-02
 5.96453241442021E-02

 2.00313734210564E-02
 2.56337346950974E-02

 2.00313734210564E-02
 2.56337346950974E-02

 5.74051425339476E-04
 4.6265765059444E-04

 .019457321995717
 2.51710770445029E-02

 5.89158041795778E-04
 5.50187476382577E-04

 5.89158041795778E-04
 5.50187476382577E-04

 5.89158041795778E-04
 5.50187476382577E-04

 3.73737691128911E-02
 2.96476024205248E-02

 3.73737691128911E-02
 2.96476024205248E-02

 7.85544055727704E-04
 4.2514486811381E-04

 3.65882250571634E-02
 .029222457552411

 .200691399621972
 .168607452989606

 .200691399621972
 .168607452989606

 8.15757288640308E-04
 5.87700258863208E-04

 1.31427563169827E-03
 6.25213041343838E-04

 8.17267950285938E-03
 7.94020562506674E-03

 2.22671526565891E-02
 1.75934949834156E-02

 2.98053542682838E-02
 1.36546528229494E-02

 3.04398321594485E-02
 2.56962559992317E-02

 .041769794501675
 4.73286272297285E-02

 6.61065536127775E-02
 5.51813030290071E-02

 .555334327550118
 .46465833232674

 .112151520571586
 8.52165375351651E-02

 1.17831608359156E-03
 6.12708780516961E-04

 4.38242943397321E-02
 .022932814356492

 6.71489101482624E-02
 6.16710143981562E-02

 .443182806978532
 .379441794791575

 3.21770930519233E-03
 2.67591181695163E-03

 4.33559892295867E-03
 4.11390181204245E-03

 4.80088270981277E-02
 5.31556127750531E-02

 5.36587016527847E-02
 4.94168387878169E-02

 .333961969999468
 .270079529599711

 6.75416821761262E-02
 7.81891429504604E-02

 6.75416821761262E-02
 7.81891429504604E-02

 1.67683442664952E-03
 9.37819562015757E-04

 3.41409531912425E-03
 1.57553686418647E-03

 6.24507524303525E-02
 7.56757865242581E-02

 .184572639863098
 .175047147315448

 .184572639863098
 .175047147315448

 1.46534179626129E-03
 1.16289625689954E-03

 1.66777045677574E-02
 2.58463071291543E-02

 4.91116100994378E-02
 3.24485568457452E-02

 .117317983399641
 .115589387083649

 .224786452869774
 .306804543648248

 .224786452869774
 .306804543648248

 1.05746315194114E-03
 1.90064764568527E-03

 6.0577531989771E-03
 5.56439606796016E-03

 3.92016697041037E-02
 .053068082949265

 .178469566814752
 .246271416985338

 4.42623862169648E-03
 2.83846720770102E-03

 4.42623862169648E-03
 2.83846720770102E-03

 4.42623862169648E-03
 2.83846720770102E-03

 .130989471292595
 .124017258880964

 .130989471292595
 .124017258880964

 4.16942614193935E-03
 3.85131233467804E-03

 5.24199591033679E-03
 6.0270537185546E-03

 5.85381387681702E-02
 5.13549992159828E-02

 6.30399104721482E-02
 6.27838936117482E-02

 .222293861154484
 .186901186579327

 .222293861154484
 .186901186579327

 7.17564281674345E-03
 5.13925119984635E-03

 7.87054717373334E-03
 7.05240310635849E-03

 1.21306130144105E-02
 9.80334048827138E-03

 .195117058149597
 .164906191784851

 2.91859829935755E-02
 3.66749970052295E-02

 2.91859829935755E-02
 3.66749970052295E-02

 2.91859829935755E-02
 3.66749970052295E-02

 7.74969424208292E-02
 8.51790247526845E-02

 7.74969424208292E-02
 8.51790247526845E-02

 3.71471698660466E-02
 3.72877057857465E-02

 4.03497725547826E-02
 .047891318966938

 .042721511338422
 3.33988806685878E-02

 .042721511338422
 3.33988806685878E-02

 .042721511338422
 3.33988806685878E-02

 8.10318906716039E-02
 7.58758546974882E-02

 8.10318906716039E-02
 7.58758546974882E-02

 8.10318906716039E-02
 7.58758546974882E-02

 4.56663951504136
 4.59265244632108

 1.37504954970198
 1.32027488366661

 .183409430395963
 .190414883871679

 1.5106616456302E-05
 1.25042608268768E-05

 4.22985260776456E-04
 2.50085216537535E-04

 5.74051425339476E-04
 3.00102259845042E-04

 1.19493336169349E-02
 1.72308714194362E-02

 2.48050642212479E-02
 2.73343141675526E-02

 3.54854420558534E-02
 3.35739403201641E-02

 5.36587016527847E-02
 5.30055616451306E-02

 5.64987455465695E-02
 5.87075045821864E-02

 .057828127794724
 5.75696168469406E-02

 9.0639698737812E-05
 0

 6.64691124077288E-04
 8.75298257881373E-04

 6.7979774053359E-04
 2.48834790454847E-03

 4.5168783204343E-03
 1.82562208072401E-03

 5.18761209109411E-02
 5.23803486037867E-02

 6.19371274708382E-04
 1.50051129922521E-04

 0
 7.50255649612605E-05

 6.19371274708382E-04
 7.50255649612605E-05

 2.04996785312018E-02
 7.75264171266359E-03

 1.05746315194114E-04
 0

 6.0426465825208E-04
 0

 6.49584507620986E-04
 0

 9.97036686115932E-04
 0

 1.60130134436801E-03
 0

 2.68897772922176E-03
 0

 4.27517245713347E-03
 3.12606520671919E-04

 5.40816869135612E-03
 0

 4.16942614193935E-03
 7.44003519199167E-03

 .610549010697901
 .550287510469192

 7.553308228151E-05
 2.50085216537535E-05

 3.17238945582342E-04
 1.25042608268768E-04

 8.91290370921818E-04
 4.87666172248194E-04

 1.23874254941676E-03
 6.62725823824468E-04

 1.99407337223186E-03
 2.01318599312716E-03

 3.18749607227972E-03
 1.6130496466671E-03

 2.84004389378478E-03
 2.37580955710658E-03

 4.00325336092003E-03
 3.66374842227489E-03

 6.8432972547048E-03
 5.35182363390325E-03

 1.03329256561106E-02
 9.54075101090697E-03

 3.17390011746905E-02
 3.07979944165975E-02

 4.85979851399235E-02
 4.39399725456449E-02

 6.17860613062752E-02
 3.82630381302429E-02

 .066680605038117
 5.99204178823934E-02

 9.34344227822279E-02
 8.36284964101518E-02

 9.60025475797992E-02
 9.02057376050889E-02

 .180584493118634
 .177673042089092

 .117756075276874
 8.72922448324266E-02

 0
 2.00068173230028E-04

 3.47452178494946E-04
 3.00102259845042E-04

 7.85544055727704E-04
 3.87632085633179E-04

 1.13299623422265E-03
 7.25247127958852E-04

 1.10278300131005E-03
 9.75332344496387E-04

 2.09981968742598E-03
 9.75332344496387E-04

 2.55301818111504E-03
 1.81311781989713E-03

 2.43216524946462E-03
 1.9381604281659E-03

 3.38388208621165E-03
 2.87597999018165E-03

 5.58944808883174E-03
 5.31431085142262E-03

 7.26628251548126E-03
 5.98954093607397E-03

 8.12735965349047E-03
 5.27679806894199E-03

 1.11637895612072E-02
 7.84017153845173E-03

 1.04839918206736E-02
 9.35318709850382E-03

 1.03329256561106E-02
 .009740819184137

 1.63000391563499E-02
 8.5028973622762E-03

 1.40491533043609E-02
 1.25292693485305E-02

 2.06054248463959E-02
 1.25542778701843E-02

 8.03823061639829E-02
 .1194281951575

 9.0639698737812E-05
 1.37546869095644E-04

 3.92772027863852E-04
 1.62555390749398E-04

 5.89158041795778E-04
 5.62691737209454E-04

 8.11225303703417E-03
 9.61577657586823E-03

 7.11974833585513E-02
 .108949624584577

 1.66172781019322E-04
 1.37546869095644E-04

 1.66172781019322E-04
 1.37546869095644E-04

 2.83400124720225E-02
 3.07854901557706E-02

 3.0213232912604E-04
 2.25076694883782E-04

 3.65580118242508E-03
 3.43867172739111E-03

 5.54412823946283E-03
 4.38899555023374E-03

 1.88379507210086E-02
 2.27327461832619E-02

 .190539753363337
 .192190488909096

 3.7766541140755E-04
 3.62623563979426E-04

 1.25384916587307E-03
 8.37785475400743E-04

 2.46237848237723E-03
 2.92599703348916E-03

 .017206436143728
 .014817549079849

 1.56504546487289E-02
 1.64055902048623E-02

 1.92307227488724E-02
 1.45674638633114E-02

 2.94881153227015E-02
 3.64749288319995E-02

 4.79030807829336E-02
 5.24428699079211E-02

 5.69670506567148E-02
 5.33556809482831E-02

 7.70890637765091E-02
 8.42662137123225E-02

 5.89158041795778E-04
 1.87563912403151E-04

 9.8193006965963E-04
 4.50153389767563E-04

 5.61966132174434E-03
 0

 2.88536374315368E-03
 2.46333938289472E-03

 1.42908591676617E-02
 1.78435801999531E-02

 1.99407337223186E-02
 2.34079762679133E-02

 3.27813577101753E-02
 3.99136005593906E-02

 7.87054717373334E-03
 0

 2.73429757859066E-03
 0

 5.13624959514268E-03
 0

 .598569463848054
 .586975011735249

 .272916132899552
 .263289715970717

 1.5106616456302E-05
 1.25042608268768E-05

 1.96386013931926E-04
 0

 2.71919096213436E-04
 3.75127824806303E-05

 2.41705863300832E-04
 2.75093738191289E-04

 5.13624959514268E-04
 5.25178954728824E-04

 1.32938224815458E-03
 0

 8.91290370921818E-04
 7.50255649612605E-04

 1.28406239878567E-03
 5.75195998036331E-04

 1.19342270004786E-03
 6.50221562997591E-04

 1.11788961776635E-03
 8.75298257881373E-04

 1.31427563169827E-03
 8.62793997054496E-04

 2.35663216718311E-03
 2.37580955710658E-04

 1.31427563169827E-03
 1.10037495276515E-03

 6.52605830912246E-03
 6.25213041343838E-04

 1.04688852042173E-02
 7.74013745183671E-03

 1.08012307662559E-02
 1.31794909115281E-02

 3.62407728786685E-02
 3.26111122364946E-02

 3.90506035395407E-02
 4.63908076677128E-02

 5.20120804590478E-02
 4.86790873990312E-02

 .105776528427027
 .108161856152484

 .185720742713777
 .194878904986874

 1.05746315194114E-04
 7.50255649612605E-05

 1.02724991902854E-03
 6.75230084651345E-04

 1.5408748785428E-03
 1.22541756103392E-03

 4.5470915533469E-03
 4.11390181204245E-03

 6.31456567873423E-03
 .003551210074833

 1.05595249029551E-02
 9.67829788000261E-03

 2.99866336657595E-02
 4.19142822916909E-02

 5.36587016527847E-02
 .052455374168748

 7.79803541474309E-02
 8.11901655489108E-02

 .139932588234725
 .128806390777657

 .139932588234725
 .128806390777657

 .357150626259892
 .360722916333741

 .15617220092525
 .141923360385051

 3.0213232912604E-05
 0

 8.00650672184006E-04
 0

 9.97036686115932E-04
 0

 6.64691124077288E-04
 4.37649128940687E-04

 1.61640796082431E-03
 0

 1.84300720766884E-03
 0

 6.85840387116111E-03
 1.00034086615014E-03

 5.43838192426872E-03
 3.46368024904486E-03

 7.11521635091824E-03
 9.14061466444691E-03

 3.99116806775499E-02
 4.16892055968071E-02

 4.43832391486153E-02
 3.96385068211993E-02

 4.65132720689539E-02
 4.65533630584622E-02

 8.33280963729618E-02
 8.73047490932535E-02

 6.49584507620986E-04
 4.12640607286933E-04

 7.25117589902496E-04
 5.12674693901947E-04

 4.07878644320154E-03
 2.08821155808842E-03

 8.85247724339297E-03
 1.22916883928199E-02

 9.23014265480052E-03
 1.27168332609337E-02

 5.97919879340433E-02
 5.92827005802227E-02

 .11765032896168
 .131494806855436

 .038265059483813
 .03967601960368

 3.92923094028415E-02
 3.99886261243519E-02

 4.00929600750255E-02
 5.18301611274042E-02

 .916775232883599
 .997289826508383

 9.21503603834422E-04
 3.75127824806303E-04

 0
 2.50085216537535E-05

 7.553308228151E-05
 2.50085216537535E-05

 6.0426465825208E-05
 7.50255649612605E-05

 2.11492630388228E-04
 2.50085216537535E-05

 1.96386013931926E-04
 1.50051129922521E-04

 3.7766541140755E-04
 7.50255649612605E-05

 .440735535112611
 .526879534201279

 3.0213232912604E-05
 0

 6.0426465825208E-05
 3.75127824806303E-05

 4.5319849368906E-05
 5.0017043307507E-05

 2.41705863300832E-04
 0

 9.0639698737812E-05
 1.37546869095644E-04

 3.32345562038644E-04
 3.00102259845042E-04

 4.83411726601664E-04
 2.25076694883782E-04

 2.71919096213436E-04
 5.12674693901947E-04

 1.34448886461088E-03
 1.25042608268768E-05

 1.22363593296046E-03
 7.12742867131975E-04

 2.44727186592092E-03
 2.50085216537535E-05

 2.47748509883353E-03
 1.37546869095644E-04

 1.97896675577556E-03
 2.01318599312716E-03

 3.68601441533769E-03
 4.02637198625432E-03

 4.81901064956034E-03
 3.90132937798555E-03

 6.25413921290903E-03
 5.50187476382577E-03

 9.19992942188792E-03
 9.52824675008009E-03

 8.42949198261651E-03
 1.09787410059978E-02

 1.46987378119818E-02
 1.23166969144736E-02

 1.33240357144584E-02
 1.59429325542679E-02

 1.35204217283903E-02
 1.81561867206251E-02

 1.82487926792128E-02
 2.16198669696699E-02

 1.65115317867381E-02
 2.34454890503939E-02

 1.93515756805229E-02
 2.15698499263624E-02

 1.66625979513011E-02
 2.40331893092571E-02

 2.88234241986242E-02
 2.69091692994388E-02

 2.54848619617815E-02
 2.98476705937548E-02

 2.58323141402764E-02
 .032160958846727

 .032766251093719
 3.78003804796484E-02

 3.42466995064366E-02
 4.52404156716401E-02

 4.45796251625472E-02
 5.77696850201706E-02

 4.33408826131304E-02
 6.26963637859601E-02

 4.99273673880781E-02
 5.92701963193958E-02

 .235179804991709
 .212947561881711

 3.0213232912604E-05
 1.25042608268768E-05

 1.20852931650416E-04
 1.25042608268768E-05

 2.87025712669738E-04
 0

 2.56812479757134E-04
 2.12572434056905E-04

 4.07878644320154E-04
 2.25076694883782E-04

 4.83411726601664E-04
 2.87597999018165E-04

 2.52280494820243E-03
 1.4004772126102E-03

 2.20556600262009E-03
 1.87563912403151E-03

 2.08471307096968E-03
 2.01318599312716E-03

 1.29916901524197E-03
 2.6884160777785E-03

 3.21770930519233E-03
 2.60088625199037E-03

 7.87054717373334E-03
 4.75161911421317E-03

 6.0426465825208E-03
 7.61509484356795E-03

 .013505315111934
 .011466407178246

 1.32182893992642E-02
 1.35171059538538E-02

 2.10133034907161E-02
 1.59929495975754E-02

 4.46400516283724E-02
 4.82789510525712E-02

 .11597349453503
 9.99965738325334E-02

 3.30381701899325E-02
 3.91383363881243E-02

 1.66172781019322E-04
 1.62555390749398E-04

 1.5106616456302E-04
 3.37615042325672E-04

 1.81279397475624E-04
 3.25110781498796E-04

 8.45970521552912E-04
 4.37649128940687E-04

 3.16936813253216E-02
 3.78754060446097E-02

 .206900218985512
 .217949266212462

 8.79205077756776E-03
 8.21529936325803E-03

 9.13950295606271E-03
 9.14061466444691E-03

 .012855730604313
 1.58804112501335E-02

 1.83998588437758E-02
 1.84062719371626E-02

 1.97141344754741E-02
 2.58838199116349E-02

 3.95491218825986E-02
 .037037620569209

 4.84015991259916E-02
 .051742631301616

 5.00482203197285E-02
 .051642597215001

 .140793665372735
 .155290415208982

 7.17564281674345E-03
 3.96385068211993E-03

 1.5106616456302E-05
 2.50085216537535E-05

 5.89158041795778E-04
 6.62725823824468E-04

 1.05746315194114E-03
 5.50187476382577E-04

 5.51391500655023E-03
 2.72592886025913E-03

 3.09685637354191E-03
 2.75093738191289E-03

 3.0213232912604E-05
 2.50085216537535E-05

 3.06664314062931E-03
 2.72592886025913E-03

 .119689722183281
 .140460361868307

 4.5319849368906E-05
 2.50085216537535E-05

 5.89158041795778E-04
 3.37615042325672E-04

 1.21306130144105E-02
 1.12788432658428E-02

 1.06501646016929E-02
 1.29043971733368E-02

 1.94120021463481E-02
 .024758436437216

 2.13456490527547E-02
 2.75218780799557E-02

 2.66782846618293E-02
 .029622593898871

 2.88385308150805E-02
 3.40115894491048E-02

 2.87025712669738E-04
 1.25042608268768E-05

 2.87025712669738E-04
 1.25042608268768E-05

 1.05444182864988E-02
 8.10276101581614E-03

 1.81279397475624E-04
 2.62589477364412E-04

 3.0213232912604E-04
 2.75093738191289E-04

 1.75236750893103E-03
 1.31294738682206E-03

 4.00325336092003E-03
 2.81345868604727E-03

 4.30538569004607E-03
 3.43867172739111E-03

 .435327366421255
 .408164081910911

 1.5106616456302E-04
 7.50255649612605E-05

 1.5106616456302E-05
 3.75127824806303E-05

 1.35959548106718E-04
 3.75127824806303E-05

 .435176300256692
 .40808905634595

 5.58944808883174E-04
 6.37717302170715E-04

 9.19992942188792E-03
 7.11492441049288E-03

 1.15263483561584E-02
 8.09025675498926E-03

 1.54691752512532E-02
 1.31169696073937E-02

 2.27505643831908E-02
 1.07661685719409E-02

 2.40044135490639E-02
 2.35830359194896E-02

 3.35820083823593E-02
 4.79788487927261E-02

 .114916031383089
 9.17062489043141E-02

 .203168884720806
 .205094886082433

 .267568390674021
 .298826825240701

 .265770703315721
 .29562573446902

 7.553308228151E-05
 0

 3.0213232912604E-04
 8.75298257881373E-05

 4.68305110145362E-04
 2.50085216537535E-05

 3.0213232912604E-04
 4.00136346460056E-04

 6.49584507620986E-04
 5.0017043307507E-04

 8.76183754465516E-04
 9.00306779535127E-04

 1.60130134436801E-03
 1.35046016930269E-03

 2.96089682543519E-03
 1.87563912403151E-03

 2.77961742795957E-03
 3.7637825088899E-03

 3.36877546975535E-03
 3.42616746656423E-03

 6.8281906382485E-03
 4.63908076677128E-03

 1.08767638485374E-02
 6.37717302170715E-03

 1.45627782638751E-02
 1.93941085424859E-02

 .036814824304008
 4.39399725456449E-02

 4.23287393105582E-02
 4.13390862936546E-02

 7.24060126750555E-02
 7.74513915616746E-02

 6.85689320951548E-02
 9.01557205617814E-02

 1.79768735829994E-03
 3.20109077168045E-03

 1.5106616456302E-05
 2.62589477364412E-04

 2.11492630388228E-04
 5.37683215555701E-04

 2.11492630388228E-04
 5.50187476382577E-04

 3.7766541140755E-04
 5.25178954728824E-04

 9.8193006965963E-04
 1.32545164764894E-03

 .265030479109362
 .264352578141002

 .265030479109362
 .264352578141002

 5.2873157597057E-04
 0

 5.89158041795778E-04
 0

 1.10278300131005E-03
 0

 2.9911100583478E-03
 0

 2.85515051024108E-03
 1.72558799410899E-03

 3.88240042926961E-03
 3.88882511715867E-03

 1.53332157031465E-02
 1.49050789056371E-02

 2.27656709996471E-02
 1.53302237737509E-02

 1.88379507210086E-02
 .025471179304348

 2.62250861681403E-02
 3.06854560691556E-02

 3.02736593784292E-02
 2.83596635553565E-02

 2.70710566896932E-02
 3.51869899668312E-02

 3.54854420558534E-02
 3.41991533615079E-02

 7.70890637765091E-02
 7.46004200931467E-02

 1.16774145207214E-02
 9.12811040362003E-04

 1.16774145207214E-02
 9.12811040362003E-04

 7.40224206358798E-04
 0

 8.00650672184006E-04
 0

 1.81279397475624E-03
 3.75127824806303E-05

 2.19045938616379E-03
 0

 6.13328628125861E-03
 8.75298257881373E-04

 1.33089290980021E-02
 3.37615042325672E-04

 1.33089290980021E-02
 3.37615042325672E-04

 1.11788961776635E-03
 6.25213041343838E-05

 4.93986358121075E-03
 1.25042608268768E-05

 7.25117589902496E-03
 2.62589477364412E-04

 .142802845361423
 .165031234393119

 .142802845361423
 .165031234393119

 5.74051425339476E-04
 8.5028973622762E-04

 1.34448886461088E-03
 1.18790477855329E-03

 3.53494825077467E-03
 2.48834790454847E-03

 6.51095169266616E-03
 1.13163560483235E-02

 1.08616572320811E-02
 1.69182648987643E-02

 3.35215819165341E-02
 3.78378932621291E-02

 4.52443162866245E-02
 4.41650492405287E-02

 4.12108496927918E-02
 5.02671285240446E-02

 2.59078472225579E-02
 3.43617087522573E-02

 2.59078472225579E-02
 3.43617087522573E-02

 9.66823453203328E-04
 5.50187476382577E-04

 2.49410237693546E-02
 3.38115212758748E-02

 1.66777045677574E-02
 1.12538347441891E-04

 1.66777045677574E-02
 1.12538347441891E-04

 2.97600344189149E-03
 0

 3.45941516849316E-03
 1.25042608268768E-05

 4.92475696475445E-03
 7.50255649612605E-05

 5.3175289926183E-03
 2.50085216537535E-05

 6.4965550871833
 5.94698893648011

 .163831255468595
 .186588580058655

 .163423376824275
 .186238460755502

 1.5106616456302E-05
 1.25042608268768E-05

 1.5106616456302E-04
 1.25042608268768E-05

 6.0426465825208E-04
 3.37615042325672E-04

 7.40224206358798E-04
 7.12742867131975E-04

 2.49259171528983E-03
 2.45083512206784E-03

 2.09981968742598E-03
 3.1760822500267E-03

 5.72540763693846E-03
 2.76344164273976E-03

 4.65283786854102E-03
 4.22644015948434E-03

 1.37772342081474E-02
 1.39547550827945E-02

 1.36865945094096E-02
 1.57678729026916E-02

 1.90041235020279E-02
 1.84312804588163E-02

 2.18894872451816E-02
 2.05069877560779E-02

 2.06960645451337E-02
 2.30578569647607E-02

 1.73423956918347E-02
 2.61839221714799E-02

 4.05461585687146E-02
 5.46436198134514E-02

 4.07878644320154E-04
 3.50119303152549E-04

 4.5319849368906E-05
 0

 4.5319849368906E-05
 1.25042608268768E-05

 6.0426465825208E-05
 1.25042608268768E-05

 7.553308228151E-05
 6.25213041343838E-05

 4.5319849368906E-05
 1.25042608268768E-04

 1.35959548106718E-04
 1.37546869095644E-04

 2.8300433136907
 2.67192295774785

 .135219323900359
 .112650885789333

 0
 2.50085216537535E-05

 4.38091877232758E-04
 3.37615042325672E-04

 6.64691124077288E-04
 3.75127824806303E-04

 6.0426465825208E-04
 5.25178954728824E-04

 1.79768735829994E-03
 1.35046016930269E-03

 3.44430855203686E-03
 0

 3.39898870266795E-03
 2.62589477364412E-03

 9.0639698737812E-03
 2.71342459943226E-03

 1.40491533043609E-02
 8.16528231995052E-03

 4.47609045600228E-02
 3.71501589166508E-02

 5.69972638896274E-02
 5.93827346668377E-02

 .374855580746678
 .407113724001454

 1.5106616456302E-05
 3.75127824806303E-05

 4.63622059043908E-02
 3.90508065623361E-02

 5.17099481299217E-02
 6.11083226609467E-02

 6.12422231138483E-02
 6.32465512623426E-02

 .215526096982061
 .243670530733347

 5.04107791146798E-02
 4.73161229689017E-02

 4.5319849368906E-05
 1.25042608268768E-05

 1.81279397475624E-04
 1.25042608268768E-05

 4.98518343057966E-04
 1.62555390749398E-04

 2.11492630388228E-04
 6.50221562997591E-04

 9.36610220290724E-03
 5.95202815359334E-03

 1.58166274297482E-02
 1.80436483731832E-02

 2.42914392617336E-02
 2.24826609667244E-02

 .33166576429811
 .342154089005829

 9.0639698737812E-05
 2.50085216537535E-05

 1.5106616456302E-05
 8.75298257881373E-05

 2.11492630388228E-04
 1.12538347441891E-04

 1.57108811145541E-03
 1.13788773524579E-03

 4.62866728221093E-02
 6.74479829001732E-02

 .104900344672561
 .103197664604214

 .178590419746402
 .170145477071312

 1.35959548106718E-04
 0

 1.35959548106718E-04
 0

 .102634352204116
 6.95486987190885E-02

 2.41705863300832E-04
 1.25042608268768E-05

 4.07878644320154E-04
 4.37649128940687E-04

 4.83411726601664E-04
 4.12640607286933E-04

 3.15728283936712E-03
 3.16357798919982E-03

 .011556561589071
 3.50119303152549E-03

 1.17831608359156E-02
 1.39297465611407E-02

 7.50043507055394E-02
 .048091387140168

 .865578909713192
 .850752393878214

 2.41705863300832E-04
 2.87597999018165E-04

 2.56812479757134E-04
 3.12606520671919E-04

 5.58944808883174E-04
 3.87632085633179E-04

 8.45970521552912E-04
 4.50153389767563E-04

 1.26895578232937E-03
 2.50085216537535E-04

 2.64365787985285E-03
 2.62589477364412E-03

 4.69815771790992E-03
 4.9516872874432E-03

 9.54738160038286E-03
 8.41536753648806E-03

 1.26442379739248E-02
 8.45288031896869E-03

 2.18743806287253E-02
 2.68216394736506E-02

 4.16640481864809E-02
 4.45026642828544E-02

 3.87937910597835E-02
 5.94827687534527E-02

 5.94898556049173E-02
 4.66283886234234E-02

 5.72993962187535E-02
 5.42309792061645E-02

 5.84172858365198E-02
 5.40934323370689E-02

 6.34931089658373E-02
 6.25213041343838E-02

 6.78438145052523E-02
 6.91610666334554E-02

 6.53210095570498E-02
 7.85767750360935E-02

 7.93097363955855E-02
 7.50755820045681E-02

 7.93097363955855E-02
 8.03773885951638E-02

 8.37057617843694E-02
 7.98146968579544E-02

 .116351159946438
 9.33318028118081E-02

 .741326989360108
 .625200537083011

 5.74051425339476E-04
 2.50085216537535E-04

 4.47155847106539E-03
 2.98851833762355E-03

 8.67119784591735E-03
 4.71410633173254E-03

 5.02899261830293E-02
 6.00704690123159E-02

 .140672812441084
 .143561418553372

 .201733756157457
 .154640193645985

 .334913686836215
 .258975745985445

 5.51995765313275E-02
 5.21302633872492E-02

 8.45970521552912E-04
 6.25213041343838E-05

 5.89158041795778E-04
 3.25110781498796E-04

 5.89158041795778E-04
 4.00136346460056E-04

 7.553308228151E-04
 4.00136346460056E-04

 9.21503603834422E-04
 5.50187476382577E-04

 1.10278300131005E-03
 4.37649128940687E-04

 1.40491533043609E-03
 9.2531530118888E-04

 4.89907571677874E-02
 4.90292067021838E-02

 1.16320946713525E-03
 3.75127824806303E-04

 1.16320946713525E-03
 3.75127824806303E-04

 1.16320946713525E-03
 8.37785475400743E-04

 1.16320946713525E-03
 8.37785475400743E-04

 1.33391423309147E-02
 5.93952389276646E-03

 1.23874254941676E-03
 8.5028973622762E-04

 1.21003997814979E-02
 5.08923415653884E-03

 5.46708449553569E-02
 5.78947276284394E-02

 1.07256976839744E-03
 1.01284512697702E-03

 4.5319849368906E-03
 1.81311781989713E-03

 3.06664314062931E-03
 3.05103964175793E-03

 3.56516148368727E-03
 2.70092033860538E-03

 5.78583410276366E-03
 2.25076694883782E-03

 3.66486515229886E-02
 4.70660377523641E-02

 4.46098383954598E-02
 4.48027665426994E-02

 5.52902162300653E-03
 6.67727528155219E-03

 1.52274693879524E-02
 2.04069536694629E-02

 2.38533473845009E-02
 1.77185375916844E-02

 5.80698336580249E-02
 5.52063115506609E-02

 1.35355283448466E-02
 1.64431029873429E-02

 4.45343053131783E-02
 3.87632085633179E-02

 2.0439100999212
 1.86789898657968

 .413030000531753
 .355608673655548

 3.0213232912604E-05
 0

 6.64691124077288E-03
 0

 .11695542460469
 .125330206267786

 .289397451453377
 .230278467387762

 .304791093622349
 .31860856586882

 4.5319849368906E-05
 0

 1.26895578232937E-03
 2.87597999018165E-04

 1.99407337223186E-03
 2.35080103545283E-03

 5.06071651286117E-03
 2.11322007974217E-03

 1.40189400714483E-02
 2.50085216537535E-05

 1.31880761663516E-02
 1.96066809765428E-02

 .10686420481188
 .137584381878125

 .162350807055878
 .156640875378285

 7.553308228151E-05
 2.50085216537535E-05

 7.553308228151E-05
 2.50085216537535E-05

 .949299778114018
 .897893457195539

 1.35959548106718E-04
 1.00034086615014E-04

 1.96386013931926E-04
 7.50255649612605E-05

 1.81279397475624E-04
 1.62555390749398E-04

 2.2659924684453E-04
 2.62589477364412E-04

 4.5319849368906E-04
 3.62623563979426E-04

 7.553308228151E-04
 4.75161911421317E-04

 6.19371274708382E-04
 7.50255649612605E-04

 9.66823453203328E-04
 9.12811040362003E-04

 4.59241140271581E-03
 5.75195998036331E-03

 1.07710175333433E-02
 9.72831492331012E-03

 .029261516075857
 2.66215713004206E-02

 3.03340858442544E-02
 3.06854560691556E-02

 3.53494825077467E-02
 3.51619814451774E-02

 3.70263169343962E-02
 4.23644356814585E-02

 4.39753605042951E-02
 5.33181681658025E-02

 8.23008464539333E-02
 .068585870635419

 8.43100264426214E-02
 7.69887339110802E-02

 .111124270652557
 .113238586048196

 .123315310132793
 .12539272757192

 .163453590057188
 .145199476721693

 .189950595321541
 .161755118056478

 .109794888404403
 4.29896487228023E-02

 1.31427563169827E-03
 0

 1.40491533043609E-03
 0

 3.18749607227972E-03
 0

 3.39898870266795E-03
 0

 3.59537471659988E-03
 0

 3.97304012800743E-03
 0

 4.5168783204343E-03
 0

 5.24199591033679E-03
 0

 5.34774222553091E-03
 0

 6.8130840217922E-03
 0

 2.06054248463959E-02
 0

 5.03956724982235E-02
 4.29896487228023E-02

 .2669188061664
 .252773632615314

 .123466376297356
 .117102402643701

 .143452429869044
 .135671229971613

 3.73284492635222E-02
 2.33954720070864E-02

 3.0213232912604E-04
 7.50255649612605E-05

 6.0426465825208E-05
 0

 2.41705863300832E-04
 7.50255649612605E-05

 1.32333960157205E-02
 7.62759910439482E-03

 4.5319849368906E-05
 5.0017043307507E-05

 7.553308228151E-05
 3.75127824806303E-05

 1.05746315194114E-04
 6.25213041343838E-05

 7.553308228151E-05
 1.00034086615014E-04

 1.66172781019322E-04
 6.25213041343838E-05

 1.81279397475624E-04
 6.25213041343838E-05

 1.5106616456302E-04
 1.25042608268768E-04

 2.41705863300832E-04
 1.37546869095644E-04

 2.41705863300832E-04
 3.50119303152549E-04

 1.11788961776635E-03
 0

 3.7464408811629E-03
 2.60088625199037E-03

 7.08500311800564E-03
 4.03887624708119E-03

 1.5106616456302E-03
 1.47550277757146E-03

 6.0426465825208E-05
 6.25213041343838E-05

 1.5106616456302E-04
 1.12538347441891E-04

 1.05746315194114E-04
 1.87563912403151E-04

 2.11492630388228E-04
 1.50051129922521E-04

 1.66172781019322E-04
 1.87563912403151E-04

 3.47452178494946E-04
 2.87597999018165E-04

 4.68305110145362E-04
 4.87666172248194E-04

 1.37017011258659E-02
 9.06558909948565E-03

 9.0639698737812E-05
 7.50255649612605E-05

 1.93364690640666E-03
 5.0017043307507E-04

 1.16774145207214E-02
 8.49039310144932E-03

 8.58055814717953E-03
 5.15175546067322E-03

 2.41705863300832E-04
 1.62555390749398E-04

 7.553308228151E-04
 6.87734345478222E-04

 1.93364690640666E-03
 7.50255649612605E-04

 2.67387111276545E-03
 1.43798999509083E-03

 2.97600344189149E-03
 2.11322007974217E-03

 1.34648293798311
 1.13723751368279

 1.61640796082431E-03
 1.51301556005209E-03

 1.5106616456302E-05
 1.25042608268768E-04

 1.66172781019322E-04
 3.75127824806303E-05

 2.71919096213436E-04
 1.50051129922521E-04

 5.43838192426872E-04
 6.00204519690084E-04

 6.19371274708382E-04
 6.00204519690084E-04

 .614658010374016
 .53091841044836

 1.05746315194114E-04
 2.25076694883782E-04

 3.24792253810493E-03
 2.56337346950974E-03

 6.17860613062752E-03
 1.28793886516831E-03

 .103812668287707
 6.76230425517495E-02

 9.05792722719868E-02
 9.95089076602852E-02

 .410733794830395
 .359710071206764

 5.69519440402585E-02
 5.43935345969139E-02

 4.07878644320154E-04
 2.37580955710658E-04

 5.74051425339476E-04
 4.00136346460056E-04

 2.06960645451337E-03
 1.92565616733902E-03

 2.82493727732847E-03
 1.80061355907025E-03

 1.74330353905725E-02
 1.21541415237242E-02

 3.36424348481845E-02
 3.78754060446097E-02

 6.46412118165162E-02
 6.15459717898874E-02

 5.89158041795778E-04
 3.75127824806303E-04

 3.7766541140755E-04
 5.62691737209454E-04

 9.94015362824671E-03
 1.41423189951976E-02

 5.37342347350662E-02
 .046465833232674

 3.01074865974099E-02
 1.82187080247594E-02

 9.21503603834422E-04
 1.50051129922521E-04

 1.02724991902854E-03
 8.00272692920113E-04

 3.62558794951248E-03
 6.50221562997591E-04

 6.46563184329725E-03
 3.60122711814051E-03

 1.80675132817372E-02
 1.30169355207787E-02

 .578507877194085
 .470647873262814

 .284004389378478
 .230015877910398

 .294503487815607
 .240631995352416

 7.49590308561705E-02
 5.99454264040472E-02

 6.0124333496082E-03
 0

 2.11492630388228E-03
 0

 3.89750704572592E-03
 0

 6.89465975065623E-02
 5.99454264040472E-02

 6.89465975065623E-02
 5.99454264040472E-02

 .375535378487211
 .278857520700179

 .3449444801632
 .258037926423429

 2.75242551833822E-02
 1.42673616034664E-02

 3.0213232912604E-05
 0

 1.5106616456302E-05
 1.25042608268768E-05

 4.5319849368906E-05
 1.25042608268768E-05

 7.553308228151E-05
 0

 7.553308228151E-05
 1.25042608268768E-05

 6.0426465825208E-05
 5.0017043307507E-05

 7.553308228151E-05
 7.50255649612605E-05

 1.81279397475624E-04
 2.50085216537535E-05

 5.58944808883174E-04
 3.37615042325672E-04

 7.10010973446194E-04
 8.37785475400743E-04

 2.05449983805707E-03
 3.12606520671919E-04

 7.14542958383084E-03
 6.2396261526115E-03

 1.64964251702818E-02
 6.35216450005339E-03

 .197790929262362
 .130632012858381

 3.0213232912604E-05
 0

 3.0213232912604E-05
 0

 1.5106616456302E-05
 3.75127824806303E-05

 3.0213232912604E-05
 8.75298257881373E-05

 8.76183754465516E-04
 4.00136346460056E-04

 1.07256976839744E-03
 1.46299851674458E-03

 3.33856223684274E-03
 1.70057947245524E-03

 4.09389305965784E-03
 2.30078399214532E-03

 4.16942614193935E-03
 3.91383363881243E-03

 4.62262463562841E-03
 3.81379955219741E-03

 6.60159139140397E-03
 4.21393589865747E-03

 8.85247724339297E-03
 5.25178954728824E-03

 1.00156867105282E-02
 6.26463467426526E-03

 1.09674035472752E-02
 6.22712189178463E-03

 .010106326409266
 8.09025675498926E-03

 1.16623079042651E-02
 8.02773545085488E-03

 1.34146754131962E-02
 8.25281214573866E-03

 2.36720679870252E-02
 1.55678047294616E-02

 2.45935715908597E-02
 1.49050789056371E-02

 2.95032219391578E-02
 1.76435120267231E-02

 3.01225932138662E-02
 2.24701567058975E-02

 2.09981968742598E-03
 2.87597999018165E-03

 3.0213232912604E-05
 0

 4.5319849368906E-05
 0

 1.5106616456302E-04
 2.50085216537535E-05

 7.553308228151E-05
 1.37546869095644E-04

 1.05746315194114E-04
 1.25042608268768E-04

 1.69194104310582E-03
 2.58838199116349E-03

 7.40224206358798E-04
 3.50119303152549E-04

 4.5319849368906E-05
 0

 6.94904356989892E-04
 3.50119303152549E-04

 3.0213232912604E-05
 3.75127824806303E-05

 3.0213232912604E-05
 3.75127824806303E-05

 1.5106616456302E-05
 5.0017043307507E-05

 1.5106616456302E-05
 5.0017043307507E-05

 3.17238945582342E-04
 3.75127824806303E-05

 9.0639698737812E-05
 3.75127824806303E-05

 2.2659924684453E-04
 0

 1.37017011258659E-02
 1.40172763869288E-02

 1.20852931650416E-04
 6.25213041343838E-05

 1.66172781019322E-04
 1.62555390749398E-04

 1.34146754131962E-02
 1.37921996920451E-02

 5.01539666349226E-03
 2.6884160777785E-03

 4.22985260776456E-04
 3.00102259845042E-04

 8.15757288640308E-04
 2.50085216537535E-04

 9.36610220290724E-04
 5.25178954728824E-04

 2.84004389378478E-03
 1.6130496466671E-03

 1.59374803613986E-02
 1.31044653465668E-02

 6.49584507620986E-04
 4.50153389767563E-04

 1.07256976839744E-03
 1.05035790945765E-03

 1.45023517980499E-03
 1.66306668997461E-03

 4.01835997737633E-03
 4.35148276775311E-03

 8.74673092819886E-03
 5.58940458961391E-03

 1.65115317867381E-02
 1.35671229971613E-02

 9.97036686115932E-04
 9.50323822842634E-04

 8.3086390509661E-04
 1.57553686418647E-03

 4.86433049892924E-03
 3.7137654655824E-03

 9.8193006965963E-03
 7.32749684454978E-03

 5.00482203197285E-02
 5.63441992859067E-02

 4.71326433436622E-03
 3.51369729235237E-03

 4.72837095082252E-03
 5.40184067721076E-03

 1.29010504536819E-02
 7.79015449514422E-03

 9.21503603834422E-03
 1.55177876861541E-02

 1.84904985425136E-02
 2.41207191350453E-02

 1.52123627714961E-02
 1.00659299656358E-02

 1.52123627714961E-02
 1.00659299656358E-02

 3.05908983240115E-02
 2.08195942767498E-02

 2.19499137110068E-02
 1.41423189951976E-02

 3.0213232912604E-05
 1.25042608268768E-05

 2.41705863300832E-04
 6.25213041343838E-05

 9.66823453203328E-04
 2.87597999018165E-04

 3.55005486723097E-03
 1.82562208072401E-03

 3.58026810014357E-03
 2.62589477364412E-03

 1.35808481942155E-02
 9.32817857685006E-03

 8.64098461300474E-03
 6.67727528155219E-03

 8.64098461300474E-03
 6.67727528155219E-03

 1.28454581051227
 1.33059089884878

 .329520624761315
 .326386216103137

 3.00168468986721E-02
 4.38899555023374E-02

 3.0213232912604E-05
 0

 3.0213232912604E-05
 0

 4.5319849368906E-05
 3.75127824806303E-05

 9.0639698737812E-05
 2.50085216537535E-05

 2.41705863300832E-04
 3.00102259845042E-04

 2.55301818111504E-03
 1.20040903938017E-03

 1.16472012878088E-02
 1.17540051772642E-02

 1.53785355525154E-02
 3.05729177217137E-02

 .045818367711964
 .043227229678513

 4.5319849368906E-05
 3.75127824806303E-05

 2.11492630388228E-04
 0

 9.0639698737812E-05
 1.25042608268768E-04

 2.87025712669738E-04
 1.62555390749398E-04

 4.51838898207993E-02
 4.29021188970142E-02

 5.14531356501646E-02
 4.97794623517964E-02

 1.35959548106718E-04
 1.25042608268768E-04

 1.96386013931926E-04
 1.37546869095644E-04

 3.92772027863852E-03
 1.25042608268768E-04

 6.0728598154334E-03
 5.37683215555701E-04

 .041120209994054
 4.88541470506075E-02

 .1088280649512
 8.59167761414702E-02

 3.7766541140755E-04
 0

 4.74347756727883E-03
 1.25042608268768E-05

 4.5470915533469E-03
 4.87666172248194E-04

 1.48649105930012E-02
 1.22166628278586E-02

 8.42949198261651E-02
 7.31999428805365E-02

 8.91290370921818E-04
 1.75059651576275E-04

 3.7766541140755E-04
 1.00034086615014E-04

 5.13624959514268E-04
 7.50255649612605E-05

 7.00040606585035E-02
 .072849823577384

 1.82941125285817E-02
 .022932814356492

 5.17099481299217E-02
 .049917009220892

 .02250885851989
 3.05479092000599E-02

 .02250885851989
 3.05479092000599E-02

 .650521117841277
 .715631351382984

 4.81901064956034E-03
 4.48902963684876E-03

 4.5319849368906E-05
 0

 2.41705863300832E-04
 3.75127824806303E-05

 7.553308228151E-05
 3.00102259845042E-04

 2.71919096213436E-04
 1.87563912403151E-04

 4.18453275839565E-03
 3.96385068211993E-03

 7.553308228151E-04
 5.50187476382577E-04

 3.0213232912604E-05
 1.25042608268768E-05

 7.25117589902496E-04
 5.37683215555701E-04

 4.75254153715261E-02
 4.76162252287467E-02

 9.0639698737812E-05
 3.75127824806303E-05

 3.7766541140755E-04
 2.37580955710658E-04

 4.22985260776456E-04
 2.12572434056905E-04

 2.52280494820243E-03
 1.73809225493587E-03

 4.41113200524018E-02
 4.53904668015626E-02

 7.00947003572413E-03
 .006189609109304

 1.35959548106718E-04
 6.25213041343838E-05

 1.81279397475624E-04
 1.25042608268768E-04

 2.11492630388228E-04
 1.25042608268768E-04

 7.10010973446194E-04
 3.37615042325672E-04

 5.89158041795778E-04
 6.75230084651345E-04

 9.36610220290724E-04
 8.12776953746989E-04

 1.14810285067895E-03
 9.50323822842634E-04

 1.69194104310582E-03
 1.38797295178332E-03

 1.40491533043609E-03
 1.71308373328212E-03

 2.2810990849016E-03
 1.05035790945765E-03

 2.41705863300832E-04
 0

 4.68305110145362E-04
 0

 1.57108811145541E-03
 1.05035790945765E-03

 8.85247724339297E-03
 8.00272692920112E-03

 2.41705863300832E-04
 0

 1.5106616456302E-05
 2.12572434056905E-04

 2.87025712669738E-04
 0

 3.0213232912604E-04
 5.0017043307507E-05

 1.35959548106718E-04
 2.12572434056905E-04

 6.0426465825208E-05
 3.12606520671919E-04

 4.68305110145362E-04
 0

 1.96386013931926E-04
 2.37580955710658E-04

 1.66172781019322E-04
 3.12606520671919E-04

 4.22985260776456E-04
 1.25042608268768E-04

 1.5106616456302E-05
 4.6265765059444E-04

 2.87025712669738E-04
 4.50153389767563E-04

 5.13624959514268E-04
 3.37615042325672E-04

 4.07878644320154E-04
 5.87700258863208E-04

 8.61077138009214E-04
 7.37751388785729E-04

 7.40224206358798E-04
 8.5028973622762E-04

 1.82790059121254E-03
 1.08787069193828E-03

 1.90343367349405E-03
 2.02569025395404E-03

 1.20852931650416E-04
 1.00034086615014E-04

 1.20852931650416E-04
 1.00034086615014E-04

 .579157461701706
 .647633181006428

 1.66172781019322E-04
 2.12572434056905E-04

 5.13624959514268E-04
 1.87563912403151E-04

 9.21503603834422E-04
 3.25110781498796E-04

 1.17831608359156E-03
 1.18790477855329E-03

 1.64662119373692E-03
 1.86313486320464E-03

 2.43216524946462E-03
 1.87563912403151E-03

 1.53634289360591E-02
 1.97192193239846E-02

 2.59682736883831E-02
 2.05445005385585E-02

 3.73737691128911E-02
 3.53245368359268E-02

 4.95799152095832E-02
 5.16551014758279E-02

 5.88553777137526E-02
 7.04990225419312E-02

 6.79344542039901E-02
 7.83642026020367E-02

 6.74359358609321E-02
 8.15527891128902E-02

 .129524129496333
 .132557669025721

 .12026377360862
 .151764213655803

 2.11341564223665E-02
 .013904738039487

 2.11341564223665E-02
 .013904738039487

 6.0426465825208E-05
 3.75127824806303E-05

 5.58944808883174E-04
 2.50085216537535E-05

 1.66172781019322E-04
 4.00136346460056E-04

 2.91557697606629E-03
 2.35080103545283E-03

 7.44756191295688E-03
 2.3132882529722E-03

 4.26006584067716E-03
 4.96419154827007E-03

 5.72540763693846E-03
 3.81379955219741E-03

 .269864596375379
 .264665184661673

 .269864596375379
 .264665184661673

 6.0426465825208E-05
 1.00034086615014E-04

 2.2659924684453E-04
 2.25076694883782E-04

 4.98518343057966E-04
 4.87666172248194E-04

 2.17535276970749E-03
 1.70057947245524E-03

 2.82493727732847E-03
 1.58804112501335E-03

 3.42920193558055E-03
 1.82562208072401E-03

 5.18156944451159E-03
 3.2260992933342E-03

 5.12114297868638E-03
 4.32647424609936E-03

 4.65283786854102E-03
 5.42684919886451E-03

 9.95526024470302E-03
 8.26531640656554E-03

 1.41095797701861E-02
 9.55325527173384E-03

 1.65115317867381E-02
 1.01534597914239E-02

 1.26895578232937E-02
 1.78935972432606E-02

 3.36877546975535E-02
 2.03319281045016E-02

 2.68746706757613E-02
 2.85597317285865E-02

 3.59235339330862E-02
 4.82789510525712E-02

 4.69060440968177E-02
 .050729786174639

 4.90360770171563E-02
 5.19927165181536E-02

 .013505315111934
 1.00034086615014E-02

 .013505315111934
 1.00034086615014E-02

 2.71919096213436E-04
 5.87700258863208E-04

 1.19342270004786E-03
 5.50187476382577E-04

 3.21770930519233E-03
 2.38831381793346E-03

 8.82226401048037E-03
 6.47720710832216E-03

 3.23337486645751
 4.31618323943884

 3.18232960945167
 4.25422462704166

 .154767285594814
 .125842880961688

 3.0213232912604E-05
 0

 3.0213232912604E-05
 0

 4.5319849368906E-05
 0

 6.0426465825208E-05
 1.25042608268768E-05

 9.0639698737812E-05
 0

 0
 1.25042608268768E-04

 3.0213232912604E-05
 1.00034086615014E-04

 6.0426465825208E-05
 8.75298257881373E-05

 1.5106616456302E-04
 8.75298257881373E-05

 2.71919096213436E-04
 2.50085216537535E-05

 2.87025712669738E-04
 1.87563912403151E-04

 5.13624959514268E-04
 1.50051129922521E-04

 6.49584507620986E-04
 3.00102259845042E-04

 8.3086390509661E-04
 2.00068173230028E-04

 7.70437439271402E-04
 3.37615042325672E-04

 9.8193006965963E-04
 3.00102259845042E-04

 7.553308228151E-04
 5.50187476382577E-04

 9.51716836747026E-04
 7.00238606305098E-04

 1.14810285067895E-03
 6.12708780516961E-04

 3.70112103179399E-03
 3.40115894491048E-03

 7.61373469397621E-03
 5.92701963193958E-03

 8.88269047630557E-03
 1.14288943957654E-02

 .013006796768876
 1.10037495276515E-02

 1.26895578232937E-02
 1.13538688308041E-02

 1.48951238259138E-02
 1.11788091792278E-02

 1.53634289360591E-02
 1.14288943957654E-02

 2.73278691694503E-02
 2.29203100956651E-02

 4.36279083258002E-02
 3.34238891902416E-02

 1.31427563169827E-03
 9.12811040362003E-04

 4.5319849368906E-05
 0

 1.05746315194114E-04
 3.75127824806303E-05

 7.40224206358798E-04
 2.50085216537535E-04

 4.22985260776456E-04
 6.25213041343838E-04

 2.47293800728018
 3.5137598136565

 3.0213232912604E-05
 1.25042608268768E-05

 0
 5.0017043307507E-05

 6.0426465825208E-05
 7.50255649612605E-05

 0
 1.25042608268768E-04

 7.553308228151E-05
 1.12538347441891E-04

 1.5106616456302E-04
 1.25042608268768E-04

 4.98518343057966E-04
 5.0017043307507E-04

 1.04235653548484E-03
 2.25076694883782E-04

 8.91290370921818E-04
 1.50051129922521E-03

 3.44430855203686E-03
 3.11356094589231E-03

 3.53494825077467E-03
 4.01386772542744E-03

 8.39927874970391E-03
 .007102420149666

 1.14357086574206E-02
 8.15277805912365E-03

 1.73121824589221E-02
 .010553596137884

 2.42606217641628
 3.47809766177825

 .303899803251427
 .306204339128558

 4.5319849368906E-05
 1.25042608268768E-05

 6.0426465825208E-05
 5.0017043307507E-05

 9.0639698737812E-05
 8.75298257881373E-05

 3.17238945582342E-04
 3.75127824806303E-04

 1.07256976839744E-03
 5.12674693901947E-04

 1.97896675577556E-03
 2.26327120966469E-03

 6.42484397886524E-02
 7.76014426915972E-02

 .236086201979088
 .225301771578665

 6.70733770659809E-03
 5.87700258863208E-03

 3.0213232912604E-05
 6.25213041343838E-05

 1.96386013931926E-04
 3.12606520671919E-04

 6.48073845975356E-03
 5.50187476382577E-03

 8.03823061639829E-02
 .104485603469382

 7.553308228151E-05
 3.75127824806303E-05

 1.05746315194114E-04
 3.75127824806303E-05

 2.2659924684453E-04
 0

 3.0213232912604E-05
 2.37580955710658E-04

 6.0426465825208E-05
 3.00102259845042E-04

 9.0639698737812E-05
 3.12606520671919E-04

 1.35959548106718E-04
 4.37649128940687E-04

 1.01214330257223E-03
 9.00306779535127E-04

 1.46534179626129E-03
 8.62793997054496E-04

 6.94904356989892E-04
 1.78810929824338E-03

 2.93068359252259E-03
 2.95100555514291E-03

 7.35541155257344E-02
 9.66204234092767E-02

 6.68618844355926E-02
 9.51824334141859E-02

 9.0639698737812E-05
 3.75127824806303E-05

 5.13624959514268E-04
 1.50051129922521E-04

 9.0639698737812E-04
 3.62623563979426E-04

 6.94904356989892E-04
 9.6282808366951E-04

 6.46563184329726E-02
 9.36694178541338E-02

 1.87322044058145E-02
 2.28577887915307E-02

 1.5106616456302E-04
 7.50255649612605E-05

 1.14810285067895E-03
 1.33795590847581E-03

 1.74330353905725E-02
 2.14448073180936E-02

 4.84620255918168E-02
 4.60281841037333E-02

 1.5106616456302E-04
 1.37546869095644E-04

 3.47452178494946E-04
 3.75127824806303E-05

 1.23874254941676E-03
 3.62623563979426E-04

 7.70437439271402E-04
 9.12811040362003E-04

 9.1092897231501E-03
 5.15175546067322E-03

 1.70704765956213E-02
 2.07195601901348E-02

 1.97745609412993E-02
 1.87063741970076E-02

 1.81279397475624E-03
 9.75332344496387E-04

 1.5106616456302E-04
 3.00102259845042E-04

 1.66172781019322E-03
 6.75230084651345E-04

 2.26448180679967E-02
 2.92474660740647E-02

 7.553308228151E-05
 4.50153389767563E-04

 2.71919096213436E-04
 4.6265765059444E-04

 2.87025712669738E-04
 5.50187476382577E-04

 1.94875352286296E-03
 1.52551982087896E-03

 1.76747412538733E-03
 3.67625268310177E-03

 1.82941125285817E-02
 2.25826950533394E-02

 1.63151457728062E-03
 9.75332344496387E-04

 6.94904356989892E-04
 4.87666172248194E-04

 9.36610220290724E-04
 4.87666172248194E-04

 2.17535276970749E-03
 1.87563912403151E-03

 6.0426465825208E-04
 5.62691737209454E-04

 7.40224206358798E-04
 6.00204519690084E-04

 8.3086390509661E-04
 7.12742867131975E-04

 5.10452570058445E-02
 6.19586123971743E-02

 7.553308228151E-05
 3.75127824806303E-05

 3.0213232912604E-05
 0

 4.5319849368906E-05
 3.75127824806303E-05

 8.45970521552912E-04
 1.25042608268768E-03

 1.5106616456302E-05
 1.25042608268768E-05

 3.0213232912604E-05
 0

 1.20852931650416E-04
 1.00034086615014E-04

 1.05746315194114E-04
 2.37580955710658E-04

 4.5319849368906E-04
 5.0017043307507E-05

 1.20852931650416E-04
 8.5028973622762E-04

 2.53791156465874E-03
 8.5028973622762E-04

 3.0213232912604E-05
 0

 4.5319849368906E-05
 0

 3.0213232912604E-05
 2.50085216537535E-05

 7.553308228151E-05
 0

 1.05746315194114E-04
 1.25042608268768E-05

 1.81279397475624E-04
 0

 1.66172781019322E-04
 3.75127824806303E-05

 2.11492630388228E-04
 3.75127824806303E-05

 2.2659924684453E-04
 5.0017043307507E-05

 1.20852931650416E-04
 1.37546869095644E-04

 2.2659924684453E-04
 1.87563912403151E-04

 5.58944808883174E-04
 6.25213041343838E-05

 5.58944808883174E-04
 3.00102259845042E-04

 0
 2.50085216537535E-05

 0
 2.50085216537535E-05

 9.0639698737812E-05
 3.75127824806303E-05

 1.5106616456302E-05
 1.25042608268768E-05

 3.0213232912604E-05
 1.25042608268768E-05

 4.5319849368906E-05
 1.25042608268768E-05

 7.553308228151E-05
 2.50085216537535E-05

 3.0213232912604E-05
 1.25042608268768E-05

 4.5319849368906E-05
 1.25042608268768E-05

 3.17238945582342E-04
 8.25281214573866E-04

 1.5106616456302E-05
 5.0017043307507E-05

 9.0639698737812E-05
 2.50085216537535E-05

 3.0213232912604E-05
 1.00034086615014E-04

 1.5106616456302E-05
 1.12538347441891E-04

 1.66172781019322E-04
 5.37683215555701E-04

 7.03968326863673E-03
 7.81516301679797E-03

 6.0426465825208E-05
 2.50085216537535E-05

 1.81279397475624E-04
 0

 6.0426465825208E-05
 1.50051129922521E-04

 2.41705863300832E-04
 0

 1.35959548106718E-04
 2.50085216537535E-04

 3.92772027863852E-04
 5.0017043307507E-05

 1.20852931650416E-04
 3.37615042325672E-04

 1.20852931650416E-04
 3.50119303152549E-04

 1.35959548106718E-04
 4.2514486811381E-04

 8.00650672184006E-04
 4.87666172248194E-04

 6.34477891164684E-04
 8.37785475400743E-04

 3.47452178494946E-04
 1.13788773524579E-03

 8.00650672184006E-04
 8.75298257881373E-04

 1.32938224815458E-03
 1.56303260335959E-03

 1.67683442664952E-03
 1.32545164764894E-03

 7.68926777625772E-03
 6.57724119493718E-03

 7.553308228151E-05
 1.25042608268768E-05

 3.0213232912604E-05
 7.50255649612605E-05

 2.2659924684453E-04
 1.25042608268768E-04

 2.2659924684453E-04
 1.37546869095644E-04

 1.20852931650416E-04
 2.37580955710658E-04

 2.87025712669738E-04
 1.50051129922521E-04

 3.17238945582342E-04
 1.25042608268768E-04

 2.11492630388228E-04
 2.75093738191289E-04

 2.56812479757134E-04
 3.50119303152549E-04

 4.38091877232758E-04
 2.25076694883782E-04

 2.71919096213436E-04
 3.75127824806303E-04

 3.7766541140755E-04
 3.37615042325672E-04

 3.47452178494946E-04
 4.12640607286933E-04

 6.49584507620986E-04
 5.12674693901947E-04

 8.61077138009214E-04
 4.6265765059444E-04

 5.2873157597057E-04
 7.50255649612605E-04

 9.0639698737812E-04
 5.50187476382577E-04

 1.55598149499911E-03
 1.46299851674458E-03

 6.7979774053359E-04
 6.62725823824468E-04

 9.0639698737812E-05
 2.50085216537535E-05

 3.32345562038644E-04
 2.37580955710658E-04

 2.56812479757134E-04
 4.00136346460056E-04

 8.00650672184006E-03
 1.36046357796419E-02

 1.5106616456302E-05
 1.37546869095644E-04

 9.0639698737812E-05
 2.50085216537535E-04

 1.5106616456302E-04
 4.50153389767563E-04

 1.66172781019322E-04
 5.62691737209454E-04

 1.35959548106718E-04
 6.00204519690084E-04

 1.90343367349405E-03
 2.62589477364412E-03

 5.54412823946283E-03
 8.97805927369751E-03

 1.02120727244602E-02
 1.81436824597982E-02

 2.41705863300832E-04
 0

 9.97036686115932E-03
 1.81436824597982E-02

 5.13624959514268E-04
 1.46299851674458E-03

 4.5319849368906E-05
 1.75059651576275E-04

 9.0639698737812E-05
 7.12742867131975E-04

 3.7766541140755E-04
 5.75195998036331E-04

 1.02724991902854E-03
 0

 4.07878644320154E-04
 0

 6.19371274708382E-04
 0

 1.19342270004786E-02
 1.06411259636721E-02

 8.45970521552912E-04
 3.87632085633179E-04

 .003051536524173
 2.41332233958721E-03

 8.03671995475266E-03
 7.84017153845173E-03

 3.07122044541556
 2.89543662002827

 .719150476402257
 .778515279081347

 2.56812479757134E-04
 1.25042608268768E-04

 3.0213232912604E-05
 0

 4.5319849368906E-05
 0

 1.5106616456302E-05
 3.75127824806303E-05

 4.5319849368906E-05
 2.50085216537535E-05

 6.0426465825208E-05
 2.50085216537535E-05

 6.0426465825208E-05
 3.75127824806303E-05

 .218758912903709
 .16583150708604

 1.5106616456302E-05
 1.25042608268768E-05

 3.0213232912604E-05
 0

 3.0213232912604E-05
 0

 0
 2.50085216537535E-05

 7.553308228151E-05
 5.0017043307507E-05

 6.0426465825208E-05
 6.25213041343838E-05

 1.5106616456302E-05
 1.37546869095644E-04

 9.0639698737812E-05
 7.50255649612605E-05

 4.5319849368906E-05
 1.62555390749398E-04

 1.96386013931926E-04
 1.87563912403151E-04

 2.11492630388228E-04
 3.00102259845042E-04

 3.47452178494946E-04
 3.37615042325672E-04

 5.58944808883174E-04
 4.00136346460056E-04

 5.2873157597057E-04
 5.75195998036331E-04

 7.10010973446194E-04
 4.6265765059444E-04

 6.64691124077288E-04
 5.50187476382577E-04

 8.00650672184006E-04
 6.87734345478222E-04

 7.40224206358798E-04
 7.62759910439482E-04

 5.13624959514268E-04
 1.0753664311114E-03

 6.94904356989892E-04
 1.03785364863077E-03

 1.43512856334869E-03
 9.2531530118888E-04

 1.40491533043609E-03
 9.87836605323264E-04

 1.88832705703775E-03
 6.25213041343838E-04

 1.23874254941676E-03
 1.20040903938017E-03

 1.32938224815458E-03
 1.32545164764894E-03

 1.31427563169827E-03
 1.52551982087896E-03

 2.2357792355327E-03
 1.47550277757146E-03

 2.55301818111504E-03
 1.65056242914773E-03

 2.73429757859066E-03
 1.83812634155088E-03

 4.09389305965784E-03
 2.1007158189153E-03

 3.82197396344441E-03
 2.8509714685279E-03

 6.61669800786027E-03
 2.25076694883782E-03

 5.72540763693846E-03
 4.53904668015626E-03

 7.31160236485017E-03
 5.52688328547953E-03

 8.38417213324761E-03
 4.9516872874432E-03

 .010604844752324
 9.2531530118888E-03

 1.48800172094575E-02
 1.02159810955583E-02

 1.82336860627565E-02
 1.72808884627437E-02

 2.28865239312975E-02
 1.55427962078078E-02

 .034065420108961
 3.15357458053832E-02

 5.96711350023929E-02
 4.13265820328277E-02

 1.37470209752348E-03
 1.21291330020705E-03

 0
 2.50085216537535E-05

 1.5106616456302E-04
 6.25213041343838E-05

 7.553308228151E-05
 1.87563912403151E-04

 3.7766541140755E-04
 3.75127824806303E-05

 2.2659924684453E-04
 3.50119303152549E-04

 5.43838192426872E-04
 5.50187476382577E-04

 .165931075156021
 .213347698228171

 1.5106616456302E-05
 1.25042608268768E-05

 3.0213232912604E-05
 1.25042608268768E-05

 1.5106616456302E-05
 2.50085216537535E-05

 4.5319849368906E-05
 0

 1.5106616456302E-04
 3.75127824806303E-05

 5.2873157597057E-04
 1.25042608268768E-05

 7.70437439271402E-04
 6.87734345478222E-04

 8.15757288640308E-04
 7.75264171266359E-04

 1.19342270004786E-03
 1.4004772126102E-03

 1.37470209752348E-03
 1.6130496466671E-03

 1.99407337223186E-03
 1.17540051772642E-03

 3.17238945582342E-03
 2.58838199116349E-03

 6.85840387116111E-03
 3.48868877069862E-03

 5.77072748630736E-03
 5.75195998036331E-03

 6.60159139140397E-03
 6.33966023922652E-03

 1.33240357144584E-02
 1.69307691595911E-02

 .123269990283424
 .172496278106765

 2.21009798755698E-02
 2.15573456655355E-02

 3.0213232912604E-05
 0

 6.0426465825208E-05
 2.50085216537535E-05

 4.22985260776456E-04
 1.25042608268768E-04

 2.41705863300832E-03
 .002638399034471

 1.91702962830472E-02
 .018768895501142

 3.0213232912604E-05
 2.50085216537535E-05

 3.0213232912604E-05
 2.50085216537535E-05

 .038068673469881
 4.40900236755674E-02

 0
 6.25213041343838E-05

 7.85544055727704E-04
 5.87700258863208E-04

 6.7979774053359E-04
 8.25281214573866E-04

 1.45023517980499E-03
 1.25042608268768E-03

 1.34448886461088E-03
 1.62555390749398E-03

 2.41705863300832E-03
 2.20074990553031E-03

 3.13915489961955E-02
 .037537791002284

 1.10278300131005E-03
 1.63805816832086E-03

 4.5319849368906E-05
 3.75127824806303E-05

 9.0639698737812E-05
 6.25213041343838E-05

 9.66823453203328E-04
 1.53802408170584E-03

 3.69205706192021E-02
 .051542563128386

 4.5319849368906E-05
 3.75127824806303E-05

 6.0426465825208E-05
 1.25042608268768E-04

 9.21503603834422E-04
 1.25042608268768E-05

 1.02724991902854E-03
 3.62623563979426E-04

 1.90343367349405E-03
 1.76310077658962E-03

 .032962637107651
 4.92417791362407E-02

 .233487863948604
 .277607094617491

 9.0639698737812E-05
 1.12538347441891E-04

 2.11492630388228E-04
 1.00034086615014E-04

 1.20852931650416E-04
 3.62623563979426E-04

 7.10010973446194E-04
 0

 8.45970521552912E-04
 1.25042608268768E-05

 5.2873157597057E-04
 3.62623563979426E-04

 8.45970521552912E-04
 3.25110781498796E-04

 1.26895578232937E-03
 6.37717302170715E-04

 6.34477891164684E-04
 1.32545164764894E-03

 2.43216524946462E-03
 4.6265765059444E-04

 1.82790059121254E-03
 1.25042608268768E-03

 2.43216524946462E-03
 1.20040903938017E-03

 3.36877546975535E-03
 1.43798999509083E-03

 2.20556600262009E-03
 3.35114190160297E-03

 9.62291468266437E-03
 5.1642597215001E-03

 8.64098461300474E-03
 7.13993293214663E-03

 1.39887268385356E-02
 1.65556413347848E-02

 1.85962448577078E-02
 2.35455231370089E-02

 2.38835606174135E-02
 .021207226362383

 4.24646988586649E-02
 .045452988105697

 4.58334743284203E-02
 .071937012537022

 5.29335840628822E-02
 7.56632822634313E-02

 1.11788961776635E-03
 1.53802408170584E-03

 5.43838192426872E-04
 5.87700258863208E-04

 5.74051425339476E-04
 9.50323822842634E-04

 1.16320946713525E-03
 1.43798999509083E-03

 4.5319849368906E-04
 1.87563912403151E-04

 1.5106616456302E-05
 1.25042608268768E-05

 7.553308228151E-05
 1.25042608268768E-05

 1.81279397475624E-04
 5.0017043307507E-05

 1.81279397475624E-04
 1.12538347441891E-04

 7.10010973446194E-04
 1.25042608268768E-03

 0
 2.50085216537535E-05

 3.0213232912604E-05
 2.50085216537535E-05

 1.81279397475624E-04
 2.37580955710658E-04

 4.98518343057966E-04
 9.6282808366951E-04

 6.67712447368548E-03
 7.11492441049288E-03

 4.18453275839565E-03
 3.50119303152549E-03

 1.5106616456302E-05
 1.25042608268768E-05

 1.57108811145541E-03
 1.48800703839833E-03

 2.59833803048394E-03
 2.00068173230028E-03

 5.13624959514268E-04
 2.43833086124097E-03

 1.66172781019322E-04
 3.75127824806303E-04

 3.47452178494946E-04
 2.06320303643467E-03

 1.97896675577556E-03
 1.17540051772642E-03

 1.97896675577556E-03
 1.17540051772642E-03

 .955523704094014
 .918237889560868

 5.00029004703596E-03
 5.60190885044079E-03

 1.5106616456302E-05
 1.25042608268768E-05

 3.7766541140755E-04
 1.62555390749398E-04

 4.60751801917211E-03
 5.42684919886451E-03

 1.68891971981456E-02
 1.68557435946299E-02

 7.553308228151E-05
 0

 4.5319849368906E-05
 7.50255649612605E-05

 2.41705863300832E-04
 1.12538347441891E-04

 2.87025712669738E-04
 1.12538347441891E-04

 6.34477891164684E-04
 8.75298257881373E-05

 6.0426465825208E-04
 9.87836605323264E-04

 9.66823453203328E-04
 2.66340755612475E-03

 1.40340466879046E-02
 1.28168673475487E-02

 1.05746315194114E-04
 3.75127824806303E-05

 1.05746315194114E-04
 3.75127824806303E-05

 5.68008778756955E-03
 1.01284512697702E-02

 1.5106616456302E-04
 0

 3.0213232912604E-04
 8.62793997054496E-04

 7.553308228151E-04
 1.55052834253272E-03

 9.0639698737812E-04
 2.40081807876034E-03

 1.26895578232937E-03
 2.42582660041409E-03

 2.2962057013579E-03
 2.88848425100853E-03

 5.50031905173956E-02
 6.43719347367615E-02

 1.96386013931926E-04
 1.25042608268768E-04

 1.42002194689239E-03
 8.37785475400743E-04

 2.55301818111504E-03
 9.00306779535127E-04

 1.32787158650895E-02
 1.51301556005209E-02

 3.75550485103668E-02
 .047378644273036

 .751795874564325
 .732899735584901

 3.32345562038644E-03
 2.4758436437216E-03

 2.02428660514447E-02
 2.51210600011954E-02

 4.08633975142969E-02
 5.43560218144333E-02

 6.87048916432615E-02
 7.98647139012619E-02

 .13378419533701
 .151589154004227

 .180644919584459
 .15437760416862

 .304232148813466
 .265115338051441

 .121049317664348
 8.83426027418843E-02

 .121049317664348
 8.83426027418843E-02

 .23088952591812
 .204632228431838

 3.92772027863852E-02
 4.83164638350518E-02

 1.5106616456302E-05
 2.50085216537535E-05

 4.5319849368906E-05
 0

 1.05746315194114E-04
 1.25042608268768E-05

 1.35959548106718E-04
 2.50085216537535E-05

 4.5319849368906E-05
 1.12538347441891E-04

 3.0213232912604E-05
 2.12572434056905E-04

 1.35959548106718E-04
 1.25042608268768E-04

 2.56812479757134E-04
 5.0017043307507E-05

 2.71919096213436E-04
 5.0017043307507E-05

 1.05746315194114E-04
 2.00068173230028E-04

 1.5106616456302E-05
 2.75093738191289E-04

 7.553308228151E-05
 2.37580955710658E-04

 4.5319849368906E-05
 2.75093738191289E-04

 1.35959548106718E-04
 2.12572434056905E-04

 3.0213232912604E-05
 3.12606520671919E-04

 1.96386013931926E-04
 1.75059651576275E-04

 1.35959548106718E-04
 3.12606520671919E-04

 3.7766541140755E-04
 1.25042608268768E-04

 2.11492630388228E-04
 2.75093738191289E-04

 1.81279397475624E-04
 3.00102259845042E-04

 1.5106616456302E-05
 4.75161911421317E-04

 2.87025712669738E-04
 2.50085216537535E-04

 1.20852931650416E-04
 4.2514486811381E-04

 1.5106616456302E-04
 4.37649128940687E-04

 1.35959548106718E-04
 4.75161911421317E-04

 2.41705863300832E-04
 4.12640607286933E-04

 2.87025712669738E-04
 3.87632085633179E-04

 6.0426465825208E-05
 5.75195998036331E-04

 7.85544055727704E-04
 8.75298257881373E-05

 1.5106616456302E-04
 6.25213041343838E-04

 3.92772027863852E-04
 4.50153389767563E-04

 3.0213232912604E-04
 5.62691737209454E-04

 3.32345562038644E-04
 5.75195998036331E-04

 3.7766541140755E-04
 5.50187476382577E-04

 9.97036686115932E-04
 1.00034086615014E-04

 8.15757288640308E-04
 3.12606520671919E-04

 3.62558794951248E-04
 9.37819562015757E-04

 1.61640796082431E-03
 0

 3.7766541140755E-04
 1.06286217028452E-03

 1.5408748785428E-03
 1.37546869095644E-04

 6.64691124077288E-04
 8.75298257881373E-04

 1.08767638485374E-03
 1.87563912403151E-03

 1.17831608359156E-03
 1.83812634155088E-03

 4.83411726601664E-04
 2.46333938289472E-03

 1.08767638485374E-03
 2.65090329529787E-03

 4.29027907358977E-03
 2.06320303643467E-03

 2.17535276970749E-03
 4.37649128940687E-03

 7.19074943319975E-03
 5.97703667524709E-03

 9.21503603834422E-03
 1.40422849085826E-02

 8.12735965349047E-03
 1.68057265513224E-02

 0
 5.0017043307507E-05

 3.0213232912604E-05
 7.50255649612605E-05

 7.553308228151E-05
 5.0017043307507E-05

 7.553308228151E-05
 5.0017043307507E-05

 6.0426465825208E-05
 1.00034086615014E-04

 4.5319849368906E-05
 1.50051129922521E-04

 3.0213232912604E-05
 1.75059651576275E-04

 1.5106616456302E-05
 2.12572434056905E-04

 1.20852931650416E-04
 1.37546869095644E-04

 7.553308228151E-05
 2.25076694883782E-04

 9.0639698737812E-05
 2.12572434056905E-04

 1.81279397475624E-04
 1.37546869095644E-04

 6.0426465825208E-05
 2.50085216537535E-04

 9.0639698737812E-05
 2.37580955710658E-04

 1.5106616456302E-04
 2.00068173230028E-04

 7.553308228151E-05
 2.87597999018165E-04

 1.05746315194114E-04
 2.75093738191289E-04

 1.35959548106718E-04
 2.62589477364412E-04

 1.5106616456302E-04
 2.50085216537535E-04

 1.5106616456302E-04
 2.75093738191289E-04

 1.96386013931926E-04
 2.37580955710658E-04

 9.0639698737812E-05
 3.25110781498796E-04

 1.5106616456302E-04
 3.12606520671919E-04

 1.5106616456302E-04
 3.37615042325672E-04

 1.05746315194114E-04
 3.75127824806303E-04

 1.81279397475624E-04
 3.50119303152549E-04

 1.20852931650416E-04
 4.50153389767563E-04

 2.2659924684453E-04
 3.87632085633179E-04

 2.2659924684453E-04
 4.2514486811381E-04

 1.81279397475624E-04
 4.6265765059444E-04

 1.66172781019322E-04
 5.0017043307507E-04

 2.56812479757134E-04
 4.2514486811381E-04

 3.92772027863852E-04
 3.87632085633179E-04

 1.96386013931926E-04
 6.87734345478222E-04

 3.17238945582342E-04
 8.00272692920113E-04

 3.92772027863852E-04
 8.5028973622762E-04

 4.98518343057966E-04
 9.75332344496387E-04

 5.89158041795778E-04
 1.00034086615014E-03

 6.0426465825208E-04
 1.18790477855329E-03

 6.64691124077288E-04
 1.21291330020705E-03

 6.94904356989892E-04
 1.50051129922521E-03

 1.81279397475624E-04
 1.50051129922521E-04

 4.5319849368906E-05
 5.0017043307507E-05

 1.35959548106718E-04
 1.00034086615014E-04

 .005302422376162
 1.51301556005209E-03

 1.66172781019322E-04
 5.0017043307507E-05

 0
 2.25076694883782E-04

 6.0426465825208E-04
 1.75059651576275E-04

 1.42002194689239E-03
 2.00068173230028E-04

 3.11196298999821E-03
 8.62793997054496E-04

 .178001261704606
 .137846971355489

 4.22985260776456E-04
 2.12572434056905E-04

 7.553308228151E-04
 2.62589477364412E-04

 2.09981968742598E-03
 2.06320303643467E-03

 .174723125933589
 .135308606407633

 .150129554342729
 .139960191435232

 2.59078472225579E-02
 2.40832063525646E-02

 3.0213232912604E-05
 1.25042608268768E-05

 0
 3.75127824806303E-05

 6.0426465825208E-05
 2.50085216537535E-05

 3.0213232912604E-05
 5.0017043307507E-05

 7.553308228151E-05
 1.50051129922521E-04

 3.0213232912604E-04
 2.37580955710658E-04

 2.50769833174613E-03
 1.96316894981965E-03

 2.32641893427051E-03
 4.48902963684876E-03

 2.05752116134833E-02
 1.71183330719943E-02

 9.51716836747026E-02
 9.82084645342901E-02

 1.35959548106718E-04
 2.75093738191289E-04

 .035515655288766
 4.60531926253871E-02

 5.95200688378299E-02
 5.18801781707117E-02

 4.22985260776456E-04
 1.37546869095644E-04

 4.22985260776456E-04
 1.37546869095644E-04

 2.86270381846923E-02
 1.75309736792812E-02

 5.43838192426872E-04
 3.12606520671919E-04

 6.19371274708382E-04
 4.6265765059444E-04

 7.553308228151E-04
 5.62691737209454E-04

 8.76183754465516E-04
 5.62691737209454E-04

 8.15757288640308E-04
 6.62725823824468E-04

 1.02724991902854E-03
 8.5028973622762E-04

 1.07256976839744E-03
 8.12776953746989E-04

 1.08767638485374E-03
 8.8780251870825E-04

 1.57108811145541E-03
 9.37819562015757E-04

 1.76747412538733E-03
 1.08787069193828E-03

 4.59241140271581E-03
 2.60088625199037E-03

 1.38980871397978E-02
 7.79015449514422E-03

 1.45023517980499E-02
 1.17790136989179E-02

 4.98518343057966E-03
 4.11390181204245E-03

 1.5106616456302E-05
 2.50085216537535E-05

 3.0213232912604E-05
 1.25042608268768E-05

 3.0213232912604E-05
 2.50085216537535E-05

 7.553308228151E-05
 0

 1.05746315194114E-04
 0

 6.0426465825208E-05
 7.50255649612605E-05

 7.553308228151E-05
 6.25213041343838E-05

 1.5106616456302E-04
 1.50051129922521E-04

 4.5319849368906E-05
 3.62623563979426E-04

 3.0213232912604E-04
 2.87597999018165E-04

 3.7766541140755E-04
 4.6265765059444E-04

 7.40224206358798E-04
 5.50187476382577E-04

 1.32938224815458E-03
 7.25247127958852E-04

 1.64662119373692E-03
 1.37546869095644E-03

 6.49584507620986E-03
 5.81448128449769E-03

 9.0639698737812E-05
 0

 9.0639698737812E-05
 3.75127824806303E-05

 1.05746315194114E-04
 2.12572434056905E-04

 9.0639698737812E-05
 3.37615042325672E-04

 6.0426465825208E-05
 3.62623563979426E-04

 2.71919096213436E-04
 2.12572434056905E-04

 2.56812479757134E-04
 2.25076694883782E-04

 9.21503603834422E-04
 3.00102259845042E-04

 8.3086390509661E-04
 6.37717302170715E-04

 1.69194104310582E-03
 2.75093738191289E-04

 5.58944808883174E-04
 1.47550277757146E-03

 1.5257682620865E-03
 1.73809225493587E-03

 3.0213232912604E-03
 1.85063060237776E-03

 3.0213232912604E-03
 1.85063060237776E-03

 .025212942865568
 2.88598339884316E-02

 3.88240042926961E-03
 2.37580955710658E-03

 6.0426465825208E-05
 0

 2.11492630388228E-04
 1.37546869095644E-04

 4.07878644320154E-04
 1.50051129922521E-04

 4.98518343057966E-04
 2.75093738191289E-04

 4.83411726601664E-04
 3.12606520671919E-04

 8.3086390509661E-04
 7.87768432093236E-04

 1.38980871397978E-03
 7.12742867131975E-04

 4.38091877232758E-04
 6.25213041343838E-05

 9.0639698737812E-05
 0

 1.5106616456302E-05
 6.25213041343838E-05

 3.32345562038644E-04
 0

 4.5319849368906E-04
 8.75298257881373E-05

 9.0639698737812E-05
 1.25042608268768E-05

 1.35959548106718E-04
 3.75127824806303E-05

 2.2659924684453E-04
 3.75127824806303E-05

 6.0426465825208E-05
 3.75127824806303E-05

 6.0426465825208E-05
 3.75127824806303E-05

 6.46563184329725E-03
 1.11913134400547E-02

 6.0426465825208E-05
 1.87563912403151E-04

 4.5319849368906E-05
 2.25076694883782E-04

 2.2659924684453E-04
 6.00204519690084E-04

 2.41705863300832E-04
 6.00204519690084E-04

 5.74051425339476E-04
 4.6265765059444E-04

 5.13624959514268E-04
 7.00238606305098E-04

 8.45970521552912E-04
 1.15039199607266E-03

 4.38091877232758E-04
 1.62555390749398E-03

 3.51984163431837E-03
 5.63942163292142E-03

 1.14810285067895E-03
 3.72626972640927E-03

 1.35959548106718E-04
 2.25076694883782E-04

 7.553308228151E-05
 4.75161911421317E-04

 1.5106616456302E-04
 4.12640607286933E-04

 1.81279397475624E-04
 7.37751388785729E-04

 2.71919096213436E-04
 8.62793997054496E-04

 3.32345562038644E-04
 1.01284512697702E-03

 3.17238945582342E-03
 1.81311781989713E-03

 1.35959548106718E-04
 2.62589477364412E-04

 6.34477891164684E-04
 3.75127824806303E-04

 8.45970521552912E-04
 4.2514486811381E-04

 1.55598149499911E-03
 7.50255649612605E-04

 3.55005486723097E-03
 3.45117598821799E-03

 1.96386013931926E-04
 3.00102259845042E-04

 2.56812479757134E-04
 2.50085216537535E-04

 4.07878644320154E-04
 2.50085216537535E-04

 4.98518343057966E-04
 2.62589477364412E-04

 4.22985260776456E-04
 4.2514486811381E-04

 3.92772027863852E-04
 5.75195998036331E-04

 1.37470209752348E-03
 1.38797295178332E-03

 6.0426465825208E-03
 6.11458354434273E-03

 4.38091877232758E-04
 1.37546869095644E-04

 3.92772027863852E-04
 6.12708780516961E-04

 6.7979774053359E-04
 6.12708780516961E-04

 7.553308228151E-04
 6.25213041343838E-04

 7.70437439271402E-04
 9.00306779535127E-04

 1.67683442664952E-03
 1.08787069193828E-03

 1.32938224815458E-03
 2.13822860139593E-03

 .211477523771772
 .203356793827497

 .211477523771772
 .203356793827497

 6.0426465825208E-05
 1.25042608268768E-05

 4.5319849368906E-05
 3.75127824806303E-05

 1.81279397475624E-04
 0

 1.5106616456302E-05
 1.62555390749398E-04

 4.5319849368906E-05
 2.75093738191289E-04

 6.0426465825208E-05
 7.50255649612605E-04

 2.87025712669738E-04
 7.75264171266359E-04

 9.36610220290724E-04
 5.0017043307507E-04

 7.70437439271402E-04
 1.15039199607266E-03

 7.553308228151E-04
 2.12572434056905E-03

 .208320240932405
 .197567321064653

 .589248681494516
 .430959349398307

 1.04235653548484E-03
 3.00102259845042E-04

 1.05746315194114E-04
 0

 2.2659924684453E-04
 3.75127824806303E-05

 3.62558794951248E-04
 6.25213041343838E-05

 3.47452178494946E-04
 2.00068173230028E-04

 7.25117589902496E-04
 6.50221562997591E-04

 9.0639698737812E-05
 3.75127824806303E-05

 9.0639698737812E-05
 6.25213041343838E-05

 1.35959548106718E-04
 5.0017043307507E-05

 1.5106616456302E-04
 7.50255649612605E-05

 2.56812479757134E-04
 4.2514486811381E-04

 4.33559892295867E-03
 2.13822860139593E-03

 5.13624959514268E-04
 3.50119303152549E-04

 6.7979774053359E-04
 2.62589477364412E-04

 6.94904356989892E-04
 3.37615042325672E-04

 1.01214330257223E-03
 5.12674693901947E-04

 1.43512856334869E-03
 6.75230084651345E-04

 2.84004389378478E-03
 1.46299851674458E-03

 5.13624959514268E-04
 7.50255649612605E-04

 1.05746315194114E-03
 3.00102259845042E-04

 1.26895578232937E-03
 4.12640607286933E-04

 3.06664314062931E-02
 2.06195261035198E-02

 7.85544055727704E-04
 5.50187476382577E-04

 5.69519440402585E-03
 3.12606520671919E-03

 5.21178267742419E-03
 3.86381659550492E-03

 6.39009876101574E-03
 5.73945571953643E-03

 1.25838115080996E-02
 7.34000110537666E-03

 2.92464094594007E-02
 1.90564935001602E-02

 8.15757288640308E-04
 7.37751388785729E-04

 3.65580118242508E-03
 2.33829677462595E-03

 6.0426465825208E-03
 3.27611633664171E-03

 9.12439633960641E-03
 5.92701963193958E-03

 9.60780806620807E-03
 6.7773093681672E-03

 .127061751013956
 9.10310188196628E-02

 1.01969661080038E-02
 7.50255649612605E-03

 2.42310127959084E-02
 1.74934608968006E-02

 .029261516075857
 2.03194238436747E-02

 3.00923799809536E-02
 2.06695431468273E-02

 3.32798760532333E-02
 2.50460344362341E-02

 8.97484083668902E-02
 7.46129243539736E-02

 1.19191203840223E-02
 9.89087031405952E-03

 1.34751018790214E-02
 1.23041926536467E-02

 1.74783552399414E-02
 1.57553686418647E-02

 2.34152555072681E-02
 1.81811952422788E-02

 .023460575356637
 1.84812975021239E-02

 .303582564305845
 .221087835680008

 4.29027907358977E-02
 3.00852515494655E-02

 4.35976950928876E-02
 3.38365297975285E-02

 5.39608339819107E-02
 4.49653219334488E-02

 7.48683911574327E-02
 4.92542833970675E-02

 8.82528533377163E-02
 6.29464490024976E-02

 8.08808245070409E-02
 9.44446820254002E-02

 6.11817966480231E-03
 1.65056242914773E-03

 1.05746315194114E-04
 0

 1.81279397475624E-04
 2.50085216537535E-05

 4.83411726601664E-04
 5.0017043307507E-05

 4.22985260776456E-04
 2.62589477364412E-04

 6.0426465825208E-04
 2.87597999018165E-04

 1.08767638485374E-03
 6.37717302170715E-04

 3.23281592164863E-03
 3.87632085633179E-04

 1.34599952625651E-02
 9.4157084026382E-03

 0
 8.75298257881373E-05

 7.10010973446194E-04
 4.37649128940687E-04

 4.26006584067716E-03
 3.36364616242985E-03

 8.48991844844172E-03
 5.52688328547953E-03

 6.13026495796735E-02
 8.33784111936142E-02

 3.47452178494946E-04
 5.87700258863208E-04

 9.21503603834422E-04
 7.37751388785729E-04

 9.51716836747026E-04
 7.62759910439482E-04

 8.15757288640308E-04
 9.37819562015757E-04

 5.64987455465695E-03
 5.21427676480761E-03

 5.26163451172999E-02
 7.51381033087024E-02

 7.40677404852487E-02
 5.73945571953643E-02

 .063780134678507
 4.81789169659562E-02

 7.553308228151E-05
 3.75127824806303E-05

 3.17238945582342E-04
 3.62623563979426E-04

 6.0426465825208E-04
 3.12606520671919E-04

 1.01214330257223E-03
 1.12538347441891E-03

 1.4955550291739E-03
 1.87563912403151E-03

 2.08471307096968E-03
 1.47550277757146E-03

 3.23281592164863E-03
 1.60054538584023E-03

 2.56812479757134E-03
 2.21325416635719E-03

 3.50473501786206E-03
 2.40081807876034E-03

 3.97304012800743E-03
 3.18858651085357E-03

 4.49119707245858E-02
 .033586444580991

 9.38120881936354E-03
 8.35284623235368E-03

 3.7766541140755E-04
 3.00102259845042E-04

 3.59537471659988E-03
 2.07570729726154E-03

 2.85515051024108E-03
 2.71342459943226E-03

 2.55301818111504E-03
 3.26361207581483E-03

 9.0639698737812E-04
 8.62793997054496E-04

 9.0639698737812E-04
 8.62793997054496E-04

 1.22967857954298E-02
 1.87438869794883E-02

 3.0213232912604E-03
 3.20109077168045E-03

 1.66172781019322E-04
 3.00102259845042E-04

 3.47452178494946E-04
 2.37580955710658E-04

 3.17238945582342E-04
 2.87597999018165E-04

 4.5319849368906E-04
 3.37615042325672E-04

 4.38091877232758E-04
 6.62725823824468E-04

 4.98518343057966E-04
 6.75230084651345E-04

 8.00650672184006E-04
 7.00238606305098E-04

 9.27546250416943E-03
 1.55427962078078E-02

 9.27546250416943E-03
 1.55427962078078E-02

 .564096165094773
 .620461422229625

 4.93835291956512E-02
 4.08514201214064E-02

 7.553308228151E-05
 1.37546869095644E-04

 0
 2.50085216537535E-05

 0
 3.75127824806303E-05

 3.0213232912604E-05
 2.50085216537535E-05

 4.5319849368906E-05
 5.0017043307507E-05

 2.11492630388228E-04
 6.75230084651345E-04

 1.5106616456302E-05
 2.50085216537535E-05

 4.5319849368906E-05
 1.25042608268768E-04

 1.5106616456302E-04
 5.25178954728824E-04

 4.90965034829815E-02
 4.00386431676594E-02

 2.41705863300832E-04
 2.50085216537535E-05

 1.96386013931926E-04
 2.00068173230028E-04

 3.32345562038644E-04
 1.50051129922521E-04

 3.32345562038644E-04
 3.75127824806303E-04

 1.02724991902854E-03
 8.62793997054496E-04

 3.47452178494946E-03
 2.72592886025913E-03

 1.40340466879046E-02
 1.52927109912703E-02

 2.94579020897889E-02
 2.04069536694629E-02

 8.87664782972305E-02
 8.74422959623492E-02

 4.5319849368906E-05
 1.25042608268768E-05

 3.0213232912604E-05
 0

 1.5106616456302E-05
 1.25042608268768E-05

 1.69949435133397E-02
 1.30794568249131E-02

 1.5106616456302E-05
 2.50085216537535E-05

 1.5106616456302E-05
 1.00034086615014E-04

 1.35959548106718E-04
 1.25042608268768E-04

 1.81279397475624E-04
 2.12572434056905E-04

 2.2659924684453E-04
 3.12606520671919E-04

 1.14810285067895E-03
 7.75264171266359E-04

 1.52727892373213E-02
 1.15289284823804E-02

 1.16169880548962E-02
 1.54427621211928E-02

 0
 5.0017043307507E-05

 6.0426465825208E-05
 1.12538347441891E-04

 7.553308228151E-05
 1.75059651576275E-04

 1.14810285067895E-02
 1.51051470788671E-02

 2.54697553453252E-02
 2.06195261035198E-02

 9.0639698737812E-05
 0

 1.5106616456302E-04
 0

 2.41705863300832E-04
 0

 3.17238945582342E-04
 1.50051129922521E-04

 4.07878644320154E-04
 1.12538347441891E-04

 4.98518343057966E-04
 3.12606520671919E-04

 .023762707685763
 2.00443301054834E-02

 2.47446377554227E-02
 3.07229688516362E-02

 4.07878644320154E-04
 1.12538347441891E-04

 1.12695358764013E-02
 1.21541415237242E-02

 1.30672232347012E-02
 1.84562889804701E-02

 9.89483377887781E-03
 7.56507780026044E-03

 8.61077138009214E-04
 4.37649128940687E-04

 1.64662119373692E-03
 1.41298147343707E-03

 7.38713544713168E-03
 5.71444719788268E-03

 6.46109985836036E-02
 6.79106405507677E-02

 3.26302915456123E-02
 3.31362911912234E-02

 0
 2.50085216537535E-05

 1.5106616456302E-05
 1.25042608268768E-05

 1.5106616456302E-05
 3.75127824806303E-05

 7.553308228151E-05
 2.50085216537535E-05

 1.66172781019322E-04
 1.00034086615014E-04

 3.17238945582342E-04
 2.87597999018165E-04

 4.81901064956034E-03
 3.87632085633179E-03

 2.72221228542562E-02
 2.87723041626434E-02

 3.0213232912604E-05
 0

 3.0213232912604E-05
 0

 3.17238945582342E-04
 0

 1.35959548106718E-04
 0

 1.81279397475624E-04
 0

 2.42007995629958E-02
 .030235302679388

 9.0639698737812E-05
 7.50255649612605E-05

 .024110159864258
 3.01602771144267E-02

 6.48073845975356E-03
 2.80095442522039E-03

 1.66172781019322E-04
 2.50085216537535E-05

 4.07878644320154E-04
 3.75127824806303E-05

 3.17238945582342E-04
 1.25042608268768E-04

 4.07878644320154E-04
 6.25213041343838E-05

 3.32345562038644E-04
 1.37546869095644E-04

 2.2659924684453E-04
 2.37580955710658E-04

 4.83411726601664E-04
 8.75298257881373E-05

 5.13624959514268E-04
 1.00034086615014E-04

 5.2873157597057E-04
 1.25042608268768E-04

 4.38091877232758E-04
 2.12572434056905E-04

 6.7979774053359E-04
 8.75298257881373E-05

 5.58944808883174E-04
 2.12572434056905E-04

 3.47452178494946E-04
 4.6265765059444E-04

 7.25117589902496E-04
 2.75093738191289E-04

 3.47452178494946E-04
 6.12708780516961E-04

 9.51716836747026E-04
 1.73809225493587E-03

 1.66172781019322E-04
 1.00034086615014E-04

 7.85544055727704E-04
 1.63805816832086E-03

 .361335159018287
 .424257065595102

 0
 2.50085216537535E-05

 0
 2.50085216537535E-05

 2.87025712669738E-04
 3.62623563979426E-04

 3.0213232912604E-05
 0

 3.0213232912604E-05
 0

 1.5106616456302E-05
 1.25042608268768E-05

 0
 5.0017043307507E-05

 1.66172781019322E-04
 3.75127824806303E-05

 4.5319849368906E-05
 2.62589477364412E-04

 1.96386013931926E-04
 7.50255649612605E-05

 1.5106616456302E-05
 1.25042608268768E-05

 3.0213232912604E-05
 0

 1.5106616456302E-05
 2.50085216537535E-05

 1.35959548106718E-04
 3.75127824806303E-05

 3.7917607305318E-03
 1.00034086615014E-03

 0
 3.75127824806303E-05

 1.81279397475624E-04
 0

 5.74051425339476E-04
 1.25042608268768E-05

 1.42002194689239E-03
 4.87666172248194E-04

 1.61640796082431E-03
 4.6265765059444E-04

 1.5257682620865E-03
 1.75059651576275E-03

 6.0426465825208E-05
 0

 8.91290370921818E-04
 4.87666172248194E-04

 5.74051425339476E-04
 1.26293034351455E-03

 3.0213232912604E-05
 2.50085216537535E-05

 3.0213232912604E-05
 2.50085216537535E-05

 7.70437439271402E-04
 3.75127824806303E-05

 3.0213232912604E-05
 3.75127824806303E-05

 3.0213232912604E-04
 0

 4.38091877232758E-04
 0

 3.0213232912604E-04
 2.50085216537535E-05

 1.35959548106718E-04
 1.25042608268768E-05

 1.66172781019322E-04
 1.25042608268768E-05

 1.17831608359156E-03
 5.50187476382577E-04

 1.05746315194114E-04
 7.50255649612605E-05

 1.96386013931926E-04
 7.50255649612605E-05

 8.76183754465516E-04
 4.00136346460056E-04

 .350307329005187
 .419255361264351

 2.41705863300832E-04
 1.12538347441891E-04

 .350065623141886
 .419142822916909

 1.81279397475624E-04
 2.87597999018165E-04

 1.81279397475624E-04
 2.87597999018165E-04

 9.21503603834422E-04
 2.75093738191289E-04

 3.47452178494946E-04
 2.25076694883782E-04

 5.74051425339476E-04
 5.0017043307507E-05

 1.84300720766884E-03
 5.87700258863208E-04

 1.84300720766884E-03
 5.87700258863208E-04

 4.94270362510454
 4.87588645831067

 4.51097163339988
 4.42308216524781

 6.34477891164684E-04
 8.8780251870825E-04

 0
 2.50085216537535E-05

 2.41705863300832E-04
 1.25042608268768E-04

 3.92772027863852E-04
 7.37751388785729E-04

 4.98518343057966E-04
 2.50085216537535E-05

 3.0213232912604E-05
 0

 4.5319849368906E-05
 1.25042608268768E-05

 1.05746315194114E-04
 0

 3.17238945582342E-04
 1.25042608268768E-05

 1.01667528750912E-02
 1.06661344853259E-02

 1.5106616456302E-05
 1.25042608268768E-05

 6.0426465825208E-05
 2.50085216537535E-05

 7.553308228151E-05
 8.75298257881373E-05

 3.32345562038644E-04
 1.87563912403151E-04

 4.38091877232758E-04
 1.37546869095644E-04

 9.24524927125682E-03
 1.02159810955583E-02

 1.44872451815936E-02
 .021507328622228

 0
 2.50085216537535E-05

 4.5319849368906E-05
 0

 0
 5.0017043307507E-05

 2.71919096213436E-04
 1.50051129922521E-04

 1.35959548106718E-04
 2.75093738191289E-04

 2.41705863300832E-04
 2.25076694883782E-04

 4.68305110145362E-04
 3.62623563979426E-04

 4.83411726601664E-04
 5.75195998036331E-04

 6.64691124077288E-04
 4.75161911421317E-04

 7.70437439271402E-04
 5.87700258863208E-04

 .011405495424508
 1.87813997619689E-02

 .339566524704756
 .331950612171097

 3.0213232912604E-05
 0

 7.553308228151E-05
 0

 7.553308228151E-05
 6.25213041343838E-05

 1.07256976839744E-03
 6.25213041343838E-04

 2.68897772922176E-03
 1.75059651576275E-04

 2.77961742795957E-03
 1.87563912403151E-03

 3.62558794951248E-03
 1.53802408170584E-03

 6.37499214455944E-03
 2.50085216537535E-05

 7.96118687247115E-03
 6.01454945772772E-03

 1.91249764336783E-02
 1.74434438534931E-02

 2.37173878363941E-02
 2.37956083535465E-02

 2.74336154846444E-02
 3.34488977118953E-02

 4.14072357067238E-02
 3.39740766666242E-02

 3.88844307585213E-02
 4.11140095987708E-02

 .038068673469881
 4.64158161893665E-02

 4.84922388247294E-02
 4.94918643527782E-02

 7.77537549005864E-02
 7.59508802624494E-02

 .471266006970797
 .545798480832344

 1.5106616456302E-05
 1.25042608268768E-05

 1.35959548106718E-04
 5.0017043307507E-05

 1.96386013931926E-04
 3.75127824806303E-05

 6.34477891164684E-04
 0

 1.20852931650416E-03
 1.08787069193828E-03

 3.17238945582342E-03
 2.01318599312716E-03

 2.37173878363941E-03
 4.37649128940687E-03

 4.75858418373513E-03
 3.27611633664171E-03

 6.0728598154334E-03
 3.96385068211993E-03

 5.3175289926183E-03
 5.73945571953643E-03

 1.01969661080038E-02
 7.40252240951104E-03

 1.13903888080517E-02
 1.07161515286334E-02

 1.12997491093139E-02
 1.44549255158695E-02

 1.61942928411557E-02
 1.05410918770571E-02

 1.09976167801879E-02
 1.53677365562315E-02

 1.69798368968834E-02
 1.40923019518901E-02

 2.37778143022193E-02
 .027897005904762

 3.28568907924568E-02
 4.39774853281256E-02

 4.19057540497817E-02
 5.93452218843571E-02

 5.74202491504039E-02
 9.37819562015757E-02

 7.69682108448587E-02
 8.69671340509278E-02

 .137394676670067
 .140697942824017

 .120686758869397
 .12695576017528

 3.0213232912604E-05
 0

 9.0639698737812E-05
 2.50085216537535E-05

 3.7766541140755E-04
 8.75298257881373E-05

 3.90959233889096E-02
 3.53370410967537E-02

 3.82801661002693E-02
 4.52779284541207E-02

 4.28121510371599E-02
 4.62282522769634E-02

 .011103363095382
 9.89087031405951E-03

 3.0213232912604E-05
 0

 1.66172781019322E-04
 1.37546869095644E-04

 9.0639698737812E-04
 7.87768432093236E-04

 7.40224206358798E-04
 9.87836605323264E-04

 1.91854028995035E-03
 2.06320303643467E-03

 4.47155847106539E-03
 1.33795590847581E-03

 2.87025712669738E-03
 4.57655946263689E-03

 4.68456176309925E-02
 2.89598680750466E-02

 3.0213232912604E-05
 1.25042608268768E-05

 8.00650672184006E-04
 2.87597999018165E-04

 1.61338663753305E-02
 9.20313596858129E-03

 2.98808873505653E-02
 1.94566298466202E-02

 1.71762229108154E-02
 1.81436824597982E-02

 6.0426465825208E-05
 0

 1.71157964449902E-02
 1.81436824597982E-02

 9.45372057835379E-02
 .108474462673156

 1.5106616456302E-05
 3.75127824806303E-05

 4.07878644320154E-04
 5.12674693901947E-04

 1.45023517980499E-03
 3.12606520671919E-04

 2.13305424362984E-02
 .02658405851794

 .071333442906658
 8.10276101581614E-02

 .124629585764491
 .120853680891764

 1.5106616456302E-05
 3.75127824806303E-05

 2.14513953679488E-03
 1.85063060237776E-03

 3.36877546975535E-03
 4.42650833271437E-03

 1.14357086574206E-02
 8.26531640656554E-03

 4.19057540497817E-02
 4.75537039246123E-02

 6.57591014342826E-02
 5.87200088430133E-02

 1.97896675577556E-03
 6.50221562997591E-04

 6.0426465825208E-05
 0

 0
 6.25213041343838E-05

 2.87025712669738E-04
 3.75127824806303E-05

 6.19371274708382E-04
 0

 6.49584507620986E-04
 1.25042608268768E-05

 3.62558794951248E-04
 5.37683215555701E-04

 5.74051425339476E-04
 6.25213041343838E-05

 9.0639698737812E-05
 0

 4.83411726601664E-04
 6.25213041343838E-05

 5.54412823946283E-03
 4.20143163783059E-03

 1.5106616456302E-05
 7.50255649612605E-05

 4.68305110145362E-04
 1.12538347441891E-04

 1.23874254941676E-03
 1.36296443012957E-03

 3.82197396344441E-03
 2.65090329529787E-03

 7.553308228151E-05
 3.75127824806303E-05

 7.553308228151E-05
 3.75127824806303E-05

 5.12265364033201E-02
 5.09923756520034E-02

 1.5106616456302E-05
 8.75298257881373E-05

 1.05746315194114E-04
 1.50051129922521E-04

 6.94904356989892E-04
 9.50323822842634E-04

 1.37470209752348E-03
 1.13788773524579E-03

 1.88832705703775E-03
 7.75264171266359E-04

 4.71477499601185E-02
 .047891318966938

 1.76143147880481E-02
 1.40422849085826E-02

 1.5106616456302E-05
 8.75298257881373E-05

 1.66172781019322E-04
 1.25042608268768E-05

 3.0213232912604E-04
 2.87597999018165E-04

 1.71309030614465E-02
 1.36546528229494E-02

 .012553598275187
 1.35296102146807E-02

 7.553308228151E-05
 5.0017043307507E-05

 5.3326356090746E-03
 5.45185772051827E-03

 7.14542958383084E-03
 8.02773545085488E-03

 6.95206489319018E-02
 6.99613393263755E-02

 4.5319849368906E-05
 8.75298257881373E-05

 2.80831999922654E-02
 2.54586750435211E-02

 4.13921290902675E-02
 4.44151344570662E-02

 4.22985260776456E-04
 0

 1.66172781019322E-04
 0

 2.56812479757134E-04
 0

 .433333293049023
 .405000503921711

 9.0639698737812E-05
 7.50255649612605E-05

 3.7766541140755E-04
 7.50255649612605E-05

 7.10010973446194E-04
 3.87632085633179E-04

 1.11788961776635E-03
 6.00204519690084E-04

 4.01835997737633E-03
 7.50255649612605E-04

 1.04990984371299E-02
 2.26327120966469E-03

 6.94904356989892E-03
 5.71444719788268E-03

 .029865780734109
 2.04694749735973E-02

 3.76910080584735E-02
 4.04262752532926E-02

 8.16512619463123E-02
 7.30373874897872E-02

 8.87513716807742E-02
 8.31158217162498E-02

 .171611162943591
 .178085682696379

 1.96990278590178E-02
 2.22325757501869E-02

 1.66172781019322E-04
 2.50085216537535E-05

 1.95328550779985E-02
 2.22075672285331E-02

 2.11492630388228E-04
 0

 2.11492630388228E-04
 0

 .140763452139822
 .118602913942926

 1.5106616456302E-04
 7.50255649612605E-05

 2.41705863300832E-04
 1.25042608268768E-04

 3.62558794951248E-04
 3.12606520671919E-04

 6.54116492557877E-03
 4.88916598330881E-03

 9.32078235353833E-03
 7.00238606305098E-03

 .12414617403789
 .106198687202664

 1.28406239878567E-03
 6.75230084651345E-04

 1.81279397475624E-04
 5.0017043307507E-05

 3.0213232912604E-04
 1.25042608268768E-05

 3.47452178494946E-04
 2.00068173230028E-04

 4.5319849368906E-04
 4.12640607286933E-04

 7.15449355370462E-02
 .066560180381465

 2.56812479757134E-04
 0

 1.4804484127176E-03
 1.01284512697702E-03

 6.98076746445715E-02
 .065547335254488

 .178665952828684
 .161880160664747

 1.20852931650416E-04
 1.25042608268768E-04

 7.70437439271402E-04
 0

 1.87322044058145E-03
 0

 2.13003292033858E-03
 0

 2.68897772922176E-03
 0

 4.35070553941498E-03
 2.48834790454847E-03

 6.63633660925347E-02
 6.10332970959855E-02

 .10036835973567
 9.82334730559438E-02

 .054972977284483
 5.69318995447699E-02

 2.41705863300832E-04
 3.75127824806303E-05

 1.16320946713525E-03
 1.12538347441891E-03

 9.8344073130526E-03
 5.17676398232698E-03

 8.27842581805349E-03
 8.02773545085488E-03

 1.03480322725669E-02
 8.66545275302559E-03

 2.51071965503739E-02
 3.38990511016629E-02

 8.00650672184006E-04
 1.11287921359203E-03

 2.2659924684453E-04
 8.75298257881373E-05

 1.20852931650416E-04
 2.12572434056905E-04

 4.5319849368906E-04
 8.12776953746989E-04

 .251766869860729
 .227577547049157

 1.96386013931926E-04
 1.87563912403151E-04

 2.71919096213436E-04
 2.62589477364412E-04

 5.18156944451159E-03
 3.10105668506544E-03

 .112468759517168
 .102735006953619

 .133648235788904
 .121291330020705

 2.41705863300832E-04
 2.12572434056905E-04

 2.41705863300832E-04
 2.12572434056905E-04

 .178091901403344
 .215223337352203

 3.32345562038644E-04
 1.62555390749398E-04

 1.45023517980499E-03
 4.12640607286933E-04

 1.42002194689239E-03
 7.37751388785729E-04

 1.96386013931926E-03
 5.62691737209454E-04

 2.43216524946462E-03
 1.38797295178332E-03

 2.65876449630915E-03
 2.71342459943226E-03

 3.68601441533769E-03
 1.97567321064653E-03

 1.48951238259138E-02
 1.72558799410899E-02

 1.42153260853802E-02
 1.89564594135452E-02

 3.47603244659509E-02
 4.61032096686946E-02

 4.94137424285638E-02
 6.17335357022906E-02

 5.08639776083688E-02
 6.32215427406889E-02

 .242627366904666
 .206357816425947

 2.11492630388228E-04
 2.75093738191289E-04

 2.68897772922176E-03
 8.25281214573866E-04

 8.85247724339297E-03
 5.66443015457517E-03

 1.09371903143626E-02
 4.91417450496257E-03

 1.83847522273195E-02
 1.49801044705984E-02

 2.03486123666388E-02
 2.24576524450707E-02

 .181203864393342
 .157241079897975

 3.83708057990071E-03
 3.30112485829546E-03

 3.7766541140755E-04
 1.87563912403151E-04

 4.38091877232758E-04
 5.50187476382577E-04

 3.0213232912604E-03
 2.56337346950974E-03

 .146624819324867
 .134133205889907

 4.5319849368906E-04
 1.87563912403151E-04

 2.05449983805707E-03
 1.72558799410899E-03

 2.71919096213436E-03
 2.53836494785598E-03

 1.38225540575163E-02
 1.36421485621225E-02

 .027161696388431
 2.61088966065187E-02

 .100413679585039
 8.99306438668976E-02

 3.20260268873602E-03
 2.71342459943226E-03

 4.98518343057966E-04
 2.12572434056905E-04

 1.96386013931926E-04
 9.37819562015757E-04

 2.50769833174613E-03
 1.56303260335959E-03

 1.02271793409165E-02
 7.4775479744723E-03

 3.47452178494946E-04
 3.50119303152549E-04

 4.22985260776456E-04
 6.25213041343838E-04

 3.32345562038644E-03
 2.32579251379908E-03

 6.13328628125861E-03
 4.17642311617684E-03

 8.68328313908239E-02
 7.86768091227086E-02

 4.07878644320154E-04
 3.12606520671919E-04

 7.10010973446194E-04
 5.50187476382577E-04

 1.79768735829994E-03
 6.75230084651345E-04

 1.18889071511097E-02
 8.11526527664302E-03

 1.83243257614943E-02
 7.6401033652217E-03

 .019306255831154
 2.37330870494121E-02

 3.43977656709996E-02
 3.76503293497259E-02

 8.99750076137347E-02
 7.83767068628635E-02

 3.92772027863852E-04
 3.37615042325672E-04

 4.36581215587128E-02
 .031960890673497

 4.59241140271581E-02
 4.60782011470409E-02

 6.51548367760305E-02
 8.96430458678795E-02

 8.61077138009214E-04
 3.75127824806303E-05

 5.54412823946283E-03
 9.86586179240576E-03

 2.18894872451816E-02
 3.08104986774243E-02

 3.68601441533769E-02
 4.89291726155688E-02

 .139826841919531
 .11886550342029

 5.13624959514268E-04
 3.37615042325672E-04

 7.50798837878209E-03
 3.65124416144801E-03

 6.8281906382485E-03
 4.47652537602188E-03

 1.95026418450859E-02
 1.00159129223283E-02

 1.83696456108632E-02
 1.82312122855863E-02

 2.80076669099839E-02
 .024758436437216

 5.90970835770534E-02
 5.73945571953643E-02

 6.94904356989892E-04
 2.25076694883782E-04

 6.94904356989892E-04
 2.25076694883782E-04

 .134448886461088
 .148963259230583

 6.0426465825208E-04
 3.50119303152549E-04

 2.03939322160077E-03
 1.0753664311114E-03

 2.61344464694025E-03
 2.1507328622228E-03

 1.41851128524676E-02
 5.93952389276646E-03

 9.41142205227614E-03
 1.32295079548356E-02

 1.14659218903332E-02
 1.66556754213998E-02

 1.81279397475624E-02
 1.78935972432606E-02

 .032615184929156
 3.66875012660564E-02

 4.33862024624993E-02
 5.49812348557771E-02

 .172970758424658
 .143173786467739

 3.92772027863852E-04
 5.37683215555701E-04

 1.04235653548484E-03
 9.12811040362003E-04

 8.55034491426693E-03
 4.15141459452308E-03

 1.25687048916433E-02
 .010040921443982

 2.24333254376085E-02
 1.41673275168514E-02

 .127983254617791
 .113363628656465

 3.89750704572591E-03
 3.90132937798555E-03

 5.43838192426872E-04
 4.37649128940687E-04

 1.28406239878567E-03
 1.57553686418647E-03

 2.06960645451337E-03
 1.88814338485839E-03

 6.19371274708382E-04
 4.12640607286933E-04

 6.19371274708382E-04
 4.12640607286933E-04

 .153588969511222
 .172958935757359

 1.31427563169827E-03
 0

 7.40224206358798E-04
 6.37717302170715E-04

 1.42002194689239E-02
 1.79686228082219E-02

 2.28563106983849E-02
 2.55086920868286E-02

 .114477939505857
 .128843903560138

 .241237558190687
 .223888790105228

 6.34477891164684E-04
 5.87700258863208E-04

 1.88832705703775E-03
 1.81311781989713E-03

 2.77508544302268E-02
 1.62555390749398E-02

 3.07419644885746E-02
 1.47925405581952E-02

 .038068673469881
 2.89723723358735E-02

 6.00488004138004E-02
 7.58758546974882E-02

 8.21044604400014E-02
 8.55916653599714E-02

 3.64069456596878E-03
 3.58872285731363E-03

 8.3086390509661E-04
 7.12742867131975E-04

 8.45970521552912E-04
 1.10037495276515E-03

 1.96386013931926E-03
 1.7756050374165E-03

 1.4804484127176E-03
 8.12776953746989E-04

 1.4804484127176E-03
 8.12776953746989E-04

 3.53192692748341E-02
 2.79970399913771E-02

 6.0426465825208E-03
 0

 2.92766226923133E-02
 2.79970399913771E-02

 4.74045624398757E-02
 3.29112144963396E-02

 2.58323141402764E-03
 3.02603112010418E-03

 1.00610065598971E-02
 4.61407224511752E-03

 .011556561589071
 9.00306779535127E-03

 2.32037628768799E-02
 1.62680433357667E-02

 1.80826198981935E-02
 1.33670548239313E-02

 5.75562086985106E-03
 1.9881774714734E-03

 1.23269990283424E-02
 1.13788773524579E-02

 5.85532453846265E-02
 6.29339447416707E-02

 4.09389305965784E-03
 6.85233493312846E-03

 5.44593523249687E-02
 5.60816098085423E-02

 5.10150437729318E-02
 5.22052889522105E-02

 1.57410943474667E-02
 1.84562889804701E-02

 3.52739494254652E-02
 3.37489999717404E-02

 2.42310127959084E-02
 2.95350640730829E-02

 2.42310127959084E-02
 2.95350640730829E-02

 3.29928503405636E-02
 3.30612656262621E-02

 3.29928503405636E-02
 3.30612656262621E-02

 .431731991704655
 .452804293062861

 .265831129781546
 .255186954954901

 0
 3.75127824806303E-05

 3.0213232912604E-05
 1.25042608268768E-05

 3.0213232912604E-05
 2.50085216537535E-05

 1.81279397475624E-04
 0

 3.17238945582342E-04
 0

 1.5106616456302E-04
 1.62555390749398E-04

 3.0213232912604E-04
 1.75059651576275E-04

 6.34477891164684E-04
 0

 7.40224206358798E-04
 0

 5.2873157597057E-04
 4.75161911421317E-04

 7.10010973446194E-04
 4.87666172248194E-04

 6.64691124077288E-04
 6.25213041343838E-04

 8.00650672184006E-04
 6.12708780516961E-04

 7.10010973446194E-04
 9.6282808366951E-04

 1.35959548106718E-03
 7.75264171266359E-04

 1.19342270004786E-03
 1.02534938780389E-03

 1.25384916587307E-03
 1.01284512697702E-03

 2.59833803048394E-03
 2.37580955710658E-04

 8.91290370921818E-04
 1.67557095080149E-03

 2.68897772922176E-03
 6.00204519690084E-04

 2.53791156465874E-03
 1.67557095080149E-03

 3.27813577101753E-03
 3.02603112010418E-03

 4.39602538878388E-03
 2.46333938289472E-03

 1.05746315194114E-02
 8.75298257881373E-05

 6.35988552810314E-03
 3.63873990062114E-03

 1.67683442664952E-02
 0

 1.08012307662559E-02
 7.01489032387786E-03

 4.37336546409943E-02
 6.27463808292676E-02

 .151594896138991
 .16563143891281

 7.553308228151E-05
 0

 7.553308228151E-05
 0

 3.28266775595442E-02
 4.81414041834755E-02

 7.553308228151E-05
 2.87597999018165E-04

 3.47452178494946E-04
 2.12572434056905E-04

 4.68305110145362E-04
 1.87563912403151E-04

 4.22985260776456E-04
 2.37580955710658E-04

 3.62558794951248E-04
 5.0017043307507E-04

 1.10278300131005E-03
 6.87734345478222E-04

 3.32345562038644E-03
 1.63805816832086E-03

 7.20585604965605E-03
 1.26918247392799E-02

 1.95177484615422E-02
 3.16983011961326E-02

 1.42757525512054E-02
 1.46549936890996E-02

 2.2659924684453E-04
 1.87563912403151E-04

 2.56812479757134E-04
 2.50085216537535E-04

 3.0213232912604E-04
 2.62589477364412E-04

 5.13624959514268E-04
 3.00102259845042E-04

 5.58944808883174E-04
 4.37649128940687E-04

 8.15757288640308E-04
 9.2531530118888E-04

 1.16018814384399E-02
 1.22916883928199E-02

 6.47620647481667E-02
 7.89769113825536E-02

 1.05746315194114E-04
 3.62623563979426E-04

 1.01214330257223E-03
 1.75059651576275E-04

 1.38980871397978E-03
 2.50085216537535E-04

 1.61640796082431E-03
 1.02534938780389E-03

 1.96386013931926E-03
 1.21291330020705E-03

 2.86572514176049E-02
 2.86472615543747E-02

 3.00168468986721E-02
 4.73036187080748E-02

 3.62558794951248E-04
 2.50085216537535E-04

 3.62558794951248E-04
 2.50085216537535E-04

 .012206146096692
 8.86552092625562E-03

 3.61048133305618E-03
 2.72592886025913E-03

 8.59566476363584E-03
 6.13959206599649E-03

 4.13921290902675E-02
 4.67284227100384E-02

 4.13921290902675E-02
 4.67284227100384E-02

 .630232931940463
 .557002298533225

 .630232931940463
 .557002298533225

 6.49584507620986E-03
 7.05240310635849E-03

 3.0213232912604E-05
 0

 2.87025712669738E-04
 1.62555390749398E-04

 5.58944808883174E-04
 3.12606520671919E-04

 5.58944808883174E-04
 3.87632085633179E-04

 6.19371274708382E-04
 6.87734345478222E-04

 7.10010973446194E-04
 7.37751388785729E-04

 1.31427563169827E-03
 9.50323822842634E-04

 1.10278300131005E-03
 1.27543460434143E-03

 1.31427563169827E-03
 2.53836494785598E-03

 7.5835214610636E-03
 5.60190885044079E-03

 3.0213232912604E-05
 0

 9.0639698737812E-05
 1.87563912403151E-04

 1.20852931650416E-04
 2.25076694883782E-04

 3.32345562038644E-04
 3.25110781498796E-04

 8.91290370921818E-04
 7.50255649612605E-05

 1.01214330257223E-03
 3.00102259845042E-04

 1.23874254941676E-03
 3.62623563979426E-04

 9.97036686115932E-04
 1.06286217028452E-03

 1.07256976839744E-03
 1.13788773524579E-03

 1.79768735829994E-03
 1.92565616733902E-03

 0
 2.50085216537535E-05

 0
 2.50085216537535E-05

 4.99877938539033E-02
 4.15141459452308E-02

 3.0213232912604E-05
 0

 4.5319849368906E-05
 0

 3.0213232912604E-05
 2.50085216537535E-05

 9.0639698737812E-05
 3.75127824806303E-05

 9.0639698737812E-05
 1.25042608268768E-04

 1.5106616456302E-04
 1.62555390749398E-04

 4.68305110145362E-04
 3.75127824806303E-05

 3.32345562038644E-04
 3.12606520671919E-04

 4.38091877232758E-04
 3.25110781498796E-04

 4.07878644320154E-04
 3.62623563979426E-04

 4.98518343057966E-04
 3.12606520671919E-04

 6.19371274708382E-04
 2.75093738191289E-04

 6.49584507620986E-04
 2.75093738191289E-04

 5.58944808883174E-04
 5.25178954728824E-04

 8.45970521552912E-04
 4.37649128940687E-04

 6.64691124077288E-04
 6.25213041343838E-04

 8.45970521552912E-04
 7.25247127958852E-04

 7.553308228151E-04
 8.00272692920113E-04

 1.57108811145541E-03
 5.50187476382577E-04

 1.45023517980499E-03
 1.56303260335959E-03

 1.66172781019322E-03
 1.65056242914773E-03

 2.2659924684453E-03
 1.15039199607266E-03

 2.76451081150327E-03
 9.75332344496387E-04

 1.99407337223186E-03
 1.70057947245524E-03

 2.09981968742598E-03
 1.82562208072401E-03

 2.41705863300832E-03
 2.07570729726154E-03

 2.77961742795957E-03
 2.13822860139593E-03

 2.37173878363941E-03
 2.61339051281724E-03

 3.47452178494946E-03
 1.70057947245524E-03

 2.96089682543519E-03
 3.16357798919982E-03

 5.25710252679309E-03
 4.60156798429065E-03

 3.68601441533769E-03
 5.98954093607397E-03

 5.71030102048216E-03
 4.45151685436813E-03

 9.0639698737812E-05
 0

 3.0213232912604E-05
 0

 6.0426465825208E-05
 0

 1.11637895612072E-02
 5.11424267819259E-03

 1.5106616456302E-05
 1.25042608268768E-05

 3.0213232912604E-05
 2.50085216537535E-05

 3.0213232912604E-05
 2.50085216537535E-05

 3.0213232912604E-05
 1.12538347441891E-04

 2.11492630388228E-04
 1.75059651576275E-04

 2.56812479757134E-04
 2.25076694883782E-04

 9.51716836747026E-04
 2.25076694883782E-04

 1.43512856334869E-03
 3.87632085633179E-04

 1.46534179626129E-03
 1.06286217028452E-03

 2.79472404441587E-03
 1.25042608268768E-05

 3.94282689509482E-03
 2.8509714685279E-03

 .155839855363211
 .174421934274104

 1.5106616456302E-05
 1.25042608268768E-05

 3.0213232912604E-04
 4.00136346460056E-04

 4.83411726601664E-04
 5.50187476382577E-04

 1.25384916587307E-03
 8.37785475400743E-04

 4.83411726601664E-03
 1.06286217028452E-03

 4.68305110145362E-03
 2.86347572935478E-03

 4.38242943397321E-02
 6.23837572652881E-02

 .100443892817952
 .106311225550106

 2.52280494820243E-03
 1.27543460434143E-03

 4.5319849368906E-05
 0

 1.81279397475624E-04
 0

 1.96386013931926E-04
 0

 1.66172781019322E-04
 7.50255649612605E-05

 6.7979774053359E-04
 1.62555390749398E-04

 6.0426465825208E-04
 3.87632085633179E-04

 6.49584507620986E-04
 6.50221562997591E-04

 .396548681977927
 .321997220552903

 1.96386013931926E-04
 7.50255649612605E-05

 1.82790059121254E-03
 9.87836605323264E-04

 3.08174975708561E-03
 1.2379218218608E-03

 4.87943711538555E-03
 3.10105668506544E-03

 5.93690026732668E-03
 3.73877398723615E-03

 8.73162431174255E-03
 1.29169014341637E-02

 2.06809579286774E-02
 1.91065105434677E-02

 6.46865316658851E-02
 4.78162934019767E-02

 9.40991139063051E-02
 .085229041795992

 .192428080420375
 .147787858712856

 3.09407675611395
 2.83452836554056

 .140008121317007
 .131544823898744

 8.43704529084466E-02
 7.86643048618817E-02

 3.0213232912604E-05
 0

 7.553308228151E-05
 0

 1.81279397475624E-04
 1.50051129922521E-04

 2.11492630388228E-04
 1.87563912403151E-04

 2.41705863300832E-04
 2.25076694883782E-04

 2.71919096213436E-04
 2.00068173230028E-04

 3.32345562038644E-04
 2.25076694883782E-04

 7.40224206358798E-04
 4.37649128940687E-04

 1.67683442664952E-03
 5.75195998036331E-04

 2.34152555072681E-03
 6.62725823824468E-04

 7.90076040664594E-03
 5.7019429370558E-03

 9.12439633960641E-03
 6.42719006501465E-03

 8.92801032567448E-03
 8.16528231995052E-03

 1.91702962830472E-02
 2.53086239135986E-02

 3.31439165051266E-02
 3.03978580701374E-02

 1.66625979513011E-02
 1.10912793534397E-02

 2.71919096213436E-04
 6.25213041343838E-05

 4.98518343057966E-04
 3.12606520671919E-04

 1.63151457728062E-03
 7.87768432093236E-04

 1.69194104310582E-03
 1.35046016930269E-03

 1.25687048916433E-02
 8.57792292723746E-03

 3.89750704572592E-02
 4.17892396834221E-02

 3.89750704572592E-02
 4.17892396834221E-02

 1.01916787922441
 .93364313889958

 .704572591521925
 .668340236935736

 3.0213232912604E-05
 3.75127824806303E-05

 4.77369080019143E-03
 2.32579251379908E-03

 4.89454373184185E-03
 4.72661059255941E-03

 7.52309499523839E-03
 3.52620155317925E-03

 7.81012070790813E-03
 4.7891318966938E-03

 1.59525869778549E-02
 .016330564639901

 1.55144951006222E-02
 1.69182648987643E-02

 1.90041235020279E-02
 1.50551300355596E-02

 2.33548290414429E-02
 1.17790136989179E-02

 2.32641893427051E-02
 2.23826268801094E-02

 .084838758018592
 8.19529254593503E-02

 9.59723343468866E-02
 .101534597914239

 .107800815032171
 .105723525291243

 9.39027278923732E-02
 .120816168109283

 .199936068799157
 .160442170669656

 6.14839289771491E-03
 4.26395294196497E-03

 6.0426465825208E-05
 2.50085216537535E-05

 2.2659924684453E-04
 6.25213041343838E-05

 1.96386013931926E-04
 1.37546869095644E-04

 3.0213232912604E-04
 2.12572434056905E-04

 4.22985260776456E-04
 2.25076694883782E-04

 6.34477891164684E-04
 1.12538347441891E-04

 5.43838192426872E-04
 2.00068173230028E-04

 3.62558794951248E-04
 4.2514486811381E-04

 5.13624959514268E-04
 4.2514486811381E-04

 5.74051425339476E-04
 4.00136346460056E-04

 6.19371274708382E-04
 3.87632085633179E-04

 7.25117589902496E-04
 8.8780251870825E-04

 9.66823453203328E-04
 7.62759910439482E-04

 .295968829611869
 .251848317314125

 2.11492630388228E-04
 2.12572434056905E-04

 3.17238945582342E-04
 2.25076694883782E-04

 1.14810285067895E-03
 5.37683215555701E-04

 1.42757525512054E-02
 9.16562318610066E-03

 2.29167371642101E-02
 2.53461366960792E-02

 3.04700453923611E-02
 2.24576524450707E-02

 3.79931403875995E-02
 3.33238551036266E-02

 6.23903259645272E-02
 4.40900236755674E-02

 5.54866022439972E-02
 5.45310814660095E-02

 7.07593914813186E-02
 6.19586123971743E-02

 1.24780651929054E-02
 9.19063170775442E-03

 2.03939322160077E-03
 1.42548573426395E-03

 3.09685637354191E-03
 2.71342459943226E-03

 7.34181559776277E-03
 5.05172137405821E-03

 1.46439007942455
 1.34833444496212

 .618132532158965
 .58288611844486

 4.5319849368906E-05
 3.75127824806303E-05

 1.5106616456302E-05
 7.50255649612605E-05

 3.7766541140755E-04
 3.25110781498796E-04

 6.64691124077288E-04
 5.37683215555701E-04

 9.66823453203328E-04
 1.10037495276515E-03

 1.93364690640666E-03
 1.72558799410899E-03

 2.17535276970749E-03
 1.56303260335959E-03

 1.70704765956213E-03
 2.20074990553031E-03

 5.81604733567627E-03
 7.87768432093236E-04

 5.19667606096789E-03
 3.36364616242985E-03

 6.64691124077288E-03
 5.75195998036331E-03

 9.48695513455765E-03
 9.0405805778319E-03

 2.17384210806186E-02
 1.66931882038805E-02

 2.14211821350362E-02
 2.77469547748395E-02

 2.22218328072202E-02
 2.84221848594909E-02

 3.17843210240594E-02
 2.80970740779921E-02

 .035515655288766
 2.50710429578879E-02

 3.10743100506132E-02
 3.21984716292077E-02

 4.70873234942933E-02
 5.46561240742783E-02

 5.94143225226358E-02
 6.11333311826005E-02

 .148951238259138
 .145737159937249

 .16389168193442
 .136621553794455

 .135143790818078
 .112113202573777

 1.66172781019322E-04
 2.37580955710658E-04

 1.22363593296046E-03
 7.87768432093236E-04

 3.44430855203686E-03
 2.65090329529787E-03

 3.58026810014357E-03
 2.56337346950974E-03

 5.63476793820064E-03
 5.81448128449769E-03

 6.51095169266616E-03
 6.83983067230159E-03

 1.17982674523719E-02
 8.91553796956313E-03

 .014608098113244
 1.17790136989179E-02

 4.84015991259916E-02
 3.06354390258481E-02

 3.97757211294432E-02
 4.18892737700371E-02

 .692154952794845
 .634928852006321

 3.32345562038644E-04
 2.62589477364412E-04

 2.2659924684453E-04
 3.75127824806303E-04

 1.82790059121254E-03
 2.50085216537535E-05

 1.61640796082431E-03
 8.62793997054496E-04

 3.73133426470659E-03
 0

 6.67712447368548E-03
 4.82664467917443E-03

 1.31125430840701E-02
 1.51301556005209E-03

 2.97449278024586E-02
 2.66465798220744E-02

 2.81436264580906E-02
 2.88723382492584E-02

 2.59078472225579E-02
 3.39990851882779E-02

 3.31892363544955E-02
 3.23360184983033E-02

 5.87949512479274E-02
 .059257692058569

 7.12125899750076E-02
 7.01363989779517E-02

 7.80558872297124E-02
 7.00738776738174E-02

 .114810285067895
 .102022264086487

 .224771346253317
 .203719417391476

 1.61187597588742E-02
 1.67307009863611E-02

 3.47452178494946E-04
 3.50119303152549E-04

 1.57713075803793E-02
 1.63805816832086E-02

 1.16320946713525E-03
 4.75161911421317E-04

 1.16320946713525E-03
 4.75161911421317E-04

 1.67683442664952E-03
 1.20040903938017E-03

 1.67683442664952E-03
 1.20040903938017E-03

 .470510676147982
 .421005957780114

 1.96386013931926E-04
 2.00068173230028E-04

 4.5319849368906E-05
 6.25213041343838E-05

 1.5106616456302E-04
 1.37546869095644E-04

 3.0213232912604E-05
 3.12606520671919E-04

 3.0213232912604E-05
 3.12606520671919E-04

 5.69519440402585E-03
 5.23928528646136E-03

 3.62558794951248E-04
 6.25213041343838E-05

 3.32345562038644E-04
 3.50119303152549E-04

 5.74051425339476E-04
 5.50187476382577E-04

 6.94904356989892E-04
 5.87700258863208E-04

 8.45970521552912E-04
 5.25178954728824E-04

 6.0426465825208E-04
 7.50255649612605E-04

 9.66823453203328E-04
 9.87836605323264E-04

 1.31427563169827E-03
 1.42548573426395E-03

 .43550864581873
 .398835903334061

 2.87025712669738E-04
 2.62589477364412E-04

 7.40224206358798E-04
 7.87768432093236E-04

 1.84300720766884E-03
 1.62555390749398E-03

 3.32345562038644E-03
 2.6884160777785E-03

 1.96537080096489E-02
 1.50926428180402E-02

 3.82499528673567E-02
 3.41116235357198E-02

 .110671072158868
 .120240972111247

 .260740200035772
 .224026336974324

 2.90802366783813E-02
 1.64180944656892E-02

 4.5319849368906E-04
 2.62589477364412E-04

 4.68305110145362E-04
 3.25110781498796E-04

 8.00650672184006E-04
 5.0017043307507E-04

 3.33856223684274E-03
 3.00102259845042E-03

 2.40195201655202E-02
 1.23292011753005E-02

 4.01002112509245
 4.44712785881789

 9.0639698737812E-05
 6.25213041343838E-05

 9.0639698737812E-05
 6.25213041343838E-05

 0
 2.50085216537535E-05

 1.5106616456302E-05
 3.75127824806303E-05

 7.553308228151E-05
 0

 3.47452178494946E-04
 7.12742867131975E-04

 3.47452178494946E-04
 7.12742867131975E-04

 0
 2.50085216537535E-05

 0
 1.00034086615014E-04

 3.47452178494946E-04
 5.87700258863208E-04

 .178454460198295
 .163755799788778

 .158332447078501
 .13392063345585

 3.0213232912604E-05
 0

 1.20852931650416E-04
 1.00034086615014E-04

 1.35959548106718E-04
 1.50051129922521E-04

 3.62558794951248E-04
 3.00102259845042E-04

 5.43838192426872E-04
 5.87700258863208E-04

 .157139024378453
 .132782745720604

 8.00650672184006E-04
 9.50323822842634E-04

 0
 2.50085216537535E-05

 4.5319849368906E-05
 0

 7.553308228151E-04
 9.2531530118888E-04

 1.93213624476103E-02
 2.88848425100853E-02

 7.40224206358798E-04
 4.2514486811381E-04

 8.21799935222829E-03
 1.13413645699772E-02

 1.03631388890232E-02
 1.71183330719943E-02

 2.47253012863586
 2.58468072995873

 6.57137815849137E-03
 6.68977954237907E-03

 1.5106616456302E-05
 1.25042608268768E-05

 1.5106616456302E-05
 1.25042608268768E-05

 3.0213232912604E-05
 0

 3.0213232912604E-05
 0

 1.5106616456302E-05
 2.50085216537535E-05

 3.0213232912604E-05
 1.25042608268768E-05

 4.5319849368906E-05
 1.00034086615014E-04

 1.20852931650416E-04
 8.75298257881373E-05

 1.35959548106718E-04
 2.37580955710658E-04

 3.32345562038644E-04
 1.37546869095644E-04

 3.0213232912604E-04
 2.00068173230028E-04

 3.32345562038644E-04
 1.87563912403151E-04

 3.47452178494946E-04
 3.00102259845042E-04

 3.47452178494946E-04
 1.15039199607266E-03

 1.19342270004786E-03
 8.12776953746989E-04

 1.82790059121254E-03
 1.35046016930269E-03

 1.45023517980499E-03
 2.06320303643467E-03

 9.0639698737812E-04
 5.87700258863208E-04

 0
 2.50085216537535E-05

 1.5106616456302E-05
 3.75127824806303E-05

 7.553308228151E-05
 1.25042608268768E-05

 1.35959548106718E-04
 2.50085216537535E-05

 1.20852931650416E-04
 5.0017043307507E-05

 2.56812479757134E-04
 0

 1.5106616456302E-05
 3.00102259845042E-04

 2.87025712669738E-04
 1.37546869095644E-04

 .333448345039954
 .367150106398755

 3.0213232912604E-05
 0

 1.66172781019322E-04
 7.50255649612605E-05

 9.0639698737812E-05
 3.25110781498796E-04

 2.87025712669738E-04
 2.12572434056905E-04

 3.62558794951248E-04
 3.62623563979426E-04

 1.02724991902854E-03
 4.87666172248194E-04

 4.5319849368906E-05
 1.55052834253272E-03

 3.47452178494946E-04
 7.25247127958852E-03

 1.99709469552312E-02
 2.06695431468273E-02

 2.53186891807621E-02
 2.29703271389726E-02

 5.67102381769577E-02
 5.45185772051827E-02

 .22909183855982
 .258725660768907

 .253579663835485
 .225726916446779

 3.0213232912604E-05
 0

 1.5106616456302E-05
 3.75127824806303E-05

 6.0426465825208E-05
 2.50085216537535E-05

 9.0639698737812E-05
 0

 1.35959548106718E-04
 1.25042608268768E-05

 6.0426465825208E-05
 1.00034086615014E-04

 1.35959548106718E-04
 1.12538347441891E-04

 2.87025712669738E-04
 1.37546869095644E-04

 3.7766541140755E-04
 2.37580955710658E-04

 8.3086390509661E-04
 6.62725823824468E-04

 5.74051425339476E-04
 1.05035790945765E-03

 8.3086390509661E-03
 3.95134642129306E-03

 6.58648477494767E-03
 5.72695145870956E-03

 9.0639698737812E-03
 5.25178954728824E-03

 1.07256976839744E-02
 .011566441264861

 1.43814988663995E-02
 1.09912452668247E-02

 2.01673329691632E-02
 2.24576524450707E-02

 2.88838506644494E-02
 2.80720655563383E-02

 3.82499528673567E-02
 2.70092033860538E-02

 .114613899053963
 .108324411543233

 .115837534986924
 9.83585156642126E-02

 1.5106616456302E-05
 1.25042608268768E-05

 3.0213232912604E-05
 0

 1.5106616456302E-05
 1.25042608268768E-05

 6.0426465825208E-05
 5.0017043307507E-05

 1.66172781019322E-04
 1.25042608268768E-05

 3.0213232912604E-05
 3.37615042325672E-04

 1.35959548106718E-03
 2.75093738191289E-04

 .114160700560274
 9.76582770579075E-02

 6.0426465825208E-04
 2.87597999018165E-04

 3.0213232912604E-05
 1.25042608268768E-05

 1.66172781019322E-04
 2.37580955710658E-04

 4.07878644320154E-04
 3.75127824806303E-05

 9.0639698737812E-05
 7.12742867131975E-04

 1.5106616456302E-05
 3.75127824806303E-05

 4.5319849368906E-05
 1.00034086615014E-04

 3.0213232912604E-05
 5.75195998036331E-04

 1.16774145207214E-02
 9.49073396759946E-03

 0
 5.0017043307507E-05

 9.0639698737812E-05
 2.50085216537535E-05

 1.05746315194114E-04
 2.50085216537535E-05

 1.96386013931926E-04
 1.00034086615014E-04

 2.94579020897889E-03
 1.48800703839833E-03

 8.3388522838787E-03
 7.8026587559711E-03

 2.43216524946462E-03
 2.05069877560779E-03

 0
 6.25213041343838E-05

 6.0426465825208E-05
 1.75059651576275E-04

 2.37173878363941E-03
 1.81311781989713E-03

 2.67991375934797E-02
 2.44333256557172E-02

 1.5106616456302E-05
 5.0017043307507E-05

 0
 2.37580955710658E-04

 3.47452178494946E-04
 4.50153389767563E-04

 1.69194104310582E-03
 2.37580955710658E-03

 3.35366885329904E-03
 1.83812634155088E-03

 3.61048133305618E-03
 1.92565616733902E-03

 1.77804875690675E-02
 .017555982200935

 4.88094777703117E-02
 4.08514201214064E-02

 4.5319849368906E-05
 3.75127824806303E-05

 0
 1.37546869095644E-04

 6.0426465825208E-05
 1.87563912403151E-04

 2.87025712669738E-04
 0

 5.2873157597057E-04
 1.50051129922521E-04

 2.71919096213436E-04
 9.87836605323264E-04

 3.62558794951248E-04
 9.50323822842634E-04

 1.22363593296046E-03
 8.12776953746989E-04

 5.12114297868638E-03
 3.31362911912234E-03

 6.96415018635522E-03
 5.27679806894199E-03

 1.51217230727583E-02
 1.00534257048089E-02

 1.88228441045523E-02
 1.89439551527183E-02

 .12757537597347
 .128143664953833

 6.0426465825208E-05
 5.0017043307507E-05

 1.66172781019322E-04
 3.75127824806303E-05

 1.05746315194114E-04
 1.12538347441891E-04

 3.0213232912604E-04
 1.50051129922521E-04

 2.87025712669738E-04
 2.00068173230028E-04

 6.94904356989892E-04
 4.6265765059444E-04

 1.99407337223186E-03
 2.1007158189153E-03

 6.19371274708382E-04
 3.73877398723615E-03

 9.0790764902375E-03
 9.60327231504135E-03

 1.50461899904768E-02
 1.71558458544749E-02

 1.55900281829037E-02
 1.74059310710124E-02

 8.36302287020879E-02
 7.71262807801758E-02

 7.25117589902496E-04
 7.75264171266359E-04

 1.05746315194114E-04
 2.50085216537535E-05

 6.19371274708382E-04
 7.50255649612605E-04

 6.12573297303046E-02
 5.91201451894733E-02

 1.20852931650416E-04
 1.25042608268768E-05

 9.0639698737812E-05
 7.50255649612605E-05

 4.07878644320154E-04
 5.0017043307507E-05

 1.20852931650416E-04
 4.6265765059444E-04

 6.94904356989892E-04
 6.75230084651345E-04

 6.69223109014178E-03
 7.06490736718537E-03

 1.35808481942155E-02
 1.24667480443961E-02

 1.44117120993121E-02
 1.37296783879107E-02

 2.51374097832865E-02
 2.45833767856397E-02

 1.51821495385835E-02
 1.17164923947835E-02

 1.5106616456302E-05
 1.50051129922521E-04

 1.05746315194114E-04
 7.50255649612605E-05

 2.56812479757134E-04
 1.75059651576275E-04

 8.91290370921818E-04
 5.0017043307507E-04

 9.97036686115932E-04
 4.37649128940687E-04

 8.00650672184006E-04
 6.62725823824468E-04

 2.73429757859066E-03
 7.37751388785729E-04

 3.70112103179399E-03
 2.8509714685279E-03

 5.68008778756955E-03
 6.12708780516961E-03

 1.41899469697336
 1.53419777789282

 2.41705863300832E-04
 3.75127824806303E-05

 1.66172781019322E-04
 2.00068173230028E-04

 3.62558794951248E-04
 1.50051129922521E-04

 2.2659924684453E-04
 3.37615042325672E-04

 4.5319849368906E-04
 3.00102259845042E-04

 3.62558794951248E-04
 1.53802408170584E-03

 1.74481420070288E-02
 1.45674638633114E-02

 .11474985860207
 .127143324087683

 .240376481052677
 .211659623016543

 1.04460742133683
 1.17826399345577

 2.42914392617336E-02
 1.96566980198503E-02

 4.38091877232758E-04
 1.37546869095644E-04

 3.17238945582342E-04
 4.00136346460056E-04

 1.94875352286296E-03
 1.16289625689954E-03

 4.12410629257045E-03
 2.28827973131845E-03

 1.74632486234851E-02
 1.56678388160766E-02

 2.50769833174613E-03
 1.22541756103392E-03

 8.91290370921818E-04
 3.37615042325672E-04

 1.61640796082431E-03
 8.8780251870825E-04

 2.03335057501825E-02
 3.15732585878638E-02

 1.25384916587307E-03
 4.2514486811381E-04

 5.06071651286117E-03
 6.65226675989844E-03

 1.40189400714483E-02
 2.44958469598516E-02

 9.0639698737812E-04
 2.19324734903418E-02

 1.05746315194114E-04
 2.01318599312716E-03

 2.56812479757134E-04
 2.83846720770102E-03

 1.05746315194114E-04
 4.12640607286933E-03

 1.5106616456302E-04
 5.72695145870956E-03

 2.87025712669738E-04
 7.22746275793477E-03

 9.36610220290724E-03
 6.21461763095775E-03

 7.79501409145183E-03
 2.72592886025913E-03

 3.0213232912604E-05
 0

 6.49584507620986E-04
 2.00068173230028E-04

 7.10010973446194E-04
 6.00204519690084E-04

 1.16320946713525E-03
 3.00102259845042E-04

 2.02428660514447E-03
 2.62589477364412E-04

 3.21770930519233E-03
 1.36296443012957E-03

 1.57108811145541E-03
 3.48868877069862E-03

 3.0213232912604E-05
 5.0017043307507E-05

 1.5106616456302E-05
 1.25042608268768E-04

 1.96386013931926E-04
 2.50085216537535E-05

 0
 2.75093738191289E-04

 4.5319849368906E-05
 2.50085216537535E-04

 1.20852931650416E-04
 3.12606520671919E-04

 1.81279397475624E-04
 5.62691737209454E-04

 1.5106616456302E-04
 9.6282808366951E-04

 8.3086390509661E-04
 9.2531530118888E-04

 1.34923234217816
 1.69170144726816

 .123013177803667
 9.98465227026109E-02

 3.0213232912604E-05
 3.75127824806303E-05

 2.87025712669738E-04
 1.25042608268768E-05

 2.56812479757134E-04
 3.75127824806303E-05

 5.13624959514268E-04
 5.75195998036331E-04

 7.10010973446194E-04
 5.25178954728824E-04

 3.50473501786206E-03
 5.25178954728824E-04

 1.38376606739726E-02
 .010040921443982

 3.29928503405636E-02
 2.45708725248128E-02

 .070880244412969
 6.35216450005339E-02

 1.60130134436801E-03
 2.4758436437216E-03

 2.71919096213436E-04
 2.50085216537535E-05

 7.85544055727704E-04
 1.08787069193828E-03

 5.43838192426872E-04
 1.36296443012957E-03

 1.22461786303012
 1.58937908092182

 7.62884131043251E-03
 9.21564022940817E-03

 9.62291468266437E-03
 1.06036131811915E-02

 1.39887268385356E-02
 1.62430348141129E-02

 1.79164471171742E-02
 2.32704293988176E-02

 2.04543586818329E-02
 2.75593908624364E-02

 2.78717073618772E-02
 3.41366320573736E-02

 2.90047035960998E-02
 3.94509429087962E-02

 3.73133426470659E-02
 4.28270933320529E-02

 3.71320632495903E-02
 4.90667194846644E-02

 4.95799152095832E-02
 4.89166683547419E-02

 4.26761914890531E-02
 5.54188839847178E-02

 7.04119393028236E-02
 9.21188895116011E-02

 .075684148446073
 .105848567899512

 8.53977028274752E-02
 9.96714630510346E-02

 8.74068828161634E-02
 .114564037695845

 9.69391578000899E-02
 .131507311116263

 .105398863015619
 .138784790917505

 .17868105944514
 .231428859383835

 .231508897192828
 .318746112737915

 1.20041706346713
 1.29635423270479

 1.20041706346713
 1.29635423270479

 1.72668626095532E-02
 1.70683160286868E-02

 0
 2.50085216537535E-05

 3.62558794951248E-04
 2.62589477364412E-04

 3.62558794951248E-04
 4.12640607286933E-04

 2.71919096213436E-04
 5.25178954728824E-04

 4.98518343057966E-04
 4.6265765059444E-04

 8.45970521552912E-04
 4.37649128940687E-04

 7.25117589902496E-04
 6.25213041343838E-04

 1.34448886461088E-03
 9.6282808366951E-04

 1.07256976839744E-03
 1.2379218218608E-03

 1.43512856334869E-03
 1.56303260335959E-03

 1.73726089247473E-03
 1.41298147343707E-03

 1.91854028995035E-03
 1.51301556005209E-03

 6.69223109014178E-03
 7.62759910439482E-03

 0
 2.50085216537535E-05

 0
 2.50085216537535E-05

 .712473351928571
 .773676130141346

 4.5319849368906E-05
 2.50085216537535E-05

 9.0639698737812E-05
 5.0017043307507E-05

 1.66172781019322E-04
 7.50255649612605E-05

 3.47452178494946E-04
 4.6265765059444E-04

 1.4804484127176E-03
 7.62759910439482E-04

 1.64662119373692E-03
 1.05035790945765E-03

 1.84300720766884E-03
 1.75059651576275E-03

 1.91854028995035E-03
 5.72695145870956E-03

 8.39927874970391E-03
 2.07570729726154E-03

 1.04386719713047E-02
 3.52620155317925E-03

 5.3326356090746E-03
 .011766509438091

 7.85544055727704E-03
 9.90337457488639E-03

 1.33995687967399E-02
 6.85233493312846E-03

 1.85660316247952E-02
 3.12856605888456E-02

 .031814534256972
 2.62964605189218E-02

 2.53035825643058E-02
 3.35364275376835E-02

 4.20266069814322E-02
 4.00511474284863E-02

 .039866360828181
 6.15459717898874E-02

 .053975940598367
 5.23928528646136E-02

 5.76015285478795E-02
 5.32556468616681E-02

 7.99593209032065E-02
 9.10810358629703E-02

 .084340239675534
 9.41445797655551E-02

 9.26639853429564E-02
 .10297258790933

 .133391423309147
 .143086256641951

 1.93968955298918E-02
 2.81470911212996E-02

 6.0426465825208E-05
 5.0017043307507E-05

 4.07878644320154E-04
 3.37615042325672E-04

 1.89285904197464E-02
 2.77594590356664E-02

 6.43239728709339E-02
 8.08900632890658E-02

 1.41851128524676E-02
 1.62430348141129E-02

 1.62698259234373E-02
 1.80686568948369E-02

 3.38690340950291E-02
 4.65783715801159E-02

 .137107650957397
 .161217434840922

 1.83696456108632E-02
 1.80811611556638E-02

 .118738005346534
 .143136273685258

 .249848329570779
 .235330188761821

 4.45796251625472E-02
 3.89757809973749E-02

 4.71628565765748E-02
 4.84290021824937E-02

 7.23909060585992E-02
 5.85199406697832E-02

 8.57149417730575E-02
 8.94054649121688E-02

 2.41705863300832E-04
 1.62555390749398E-04

 4.5319849368906E-05
 1.25042608268768E-05

 4.5319849368906E-05
 1.25042608268768E-05

 1.5106616456302E-05
 1.25042608268768E-05

 3.0213232912604E-05
 0

 1.5106616456302E-05
 2.50085216537535E-05

 1.5106616456302E-05
 2.50085216537535E-05

 1.5106616456302E-05
 2.50085216537535E-05

 1.81279397475624E-04
 1.25042608268768E-04

 1.81279397475624E-04
 1.25042608268768E-04

 3.0213232912604E-05
 3.75127824806303E-05

 1.5106616456302E-04
 8.75298257881373E-05

 8.50253247318274
 7.60261559126272

 6.30270698481604
 5.34145760167777

 3.32345562038644E-04
 2.25076694883782E-04

 3.0213232912604E-05
 0

 4.5319849368906E-05
 3.75127824806303E-05

 1.20852931650416E-04
 1.00034086615014E-04

 1.35959548106718E-04
 8.75298257881373E-05

 1.20852931650416E-04
 1.12538347441891E-04

 1.5106616456302E-05
 1.25042608268768E-05

 4.5319849368906E-05
 0

 0
 7.50255649612605E-05

 6.0426465825208E-05
 2.50085216537535E-05

 3.0213232912604E-05
 0

 3.0213232912604E-05
 0

 2.89744903631872E-02
 1.59429325542679E-02

 3.0213232912604E-05
 0

 3.0213232912604E-05
 0

 9.0639698737812E-05
 0

 1.10278300131005E-03
 0

 1.17831608359156E-03
 0

 3.23281592164863E-03
 0

 3.50473501786206E-03
 0

 7.28138913193756E-03
 0

 1.25233850422744E-02
 1.59429325542679E-02

 8.18174347273316E-02
 6.12708780516961E-02

 3.0213232912604E-05
 0

 .032161986435467
 2.89098510317391E-02

 4.96252350589521E-02
 3.23610270199571E-02

 3.0213232912604E-05
 0

 3.0213232912604E-05
 0

 .180720452666741
 .14384901655239

 3.0213232912604E-05
 0

 .180690239433828
 .14384901655239

 3.0213232912604E-05
 1.25042608268768E-05

 3.0213232912604E-05
 1.25042608268768E-05

 5.69066241908896E-02
 3.75127824806303E-05

 4.5319849368906E-05
 0

 4.5319849368906E-05
 0

 7.553308228151E-05
 0

 9.0639698737812E-05
 0

 2.56812479757134E-04
 0

 3.17238945582342E-04
 0

 3.32345562038644E-04
 0

 3.92772027863852E-04
 0

 4.07878644320154E-04
 0

 4.07878644320154E-04
 0

 4.5319849368906E-04
 1.25042608268768E-05

 5.13624959514268E-04
 0

 7.25117589902496E-04
 0

 7.85544055727704E-04
 0

 1.02724991902854E-03
 0

 1.25384916587307E-03
 0

 1.34448886461088E-03
 0

 1.46534179626129E-03
 0

 1.5106616456302E-03
 0

 1.76747412538733E-03
 0

 1.79768735829994E-03
 0

 1.91854028995035E-03
 0

 1.93364690640666E-03
 0

 2.2962057013579E-03
 0

 2.38684540009572E-03
 0

 2.58323141402764E-03
 0

 2.67387111276545E-03
 0

 2.74940419504696E-03
 0

 2.76451081150327E-03
 0

 2.93068359252259E-03
 0

 3.50473501786206E-03
 0

 4.42623862169648E-03
 2.50085216537535E-05

 4.5168783204343E-03
 0

 7.20585604965605E-03
 0

 .612497764220765
 .52315326447487

 4.5319849368906E-05
 1.25042608268768E-05

 2.11492630388228E-04
 1.12538347441891E-04

 2.65876449630915E-03
 7.00238606305098E-04

 .010604844752324
 1.25042608268768E-05

 1.75992081715918E-02
 9.94088735736702E-03

 3.94131623344919E-02
 3.30612656262621E-02

 7.57143616789856E-02
 .082990779107981

 .101138797174942
 .100246659049071

 .365111813132363
 .296075887858788

 .183454750245331
 .143273820554354

 3.0213232912604E-05
 2.50085216537535E-05

 7.553308228151E-05
 2.50085216537535E-05

 1.59525869778549E-02
 6.75230084651345E-04

 .167396416952282
 .142548573426395

 6.19371274708382E-03
 2.86347572935478E-03

 6.0426465825208E-05
 0

 2.56812479757134E-04
 3.00102259845042E-04

 9.36610220290724E-04
 6.37717302170715E-04

 1.02724991902854E-03
 5.75195998036331E-04

 1.14810285067895E-03
 1.21291330020705E-03

 2.76451081150327E-03
 1.37546869095644E-04

 1.67834508829515E-02
 .013904738039487

 3.0213232912604E-05
 3.75127824806303E-05

 3.0213232912604E-04
 3.12606520671919E-04

 1.64511053209129E-02
 1.35546187363344E-02

 1.23736784731924
 1.09563583791177

 4.5319849368906E-05
 2.50085216537535E-05

 1.33844621802836E-02
 1.32044994331819E-02

 1.36865945094096E-02
 1.45299510808308E-02

 1.53030024702339E-02
 1.32295079548356E-02

 1.53483223196028E-02
 1.40422849085826E-02

 2.57265678250823E-02
 3.34989147552028E-02

 3.33100892861459E-02
 3.46493067512755E-02

 4.02591328560448E-02
 3.39615724057973E-02

 4.27366179548783E-02
 3.35739403201641E-02

 4.11655298434229E-02
 3.62623563979426E-02

 4.44285589979842E-02
 3.77753719579947E-02

 4.65736985347791E-02
 3.81129870003204E-02

 4.51536765878867E-02
 4.26020166371691E-02

 4.66945514664295E-02
 4.18892737700371E-02

 4.85828785234672E-02
 4.29896487228023E-02

 5.30997568439015E-02
 4.00511474284863E-02

 4.87792645373991E-02
 4.39649810672987E-02

 5.35529553375906E-02
 4.47152367169113E-02

 5.32357163920082E-02
 .04860406183407

 5.76619550137047E-02
 4.79663445318992E-02

 5.88855909466652E-02
 5.09673671303497E-02

 7.33577295118025E-02
 .06666021446808

 6.78438145052523E-02
 7.92144923382643E-02

 9.08058715188313E-02
 7.33499940104591E-02

 9.69391578000899E-02
 7.16994315813113E-02

 .110807031706975
 9.80959261868482E-02

 2.71768030048873E-02
 3.75127824806303E-05

 7.553308228151E-05
 0

 9.0639698737812E-05
 0

 9.0639698737812E-05
 0

 1.20852931650416E-04
 0

 2.11492630388228E-04
 2.50085216537535E-05

 2.71919096213436E-04
 0

 3.17238945582342E-04
 0

 4.38091877232758E-04
 0

 4.5319849368906E-04
 0

 5.89158041795778E-04
 0

 6.7979774053359E-04
 0

 1.02724991902854E-03
 0

 1.02724991902854E-03
 0

 1.08767638485374E-03
 0

 1.10278300131005E-03
 1.25042608268768E-05

 1.42002194689239E-03
 0

 1.4955550291739E-03
 0

 1.55598149499911E-03
 0

 1.90343367349405E-03
 0

 2.00917998868817E-03
 0

 2.14513953679488E-03
 0

 2.35663216718311E-03
 0

 2.79472404441587E-03
 0

 3.91261366218222E-03
 0

 4.5319849368906E-05
 2.50085216537535E-05

 4.5319849368906E-05
 2.50085216537535E-05

 9.72110768963033E-02
 .116877325948817

 0
 1.12538347441891E-04

 3.0213232912604E-04
 7.50255649612605E-05

 5.74051425339476E-04
 1.62555390749398E-04

 8.3086390509661E-04
 5.50187476382577E-04

 7.40224206358798E-04
 7.00238606305098E-04

 1.10278300131005E-03
 9.00306779535127E-04

 1.82790059121254E-03
 1.47550277757146E-03

 1.40340466879046E-02
 1.13538688308041E-02

 1.37923408246037E-02
 2.41332233958721E-02

 2.56510347428008E-02
 2.54461707826942E-02

 3.83556991825508E-02
 5.19677079964998E-02

 1.5106616456302E-04
 6.25213041343838E-05

 1.5106616456302E-04
 6.25213041343838E-05

 .259697843500288
 .24462085455619

 1.96386013931926E-04
 3.37615042325672E-04

 6.0426465825208E-04
 1.41298147343707E-03

 9.53378564557219E-02
 .094769792806899

 .163559336372382
 .148100465233528

 5.89611240289467E-02
 .049616906961047

 9.97036686115932E-04
 0

 4.06367982674524E-03
 0

 5.39004075160855E-02
 .049616906961047

 .163514016523013
 .125567787223496

 8.00650672184006E-04
 4.2514486811381E-04

 3.58026810014357E-03
 5.0017043307507E-05

 7.14542958383084E-03
 1.25042608268768E-05

 2.54697553453252E-02
 1.54052493387122E-02

 .126517912821529
 .109674871712536

 6.60159139140397E-03
 8.44037605814181E-03

 9.97036686115932E-04
 8.75298257881373E-04

 5.60455470528804E-03
 7.56507780026044E-03

 1.16320946713525E-03
 4.33897850692623E-03

 1.16320946713525E-03
 4.33897850692623E-03

 8.81622136389785E-02
 5.67693441540205E-02

 6.26924582936533E-03
 1.95066468899277E-03

 8.18929678096131E-02
 5.48186794650277E-02

 .111139377269014
 9.84335412291738E-02

 5.13624959514268E-03
 3.86381659550492E-03

 .106003127673871
 9.45697246336689E-02

 .152909171770689
 .125817872440034

 1.16925211371777E-02
 6.38967728253402E-03

 .015408748785428
 8.44037605814181E-03

 .125807901848083
 .110987819099358

 .353600571392661
 .308955276510471

 8.67119784591735E-03
 1.36046357796419E-02

 2.20556600262009E-02
 1.54552663820197E-02

 2.81889463074595E-02
 3.20359162384583E-02

 5.46557383389006E-02
 4.39024597631643E-02

 .060879664318897
 .058644983278052

 9.04282061074237E-02
 7.17244401029651E-02

 8.87211584478616E-02
 7.35875749661697E-02

 1.45325650309625E-02
 1.82437165464132E-02

 1.45325650309625E-02
 1.82437165464132E-02

 .776132633675428
 .70451506350789

 2.17535276970749E-02
 .020694551668481

 2.09981968742598E-02
 2.16573797521505E-02

 .021708207847706
 2.17199010562849E-02

 2.96542881037208E-02
 3.00852515494655E-02

 3.26756113949812E-02
 3.27486591055902E-02

 4.18755408168691E-02
 3.01977898969074E-02

 4.16791548029372E-02
 .033073769887089

 4.61960331233715E-02
 .032973735800474

 5.52146831477838E-02
 4.52654241932939E-02

 5.86287784669081E-02
 4.70285249698835E-02

 7.17866414003471E-02
 5.74195657170181E-02

 6.31003369379734E-02
 6.57849162101986E-02

 6.93393695344262E-02
 6.34716279572264E-02

 9.93713230495545E-02
 8.73672703973879E-02

 .102150940477514
 .115026695346439

 .274683607024939
 .250435335840688

 3.22073062848359E-02
 3.19233778910164E-02

 3.89297506078902E-02
 3.05103964175793E-02

 .203546550132213
 .188001561532092

 9.35703823303346E-02
 7.59258717407957E-02

 9.35703823303346E-02
 7.59258717407957E-02

 .099643242145768
 7.93645434681868E-02

 .099643242145768
 7.93645434681868E-02

 .291844723319298
 .253461366960792

 .10274009851931
 8.16903359819859E-02

 .189104624799988
 .171771030978806

 .131790121964779
 .113576201090522

 .131790121964779
 .113576201090522

 .154721965745445
 .131969968766857

 .154721965745445
 .131969968766857

 .202519300213185
 .15414002321291

 .202519300213185
 .15414002321291

 .251223031668302
 .197654850890441

 .251223031668302
 .197654850890441

 .286451661244398
 .222325757501869

 .286451661244398
 .222325757501869

 1.22159653973886
 1.24936322051739

 2.87025712669738E-04
 0

 3.0213232912604E-05
 0

 4.5319849368906E-05
 0

 6.0426465825208E-05
 0

 7.553308228151E-05
 0

 7.553308228151E-05
 0

 .331620444448741
 .395034608042691

 3.0213232912604E-05
 0

 3.0213232912604E-05
 3.75127824806303E-05

 1.37470209752348E-03
 6.50221562997591E-04

 5.71030102048216E-03
 7.4775479744723E-03

 1.13752821915954E-02
 4.23894442031122E-03

 1.64208920880003E-02
 .013592131518815

 1.88530573374649E-02
 1.31544823898744E-02

 1.71762229108154E-02
 .018368759154682

 .120218453759251
 .160079547105676

 .140431106577783
 .177435461133381

 1.04235653548484E-03
 8.25281214573866E-04

 3.0213232912604E-05
 0

 4.22985260776456E-04
 4.75161911421317E-04

 5.89158041795778E-04
 3.50119303152549E-04

 .019759454324843
 1.69307691595911E-02

 9.0639698737812E-05
 2.50085216537535E-05

 1.66172781019322E-04
 6.25213041343838E-05

 1.08465506156248E-02
 7.2649755404154E-03

 8.65609122946105E-03
 9.5782637933876E-03

 2.38080275351319E-02
 2.11196965365948E-02

 2.56812479757134E-04
 5.0017043307507E-05

 9.21503603834422E-04
 2.00068173230028E-04

 8.59566476363584E-03
 5.20177250398073E-03

 1.40340466879046E-02
 1.56678388160766E-02

 7.85544055727704E-04
 1.66306668997461E-03

 2.56812479757134E-04
 2.75093738191289E-04

 5.2873157597057E-04
 1.38797295178332E-03

 .26663178045373
 .235742829369108

 4.83411726601664E-04
 2.12572434056905E-04

 9.66823453203328E-04
 7.87768432093236E-04

 2.47748509883353E-03
 7.12742867131975E-04

 2.94579020897889E-03
 2.40081807876034E-03

 3.45941516849316E-03
 2.23826268801094E-03

 3.82197396344441E-03
 2.17574138387656E-03

 1.78107008019801E-02
 6.46470284749528E-03

 2.75544684162948E-02
 1.58679069893066E-02

 7.96118687247115E-02
 .053980893989627

 .127499842891189
 .150901419658749

 2.25541783692589E-02
 2.27827632265695E-02

 6.64691124077288E-04
 6.12708780516961E-04

 6.7979774053359E-03
 4.15141459452308E-03

 8.02161333829636E-03
 4.56405520181002E-03

 7.06989650154933E-03
 1.34545846497194E-02

 6.46563184329725E-03
 1.25042608268768E-05

 1.69194104310582E-03
 0

 4.77369080019143E-03
 1.25042608268768E-05

 .319912816695107
 .2813333643439

 1.07256976839744E-03
 6.50221562997591E-04

 1.91854028995035E-03
 3.62623563979426E-04

 1.19342270004786E-03
 1.01284512697702E-03

 1.4955550291739E-03
 1.06286217028452E-03

 3.26302915456123E-03
 2.98851833762355E-03

 8.18778611931568E-03
 0

 6.51095169266616E-03
 2.58838199116349E-03

 1.44268187157684E-02
 8.77799110046748E-03

 .134902084954777
 .127018281479414

 .14694205827045
 .136871639010993

 .223502390470988
 .269579359166636

 3.35366885329904E-03
 6.75230084651345E-04

 1.19644402333912E-02
 7.84017153845173E-03

 1.35506349613029E-02
 1.80311441123563E-02

 4.55766618486631E-02
 6.06331607495254E-02

 .149056984574332
 .182399652681651

 5.22688929388049E-03
 4.33897850692623E-03

 5.22688929388049E-03
 4.33897850692623E-03

 .32006388285967
 .287935614060491

 5.72540763693846E-03
 1.13788773524579E-03

 3.0213232912604E-05
 0

 1.40491533043609E-03
 4.87666172248194E-04

 4.29027907358977E-03
 6.50221562997591E-04

 1.5106616456302E-05
 1.25042608268768E-05

 1.5106616456302E-05
 1.25042608268768E-05

 .111985347790567
 .118577905421272

 2.2659924684453E-04
 8.75298257881373E-05

 8.61077138009214E-04
 6.62725823824468E-04

 1.11788961776635E-03
 1.18790477855329E-03

 3.65580118242508E-03
 1.50051129922521E-03

 1.23269990283424E-02
 7.41502667033792E-03

 4.10446769117725E-02
 5.22928187779986E-02

 5.27523046654066E-02
 5.54313882455447E-02

 1.91854028995035E-03
 1.55052834253272E-03

 2.87025712669738E-04
 2.75093738191289E-04

 3.7766541140755E-04
 2.12572434056905E-04

 5.13624959514268E-04
 5.50187476382577E-04

 7.40224206358798E-04
 5.12674693901947E-04

 5.68008778756955E-03
 4.87666172248194E-03

 1.45023517980499E-03
 1.06286217028452E-03

 4.22985260776456E-03
 3.81379955219741E-03

 6.49735573785549E-02
 5.04671966972746E-02

 1.20852931650416E-03
 1.86313486320464E-03

 1.04084587383921E-02
 8.12776953746989E-04

 1.05897381358677E-02
 8.31533344987304E-03

 1.22212527131483E-02
 .015730360120211

 3.05455784746426E-02
 .023745591310239

 7.58805344600049E-02
 6.84733322879771E-02

 1.52274693879524E-02
 1.91940403692558E-02

 6.06530650720525E-02
 4.92792919187213E-02

 5.38853008996292E-02
 4.28395975928798E-02

 5.38853008996292E-02
 4.28395975928798E-02

 .147954201573022
 .160479683452136

 .119583975868087
 .137271775357453

 1.5106616456302E-05
 1.25042608268768E-05

 9.0639698737812E-05
 1.87563912403151E-04

 2.56812479757134E-04
 6.25213041343838E-05

 1.96386013931926E-04
 2.75093738191289E-04

 5.43838192426872E-04
 3.37615042325672E-04

 7.70437439271402E-04
 3.37615042325672E-04

 1.04235653548484E-03
 6.25213041343838E-04

 1.79768735829994E-03
 1.22541756103392E-03

 1.84300720766884E-03
 1.28793886516831E-03

 2.50769833174613E-03
 1.75059651576275E-03

 .110520005994305
 .131169696073937

 1.5408748785428E-03
 5.62691737209454E-04

 1.5408748785428E-03
 5.62691737209454E-04

 2.68293508263923E-02
 2.26452163574738E-02

 5.3175289926183E-03
 4.0888932903887E-03

 5.81604733567627E-03
 4.90167024413569E-03

 5.93690026732668E-03
 4.87666172248194E-03

 9.75887423077109E-03
 8.77799110046748E-03

 .510210864195144
 .563379471554932

 .121925501418813
 .156065679380249

 3.0213232912604E-05
 1.25042608268768E-05

 5.74051425339476E-04
 3.25110781498796E-04

 6.7979774053359E-04
 4.75161911421317E-04

 1.26895578232937E-03
 9.2531530118888E-04

 3.64069456596878E-03
 2.27577547049157E-03

 1.82941125285817E-02
 2.48459662630041E-02

 2.12399027375606E-02
 2.88348254667778E-02

 3.08477108037687E-02
 3.64749288319995E-02

 4.53500626018186E-02
 .06189609109304

 .264411107834654
 .285397249112635

 1.20852931650416E-04
 1.62555390749398E-04

 8.3086390509661E-04
 6.37717302170715E-04

 3.83708057990071E-03
 2.91349277266228E-03

 .022962057013579
 3.06354390258481E-02

 4.19359672826943E-02
 4.65408587976353E-02

 .194724286121733
 .204507185823569

 7.29498508674823E-02
 9.68329958433336E-02

 4.22985260776456E-04
 3.25110781498796E-04

 7.10010973446194E-04
 7.75264171266359E-04

 1.14810285067895E-03
 6.00204519690084E-04

 4.68305110145362E-04
 1.53802408170584E-03

 7.02004466724354E-02
 9.35943922891725E-02

 2.32037628768799E-02
 2.50085216537535E-05

 2.32037628768799E-02
 2.50085216537535E-05

 2.77206411973142E-02
 .025058538697061

 2.77206411973142E-02
 .025058538697061

 .914025828688552
 1.02857548709723

 .755013583869518
 .874447968145145

 1.39434069891667E-02
 9.17812744692754E-03

 0
 2.50085216537535E-05

 1.35959548106718E-04
 1.00034086615014E-04

 2.87025712669738E-04
 6.25213041343838E-05

 1.96386013931926E-04
 1.87563912403151E-04

 4.22985260776456E-04
 3.25110781498796E-04

 4.98518343057966E-04
 3.37615042325672E-04

 4.68305110145362E-04
 4.2514486811381E-04

 4.68305110145362E-04
 5.62691737209454E-04

 6.19371274708382E-04
 4.50153389767563E-04

 5.74051425339476E-04
 6.00204519690084E-04

 6.64691124077288E-04
 5.50187476382577E-04

 6.7979774053359E-04
 6.75230084651345E-04

 8.76183754465516E-04
 5.62691737209454E-04

 9.8193006965963E-04
 5.37683215555701E-04

 1.13299623422265E-03
 4.37649128940687E-04

 1.13299623422265E-03
 5.0017043307507E-04

 1.04235653548484E-03
 9.87836605323264E-04

 1.96386013931926E-03
 7.50255649612605E-04

 1.79768735829994E-03
 1.10037495276515E-03

 4.15885151041994E-02
 .048504027747455

 3.0213232912604E-05
 0

 1.5106616456302E-05
 1.25042608268768E-05

 3.0213232912604E-05
 0

 1.5106616456302E-05
 2.50085216537535E-05

 1.5106616456302E-05
 2.50085216537535E-05

 3.0213232912604E-05
 2.50085216537535E-05

 4.5319849368906E-05
 3.75127824806303E-05

 1.66172781019322E-04
 0

 4.38091877232758E-04
 2.25076694883782E-04

 2.58323141402764E-03
 4.12640607286933E-03

 8.02161333829636E-03
 9.36569135933069E-03

 3.01981262961477E-02
 3.46618110121024E-02

 7.553308228151E-05
 6.25213041343838E-05

 3.0213232912604E-05
 1.25042608268768E-05

 4.5319849368906E-05
 5.0017043307507E-05

 3.75550485103668E-02
 4.51403815850251E-02

 1.5106616456302E-05
 2.50085216537535E-05

 4.5319849368906E-05
 0

 1.20852931650416E-04
 2.50085216537535E-05

 3.47452178494946E-04
 1.62555390749398E-04

 3.7766541140755E-04
 2.62589477364412E-04

 5.43838192426872E-04
 2.62589477364412E-04

 1.11788961776635E-03
 7.25247127958852E-04

 2.37173878363941E-03
 1.38797295178332E-03

 1.44419253322247E-02
 2.01193556704447E-02

 1.81732595969313E-02
 2.21700544460525E-02

 5.2873157597057E-04
 5.75195998036331E-04

 4.5319849368906E-05
 1.25042608268768E-05

 1.35959548106718E-04
 6.25213041343838E-05

 1.35959548106718E-04
 2.50085216537535E-04

 2.11492630388228E-04
 2.50085216537535E-04

 1.12242160270324E-02
 1.06911430069796E-02

 4.5319849368906E-05
 3.75127824806303E-05

 1.37470209752348E-03
 1.17540051772642E-03

 1.94875352286296E-03
 1.66306668997461E-03

 2.05449983805707E-03
 2.16323712304968E-03

 2.34152555072681E-03
 2.97601407679667E-03

 3.45941516849316E-03
 2.67591181695163E-03

 .173469276767716
 .200880950183775

 6.0426465825208E-05
 3.75127824806303E-05

 1.5106616456302E-05
 1.50051129922521E-04

 2.41705863300832E-04
 6.25213041343838E-05

 1.96386013931926E-04
 2.87597999018165E-04

 9.8193006965963E-04
 5.75195998036331E-04

 1.13299623422265E-03
 5.12674693901947E-04

 2.50769833174613E-03
 5.0017043307507E-04

 1.60130134436801E-03
 1.27543460434143E-03

 7.67416115980141E-03
 .008015231190028

 8.28295780299038E-02
 7.15243719297351E-02

 7.62279866384999E-02
 .117940188119102

 9.8193006965963E-04
 1.88814338485839E-03

 1.35959548106718E-04
 3.75127824806303E-05

 9.0639698737812E-05
 1.37546869095644E-04

 3.0213232912604E-04
 6.37717302170715E-04

 4.5319849368906E-04
 1.0753664311114E-03

 2.40799466313454E-02
 2.14448073180936E-02

 1.20852931650416E-04
 1.00034086615014E-04

 2.87025712669738E-04
 2.25076694883782E-04

 3.7766541140755E-04
 2.12572434056905E-04

 3.62558794951248E-04
 2.37580955710658E-04

 2.87025712669738E-04
 3.25110781498796E-04

 4.68305110145362E-04
 3.37615042325672E-04

 3.17238945582342E-04
 4.75161911421317E-04

 3.7766541140755E-04
 4.2514486811381E-04

 4.68305110145362E-04
 4.6265765059444E-04

 6.64691124077288E-04
 5.12674693901947E-04

 1.02724991902854E-03
 3.87632085633179E-04

 7.85544055727704E-04
 7.25247127958852E-04

 6.49584507620986E-04
 8.37785475400743E-04

 9.0639698737812E-04
 9.00306779535127E-04

 1.11788961776635E-03
 9.2531530118888E-04

 1.45023517980499E-03
 7.25247127958852E-04

 1.45023517980499E-03
 1.08787069193828E-03

 1.25384916587307E-03
 1.56303260335959E-03

 1.55598149499911E-03
 1.41298147343707E-03

 1.85811382412515E-03
 1.72558799410899E-03

 2.49259171528983E-03
 1.37546869095644E-03

 2.11492630388228E-03
 1.72558799410899E-03

 3.68601441533769E-03
 4.73911485338629E-03

 .310712887273219
 .408776790691428

 4.5319849368906E-05
 1.87563912403151E-04

 3.62558794951248E-04
 1.00034086615014E-04

 6.19371274708382E-04
 2.87597999018165E-04

 7.38713544713168E-03
 4.57655946263689E-03

 1.65115317867381E-02
 2.71467502551494E-02

 2.68293508263923E-02
 2.33704634854327E-02

 .258957619293929
 .353107821490173

 3.65580118242508E-03
 3.12606520671919E-03

 2.87025712669738E-04
 1.37546869095644E-04

 4.5319849368906E-04
 2.87597999018165E-04

 4.5319849368906E-04
 5.62691737209454E-04

 1.13299623422265E-03
 1.15039199607266E-03

 1.32938224815458E-03
 9.87836605323264E-04

 3.47452178494946E-04
 2.25076694883782E-04

 3.47452178494946E-04
 2.25076694883782E-04

 1.53181090866902E-02
 1.05285876162302E-02

 1.37470209752348E-03
 1.15039199607266E-03

 1.39434069891667E-02
 9.37819562015757E-03

 3.89146439914339E-02
 3.23235142374764E-02

 1.96386013931926E-03
 1.67557095080149E-03

 5.19667606096789E-03
 3.15107372837294E-03

 4.42623862169648E-03
 8.86552092625562E-03

 2.73278691694503E-02
 1.86313486320464E-02

 3.33856223684274E-03
 3.47618450987174E-03

 3.33856223684274E-03
 3.47618450987174E-03

 7.92795231626729E-02
 7.76264512132509E-02

 7.92795231626729E-02
 7.76264512132509E-02

 2.21765129578513E-02
 1.92065446300827E-02

 1.90343367349405E-03
 2.53836494785598E-03

 1.5106616456302E-05
 1.25042608268768E-05

 1.88832705703775E-03
 2.52586068702911E-03

 2.87025712669738E-04
 2.00068173230028E-04

 3.0213232912604E-05
 0

 1.5106616456302E-04
 7.50255649612605E-05

 1.05746315194114E-04
 1.25042608268768E-04

 3.7766541140755E-04
 5.50187476382577E-04

 4.5319849368906E-05
 6.25213041343838E-05

 1.5106616456302E-04
 1.25042608268768E-04

 1.81279397475624E-04
 3.62623563979426E-04

 6.64691124077288E-04
 4.75161911421317E-04

 1.66172781019322E-04
 1.00034086615014E-04

 1.81279397475624E-04
 2.37580955710658E-04

 3.17238945582342E-04
 1.37546869095644E-04

 8.3237456674224E-03
 7.56507780026044E-03

 3.0213232912604E-04
 7.50255649612605E-05

 2.87025712669738E-04
 1.50051129922521E-04

 3.47452178494946E-04
 1.75059651576275E-04

 4.07878644320154E-04
 3.50119303152549E-04

 3.92772027863852E-04
 4.12640607286933E-04

 8.3086390509661E-04
 4.75161911421317E-04

 9.0639698737812E-04
 7.12742867131975E-04

 9.66823453203328E-04
 1.12538347441891E-03

 3.88240042926961E-03
 4.0888932903887E-03

 4.77369080019143E-03
 4.28896146361873E-03

 2.56812479757134E-04
 1.25042608268768E-04

 2.41705863300832E-04
 2.87597999018165E-04

 4.68305110145362E-04
 2.62589477364412E-04

 4.98518343057966E-04
 2.75093738191289E-04

 5.2873157597057E-04
 2.62589477364412E-04

 4.68305110145362E-04
 3.25110781498796E-04

 1.04235653548484E-03
 1.06286217028452E-03

 1.26895578232937E-03
 1.68807521162836E-03

 4.87943711538555E-03
 3.08855242423856E-03

 2.56812479757134E-04
 1.50051129922521E-04

 1.02724991902854E-03
 5.50187476382577E-04

 1.60130134436801E-03
 9.87836605323264E-04

 1.99407337223186E-03
 1.4004772126102E-03

 9.66823453203328E-04
 5.0017043307507E-04

 9.66823453203328E-04
 5.0017043307507E-04

 .132923118199001
 .132945301111354

 1.5106616456302E-05
 2.50085216537535E-05

 1.5106616456302E-05
 2.50085216537535E-05

 2.88536374315368E-03
 1.22541756103392E-03

 3.0213232912604E-05
 1.25042608268768E-05

 4.5319849368906E-05
 5.0017043307507E-05

 4.5319849368906E-05
 6.25213041343838E-05

 6.49584507620986E-04
 1.25042608268768E-04

 8.45970521552912E-04
 7.50255649612605E-05

 1.26895578232937E-03
 9.00306779535127E-04

 8.3086390509661E-04
 8.12776953746989E-04

 1.5106616456302E-05
 3.75127824806303E-05

 1.5106616456302E-04
 6.25213041343838E-05

 6.64691124077288E-04
 7.12742867131975E-04

 .048869904236137
 4.86915916598581E-02

 3.0213232912604E-05
 2.50085216537535E-05

 6.0426465825208E-05
 6.25213041343838E-05

 3.17238945582342E-04
 1.25042608268768E-04

 6.7979774053359E-04
 7.00238606305098E-04

 1.08767638485374E-03
 3.87632085633179E-04

 1.99407337223186E-03
 5.87700258863208E-04

 2.61344464694025E-03
 2.3132882529722E-03

 1.10127233966442E-02
 1.26167991743186E-02

 3.10743100506132E-02
 3.18733608477089E-02

 9.8193006965963E-04
 7.62759910439482E-04

 3.0213232912604E-05
 3.75127824806303E-05

 3.0213232912604E-05
 1.00034086615014E-04

 1.5106616456302E-04
 2.50085216537535E-05

 1.35959548106718E-04
 5.0017043307507E-05

 3.0213232912604E-05
 1.37546869095644E-04

 2.2659924684453E-04
 6.25213041343838E-05

 3.7766541140755E-04
 3.50119303152549E-04

 3.14217622291082E-02
 2.48834790454847E-02

 4.5319849368906E-05
 1.12538347441891E-04

 2.87025712669738E-04
 2.50085216537535E-05

 2.56812479757134E-04
 2.00068173230028E-04

 6.0426465825208E-04
 5.0017043307507E-05

 4.38091877232758E-04
 2.87597999018165E-04

 5.74051425339476E-04
 1.87563912403151E-04

 7.10010973446194E-04
 2.25076694883782E-04

 5.43838192426872E-04
 4.12640607286933E-04

 8.61077138009214E-04
 5.50187476382577E-04

 8.45970521552912E-04
 6.62725823824468E-04

 1.02724991902854E-03
 8.5028973622762E-04

 1.5257682620865E-03
 4.75161911421317E-04

 1.67683442664952E-03
 1.90064764568527E-03

 9.44163528518875E-03
 9.05308483865877E-03

 1.25838115080996E-02
 9.89087031405952E-03

 6.64691124077288E-04
 9.00306779535127E-04

 1.66172781019322E-04
 1.25042608268768E-04

 2.11492630388228E-04
 2.00068173230028E-04

 2.87025712669738E-04
 5.75195998036331E-04

 4.72534962753126E-02
 5.56439606796016E-02

 2.2659924684453E-04
 4.87666172248194E-04

 4.42623862169648E-03
 7.90269284258611E-03

 1.61791862246994E-02
 2.68716565169582E-02

 2.64214721820722E-02
 2.03819451478091E-02

 3.59537471659988E-03
 1.97567321064653E-03

 2.71919096213436E-04
 1.25042608268768E-04

 3.0213232912604E-05
 1.25042608268768E-05

 2.41705863300832E-04
 1.12538347441891E-04

 7.25117589902496E-04
 4.12640607286933E-04

 1.66172781019322E-04
 0

 2.56812479757134E-04
 1.50051129922521E-04

 3.0213232912604E-04
 2.62589477364412E-04

 2.41705863300832E-04
 5.0017043307507E-05

 2.41705863300832E-04
 5.0017043307507E-05

 7.553308228151E-04
 4.50153389767563E-04

 2.2659924684453E-04
 1.37546869095644E-04

 5.2873157597057E-04
 3.12606520671919E-04

 1.60130134436801E-03
 9.37819562015757E-04

 6.64691124077288E-04
 4.2514486811381E-04

 9.36610220290724E-04
 5.12674693901947E-04

 3.17238945582342E-04
 0

 1.05746315194114E-04
 0

 1.05746315194114E-04
 0

 2.11492630388228E-04
 0

 2.11492630388228E-04
 0

 2.8241668398892
 2.58212986075005

 .163453590057188
 .154702714950119

 8.05786921779149E-02
 7.59633845232763E-02

 0
 2.50085216537535E-05

 4.5319849368906E-05
 0

 1.5106616456302E-05
 2.50085216537535E-05

 4.5319849368906E-05
 3.75127824806303E-05

 3.0213232912604E-05
 8.75298257881373E-05

 1.05746315194114E-04
 3.75127824806303E-05

 9.0639698737812E-05
 1.00034086615014E-04

 1.96386013931926E-04
 1.25042608268768E-04

 1.20852931650416E-04
 2.25076694883782E-04

 1.5106616456302E-04
 2.87597999018165E-04

 3.7766541140755E-04
 1.00034086615014E-04

 1.81279397475624E-04
 2.62589477364412E-04

 4.98518343057966E-04
 4.75161911421317E-04

 6.34477891164684E-04
 3.87632085633179E-04

 6.94904356989892E-04
 5.62691737209454E-04

 6.64691124077288E-04
 6.62725823824468E-04

 7.553308228151E-04
 1.01284512697702E-03

 2.35663216718311E-03
 0

 1.05746315194114E-03
 1.41298147343707E-03

 1.23874254941676E-03
 1.32545164764894E-03

 1.55598149499911E-03
 1.15039199607266E-03

 1.42002194689239E-03
 1.65056242914773E-03

 4.77369080019143E-03
 2.50085216537535E-05

 2.70408434567806E-03
 2.73843312108601E-03

 2.32641893427051E-03
 3.11356094589231E-03

 4.32049230650237E-03
 4.4140040718875E-03

 7.82522732436444E-03
 2.72592886025913E-03

 4.57730478625951E-03
 5.45185772051827E-03

 6.99436341926782E-03
 7.49005223529918E-03

 1.04084587383921E-02
 1.13913816132847E-02

 .024412292193384
 2.86597658152015E-02

 1.35959548106718E-04
 8.75298257881373E-05

 0
 2.50085216537535E-05

 0
 2.50085216537535E-05

 3.0213232912604E-05
 0

 4.5319849368906E-05
 0

 1.5106616456302E-05
 2.50085216537535E-05

 4.5319849368906E-05
 1.25042608268768E-05

 1.66172781019322E-04
 7.50255649612605E-05

 4.5319849368906E-05
 0

 6.0426465825208E-05
 1.25042608268768E-05

 6.0426465825208E-05
 6.25213041343838E-05

 2.22067261907639E-03
 1.10037495276515E-03

 6.0426465825208E-05
 0

 4.5319849368906E-05
 7.50255649612605E-05

 2.11492630388228E-04
 2.50085216537535E-05

 1.5106616456302E-04
 2.25076694883782E-04

 7.10010973446194E-04
 3.00102259845042E-04

 1.04235653548484E-03
 4.75161911421317E-04

 3.7917607305318E-03
 1.80061355907025E-03

 7.553308228151E-05
 2.50085216537535E-05

 4.22985260776456E-04
 2.37580955710658E-04

 6.7979774053359E-04
 1.25042608268768E-04

 1.05746315194114E-03
 6.37717302170715E-04

 1.55598149499911E-03
 7.75264171266359E-04

 8.47481183198542E-03
 6.46470284749528E-03

 2.87025712669738E-04
 1.62555390749398E-04

 6.19371274708382E-04
 2.87597999018165E-04

 1.67683442664952E-03
 1.56303260335959E-03

 5.89158041795778E-03
 4.45151685436813E-03

 6.46412118165162E-02
 6.68477783804831E-02

 8.76183754465516E-04
 5.0017043307507E-05

 8.27842581805349E-03
 7.62759910439482E-03

 .014457031948681
 1.32795249981431E-02

 4.10295702953162E-02
 4.58906372346377E-02

 1.94875352286296E-03
 7.75264171266359E-04

 1.94875352286296E-03
 7.75264171266359E-04

 1.4955550291739E-03
 1.58804112501335E-03

 1.4955550291739E-03
 1.58804112501335E-03

 1.37514018940071
 1.25944165474385

 7.40224206358798E-04
 6.25213041343838E-04

 1.5106616456302E-05
 1.25042608268768E-05

 7.553308228151E-05
 8.75298257881373E-05

 9.0639698737812E-05
 1.00034086615014E-04

 2.2659924684453E-04
 5.0017043307507E-05

 3.32345562038644E-04
 3.75127824806303E-04

 .309806490285841
 .360672899290433

 3.0213232912604E-05
 0

 3.0213232912604E-05
 0

 4.5319849368906E-05
 0

 6.0426465825208E-05
 1.25042608268768E-05

 1.5106616456302E-04
 1.25042608268768E-05

 1.81279397475624E-04
 1.25042608268768E-05

 2.56812479757134E-04
 2.87597999018165E-04

 1.37470209752348E-03
 7.25247127958852E-04

 3.06664314062931E-03
 1.83812634155088E-03

 3.70112103179399E-03
 1.65056242914773E-03

 2.80983066087217E-03
 4.00136346460056E-03

 1.04084587383921E-02
 9.05308483865877E-03

 9.30567573708203E-03
 1.12413304833622E-02

 .010302712423198
 1.19040563071867E-02

 1.22514659460609E-02
 1.49801044705984E-02

 1.51519363056709E-02
 .016330564639901

 1.64511053209129E-02
 2.07695772334423E-02

 .028460865403673
 .030035234506158

 3.87333645939583E-02
 .051542563128386

 4.43983457650716E-02
 4.94793600919513E-02

 5.08488709919125E-02
 6.16960229198099E-02

 6.17860613062752E-02
 7.51005905262218E-02

 .297766516970169
 .251685761923375

 3.0213232912604E-05
 2.50085216537535E-05

 4.5319849368906E-05
 3.75127824806303E-05

 3.0213232912604E-05
 7.50255649612605E-05

 6.0426465825208E-05
 5.0017043307507E-05

 9.0639698737812E-05
 3.75127824806303E-05

 3.0213232912604E-05
 8.75298257881373E-05

 9.0639698737812E-05
 5.0017043307507E-05

 1.5106616456302E-04
 1.25042608268768E-05

 9.0639698737812E-05
 7.50255649612605E-05

 1.35959548106718E-04
 1.00034086615014E-04

 4.68305110145362E-04
 0

 4.38091877232758E-04
 3.25110781498796E-04

 9.51716836747026E-04
 4.00136346460056E-04

 1.35959548106718E-03
 5.0017043307507E-04

 1.07256976839744E-03
 1.00034086615014E-03

 1.58619472791171E-03
 8.62793997054496E-04

 1.87322044058145E-03
 1.20040903938017E-03

 1.61640796082431E-03
 1.80061355907025E-03

 2.37173878363941E-03
 2.27577547049157E-03

 2.70408434567806E-03
 2.21325416635719E-03

 3.82197396344441E-03
 2.12572434056905E-03

 3.53494825077467E-03
 2.67591181695163E-03

 3.56516148368727E-03
 3.28862059746859E-03

 6.70733770659809E-03
 6.76480510734033E-03

 1.10731498624694E-02
 1.14288943957654E-02

 1.33089290980021E-02
 1.42798658642933E-02

 6.43843993367591E-02
 5.17801440840967E-02

 7.41885934168991E-02
 5.32056298183606E-02

 .101984767696495
 9.50073737626096E-02

 4.27970444207036E-02
 4.18392567267296E-02

 3.0213232912604E-05
 3.75127824806303E-05

 1.5106616456302E-05
 1.00034086615014E-04

 1.35959548106718E-04
 1.50051129922521E-04

 2.11492630388228E-04
 1.50051129922521E-04

 2.2659924684453E-04
 1.87563912403151E-04

 3.0213232912604E-04
 1.87563912403151E-04

 4.22985260776456E-04
 1.12538347441891E-04

 3.92772027863852E-04
 1.75059651576275E-04

 5.74051425339476E-04
 3.50119303152549E-04

 1.56806678816415E-02
 1.94566298466202E-02

 2.48050642212479E-02
 2.09321326241917E-02

 8.26936184817971E-02
 2.29703271389726E-02

 4.5319849368906E-05
 3.75127824806303E-05

 3.92772027863852E-04
 3.50119303152549E-04

 .010453778587761
 0

 1.06803778346055E-02
 0

 8.17267950285938E-03
 3.62623563979426E-03

 1.40189400714483E-02
 0

 3.89297506078902E-02
 1.89564594135452E-02

 2.33548290414429E-02
 1.37546869095644E-04

 0
 1.12538347441891E-04

 1.85811382412515E-03
 0

 5.68008778756955E-03
 0

 7.20585604965605E-03
 2.50085216537535E-05

 8.61077138009214E-03
 0

 .433272866583198
 .438686982589317

 1.20852931650416E-04
 3.75127824806303E-05

 1.77955941855238E-02
 .01785608446078

 1.63906788550877E-02
 2.12947561881711E-02

 2.12852225869295E-02
 2.59838539982499E-02

 .025212942865568
 2.43708043515828E-02

 2.84306521707604E-02
 2.79095101655889E-02

 3.27511444772627E-02
 3.56871603999063E-02

 3.57573611520668E-02
 3.77003463930334E-02

 .03921677632056
 3.88757469107598E-02

 .043869614189101
 4.47652537602188E-02

 4.64830588360412E-02
 4.27020507237841E-02

 5.34018891730276E-02
 5.27054593852855E-02

 7.25570788396185E-02
 6.87984430694759E-02

 2.17837409299875E-02
 2.23826268801094E-03

 6.0426465825208E-05
 1.50051129922521E-04

 3.17238945582342E-04
 1.25042608268768E-04

 1.96386013931926E-03
 1.83812634155088E-03

 9.35099558645094E-03
 0

 1.00912197928097E-02
 1.25042608268768E-04

 1.5106616456302E-04
 1.12538347441891E-04

 1.5106616456302E-04
 1.12538347441891E-04

 9.55040292367412E-02
 9.76832855795612E-02

 2.11492630388228E-04
 7.50255649612605E-05

 1.13299623422265E-03
 6.50221562997591E-04

 1.55598149499911E-03
 6.87734345478222E-04

 6.57137815849137E-03
 1.72558799410899E-03

 2.03032925172699E-02
 .019281570195044

 .06572888820137
 7.52631459169712E-02

 5.93690026732668E-02
 3.66624927444027E-02

 1.96386013931926E-04
 1.37546869095644E-04

 2.52280494820243E-03
 1.05035790945765E-03

 1.62849325398936E-02
 8.44037605814181E-03

 1.69194104310582E-02
 1.28793886516831E-02

 2.34454687401807E-02
 1.41548232560245E-02

 7.90076040664594E-03
 6.12708780516961E-03

 2.41705863300832E-04
 2.37580955710658E-04

 2.37173878363941E-03
 1.81311781989713E-03

 5.2873157597057E-03
 4.07638902956182E-03

 .31843236828239
 .274868661496405

 1.42002194689239E-03
 9.12811040362003E-04

 1.5106616456302E-05
 1.25042608268768E-05

 1.5106616456302E-04
 1.75059651576275E-04

 2.2659924684453E-04
 2.75093738191289E-04

 1.02724991902854E-03
 4.50153389767563E-04

 2.31282297945984E-02
 .025158572783676

 3.0213232912604E-05
 0

 3.0213232912604E-05
 0

 1.20852931650416E-04
 1.62555390749398E-04

 1.20852931650416E-04
 2.12572434056905E-04

 2.41705863300832E-04
 1.75059651576275E-04

 4.38091877232758E-04
 3.87632085633179E-04

 5.43838192426872E-04
 3.87632085633179E-04

 1.17831608359156E-03
 1.2379218218608E-03

 1.4955550291739E-03
 1.6130496466671E-03

 2.03939322160077E-03
 2.06320303643467E-03

 1.97896675577556E-03
 2.53836494785598E-03

 2.74940419504696E-03
 2.8509714685279E-03

 2.61344464694025E-03
 3.0135268592773E-03

 3.64069456596878E-03
 2.92599703348916E-03

 5.90668703441408E-03
 7.59008632191419E-03

 4.65888051512354E-02
 3.29237187571665E-02

 1.5106616456302E-05
 1.25042608268768E-05

 1.5106616456302E-05
 1.25042608268768E-05

 1.5106616456302E-05
 2.50085216537535E-05

 1.5106616456302E-05
 5.0017043307507E-05

 3.0213232912604E-05
 6.25213041343838E-05

 9.0639698737812E-05
 3.75127824806303E-05

 7.553308228151E-05
 5.0017043307507E-05

 1.20852931650416E-04
 2.50085216537535E-05

 1.5106616456302E-04
 7.50255649612605E-05

 2.11492630388228E-04
 1.62555390749398E-04

 3.62558794951248E-04
 1.00034086615014E-04

 2.71919096213436E-04
 1.87563912403151E-04

 3.7766541140755E-04
 1.50051129922521E-04

 4.22985260776456E-04
 3.37615042325672E-04

 3.92772027863852E-04
 3.62623563979426E-04

 5.89158041795778E-04
 2.25076694883782E-04

 4.07878644320154E-04
 4.00136346460056E-04

 6.34477891164684E-04
 4.37649128940687E-04

 7.25117589902496E-04
 4.75161911421317E-04

 7.70437439271402E-04
 4.37649128940687E-04

 8.3086390509661E-04
 9.2531530118888E-04

 1.42002194689239E-03
 9.12811040362003E-04

 1.40491533043609E-03
 9.6282808366951E-04

 2.06960645451337E-03
 1.25042608268768E-03

 1.94875352286296E-03
 1.41298147343707E-03

 1.97896675577556E-03
 1.4504942559177E-03

 2.34152555072681E-03
 1.83812634155088E-03

 2.56812479757134E-03
 1.72558799410899E-03

 2.70408434567806E-03
 1.90064764568527E-03

 3.18749607227972E-03
 2.37580955710658E-03

 4.59241140271581E-03
 2.21325416635719E-03

 3.7615474976192E-03
 3.05103964175793E-03

 5.01539666349226E-03
 3.72626972640927E-03

 7.06989650154933E-03
 5.55189180713328E-03

 4.74800955221572E-02
 2.73843312108601E-03

 3.0213232912604E-05
 0

 1.20852931650416E-04
 1.25042608268768E-05

 1.20852931650416E-04
 2.87597999018165E-04

 5.58944808883174E-04
 0

 6.94904356989892E-04
 0

 7.40224206358798E-04
 0

 8.61077138009214E-04
 0

 8.76183754465516E-04
 0

 1.07256976839744E-03
 0

 1.11788961776635E-03
 0

 6.49584507620986E-04
 5.37683215555701E-04

 1.31427563169827E-03
 0

 1.31427563169827E-03
 0

 5.74051425339476E-04
 7.62759910439482E-04

 1.73726089247473E-03
 0

 2.14513953679488E-03
 0

 2.22067261907639E-03
 0

 1.37470209752348E-03
 9.37819562015757E-04

 3.0213232912604E-03
 0

 4.21474599130826E-03
 0

 1.14659218903332E-02
 0

 .011254429259945
 2.00068173230028E-04

 3.38388208621165E-03
 1.72558799410899E-03

 0
 2.50085216537535E-05

 4.5319849368906E-05
 0

 5.58944808883174E-04
 3.75127824806303E-05

 6.49584507620986E-04
 2.87597999018165E-04

 2.13003292033858E-03
 1.37546869095644E-03

 .15768286257088
 .166819343691363

 3.0213232912604E-05
 0

 1.5106616456302E-04
 2.75093738191289E-04

 3.17238945582342E-04
 2.00068173230028E-04

 3.7766541140755E-04
 4.50153389767563E-04

 2.35663216718311E-03
 2.53836494785598E-03

 2.2357792355327E-03
 4.31396998527248E-03

 1.57410943474667E-02
 2.24576524450707E-02

 .136473173066232
 .136584041011975

 1.5106616456302E-05
 2.50085216537535E-05

 1.5106616456302E-05
 2.50085216537535E-05

 3.55005486723097E-03
 4.7891318966938E-03

 0
 5.0017043307507E-05

 3.55005486723097E-03
 4.73911485338629E-03

 3.51379898773584E-02
 3.97135323861606E-02

 0
 7.50255649612605E-05

 7.553308228151E-05
 8.75298257881373E-05

 2.56812479757134E-04
 1.25042608268768E-04

 3.7766541140755E-04
 2.25076694883782E-04

 4.83411726601664E-04
 3.25110781498796E-04

 7.10010973446194E-04
 2.87597999018165E-04

 4.83411726601664E-04
 6.50221562997591E-04

 1.10278300131005E-03
 9.2531530118888E-04

 1.75236750893103E-03
 1.12538347441891E-03

 1.37470209752348E-03
 1.43798999509083E-03

 2.03939322160077E-03
 1.68807521162836E-03

 1.63151457728062E-03
 2.05069877560779E-03

 2.31131231781421E-03
 2.36330529627971E-03

 5.46859515718132E-03
 4.58906372346377E-03

 1.70704765956213E-02
 2.37580955710658E-02

 4.5319849368906E-05
 6.25213041343838E-05

 4.5319849368906E-05
 6.25213041343838E-05

 7.04119393028236E-02
 6.57724119493717E-03

 2.76753213479453E-02
 6.42719006501465E-03

 3.0213232912604E-05
 0

 3.0213232912604E-05
 0

 4.5319849368906E-05
 0

 4.5319849368906E-05
 0

 1.05746315194114E-04
 0

 1.66172781019322E-04
 0

 2.11492630388228E-04
 0

 2.87025712669738E-04
 0

 3.0213232912604E-04
 0

 3.17238945582342E-04
 0

 3.47452178494946E-04
 0

 4.22985260776456E-04
 0

 5.58944808883174E-04
 0

 6.19371274708382E-04
 0

 1.04235653548484E-03
 0

 1.58619472791171E-03
 0

 1.76747412538733E-03
 0

 2.02428660514447E-03
 0

 2.71919096213436E-03
 0

 3.44430855203686E-03
 0

 4.27517245713347E-03
 0

 3.71622764825029E-03
 3.10105668506544E-03

 3.61048133305618E-03
 3.32613337994922E-03

 .024910810536442
 1.50051129922521E-04

 3.0213232912604E-05
 0

 1.05746315194114E-04
 0

 1.05746315194114E-04
 0

 1.20852931650416E-04
 0

 9.0639698737812E-05
 2.50085216537535E-05

 2.2659924684453E-04
 1.12538347441891E-04

 9.51716836747026E-04
 0

 1.29916901524197E-03
 0

 1.94875352286296E-03
 0

 3.14217622291082E-03
 0

 3.61048133305618E-03
 0

 5.60455470528804E-03
 0

 7.67416115980141E-03
 1.25042608268768E-05

 1.78258074184364E-02
 0

 1.02724991902854E-03
 0

 1.5408748785428E-03
 0

 1.60130134436801E-03
 0

 3.12706960645451E-03
 0

 3.50473501786206E-03
 0

 7.02457665218043E-03
 0

 .15727498392656
 .166244147693327

 .15727498392656
 .166244147693327

 3.0213232912604E-05
 5.0017043307507E-05

 9.0639698737812E-05
 0

 1.5106616456302E-04
 1.75059651576275E-04

 2.56812479757134E-04
 1.37546869095644E-04

 3.7766541140755E-04
 4.00136346460056E-04

 4.5319849368906E-04
 4.2514486811381E-04

 2.74940419504696E-03
 2.72592886025913E-03

 1.26140247410122E-02
 1.34670889105463E-02

 .017055369979165
 1.59929495975754E-02

 1.67381310335826E-02
 .021919969229515

 3.71924897154155E-02
 3.88757469107598E-02

 6.95659687812707E-02
 7.20745594061176E-02

 .739453768919526
 .720295440671409

 .282508834349304
 .29462539360287

 5.43838192426872E-04
 1.25042608268768E-05

 2.2357792355327E-03
 1.90064764568527E-03

 1.64511053209129E-02
 2.09196283633648E-02

 1.88379507210086E-02
 2.72092715592838E-02

 2.65725383466352E-02
 2.47959492196966E-02

 5.47614846540947E-02
 6.36716961304565E-02

 7.24211192915118E-02
 6.82232470714396E-02

 9.06850185871809E-02
 8.78924493521167E-02

 .456944934570223
 .425670047068539

 2.85515051024108E-03
 3.2260992933342E-03

 3.50473501786206E-03
 4.50153389767563E-03

 1.71460096779028E-02
 1.54552663820197E-02

 1.67532376500389E-02
 .01745594811432

 1.84753919260573E-02
 1.70808202895137E-02

 1.69345170475145E-02
 2.04819792344241E-02

 4.97309813741462E-02
 5.57189862445628E-02

 5.83417527542383E-02
 5.61316268518498E-02

 .129070931002644
 .112350783529488

 .144132227609577
 .123267003231351

 4.57024999637441
 4.12331752044509

 2.91139244330789
 2.64027467359503

 .294050289321918
 .323472723330475

 0
 2.50085216537535E-05

 1.5106616456302E-05
 2.50085216537535E-05

 1.61640796082431E-03
 0

 2.13003292033858E-03
 0

 8.76183754465516E-04
 1.21291330020705E-03

 4.00325336092003E-03
 1.68807521162836E-03

 7.79501409145183E-03
 2.02569025395404E-03

 1.98198807906682E-02
 1.97442278456384E-02

 1.93213624476103E-02
 2.02694068003672E-02

 2.69804169909554E-02
 2.78219803398008E-02

 4.64075257537597E-02
 4.29271274186679E-02

 .165085104634468
 .207733285116904

 1.14508152738769E-02
 2.05069877560779E-03

 3.0213232912604E-05
 0

 3.17238945582342E-04
 0

 3.62558794951248E-04
 0

 4.22985260776456E-04
 0

 4.68305110145362E-04
 0

 6.94904356989892E-04
 0

 1.63151457728062E-03
 1.25042608268768E-05

 3.88240042926961E-03
 3.75127824806303E-05

 3.64069456596878E-03
 2.00068173230028E-03

 .392953307261328
 .319121240562722

 1.5106616456302E-05
 1.25042608268768E-05

 0
 3.75127824806303E-05

 7.553308228151E-05
 0

 1.35959548106718E-04
 0

 5.2873157597057E-04
 0

 4.98518343057966E-04
 7.50255649612605E-05

 9.51716836747026E-04
 0

 1.14810285067895E-03
 0

 5.89158041795778E-04
 4.75161911421317E-04

 1.5257682620865E-03
 0

 1.08767638485374E-03
 5.12674693901947E-04

 1.79768735829994E-03
 0

 1.17831608359156E-03
 5.62691737209454E-04

 1.87322044058145E-03
 0

 2.09981968742598E-03
 0

 3.0213232912604E-04
 1.52551982087896E-03

 2.71919096213436E-03
 0

 3.14217622291082E-03
 0

 3.20260268873602E-03
 0

 3.70112103179399E-03
 3.75127824806303E-05

 3.91261366218222E-03
 0

 3.50473501786206E-03
 5.75195998036331E-04

 3.61048133305618E-03
 7.37751388785729E-04

 3.98814674446373E-03
 7.50255649612605E-04

 5.49880839009393E-03
 1.25042608268768E-05

 4.22985260776456E-03
 2.38831381793346E-03

 8.23310596868459E-03
 4.87666172248194E-04

 1.17680542194593E-02
 1.18415350030523E-02

 1.61036531424179E-02
 1.09287239626903E-02

 2.64365787985285E-02
 1.97317235848115E-02

 3.21468798190106E-02
 4.19142822916909E-02

 3.91412432382785E-02
 3.79379273487441E-02

 8.65306990616978E-02
 8.64169465745453E-02

 .121275916911192
 .102159810955583

 2.22973658895017E-02
 1.52802067304434E-02

 3.0213232912604E-05
 0

 1.43512856334869E-03
 1.08787069193828E-03

 1.76747412538733E-03
 8.75298257881373E-04

 1.90645499678531E-02
 1.33170377806237E-02

 .143709242348801
 .152001794611514

 4.5319849368906E-05
 0

 1.5106616456302E-05
 3.75127824806303E-05

 7.553308228151E-05
 6.25213041343838E-05

 7.553308228151E-05
 6.25213041343838E-05

 1.96386013931926E-04
 5.0017043307507E-05

 1.81279397475624E-04
 7.50255649612605E-05

 5.89158041795778E-04
 2.25076694883782E-04

 7.553308228151E-04
 6.50221562997591E-04

 1.13299623422265E-03
 3.50119303152549E-04

 1.02724991902854E-03
 8.8780251870825E-04

 1.32938224815458E-03
 6.75230084651345E-04

 1.16320946713525E-03
 8.37785475400743E-04

 8.00650672184006E-04
 1.17540051772642E-03

 1.63151457728062E-03
 7.25247127958852E-04

 1.97896675577556E-03
 9.12811040362003E-04

 2.00917998868817E-03
 9.87836605323264E-04

 4.15280886383742E-02
 5.07172819138121E-02

 8.91743569415507E-02
 9.35693837675188E-02

 .164390200277478
 .166819343691363

 4.5319849368906E-05
 0

 7.13787627560269E-02
 .074875513831338

 9.29661176720825E-02
 9.19438298600248E-02

 .148618892697099
 .129744210339673

 4.5319849368906E-05
 0

 4.22985260776456E-04
 2.50085216537535E-04

 4.07878644320154E-04
 2.75093738191289E-04

 2.55301818111504E-03
 3.25110781498796E-04

 2.14513953679488E-03
 1.78810929824338E-03

 7.34181559776277E-03
 2.93850129431604E-03

 9.50206175101396E-03
 4.92667876578944E-03

 9.33588896999463E-03
 5.80197702367082E-03

 .116864784905952
 .113438654221426

 1.73726089247473E-03
 9.00306779535127E-04

 6.0426465825208E-05
 2.50085216537535E-05

 3.0213232912604E-05
 6.25213041343838E-05

 2.41705863300832E-04
 2.12572434056905E-04

 1.40491533043609E-03
 6.00204519690084E-04

 1.26291313574685E-02
 2.75093738191289E-04

 1.05746315194114E-04
 0

 6.49584507620986E-04
 2.62589477364412E-04

 1.18738005346534E-02
 1.25042608268768E-05

 2.43367591111025E-02
 2.49460003496191E-02

 1.05746315194114E-04
 0

 4.38091877232758E-04
 5.62691737209454E-04

 1.4955550291739E-03
 7.12742867131975E-04

 3.91261366218222E-03
 5.68943867622893E-03

 1.83847522273195E-02
 1.79811270690488E-02

 .139947694851182
 .122716815754969

 1.5106616456302E-04
 0

 9.36610220290724E-04
 4.00136346460056E-04

 8.3086390509661E-04
 6.37717302170715E-04

 1.40491533043609E-03
 8.00272692920113E-04

 2.67538177441108E-02
 2.09196283633648E-02

 3.20109202709039E-02
 3.33863764077609E-02

 7.78595012157805E-02
 6.65726846422919E-02

 2.2659924684453E-04
 1.25042608268768E-05

 2.2659924684453E-04
 1.25042608268768E-05

 .588508457288157
 .570606934312867

 1.81279397475624E-04
 5.0017043307507E-05

 4.5319849368906E-04
 0

 3.7766541140755E-04
 1.37546869095644E-04

 5.58944808883174E-04
 4.87666172248194E-04

 7.10010973446194E-04
 4.12640607286933E-04

 1.4804484127176E-03
 4.2514486811381E-04

 2.38684540009572E-03
 1.0753664311114E-03

 4.86433049892924E-03
 5.02671285240446E-03

 6.70733770659809E-03
 6.90235197643597E-03

 9.29056912062573E-03
 6.06456650103523E-03

 1.49404436752827E-02
 1.12288262225353E-02

 2.26448180679967E-02
 1.66681796822267E-02

 2.31886562604236E-02
 1.65931541172655E-02

 2.83097992391099E-02
 2.67216053870356E-02

 3.21317732025544E-02
 4.19768035958253E-02

 4.92173564146319E-02
 .048291455313398

 .05192144076031
 5.33931937307638E-02

 5.50334037503082E-02
 5.21802804305567E-02

 7.24211192915118E-02
 6.23212359611538E-02

 8.53070631287374E-02
 9.09559932547015E-02

 .126381953273422
 .129694193296366

 1.82790059121254E-03
 1.4504942559177E-03

 1.05746315194114E-04
 2.87597999018165E-04

 4.07878644320154E-04
 1.12538347441891E-04

 1.31427563169827E-03
 1.05035790945765E-03

 .218079115163176
 .171083296633328

 2.56812479757134E-04
 2.00068173230028E-04

 9.36610220290724E-04
 6.62725823824468E-04

 7.49288176232579E-03
 2.70092033860538E-03

 8.2935324345098E-03
 5.61441311126766E-03

 .201099278266292
 .1619051691864

 .128617732508955
 .097708294101215

 5.74051425339476E-04
 0

 6.94904356989892E-04
 4.12640607286933E-04

 1.08767638485374E-03
 9.12811040362003E-04

 .126261100341772
 9.63828424535661E-02

 .21197604211483
 .186038392582272

 3.17238945582342E-04
 2.87597999018165E-04

 6.34477891164684E-04
 1.12538347441891E-04

 3.92772027863852E-04
 4.75161911421317E-04

 2.56812479757134E-04
 6.25213041343838E-04

 7.25117589902496E-04
 5.25178954728824E-04

 1.84300720766884E-03
 7.75264171266359E-04

 4.45645185460909E-03
 2.07570729726154E-03

 3.65580118242508E-03
 2.93850129431604E-03

 4.69815771790992E-03
 9.81584474909825E-03

 1.74934618563977E-02
 1.21291330020705E-02

 1.97745609412993E-02
 1.56053175119422E-02

 2.58927406061016E-02
 2.65965627787669E-02

 2.86421448011486E-02
 .029422525725641

 3.44884053697375E-02
 2.94975512906023E-02

 6.87048916432615E-02
 5.51562945073534E-02

 4.50026104233236E-02
 4.01386772542744E-02

 6.7979774053359E-04
 0

 4.43228126827901E-02
 4.01386772542744E-02

 7.31160236485017E-03
 4.88916598330881E-03

 8.3086390509661E-04
 0

 2.44727186592092E-03
 1.41298147343707E-03

 4.03346659383263E-03
 3.47618450987174E-03

 9.77095952393613E-02
 .077013742432734

 5.43838192426872E-04
 2.87597999018165E-04

 1.42002194689239E-03
 8.12776953746989E-04

 1.85056051589699E-02
 9.07809336031253E-03

 7.72401299410721E-02
 6.68352741196563E-02

 1.69194104310582E-03
 2.41332233958721E-03

 3.0213232912604E-04
 5.0017043307507E-04

 4.83411726601664E-04
 5.75195998036331E-04

 9.0639698737812E-04
 1.33795590847581E-03

 2.90198102125561E-02
 2.14323030572668E-02

 5.13624959514268E-04
 7.37751388785729E-04

 1.07256976839744E-03
 5.62691737209454E-04

 5.57434147237544E-03
 4.45151685436813E-03

 .021859274012269
 1.56803430769035E-02

 7.85544055727704E-04
 6.00204519690084E-04

 7.85544055727704E-04
 6.00204519690084E-04

 8.21951001387392E-02
 .075688290785085

 1.05746315194114E-03
 3.75127824806303E-04

 1.78258074184364E-03
 3.37615042325672E-04

 4.41113200524018E-03
 2.17574138387656E-03

 3.84312322648323E-02
 3.32613337994922E-02

 3.65126919748819E-02
 3.95384727345843E-02

 .130158607387498
 .133845607890889

 2.68897772922176E-03
 0

 .127469629658276
 .133845607890889

 2.91557697606629E-03
 1.25042608268768E-05

 2.91557697606629E-03
 1.25042608268768E-05

 3.70112103179399E-03
 0

 3.70112103179399E-03
 0

 5.54412823946283E-03
 1.25042608268768E-05

 5.54412823946283E-03
 1.25042608268768E-05

 1.29654046397858
 1.11130367672785

 5.60002272035115E-02
 0

 3.0213232912604E-05
 0

 4.5319849368906E-05
 0

 7.553308228151E-05
 0

 1.5106616456302E-04
 0

 2.41705863300832E-04
 0

 3.7766541140755E-04
 0

 7.25117589902496E-04
 0

 2.31131231781421E-03
 0

 7.34181559776277E-03
 0

 8.09714642057787E-03
 0

 1.32333960157205E-02
 0

 2.33699356578992E-02
 0

 6.0426465825208E-05
 1.25042608268768E-05

 1.5106616456302E-05
 1.25042608268768E-05

 4.5319849368906E-05
 0

 6.49584507620986E-04
 3.37615042325673E-04

 0
 2.50085216537535E-05

 4.5319849368906E-05
 2.50085216537535E-05

 1.35959548106718E-04
 1.25042608268768E-05

 7.553308228151E-05
 2.50085216537535E-04

 3.92772027863852E-04
 2.50085216537535E-05

 3.59386405495424E-02
 2.88598339884316E-02

 3.0213232912604E-05
 0

 3.0213232912604E-05
 1.25042608268768E-05

 9.0639698737812E-05
 1.00034086615014E-04

 4.07878644320154E-04
 3.12606520671919E-04

 8.00650672184006E-04
 5.12674693901947E-04

 1.69194104310582E-03
 8.8780251870825E-04

 3.33856223684274E-03
 1.58804112501335E-03

 5.63476793820064E-03
 .002638399034471

 3.32345562038644E-03
 7.56507780026044E-03

 2.05903182299396E-02
 1.52426939479628E-02

 3.7917607305318E-03
 3.88882511715867E-03

 3.0213232912604E-05
 1.25042608268768E-05

 4.5319849368906E-05
 2.50085216537535E-05

 9.0639698737812E-05
 0

 3.47452178494946E-04
 2.62589477364412E-04

 3.17238945582342E-04
 3.75127824806303E-04

 5.43838192426872E-04
 3.00102259845042E-04

 2.41705863300832E-03
 2.91349277266228E-03

 1.5106616456302E-04
 0

 6.0426465825208E-05
 0

 9.0639698737812E-05
 0

 6.0426465825208E-05
 0

 6.0426465825208E-05
 0

 6.88861710407371E-03
 0

 2.41705863300832E-04
 0

 2.19045938616379E-03
 0

 4.45645185460909E-03
 0

 4.83411726601664E-04
 1.50051129922521E-04

 1.66172781019322E-04
 1.12538347441891E-04

 3.17238945582342E-04
 3.75127824806303E-05

 3.45941516849316E-03
 1.25042608268768E-05

 2.87025712669738E-04
 1.25042608268768E-05

 1.5257682620865E-03
 0

 1.64662119373692E-03
 0

 .242476300740103
 .263039630754179

 1.20852931650416E-04
 1.87563912403151E-04

 2.56812479757134E-04
 1.62555390749398E-04

 5.13624959514268E-04
 3.12606520671919E-04

 3.29475304911947E-02
 3.82130210869354E-02

 5.22537863223486E-02
 4.22393930731897E-02

 7.58201079941797E-02
 7.16494145380038E-02

 8.05635855614586E-02
 .110275076232226

 .94491885934169
 .815002712174173

 4.07878644320154E-04
 3.12606520671919E-04

 2.00917998868817E-03
 5.87700258863208E-04

 2.97600344189149E-03
 1.55052834253272E-03

 4.83411726601664E-03
 3.40115894491048E-03

 .464075257537597
 .395884897778918

 .470616422463176
 .413265820328277

 1.66172781019322E-03
 0

 1.66172781019322E-03
 0

 .362317089087947
 .371739170122219

 .362317089087947
 .371739170122219

 .362317089087947
 .371739170122219

 .139449176508124
 .109799914320805

 7.10010973446194E-04
 3.75127824806303E-04

 1.5106616456302E-05
 1.25042608268768E-05

 1.5106616456302E-05
 1.25042608268768E-05

 1.5106616456302E-05
 1.25042608268768E-05

 1.5106616456302E-05
 1.25042608268768E-05

 6.7979774053359E-04
 3.50119303152549E-04

 6.7979774053359E-04
 3.50119303152549E-04

 3.12706960645451E-03
 1.62555390749398E-03

 4.5319849368906E-05
 0

 4.5319849368906E-05
 0

 7.553308228151E-04
 5.75195998036331E-04

 1.35959548106718E-04
 8.75298257881373E-05

 6.19371274708382E-04
 4.87666172248194E-04

 2.32641893427051E-03
 1.05035790945765E-03

 2.32641893427051E-03
 1.05035790945765E-03

 .116064134233768
 9.57576294122222E-02

 .116064134233768
 9.57576294122222E-02

 0
 3.75127824806303E-05

 3.64522655090567E-02
 3.27236505839365E-02

 7.96118687247115E-02
 6.29964660458051E-02

 5.04560989640487E-03
 2.45083512206784E-03

 5.04560989640487E-03
 2.45083512206784E-03

 1.66172781019322E-04
 8.75298257881373E-05

 1.66172781019322E-04
 1.25042608268768E-04

 2.56812479757134E-04
 2.50085216537535E-04

 4.07878644320154E-04
 1.87563912403151E-04

 6.7979774053359E-04
 2.50085216537535E-04

 8.76183754465516E-04
 2.37580955710658E-04

 6.34477891164684E-04
 5.0017043307507E-04

 9.51716836747026E-04
 3.00102259845042E-04

 9.0639698737812E-04
 5.12674693901947E-04

 2.2659924684453E-04
 3.50119303152549E-04

 2.2659924684453E-04
 3.50119303152549E-04

 2.2659924684453E-04
 3.50119303152549E-04

 6.0426465825208E-04
 6.75230084651345E-04

 6.0426465825208E-04
 6.75230084651345E-04

 6.0426465825208E-04
 6.75230084651345E-04

 4.78879741664773E-03
 1.0753664311114E-03

 4.78879741664773E-03
 1.0753664311114E-03

 4.78879741664773E-03
 1.0753664311114E-03

 8.88269047630557E-03
 7.49005223529918E-03

 8.88269047630557E-03
 7.49005223529918E-03

 8.88269047630557E-03
 7.49005223529918E-03

 1.74788084384351
 1.57972579156348

 .185886915494796
 .214660645614993

 4.16942614193935E-02
 3.30237528437815E-02

 3.0213232912604E-05
 0

 6.34477891164684E-04
 4.6265765059444E-04

 6.0426465825208E-04
 8.00272692920113E-04

 1.5106616456302E-03
 1.88814338485839E-03

 4.07878644320154E-03
 8.62793997054496E-04

 1.90343367349405E-03
 2.8509714685279E-03

 2.62855126339655E-03
 2.76344164273976E-03

 1.69949435133397E-02
 6.47720710832216E-03

 1.33089290980021E-02
 1.69182648987643E-02

 .119100564141485
 .162055220316323

 3.0213232912604E-05
 1.25042608268768E-05

 7.553308228151E-05
 3.75127824806303E-05

 6.0426465825208E-05
 5.0017043307507E-05

 7.553308228151E-05
 1.25042608268768E-04

 6.49584507620986E-04
 2.87597999018165E-04

 4.07878644320154E-04
 6.87734345478222E-04

 8.76183754465516E-04
 8.12776953746989E-04

 1.84300720766884E-03
 1.38797295178332E-03

 2.31131231781421E-03
 1.08787069193828E-03

 2.47748509883353E-03
 3.68875694392864E-03

 2.68897772922176E-03
 4.30146572444561E-03

 4.92475696475445E-03
 4.33897850692623E-03

 8.91290370921818E-03
 1.84312804588163E-02

 1.68438773487767E-02
 1.46925064715802E-02

 9.87972716242151E-03
 2.05319962777316E-02

 1.64662119373692E-02
 3.08104986774243E-02

 5.05769518956991E-02
 .060770707618621

 3.0062166748041E-03
 1.4004772126102E-03

 4.5319849368906E-05
 7.50255649612605E-05

 2.96089682543519E-03
 1.32545164764894E-03

 2.44727186592092E-03
 2.62589477364412E-03

 1.20852931650416E-04
 1.00034086615014E-04

 3.17238945582342E-04
 2.50085216537535E-04

 9.0639698737812E-04
 9.2531530118888E-04

 1.10278300131005E-03
 1.35046016930269E-03

 1.58619472791171E-03
 1.73809225493587E-03

 2.87025712669738E-04
 3.25110781498796E-04

 4.38091877232758E-04
 3.25110781498796E-04

 3.92772027863852E-04
 4.87666172248194E-04

 4.68305110145362E-04
 6.00204519690084E-04

 1.80524066652809E-02
 1.38172082136988E-02

 3.47452178494946E-04
 4.12640607286933E-04

 3.92772027863852E-04
 4.2514486811381E-04

 1.73121824589221E-02
 1.29794227382981E-02

 .58396136573481
 .537133028079318

 1.29463703030508E-02
 7.32749684454978E-03

 1.5106616456302E-05
 1.25042608268768E-05

 3.0213232912604E-05
 0

 3.0213232912604E-05
 5.0017043307507E-05

 9.0639698737812E-05
 1.25042608268768E-05

 2.62855126339655E-03
 1.73809225493587E-03

 1.01516462586349E-02
 5.51437902465265E-03

 2.66178581960041E-02
 2.78094760789739E-02

 3.0213232912604E-05
 0

 2.56812479757134E-04
 7.50255649612605E-05

 7.553308228151E-05
 3.87632085633179E-04

 2.56812479757134E-04
 3.25110781498796E-04

 3.92772027863852E-04
 2.62589477364412E-04

 3.32345562038644E-04
 3.25110781498796E-04

 3.62558794951248E-04
 3.37615042325672E-04

 5.2873157597057E-04
 2.25076694883782E-04

 1.23874254941676E-03
 2.00068173230028E-04

 9.8646205459652E-03
 1.23417054361274E-02

 1.32787158650895E-02
 1.33295420414506E-02

 .119931428046582
 .13215753267926

 4.5319849368906E-05
 0

 7.553308228151E-05
 2.50085216537535E-05

 1.05746315194114E-04
 5.0017043307507E-05

 1.20852931650416E-04
 6.25213041343838E-05

 1.35959548106718E-04
 5.0017043307507E-05

 3.0213232912604E-04
 3.75127824806303E-05

 2.2659924684453E-04
 1.75059651576275E-04

 3.32345562038644E-04
 2.25076694883782E-04

 1.66172781019322E-04
 4.12640607286933E-04

 4.07878644320154E-04
 2.25076694883782E-04

 9.66823453203328E-04
 4.2514486811381E-04

 1.32938224815458E-03
 6.37717302170715E-04

 1.79768735829994E-03
 1.58804112501335E-03

 2.73429757859066E-03
 1.87563912403151E-03

 2.76451081150327E-03
 2.81345868604727E-03

 3.7615474976192E-03
 3.26361207581483E-03

 6.29945906227793E-03
 9.47822970677258E-03

 1.96537080096489E-02
 2.55462048693092E-02

 .039866360828181
 4.05263093399076E-02

 3.88391109091524E-02
 .044740245238565

 3.71622764825029E-03
 1.58804112501335E-03

 3.0213232912604E-05
 2.50085216537535E-05

 2.87025712669738E-04
 1.25042608268768E-04

 1.23874254941676E-03
 4.37649128940687E-04

 2.16024615325119E-03
 1.00034086615014E-03

 1.5106616456302E-04
 1.75059651576275E-04

 6.0426465825208E-05
 0

 1.5106616456302E-05
 6.25213041343838E-05

 1.5106616456302E-05
 6.25213041343838E-05

 6.0426465825208E-05
 5.0017043307507E-05

 7.25117589902496E-04
 2.00068173230028E-04

 9.0639698737812E-05
 0

 1.5106616456302E-04
 1.12538347441891E-04

 4.83411726601664E-04
 8.75298257881373E-05

 .10645632616756
 9.63203211494317E-02

 3.0213232912604E-05
 7.50255649612605E-05

 1.05746315194114E-04
 5.0017043307507E-05

 3.32345562038644E-04
 3.75127824806303E-04

 9.51716836747026E-04
 4.6265765059444E-04

 1.69194104310582E-03
 1.62555390749398E-03

 1.58619472791171E-02
 1.89064423702377E-02

 8.74824158984449E-02
 7.48254967880305E-02

 .133527382857253
 .113951328915328

 3.0213232912604E-05
 1.00034086615014E-04

 2.87025712669738E-04
 1.87563912403151E-04

 4.87490513044865E-02
 4.34898191558774E-02

 8.44610926071845E-02
 7.01739117604324E-02

 9.60780806620807E-03
 1.12788432658428E-02

 6.0426465825208E-05
 2.62589477364412E-04

 5.2873157597057E-04
 5.25178954728824E-04

 1.42002194689239E-03
 8.62793997054496E-04

 7.5986280775199E-03
 9.6282808366951E-03

 .105716101961201
 .091218582732066

 4.83411726601664E-04
 1.25042608268768E-05

 5.74051425339476E-04
 5.12674693901947E-04

 1.90343367349405E-03
 1.03785364863077E-03

 6.66201785722918E-03
 3.43867172739111E-03

 8.58055814717953E-03
 7.71512893018296E-03

 1.41397930030987E-02
 1.31544823898744E-02

 7.33728361282588E-02
 6.53472670812579E-02

 1.45023517980499E-03
 5.50187476382577E-04

 5.74051425339476E-04
 1.75059651576275E-04

 8.76183754465516E-04
 3.75127824806303E-04

 3.61048133305618E-03
 2.52586068702911E-03

 5.89158041795778E-04
 8.5028973622762E-04

 3.0213232912604E-03
 1.67557095080149E-03

 5.95049622213736E-02
 5.20302293006342E-02

 6.17860613062752E-03
 4.00136346460056E-03

 2.03788255995514E-02
 1.43548914292545E-02

 3.29475304911947E-02
 3.36739744067791E-02

 .107045484209356
 .102797528257754

 1.92307227488724E-02
 1.81186739381444E-02

 4.5319849368906E-05
 1.25042608268768E-05

 1.20852931650416E-04
 2.87597999018165E-04

 4.98518343057966E-04
 2.87597999018165E-04

 1.5408748785428E-03
 9.75332344496387E-04

 1.70251567462524E-02
 1.65556413347848E-02

 2.14513953679488E-03
 1.20040903938017E-03

 1.5106616456302E-05
 3.75127824806303E-05

 6.34477891164684E-04
 5.62691737209454E-04

 1.4955550291739E-03
 6.00204519690084E-04

 5.84626056858887E-03
 .004464021115195

 3.0213232912604E-05
 2.50085216537535E-05

 4.5319849368906E-05
 5.0017043307507E-05

 9.0639698737812E-05
 5.0017043307507E-05

 4.38091877232758E-04
 3.25110781498796E-04

 8.45970521552912E-04
 3.87632085633179E-04

 4.39602538878388E-03
 3.62623563979426E-03

 1.31276497005264E-02
 .014404908472562

 6.0426465825208E-05
 0

 1.35959548106718E-04
 2.00068173230028E-04

 2.17535276970749E-03
 2.00068173230028E-03

 2.44727186592092E-03
 3.48868877069862E-03

 8.3086390509661E-03
 8.7154697963331E-03

 4.18151143510439E-02
 3.66624927444026E-02

 1.5106616456302E-04
 1.00034086615014E-04

 2.76451081150327E-03
 8.5028973622762E-04

 3.88995373749776E-02
 .03571216892156

 1.01214330257223E-02
 6.76480510734033E-03

 1.66172781019322E-04
 1.25042608268768E-04

 2.56812479757134E-04
 5.50187476382577E-04

 8.61077138009214E-04
 9.6282808366951E-04

 1.37470209752348E-03
 8.8780251870825E-04

 1.99407337223186E-03
 4.6265765059444E-04

 2.56812479757134E-03
 1.72558799410899E-03

 2.90047035960998E-03
 2.05069877560779E-03

 3.94282689509482E-03
 2.97601407679667E-03

 1.96386013931926E-04
 2.50085216537535E-04

 2.87025712669738E-04
 2.37580955710658E-04

 2.2659924684453E-04
 3.50119303152549E-04

 4.07878644320154E-04
 2.62589477364412E-04

 3.32345562038644E-04
 4.2514486811381E-04

 5.89158041795778E-04
 2.87597999018165E-04

 8.45970521552912E-04
 4.12640607286933E-04

 1.05746315194114E-03
 7.50255649612605E-04

 1.32938224815458E-03
 6.50221562997591E-04

 4.38091877232758E-04
 2.37580955710658E-04

 8.91290370921818E-04
 4.12640607286933E-04

 9.51716836747026E-04
 8.25281214573866E-04

 9.51716836747026E-04
 8.25281214573866E-04

 8.53523829781063E-03
 1.67307009863611E-02

 8.53523829781063E-03
 1.67307009863611E-02

 1.05746315194114E-02
 8.81550388294811E-03

 4.33559892295867E-03
 5.43935345969139E-03

 6.0426465825208E-05
 3.75127824806303E-05

 7.553308228151E-05
 1.12538347441891E-04

 6.64691124077288E-04
 5.0017043307507E-04

 9.0639698737812E-04
 6.62725823824468E-04

 1.14810285067895E-03
 9.2531530118888E-04

 7.553308228151E-04
 1.46299851674458E-03

 7.25117589902496E-04
 1.73809225493587E-03

 3.7615474976192E-03
 1.78810929824338E-03

 1.81279397475624E-04
 1.37546869095644E-04

 1.82790059121254E-03
 6.37717302170715E-04

 1.75236750893103E-03
 1.01284512697702E-03

 2.47748509883353E-03
 1.58804112501335E-03

 3.17238945582342E-04
 3.00102259845042E-04

 4.22985260776456E-04
 3.50119303152549E-04

 4.68305110145362E-04
 4.2514486811381E-04

 1.26895578232937E-03
 5.12674693901947E-04

 .032116666586098
 3.50119303152549E-02

 .032116666586098
 3.50119303152549E-02

 4.5319849368906E-05
 6.25213041343838E-05

 9.0639698737812E-05
 6.25213041343838E-05

 1.96386013931926E-04
 1.25042608268768E-04

 3.62558794951248E-04
 1.75059651576275E-04

 9.8193006965963E-04
 5.62691737209454E-04

 2.2810990849016E-03
 1.06286217028452E-03

 2.22067261907639E-03
 2.56337346950974E-03

 4.78879741664773E-03
 2.60088625199037E-03

 2.11492630388228E-02
 .027796971818147

 .740662298236031
 .594477568231375

 .740662298236031
 .594477568231375

 9.0639698737812E-05
 1.37546869095644E-04

 1.26895578232937E-03
 9.6282808366951E-04

 5.2873157597057E-03
 1.28793886516831E-03

 4.03346659383263E-03
 2.95100555514291E-03

 1.26291313574685E-02
 8.5028973622762E-03

 1.65417450196507E-02
 1.29544142166443E-02

 .027463828717557
 2.05445005385585E-02

 4.60147537258959E-02
 4.15766672493652E-02

 4.85375586740983E-02
 4.75787124462661E-02

 5.02899261830293E-02
 5.09548628695228E-02

 .117635222345224
 9.69830469732561E-02

 .173303103986697
 .131244721638898

 .237566650391805
 .178798425563511

 6.87351048761741E-03
 7.90269284258611E-03

 6.87351048761741E-03
 7.90269284258611E-03

 6.0426465825208E-05
 2.00068173230028E-04

 5.43838192426872E-04
 4.6265765059444E-04

 4.38091877232758E-04
 5.62691737209454E-04

 6.19371274708382E-04
 7.37751388785729E-04

 8.15757288640308E-04
 6.87734345478222E-04

 6.34477891164684E-04
 9.00306779535127E-04

 7.40224206358798E-04
 8.62793997054496E-04

 8.61077138009214E-04
 9.12811040362003E-04

 1.02724991902854E-03
 1.25042608268768E-03

 1.13299623422265E-03
 1.32545164764894E-03

 1.57259877310104E-02
 .0134920974322

 1.78258074184364E-03
 4.60156798429065E-03

 1.66172781019322E-04
 2.62589477364412E-04

 2.2659924684453E-04
 3.75127824806303E-04

 9.0639698737812E-05
 4.87666172248194E-04

 5.43838192426872E-04
 9.87836605323264E-04

 7.553308228151E-04
 2.48834790454847E-03

 1.39434069891667E-02
 8.89052944790937E-03

 5.58944808883174E-04
 4.37649128940687E-04

 7.85544055727704E-04
 5.25178954728824E-04

 3.08174975708561E-03
 1.02534938780389E-03

 9.51716836747026E-03
 6.90235197643597E-03

 6.50339838443801E-02
 6.54347969070461E-02

 1.51670429221272E-02
 1.12538347441891E-02

 3.7766541140755E-04
 2.25076694883782E-04

 1.16320946713525E-03
 6.25213041343838E-04

 1.72215427601843E-03
 3.62623563979426E-04

 .011904013767566
 .010040921443982

 4.98669409222529E-02
 .054180962162857

 1.99407337223186E-03
 1.36296443012957E-03

 7.64394792688881E-03
 4.66408928842503E-03

 1.63453590057188E-02
 2.08320985375767E-02

 2.38835606174135E-02
 2.73218099067257E-02

 .682864383674219
 .763072516960154

 .11393410131343
 .107686694241063

 .11393410131343
 .107686694241063

 3.0213232912604E-05
 0

 1.5106616456302E-04
 0

 3.47452178494946E-04
 8.75298257881373E-05

 3.47452178494946E-04
 1.87563912403151E-04

 6.34477891164684E-04
 7.62759910439482E-04

 1.10278300131005E-03
 3.87632085633179E-04

 1.17831608359156E-03
 5.50187476382577E-04

 1.28406239878567E-03
 5.37683215555701E-04

 2.52280494820243E-03
 2.25076694883782E-04

 1.88832705703775E-03
 1.08787069193828E-03

 1.96386013931926E-03
 1.13788773524579E-03

 2.97600344189149E-03
 9.87836605323264E-04

 2.44727186592092E-03
 1.68807521162836E-03

 7.93097363955855E-03
 7.81516301679797E-03

 8.2935324345098E-03
 7.87768432093236E-03

 9.35099558645094E-03
 8.60293144889121E-03

 1.16169880548962E-02
 9.87836605323264E-03

 .011904013767566
 1.12913475266697E-02

 1.25989181245559E-02
 1.47925405581952E-02

 1.28255173714004E-02
 1.65931541172655E-02

 2.25390717528026E-02
 2.31954038338564E-02

 .481070201050937
 .582861109923206

 2.16024615325119E-03
 1.85063060237776E-03

 1.5106616456302E-05
 1.25042608268768E-05

 3.0213232912604E-05
 2.50085216537535E-05

 1.5106616456302E-05
 7.50255649612605E-05

 3.47452178494946E-04
 4.50153389767563E-04

 1.75236750893103E-03
 1.28793886516831E-03

 .386472568801574
 .465145998498989

 4.5319849368906E-05
 6.25213041343838E-05

 .013157862933439
 1.17540051772642E-02

 1.30219033853323E-02
 1.54802749036734E-02

 2.87025712669738E-02
 2.80470570346846E-02

 3.31439165051266E-02
 5.03421540890058E-02

 4.18604342004128E-02
 5.07422904354659E-02

 5.15588819653587E-02
 6.99363308047217E-02

 .204981678695562
 .238781364750039

 4.12410629257045E-03
 3.52620155317925E-03

 1.66172781019322E-04
 1.12538347441891E-04

 1.81279397475624E-04
 1.75059651576275E-04

 3.32345562038644E-04
 1.62555390749398E-04

 4.07878644320154E-04
 2.37580955710658E-04

 4.22985260776456E-04
 2.62589477364412E-04

 6.94904356989892E-04
 6.25213041343838E-04

 8.00650672184006E-04
 1.03785364863077E-03

 1.11788961776635E-03
 9.12811040362003E-04

 8.83132798035415E-02
 .112338279268661

 1.23421056447987E-02
 1.40547891694095E-02

 1.52274693879524E-02
 1.60429666408829E-02

 2.13003292033858E-02
 2.68091352128238E-02

 3.94433755674045E-02
 5.54313882455447E-02

 8.78600813098524E-02
 7.25247127958852E-02

 3.12706960645451E-03
 1.86313486320464E-03

 1.66172781019322E-04
 1.00034086615014E-04

 3.0213232912604E-04
 7.50255649612605E-05

 2.71919096213436E-04
 1.62555390749398E-04

 2.71919096213436E-04
 2.25076694883782E-04

 4.83411726601664E-04
 4.00136346460056E-04

 1.63151457728062E-03
 9.00306779535127E-04

 3.0364299077167E-03
 2.58838199116349E-03

 2.11492630388228E-04
 8.75298257881373E-05

 1.96386013931926E-04
 2.50085216537535E-04

 1.17831608359156E-03
 1.16289625689954E-03

 1.45023517980499E-03
 1.08787069193828E-03

 .068478292396417
 5.92952048410496E-02

 6.34477891164684E-04
 1.25042608268768E-05

 7.25117589902496E-04
 2.50085216537535E-05

 3.17238945582342E-03
 2.20074990553031E-03

 5.78583410276366E-03
 3.52620155317925E-03

 1.03933521219358E-02
 5.62691737209454E-03

 1.43361790170306E-02
 1.02034768347314E-02

 3.34309422177963E-02
 3.77003463930334E-02

 9.1092897231501E-03
 6.13959206599649E-03

 5.58944808883174E-04
 2.12572434056905E-04

 7.553308228151E-04
 5.12674693901947E-04

 2.46237848237723E-03
 2.20074990553031E-03

 5.3326356090746E-03
 3.21359503250733E-03

 4.10899967611414E-03
 .002638399034471

 5.13624959514268E-04
 3.25110781498796E-04

 5.89158041795778E-04
 4.37649128940687E-04

 8.15757288640308E-04
 5.25178954728824E-04

 2.19045938616379E-03
 1.35046016930269E-03

 2.82812477340075
 3.06029279476982

 .152259587263068
 .167932222904955

 3.30230635734762E-02
 4.27520677670916E-02

 1.5106616456302E-05
 1.25042608268768E-05

 1.5106616456302E-05
 2.50085216537535E-05

 1.5106616456302E-05
 2.50085216537535E-05

 6.0426465825208E-05
 3.75127824806303E-05

 6.0426465825208E-05
 5.0017043307507E-05

 9.66823453203328E-04
 5.0017043307507E-04

 4.47155847106539E-03
 6.91485623726285E-03

 2.74185088681881E-02
 3.51869899668312E-02

 .048069253563953
 4.58531244521571E-02

 0
 8.75298257881373E-05

 1.01214330257223E-03
 2.50085216537535E-04

 5.49880839009393E-03
 1.53802408170584E-03

 5.89158041795778E-03
 1.7756050374165E-03

 2.07413843945026E-02
 1.84938017629507E-02

 1.49253370588264E-02
 2.37080785277583E-02

 5.19667606096789E-03
 4.06388476873495E-03

 2.2659924684453E-04
 2.50085216537535E-05

 3.62558794951248E-04
 1.25042608268768E-04

 3.7766541140755E-04
 2.62589477364412E-04

 7.25117589902496E-04
 5.75195998036331E-04

 6.64691124077288E-04
 7.87768432093236E-04

 9.21503603834422E-04
 9.37819562015757E-04

 1.91854028995035E-03
 1.35046016930269E-03

 6.54418624887003E-02
 7.31124130547484E-02

 2.2659924684453E-04
 2.50085216537535E-05

 7.10010973446194E-04
 5.25178954728824E-04

 4.81901064956034E-03
 .002638399034471

 5.96862416188492E-02
 6.99238265438948E-02

 5.2873157597057E-04
 2.1507328622228E-03

 5.2873157597057E-04
 2.1507328622228E-03

 5.20574003084167E-02
 4.75411996637854E-02

 2.66178581960041E-02
 2.24076354017632E-02

 3.0213232912604E-05
 0

 3.30834900393014E-03
 5.25178954728824E-03

 2.32792959591614E-02
 1.71558458544749E-02

 1.88530573374649E-02
 2.05194920169048E-02

 1.35959548106718E-04
 6.25213041343838E-05

 5.2873157597057E-04
 5.0017043307507E-05

 1.07256976839744E-03
 6.00204519690084E-04

 1.01214330257223E-03
 7.00238606305098E-04

 7.85544055727704E-04
 9.12811040362003E-04

 6.0275399660645E-03
 2.95100555514291E-03

 9.29056912062573E-03
 1.52426939479628E-02

 6.58648477494767E-03
 4.61407224511752E-03

 2.41705863300832E-04
 2.25076694883782E-04

 4.68305110145362E-04
 3.37615042325672E-04

 5.58944808883174E-04
 3.25110781498796E-04

 1.57108811145541E-03
 1.20040903938017E-03

 1.55598149499911E-03
 1.47550277757146E-03

 2.19045938616379E-03
 1.05035790945765E-03

 6.47469581317104E-02
 4.53904668015626E-02

 2.2659924684453E-04
 1.25042608268768E-04

 1.5106616456302E-05
 1.25042608268768E-05

 2.11492630388228E-04
 1.12538347441891E-04

 1.07256976839744E-02
 6.0770707618621E-03

 3.0213232912604E-05
 0

 3.0213232912604E-05
 0

 1.5106616456302E-05
 1.25042608268768E-05

 0
 2.50085216537535E-05

 3.0213232912604E-05
 1.25042608268768E-05

 3.0213232912604E-05
 2.50085216537535E-05

 6.0426465825208E-05
 0

 6.0426465825208E-05
 0

 1.5106616456302E-05
 3.75127824806303E-05

 9.0639698737812E-05
 1.25042608268768E-05

 7.553308228151E-05
 3.75127824806303E-05

 1.20852931650416E-04
 0

 1.05746315194114E-04
 5.0017043307507E-05

 7.85544055727704E-04
 2.87597999018165E-04

 6.64691124077288E-04
 6.00204519690084E-04

 4.62262463562841E-03
 3.37615042325672E-04

 3.98814674446373E-03
 4.63908076677128E-03

 6.94904356989892E-04
 1.37546869095644E-03

 3.0213232912604E-05
 0

 0
 2.50085216537535E-04

 5.58944808883174E-04
 6.25213041343838E-05

 1.05746315194114E-04
 1.06286217028452E-03

 5.19667606096789E-02
 3.71626631774777E-02

 7.553308228151E-05
 3.75127824806303E-05

 1.20852931650416E-04
 1.25042608268768E-05

 1.35959548106718E-04
 1.25042608268768E-05

 3.0213232912604E-05
 1.25042608268768E-04

 3.47452178494946E-04
 2.25076694883782E-04

 7.19074943319975E-03
 4.7891318966938E-03

 4.40660002030329E-02
 .031960890673497

 1.13299623422265E-03
 6.50221562997591E-04

 1.20852931650416E-04
 1.62555390749398E-04

 1.01214330257223E-03
 4.87666172248194E-04

 .18268431280606
 .16646922438821

 4.51083567385178E-02
 3.57996987473482E-02

 1.5106616456302E-05
 1.25042608268768E-05

 1.5106616456302E-04
 0

 4.22985260776456E-04
 2.50085216537535E-05

 1.81279397475624E-04
 2.75093738191289E-04

 9.66823453203328E-03
 5.42684919886451E-03

 9.66823453203328E-03
 7.72763319100984E-03

 9.72866099785849E-03
 8.19029084160428E-03

 1.52727892373213E-02
 1.41423189951976E-02

 3.41107399583299E-02
 2.90348936400078E-02

 3.0213232912604E-05
 0

 3.0213232912604E-05
 5.0017043307507E-05

 1.5106616456302E-04
 1.12538347441891E-04

 3.47452178494946E-04
 5.0017043307507E-05

 2.2659924684453E-04
 4.87666172248194E-04

 5.74051425339476E-04
 3.00102259845042E-04

 5.2873157597057E-04
 4.75161911421317E-04

 6.19371274708382E-04
 9.37819562015757E-04

 1.13299623422265E-03
 1.11287921359203E-03

 4.84922388247294E-03
 2.36330529627971E-03

 2.56208215098882E-02
 2.31453867905489E-02

 .031164949749351
 .033173803973704

 2.56812479757134E-04
 3.00102259845042E-04

 4.22985260776456E-04
 1.75059651576275E-04

 9.8193006965963E-04
 5.75195998036331E-04

 1.02724991902854E-03
 7.50255649612605E-04

 6.23903259645273E-03
 6.7773093681672E-03

 2.22369394236765E-02
 2.45958810464666E-02

 7.23002663598614E-02
 6.84608280271503E-02

 6.7979774053359E-04
 1.08787069193828E-03

 3.12706960645451E-03
 1.97567321064653E-03

 3.0213232912604E-03
 2.98851833762355E-03

 3.92772027863852E-03
 4.07638902956182E-03

 9.01865002441229E-03
 9.31567431602318E-03

 9.59270144975177E-03
 1.17289966556104E-02

 4.29330039688103E-02
 3.72877057857465E-02

 2.31221871480158
 2.57057592374602

 .101561782435718
 .110375110318841

 3.0213232912604E-05
 0

 7.553308228151E-05
 5.0017043307507E-05

 2.16024615325119E-03
 2.01318599312716E-03

 8.50502506489802E-03
 4.33897850692623E-03

 1.79013405007179E-02
 2.40081807876034E-02

 2.50316634680924E-02
 2.25576865316857E-02

 4.78577609335647E-02
 5.74070614561912E-02

 5.27523046654066E-02
 5.66317972849248E-02

 3.0213232912604E-05
 0

 7.553308228151E-05
 3.75127824806303E-05

 7.553308228151E-05
 2.50085216537535E-04

 6.0426465825208E-05
 3.12606520671919E-04

 7.40224206358798E-04
 6.25213041343838E-04

 3.7464408811629E-03
 1.53802408170584E-03

 3.88240042926961E-03
 6.68977954237907E-03

 1.43663922499432E-02
 7.69012040852921E-03

 2.97751410353712E-02
 3.94884556912768E-02

 .187367363907514
 .148825712361487

 1.5106616456302E-05
 1.25042608268768E-05

 6.0426465825208E-05
 2.50085216537535E-05

 1.05746315194114E-04
 3.75127824806303E-05

 6.0426465825208E-05
 1.12538347441891E-04

 1.20852931650416E-04
 1.75059651576275E-04

 4.22985260776456E-04
 0

 3.0213232912604E-04
 1.12538347441891E-04

 9.21503603834422E-04
 2.50085216537535E-05

 8.00650672184006E-04
 3.25110781498796E-04

 7.10010973446194E-04
 5.62691737209454E-04

 1.31427563169827E-03
 8.62793997054496E-04

 3.36877546975535E-03
 4.75161911421317E-04

 .179164471171742
 .146099783501228

 3.0213232912604E-05
 0

 3.0213232912604E-05
 0

 1.05746315194114E-03
 7.75264171266359E-04

 1.5106616456302E-05
 1.25042608268768E-05

 3.0213232912604E-05
 1.25042608268768E-05

 1.01214330257223E-03
 7.50255649612605E-04

 1.5106616456302E-05
 1.25042608268768E-05

 1.5106616456302E-05
 1.25042608268768E-05

 .011752947603003
 1.38797295178332E-02

 4.5319849368906E-05
 0

 4.5319849368906E-05
 0

 3.0213232912604E-05
 2.50085216537535E-05

 1.05746315194114E-04
 0

 1.35959548106718E-04
 0

 4.22985260776456E-04
 1.62555390749398E-04

 1.09674035472752E-02
 1.36921656054301E-02

 5.61059735187056E-02
 .053268151122495

 3.0213232912604E-05
 2.50085216537535E-05

 7.553308228151E-05
 5.0017043307507E-05

 5.60002272035115E-02
 5.31931255575337E-02

 .64997727964885
 .748617591444285

 6.0426465825208E-05
 3.75127824806303E-05

 2.71919096213436E-04
 2.37580955710658E-04

 4.98518343057966E-04
 7.50255649612605E-05

 4.68305110145362E-04
 1.25042608268768E-04

 4.07878644320154E-04
 2.62589477364412E-04

 6.49584507620986E-04
 6.50221562997591E-04

 1.75236750893103E-03
 5.12674693901947E-04

 1.45023517980499E-03
 1.32545164764894E-03

 2.82191595403721E-02
 1.57553686418647E-02

 .020257972667901
 2.94975512906023E-02

 2.93672623910511E-02
 3.11856265022306E-02

 3.17390011746905E-02
 4.65533630584622E-02

 .271088232308339
 .285284710765193

 .263746416710577
 .337114871892597

 6.34477891164684E-04
 5.0017043307507E-05

 1.35959548106718E-04
 0

 4.98518343057966E-04
 5.0017043307507E-05

 .109266156828432
 .106411259636721

 1.5106616456302E-04
 0

 .109115090663869
 .106411259636721

 3.31439165051266E-02
 2.15198328830549E-02

 1.05746315194114E-04
 5.0017043307507E-05

 .014804484127176
 1.08912111802097E-02

 1.82336860627565E-02
 1.05786046595377E-02

 8.93707429554826E-02
 .111763083270624

 1.05746315194114E-04
 1.37546869095644E-04

 1.05746315194114E-04
 1.75059651576275E-04

 2.2659924684453E-04
 1.12538347441891E-04

 7.553308228151E-05
 3.12606520671919E-04

 3.47452178494946E-04
 4.75161911421317E-04

 1.78258074184364E-03
 1.22541756103392E-03

 1.01667528750912E-02
 1.09162197018634E-02

 1.84300720766884E-02
 2.65465457354594E-02

 5.81302601238501E-02
 7.18619869720607E-02

 5.13927091843394E-02
 5.07422904354659E-02

 1.5106616456302E-05
 2.12572434056905E-04

 6.19371274708382E-03
 5.57690032878703E-03

 4.51838898207993E-02
 4.49528176726219E-02

 3.0213232912604E-05
 2.00068173230028E-04

 3.0213232912604E-05
 2.00068173230028E-04

 .469030227735264
 .588438010251993

 2.56812479757134E-04
 3.75127824806303E-05

 2.11492630388228E-04
 1.87563912403151E-04

 5.13624959514268E-04
 4.37649128940687E-04

 4.68305110145362E-04
 7.62759910439482E-04

 7.40224206358798E-04
 5.87700258863208E-04

 9.21503603834422E-04
 4.37649128940687E-04

 1.97896675577556E-03
 1.37546869095644E-04

 1.46534179626129E-03
 8.25281214573866E-04

 4.44134523815279E-03
 4.71410633173254E-03

 1.91551896665909E-02
 1.92190488909096E-02

 1.33089290980021E-02
 2.45458640031591E-02

 3.74493021951727E-02
 .038250533869416

 5.61210801351619E-02
 5.87325131038401E-02

 .092543132411306
 .123179473405563

 .239454977448843
 .316382807441636

 3.0213232912604E-04
 0

 3.0213232912604E-04
 0

 4.70571102613807E-02
 4.25019825505541E-02

 1.5106616456302E-04
 1.37546869095644E-04

 2.20556600262009E-03
 8.25281214573866E-04

 2.44727186592092E-03
 2.77594590356664E-03

 3.35366885329904E-03
 2.18824564470343E-03

 1.53332157031465E-02
 1.57053515985572E-02

 2.35663216718311E-02
 2.08696113200573E-02

 1.46534179626129E-03
 1.15039199607266E-03

 2.41705863300832E-04
 1.12538347441891E-04

 2.56812479757134E-04
 2.00068173230028E-04

 3.32345562038644E-04
 2.62589477364412E-04

 6.34477891164684E-04
 5.75195998036331E-04

 5.74051425339476E-04
 3.00102259845042E-04

 2.41705863300832E-04
 1.25042608268768E-04

 3.32345562038644E-04
 1.75059651576275E-04

 4.16942614193935E-03
 3.82630381302429E-03

 2.56812479757134E-04
 1.87563912403151E-04

 1.16320946713525E-03
 1.37546869095644E-03

 2.74940419504696E-03
 2.26327120966469E-03

 6.42031199392835E-02
 3.69750992650746E-02

 1.81279397475624E-04
 2.87597999018165E-04

 4.5319849368906E-04
 1.37546869095644E-04

 5.2873157597057E-04
 1.37546869095644E-04

 1.56957744980978E-02
 8.76548683964061E-03

 2.08471307096968E-02
 1.26918247392799E-02

 2.64970052643537E-02
 1.49550959489446E-02

 5.98222011669559E-03
 5.60190885044079E-03

 3.7766541140755E-04
 1.37546869095644E-04

 3.17238945582342E-04
 2.25076694883782E-04

 1.23874254941676E-03
 8.75298257881373E-04

 2.2962057013579E-03
 1.52551982087896E-03

 1.75236750893103E-03
 2.83846720770102E-03

 1.76898478703296E-02
 2.16448754913237E-02

 3.17238945582342E-04
 2.75093738191289E-04

 2.19045938616379E-03
 1.87563912403151E-03

 1.51821495385835E-02
 1.94941426291009E-02

 4.15431952548305E-03
 3.53870581400612E-03

 4.98518343057966E-04
 3.00102259845042E-04

 6.49584507620986E-04
 8.75298257881373E-04

 3.0062166748041E-03
 2.36330529627971E-03

 .1560513479936
 .210384188412201

 7.40224206358798E-04
 5.62691737209454E-04

 8.76183754465516E-04
 7.00238606305098E-04

 8.3086390509661E-04
 1.02534938780389E-03

 1.84300720766884E-03
 1.20040903938017E-03

 1.94875352286296E-03
 1.27543460434143E-03

 .149812315397147
 .205620065037161

 3.82197396344441E-03
 3.90132937798555E-03

 3.82197396344441E-03
 3.90132937798555E-03

 3.29324238747384E-03
 4.92667876578944E-03

 3.29324238747384E-03
 4.92667876578944E-03

 .189965701937998
 .224313934973342

 5.83719659871509E-02
 8.10151058973345E-02

 .131593735950847
 .143298829076008

 1.51519363056709E-02
 1.83562548938551E-02

 1.23874254941676E-02
 1.83562548938551E-02

 6.0426465825208E-05
 2.50085216537535E-05

 1.5106616456302E-05
 1.12538347441891E-04

 1.20852931650416E-04
 8.75298257881373E-05

 1.21910394802357E-02
 1.81311781989713E-02

 1.5106616456302E-04
 0

 1.5106616456302E-04
 0

 2.61344464694025E-03
 0

 2.61344464694025E-03
 0

 4.90058637842437E-02
 4.40275023714331E-02

 4.90058637842437E-02
 4.40275023714331E-02

 4.90058637842437E-02
 4.40275023714331E-02

 .781208456804745
 .758771051235709

 .781208456804745
 .758771051235709

 1.85811382412515E-03
 1.18790477855329E-03

 3.0213232912604E-05
 1.25042608268768E-05

 2.11492630388228E-04
 3.37615042325672E-04

 5.58944808883174E-04
 3.00102259845042E-04

 1.05746315194114E-03
 5.37683215555701E-04

 .364175202912072
 .360672899290433

 7.553308228151E-05
 5.0017043307507E-05

 2.71919096213436E-04
 4.12640607286933E-04

 6.19371274708382E-04
 2.00068173230028E-04

 7.40224206358798E-04
 1.75059651576275E-04

 4.38091877232758E-04
 5.0017043307507E-04

 2.2962057013579E-03
 2.02569025395404E-03

 9.62291468266437E-03
 8.47788884062244E-03

 2.73882956352755E-02
 2.37956083535465E-02

 3.81593131686188E-02
 .034499255621353

 6.17558480733626E-02
 .056919395283943

 6.01696533454509E-02
 .068285768375574

 .162637832768547
 .165331336652965

 2.54395421124126E-02
 1.70558117678599E-02

 9.0639698737812E-05
 5.0017043307507E-05

 1.05746315194114E-04
 8.75298257881373E-05

 3.32345562038644E-04
 7.50255649612605E-05

 5.46859515718132E-03
 1.18790477855329E-03

 7.65905454334511E-03
 5.88950684945895E-03

 1.17831608359156E-02
 9.76582770579075E-03

 8.77694416111146E-03
 5.66443015457517E-03

 1.20852931650416E-04
 1.00034086615014E-04

 2.87025712669738E-04
 1.25042608268768E-04

 8.00650672184006E-04
 9.75332344496387E-04

 1.5408748785428E-03
 1.05035790945765E-03

 6.0275399660645E-03
 3.41366320573736E-03

 2.44727186592092E-03
 1.01284512697702E-03

 2.11492630388228E-04
 2.12572434056905E-04

 9.0639698737812E-04
 4.75161911421317E-04

 1.32938224815458E-03
 3.25110781498796E-04

 5.43536060097746E-02
 4.64158161893665E-02

 2.14513953679488E-03
 1.2379218218608E-03

 4.63773125208471E-03
 4.43901259354125E-03

 1.14810285067895E-02
 8.91553796956313E-03

 3.60897067141055E-02
 3.18233438044013E-02

 .324157775919328
 .326761343927943

 1.08767638485374E-02
 1.21541415237242E-02

 .313281012070791
 .314607202404219

 .622105572286973
 .759133674799688

 .622105572286973
 .759133674799688

 .522598289689311
 .65114687829878

 4.5319849368906E-05
 0

 7.553308228151E-05
 2.50085216537535E-05

 1.5106616456302E-04
 7.50255649612605E-05

 3.0213232912604E-04
 1.62555390749398E-04

 3.0213232912604E-04
 2.37580955710658E-04

 3.47452178494946E-04
 3.25110781498796E-04

 4.22985260776456E-04
 4.6265765059444E-04

 1.32938224815458E-03
 1.62555390749398E-04

 8.76183754465516E-04
 6.25213041343838E-04

 1.72215427601843E-03
 1.15039199607266E-03

 2.70408434567806E-03
 1.32545164764894E-03

 3.06664314062931E-03
 2.30078399214532E-03

 2.2810990849016E-03
 3.07604816341168E-03

 1.38678739068852E-02
 9.76582770579075E-03

 1.26744512068374E-02
 .015417753599539

 1.47742708942634E-02
 1.57928814243453E-02

 1.80524066652809E-02
 1.44549255158695E-02

 1.68438773487767E-02
 .018868929587757

 1.99709469552312E-02
 1.64305987265161E-02

 2.94881153227015E-02
 4.40024938497793E-02

 5.89158041795778E-02
 6.62600781216199E-02

 6.91883033698631E-02
 7.38501644435341E-02

 6.20730870189449E-02
 8.34784452802292E-02

 7.05630054673866E-02
 9.12685997753735E-02

 .122559979309978
 .191627797171886

 .017010050129796
 9.85335753157889E-03

 6.7979774053359E-04
 3.37615042325672E-04

 2.64365787985285E-03
 9.87836605323264E-04

 4.97007681412336E-03
 2.51335642620223E-03

 8.71651769528625E-03
 6.01454945772772E-03

 8.24972324678652E-02
 9.81334389693288E-02

 .019109869817222
 2.56212304342705E-02

 6.33873626506432E-02
 7.25122085350583E-02

 .797297003330707
 .545998549005574

 .679299222190532
 .459344021475318

 .154178127553018
 .12915651008081

 4.5319849368906E-05
 2.50085216537535E-05

 1.20852931650416E-03
 9.37819562015757E-04

 5.03050327994856E-03
 3.25110781498796E-03

 6.40520537747205E-03
 2.35080103545283E-03

 5.12114297868638E-03
 3.85131233467804E-03

 1.66777045677574E-02
 1.27168332609337E-02

 2.02428660514447E-02
 1.37421826487376E-02

 3.36122216152719E-02
 2.02318940178866E-02

 6.58346345165641E-02
 7.20495508844639E-02

 .360730894360035
 .180148885732813

 7.553308228151E-05
 5.0017043307507E-05

 1.35959548106718E-04
 1.00034086615014E-04

 1.35959548106718E-04
 1.25042608268768E-04

 1.20852931650416E-04
 2.12572434056905E-04

 3.62558794951248E-04
 1.00034086615014E-04

 2.41705863300832E-04
 2.12572434056905E-04

 3.32345562038644E-04
 1.87563912403151E-04

 3.17238945582342E-04
 2.25076694883782E-04

 3.17238945582342E-04
 2.75093738191289E-04

 2.71919096213436E-04
 3.50119303152549E-04

 8.00650672184006E-04
 2.75093738191289E-04

 6.49584507620986E-04
 4.6265765059444E-04

 6.49584507620986E-04
 6.25213041343838E-04

 1.5106616456302E-03
 3.25110781498796E-04

 1.11788961776635E-03
 1.06286217028452E-03

 1.31427563169827E-03
 9.6282808366951E-04

 2.62855126339655E-03
 1.12538347441891E-04

 3.12706960645451E-03
 2.26327120966469E-03

 4.44134523815279E-03
 2.20074990553031E-03

 4.5470915533469E-03
 2.62589477364412E-03

 6.61669800786027E-03
 2.50085216537535E-03

 5.87647380150148E-03
 3.41366320573736E-03

 8.53523829781063E-03
 1.43798999509083E-03

 6.75265755596699E-03
 3.11356094589231E-03

 5.60455470528804E-03
 .004464021115195

 6.48073845975356E-03
 3.96385068211993E-03

 6.57137815849137E-03
 5.20177250398073E-03

 6.45052522684095E-03
 6.51471989080279E-03

 9.03375664086859E-03
 6.57724119493718E-03

 9.21503603834422E-03
 6.58974545576405E-03

 1.09220836979063E-02
 6.11458354434273E-03

 1.30823298511575E-02
 7.00238606305098E-03

 1.27197710562063E-02
 7.65260762604858E-03

 1.64359987044566E-02
 8.99056353452439E-03

 1.62245060740683E-02
 1.40297806477557E-02

 1.42304327018365E-02
 1.91315190651214E-02

 3.42013796570677E-02
 2.26952334007813E-02

 4.24798054751212E-02
 2.74218439933407E-02

 .106199513687803
 1.05786046595377E-02

 1.07256976839744E-03
 0

 1.07256976839744E-03
 0

 .163317630509081
 .150038625661694

 6.7979774053359E-04
 4.37649128940687E-04

 2.43216524946462E-03
 2.12572434056905E-03

 2.85061852530419E-02
 2.23076013151481E-02

 .131699482266041
 .125167650877036

 3.0213232912604E-05
 8.75298257881373E-05

 3.0213232912604E-05
 8.75298257881373E-05

 3.0213232912604E-05
 8.75298257881373E-05

 .117967567907262
 8.65669977044678E-02

 .117967567907262
 8.65669977044678E-02

 7.70437439271402E-04
 4.75161911421317E-04

 7.85544055727704E-04
 5.37683215555701E-04

 1.22363593296046E-03
 2.87597999018165E-04

 7.10010973446194E-04
 8.62793997054496E-04

 1.38980871397978E-03
 5.37683215555701E-04

 1.22363593296046E-03
 8.5028973622762E-04

 1.60130134436801E-03
 1.36296443012957E-03

 6.19371274708382E-03
 4.52654241932939E-03

 6.67712447368548E-03
 5.08923415653884E-03

 7.03968326863673E-03
 1.11162878750934E-02

 1.13450689586828E-02
 9.10310188196628E-03

 1.38376606739726E-02
 7.30248832289603E-03

 1.49102304423701E-02
 6.67727528155219E-03

 1.42606459347491E-02
 1.21666457845511E-02

 1.70251567462524E-02
 1.15789455256879E-02

 1.89739102691153E-02
 1.40923019518901E-02
