## Supplementary Item 4 for "Methane emission of humans is explained by dietary habits, host genetics, local formate availability and a uniform archaeome"

Javascript must be enabled to view this page.

magnitude

HE
LE

 100
 100

 98.6352484534303
 99.8086098878944

 1.90885936340482
 1.57809966148591

 1.90885936340482
 1.57809966148591

 .348797803343258
 .235683000104947

 2.56647529374802E-02
 1.73725925389504E-02

 1.51527424254697E-02
 1.07943349083253E-02

 3.27273054545782E-06
 2.6751759376271E-06

 2.35636599272963E-04
 5.93889058153215E-04

 2.51018432836615E-03
 1.0165668562983E-03

 2.38582056763875E-03
 2.18026838916608E-03

 4.43127715854989E-03
 3.06842680045828E-03

 5.5865510410965E-03
 3.93250862831183E-03

 1.44654690109236E-03
 1.08879660661423E-03

 3.27273054545782E-06
 8.02552781288129E-06

 1.4432741705469E-03
 1.08077107880135E-03

 1.34836498472862E-03
 1.07274555098847E-03

 1.34836498472862E-03
 1.07274555098847E-03

 2.05200205200205E-03
 1.27070857037287E-03

 2.05200205200205E-03
 1.27070857037287E-03

 5.66509657418748E-03
 3.14600690264946E-03

 2.0094565549111E-03
 1.38574113569084E-03

 3.65564001927638E-03
 1.76026576695863E-03

 1.54145608691063E-02
 1.10377759186494E-02

 3.78327651054924E-03
 2.74740568794303E-03

 0
 8.02552781288129E-06

 3.78327651054924E-03
 2.73938016013015E-03

 1.54472881745609E-03
 1.07542072692609E-03

 1.54472881745609E-03
 1.07542072692609E-03

 3.31200331200331E-03
 2.39428246417625E-03

 3.31200331200331E-03
 2.39428246417625E-03

 6.77455222909768E-03
 4.82066703960403E-03

 6.77455222909768E-03
 4.82066703960403E-03

 3.82582200764019E-02
 2.62140490128079E-02

 3.82582200764019E-02
 2.62140490128079E-02

 0
 8.02552781288129E-06

 9.81819163637346E-06
 2.6751759376271E-06

 1.30909221818313E-05
 1.07007037505084E-05

 2.61818443636625E-05
 4.01276390644064E-05

 3.6000036000036E-04
 2.59492065949828E-04

 4.6472773745501E-04
 2.43441010324066E-04

 6.18546073091528E-04
 1.73886435945761E-04

 4.15636779273143E-04
 3.5044804782915E-04

 5.00727773455046E-04
 3.39747344078641E-04

 5.20364156727793E-04
 3.26371464390506E-04

 6.25091534182443E-04
 2.62167241887455E-04

 5.46546001091456E-04
 3.85225335018302E-04

 5.30182348364167E-04
 4.01276390644064E-04

 4.90909581818673E-04
 4.41404029708471E-04

 6.77455222909768E-04
 2.99619705014235E-04

 6.51273378546106E-04
 5.00257900336267E-04

 6.87273414546142E-04
 5.53761419088809E-04

 7.69091678182587E-04
 5.91213882215588E-04

 1.05381923563742E-03
 7.00896095658299E-04

 1.11927384654657E-03
 7.62425142223722E-04

 1.14872842145569E-03
 9.09559818793213E-04

 1.32872860145587E-03
 8.69432179728806E-04

 1.58072885345613E-03
 8.50705948165417E-04

 2.48400248400248E-03
 1.51949993257219E-03

 2.95527568254841E-03
 1.89669973977761E-03

 4.75527748255021E-03
 3.48575424672811E-03

 6.11018792836975E-03
 4.11174541613285E-03

 7.81528054255327E-03
 5.91748917403114E-03

 .034177125086216
 2.19337675126046E-02

 .034177125086216
 2.19337675126046E-02

 6.54546109091564E-06
 7.22297503159316E-05

 1.47272874545602E-04
 1.95287843446778E-04

 3.43636707273071E-04
 1.49809852507117E-04

 4.32000432000432E-04
 1.55160204382372E-04

 3.69818551636733E-04
 2.46116186261693E-04

 5.89091498182407E-04
 2.38090658448812E-04

 4.87636851273215E-04
 3.45097695953895E-04

 6.905461450916E-04
 3.02294880951862E-04

 5.95636959273323E-04
 4.20002622207454E-04

 7.2000072000072E-04
 3.61148751579658E-04

 7.00364336727973E-04
 3.98601214706437E-04

 9.68728241455514E-04
 7.24972679096943E-04

 1.38763775127411E-03
 8.23954188789146E-04

 2.16654762109308E-03
 8.80132883479315E-04

 1.89163825527462E-03
 1.54625169194846E-03

 1.89491098582008E-03
 1.68536084070507E-03

 3.68182186364005E-03
 2.31135201010981E-03

 3.76691285782195E-03
 2.31670236198507E-03

 4.1661859843678E-03
 2.79823403075794E-03

 9.17019098837281E-03
 5.9629671649708E-03

 1.68185622731077E-02
 1.18751059871267E-02

 1.20731029821939E-02
 8.79330330698026E-03

 5.56364192727829E-05
 5.6178694690169E-05

 1.31236494872859E-03
 9.60388161608127E-04

 .00182291091382
 1.61045591445151E-03

 3.51818533636715E-03
 2.35415482511184E-03

 5.36400536400536E-03
 3.81212571111861E-03

 4.74545929091384E-03
 3.08180268014641E-03

 4.74545929091384E-03
 3.08180268014641E-03

 2.22447495174768E-02
 1.63051973398371E-02

 2.22447495174768E-02
 1.63051973398371E-02

 3.92727665454938E-05
 1.04331861567457E-04

 1.0472737745465E-04
 1.04331861567457E-04

 1.11272838545566E-04
 2.40765834386439E-04

 2.78182096363915E-04
 2.24714778760676E-04

 3.63273090545818E-04
 2.19364426885422E-04

 7.13455258909804E-04
 3.53123223766777E-04

 9.2945547491002E-04
 5.53761419088809E-04

 5.23636887273251E-04
 9.63063337545755E-04

 1.0243646607283E-03
 7.81151373787112E-04

 1.1945466490921E-03
 6.98220919720672E-04

 1.36800136800137E-03
 7.41023734722706E-04

 1.57091066181975E-03
 1.39644183944134E-03

 1.82945637491092E-03
 1.44994535819389E-03

 2.24182042363861E-03
 .001599755210701

 2.36945691491146E-03
 1.87797350821422E-03

 3.00763937127573E-03
 2.21237050041761E-03

 4.57527730255003E-03
 2.78485815106981E-03

 3.37745792291247E-03
 1.75491541508337E-03

 3.37745792291247E-03
 1.75491541508337E-03

 3.40363976727613E-04
 1.81911963758643E-04

 3.03709394618486E-03
 1.57300345132473E-03

 2.22872950145677E-03
 1.44994535819389E-03

 2.94545749091204E-04
 2.32740306573557E-04

 2.94545749091204E-04
 2.32740306573557E-04

 1.93418375236557E-03
 1.21720505162033E-03

 1.93418375236557E-03
 1.21720505162033E-03

 1.03974649429195E-02
 6.94475673407994E-03

 4.74545929091384E-04
 1.28408445006101E-04

 4.74545929091384E-04
 1.28408445006101E-04

 3.24000324000324E-03
 2.46116186261693E-03

 3.24000324000324E-03
 2.46116186261693E-03

 6.68291577382486E-03
 4.35518642645691E-03

 6.68291577382486E-03
 4.35518642645691E-03

 1.57091066181975E-02
 1.02218472576731E-02

 1.57091066181975E-02
 1.02218472576731E-02

 5.43273270545998E-04
 2.86243825326099E-04

 7.26546181091636E-04
 3.55798399704404E-04

 2.32036595672959E-03
 1.57835380319999E-03

 3.32509423418514E-03
 2.37555623261286E-03

 3.83236746873111E-03
 2.43441010324066E-03

 4.96145950691405E-03
 3.19148489358913E-03

 4.82564118927755E-02
 3.31855575062641E-02

 1.25051034141943E-02
 7.45571533816672E-03

 6.31636995273359E-04
 3.29046640328133E-04

 1.14545569091024E-03
 6.79494688157282E-04

 1.49891058981968E-03
 6.92870567845418E-04

 1.46945601491056E-03
 1.02994273598643E-03

 2.36945691491146E-03
 1.23860645912135E-03

 5.39018720836903E-03
 3.48575424672811E-03

 2.33934779389325E-02
 1.74073698261395E-02

 .004732368368732
 3.06842680045828E-03

 1.86611095702005E-02
 1.43389430256812E-02

 1.23578305396487E-02
 8.3224723419579E-03

 1.23578305396487E-02
 8.3224723419579E-03

 1.11011020101929E-02
 6.70666607563113E-03

 5.25927798655071E-03
 3.15403243046235E-03

 9.68728241455514E-04
 4.146522703322E-04

 1.14218296036478E-03
 8.48030772227789E-04

 3.14836678473042E-03
 1.89134938790236E-03

 5.84182402364221E-03
 3.55263364516878E-03

 1.93091102182011E-03
 1.13962494942914E-03

 3.91091300182209E-03
 2.41300869573964E-03

 9.52364588728225E-04
 5.29684835650165E-04

 9.52364588728225E-04
 5.29684835650165E-04

 9.52364588728225E-04
 5.29684835650165E-04

 8.57455402909948E-04
 6.47392576905757E-04

 8.57455402909948E-04
 6.47392576905757E-04

 8.57455402909948E-04
 6.47392576905757E-04

 2.82763919127556E-03
 1.67198496101693E-03

 1.15527388254661E-03
 7.14271975346435E-04

 1.15527388254661E-03
 7.14271975346435E-04

 1.67236530872895E-03
 9.577129856705E-04

 1.67236530872895E-03
 9.577129856705E-04

 1.64291073381982E-03
 8.42680420352535E-04

 1.64291073381982E-03
 8.42680420352535E-04

 1.64291073381982E-03
 8.42680420352535E-04

 1.58727431454704E-03
 1.01924203223592E-03

 1.58727431454704E-03
 1.01924203223592E-03

 1.58727431454704E-03
 1.01924203223592E-03

 2.16000216000216E-03
 2.17491803729083E-03

 2.16000216000216E-03
 2.17491803729083E-03

 2.16000216000216E-03
 2.17491803729083E-03

 8.39782657964476E-03
 5.36372775494233E-03

 3.03054848509394E-03
 1.55160204382372E-03

 3.03054848509394E-03
 1.55160204382372E-03

 5.36727809455082E-03
 3.81212571111861E-03

 5.36727809455082E-03
 3.81212571111861E-03

 2.92582110763929E-03
 2.03045853665897E-03

 2.92582110763929E-03
 2.03045853665897E-03

 2.92582110763929E-03
 2.03045853665897E-03

 7.68437132073496E-03
 5.70079992308334E-03

 3.0567303294576E-03
 2.00638195322032E-03

 3.0567303294576E-03
 2.00638195322032E-03

 4.62764099127736E-03
 3.69441796986302E-03

 4.62764099127736E-03
 3.69441796986302E-03

 2.86363922727559E-03
 2.17759321322846E-03

 2.86363922727559E-03
 2.17759321322846E-03

 2.86363922727559E-03
 2.17759321322846E-03

 9.61855507310053E-03
 6.06462385060063E-03

 9.61855507310053E-03
 6.06462385060063E-03

 3.48873076145803E-03
 2.05453512009761E-03

 6.12982431164249E-03
 4.01008873050302E-03

 3.97964034327671E-03
 2.28995060260879E-03

 3.97964034327671E-03
 2.28995060260879E-03

 3.97964034327671E-03
 2.28995060260879E-03

 3.89127661854935E-03
 2.7153035766915E-03

 3.89127661854935E-03
 2.7153035766915E-03

 3.89127661854935E-03
 2.7153035766915E-03

 4.20218602036784E-03
 2.66180005793896E-03

 4.20218602036784E-03
 2.66180005793896E-03

 4.20218602036784E-03
 2.66180005793896E-03

 4.40836804473168E-03
 2.66447523387659E-03

 4.40836804473168E-03
 2.66447523387659E-03

 4.40836804473168E-03
 2.66447523387659E-03

 4.38873166145893E-03
 2.9025658923254E-03

 4.38873166145893E-03
 2.9025658923254E-03

 4.38873166145893E-03
 2.9025658923254E-03

 4.38545893091348E-03
 3.10855443952269E-03

 4.38545893091348E-03
 3.10855443952269E-03

 4.38545893091348E-03
 3.10855443952269E-03

 4.3756407392771E-03
 3.5044804782915E-03

 4.3756407392771E-03
 3.5044804782915E-03

 4.3756407392771E-03
 3.5044804782915E-03

 5.02036865673229E-03
 4.0475411936298E-03

 5.02036865673229E-03
 4.0475411936298E-03

 5.02036865673229E-03
 4.0475411936298E-03

 5.23636887273251E-03
 4.05556672144268E-03

 5.23636887273251E-03
 4.05556672144268E-03

 5.23636887273251E-03
 4.05556672144268E-03

 7.02982521164339E-03
 2.92664247576404E-03

 7.02982521164339E-03
 2.92664247576404E-03

 7.02982521164339E-03
 2.92664247576404E-03

 8.79709970619062E-03
 5.39582986619385E-03

 8.79709970619062E-03
 5.39582986619385E-03

 8.79709970619062E-03
 5.39582986619385E-03

 7.91673518946246E-03
 6.18500676779385E-03

 7.91673518946246E-03
 6.18500676779385E-03

 7.91673518946246E-03
 6.18500676779385E-03

 .907515089333271
 .803962599007258

 .907515089333271
 .803962599007258

 .870376143103416
 .726296891186068

 7.2000072000072E-05
 1.09682213442711E-04

 3.00763937127573E-03
 1.59172968288812E-03

 6.43418825237007E-03
 4.95977618836064E-03

 1.88574734029279E-02
 5.70347509902097E-03

 2.78116641753005E-02
 1.49515583153978E-02

 3.47498529316711E-02
 1.81778204961761E-02

 4.60702278884097E-02
 1.73618918351999E-02

 4.78211387302296E-02
 6.72432223681947E-02

 .093102638557184
 5.39368972544375E-02

 .268075904439541
 .287543960831786

 .324373415282506
 .254716876901164

 4.58836822473186E-03
 2.56549372418439E-03

 4.58836822473186E-03
 2.56549372418439E-03

 5.10218692036874E-03
 3.31454298671997E-03

 5.10218692036874E-03
 3.31454298671997E-03

 2.74483910847547E-02
 7.17856711102855E-02

 2.74483910847547E-02
 7.17856711102855E-02

 1.03418285236467E-03
 9.63063337545755E-04

 1.03418285236467E-03
 9.63063337545755E-04

 1.03418285236467E-03
 9.63063337545755E-04

 1.03418285236467E-03
 9.63063337545755E-04

 1.78691087781997E-03
 1.00586615254779E-03

 1.78691087781997E-03
 1.00586615254779E-03

 1.78691087781997E-03
 1.00586615254779E-03

 1.78691087781997E-03
 1.00586615254779E-03

 5.07273234545962E-03
 2.99887222607997E-03

 5.07273234545962E-03
 2.99887222607997E-03

 5.07273234545962E-03
 2.99887222607997E-03

 5.07273234545962E-03
 2.99887222607997E-03

 .631993722902814
 .525348375455271

 .631993722902814
 .525348375455271

 .144844508480872
 .146746776058534

 1.01454646909192E-02
 8.21546530445281E-03

 1.37487410214683E-02
 9.36846613357009E-03

 .120950302768485
 .129162844620511

 .102145193054284
 8.03729858700685E-02

 9.4418276236458E-03
 9.06884642855586E-03

 1.25803762167399E-02
 7.60017483879858E-03

 8.01229892138983E-02
 .063703964602714

 .137667410394683
 .100492984096962

 .012217103126194
 8.92171175198636E-03

 .125450307268489
 9.15712723449755E-02

 3.71258553076735E-02
 2.77174978897543E-02

 3.71258553076735E-02
 2.77174978897543E-02

 .076424803697531
 5.98169339653419E-02

 .076424803697531
 5.98169339653419E-02

 .13378595196777
 .110201197574611

 .13378595196777
 .110201197574611

 1.26589217498308E-02
 8.13788520226163E-03

 1.26589217498308E-02
 8.13788520226163E-03

 1.26589217498308E-02
 8.13788520226163E-03

 1.26589217498308E-02
 8.13788520226163E-03

 5.02496684314866
 3.13383485213326

 .428681883227338
 .425534886046467

 7.64575310029856E-02
 .106485378197247

 5.04360504360504E-02
 7.86501725662366E-02

 1.78461996643815E-02
 .029226297118576

 0
 5.35035187525419E-06

 5.97273324546052E-03
 9.18922934574908E-03

 .011873466418921
 2.00317174209517E-02

 7.28182546364365E-03
 1.13052935124121E-02

 7.28182546364365E-03
 1.13052935124121E-02

 2.53080253080253E-02
 3.81185819352485E-02

 8.06400806400806E-03
 1.38921886440975E-02

 1.72440172440172E-02
 .024226393291151

 2.60214805669351E-02
 2.78352056310099E-02

 5.05964142327779E-03
 5.85863530340334E-03

 3.92727665454938E-05
 2.6751759376271E-06

 1.05381923563742E-03
 1.44459500631863E-03

 3.96654942109488E-03
 4.41136512114708E-03

 1.86872914145641E-03
 2.7741574473193E-03

 1.86872914145641E-03
 2.7741574473193E-03

 .003181094090185
 3.10587926358506E-03

 .003181094090185
 3.10587926358506E-03

 4.54254999709545E-03
 4.1170957680081E-03

 4.54254999709545E-03
 4.1170957680081E-03

 5.29855075309621E-03
 5.66869781183182E-03

 5.29855075309621E-03
 5.66869781183182E-03

 6.07091516182425E-03
 6.31074003686232E-03

 6.07091516182425E-03
 6.31074003686232E-03

 8.39029929939021E-02
 7.70771691149119E-02

 .066024066024066
 6.12588537957229E-02

 .011595284322557
 1.00640118773531E-02

 9.81819163637346E-06
 0

 1.08000108000108E-03
 6.09940113778978E-04

 2.5756389392753E-03
 2.75008086388065E-03

 7.92982611164429E-03
 6.7039908996935E-03

 6.05455150909696E-03
 4.77786422460199E-03

 2.94545749091204E-05
 0

 2.94545749091204E-05
 1.33758796881355E-05

 5.99564235927872E-03
 4.76448834491386E-03

 8.06728079455352E-03
 7.70985705224129E-03

 3.20727593454866E-04
 1.07007037505084E-04

 3.27273054545782E-04
 1.92612667509151E-04

 5.4000054000054E-04
 6.12615289716605E-04

 5.6945511490966E-04
 6.50067752843384E-04

 3.30873058145785E-03
 2.86778860513625E-03

 3.00109391018482E-03
 3.27976569953082E-03

 1.27636491272855E-03
 1.16637670880541E-03

 3.66545821091276E-04
 3.29046640328133E-04

 4.54909545818637E-04
 3.37072168141014E-04

 4.54909545818637E-04
 5.00257900336267E-04

 3.69818551636733E-04
 3.45097695953895E-04

 3.69818551636733E-04
 3.45097695953895E-04

 9.10146364691819E-03
 8.41075314789959E-03

 4.45091354182263E-04
 3.71849455330166E-04

 4.12364048727685E-04
 4.49429557521352E-04

 5.00727773455046E-04
 6.07264937841351E-04

 7.59273486546214E-04
 5.40385539400673E-04

 1.20109211018302E-03
 1.42319359881761E-03

 1.89491098582008E-03
 1.95020325853015E-03

 3.88800388800389E-03
 3.06842680045828E-03

 3.69818551636733E-04
 4.52104733458979E-04

 3.69818551636733E-04
 4.52104733458979E-04

 5.66182384364203E-04
 4.28028150020335E-04

 5.66182384364203E-04
 4.28028150020335E-04

 5.33455078909624E-04
 5.10958604086775E-04

 5.33455078909624E-04
 5.10958604086775E-04

 1.8000018000018E-03
 1.36701490412745E-03

 6.905461450916E-04
 5.37710363463046E-04

 1.1094556549102E-03
 8.293045406644E-04

 3.40036703673067E-03
 2.99619705014235E-03

 7.06909797818889E-04
 5.9923941002847E-04

 1.21418303236485E-03
 1.27873409818575E-03

 1.47927420654693E-03
 1.11822354192813E-03

 2.27127499854773E-03
 2.10803863885015E-03

 8.60728133455406E-04
 5.35035187525419E-04

 1.41054686509232E-03
 1.57300345132473E-03

 7.10182528364347E-04
 7.86501725662366E-04

 7.10182528364347E-04
 7.86501725662366E-04

 8.64000864000864E-04
 6.87520215970164E-04

 8.64000864000864E-04
 6.87520215970164E-04

 6.74182492364311E-04
 8.58731475978298E-04

 6.74182492364311E-04
 8.58731475978298E-04

 1.5283651647288E-03
 6.50067752843384E-04

 1.5283651647288E-03
 6.50067752843384E-04

 1.31563767927404E-03
 1.49007299725829E-03

 1.31563767927404E-03
 1.49007299725829E-03

 1.06691015781925E-03
 1.92880185102914E-03

 1.06691015781925E-03
 1.92880185102914E-03

 1.66909257818349E-03
 1.4713467656949E-03

 1.66909257818349E-03
 1.4713467656949E-03

 2.59200259200259E-03
 4.32843466708064E-03

 1.40727413454686E-03
 1.95287843446778E-03

 1.18472845745573E-03
 2.37555623261286E-03

 2.09127481854755E-03
 1.90740044352812E-03

 2.09127481854755E-03
 1.90740044352812E-03

 1.87527460254733E-03
 2.1321152222888E-03

 1.87527460254733E-03
 2.1321152222888E-03

 2.92254837709383E-03
 2.42638457542778E-03

 2.92254837709383E-03
 2.42638457542778E-03

 3.30873058145785E-03
 2.25517331541964E-03

 3.30873058145785E-03
 2.25517331541964E-03

 3.23018504836687E-03
 1.69873672039321E-03

 3.23018504836687E-03
 1.69873672039321E-03

 4.61455006909552E-04
 5.53761419088809E-04

 2.76873004145731E-03
 1.1449753013044E-03

 1.46487419214692E-02
 1.41195785987958E-02

 2.79491188582098E-03
 2.82766096607184E-03

 2.79491188582098E-03
 2.82766096607184E-03

 3.15491224582134E-03
 3.57135987673217E-03

 3.15491224582134E-03
 3.57135987673217E-03

 4.42145896691351E-03
 .003673016562362

 4.42145896691351E-03
 .003673016562362

 4.27745882291337E-03
 4.0475411936298E-03

 4.27745882291337E-03
 4.0475411936298E-03

 .180235816599453
 .157634742124677

 1.24952852225579E-02
 1.00238842382887E-02

 7.16073443346171E-03
 5.23799448587385E-03

 0
 1.60510556257626E-05

 1.96363832727469E-05
 2.6751759376271E-05

 5.56364192727829E-05
 2.14014075010168E-05

 3.92727665454938E-05
 4.01276390644064E-05

 6.21818803636985E-05
 6.15290465654232E-05

 7.85455330909876E-05
 8.82808059416942E-05

 6.54546109091564E-05
 1.23058093130846E-04

 1.21091030181939E-04
 1.04331861567457E-04

 2.12727485454758E-04
 6.68793984406774E-05

 3.17454862909408E-04
 2.19364426885422E-04

 8.01818983637165E-04
 4.68155789084742E-04

 9.2945547491002E-04
 6.47392576905757E-04

 1.87200187200187E-03
 1.36433972818982E-03

 2.58545713091168E-03
 1.99033089759456E-03

 5.33455078909624E-03
 4.78588975241487E-03

 9.52364588728225E-04
 8.1057830910101E-04

 4.38218620036802E-03
 3.97531144331386E-03

 4.67378649196831E-02
 3.55530882110641E-02

 .013968013968014
 1.08638894827036E-02

 2.94545749091204E-05
 2.14014075010168E-05

 4.66691375782285E-03
 4.1866503423864E-03

 .009271645635282
 6.65583773281621E-03

 4.6472773745501E-04
 2.72867945637964E-04

 1.89818371636553E-04
 8.293045406644E-05

 2.74909365818457E-04
 1.89937491571524E-04

 1.31400131400131E-02
 .010072037405166

 2.34654780109326E-03
 1.48739782132067E-03

 4.14654960109506E-03
 3.07377715233353E-03

 6.64691573782483E-03
 5.51086243151182E-03

 .019165110074201
 1.43442933775565E-02

 7.39309830218921E-03
 5.60181841339114E-03

 1.17720117720118E-02
 8.74247496416535E-03

 5.44124180487817E-02
 4.66657690559671E-02

 5.4000054000054E-04
 5.88538706277961E-04

 5.4000054000054E-04
 5.88538706277961E-04

 1.00243736607373E-02
 8.04425404444468E-03

 2.69345723891178E-03
 1.84052104508744E-03

 3.36763973127609E-03
 3.08180268014641E-03

 3.96327669054942E-03
 3.12193031921082E-03

 2.20254765709311E-03
 2.25517331541964E-03

 2.20254765709311E-03
 2.25517331541964E-03

 1.47600147600148E-02
 1.40232722650412E-02

 3.30873058145785E-03
 3.23161253265353E-03

 4.80436844073208E-03
 4.66283165928403E-03

 6.64691573782483E-03
 6.12882807310368E-03

 .026885481430936
 2.17545307247835E-02

 .026885481430936
 2.17545307247835E-02

 2.17571126662036E-02
 2.16983520300934E-02

 6.25091534182443E-04
 7.03571271595926E-04

 6.25091534182443E-04
 7.03571271595926E-04

 8.55819037637219E-03
 7.03838789189689E-03

 1.04400104400104E-03
 9.68413689421009E-04

 7.51418933237115E-03
 6.06997420247588E-03

 6.34909725818817E-03
 6.02182103559859E-03

 6.34909725818817E-03
 6.02182103559859E-03

 6.22473349746077E-03
 7.93457183100197E-03

 6.22473349746077E-03
 7.93457183100197E-03

 1.28945583491038E-02
 1.28542203802982E-02

 1.28945583491038E-02
 1.28542203802982E-02

 8.05091714182623E-04
 8.50705948165417E-04

 1.25345579891034E-03
 1.0165668562983E-03

 2.43491152582062E-03
 2.22574638010574E-03

 2.53636617272981E-03
 2.20969532447998E-03

 2.37600237600238E-03
 2.61097171512405E-03

 3.48873076145803E-03
 3.94053415612471E-03

 3.40363976727613E-03
 3.2289373567159E-03

 7.75637139273503E-04
 9.38986754107111E-04

 7.75637139273503E-04
 9.38986754107111E-04

 2.62800262800263E-03
 2.28995060260879E-03

 2.62800262800263E-03
 2.28995060260879E-03

 8.31273558546286E-04
 9.17585346606094E-04

 8.31273558546286E-04
 9.17585346606094E-04

 8.31273558546286E-04
 9.17585346606094E-04

 6.14291523382433E-03
 .005588442533703

 6.14291523382433E-03
 .005588442533703

 9.7527370254643E-04
 9.81789569109144E-04

 1.15200115200115E-03
 8.72107355666433E-04

 1.11927384654657E-03
 9.65738513483382E-04

 1.21745576291031E-03
 1.13694977349152E-03

 1.67891076981986E-03
 1.63185732195253E-03

 4.53600453600454E-03
 4.8474187989803E-03

 1.76727449454722E-03
 1.77096647070914E-03

 1.76727449454722E-03
 1.77096647070914E-03

 2.76873004145731E-03
 3.07645232827116E-03

 2.76873004145731E-03
 3.07645232827116E-03

 4.61782279964098E-03
 4.09569436050708E-03

 1.95054740509286E-03
 2.1026882869749E-03

 1.95054740509286E-03
 2.1026882869749E-03

 2.66727539454812E-03
 1.99300607353219E-03

 2.66727539454812E-03
 1.99300607353219E-03

 1.24069214978306E-02
 1.21613498124528E-02

 2.98145752691207E-03
 2.53606678887049E-03

 2.98145752691207E-03
 2.53606678887049E-03

 3.7865492410947E-03
 4.3444857227064E-03

 3.7865492410947E-03
 4.3444857227064E-03

 5.63891472982382E-03
 5.28079730087589E-03

 5.63891472982382E-03
 5.28079730087589E-03

 3.42327615054888E-02
 2.93707566192079E-02

 2.74876638513002E-02
 .024764103654614

 6.80073407346135E-03
 5.55366524651385E-03

 2.29091138182047E-05
 1.07007037505084E-05

 1.01454646909192E-04
 2.6751759376271E-06

 1.01781919963738E-03
 1.00854132848542E-03

 5.65855111309657E-03
 4.5317480383403E-03

 3.99273126545854E-04
 3.04970056889489E-04

 3.99273126545854E-04
 3.04970056889489E-04

 2.91273018545746E-03
 3.09785373577218E-03

 9.19637283273647E-04
 7.86501725662366E-04

 1.99309290218381E-03
 2.31135201010981E-03

 5.67818749636931E-03
 4.21072692582505E-03

 5.67818749636931E-03
 4.21072692582505E-03

 5.28218710036892E-03
 5.95761681309554E-03

 5.28218710036892E-03
 5.95761681309554E-03

 6.41455186909732E-03
 5.63927087651792E-03

 6.41455186909732E-03
 5.63927087651792E-03

 6.74509765418856E-03
 4.60665296459386E-03

 6.74509765418856E-03
 4.60665296459386E-03

 6.64364300727937E-04
 5.8051317846508E-04

 2.37600237600238E-03
 1.49274817319592E-03

 3.70473097745825E-03
 2.53339161293286E-03

 1.97738379556561E-02
 1.92131135840378E-02

 7.05600705600706E-03
 6.62373562156469E-03

 3.83564019927656E-03
 4.10104471238234E-03

 1.96363832727469E-05
 1.87262315633897E-05

 2.61818443636625E-05
 2.14014075010168E-05

 3.6000036000036E-05
 1.60510556257626E-05

 2.094547549093E-04
 2.62167241887455E-04

 4.25454970909516E-04
 4.52104733458979E-04

 6.97091606182515E-04
 6.74144336282028E-04

 8.11637175273539E-04
 1.04064343973694E-03

 1.61018342836525E-03
 1.61580626632677E-03

 6.31636995273359E-04
 5.29684835650165E-04

 1.11272838545566E-04
 5.35035187525419E-05

 5.20364156727793E-04
 4.76181316897623E-04

 4.77818659636841E-04
 4.76181316897623E-04

 1.27636491272855E-04
 1.60510556257626E-04

 3.50182168363987E-04
 3.15670760639997E-04

 2.11091120182029E-03
 1.51682475663456E-03

 4.54909545818637E-04
 4.57455085334233E-04

 7.75637139273503E-04
 4.7885649283525E-04

 8.80364516728153E-04
 5.8051317846508E-04

 9.5661913843732E-03
 9.48082352295043E-03

 .008728372364736
 8.65954451009891E-03

 2.61818443636625E-05
 2.40765834386439E-05

 9.81819163637345E-05
 6.68793984406774E-05

 1.34181952363771E-04
 1.81911963758643E-04

 1.14545569091024E-04
 2.08663723134913E-04

 4.6472773745501E-04
 4.17327446269827E-04

 6.25091534182443E-04
 3.39747344078641E-04

 6.57818839637021E-04
 3.15670760639997E-04

 4.87636851273215E-04
 4.92232372523386E-04

 3.46909437818529E-04
 6.23315993467113E-04

 4.61455006909552E-04
 5.88538706277961E-04

 7.26546181091636E-04
 5.93889058153215E-04

 8.77091786182695E-04
 7.97202429412875E-04

 6.97091606182515E-04
 9.63063337545755E-04

 3.01091210182119E-03
 3.04702539295726E-03

 8.37819019637201E-04
 8.21279012851518E-04

 8.37819019637201E-04
 8.21279012851518E-04

 3.15163951527588E-03
 3.10855443952269E-03

 3.40363976727613E-04
 7.00896095658299E-04

 3.40363976727613E-04
 7.00896095658299E-04

 2.81127553854827E-03
 2.40765834386439E-03

 5.59636923273287E-04
 6.15290465654232E-04

 2.25163861527498E-03
 1.79236787821015E-03

 5.23964160327797E-03
 7.97469947006637E-03

 5.23964160327797E-03
 7.97469947006637E-03

 3.42000342000342E-03
 5.79443108090029E-03

 2.32363868727505E-04
 6.42042225030503E-05

 3.18763955127592E-03
 5.73022685839724E-03

 1.81963818327455E-03
 2.18026838916608E-03

 1.81963818327455E-03
 2.18026838916608E-03

 2.67512994785722E-02
 2.60455129287374E-02

 2.67512994785722E-02
 2.60455129287374E-02

 2.64763901127537E-03
 2.72332910450438E-03

 6.70909761818853E-04
 7.14271975346435E-04

 1.97672924945652E-03
 2.00905712915795E-03

 1.42527415254688E-02
 1.42265856363009E-02

 2.05854751309297E-03
 1.73083883164473E-03

 6.13636977273341E-03
 6.21175852717012E-03

 6.05782423964242E-03
 6.28398827748605E-03

 2.96509387418478E-03
 2.39695764011388E-03

 2.96509387418478E-03
 2.39695764011388E-03

 6.88582506764325E-03
 6.69864054781825E-03

 6.88582506764325E-03
 6.69864054781825E-03

 2.08800208800209E-03
 1.73351400758236E-03

 2.08800208800209E-03
 1.73351400758236E-03

 2.08800208800209E-03
 1.73351400758236E-03

 2.08800208800209E-03
 1.73351400758236E-03

 .704019976747249
 .581128468930734

 .514685969231424
 .439480578209317

 6.20149711058802E-02
 5.04431174798965E-02

 3.27273054545782E-06
 2.6751759376271E-06

 3.27273054545782E-06
 2.6751759376271E-06

 5.89091498182407E-05
 8.02552781288129E-06

 5.89091498182407E-05
 8.02552781288129E-06

 7.59273486546214E-03
 6.31609038873757E-03

 7.59273486546214E-03
 6.31609038873757E-03

 1.21516485152849E-02
 9.13037547512128E-03

 1.21516485152849E-02
 9.13037547512128E-03

 1.12549203458294E-02
 1.05963718889409E-02

 1.12549203458294E-02
 1.05963718889409E-02

 .03095348549894
 2.43895790233462E-02

 .03095348549894
 2.43895790233462E-02

 .176292176292176
 .147902452063589

 5.07927780655053E-03
 .004146522703322

 3.27273054545782E-06
 2.6751759376271E-06

 5.07600507600508E-03
 4.14384752738437E-03

 8.04764441128077E-03
 6.02717138747385E-03

 8.04764441128077E-03
 6.02717138747385E-03

 8.67600867600868E-03
 7.98004982194163E-03

 8.67600867600868E-03
 7.98004982194163E-03

 .154489245398336
 .129748708150852

 .154489245398336
 .129748708150852

 .079177170086261
 6.70532848766232E-02

 4.82073209345937E-03
 4.50767145490166E-03

 3.27273054545782E-06
 1.33758796881355E-05

 4.81745936291391E-03
 4.49429557521352E-03

 2.58709349618441E-02
 2.19792455035442E-02

 1.20436484072848E-03
 9.65738513483382E-04

 1.72800172800173E-03
 1.53822616413558E-03

 1.7770926861836E-03
 1.61313109038914E-03

 .010243646607283
 9.0634960766806E-03

 1.09178290996473E-02
 8.79865365885552E-03

 2.69345723891178E-03
 2.48523844605557E-03

 2.69345723891178E-03
 2.48523844605557E-03

 4.86982305164123E-03
 4.13582199957149E-03

 4.86982305164123E-03
 4.13582199957149E-03

 .014625832807651
 1.17145954308691E-02

 6.45055190509736E-03
 .005430607153383

 8.17528090255363E-03
 6.28398827748605E-03

 6.89237052873417E-03
 5.49213619994843E-03

 6.89237052873417E-03
 5.49213619994843E-03

 7.37673464946192E-03
 6.73074265906977E-03

 7.37673464946192E-03
 6.73074265906977E-03

 1.20272847545575E-02
 .010007833182663

 1.20272847545575E-02
 .010007833182663

 .156341610887065
 .135642120741444

 8.68746323291778E-02
 7.27193075125173E-02

 3.92727665454938E-05
 5.35035187525419E-06

 5.53091462182371E-04
 5.91213882215588E-04

 6.905461450916E-04
 5.77838002527453E-04

 6.28364264727901E-04
 6.95545743783045E-04

 7.9527352254625E-04
 7.00896095658299E-04

 1.04072831345559E-03
 5.93889058153215E-04

 9.45819127637309E-04
 7.24972679096943E-04

 1.22727395454668E-03
 9.28286050356602E-04

 1.49236512872877E-03
 1.51682475663456E-03

 2.2189113098204E-03
 1.59440485882575E-03

 3.01418483236665E-03
 1.92880185102914E-03

 2.80473007745735E-03
 2.41300869573964E-03

 3.07636671273035E-03
 2.4879136219932E-03

 3.71782189964008E-03
 3.01759845764336E-03

 4.06473133745861E-03
 3.71314420142641E-03

 5.1283687647324E-03
 3.73454560892743E-03

 5.51782369964188E-03
 4.38728853770844E-03

 5.44255089709635E-03
 4.47021899177488E-03

 5.64546019091474E-03
 5.03735629055182E-03

 8.69564505928142E-03
 6.80832276126096E-03

 8.20146274691729E-03
 7.28450407815858E-03

 9.0229181138272E-03
 8.05227957225756E-03

 1.29109220018311E-02
 1.14551033649192E-02

 2.61196624832988E-02
 2.49593914980608E-02

 1.62981981163799E-03
 1.50879922882168E-03

 8.48946303491758E-03
 8.07100580382095E-03

 1.60003796367433E-02
 1.53795864654182E-02

 3.05738487556669E-02
 2.71771123503537E-02

 3.31527604254877E-03
 2.50931502949422E-03

 7.44873472146199E-03
 6.87520215970164E-03

 9.02619084437266E-03
 7.74463433943044E-03

 1.07836471472835E-02
 1.00479608217274E-02

 4.07782225964044E-03
 3.86562922987115E-03

 4.07782225964044E-03
 3.86562922987115E-03

 8.69564505928142E-03
 6.9206801506413E-03

 8.69564505928142E-03
 6.9206801506413E-03

 4.08600408600409E-02
 3.84396030477637E-02

 8.46982665164483E-03
 7.74730951536807E-03

 3.27273054545782E-05
 1.87262315633897E-05

 2.98473025745753E-03
 2.85976307732337E-03

 5.45236908873273E-03
 4.86882020648131E-03

 .032390214208396
 3.06922935323957E-02

 3.25309416218507E-03
 3.01224810576811E-03

 4.00909491818583E-03
 3.6435896270481E-03

 6.06436970073334E-03
 5.65264675620605E-03

 8.63673590946318E-03
 8.2341915360162E-03

 1.04269195178286E-02
 1.01496175073572E-02

 7.49128021855295E-03
 5.56704112620199E-03

 7.49128021855295E-03
 5.56704112620199E-03

 6.4472791745519E-04
 4.38728853770844E-04

 6.4472791745519E-04
 4.38728853770844E-04

 6.84655230109776E-03
 5.12831227243114E-03

 6.84655230109776E-03
 5.12831227243114E-03

 6.98138879957062E-02
 4.99321588758097E-02

 6.98138879957062E-02
 4.99321588758097E-02

 4.12658594476776E-02
 2.86029811251089E-02

 1.03745558291013E-03
 3.9592603876881E-04

 8.83637247273611E-04
 7.59749966286095E-04

 1.12909203818295E-03
 5.91213882215588E-04

 1.02109193018284E-03
 7.11596799408807E-04

 1.1945466490921E-03
 6.09940113778978E-04

 1.42363778727415E-03
 8.61406651915925E-04

 1.36472863745591E-03
 9.92490272859653E-04

 2.55927528654801E-03
 1.99835642540744E-03

 7.21964358327995E-03
 4.80194080804064E-03

 7.69091678182587E-03
 6.22245923092063E-03

 1.57418339236521E-02
 1.06579009355064E-02

 1.28618310436492E-03
 9.68413689421009E-04

 1.28618310436492E-03
 9.68413689421009E-04

 1.79345633891088E-03
 9.47012281919992E-04

 1.79345633891088E-03
 9.47012281919992E-04

 2.27454772909318E-03
 1.14230012536677E-03

 2.27454772909318E-03
 1.14230012536677E-03

 2.88000288000288E-03
 2.28995060260879E-03

 2.88000288000288E-03
 2.28995060260879E-03

 7.48146202691657E-03
 5.72220133058436E-03

 7.48146202691657E-03
 5.72220133058436E-03

 1.28323764687401E-02
 1.02592997207999E-02

 1.28323764687401E-02
 1.02592997207999E-02

 2.09029299938391E-02
 1.53207325947904E-02

 9.13091822182731E-04
 5.29684835650165E-04

 9.13091822182731E-04
 5.29684835650165E-04

 9.13091822182731E-04
 5.29684835650165E-04

 1.99898381716564E-02
 1.47910477591402E-02

 4.65709556618648E-03
 3.67569173829963E-03

 1.2338194156376E-03
 8.9350876316745E-04

 3.42327615054888E-03
 2.78218297513218E-03

 5.43273270545998E-03
 3.82282641486912E-03

 2.38909329818421E-03
 1.6880360166427E-03

 3.04363940727577E-03
 2.13479039822642E-03

 9.9000099000099E-03
 7.29252960597146E-03

 9.9000099000099E-03
 7.29252960597146E-03

 2.58807531534804E-02
 1.86941294521381E-02

 2.58807531534804E-02
 1.86941294521381E-02

 1.95742013923832E-02
 1.36005944668962E-02

 2.94218476036658E-03
 2.02510818478371E-03

 6.56182474364293E-03
 4.9464003086725E-03

 1.00701918883737E-02
 6.62908597343994E-03

 6.30655176109722E-03
 5.09353498524199E-03

 6.30655176109722E-03
 5.09353498524199E-03

 9.82800982800983E-03
 8.18336319320129E-03

 9.82800982800983E-03
 8.18336319320129E-03

 4.68327741055014E-03
 3.6435896270481E-03

 4.68327741055014E-03
 3.6435896270481E-03

 5.14473241745969E-03
 4.53977356615318E-03

 5.14473241745969E-03
 4.53977356615318E-03

 4.56513183785911E-02
 3.61309262135916E-02

 4.56513183785911E-02
 3.61309262135916E-02

 7.09200709200709E-03
 5.8746863590291E-03

 7.09200709200709E-03
 5.8746863590291E-03

 1.23447396174669E-02
 9.67076101452195E-03

 1.23447396174669E-02
 9.67076101452195E-03

 1.25051034141943E-02
 9.85267297828059E-03

 1.25051034141943E-02
 9.85267297828059E-03

 1.37094682549228E-02
 1.07328058617599E-02

 1.37094682549228E-02
 1.07328058617599E-02

 9.76582794764613E-03
 .007819539265684

 9.76582794764613E-03
 .007819539265684

 9.76582794764613E-03
 .007819539265684

 9.76582794764613E-03
 .007819539265684

 2.90247672065854
 1.42759693841095

 1.10359746723383
 .867594333859656

 1.10359746723383
 .867594333859656

 1.77741995923814E-02
 7.30055513378434E-03

 6.54546109091564E-06
 0

 1.13563749927386E-03
 2.54141714074574E-04

 3.48218530036712E-03
 1.41516807100473E-03

 6.23455168909714E-03
 2.90524106826303E-03

 6.91527964255237E-03
 2.72600428044201E-03

 1.20141938323757E-02
 5.69009921933283E-03

 1.30909221818313E-05
 2.6751759376271E-06

 1.7149108058199E-03
 5.24334483774911E-04

 2.97491206582116E-03
 1.51414958069694E-03

 3.6229127138218E-03
 1.68536084070507E-03

 3.68836732473096E-03
 1.96357913821829E-03

 2.61818443636625E-05
 0

 2.61818443636625E-05
 0

 3.63796727433091E-02
 1.47803470553897E-02

 4.90909581818673E-05
 1.87262315633897E-05

 9.81819163637345E-05
 3.74524631267793E-05

 2.18618400436582E-03
 1.4713467656949E-03

 1.55487428214701E-02
 7.19889844815451E-03

 1.84974730429276E-02
 6.05392314685012E-03

 1.70901989083807E-02
 6.73074265906977E-03

 5.56364192727829E-05
 5.08283428149148E-05

 1.85563821927458E-03
 7.51724438473214E-04

 1.51789242698334E-02
 5.92818987778164E-03

 2.49545704091159E-02
 .012776640278107

 1.5054560509106E-04
 2.6751759376271E-06

 2.42182060363879E-04
 1.71211260008134E-04

 6.34909725818817E-03
 1.74153953539524E-03

 1.82127454854728E-02
 .010861214306766

 2.52000252000252E-04
 6.68793984406774E-05

 2.52000252000252E-04
 6.68793984406774E-05

 1.3189104098195E-03
 4.17327446269827E-04

 3.37091246182155E-04
 1.23058093130846E-04

 9.81819163637346E-04
 2.94269353138981E-04

 5.56364192727829E-04
 7.49049262535587E-05

 5.56364192727829E-04
 7.49049262535587E-05

 2.99258481076663E-02
 1.57434103929355E-02

 5.82546037091492E-04
 1.92612667509151E-04

 2.13054758509304E-03
 1.38039078381558E-03

 2.72127544854818E-02
 1.41704069416107E-02

 4.81909572818664E-02
 2.33783625189232E-02

 7.29818911637093E-04
 2.78218297513218E-04

 1.14218296036478E-03
 3.77199807205421E-04

 1.27309218218309E-03
 4.6013026127186E-04

 1.34509225418316E-03
 4.49429557521352E-04

 1.97018378836561E-03
 7.75801021911858E-04

 2.33018414836597E-03
 1.0566944953627E-03

 3.32182150363969E-03
 1.70141189633083E-03

 3.63273090545818E-03
 2.18026838916608E-03

 9.38291847382756E-03
 5.56704112620199E-03

 1.09701927883746E-02
 5.09085980930436E-03

 1.20927393654666E-02
 5.44130785713351E-03

 1.16509207418298E-03
 3.93250862831183E-04

 1.16509207418298E-03
 3.93250862831183E-04

 6.70713397986125E-02
 5.52450582879372E-02

 1.08981927163745E-03
 6.04589761903724E-04

 7.71382589564408E-03
 5.91748917403114E-03

 1.25018306836489E-02
 1.08772653623918E-02

 1.56305610851065E-02
 1.36835249209626E-02

 3.01353028625756E-02
 2.41621890686479E-02

 1.94400194400194E-03
 9.55037809732873E-04

 1.94400194400194E-03
 9.55037809732873E-04

 .012102557557103
 3.46702801516472E-03

 2.37272964545692E-03
 6.95545743783045E-04

 3.01418483236665E-03
 6.68793984406774E-04

 2.80473007745735E-03
 1.19580364411931E-03

 3.91091300182209E-03
 9.06884642855586E-04

 3.04036667673031E-03
 7.81151373787112E-04

 3.04036667673031E-03
 7.81151373787112E-04

 .597342051887506
 .616325758742094

 2.39563875927512E-03
 1.98765572165693E-03

 2.2189113098204E-03
 2.14549110197693E-03

 3.87818569636751E-03
 2.32740306573557E-03

 6.7810976901886E-03
 4.71366000209894E-03

 2.20909311818403E-02
 3.46809808553977E-02

 .559977287250014
 .570470567995228

 8.88873616146343E-03
 2.57886960387252E-03

 4.17273144545872E-03
 1.17707741255592E-03

 4.71600471600472E-03
 1.4017921913166E-03

 1.39941958123776E-02
 6.63978667719045E-03

 3.86509477418568E-03
 1.46867158975728E-03

 1.01291010381919E-02
 5.17111508743318E-03

 .024712388348752
 .012503772332469

 3.69491278582188E-03
 2.1428159260393E-03

 6.45055190509736E-03
 3.2289373567159E-03

 1.45669236578327E-02
 7.13201904971384E-03

 1.81505636051091E-02
 9.55305327326636E-03

 5.09564145927782E-03
 2.08128687947388E-03

 1.30549221458312E-02
 7.47176639379248E-03

 5.38364174727811E-03
 2.11873934260066E-03

 5.38364174727811E-03
 2.11873934260066E-03

 .014082559537105
 6.41507189842978E-03

 5.62909653818745E-03
 3.06307644858302E-03

 8.45346299891754E-03
 3.35199544984675E-03

 2.24509315418406E-02
 8.96183939105077E-03

 6.50291559382468E-03
 2.41033351980201E-03

 1.59480159480159E-02
 6.55150587124876E-03

 6.20411529502439E-02
 1.71719543436283E-02

 9.36655482110028E-03
 1.92345149915388E-03

 8.08364444728081E-03
 4.00473837862776E-03

 2.22447495174768E-02
 2.13746557416405E-03

 .022346204164386
 9.10629889168263E-03

 6.27447900175173E-02
 3.75246928770953E-02

 6.27447900175173E-02
 3.75246928770953E-02

 .107280107280107
 1.61259605520161E-02

 .107280107280107
 1.61259605520161E-02

 6.70124306487943E-02
 1.13828736146033E-02

 3.27273054545782E-06
 5.35035187525419E-06

 3.01614847069393E-02
 6.19303229560673E-03

 3.68476732113096E-02
 5.18449096712131E-03

 .040267676631313
 4.74308693741284E-03

 .040267676631313
 4.74308693741284E-03

 .288023197114106
 .128737491646429

 .044420771693499
 2.63932858006289E-02

 1.30909221818313E-05
 0

 1.30909221818313E-05
 0

 1.60920160920161E-02
 7.84094067318502E-03

 1.50872878145605E-03
 5.18984131899657E-04

 2.04545659091114E-03
 1.0861214306766E-03

 1.25378307196489E-02
 6.23583511060876E-03

 1.10683747047383E-02
 3.33059404234573E-03

 1.10683747047383E-02
 3.33059404234573E-03

 1.72472899745627E-02
 1.52217510850982E-02

 1.72472899745627E-02
 1.52217510850982E-02

 .123463759827396
 6.16735060660551E-02

 .123463759827396
 6.16735060660551E-02

 2.45782063963882E-03
 6.17965641591859E-04

 3.47891256982166E-03
 1.36433972818982E-03

 3.70145824691279E-03
 1.70408707226846E-03

 4.54909545818637E-03
 2.75275603981828E-03

 4.81745936291391E-03
 2.56549372418439E-03

 3.93054938509484E-03
 3.33861957015862E-03

 5.00727773455046E-03
 2.72065392856676E-03

 5.54073281346009E-03
 2.53071643699523E-03

 5.85818767636949E-03
 3.01759845764336E-03

 6.40146094691549E-03
 2.9319928276393E-03

 6.72218854037036E-03
 3.19416006952675E-03

 8.47637211273575E-03
 3.02829916139387E-03

 7.79564415928052E-03
 4.05289154550505E-03

 8.88873616146343E-03
 4.22945315738844E-03

 1.00407373134646E-02
 5.37175328275521E-03

 1.06887379614652E-02
 4.90092231773284E-03

 1.14185568731023E-02
 4.61467849240674E-03

 1.36898318716501E-02
 8.7371246122901E-03

 1.18963755327392E-02
 6.18500676779385E-03

 4.45418627236809E-03
 3.53123223766777E-04

 4.45418627236809E-03
 3.53123223766777E-04

 7.44218926037108E-03
 5.83188354402707E-03

 7.44218926037108E-03
 5.83188354402707E-03

 3.70800370800371E-02
 .013135113853749

 3.70800370800371E-02
 .013135113853749

 4.50327723054996E-03
 1.08344625473897E-03

 6.6567339294612E-03
 3.13798137483658E-03

 9.88691897782807E-03
 3.08447785608404E-03

 1.60331069421979E-02
 5.82920836808944E-03

 5.56691465782375E-03
 6.28666345342367E-04

 5.56691465782375E-03
 6.28666345342367E-04

 5.56691465782375E-03
 6.28666345342367E-04

 1.54112881385609E-02
 5.03200593867657E-03

 1.54112881385609E-02
 5.03200593867657E-03

 5.36073263345991E-03
 1.78434235039727E-03

 .010050555505101
 3.24766358827929E-03

 2.62702080883899E-02
 8.54451194478095E-03

 2.62702080883899E-02
 8.54451194478095E-03

 7.18691627782537E-03
 1.59172968288812E-03

 1.90832918105645E-02
 6.95278226189282E-03

 1.10520110520111E-02
 2.28727542667117E-03

 1.10520110520111E-02
 2.28727542667117E-03

 1.10520110520111E-02
 2.28727542667117E-03

 1.28618310436492E-02
 4.85811950273081E-03

 1.28618310436492E-02
 4.85811950273081E-03

 1.28618310436492E-02
 4.85811950273081E-03

 3.44618526436708E-02
 1.89027931752731E-02

 1.30909221818313E-05
 0

 1.30909221818313E-05
 0

 1.30909221818313E-05
 0

 3.02073029345757E-03
 2.05453512009761E-03

 3.02073029345757E-03
 2.05453512009761E-03

 2.48727521454794E-04
 2.27389954698303E-04

 2.77200277200277E-03
 1.82714516539931E-03

 9.50400950400951E-03
 4.76983869678911E-03

 1.68545623091078E-03
 8.56056300040671E-04

 1.68545623091078E-03
 8.56056300040671E-04

 7.81855327309873E-03
 3.91378239674844E-03

 7.81855327309873E-03
 3.91378239674844E-03

 2.19240219240219E-02
 1.20784193583863E-02

 2.19240219240219E-02
 1.20784193583863E-02

 6.33273360546088E-03
 3.44027625578845E-03

 6.82364318727955E-03
 3.98601214706437E-03

 8.7676451312815E-03
 4.65213095553352E-03

 .666966121511576
 9.21410848196901E-02

 .109914655369201
 6.85273068182557E-02

 9.09786364331819E-02
 5.51514271301202E-02

 9.81819163637346E-06
 8.02552781288129E-06

 3.27273054545782E-05
 8.02552781288129E-06

 6.87273414546142E-05
 1.33758796881355E-05

 5.4000054000054E-04
 3.9592603876881E-04

 1.12254657709203E-03
 6.28666345342367E-04

 2.78182096363915E-03
 2.9614197629532E-03

 3.91091300182209E-03
 2.17491803729083E-03

 5.96291505382414E-03
 3.62486339548472E-03

 6.16255161709707E-03
 3.48307907079048E-03

 8.42400842400842E-03
 4.28295667614098E-03

 8.93128165855439E-03
 5.40920574588199E-03

 1.56927429654702E-02
 1.05776456573775E-02

 1.87036550672914E-02
 9.98108142328669E-03

 1.86349277258368E-02
 1.16022380414887E-02

 5.12182330364149E-03
 4.04219084175454E-03

 5.12182330364149E-03
 4.04219084175454E-03

 1.38141956323775E-02
 9.33368884638094E-03

 1.38141956323775E-02
 9.33368884638094E-03

 .557051466142375
 2.36137780014344E-02

 .536380900017264
 1.43255671459931E-02

 5.23636887273251E-05
 0

 4.48364084727721E-04
 9.89815096922026E-05

 1.8000018000018E-03
 2.80893473450845E-04

 2.39236602872967E-03
 2.56816890012201E-04

 3.11563947927584E-03
 2.22039602823049E-04

 3.49527622254895E-03
 1.4713467656949E-04

 6.59127931855205E-03
 3.04970056889489E-04

 6.51927924655197E-03
 3.79874983143048E-04

 8.55819037637219E-03
 5.6178694690169E-04

 9.95891904982814E-03
 7.59749966286095E-04

 3.61898543716726E-02
 6.68793984406774E-04

 8.91917255553619E-02
 3.2289373567159E-03

 .170129624675079
 6.86450145595113E-03

 .197938016119834
 5.51086243151182E-04

 .002637820819639
 1.11287319005287E-03

 1.8000018000018E-04
 5.35035187525419E-05

 2.45782063963882E-03
 1.05936967130033E-03

 2.11745666291121E-03
 7.19622327221689E-04

 2.11745666291121E-03
 7.19622327221689E-04

 1.59152886425614E-02
 7.45571533816672E-03

 4.56545911091366E-03
 2.14549110197693E-03

 5.43927816655089E-03
 2.15084145385219E-03

 5.91055136509682E-03
 3.1593827823376E-03

 2.30203866567503E-02
 1.40072212094155E-02

 1.71621989803808E-02
 1.09896227517721E-02

 9.81819163637346E-06
 1.60510556257626E-05

 9.81819163637346E-06
 1.60510556257626E-05

 1.71523807887444E-02
 1.09735716961463E-02

 2.19927492654765E-03
 1.03529308786169E-03

 3.04036667673031E-03
 1.43924465444338E-03

 1.19127391854665E-02
 8.49903395384128E-03

 5.85818767636949E-03
 3.01759845764336E-03

 7.06909797818889E-04
 1.8458713969627E-04

 7.06909797818889E-04
 1.8458713969627E-04

 5.15127787855061E-03
 2.83301131794709E-03

 5.15127787855061E-03
 2.83301131794709E-03

 .178242723697269
 6.31849804708144E-02

 .17667835849654
 6.27248502095425E-02

 .108556472192836
 3.93304366349936E-02

 4.25454970909516E-05
 5.35035187525419E-06

 4.25454970909516E-05
 2.94269353138981E-05

 1.57091066181975E-04
 1.39109148756609E-04

 1.47272874545602E-04
 1.49809852507117E-04

 2.32363868727505E-04
 8.293045406644E-05

 8.80364516728153E-04
 2.59492065949828E-04

 8.90182708364527E-04
 3.98601214706437E-04

 9.72000972000972E-04
 4.36053677833217E-04

 9.91637355273719E-04
 7.62425142223722E-04

 2.0323656687293E-03
 2.35415482511184E-04

 2.02909293818385E-03
 5.02933076273894E-04

 1.99309290218381E-03
 6.12615289716605E-04

 2.49709340618432E-03
 5.72487650652199E-04

 2.29418411236593E-03
 8.1057830910101E-04

 3.39382157563976E-03
 8.15928660976264E-04

 3.37418519236701E-03
 1.06739519911321E-03

 3.45927618654891E-03
 1.0272675600488E-03

 4.58182276364095E-03
 7.59749966286095E-04

 4.91564127927764E-03
 1.84052104508744E-03

 9.82800982800983E-03
 3.33861957015862E-03

 1.10781928963747E-02
 4.57990120521759E-03

 1.41643778007414E-02
 3.96996109143861E-03

 .016563289290562
 7.77138609880671E-03

 .021996021996022
 9.1624775863728E-03

 1.30909221818313E-04
 1.04331861567457E-04

 1.30909221818313E-04
 1.04331861567457E-04

 3.80225834771289E-02
 1.35497661240812E-02

 1.48581966763785E-03
 2.91594177201353E-04

 5.06291415382324E-03
 9.04209466917958E-04

 6.94800694800695E-03
 2.0438344163471E-03

 2.45258427076609E-02
 1.03101280636148E-02

 5.7960057960058E-03
 5.40385539400673E-04

 5.7960057960058E-03
 5.40385539400673E-04

 2.41723878087514E-02
 9.19993004949958E-03

 6.33600633600634E-03
 2.56816890012201E-03

 1.78363814727451E-02
 6.63176114937757E-03

 1.56436520072884E-03
 4.6013026127186E-04

 1.56436520072884E-03
 4.6013026127186E-04

 1.56436520072884E-03
 4.6013026127186E-04

 2.93858475676657E-02
 1.54277396322955E-02

 2.93858475676657E-02
 1.54277396322955E-02

 8.83637247273611E-05
 0

 8.83637247273611E-05
 0

 .019008019008019
 1.03689819342426E-02

 6.61091570182479E-04
 3.42422520016268E-04

 9.72000972000972E-04
 3.07645232827116E-04

 1.07672834945562E-03
 3.15670760639997E-04

 1.00800100800101E-03
 4.62805437209488E-04

 1.20436484072848E-03
 4.81531668772877E-04

 1.01454646909192E-03
 6.66118808469147E-04

 1.32218314036496E-03
 5.6178694690169E-04

 1.82618364436546E-03
 8.77457707541687E-04

 1.97672924945652E-03
 9.25610874418975E-04

 2.07818389636571E-03
 8.98859115042704E-04

 5.86800586800587E-03
 4.52907286240267E-03

 7.42909833818925E-04
 3.42422520016268E-04

 7.42909833818925E-04
 3.42422520016268E-04

 3.8945493490948E-03
 2.20434497260473E-03

 3.8945493490948E-03
 2.20434497260473E-03

 5.65200565200565E-03
 2.51199020543184E-03

 5.65200565200565E-03
 2.51199020543184E-03

 .293747202838112
 .105380530535007

 .293747202838112
 .105380530535007

 5.23636887273251E-04
 2.6751759376271E-06

 5.23636887273251E-04
 2.6751759376271E-06

 3.99665854211309E-02
 9.04744502105484E-03

 6.95455240909786E-03
 9.81789569109144E-04

 1.51658333476515E-02
 3.33059404234573E-03

 1.78461996643815E-02
 4.73506140959996E-03

 2.85774831229377E-02
 8.43482973133823E-03

 6.91527964255237E-03
 1.69606154445558E-03

 2.16622034803853E-02
 6.73876818688265E-03

 6.54644291007927E-02
 3.09598111261584E-02

 7.37018918837101E-03
 2.17491803729083E-03

 2.48662066843885E-02
 .011492555828046

 3.32280332280332E-02
 1.72923372608215E-02

 .106848106848107
 3.12540804792974E-02

 7.05600705600706E-03
 2.64039865043794E-03

 1.59512886785614E-02
 9.25878392012738E-03

 8.38408111135384E-02
 .019354897908732

 8.38800838800839E-03
 3.36804650547251E-03

 8.38800838800839E-03
 3.36804650547251E-03

 2.29614775069321E-02
 9.78044322796466E-03

 2.29614775069321E-02
 9.78044322796466E-03

 2.10174755629301E-02
 1.25331992677829E-02

 2.10174755629301E-02
 1.25331992677829E-02

 3.67691276782186E-02
 2.78833587978872E-02

 3.67691276782186E-02
 2.78833587978872E-02

 2.27814773269319E-02
 1.78782007911619E-02

 7.06909797818889E-04
 2.19364426885422E-04

 1.7378199196381E-03
 1.50612405288405E-03

 3.28909419818511E-03
 2.65109935418845E-03

 3.67854913309459E-03
 3.34932027390912E-03

 5.21018702836885E-03
 4.19200069426166E-03

 8.15891724982634E-03
 5.96029198903317E-03

 7.39309830218921E-03
 4.21875245363793E-03

 1.14218296036478E-03
 6.23315993467113E-04

 6.25091534182443E-03
 3.59543646017082E-03

 6.5945520490975E-03
 5.78640555308741E-03

 6.5945520490975E-03
 5.78640555308741E-03

 5.3770962861872E-03
 4.2749311483281E-03

 5.3770962861872E-03
 4.2749311483281E-03

 5.3770962861872E-03
 4.2749311483281E-03

 1.25018306836489E-03
 9.81789569109144E-04

 4.12691321782231E-03
 3.29314157921896E-03

 6.04931514022423E-02
 3.20914105477746E-02

 1.51396515032879E-02
 6.09405078591452E-03

 9.73637337273701E-03
 3.72652008111454E-03

 1.7149108058199E-03
 6.39367049092876E-04

 8.02146256691711E-03
 3.08715303202167E-03

 5.40327813055086E-03
 2.36753070479998E-03

 5.40327813055086E-03
 2.36753070479998E-03

 .019587292314565
 9.16782793824806E-03

 1.44065598611053E-02
 7.86769243256129E-03

 1.97345651891106E-03
 1.0165668562983E-03

 1.24331033421943E-02
 6.85112557626299E-03

 2.30727503454776E-03
 8.80132883479315E-04

 2.30727503454776E-03
 8.80132883479315E-04

 2.87345741891196E-03
 4.20002622207454E-04

 2.87345741891196E-03
 4.20002622207454E-04

 1.17360117360117E-02
 7.0785155309613E-03

 4.05164041527678E-03
 1.84052104508744E-03

 4.05164041527678E-03
 1.84052104508744E-03

 7.68437132073496E-03
 5.23799448587385E-03

 7.68437132073496E-03
 5.23799448587385E-03

 1.40301958483777E-02
 9.75101629265076E-03

 1.40301958483777E-02
 9.75101629265076E-03

 1.40301958483777E-02
 9.75101629265076E-03

 1.24985579531034E-02
 5.80245660871317E-03

 1.24985579531034E-02
 5.80245660871317E-03

 2.29418411236593E-03
 1.02459238411118E-03

 2.29418411236593E-03
 1.02459238411118E-03

 1.02043738407375E-02
 4.77786422460199E-03

 1.02043738407375E-02
 4.77786422460199E-03

 4.97356860993225E-02
 .029047060330755

 2.01502019683838E-02
 1.27365126390426E-02

 1.41676505312869E-02
 9.38719236513348E-03

 1.91454736909282E-03
 1.50344887694643E-03

 3.58363994727631E-03
 1.78969270227253E-03

 8.66946321491776E-03
 6.09405078591452E-03

 5.98255143709689E-03
 3.34932027390912E-03

 5.98255143709689E-03
 3.34932027390912E-03

 2.64894810349356E-02
 1.45904095638182E-02

 8.89528162255435E-03
 4.98117759586165E-03

 2.28436592072956E-03
 1.39109148756609E-03

 6.61091570182479E-03
 3.59008610829556E-03

 3.97309488218579E-03
 2.0144074810332E-03

 3.97309488218579E-03
 2.0144074810332E-03

 5.02691411782321E-03
 2.98014599451658E-03

 5.02691411782321E-03
 2.98014599451658E-03

 8.59419041237223E-03
 4.61467849240674E-03

 8.59419041237223E-03
 4.61467849240674E-03

 3.0960030960031E-03
 1.72013812789422E-03

 3.0960030960031E-03
 1.72013812789422E-03

 3.0960030960031E-03
 1.72013812789422E-03

 1.28781946963765E-02
 6.99558507689486E-03

 1.28781946963765E-02
 6.99558507689486E-03

 2.77200277200277E-03
 1.13694977349152E-03

 2.77200277200277E-03
 1.13694977349152E-03

 1.01061919243737E-02
 5.85863530340334E-03

 1.01061919243737E-02
 5.85863530340334E-03

 9.28473655746383E-03
 7.72323293192943E-03

 9.28473655746383E-03
 7.72323293192943E-03

 9.28473655746383E-03
 7.72323293192943E-03

 1.30909221818313E-05
 1.07007037505084E-05

 1.30909221818313E-05
 1.07007037505084E-05

 .009271645635282
 7.71253222817892E-03

 .009271645635282
 7.71253222817892E-03

 .211870030051848
 .145912121165995

 .189298007479826
 .129759408854602

 1.16836480472844E-03
 6.55418104718639E-04

 1.16836480472844E-03
 6.55418104718639E-04

 1.16836480472844E-03
 6.55418104718639E-04

 6.32618814436996E-02
 4.22249769995061E-02

 4.08698590516772E-02
 2.54328976390208E-02

 1.36145590691045E-03
 6.04589761903724E-04

 1.4040014040014E-03
 6.23315993467113E-04

 1.13236476872841E-03
 1.54625169194846E-03

 1.95382013563832E-03
 8.88158411292196E-04

 1.7149108058199E-03
 1.38039078381558E-03

 2.0323656687293E-03
 1.65593390539117E-03

 2.14036577672941E-03
 1.66930978507931E-03

 3.58036721673085E-03
 2.17759321322846E-03

 1.08818290636472E-02
 7.16679633690299E-03

 1.46683783047419E-02
 7.7205577559918E-03

 9.9851008941918E-03
 8.40272762008671E-03

 1.86545641091096E-03
 1.3429383206888E-03

 8.11964448328085E-03
 7.05978929939791E-03

 2.83418465236647E-03
 2.15886698166507E-03

 2.83418465236647E-03
 2.15886698166507E-03

 9.57273684546412E-03
 6.23048475873351E-03

 9.57273684546412E-03
 6.23048475873351E-03

 .124867761231398
 8.68790137503776E-02

 .103784831057558
 7.03196746964658E-02

 2.46436610072974E-03
 8.66757003791179E-04

 2.9323665687302E-03
 1.45262053413151E-03

 1.98000198000198E-03
 2.35683000104947E-03

 3.35454880909426E-03
 2.48523844605557E-03

 4.84036847673211E-03
 3.2583642920298E-03

 5.30836894473258E-03
 3.46702801516472E-03

 7.45528018255291E-03
 5.42258162557012E-03

 1.07476471112835E-02
 6.3187655646752E-03

 1.12385566931021E-02
 7.22565020753079E-03

 2.39105693651148E-02
 1.64175547292175E-02

 2.95527568254841E-02
 .02104828427725

 1.51298333116515E-02
 1.12089871786575E-02

 5.15455060909606E-03
 4.71366000209894E-03

 9.97528270255543E-03
 6.49532717655859E-03

 5.95309686218777E-03
 5.35035187525419E-03

 5.95309686218777E-03
 5.35035187525419E-03

 6.11018792836975E-03
 3.27174017171794E-03

 6.11018792836975E-03
 3.27174017171794E-03

 1.18800118800119E-03
 6.84845040032537E-04

 1.18800118800119E-03
 6.84845040032537E-04

 4.92218674036856E-03
 2.5868951316854E-03

 4.92218674036856E-03
 2.5868951316854E-03

 1.64618346436528E-02
 1.28809721396745E-02

 1.64618346436528E-02
 1.28809721396745E-02

 8.08691717782627E-03
 6.41239672249215E-03

 8.08691717782627E-03
 6.41239672249215E-03

 8.37491746582656E-03
 6.46857541718232E-03

 8.37491746582656E-03
 6.46857541718232E-03

 .766509493782221
 .544679196780565

 5.67000567000567E-02
 3.75113169974071E-02

 2.08800208800209E-03
 1.11287319005287E-03

 2.08800208800209E-03
 1.11287319005287E-03

 2.08800208800209E-03
 1.11287319005287E-03

 3.81698563516745E-02
 2.55693316118398E-02

 2.57236620872985E-02
 1.74448222892663E-02

 2.09782027963846E-03
 1.32688726506304E-03

 1.00211009301918E-02
 6.85112557626299E-03

 1.36047408774682E-02
 9.26680944794026E-03

 1.24461942643761E-02
 8.12450932257349E-03

 1.24461942643761E-02
 8.12450932257349E-03

 2.50363886727523E-03
 1.84854657290032E-03

 2.50363886727523E-03
 1.84854657290032E-03

 2.50363886727523E-03
 1.84854657290032E-03

 6.2770971861881E-03
 4.22677798145081E-03

 6.2770971861881E-03
 4.22677798145081E-03

 6.2770971861881E-03
 4.22677798145081E-03

 7.66146220691675E-03
 4.75378764116335E-03

 7.66146220691675E-03
 4.75378764116335E-03

 7.66146220691675E-03
 4.75378764116335E-03

 6.26727899455172E-03
 3.72384490517692E-03

 6.26727899455172E-03
 3.72384490517692E-03

 6.26727899455172E-03
 3.72384490517692E-03

 2.16982035163853E-03
 1.31083620943728E-03

 4.09745864291319E-03
 2.41300869573964E-03

 .4091469546015
 .294887318780572

 1.12745567291022E-02
 7.27112819847045E-03

 3.41673068945796E-03
 2.19096909291659E-03

 3.41673068945796E-03
 2.19096909291659E-03

 7.85782603964422E-03
 5.08015910555386E-03

 7.85782603964422E-03
 5.08015910555386E-03

 .383933838479293
 .277651160214441

 .354691991055627
 .250912776717858

 3.64582182764001E-03
 2.16956768541557E-03

 3.64582182764001E-03
 2.72867945637964E-03

 6.60764297127934E-03
 4.05021636956742E-03

 8.26364462728099E-03
 5.60181841339114E-03

 1.16934662389208E-02
 9.0340691413667E-03

 5.28153255425983E-02
 2.27924989885829E-02

 6.31047903775176E-02
 .047706412495704

 .102485557031012
 6.92389036176645E-02

 .102429920611739
 8.75906105497864E-02

 2.92418474236656E-02
 2.67383834965828E-02

 2.92418474236656E-02
 2.67383834965828E-02

 1.39385593931048E-02
 9.96503036766093E-03

 1.39385593931048E-02
 9.96503036766093E-03

 1.39385593931048E-02
 9.96503036766093E-03

 .161018342836525
 .112651658733477

 .10640628822447
 7.28985443003384E-02

 .10640628822447
 7.28985443003384E-02

 3.85854931309477E-03
 2.02778336072134E-03

 4.06145860691315E-03
 2.91326659607591E-03

 6.3850972941882E-03
 4.74576211335047E-03

 1.16247388974662E-02
 7.56004719973417E-03

 1.31858313676495E-02
 9.48082352295043E-03

 1.85498367316549E-02
 1.30200812884311E-02

 2.07491116582026E-02
 1.34240328550128E-02

 2.79916643553007E-02
 1.97267473640622E-02

 1.08523744887381E-02
 8.98859115042704E-03

 1.08523744887381E-02
 8.98859115042704E-03

 1.08523744887381E-02
 8.98859115042704E-03

 4.37596801233165E-02
 3.07645232827116E-02

 4.37596801233165E-02
 3.07645232827116E-02

 2.18716582352946E-02
 1.48900292688324E-02

 2.18880218880219E-02
 1.58744940138792E-02

 7.82444418808055E-02
 5.83134850883954E-02

 2.80374825829371E-02
 2.27149188863917E-02

 6.16909707818799E-03
 4.45149276021149E-03

 6.16909707818799E-03
 4.45149276021149E-03

 2.18683855047491E-02
 1.82634261261802E-02

 2.18683855047491E-02
 1.82634261261802E-02

 5.02069592978684E-02
 3.55985662020038E-02

 1.24036487672851E-02
 9.17317829012331E-03

 1.24036487672851E-02
 9.17317829012331E-03

 1.85465640011095E-02
 1.10431262705247E-02

 1.85465640011095E-02
 1.10431262705247E-02

 1.92567465294738E-02
 1.53822616413558E-02

 1.92567465294738E-02
 1.53822616413558E-02

 7.01018882837065E-03
 4.02346461019115E-03

 7.01018882837065E-03
 4.02346461019115E-03

 7.01018882837065E-03
 4.02346461019115E-03

 7.01018882837065E-03
 4.02346461019115E-03

 7.64182582364401E-03
 5.35035187525419E-03

 7.64182582364401E-03
 5.35035187525419E-03

 7.64182582364401E-03
 5.35035187525419E-03

 7.64182582364401E-03
 5.35035187525419E-03

 4.04804041167678E-02
 2.82177557900906E-02

 1.95545650091105E-02
 1.44994535819389E-02

 1.95545650091105E-02
 1.44994535819389E-02

 1.95545650091105E-02
 1.44994535819389E-02

 2.09258391076573E-02
 1.37183022081517E-02

 2.09258391076573E-02
 1.37183022081517E-02

 2.09258391076573E-02
 1.37183022081517E-02

 2.12400212400212E-03
 1.26000786662236E-03

 2.12400212400212E-03
 1.26000786662236E-03

 2.12400212400212E-03
 1.26000786662236E-03

 2.12400212400212E-03
 1.26000786662236E-03

 2.12400212400212E-03
 1.26000786662236E-03

 32.9801533437897
 25.504859871743

 2.88140360867634
 1.79672573981255

 1.4703298339662
 .904367302298278

 .904608904608905
 .551388538032134

 .632828269191905
 .398919560643015

 3.27273054545782E-06
 5.35035187525419E-06

 1.93091102182011E-04
 9.89815096922026E-05

 2.29091138182047E-04
 1.25733269068473E-04

 1.62654708109254E-03
 1.26803339443524E-03

 3.66545821091276E-03
 2.2712243710454E-03

 4.6865501410956E-03
 1.77899199852202E-03

 4.49345903891358E-03
 2.16956768541557E-03

 4.36909527818619E-03
 2.73402980825489E-03

 6.74509765418856E-03
 4.19735104613691E-03

 1.10978292796475E-02
 8.01750228506841E-03

 1.42887415614688E-02
 5.48411067213555E-03

 1.29632856905584E-02
 8.29037023070637E-03

 1.37978319796502E-02
 9.03941949324196E-03

 1.80752908025635E-02
 .010746181741448

 2.12465667011122E-02
 1.26509070090385E-02

 2.22152949425677E-02
 1.40259474409789E-02

 2.53865708411163E-02
 1.43951217203714E-02

 2.44342062523881E-02
 1.71345018805016E-02

 3.23803960167597E-02
 .017375267714888

 .033048033048033
 1.93441972049815E-02

 3.57153084425812E-02
 2.12783494078859E-02

 3.71880371880372E-02
 2.33783625189232E-02

 4.53698635516817E-02
 2.83140621238452E-02

 .044728408364772
 3.02615902064377E-02

 5.77407850135123E-02
 .037318704329898

 5.99826054371509E-02
 3.93598635703075E-02

 9.71575517030062E-02
 6.78558376579113E-02

 .16595034776853
 9.29677141844168E-02

 2.29091138182047E-05
 5.08283428149148E-05

 4.61455006909552E-04
 1.28408445006101E-04

 4.91236854873218E-03
 2.95071905920269E-03

 8.3094628549174E-03
 5.12831227243114E-03

 1.21549212458303E-02
 5.6767233396447E-03

 1.96200196200196E-02
 9.72426453327449E-03

 2.37272964545692E-02
 1.36059448187714E-02

 2.59527532254805E-02
 1.53046815391646E-02

 2.63192990465718E-02
 1.51414958069694E-02

 4.44698626516808E-02
 2.52563360271374E-02

 2.72749363658455E-02
 1.75170520395822E-02

 3.59673086945814E-03
 1.69071119258032E-03

 2.36782054963873E-02
 1.58263408470019E-02

 3.81338563156745E-02
 .021658224391029

 3.81338563156745E-02
 .021658224391029

 4.04214949669495E-02
 2.03259867740907E-02

 4.04214949669495E-02
 2.03259867740907E-02

 .112532839805567
 6.29254884048646E-02

 9.92488265215538E-02
 .055820221114527

 3.27273054545782E-06
 5.35035187525419E-06

 1.63636527272891E-05
 8.02552781288129E-06

 6.54546109091564E-06
 2.14014075010168E-05

 3.8945493490948E-04
 1.41784324694236E-04

 9.26182744364563E-04
 3.58473575642031E-04

 1.79345633891088E-03
 6.39367049092876E-04

 1.81636545272909E-03
 9.47012281919992E-04

 3.75054920509466E-03
 2.22574638010574E-03

 5.59636923273287E-03
 3.42957555203794E-03

 1.15560115560116E-02
 6.17430606404334E-03

 1.33429224338315E-02
 7.63495212598773E-03

 1.45701963883782E-02
 8.31712199008264E-03

 1.76301994483813E-02
 1.06338243520677E-02

 .027850936941846
 1.52832801316636E-02

 1.32840132840133E-02
 7.10526729033757E-03

 1.32840132840133E-02
 7.10526729033757E-03

 5.25469616378707E-02
 2.88785242466845E-02

 4.53698635516817E-02
 2.56148096027794E-02

 1.30909221818313E-04
 2.54141714074574E-04

 1.97574743029288E-02
 1.00854132848542E-02

 2.54814800269346E-02
 1.52752546038507E-02

 7.17709808618899E-03
 3.26371464390506E-03

 7.17709808618899E-03
 3.26371464390506E-03

 .107633562179017
 .069407439701735

 .107633562179017
 .069407439701735

 6.4472791745519E-04
 3.55798399704404E-04

 1.24036487672851E-03
 7.54399614410841E-04

 4.96473223745951E-03
 2.57084407605964E-03

 6.7810976901886E-03
 4.10104471238234E-03

 1.68840168840169E-02
 1.02860514801762E-02

 1.68381986563805E-02
 1.05776456573775E-02

 6.02804239167876E-02
 4.07616557616241E-02

 .246148609784973
 .164322681968744

 .214792578428942
 .144628036715934

 5.43927816655089E-03
 4.13314682363386E-03

 1.01094646549192E-02
 4.64410542772064E-03

 1.54734700189246E-02
 1.04626130920596E-02

 1.93876557512921E-02
 1.14631288927321E-02

 4.75004111367748E-02
 3.11684748492933E-02

 5.45236908873273E-02
 3.83994754086993E-02

 6.23586078131533E-02
 4.43570922217949E-02

 3.13560313560314E-02
 1.96946452528107E-02

 3.13560313560314E-02
 1.96946452528107E-02

 .046858955949865
 2.74446299441164E-02

 1.98982017163835E-02
 1.09468199367701E-02

 1.98982017163835E-02
 1.09468199367701E-02

 2.69607542334815E-02
 1.64978100073463E-02

 2.69607542334815E-02
 1.64978100073463E-02

 1.41107377471014
 .89235843751427

 .365603274694184
 .226705109658271

 .345011254102163
 .215236630413663

 6.54546109091564E-06
 2.6751759376271E-06

 2.61818443636625E-05
 1.07007037505084E-05

 6.64364300727937E-04
 4.46754381583725E-04

 7.98546253091708E-04
 5.02933076273894E-04

 1.05381923563742E-03
 7.2764785503457E-04

 2.00291109382018E-03
 1.13962494942914E-03

 2.14363850727487E-03
 1.15567600505491E-03

 2.46436610072974E-03
 9.17585346606094E-04

 2.40872968145695E-03
 1.42051842287999E-03

 2.66400266400266E-03
 1.48472264538304E-03

 4.6472773745501E-03
 2.46651221449218E-03

 7.12146166691621E-03
 4.33646019489352E-03

 7.90037153673517E-03
 5.28882282868877E-03

 9.64800964800965E-03
 7.54399614410841E-03

 1.12712839985567E-02
 6.40437119467927E-03

 1.39876503512867E-02
 6.74946889063316E-03

 1.32807405534678E-02
 7.62425142223722E-03

 1.35785590331045E-02
 7.74195916349282E-03

 1.45505600051055E-02
 9.37381648544534E-03

 1.52411061501971E-02
 9.69216242202297E-03

 1.59349250258341E-02
 9.65203478295856E-03

 1.67007439734712E-02
 1.04947152033111E-02

 .017806926897836
 1.19794378486941E-02

 1.93025647571102E-02
 1.17921755330602E-02

 .021609839791658
 1.33972810956365E-02

 2.27192954465682E-02
 .014312191266305

 2.46796610432974E-02
 1.63613760345273E-02

 2.98342116523935E-02
 1.80092844121056E-02

 5.09629600538691E-02
 3.42074747144377E-02

 2.05920205920206E-02
 1.14684792446074E-02

 1.96363832727469E-05
 0

 4.40182258364077E-03
 2.45313633480405E-03

 1.61705616251071E-02
 9.01534290980331E-03

 .232141323050414
 .152709743223505

 3.27273054545782E-05
 1.60510556257626E-05

 3.27273054545782E-05
 1.60510556257626E-05

 .232108595744959
 .152693692167879

 3.34800334800335E-03
 1.99300607353219E-03

 8.09346263891718E-03
 4.54244874209081E-03

 1.06429197338288E-02
 5.69544957120809E-03

 9.77433704706432E-02
 6.03813960881812E-02

 .112280839553567
 8.00813916928671E-02

 .654536290899927
 .415310363612856

 .601210419392238
 .383473094779156

 1.45309236218327E-03
 7.73125845974231E-04

 2.04872932145659E-03
 1.36969008006507E-03

 2.56909347818439E-03
 1.57835380319999E-03

 2.86691195782105E-03
 1.54090134007321E-03

 3.22036685673049E-03
 1.83249551727456E-03

 4.36909527818619E-03
 2.2605236672949E-03

 4.63745918291373E-03
 2.43441010324066E-03

 4.38218620036802E-03
 3.4161996723498E-03

 5.74036937673301E-03
 3.00422257795523E-03

 5.61273288546016E-03
 3.14868207858709E-03

 4.82073209345937E-03
 4.2160772777003E-03

 6.91200691200691E-03
 4.02346461019115E-03

 .013925468470923
 8.98859115042704E-03

 2.02974748429294E-02
 1.12999431605369E-02

 .021881476426931
 1.18109017646236E-02

 2.36912964185691E-02
 1.63399746270263E-02

 2.72913000185727E-02
 .014017921913166

 2.82567555294828E-02
 .019922035207509

 2.86560286560287E-02
 2.01868776253341E-02

 .03366985185167
 2.28326266276473E-02

 2.95822114003932E-02
 2.64200375600052E-02

 4.64662282844101E-02
 1.31752414928134E-02

 4.12036775673139E-02
 2.56094592509042E-02

 6.17826072371527E-02
 3.90869956246695E-02

 7.95731704822614E-02
 6.06890413210083E-02

 9.63000963000963E-02
 6.34953008795791E-02

 5.33258715076897E-02
 3.18372688337001E-02

 5.33258715076897E-02
 3.18372688337001E-02

 3.87818569636751E-02
 2.42798968099035E-02

 3.04691213782123E-03
 1.48472264538304E-03

 3.04691213782123E-03
 1.48472264538304E-03

 2.13480213480213E-02
 1.39162652275362E-02

 2.76873004145731E-03
 1.95555361040541E-03

 3.49200349200349E-03
 2.11873934260066E-03

 3.67854913309459E-03
 2.11606416666303E-03

 .011408738681466
 7.72590810786705E-03

 1.43869234778326E-02
 8.87890893698433E-03

 1.43869234778326E-02
 8.87890893698433E-03

 .101480828753556
 6.36023079170842E-02

 4.23491332582242E-03
 2.03045853665897E-03

 4.23491332582242E-03
 2.03045853665897E-03

 6.36546091091546E-03
 3.61148751579658E-03

 6.36546091091546E-03
 3.61148751579658E-03

 9.08804545168182E-02
 5.79603618646287E-02

 9.08804545168182E-02
 5.79603618646287E-02

 1.85302003483822E-02
 9.75101629265076E-03

 1.33691042781952E-02
 6.59430868625079E-03

 5.50473277746005E-03
 2.45581151074167E-03

 7.86437150073514E-03
 4.13849717550912E-03

 5.16109607018698E-03
 3.15670760639997E-03

 5.16109607018698E-03
 3.15670760639997E-03

 27.3774219228765
 21.8240103942356

 .842679024497206
 .520891532343185

 .472124108487745
 .288680910605278

 .155693610239065
 9.24005768856399E-02

 1.96363832727469E-05
 8.02552781288129E-06

 2.37927510654783E-03
 1.78166717445965E-03

 4.66036829673193E-03
 2.46918739042981E-03

 6.19527892255165E-03
 3.32524369047048E-03

 7.21637085273449E-03
 3.13798137483658E-03

 1.37814683269229E-02
 8.87890893698433E-03

 1.85596549232913E-02
 9.91420202484602E-03

 2.94840294840295E-02
 1.83276303486832E-02

 7.33975279429825E-02
 4.45577304171169E-02

 1.59938341756524E-02
 8.98324079855179E-03

 1.59938341756524E-02
 8.98324079855179E-03

 6.36153363426091E-02
 4.08098089285013E-02

 6.36153363426091E-02
 4.08098089285013E-02

 7.28673455946183E-02
 4.69118852422288E-02

 7.28673455946183E-02
 4.69118852422288E-02

 .1639539821358
 9.95753987503558E-02

 .1639539821358
 9.95753987503558E-02

 .361266906721452
 .226574026037327

 .253407526134799
 .159699977948525

 5.04655050109596E-03
 2.54141714074574E-03

 1.04301922483741E-02
 6.44182365780605E-03

 1.12025566571021E-02
 7.32463171722299E-03

 1.36276499912864E-02
 8.74515014010298E-03

 1.43247415974689E-02
 8.20208942476468E-03

 1.40498322316504E-02
 8.77190189947925E-03

 3.62094907549453E-02
 1.91703107690358E-02

 .148516512152876
 9.85026531993673E-02

 4.19007691734964E-02
 2.69497223956554E-02

 4.19007691734964E-02
 2.69497223956554E-02

 6.59586114131569E-02
 3.99243256931468E-02

 6.59586114131569E-02
 3.99243256931468E-02

 9.28800928800929E-03
 5.63659570058029E-03

 9.28800928800929E-03
 5.63659570058029E-03

 9.28800928800929E-03
 5.63659570058029E-03

 26.4355366173548
 21.2400061111719

 4.11789684516957
 2.99452774035727

 .850448486812123
 .495274047564468

 0
 2.6751759376271E-05

 .253806799261345
 .139451571276625

 .596641687550779
 .355795724528466

 5.14473241745969E-03
 2.75810639169354E-03

 5.14473241745969E-03
 2.75810639169354E-03

 4.13607686334959E-02
 2.51332779340066E-02

 4.13607686334959E-02
 2.51332779340066E-02

 9.06284542648179E-02
 5.76018882889866E-02

 4.06898588716771E-02
 2.65404204771984E-02

 4.99385953931408E-02
 3.10614678117882E-02

 7.16269807178898E-02
 4.32816714948688E-02

 7.16269807178898E-02
 4.32816714948688E-02

 7.67651676742586E-02
 6.32839619805066E-02

 7.67651676742586E-02
 6.32839619805066E-02

 .582794764612946
 .467644830480655

 .230783139874049
 .183188022680891

 .352011624738897
 .284456807799764

 .293390475208657
 .189686025033387

 .293390475208657
 .189686025033387

 .734129097765461
 .485341119308058

 .394877849423304
 .204560003246594

 .339251248342157
 .280781116061465

 1.37160791706246
 1.16452281188064

 .715801806710898
 .500059937316883

 .655806110351565
 .664462874563756

 1.70181988363807E-02
 1.03261791192406E-02

 1.70181988363807E-02
 1.03261791192406E-02

 4.58836822473186E-03
 2.50396467761896E-03

 1.24298306116488E-02
 7.82221444162163E-03

 .202807839171476
 .130532534700577

 4.90975036429582E-02
 2.87795427369923E-02

 4.62764099127736E-03
 2.5868951316854E-03

 6.89564325927962E-03
 3.61683786767183E-03

 9.58582776764595E-03
 5.25672071743724E-03

 1.33789224698316E-02
 9.04209466917958E-03

 1.46094691549237E-02
 8.27699435101823E-03

 7.86600786600787E-02
 5.54269702516958E-02

 4.86327759055032E-03
 2.97479564264133E-03

 5.13491422582332E-03
 4.27760632426573E-03

 8.34873562146289E-03
 6.30538968498707E-03

 .010744374380738
 6.56755692687452E-03

 4.95687768415041E-02
 3.53016216729272E-02

 1.16411025501935E-02
 5.79710625683792E-03

 1.16411025501935E-02
 5.79710625683792E-03

 6.34091543182452E-02
 4.05289154550505E-02

 6.34091543182452E-02
 4.05289154550505E-02

 7.85190239735694
 5.93966638255407

 7.56756429483702
 5.73580192505126

 8.93455438909984E-03
 4.73238623366233E-03

 3.27534872989418E-02
 1.81617694405504E-02

 3.45502163683982E-02
 1.95555361040541E-02

 3.40527613254886E-02
 2.02671329034629E-02

 4.07258589076771E-02
 2.49406652664974E-02

 3.77902196084014E-02
 3.76370502664756E-02

 6.39262457444276E-02
 3.59142369626438E-02

 6.54775200229746E-02
 4.24871442413935E-02

 6.75786130331585E-02
 5.28507758237609E-02

 7.26611635702545E-02
 5.63017527832999E-02

 8.92702710884529E-02
 6.01540061334829E-02

 .115056115056115
 .0786688987978

 .135913226822318
 .088834567360783

 .156063428790702
 .113406058347888

 .166362711817257
 .112277134102209

 .159074340892523
 .124804983018117

 .208960572596936
 .167206521629506

 .227261681807136
 .157987865348443

 .224214769669315
 .161668907438618

 .230033684579139
 .173881085593886

 .23447150719878
 .175948996593672

 .248216975489703
 .169156724888037

 .235289689835144
 .212144127029766

 .291165018437746
 .224452611518789

 .308841036113763
 .214094330288296

 .324206506024688
 .243336678462498

 .269741724287179
 .291069842717579

 .404578222760041
 .279363272814522

 .509361236633964
 .344479730312304

 .496181950727405
 .366788022456176

 .474647383738293
 .442104925804129

 .746169473442201
 .607337167591667

 .854032126759399
 .613787016777286

 .284338102519921
 .20386445750281

 .115802297620479
 8.13173229760508E-02

 .168535804899441
 .12254713452676

 5.47657420384693
 4.59491964293143

 5.41526941526942
 4.55408843259542

 1.09047381774655E-02
 6.04322244309961E-03

 1.58171067261976E-02
 8.31444681414501E-03

 4.49313176585904E-02
 .028982856108252

 6.75066129611584E-02
 4.47717444921271E-02

 .24187115096206
 .18048341980795

 .251476615112979
 .20327859397247

 .493904130267767
 .180314883723879

 .363842545660727
 .312011119957323

 1.66855912310458
 1.12972144810805

 2.25645607463789
 2.46016669716813

 2.42902061083879E-02
 1.71880053992541E-02

 2.42902061083879E-02
 1.71880053992541E-02

 3.70145824691279E-02
 2.36432049367483E-02

 3.70145824691279E-02
 2.36432049367483E-02

 8.85960885960886E-02
 5.92524718425025E-02

 1.21581939763758E-02
 6.10475148966503E-03

 1.21581939763758E-02
 6.10475148966503E-03

 1.79901998083816E-02
 1.08933164180175E-02

 1.79901998083816E-02
 1.08933164180175E-02

 5.84476948113312E-02
 .04225440393482

 5.84476948113312E-02
 .04225440393482

 5.01742319924138E-02
 3.45873496975807E-02

 3.32378514196696E-02
 2.44965860608513E-02

 1.18701936883755E-02
 9.43267035607314E-03

 2.13676577312941E-02
 1.50639157047782E-02

 1.69363805727442E-02
 1.00907636367294E-02

 1.69363805727442E-02
 1.00907636367294E-02

 .603095512186421
 .462658302532918

 1.87036550672914E-02
 1.04599379161219E-02

 1.87036550672914E-02
 1.04599379161219E-02

 .274339910703547
 .213428211479827

 9.11979093797276E-02
 6.01005026147303E-02

 .183142001323819
 .153327708865097

 .310051946415583
 .238770153136969

 .153818335636517
 .114072177156357

 .156233610779065
 .124697975980612

 .60687224323588
 .427161393016544

 2.07392934665662E-02
 1.45716833322548E-02

 2.07392934665662E-02
 1.45716833322548E-02

 4.45647718374991E-02
 2.47052497839862E-02

 4.45647718374991E-02
 2.47052497839862E-02

 5.96716960353324E-02
 4.01570659997203E-02

 5.96716960353324E-02
 4.01570659997203E-02

 .153674335492517
 .108430231103901

 .06548079275352
 4.54940419952864E-02

 8.81935427389973E-02
 6.29361891086151E-02

 .328222146403965
 .239297162796681

 .328222146403965
 .239297162796681

 5.40262358444177E-02
 3.50341040791644E-02

 5.40262358444177E-02
 3.50341040791644E-02

 5.40262358444177E-02
 3.50341040791644E-02

 7.29607493243857
 6.49780171430089

 7.52891661982571E-02
 3.77868601189827E-02

 7.52891661982571E-02
 3.77868601189827E-02

 3.03152048606594
 3.50546494303654

 .215064215064215
 .161896297393317

 .226364953637681
 .221836289451789

 .195830377648559
 .271872780189166

 .262531898895535
 .307332237242414

 .492297219569947
 .428825352449748

 .699935609026518
 .836524840520055

 .939496212223485
 1.27717714579005

 .26749990386354
 .189207168540552

 .26749990386354
 .189207168540552

 .40546513273786
 .29478833727088

 .40546513273786
 .29478833727088

 .595083867811141
 .301559207569014

 .595083867811141
 .301559207569014

 .505201596110687
 .391699260787359

 .505201596110687
 .391699260787359

 2.41601477965114
 1.77729593697756

 2.41601477965114
 1.77729593697756

 7.04978886797069E-02
 5.35382960397311E-02

 7.04978886797069E-02
 5.35382960397311E-02

 7.04978886797069E-02
 5.35382960397311E-02

 7.22389813298904E-02
 4.59541722565583E-02

 5.48411457502367E-02
 3.51625125241705E-02

 1.35621953803772E-02
 9.18120381793619E-03

 1.35621953803772E-02
 9.18120381793619E-03

 4.12789503698595E-02
 2.59813087062344E-02

 4.12789503698595E-02
 2.59813087062344E-02

 1.73978355796538E-02
 1.07916597323877E-02

 1.73978355796538E-02
 1.07916597323877E-02

 1.73978355796538E-02
 1.07916597323877E-02

 2.69672996945724E-02
 1.71585784639402E-02

 2.69672996945724E-02
 1.71585784639402E-02

 2.69672996945724E-02
 1.71585784639402E-02

 2.69672996945724E-02
 1.71585784639402E-02

 1.30530239621149
 1.22755530732301

 1.30530239621149
 1.22755530732301

 .970295879386788
 .932857926034007

 8.48651757742667E-02
 6.33963193698869E-02

 2.97818479636661E-04
 1.87262315633897E-05

 1.13858295676478E-02
 7.45304016222909E-03

 1.25181943363762E-02
 8.60604099134637E-03

 1.37291046381955E-02
 9.56107880107924E-03

 1.37389228298319E-02
 .010524142138625

 3.31953059225786E-02
 2.72332910450438E-02

 .245700245700246
 .197414608317192

 6.1527334254607E-03
 3.78002359986709E-03

 9.97200997200997E-03
 7.79813785818298E-03

 1.48287421014694E-02
 8.5899899357206E-03

 1.68283804647441E-02
 1.35925689390833E-02

 2.04414749869295E-02
 1.40232722650412E-02

 .177476904749632
 .149630615719296

 .516273243545971
 .571957965816548

 2.36160236160236E-02
 1.19687371449436E-02

 .492657219929947
 .559989228671605

 4.25487698214971E-02
 3.94534947281244E-02

 1.59447432174705E-02
 2.00397429487646E-02

 2.66040266040266E-02
 1.94137517793598E-02

 .026649844831663
 2.19203916329164E-02

 .026649844831663
 2.19203916329164E-02

 5.42585997131452E-02
 3.87151461693393E-02

 5.42585997131452E-02
 3.87151461693393E-02

 .335006516824699
 .294697381289001

 .335006516824699
 .294697381289001

 9.98280998280998E-02
 9.94255888978487E-02

 .235178416996599
 .195271792391152

 1.41602541602542
 .656568430371818

 1.41602541602542
 .656568430371818

 1.41602541602542
 .656568430371818

 1.41054686509232E-02
 6.69061502000537E-03

 1.41054686509232E-02
 6.69061502000537E-03

 .676813767722859
 .295087956975894

 1.97378379196561E-02
 1.26642828887267E-02

 5.25993253265981E-02
 3.29956200146926E-02

 5.21116884753248E-02
 4.44239716202356E-02

 6.07189698098789E-02
 3.99136249893963E-02

 8.81542699724518E-02
 7.26604536418896E-02

 .403491676218949
 9.24300038209538E-02

 2.46011155102064E-02
 1.21907767477667E-02

 2.46011155102064E-02
 1.21907767477667E-02

 5.10578692396874E-02
 2.51118765265055E-02

 5.10578692396874E-02
 2.51118765265055E-02

 4.69964106327743E-02
 3.09919132374099E-02

 4.69964106327743E-02
 3.09919132374099E-02

 .22774931865841
 5.91133626937459E-02

 .22774931865841
 5.91133626937459E-02

 .374701465610557
 .22738192917049

 .374701465610557
 .22738192917049

 1.57418339236521E-02
 8.98324079855179E-03

 1.57418339236521E-02
 8.98324079855179E-03

 1.57418339236521E-02
 8.98324079855179E-03

 5.84836948473312E-03
 4.00473837862776E-03

 3.68836732473096E-03
 2.62167241887455E-03

 9.81819163637346E-06
 5.35035187525419E-06

 5.85818767636949E-04
 3.10320408764743E-04

 7.16727989455262E-04
 3.71849455330166E-04

 6.93818875637057E-04
 5.64462122839317E-04

 1.68218350036532E-03
 1.36969008006507E-03

 2.16000216000216E-03
 1.38306595975321E-03

 1.22400122400122E-03
 5.59111770964063E-04

 9.36000936000936E-04
 8.23954188789146E-04

 6.7581885763704E-03
 3.41352449641217E-03

 1.72800172800173E-03
 8.56056300040671E-04

 1.72800172800173E-03
 8.56056300040671E-04

 5.03018684836867E-03
 2.5574681963715E-03

 5.03018684836867E-03
 2.5574681963715E-03

 3.13527586254859E-03
 1.56497792351185E-03

 3.13527586254859E-03
 1.56497792351185E-03

 3.13527586254859E-03
 1.56497792351185E-03

 .159005613551068
 7.87170519646773E-02

 .159005613551068
 7.87170519646773E-02

 1.02501920683739E-02
 4.42741617677284E-03

 4.90909581818673E-05
 2.6751759376271E-06

 4.90909581818673E-05
 2.6751759376271E-06

 4.90909581818673E-05
 2.6751759376271E-06

 .010201101110192
 4.42474100083522E-03

 .010201101110192
 4.42474100083522E-03

 .010201101110192
 4.42474100083522E-03

 7.92818974637157E-02
 4.55395199862261E-02

 7.92818974637157E-02
 4.55395199862261E-02

 6.85997049633413E-02
 3.91752764306112E-02

 8.60728133455406E-04
 3.3172181626576E-04

 1.08327381054654E-03
 5.88538706277961E-04

 1.37781955963774E-03
 6.68793984406774E-04

 1.44000144000144E-03
 9.30961226294229E-04

 1.68218350036532E-03
 7.41023734722706E-04

 1.87200187200187E-03
 7.03571271595926E-04

 1.64618346436528E-03
 9.30961226294229E-04

 2.33672960945688E-03
 8.53381124103044E-04

 2.93891202982112E-03
 1.00586615254779E-03

 2.77527550254823E-03
 1.30013550568677E-03

 2.74254819709365E-03
 1.57300345132473E-03

 .003452730725458
 1.53287581226033E-03

 2.23200223200223E-03
 2.88651483669964E-03

 5.2527325254598E-03
 2.53339161293286E-03

 5.50146004691459E-03
 2.41568387167727E-03

 4.7094592549138E-03
 3.12460549514845E-03

 6.36873364146091E-03
 2.98549634639184E-03

 4.41818623636805E-03
 5.37710363463046E-03

 7.00364336727973E-03
 .003515181182042

 8.90509981419072E-03
 5.17646543930843E-03

 1.06821925003743E-02
 6.36424355561486E-03

 5.38364174727811E-03
 2.98282117045421E-03

 5.29855075309621E-03
 3.38142238516065E-03

 6.94735240189786E-02
 2.87501158016784E-02

 6.94735240189786E-02
 2.87501158016784E-02

 1.89131098222007E-02
 6.56220657499927E-03

 2.07818389636571E-03
 8.85483235354569E-04

 3.2661850843669E-03
 1.0272675600488E-03

 6.1756425392789E-03
 2.36485552886235E-03

 7.39309830218921E-03
 2.28460025073354E-03

 5.05604141967778E-02
 2.21879092266791E-02

 5.05604141967778E-02
 2.21879092266791E-02

 .284845375754467
 .188990479289604

 .284845375754467
 .188990479289604

 .284845375754467
 .188990479289604

 .163001617547072
 .109224758357377

 7.37018918837101E-03
 3.98066179518912E-03

 4.90909581818673E-05
 1.87262315633897E-05

 8.83637247273611E-05
 3.74524631267793E-05

 1.27636491272855E-04
 4.28028150020335E-05

 1.96363832727469E-04
 5.6178694690169E-05

 3.53454898909444E-04
 1.44459500631863E-04

 7.33091642182551E-04
 3.53123223766777E-04

 9.45819127637309E-04
 2.86243825326099E-04

 6.31636995273359E-04
 6.58093280656266E-04

 1.28618310436492E-03
 6.15290465654232E-04

 1.32218314036496E-03
 7.03571271595926E-04

 1.63636527272891E-03
 1.06472002317558E-03

 9.87513714786442E-02
 6.63764653644035E-02

 4.21200421200421E-03
 2.43708527917828E-03

 1.53949244858336E-02
 9.72426453327449E-03

 7.91444427808064E-02
 5.42151155519507E-02

 5.68800568800569E-02
 3.88676311977841E-02

 1.25345579891034E-02
 8.06565545194569E-03

 4.43454988909534E-02
 3.08019757458384E-02

 1.59251068341977E-02
 1.05375180183131E-02

 1.59251068341977E-02
 1.05375180183131E-02

 3.08618490436672E-03
 1.90472526759049E-03

 4.93200493200493E-03
 3.70511867361353E-03

 7.90691699782609E-03
 4.92767407710911E-03

 .105918651373197
 .069228202913914

 .105918651373197
 .069228202913914

 2.04938386756569E-02
 1.37343532637775E-02

 2.73927546654819E-02
 1.69820168520568E-02

 .058032058032058
 3.85118327980797E-02

 1.33298315116497E-02
 7.88106831224942E-03

 1.33298315116497E-02
 7.88106831224942E-03

 1.33298315116497E-02
 7.88106831224942E-03

 1.33298315116497E-02
 7.88106831224942E-03

 5.89091498182407E-05
 4.01276390644064E-05

 5.89091498182407E-05
 4.01276390644064E-05

 3.5378217196399E-03
 2.39428246417625E-03

 3.5378217196399E-03
 2.39428246417625E-03

 9.73310064219155E-03
 5.44665820900877E-03

 9.73310064219155E-03
 5.44665820900877E-03

 .021917476462931
 1.45048039338141E-02

 .021917476462931
 1.45048039338141E-02

 .021917476462931
 1.45048039338141E-02

 .021917476462931
 1.45048039338141E-02

 7.57964394328031E-03
 4.70563447428606E-03

 4.58182276364095E-05
 5.35035187525419E-05

 5.1054596509142E-04
 0

 4.45091354182263E-04
 2.32740306573557E-04

 6.54546109091564E-04
 2.00638195322032E-04

 7.00364336727973E-04
 4.49429557521352E-04

 5.22327795055068E-03
 3.76932289611658E-03

 3.06982125163943E-03
 1.89402456383998E-03

 3.06982125163943E-03
 1.89402456383998E-03

 1.12680112680113E-02
 7.90514489568807E-03

 1.12680112680113E-02
 7.90514489568807E-03

 .206476570112934
 .129259150954266

 .195827104918014
 .122846754231774

 .118355027445937
 7.42976613157173E-02

 .118355027445937
 7.42976613157173E-02

 4.31673158945886E-02
 2.42424443467767E-02

 1.5054560509106E-04
 1.73886435945761E-04

 2.29091138182047E-04
 1.20382917193219E-04

 5.07273234545962E-04
 1.89937491571524E-04

 7.85455330909876E-04
 4.11977094394573E-04

 8.34546289091744E-04
 3.87900510955929E-04

 9.36000936000936E-04
 4.57455085334233E-04

 1.56763793127429E-03
 9.38986754107111E-04

 1.83600183600184E-03
 1.07809590286372E-03

 2.17309308218399E-03
 1.1155483659905E-03

 2.21563857927494E-03
 1.21988022755796E-03

 2.35963872327509E-03
 1.1744022366183E-03

 2.70654816109362E-03
 1.19847882005694E-03

 2.56254801709347E-03
 1.40446736725423E-03

 4.56545911091366E-03
 2.5574681963715E-03

 6.1134606589152E-03
 3.06307644858302E-03

 6.33600633600634E-03
 4.11977094394573E-03

 7.28837092473456E-03
 4.6307295480325E-03

 6.21818803636985E-04
 2.62167241887455E-04

 6.21818803636985E-04
 2.62167241887455E-04

 9.22910013819105E-04
 3.45097695953895E-04

 9.22910013819105E-04
 3.45097695953895E-04

 3.92204028567665E-02
 2.58850023724798E-02

 2.66727539454812E-03
 2.04650959228473E-03

 4.39200439200439E-03
 2.68320146543998E-03

 5.03018684836867E-03
 3.55263364516878E-03

 5.14473241745969E-03
 3.56600952485692E-03

 7.14764351127987E-03
 4.21875245363793E-03

 7.44873472146199E-03
 4.81531668772877E-03

 7.38982557164375E-03
 5.00257900336267E-03

 6.43418825237007E-03
 4.42741617677284E-03

 6.43418825237007E-03
 4.42741617677284E-03

 9.23891832982742E-03
 5.28614765275114E-03

 9.23891832982742E-03
 5.28614765275114E-03

 9.95891904982814E-03
 5.52958866307521E-03

 9.95891904982814E-03
 5.52958866307521E-03

 8.7905542450997E-03
 8.31979716602027E-03

 8.7905542450997E-03
 8.31979716602027E-03

 3.53258535076717E-02
 2.38545438358208E-02

 3.53258535076717E-02
 2.38545438358208E-02

 9.68728241455514E-04
 3.66499103454912E-04

 9.68728241455514E-04
 3.66499103454912E-04

 1.10618292436474E-03
 6.76819512219655E-04

 1.10618292436474E-03
 6.76819512219655E-04

 1.34836498472862E-03
 5.40385539400673E-04

 1.34836498472862E-03
 5.40385539400673E-04

 1.23087395814669E-02
 7.70183152442841E-03

 1.93091102182011E-03
 9.52362633795246E-04

 1.03778285596467E-02
 6.74946889063316E-03

 .019593837775656
 1.45690081563172E-02

 9.36655482110028E-03
 6.9902347250196E-03

 1.02272829545557E-02
 7.57877343129756E-03

 2.10403846767483E-02
 .013565817179707

 2.10403846767483E-02
 .013565817179707

 2.97818479636661E-03
 1.79771823008541E-03

 9.2945547491002E-04
 5.08283428149148E-04

 2.04872932145659E-03
 1.28943480193626E-03

 1.80981999163817E-03
 8.15928660976264E-04

 1.80981999163817E-03
 8.15928660976264E-04

 1.75091084181993E-03
 1.20917952380745E-03

 1.75091084181993E-03
 1.20917952380745E-03

 2.66072993345721E-03
 1.64523320164066E-03

 2.66072993345721E-03
 1.64523320164066E-03

 1.18407391134664E-02
 8.09775756319722E-03

 1.18407391134664E-02
 8.09775756319722E-03

 .005547278274551
 3.27441534765557E-03

 .005547278274551
 3.27441534765557E-03

 .005547278274551
 3.27441534765557E-03

 .005547278274551
 3.27441534765557E-03

 1.55585610131065E-02
 7.85431655287315E-03

 1.55585610131065E-02
 7.85431655287315E-03

 1.55585610131065E-02
 7.85431655287315E-03

 1.55585610131065E-02
 7.85431655287315E-03

 1.06494651949197E-02
 6.41239672249215E-03

 1.06494651949197E-02
 6.41239672249215E-03

 1.06494651949197E-02
 6.41239672249215E-03

 1.06494651949197E-02
 6.41239672249215E-03

 1.06494651949197E-02
 6.41239672249215E-03

 56.379666197848
 68.0868926036441

 .723705450978178
 .627406337475745

 .705836342199979
 .614035808139485

 .690775236229782
 .602952554229896

 9.03600903600904E-02
 7.60820036661146E-02

 2.22545677091132E-04
 1.49809852507117E-04

 1.15560115560116E-02
 7.91317042350095E-03

 1.10683747047383E-02
 8.9645145669884E-03

 6.75131584222493E-02
 5.90545088231181E-02

 3.14182132363951E-04
 1.81911963758643E-04

 3.14182132363951E-04
 1.81911963758643E-04

 .201829292738384
 .184747650252527

 1.23709214618306E-03
 7.30323030972197E-04

 4.33833161105888E-02
 3.08126764495889E-02

 .157208884481612
 .153204650771966

 9.06546361091816E-03
 4.88487126210708E-03

 9.06546361091816E-03
 4.88487126210708E-03

 8.38800838800839E-03
 5.65799710808131E-03

 8.38800838800839E-03
 5.65799710808131E-03

 1.15658297476479E-02
 8.92438692792399E-03

 1.15658297476479E-02
 8.92438692792399E-03

 1.66189257098348E-02
 .011134082252404

 1.66189257098348E-02
 .011134082252404

 3.80880380880381E-02
 .029499165064214

 1.77218359036541E-02
 1.17547230699335E-02

 .020366202184384
 1.77444419942805E-02

 2.02123838487475E-02
 1.59627748198209E-02

 2.02123838487475E-02
 1.59627748198209E-02

 2.22218404036586E-02
 1.57407352169978E-02

 2.22218404036586E-02
 1.57407352169978E-02

 2.43425697971153E-02
 1.87877606099551E-02

 2.43425697971153E-02
 1.87877606099551E-02

 .032511305238578
 2.42156925874005E-02

 .032511305238578
 2.42156925874005E-02

 3.72109463018554E-02
 3.24391834196662E-02

 3.72109463018554E-02
 3.24391834196662E-02

 4.21331330422239E-02
 3.81453336946248E-02

 4.21331330422239E-02
 3.81453336946248E-02

 6.17433344706072E-02
 5.57239147807724E-02

 6.17433344706072E-02
 5.57239147807724E-02

 7.41698923517105E-02
 8.08250906035275E-02

 7.41698923517105E-02
 8.08250906035275E-02

 1.19389210298301E-02
 9.06884642855586E-03

 8.11637175273539E-04
 6.79494688157282E-04

 8.11637175273539E-04
 6.79494688157282E-04

 1.11272838545566E-02
 8.38935174039857E-03

 5.18073245345973E-03
 3.89505616518505E-03

 5.94655140109686E-03
 4.49429557521352E-03

 3.12218494036676E-03
 2.0144074810332E-03

 3.12218494036676E-03
 2.0144074810332E-03

 1.2338194156376E-03
 7.19622327221689E-04

 1.88836552472916E-03
 1.29478515381151E-03

 1.78691087781997E-02
 1.33705293362602E-02

 1.78691087781997E-02
 1.33705293362602E-02

 1.78691087781997E-02
 1.33705293362602E-02

 1.89818371636553E-03
 1.21988022755796E-03

 2.4283660647297E-03
 1.48204746944541E-03

 1.35425589971045E-02
 1.06686016392569E-02

 .374334919789465
 .347893254808716

 .374334919789465
 .347893254808716

 .337042155223973
 .32103448839494

 1.18701936883755E-02
 8.58731475978298E-03

 1.15854661309207E-03
 3.63823927517285E-04

 1.07116470752834E-02
 8.22349083226569E-03

 5.97175142629688E-02
 4.30730077717339E-02

 5.97175142629688E-02
 4.30730077717339E-02

 6.12000612000612E-02
 6.17938889832483E-02

 6.12000612000612E-02
 6.17938889832483E-02

 9.97331906422815E-02
 .101062796571676

 9.97331906422815E-02
 .101062796571676

 .104521195430286
 .106517480308498

 .104521195430286
 .106517480308498

 8.78400878400878E-03
 .005588442533703

 8.78400878400878E-03
 .005588442533703

 8.78400878400878E-03
 .005588442533703

 2.85087557814831E-02
 .021270323880073

 2.85087557814831E-02
 .021270323880073

 2.85087557814831E-02
 .021270323880073

 .036000036000036
 2.59545569468581E-02

 .036000036000036
 2.59545569468581E-02

 .036000036000036
 2.59545569468581E-02

 2.93498475316657E-02
 2.04918476822236E-02

 3.15163951527588E-03
 2.95339423514031E-03

 2.61982080163898E-02
 1.75384534470832E-02

 6.65018846837029E-03
 5.46270926463453E-03

 6.65018846837029E-03
 5.46270926463453E-03

 .38931093476548
 .363117681069752

 .38931093476548
 .363117681069752

 3.26029416938508E-02
 2.70620797850357E-02

 1.08491017581927E-02
 8.75050049197823E-03

 1.08491017581927E-02
 8.75050049197823E-03

 2.17538399356581E-02
 1.83115792930575E-02

 2.17538399356581E-02
 1.83115792930575E-02

 .236873691419146
 .218345184853186

 .184366002547821
 .179009397866317

 1.63636527272891E-02
 1.34695108459524E-02

 2.09651118742028E-02
 1.74153953539524E-02

 .147037237946329
 .148124491666412

 2.24542042723861E-02
 1.65941163411009E-02

 2.24542042723861E-02
 1.65941163411009E-02

 3.00534845989391E-02
 2.27416706457679E-02

 3.00534845989391E-02
 2.27416706457679E-02

 .119834301652483
 .11771041643153

 .119834301652483
 .11771041643153

 .119834301652483
 .11771041643153

 54.8563148563149
 66.7225207733431

 54.8563148563149
 66.7225207733431

 4.10832738105465
 3.8560199979032

 .327495600222873
 .245145097396334

 1.25149216058307E-02
 8.06833062788332E-03

 2.05658387476569E-02
 1.41891331731741E-02

 2.31316594952959E-02
 1.52217510850982E-02

 .271283180374089
 .207665882510179

 .241452241452241
 .208077859604573

 .241452241452241
 .208077859604573

 3.53937953937954
 3.40279704090229

 .424446969901515
 .315317637416231

 .854271036089218
 .593656317846642

 .793358975177157
 .859943330678043

 1.46730255821165
 1.63387975496137

 3.26683963047599E-02
 2.72707435081706E-02

 3.26683963047599E-02
 2.72707435081706E-02

 3.26683963047599E-02
 2.72707435081706E-02

 6.08915408915409
 3.85666204012823

 6.08915408915409
 3.85666204012823

 3.42851251942161E-02
 3.82389648524417E-02

 .037665855847674
 3.61496524451549E-02

 4.18942237124055E-02
 3.72063469405177E-02

 5.46284182647819E-02
 3.90629190412309E-02

 4.65775011229557E-02
 5.19947195237202E-02

 5.37873265145992E-02
 4.73586396238125E-02

 5.63400563400563E-02
 5.27838964253202E-02

 6.65607938335211E-02
 4.74469204297542E-02

 6.88222506404325E-02
 6.59002840475059E-02

 7.85357148993513E-02
 8.08090395479017E-02

 8.52415397869943E-02
 .078532464824981

 .10715574351938
 .082521152147983

 .057145148054239
 .127595191521062

 .135016498652862
 .088181824432002

 .134090315908498
 .114789124307641

 .209850755305301
 .146350850019766

 .904883813974723
 .617904112545294

 1.02249593158684
 .716117846743397

 2.89417707599526
 1.38771809070874

 43.0404877677605
 57.896829111286

 43.0404877677605
 57.896829111286

 3.53945808491263E-02
 8.72000348628928E-02

 9.36295481750027E-02
 .204182803439388

 .258460622096986
 .240297678597354

 .32386614204796
 .237809764975361

 .281242099423918
 .276746950747523

 .154891791255428
 .382614363303178

 .336079972443609
 .242285334319011

 .233702415520597
 .458008846753322

 .356315265406174
 .377954206819831

 .289721744267199
 .603747081483371

 .462080098443735
 .531134781008359

 .373696737333101
 .619241700514107

 .564490382672201
 .509580888478897

 .918236554600191
 .250340289067206

 .568074022619477
 .683796371064987

 .378255650982924
 .920089311283713

 .714528714528715
 .664730392157518

 .779152051879325
 .614843711272648

 .838493202129566
 .888441979941584

 .883542338087793
 .858819756784239

 .936498391043846
 .879571096532413

 .973876246603519
 1.01627793729703

 1.43171815899089
 .806699303991451

 1.0724574360938
 1.13611779377491

 .868553232189596
 1.40270175113539

 .655118836937019
 1.96110460047598

 .818333181969546
 2.00302728259454

 1.4318850682487
 1.60035980046291

 2.32866887412342
 2.38151384942596

 1.3079958534504
 3.78394005743977

 2.66578630214994
 2.92293468191449

 2.64697791970519
 3.8642461639114

 3.04995250449796
 4.57225822756379

 3.34170661443389
 4.80595624712302

 3.87480169298351
 4.9947032854023

 6.79230351957625
 10.1135507853661

 1.58567722204086
 1.08573888051752

 1.58567722204086
 1.08573888051752

 1.58567722204086
 1.08573888051752

 .122930304748487
 8.09240721132197E-02

 4.49149540058631E-02
 3.05585347355143E-02

 4.49149540058631E-02
 3.05585347355143E-02

 4.49149540058631E-02
 3.05585347355143E-02

 3.43047615774889E-02
 2.50824495911917E-02

 6.905461450916E-04
 3.04970056889489E-04

 2.17309308218399E-03
 7.81151373787112E-04

 3.27273054545782E-03
 2.08128687947388E-03

 4.19891328982238E-03
 2.39428246417625E-03

 2.39694785149331E-02
 1.95207588168649E-02

 1.06101924283742E-02
 5.47608514432267E-03

 1.06101924283742E-02
 5.47608514432267E-03

 5.75411484502394E-02
 3.81533592224376E-02

 4.75298657116839E-02
 3.15670760639997E-02

 4.60800460800461E-02
 3.06789176527075E-02

 1.51560151560152E-02
 1.03368798229911E-02

 1.06691015781925E-03
 8.40005244414908E-04

 6.60437024073388E-03
 4.05289154550505E-03

 7.48473475746203E-03
 5.44398303307114E-03

 3.09240309240309E-02
 2.03420378297164E-02

 1.49531058621968E-02
 9.85267297828059E-03

 1.59709250618342E-02
 1.04893648514358E-02

 1.44981963163781E-03
 8.88158411292196E-04

 1.44981963163781E-03
 8.88158411292196E-04

 1.44981963163781E-03
 8.88158411292196E-04

 1.00112827385555E-02
 6.58628315843791E-03

 1.00112827385555E-02
 6.58628315843791E-03

 1.00112827385555E-02
 6.58628315843791E-03

 1.00112827385555E-02
 6.58628315843791E-03

 4.68327741055014E-03
 2.44778598292879E-03

 4.68327741055014E-03
 2.44778598292879E-03

 4.68327741055014E-03
 2.44778598292879E-03

 4.68327741055014E-03
 2.44778598292879E-03

 4.68327741055014E-03
 2.44778598292879E-03

 .015790924881834
 9.7643921723389E-03

 5.41964178327815E-03
 3.14600690264946E-03

 5.41964178327815E-03
 3.14600690264946E-03

 5.41964178327815E-03
 3.14600690264946E-03

 5.41964178327815E-03
 3.14600690264946E-03

 1.03712830985558E-02
 6.61838526968944E-03

 1.03712830985558E-02
 6.61838526968944E-03

 1.03712830985558E-02
 6.61838526968944E-03

 1.03712830985558E-02
 6.61838526968944E-03

 9.09819091637273E-04
 6.39367049092876E-04

 9.09819091637273E-04
 6.39367049092876E-04

 9.09819091637273E-04
 6.39367049092876E-04

 9.09819091637273E-04
 6.39367049092876E-04

 9.09819091637273E-04
 6.39367049092876E-04

 9.09819091637273E-04
 6.39367049092876E-04

 .333988697625061
 .241769025363049

 .333988697625061
 .241769025363049

 .333988697625061
 .241769025363049

 .333988697625061
 .241769025363049

 .175333266242357
 .130874957220593

 1.04400104400104E-03
 7.38348558785078E-04

 1.4661832843651E-03
 1.03529308786169E-03

 1.81963818327455E-03
 1.3429383206888E-03

 2.08800208800209E-03
 1.28408445006101E-03

 2.75563911927548E-03
 1.85122174883795E-03

 2.8865483410938E-03
 1.82446998946168E-03

 2.74582092763911E-03
 2.53339161293286E-03

 3.35127607854881E-03
 2.38625693636337E-03

 3.92727665454938E-03
 2.69122699325286E-03

 3.21382139563958E-03
 3.32524369047048E-03

 6.12655158109704E-03
 3.36804650547251E-03

 5.38691447782357E-03
 4.36321195426979E-03

 1.02960102960103E-02
 6.48462647280808E-03

 1.29272856545584E-02
 7.61087554254909E-03

 1.40236503872868E-02
 .010072037405166

 .101274646729192
 7.99636839516115E-02

 4.50556814193178E-02
 3.01010796501801E-02

 3.6000036000036E-03
 2.19631944479185E-03

 5.71091480182389E-03
 3.53390741360539E-03

 3.57447630174903E-02
 2.43708527917828E-02

 1.97051106142015E-02
 1.25412247955958E-02

 1.97051106142015E-02
 1.25412247955958E-02

 3.63698545516727E-02
 2.66340516350154E-02

 3.63698545516727E-02
 2.66340516350154E-02

 5.75247847975121E-02
 4.16177120616647E-02

 5.75247847975121E-02
 4.16177120616647E-02

 .118872118872119
 .079048773780943

 .118872118872119
 .079048773780943

 .118872118872119
 .079048773780943

 .115789206698298
 7.68631550399017E-02

 5.14931424022333E-02
 3.41325697881841E-02

 1.16509207418298E-03
 7.30323030972197E-04

 5.24618706436888E-03
 3.08180268014641E-03

 7.38328011055284E-03
 5.02398041086369E-03

 8.19491728582638E-03
 5.81583248840131E-03

 1.02272829545557E-02
 7.01163613252062E-03

 1.92763829127466E-02
 1.24689950452799E-02

 .010315646679283
 6.08870043403927E-03

 .010315646679283
 6.08870043403927E-03

 2.35603871967508E-02
 1.56925820501205E-02

 9.85746440291895E-03
 6.79762205751045E-03

 1.37029227938319E-02
 8.89495999261009E-03

 1.39516503152867E-02
 1.01950954982969E-02

 1.39516503152867E-02
 1.01950954982969E-02

 1.64683801047437E-02
 1.07542072692609E-02

 1.64683801047437E-02
 1.07542072692609E-02

 3.08291217382126E-03
 2.18561874104134E-03

 3.08291217382126E-03
 2.18561874104134E-03

 3.08291217382126E-03
 2.18561874104134E-03

 4.83578665396847E-02
 3.22331948724689E-02

 4.83578665396847E-02
 3.22331948724689E-02

 2.48923885287522E-02
 1.65539887020365E-02

 2.48923885287522E-02
 1.65539887020365E-02

 1.04269195178286E-02
 6.97150849345621E-03

 1.22072849345577E-03
 7.16947151284062E-04

 9.20619102437284E-03
 6.25456134217215E-03

 4.10073137345865E-03
 2.73135463231727E-03

 4.10073137345865E-03
 2.73135463231727E-03

 1.03647376374649E-02
 6.85112557626299E-03

 5.01382319564138E-03
 3.42155002422506E-03

 5.35091444182353E-03
 3.42957555203794E-03

 2.34654780109326E-02
 1.56792061704324E-02

 1.79934725389271E-02
 .012503772332469

 1.79934725389271E-02
 .012503772332469

 2.65745720291175E-03
 1.75491541508337E-03

 7.25237088873453E-03
 4.70830965022369E-03

 8.08364444728081E-03
 6.04054726716198E-03

 5.47200547200547E-03
 3.17543383796336E-03

 5.47200547200547E-03
 3.17543383796336E-03

 5.47200547200547E-03
 3.17543383796336E-03

 6.70811579902489E-02
 5.37870874019304E-02

 1.38436502072866E-02
 1.00800629329789E-02

 1.38436502072866E-02
 1.00800629329789E-02

 1.38436502072866E-02
 1.00800629329789E-02

 8.84946339491794E-03
 5.85060977559046E-03

 1.35818317636499E-03
 6.6344363253152E-04

 1.6134561589107E-03
 9.9516544879728E-04

 5.87782405964224E-03
 4.19200069426166E-03

 4.99418681236863E-03
 4.22945315738844E-03

 4.99418681236863E-03
 4.22945315738844E-03

 1.71163807527444E-02
 1.45743585081924E-02

 1.71163807527444E-02
 1.45743585081924E-02

 1.71163807527444E-02
 1.45743585081924E-02

 1.71163807527444E-02
 1.45743585081924E-02

 1.71163807527444E-02
 1.45743585081924E-02

 3.61211270302179E-02
 2.91326659607591E-02

 3.61211270302179E-02
 2.91326659607591E-02

 3.61211270302179E-02
 2.91326659607591E-02

 3.61211270302179E-02
 2.91326659607591E-02

 3.61211270302179E-02
 2.91326659607591E-02

 .04453531726259
 3.12701315349231E-02

 .04453531726259
 3.12701315349231E-02

 .04453531726259
 3.12701315349231E-02

 .04453531726259
 3.12701315349231E-02

 1.00669191578282E-02
 6.7628447703213E-03

 1.24036487672851E-03
 8.02552781288129E-04

 2.1567294294567E-03
 1.31886173725016E-03

 6.66982485164303E-03
 4.64143025178301E-03

 1.62327435054708E-03
 1.17172706068067E-03

 1.62327435054708E-03
 1.17172706068067E-03

 4.07454952909498E-03
 2.69390216919049E-03

 4.07454952909498E-03
 2.69390216919049E-03

 9.19964556328193E-03
 5.89876294246775E-03

 9.19964556328193E-03
 5.89876294246775E-03

 1.95709286618378E-02
 1.47428945922629E-02

 1.95709286618378E-02
 1.47428945922629E-02

 4.16127688854962E-02
 2.42504698745896E-02

 4.16127688854962E-02
 2.42504698745896E-02

 4.16127688854962E-02
 2.42504698745896E-02

 1.71294716749262E-02
 1.10377759186494E-02

 1.21352848625576E-02
 8.08438168350908E-03

 1.58727431454704E-03
 7.24972679096943E-04

 3.67854913309459E-03
 1.91007561946575E-03

 6.86946141491596E-03
 5.44933338494639E-03

 4.99418681236863E-03
 2.95339423514031E-03

 4.99418681236863E-03
 2.95339423514031E-03

 2.44832972105699E-02
 1.32126939559402E-02

 2.50363886727523E-03
 1.20115399599457E-03

 2.50363886727523E-03
 1.20115399599457E-03

 2.92582110763929E-03
 9.92490272859653E-04

 2.92582110763929E-03
 9.92490272859653E-04

 3.23018504836687E-03
 1.58905450695049E-03

 3.23018504836687E-03
 1.58905450695049E-03

 4.32982251164069E-03
 3.02562398545625E-03

 4.32982251164069E-03
 3.02562398545625E-03

 1.14938296756479E-02
 6.40437119467927E-03

 1.14938296756479E-02
 6.40437119467927E-03

 .126180126180126
 9.65979279317768E-02

 .126180126180126
 9.65979279317768E-02

 .126180126180126
 9.65979279317768E-02

 .126180126180126
 9.65979279317768E-02

 .06269242632879
 4.99294836998721E-02

 2.23200223200223E-03
 1.39911701537897E-03

 6.28036991673355E-03
 .004304358083642

 8.26037189673553E-03
 6.29736415717418E-03

 1.97705652251107E-02
 1.63025221638995E-02

 .026149117058208
 2.16261222797774E-02

 1.72931082021991E-02
 1.30521833996826E-02

 5.30182348364167E-03
 4.55047426990369E-03

 1.19912847185574E-02
 8.50170912977891E-03

 6.87273414546142E-03
 5.00792935523792E-03

 6.87273414546142E-03
 5.00792935523792E-03

 2.49545704091159E-02
 1.93308213252934E-02

 1.07411016501926E-02
 9.33368884638094E-03

 1.42134687589233E-02
 9.99713247891246E-03

 1.43672870945598E-02
 9.27751015169077E-03

 1.43672870945598E-02
 9.27751015169077E-03

 .357143266234175
 .179873479694171

 .297573024845752
 .140765082662

 .297573024845752
 .140765082662

 .287944651581015
 .136083524771153

 3.08618490436672E-03
 1.74688988727049E-03

 3.08618490436672E-03
 1.74688988727049E-03

 .284858466676648
 .134336634883882

 .284858466676648
 .134336634883882

 4.18254963709509E-03
 2.52269090918235E-03

 4.18254963709509E-03
 2.52269090918235E-03

 4.18254963709509E-03
 2.52269090918235E-03

 5.44582362764181E-03
 2.15886698166507E-03

 5.44582362764181E-03
 2.15886698166507E-03

 5.44582362764181E-03
 2.15886698166507E-03

 4.20218602036784E-03
 2.40498316792676E-03

 4.20218602036784E-03
 2.40498316792676E-03

 4.20218602036784E-03
 2.40498316792676E-03

 4.20218602036784E-03
 2.40498316792676E-03

 4.20218602036784E-03
 2.40498316792676E-03

 4.89993217265945E-02
 .032920715088439

 .038402220220402
 2.91085893773204E-02

 .038402220220402
 2.91085893773204E-02

 6.37200637200637E-03
 2.61364689106167E-03

 6.37200637200637E-03
 2.61364689106167E-03

 3.20302138483957E-02
 2.64949424862588E-02

 3.20302138483957E-02
 2.64949424862588E-02

 1.05971015061924E-02
 3.81212571111861E-03

 1.05971015061924E-02
 3.81212571111861E-03

 1.05971015061924E-02
 3.81212571111861E-03

 1.05971015061924E-02
 3.81212571111861E-03

 6.36873364146091E-03
 3.78269877580471E-03

 6.36873364146091E-03
 3.78269877580471E-03

 6.36873364146091E-03
 3.78269877580471E-03

 6.36873364146091E-03
 3.78269877580471E-03

 6.36873364146091E-03
 3.78269877580471E-03

 9.24611833702743E-02
 5.95199894362653E-02

 9.24611833702743E-02
 5.95199894362653E-02

 9.24611833702743E-02
 5.95199894362653E-02

 9.24611833702743E-02
 5.95199894362653E-02

 3.74400374400374E-03
 1.71478777601897E-03

 3.74400374400374E-03
 1.71478777601897E-03

 4.6472773745501E-03
 2.57619442793489E-03

 4.6472773745501E-03
 2.57619442793489E-03

 6.40146094691549E-03
 4.06359224925556E-03

 6.40146094691549E-03
 4.06359224925556E-03

 1.29927402654675E-02
 8.49368360196603E-03

 1.29927402654675E-02
 8.49368360196603E-03

 1.39974685429231E-02
 8.30909646226976E-03

 1.39974685429231E-02
 8.30909646226976E-03

 2.08211117302026E-02
 .014312191266305

 2.08211117302026E-02
 .014312191266305

 2.98571207662117E-02
 2.00504436525151E-02

 2.98571207662117E-02
 2.00504436525151E-02

 2.61360261360261E-02
 1.52030248535348E-02

 5.59309650218741E-03
 3.58206058048268E-03

 5.59309650218741E-03
 3.58206058048268E-03

 5.59309650218741E-03
 3.58206058048268E-03

 5.59309650218741E-03
 3.58206058048268E-03

 5.59309650218741E-03
 3.58206058048268E-03

 2.05429296338387E-02
 1.16209642730521E-02

 2.05429296338387E-02
 1.16209642730521E-02

 2.05429296338387E-02
 1.16209642730521E-02

 2.05429296338387E-02
 1.16209642730521E-02

 2.05429296338387E-02
 1.16209642730521E-02

 8.43513570786298E-02
 .047433544550066

 8.43513570786298E-02
 .047433544550066

 5.59309650218741E-03
 3.76397254424132E-03

 5.59309650218741E-03
 3.76397254424132E-03

 5.59309650218741E-03
 3.76397254424132E-03

 5.59309650218741E-03
 3.76397254424132E-03

 7.87582605764424E-02
 4.36695720058247E-02

 7.87582605764424E-02
 4.36695720058247E-02

 7.87582605764424E-02
 4.36695720058247E-02

 7.87582605764424E-02
 4.36695720058247E-02

 5.16436880073244E-03
 4.2053765739498E-03

 5.16436880073244E-03
 4.2053765739498E-03

 5.16436880073244E-03
 4.2053765739498E-03

 5.16436880073244E-03
 4.2053765739498E-03

 5.16436880073244E-03
 4.2053765739498E-03

 5.16436880073244E-03
 4.2053765739498E-03

 6.76800676800677E-03
 3.81480088705624E-03

 6.76800676800677E-03
 3.81480088705624E-03

 6.76800676800677E-03
 3.81480088705624E-03

 6.76800676800677E-03
 3.81480088705624E-03

 6.76800676800677E-03
 3.81480088705624E-03

 6.76800676800677E-03
 3.81480088705624E-03

 3.61996725633089E-02
 2.48550596364933E-02

 3.61996725633089E-02
 2.48550596364933E-02

 3.61996725633089E-02
 2.48550596364933E-02

 3.61996725633089E-02
 2.48550596364933E-02

 7.02327975055248E-03
 5.37977881056809E-03

 7.02327975055248E-03
 5.37977881056809E-03

 1.17654663109209E-02
 5.89073741465487E-03

 1.17654663109209E-02
 5.89073741465487E-03

 1.74109265018356E-02
 1.35845434112704E-02

 1.74109265018356E-02
 1.35845434112704E-02

 2.95527568254841E-02
 1.98605061609436E-02

 2.95527568254841E-02
 1.98605061609436E-02

 2.95527568254841E-02
 1.98605061609436E-02

 2.95527568254841E-02
 1.98605061609436E-02

 2.95527568254841E-02
 1.98605061609436E-02

 .013382195200377
 8.88158411292196E-03

 1.61705616251071E-02
 1.09789220480216E-02

 9.80411889502799E-02
 8.13012719204251E-02

 9.80411889502799E-02
 8.13012719204251E-02

 9.80411889502799E-02
 8.13012719204251E-02

 9.80411889502799E-02
 8.13012719204251E-02

 9.80411889502799E-02
 8.13012719204251E-02

 9.80411889502799E-02
 8.13012719204251E-02

 1.36475154656973
 .191390112105655

 1.21091030181939E-04
 4.01276390644064E-05

 1.21091030181939E-04
 4.01276390644064E-05

 1.21091030181939E-04
 4.01276390644064E-05

 1.21091030181939E-04
 4.01276390644064E-05

 1.21091030181939E-04
 4.01276390644064E-05

 1.21091030181939E-04
 4.01276390644064E-05

 1.30680130680131E-02
 9.05814572480535E-03

 1.30680130680131E-02
 9.05814572480535E-03

 4.70618652436834E-03
 3.46167766328946E-03

 2.9945484490939E-03
 2.27924989885829E-03

 2.38909329818421E-04
 6.68793984406774E-05

 2.38909329818421E-04
 6.68793984406774E-05

 2.32363868727505E-04
 1.95287843446778E-04

 2.32363868727505E-04
 1.95287843446778E-04

 1.88836552472916E-03
 1.3723652560027E-03

 4.02545857091312E-04
 2.86243825326099E-04

 4.41818623636805E-04
 3.23696288452879E-04

 4.05818587636769E-04
 4.3337850189559E-04

 6.38182456364275E-04
 3.29046640328133E-04

 6.34909725818817E-04
 6.4471740096813E-04

 6.34909725818817E-04
 6.4471740096813E-04

 1.71163807527444E-03
 1.18242776443118E-03

 1.71163807527444E-03
 1.18242776443118E-03

 1.71163807527444E-03
 1.18242776443118E-03

 4.27418609236791E-03
 3.06307644858302E-03

 3.37418519236701E-03
 2.45313633480405E-03

 8.47637211273575E-04
 5.29684835650165E-04

 2.38909329818421E-04
 8.293045406644E-05

 6.08727881455154E-04
 4.46754381583725E-04

 2.68363904727541E-04
 1.49809852507117E-04

 2.68363904727541E-04
 1.49809852507117E-04

 4.54909545818637E-04
 4.06626742519319E-04

 4.54909545818637E-04
 4.06626742519319E-04

 4.90909581818673E-04
 3.77199807205421E-04

 4.90909581818673E-04
 3.77199807205421E-04

 5.98909689818781E-04
 4.76181316897623E-04

 5.98909689818781E-04
 4.76181316897623E-04

 7.13455258909804E-04
 5.13633780024402E-04

 7.13455258909804E-04
 5.13633780024402E-04

 9.000009000009E-04
 6.09940113778978E-04

 9.000009000009E-04
 6.09940113778978E-04

 9.000009000009E-04
 6.09940113778978E-04

 2.8472755745483E-04
 1.04331861567457E-04

 2.8472755745483E-04
 1.04331861567457E-04

 2.8472755745483E-04
 1.04331861567457E-04

 2.8472755745483E-04
 1.04331861567457E-04

 3.80291289382198E-03
 2.4290597513654E-03

 3.80291289382198E-03
 2.4290597513654E-03

 5.4000054000054E-04
 3.66499103454912E-04

 5.4000054000054E-04
 3.66499103454912E-04

 3.26291235382144E-03
 2.06256064791049E-03

 5.53091462182371E-04
 4.146522703322E-04

 6.70909761818853E-04
 4.06626742519319E-04

 8.73819055637237E-04
 5.77838002527453E-04

 1.16509207418298E-03
 6.6344363253152E-04

 1.34850571214208
 .179894881101672

 1.98392925665653E-02
 1.20463172471348E-02

 1.98392925665653E-02
 1.20463172471348E-02

 1.98392925665653E-02
 1.20463172471348E-02

 9.74946429491884E-03
 6.01647068372334E-03

 2.74909365818457E-04
 1.25733269068473E-04

 1.19127391854665E-03
 7.16947151284062E-04

 1.19127391854665E-03
 8.13253485038637E-04

 1.45636509272873E-03
 8.90833587229823E-04

 1.62327435054708E-03
 1.07274555098847E-03

 4.01236764873129E-03
 2.39695764011388E-03

 1.00898282716465E-02
 6.02984656341147E-03

 2.91273018545746E-03
 1.84854657290032E-03

 3.58036721673085E-03
 1.97963019384405E-03

 3.59673086945814E-03
 2.2016697966671E-03

 .011752375388739
 7.95062288662773E-03

 .011752375388739
 7.95062288662773E-03

 .011752375388739
 7.95062288662773E-03

 4.25454970909516E-04
 2.62167241887455E-04

 4.25454970909516E-04
 2.62167241887455E-04

 5.07273234545962E-04
 3.69174279392539E-04

 5.07273234545962E-04
 3.69174279392539E-04

 6.38182456364275E-04
 4.81531668772877E-04

 6.38182456364275E-04
 4.81531668772877E-04

 6.25091534182443E-04
 5.00257900336267E-04

 6.25091534182443E-04
 5.00257900336267E-04

 7.62546217091672E-04
 4.06626742519319E-04

 7.62546217091672E-04
 4.06626742519319E-04

 7.56000756000756E-04
 5.1630895596203E-04

 7.56000756000756E-04
 5.1630895596203E-04

 8.80364516728153E-04
 5.85863530340334E-04

 8.80364516728153E-04
 5.85863530340334E-04

 1.00800100800101E-03
 8.15928660976264E-04

 1.00800100800101E-03
 8.15928660976264E-04

 1.16181934363753E-03
 7.57074790348468E-04

 1.16181934363753E-03
 7.57074790348468E-04

 1.31236494872859E-03
 7.35673382847451E-04

 1.31236494872859E-03
 7.35673382847451E-04

 1.1945466490921E-03
 9.63063337545755E-04

 1.1945466490921E-03
 9.63063337545755E-04

 2.48072975345703E-03
 1.55695239569897E-03

 2.48072975345703E-03
 1.55695239569897E-03

 5.83331492422401E-02
 .02436282726397

 5.83331492422401E-02
 .02436282726397

 .026806935897845
 1.84292870343131E-02

 3.43636707273071E-04
 4.20002622207454E-04

 3.43636707273071E-04
 4.20002622207454E-04

 2.64632991905719E-02
 1.80092844121056E-02

 1.40072867345595E-03
 7.43698910660333E-04

 2.8865483410938E-03
 1.95822878634303E-03

 6.86946141491596E-03
 4.9169733733586E-03

 1.53065607611062E-02
 1.03903833417436E-02

 3.15262133443952E-02
 5.9335402296569E-03

 3.15262133443952E-02
 5.9335402296569E-03

 6.41455186909732E-04
 2.24714778760676E-04

 5.95636959273323E-04
 5.29684835650165E-04

 8.31273558546286E-04
 4.11977094394573E-04

 1.33200133200133E-03
 1.20650434786982E-03

 2.81258463076645E-02
 3.56065917298166E-03

 3.86182204364023E-03
 2.51466538136947E-03

 3.86182204364023E-03
 2.51466538136947E-03

 5.56364192727829E-04
 3.10320408764743E-04

 5.56364192727829E-04
 3.10320408764743E-04

 5.56364192727829E-04
 3.10320408764743E-04

 2.22545677091132E-03
 1.3723652560027E-03

 2.22545677091132E-03
 1.3723652560027E-03

 1.07672834945562E-03
 7.03571271595926E-04

 1.14872842145569E-03
 6.68793984406774E-04

 1.08000108000108E-03
 8.31979716602027E-04

 1.08000108000108E-03
 8.31979716602027E-04

 1.08000108000108E-03
 8.31979716602027E-04

 8.03455348909894E-03
 4.26958079645284E-03

 8.03455348909894E-03
 4.26958079645284E-03

 8.03455348909894E-03
 4.26958079645284E-03

 1.05709196618288E-03
 5.6178694690169E-04

 1.05709196618288E-03
 5.6178694690169E-04

 6.97746152291607E-03
 3.70779384955115E-03

 1.34509225418316E-03
 9.01534290980331E-04

 5.6323692687329E-03
 2.80625955857082E-03

 .124854670309216
 8.59774794593972E-02

 6.21655167109713E-02
 4.12057349672702E-02

 5.69749660658752E-02
 3.73748830245882E-02

 1.53163789527426E-03
 1.18242776443118E-03

 1.53163789527426E-03
 1.18242776443118E-03

 2.89636653273017E-03
 1.92345149915388E-03

 2.89636653273017E-03
 1.92345149915388E-03

 8.81673608946336E-03
 5.8746863590291E-03

 8.81673608946336E-03
 5.8746863590291E-03

 4.37302255484074E-02
 .028394317401974

 1.05218287036469E-02
 6.87787733563926E-03

 1.53458335276517E-02
 9.51025045826433E-03

 1.78625633171088E-02
 1.20061896080704E-02

 5.1905506450961E-03
 .003830851942682

 5.1905506450961E-03
 .003830851942682

 5.1905506450961E-03
 .003830851942682

 2.95200295200295E-03
 1.6586090813288E-03

 2.95200295200295E-03
 1.6586090813288E-03

 2.95200295200295E-03
 1.6586090813288E-03

 2.95200295200295E-03
 1.6586090813288E-03

 5.97371506462415E-02
 4.31131354107983E-02

 8.4960084960085E-03
 5.50818725557419E-03

 4.0810949901859E-03
 2.65912488200133E-03

 4.0810949901859E-03
 2.65912488200133E-03

 4.4149135058226E-03
 2.84906237357286E-03

 4.4149135058226E-03
 2.84906237357286E-03

 1.77021995203813E-02
 1.49248065560216E-02

 7.30800730800731E-03
 5.43595750525826E-03

 7.30800730800731E-03
 5.43595750525826E-03

 .010394192212374
 9.48884905076331E-03

 .010394192212374
 9.48884905076331E-03

 8.78728151455424E-03
 5.73290203433487E-03

 8.78728151455424E-03
 5.73290203433487E-03

 8.78728151455424E-03
 5.73290203433487E-03

 2.47516611152975E-02
 1.69472395648677E-02

 2.47516611152975E-02
 1.69472395648677E-02

 2.47516611152975E-02
 1.69472395648677E-02

 1.11051601960693
 3.58446823882655E-02

 1.11051601960693
 3.58446823882655E-02

 2.41527514254787E-03
 7.43698910660333E-04

 2.41527514254787E-03
 7.43698910660333E-04

 2.41527514254787E-03
 7.43698910660333E-04

 1.10810074446438
 3.51009834776051E-02

 .114895751259388
 3.63823927517285E-03

 .016834925925835
 8.40005244414908E-04

 9.80608253335526E-02
 2.79823403075794E-03

 1.94465649011104E-02
 6.21443370310774E-03

 1.94465649011104E-02
 6.21443370310774E-03

 .973758428303883
 2.52483104993245E-02

 1.75156538792902E-02
 8.74247496416535E-03

 .956242774424593
 1.65058355351592E-02

 2.96509387418478E-03
 9.47012281919992E-04

 2.96509387418478E-03
 9.47012281919992E-04

 2.96509387418478E-03
 9.47012281919992E-04

 2.96509387418478E-03
 9.47012281919992E-04

 2.96509387418478E-03
 9.47012281919992E-04

 8.34873562146289E-03
 5.98169339653419E-03

 8.34873562146289E-03
 5.98169339653419E-03

 8.34873562146289E-03
 5.98169339653419E-03

 2.41527514254787E-03
 1.73886435945761E-03

 2.41527514254787E-03
 1.73886435945761E-03

 5.93346047891502E-03
 4.24282903707657E-03

 5.93346047891502E-03
 4.24282903707657E-03

 2.01600201600202E-03
 1.66395943320405E-03

 2.01600201600202E-03
 1.66395943320405E-03

 7.98546253091708E-04
 5.29684835650165E-04

 7.98546253091708E-04
 5.29684835650165E-04

 7.98546253091708E-04
 5.29684835650165E-04

 7.98546253091708E-04
 5.29684835650165E-04

 1.21745576291031E-03
 1.13427459755389E-03

 1.21745576291031E-03
 1.13427459755389E-03

 1.21745576291031E-03
 1.13427459755389E-03

 1.21745576291031E-03
 1.13427459755389E-03

 1.04072831345559E-03
 7.32998206909824E-04

 1.04072831345559E-03
 7.32998206909824E-04

 1.04072831345559E-03
 7.32998206909824E-04

 1.04072831345559E-03
 7.32998206909824E-04

 1.04072831345559E-03
 7.32998206909824E-04

 1.04072831345559E-03
 7.32998206909824E-04
