## Supplementary Item 5 for "Methane emission of humans is explained by dietary habits, host genetics, local formate availability and a uniform archaeome"

Javascript must be enabled to view this page.

magnitude

HE
LE

 100
 100

 100
 100

 8.87275273556559E-03
 2.09664118082831E-02

 8.87275273556559E-03
 2.09664118082831E-02

 8.87275273556559E-03
 2.09664118082831E-02

 8.87275273556559E-03
 2.09664118082831E-02

 8.87275273556559E-03
 2.09664118082831E-02

 8.87275273556559E-03
 2.09664118082831E-02

 .957537883057119
 4.73281802552311

 .957537883057119
 4.73281802552311

 .344838336047117
 1.80870245866122

 .219420777109257
 1.19089219071048

 1.75057013431429E-02
 3.49440196804719E-02

 1.75057013431429E-02
 3.49440196804719E-02

 1.70260930871664E-02
 .102036537466978

 1.70260930871664E-02
 .102036537466978

 .138366981849226
 .717051283843283

 2.94959077425559E-02
 .14956040423242

 .032373557278415
 .169129055253484

 2.97357118705441E-02
 .226437247529458

 4.67618049577105E-02
 .171924576827922

 4.65220008297223E-02
 .336860349719749

 4.65220008297223E-02
 .336860349719749

 .12541755893786
 .617810267950743

 .12541755893786
 .617810267950743

 .12541755893786
 .617810267950743

 .313184191152666
 1.60043610136561

 .247238055955895
 1.28174664187971

 6.21092691489591E-02
 .276756635869337

 1.75057013431429E-02
 4.33305844037851E-02

 4.46035678058162E-02
 .233426051465552

 1.96639384950373E-02
 .078274604084257

 1.96639384950373E-02
 .078274604084257

 .033332773790368
 .212459639657269

 .033332773790368
 .212459639657269

 3.59706191982389E-02
 .197084270997861

 3.59706191982389E-02
 .197084270997861

 4.38841554218514E-02
 .24880142012496

 4.38841554218514E-02
 .24880142012496

 5.22772999014405E-02
 .268370071146024

 5.22772999014405E-02
 .268370071146024

 6.59461351967713E-02
 .318689459485904

 6.59461351967713E-02
 .318689459485904

 6.59461351967713E-02
 .318689459485904

 2.08629591349785E-02
 5.45126707015361E-02

 2.08629591349785E-02
 5.45126707015361E-02

 2.08629591349785E-02
 5.45126707015361E-02

 2.08629591349785E-02
 5.45126707015361E-02

 .278652396722357
 1.26916679479474

 .278652396722357
 1.26916679479474

 3.95676811180628E-02
 .191493227848986

 3.95676811180628E-02
 .191493227848986

 .239084715604294
 1.07767356694575

 4.05268976300158E-02
 .216652922018926

 4.91598462375931E-02
 .212459639657269

 6.40277021728652E-02
 .301916330039277

 8.53702695638203E-02
 .346644675230281

 98.8096123086663
 93.9938218973205

 1.45369262386483
 6.2941168248466

 1.45369262386483
 6.2941168248466

 1.45369262386483
 6.2941168248466

 .714376497277024
 3.14356401045525

 2.01435467510138E-02
 6.56947569992871E-02

 8.72887025877263E-02
 .374599890974659

 8.72887025877263E-02
 .424919279314538

 .106712836954775
 .465454342143885

 .118942847482177
 .560502075674769

 .293999860913606
 1.25239366534811

 .739316126587803
 3.15055281439134

 .213425673909551
 .965852703968243

 .262345716019156
 1.03434298254197

 .263544736659097
 1.15035712788113

 .861136623605839
 4.1541450596145

 .861136623605839
 4.1541450596145

 .861136623605839
 4.1541450596145

 3.11745366384737E-02
 .13698055714745

 3.11745366384737E-02
 .13698055714745

 3.71696398381802E-02
 .192890988636205

 3.71696398381802E-02
 .192890988636205

 4.67618049577105E-02
 .251596941699398

 4.67618049577105E-02
 .251596941699398

 4.58025884457575E-02
 .26138126720993

 4.58025884457575E-02
 .26138126720993

 5.58743618212644E-02
 .212459639657269

 5.58743618212644E-02
 .212459639657269

 5.53947535652879E-02
 .269767831933243

 5.53947535652879E-02
 .269767831933243

 6.45073104288417E-02
 .306109612400934

 6.45073104288417E-02
 .306109612400934

 7.38596714203838E-02
 .426317040101757

 7.38596714203838E-02
 .426317040101757

 .085130465435832
 .395566302782942

 .085130465435832
 .395566302782942

 9.61614553232919E-02
 .384384216485191

 9.61614553232919E-02
 .384384216485191

 8.75285067157146E-02
 .503193883398795

 8.75285067157146E-02
 .503193883398795

 .1817715290151
 .813496778161385

 .1817715290151
 .813496778161385

 4.27426877726273
 12.7294074892023

 4.27426877726273
 12.7294074892023

 1.96423561235183
 9.62917406315083

 2.51794334387672E-02
 .219448443593363

 2.51794334387672E-02
 .219448443593363

 1.93905617891306
 9.40972561955747

 .102636166778975
 .388577498846847

 .211507240885645
 1.02316089624422

 .503348864647356
 2.56908432690829

 1.12156390660109
 5.42890289755811

 2.3100331649109
 3.10023342605147

 2.3100331649109
 3.10023342605147

 4.70016090856988E-02
 .117411906126386

 4.36443512938632E-02
 .276756635869337

 6.09102485090178E-02
 .215255161231707

 9.76002800912215E-02
 .630390115035713

 2.0608766759311
 1.86041960778832

 .282968871026146
 1.31389513998574

 .282968871026146
 1.31389513998574

 4.07667017580041E-02
 .16214025131739

 4.07667017580041E-02
 .16214025131739

 4.07667017580041E-02
 .16214025131739

 .163066807032016
 .717051283843283

 .163066807032016
 .717051283843283

 7.88955581081373E-02
 .367611087038564

 .084171248923879
 .349440196804719

 7.91353622361255E-02
 .43470360482507

 7.91353622361255E-02
 .43470360482507

 7.91353622361255E-02
 .43470360482507

 .588719134211176
 2.23082621640132

 .588719134211176
 2.23082621640132

 .588719134211176
 2.23082621640132

 7.74567333402077E-02
 .293529765315964

 7.74567333402077E-02
 .293529765315964

 .511262400870969
 1.93729645108536

 9.85594966031745E-02
 .471045385292761

 .412702904267794
 1.4662510657926

 9.14852748275209
 44.9226339404274

 4.55507941113698
 21.5297094055323

 4.1747500641476
 19.5281159582349

 .112228331898505
 .617810267950743

 .112228331898505
 .617810267950743

 .212226653269609
 1.00499000601037

 .212226653269609
 1.00499000601037

 .64603232080037
 3.06948268873265

 .64603232080037
 3.06948268873265

 3.20426275817912
 14.8358329955411

 .770970271482253
 3.59364298393973

 1.12444155613695
 4.9690395985631

 1.30885093055992
 6.27315041303831

 .380329346989379
 2.00159344729743

 .380329346989379
 2.00159344729743

 .380329346989379
 2.00159344729743

 .21630332344541
 .866611688075703

 .21630332344541
 .866611688075703

 .21630332344541
 .866611688075703

 .21630332344541
 .866611688075703

 4.37714474816969
 22.5263128468194

 .622531516257521
 2.87798946088366

 .299035747601359
 1.38937422249556

 .299035747601359
 1.38937422249556

 .323495768656162
 1.4886152383881

 .323495768656162
 1.4886152383881

 1.29710052828849
 7.79810743189411

 .535482617797783
 2.84024991962875

 .535482617797783
 2.84024991962875

 .761617910490711
 4.95785751226535

 .761617910490711
 4.95785751226535

 .643874083648476
 2.99540136701005

 .643874083648476
 2.99540136701005

 .643874083648476
 2.99540136701005

 1.8136386199752
 8.85481458703157

 1.8136386199752
 8.85481458703157

 1.8136386199752
 8.85481458703157

 81.3712959254881
 18.7285967879457

 81.3712959254881
 18.7285967879457

 .176975446455335
 .388577498846847

 .176975446455335
 .388577498846847

 .176975446455335
 .388577498846847

 81.1943204790327
 18.3400192890989

 8.41880352128381
 1.90095467061767

 1.23355243437161
 .438896887186727

 7.18525108691221
 1.46205778343094

 1.42491612850624
 3.24699830870945

 1.42491612850624
 3.24699830870945

 71.3506008292427
 13.1920663097717

 1.28343169299316
 4.56788225263128

 70.0671691362495
 8.62418405714046

 .217262539957363
 .494807318675482

 .217262539957363
 .494807318675482

 .217262539957363
 .494807318675482

 .217262539957363
 .494807318675482

 .217262539957363
 .494807318675482

 .611740330498049
 3.12539312022141

 .611740330498049
 3.12539312022141

 .611740330498049
 3.12539312022141

 .176975446455335
 .908544511692269

 .176975446455335
 .908544511692269

 .434764884042714
 2.21684860852914

 .434764884042714
 2.21684860852914

 .147719342840768
 .86940720965014

 .147719342840768
 .86940720965014

 5.85122072291352E-02
 .276756635869337

 5.85122072291352E-02
 .276756635869337

 5.85122072291352E-02
 .276756635869337

 5.85122072291352E-02
 .276756635869337

 8.92071356116324E-02
 .592650573780803

 8.92071356116324E-02
 .592650573780803

 8.92071356116324E-02
 .592650573780803

 8.92071356116324E-02
 .592650573780803

 7.62577127002664E-02
 .382986455697972

 7.62577127002664E-02
 .382986455697972

 7.62577127002664E-02
 .382986455697972

 7.62577127002664E-02
 .382986455697972

 7.62577127002664E-02
 .382986455697972

 7.62577127002664E-02
 .382986455697972
