## Supplementary figures and images for "Methane emission of humans is explained by dietary habits, host genetics, local formate availability and a uniform archaeome"

### Supplementary Item 6

Predicted exchange fluxes of amino acids from high (HE) and low (LE) methane emitters

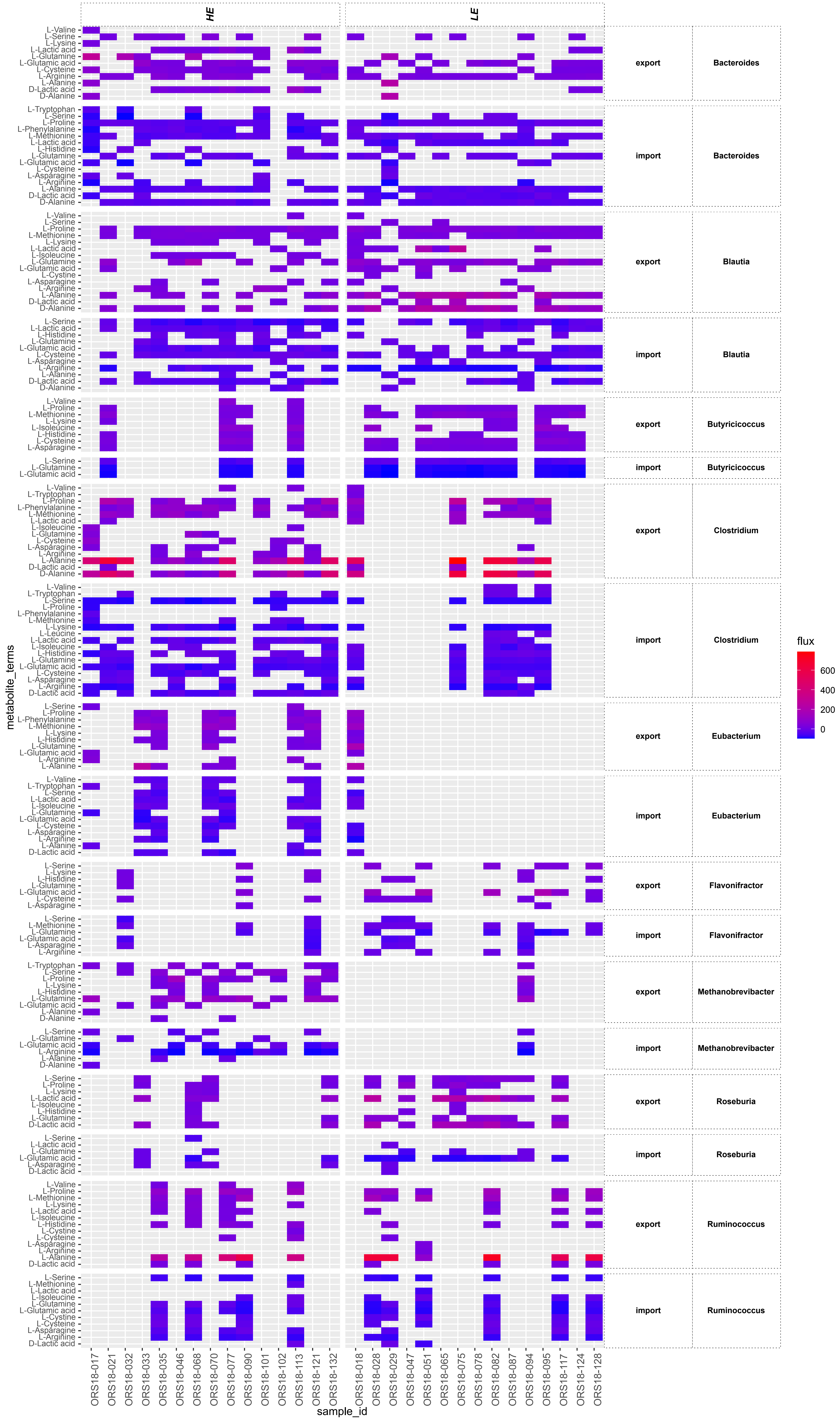

### Supplementary Item 7

Predicted exchange fluxes of C1-C4 metabolism from high (HE) and low (LE) methane emitters

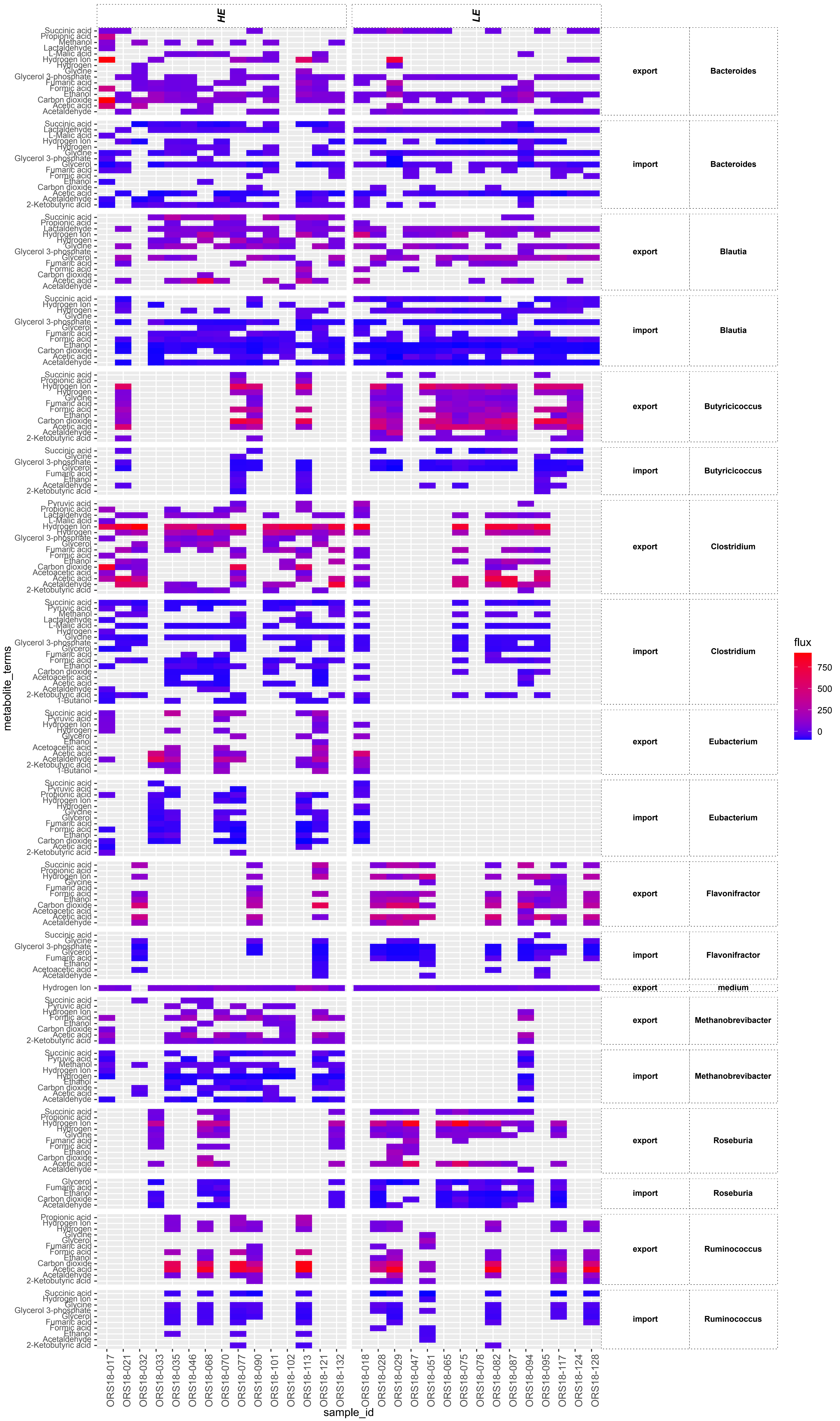

### Supplementary Item 8

#### Predicted exchange fluxes of complex compounds from high (HE) and low (LE) methane emitters

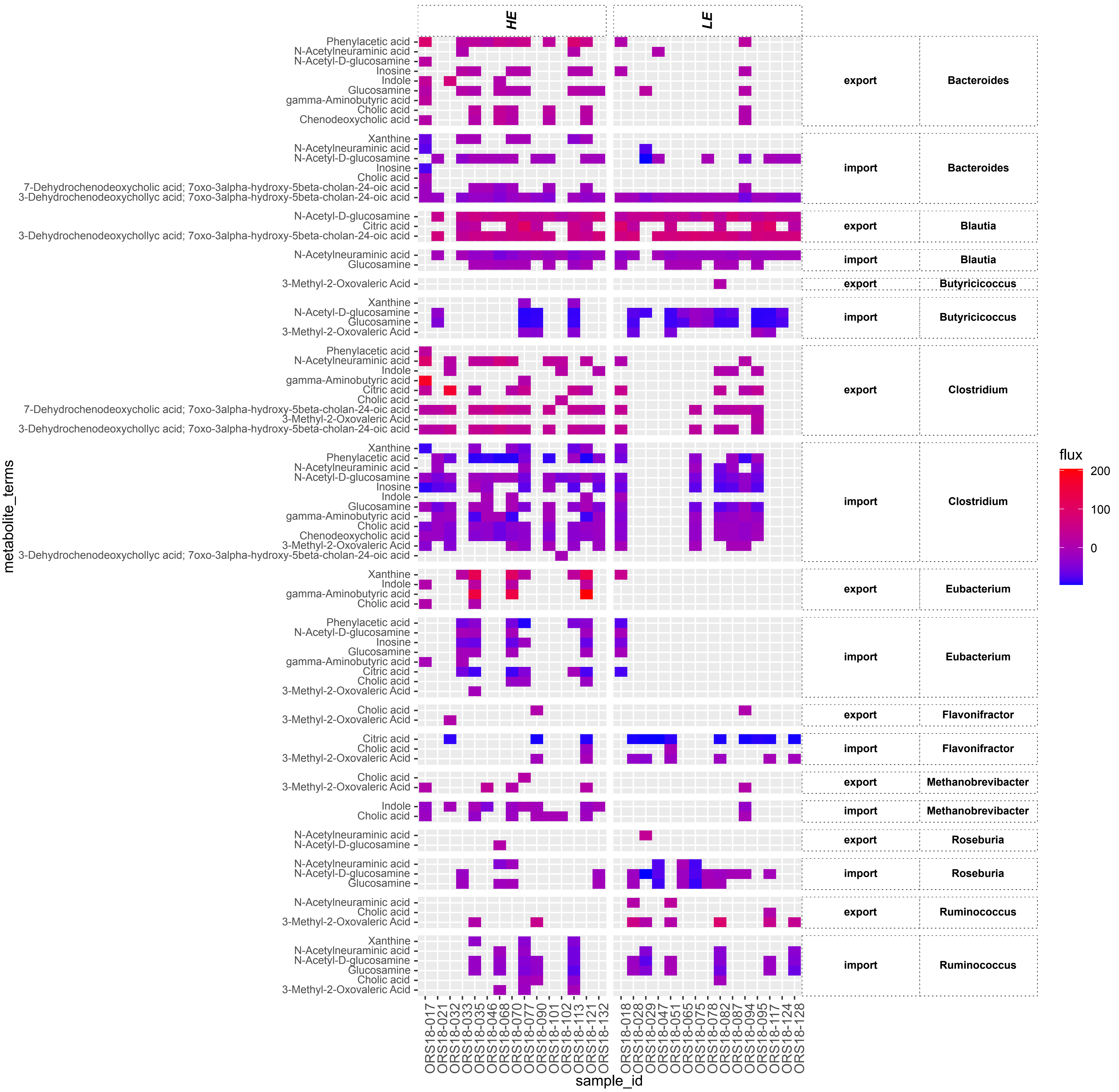

### Supplementary Item 9

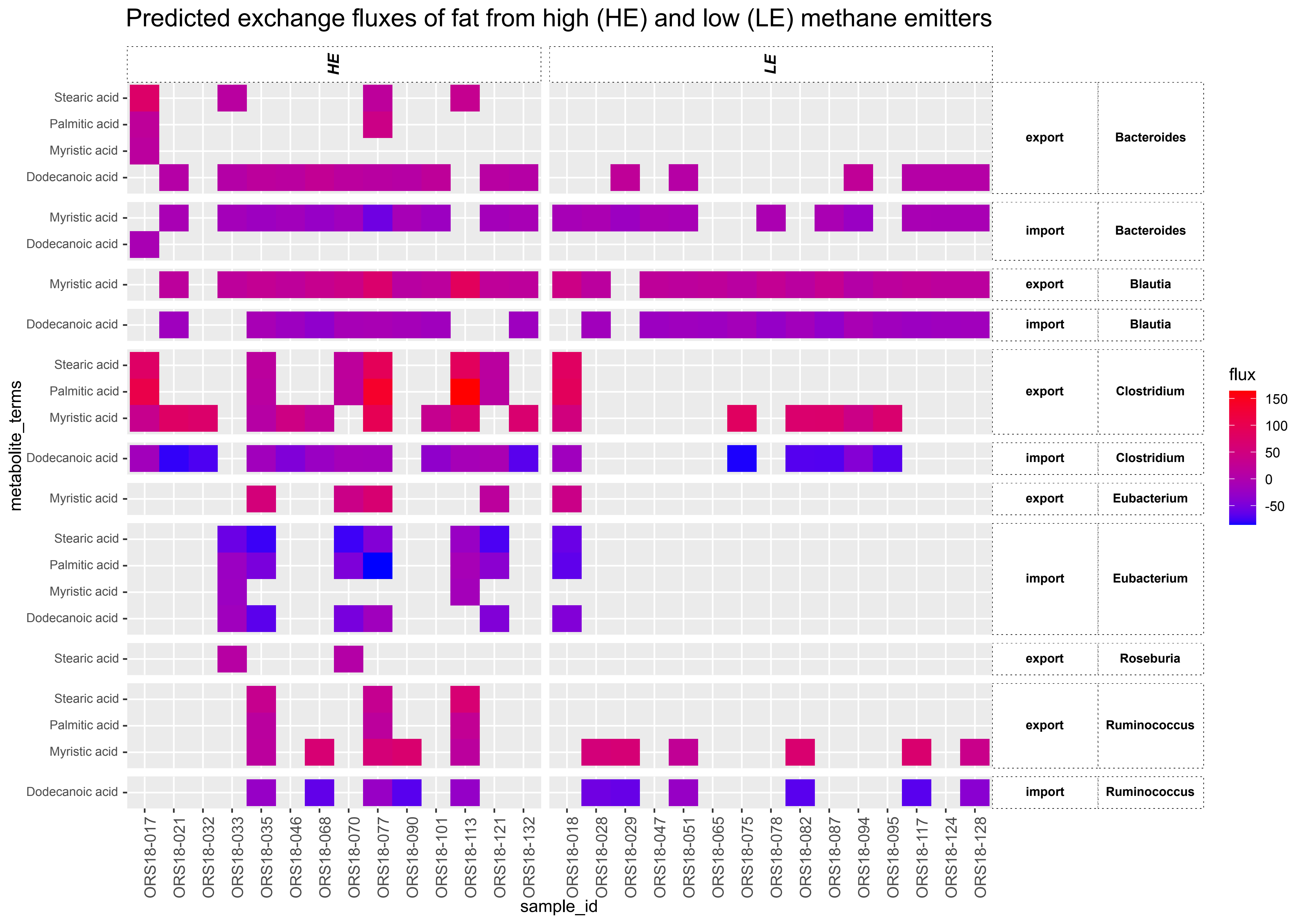

### Supplementary Item 10

#### Predicted exchange fluxes of nucleotides from high (HE) and low (LE) methane emitters

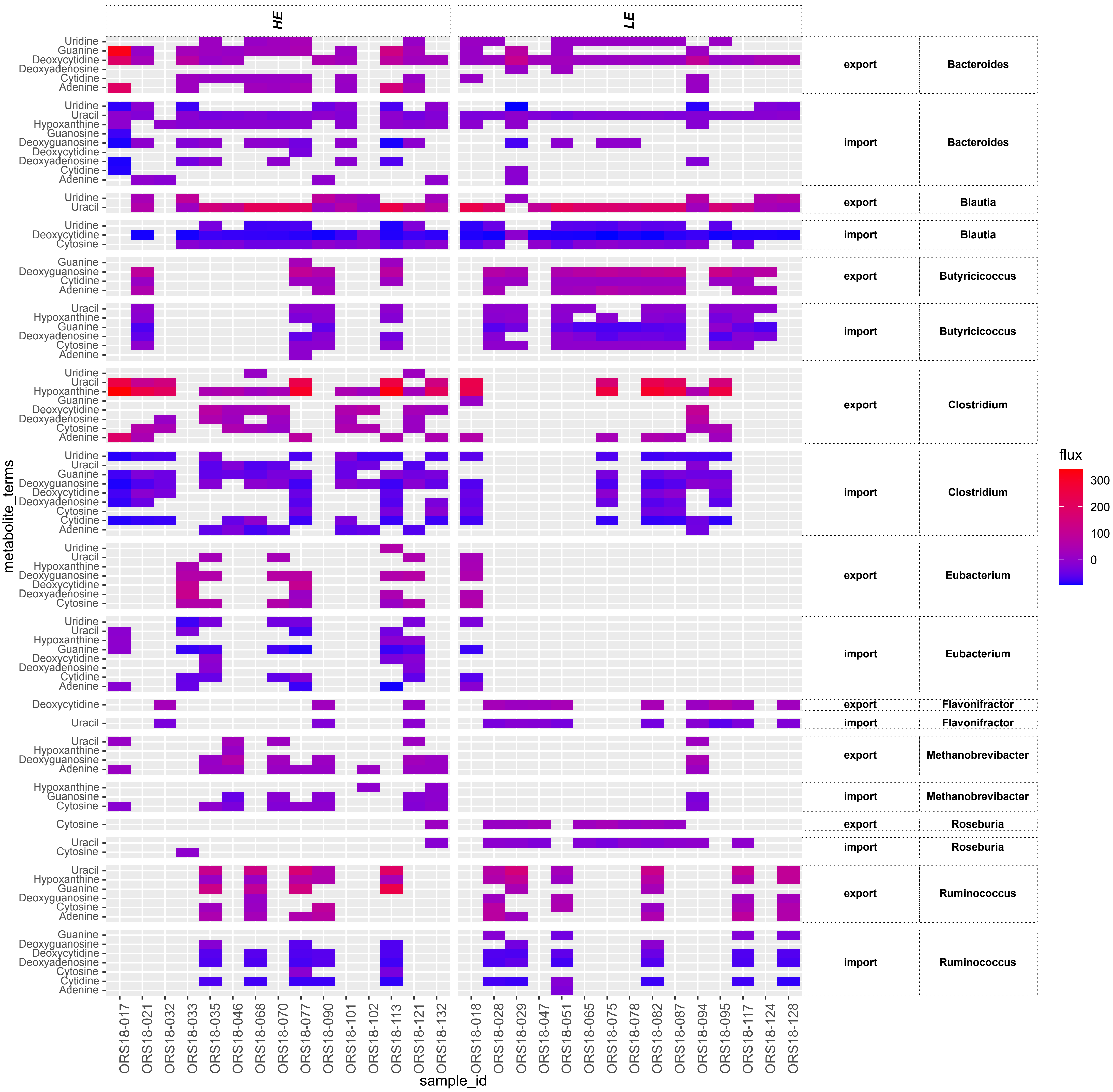

### Supplementary Item 11

Predicted exchange fluxes of non-grouped metabolites from high (HE) and low (LE) methane emitters

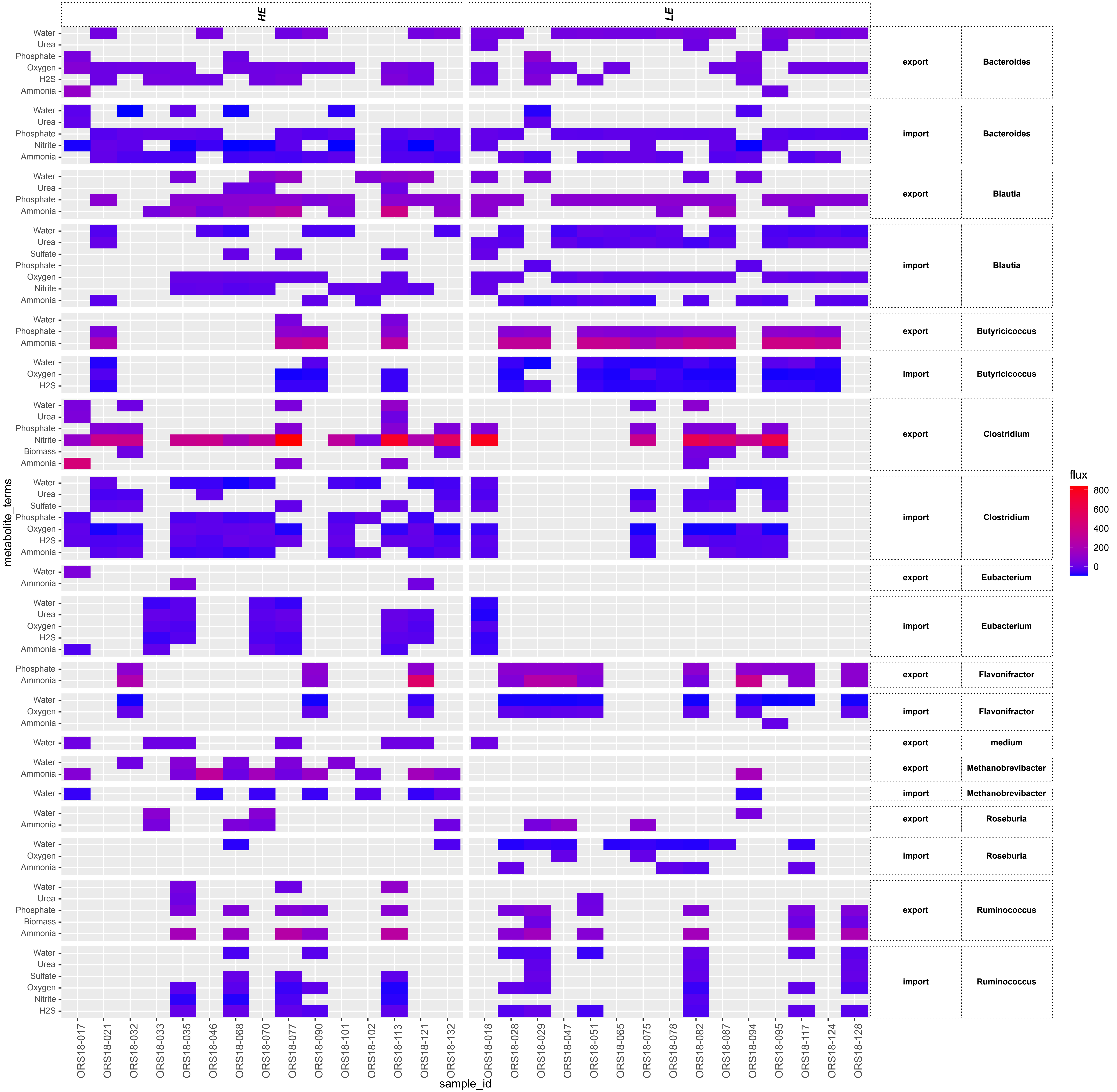

### Supplementary Item 12

Predicted exchange fluxes of sugars from high (HE) and low (LE) methane emitters

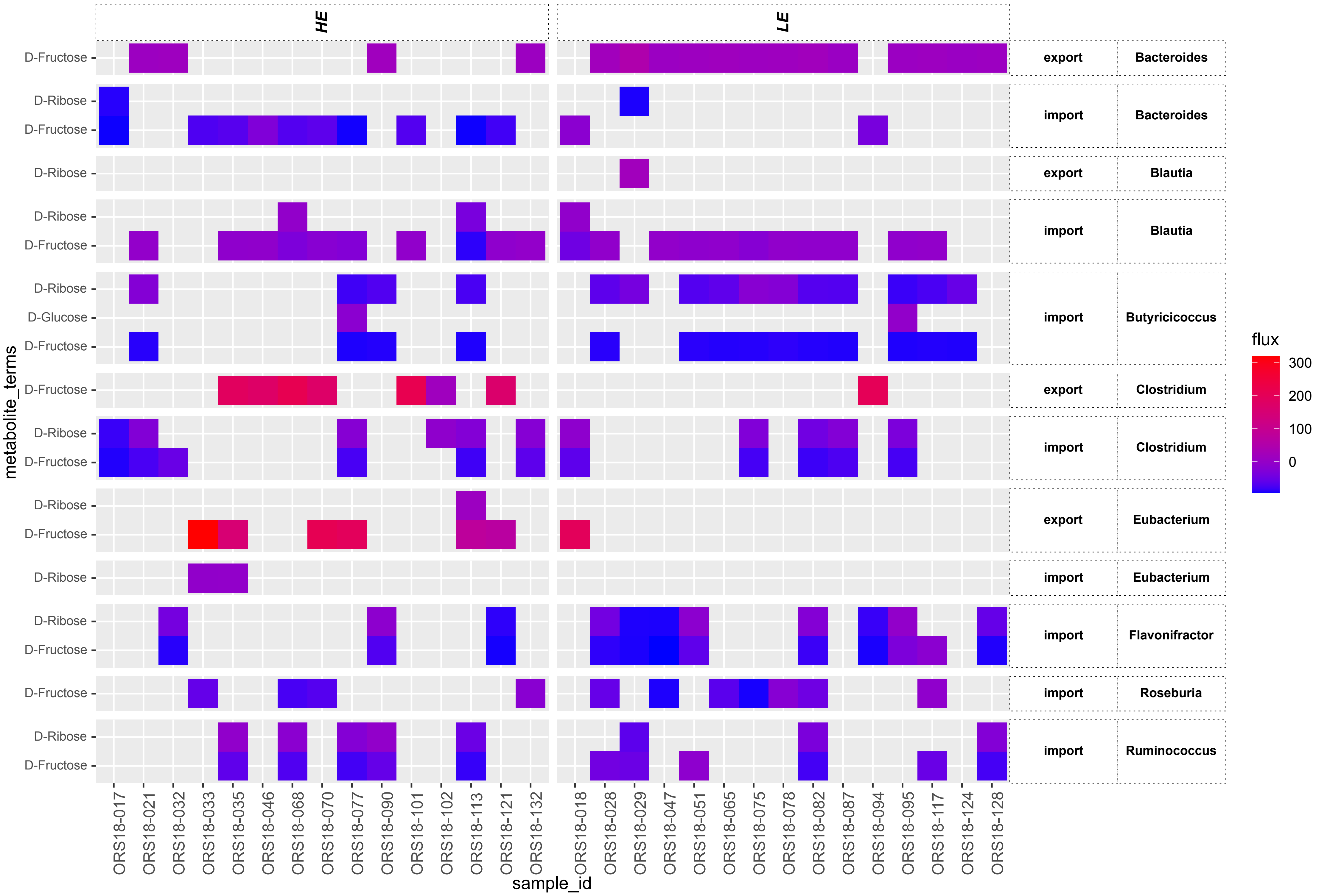

### Supplementary Item 13

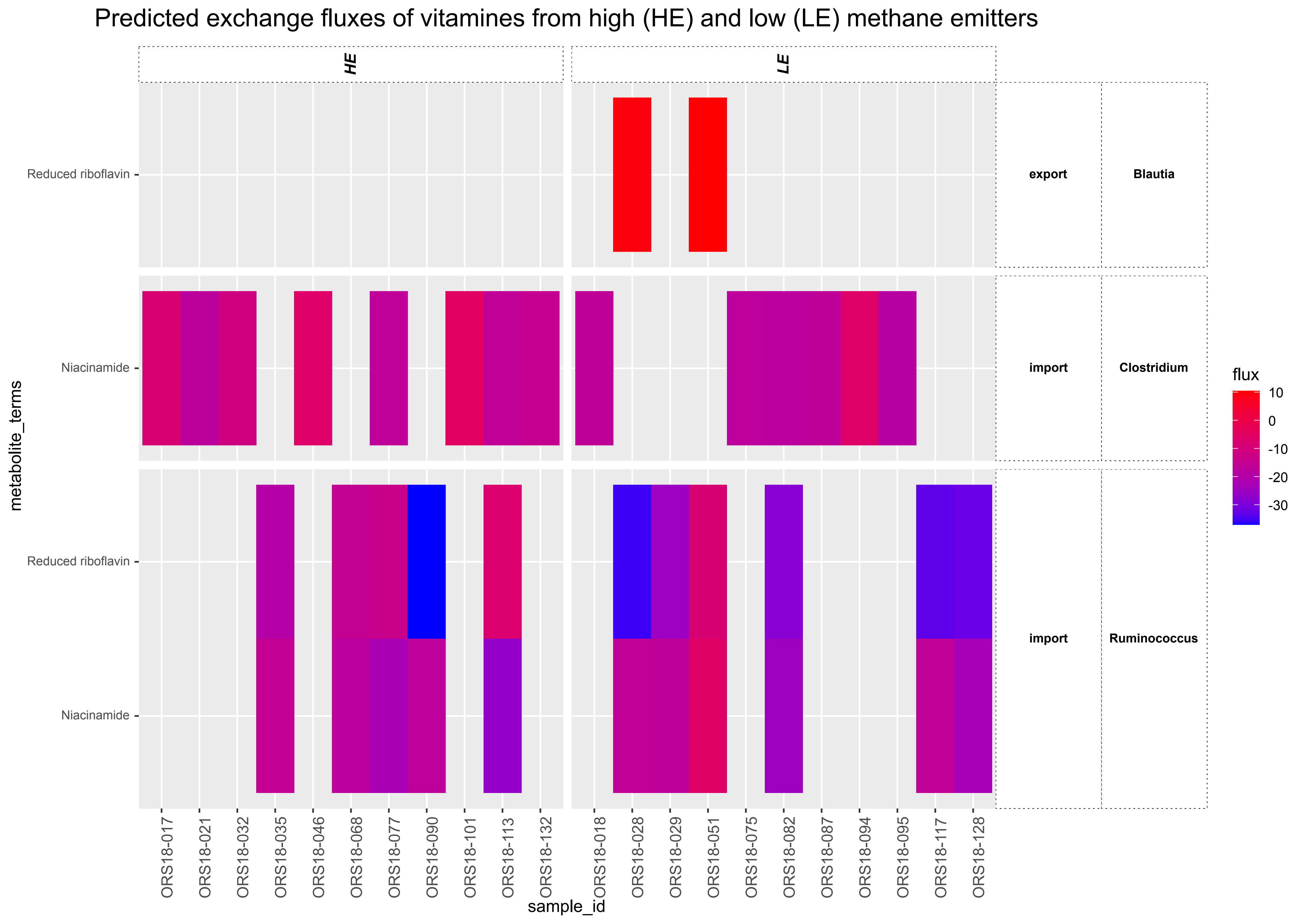
